# BNT162b2 mRNA vaccine-induced sex differences in the single-cell transcriptome of peripheral blood mononuclear cells in healthy adults

**DOI:** 10.1101/2023.10.02.560569

**Authors:** Johannes D Knapp, Aditi Bhargava

## Abstract

**Introduction:** Men reportedly experience more severe disease and adverse outcomes from COVID-19, including death. Women report more adverse events (AEs) after vaccination in general. While few studies have addressed sex-specific risk factors or molecular mechanisms behind COVID-19, none have examined sex differences in the response to COVID-19 vaccination.

**Methods:** We searched AE reporting databases to find sex differences specific to COVID-19 vaccines. We analyzed public datasets to identify baseline sex differences in gene expression across cell types and time points, and sex differences in the response to the second BNT162b2 mRNA vaccine dose.

**Results:** Sex differences in AE rates for mRNA vaccines equaled those for other non-mRNA vaccines. T cells and monocytes showed the greatest number of sexually dimorphic genes. Platelet counts in the study population differed significantly before vaccination (3.6% in females vs 1.8% in males) but not after the second BNT162b2 dose (7.2% vs 7.3%). There were no notable sex differences in the expression of key genes induced by the second dose after exclusion of platelets. BNT162b2 dose 2-specific APOBEC3A^high^ monocytes and the dose 2-induced gene signature persisted for longer in women. Glucocorticoid-responsive *TSC22D3, CEBPB/D* and *DDIT4* were specifically induced in females; the voltage-gated potassium channel regulatory subunit *KCNE3* was specifically induced in males.

**Conclusions:** This sexual dimorphism in both X-linked and autosomal gene transcriptome in PBMCs after mRNA COVID-19 vaccination might explain fatigue, autoimmune, and neurological AEs reported after vaccination at different rates in women and men.

## Introduction

Women develop greater antibody responses to vaccinations in general, but they are also more likely to be affected by adverse events compared to men.^1^ COVID-19 is a heterogenous disease with potential sex differences in symptoms and outcomes. mRNA vaccines were deployed for the first-time at large scale for COVID-19, and immune mechanisms modulated by mRNA vaccines, such as those approved under emergency use authorization and manufactured by Pfizer and Moderna, remain to be elucidated. Monocytes are increased in patients after natural infection with SARS-CoV-2,^2^ and specifically nonclassical monocytes have been associated with disease severity, but whether COVID-19 vaccines also alter monocytic cell populations is not known.

SARS-CoV-2 infection causes mild-to-moderate disease in most people. In a subset of people with severe disease, uncontrolled inflammation known as “cytokine storm” can lead to coagulation and thromboembolic events in either the arterial or the venous systems, often leading to thrombocytopenia. Platelets are produced from progenitor megakaryocytes during hematopoietic stem cell (HPC) differentiation,^3^ a process primarily driven by thrombopoietin made in the kidney and the liver. Platelets are versatile immune cells that can act as antigen presenting cells and mediators of thrombosis. Platelets play a key role in the internalization of invading pathogens, including SARS-CoV-2, and they are known to harbor infectious virus and infect macrophages during phagocytosis of infected platelets in vivo.^4^ Both low and high platelet counts are indicators of pathology, resulting in thrombocytopenia or thrombocythemia, respectively. Monocytes are bone marrow-derived leukocytes that participate in immune responses by engulfing pathogens (phagocytosis). In humans, monocytes are classified into three main subsets based on their expression of specific cell surface markers: classical (CD14^+^CD16^−^), intermediate (CD14^+^CD16^+^) and non-classical (CD14^−^CD16^+^). Classical monocytes constitute 80–95% of circulating monocytes and do not secrete pro-inflammatory mediators.

Lysine-specific demethylase 6A (UTX/*KDM6A*) is an X-linked gene that escapes X-inactivation and specifically removes repressive trimethylation from histone H3 lysine 27 (H3K27me3). Abnormal expression of *KDM* genes is known in several cancers and inflammatory diseases.^5^ In mice, *Kdm6a* was the top sexually dimorphic gene in CD4^+^ T cells and associated with autoimmunity and neuropathology.^6^ Changes in expression levels of *KDM6A* and several genes that encode histones can be a sign of acute inflammation and cellular stress. SARS-CoV-2 infection alters mitochondrial function in PBMCs and increases reliance on glycolysis to compensate for energy deficiency.^7^ SARS-CoV-2 open reading frames (ORFs) localize in host’s mitochondria,^8, 9^ where they suppress hosts’ innate immunity by manipulating mitochondrial function and the mitochondrial antiviral-signaling protein (MAVS)/tumor necrosis factor receptor-associated factor (TRAF3 and TRAF6) signaling pathways. Mitochondria are the main site of ATP production and contain their own DNA (mtDNA). Any manipulation of mtDNA can be detrimental to its function as well to the progeny, if gonadal mtDNA is altered in anyway.

Our aim was to identify potential gene targets in immune cell subtypes that might underlie mechanisms behind sex differences in COVID-19 mRNA vaccine adverse events and possibly pathways that are shared with COVID-19. To do so, we analyzed public datasets from gene expression omnibus (GEO) to investigate sex differences following vaccination.^10^

## Methods

Publicly available CITE-Sequencing data from peripheral blood mononuclear cells (PBMCs) following vaccination with BNT162b2 was used in this analysis.^10^ In the original study, 3 male and 3 female volunteers each received two doses of the BNT162b2 mRNA vaccine, and samples were taken on day 0 (before Dose 1), 1, 2, 7, 21 (before Dose 2), 22, 28 and 42 (Table 1). Due to low sample size after the first vaccine dose (only one female on Day 1 and 2 females on Days 2– 21) we focused on sex differences before vaccination (Day 0) and after the second vaccine dose (Day 22, 28 and 42). Analysis was performed using the Aseesa Stars (www.aseesa.com) analysis tool as described below.

**Table 1:** Participant characteristics including sex, age, and available time points for each subject.

| Sample ID | Sex | Age | Missing Time Points |
| --- | --- | --- | --- |
| 2047 | M | 31 |  |
| 2049 | M | 35 |  |
| 2051 | F | 24 |  |
| 2052 | M | 54 | 1 |
| 2053 | F | 30 | 1 |
| 2055 | F | 34 | 1, 2, 7, 21 |
| Average Age (F) |  | 29 |  |
| Average Age (M) |  | 40 |  |

### Data Normalization

Data was normalized using transcripts per million (TPM) because the number of increased and decreased genes was more balanced compared to transcripts per cell (TPC), transcripts per positive cell (TPPC) and percent positive (Supplementary Fig. S1a). TPM was calculated for every cell individually, summed, and divided by the total cell count for each sample. TPC was calculated by dividing the transcript count by the total cell count for each sample. TPPC was calculated by dividing the transcript count by the number of cells in which at least one transcript of the symbol was present. Percent positive was calculated by dividing the number of cells in which at least one transcript of the symbol was present by the total cell count. If a symbol was present in multiple cell types, the number of transcripts, cells, positive cells, and TPM were summed before normalization. Cell counts were normalized to the total included cell count. For symbols that were detected in a test group but not in a control group, the value 2^−10^ was imputed as the normalized control average. Due to suspected cross-contamination in a small number of cells, female test groups were further restricted to cells that were positive for *XIST* and negative for *PRKY*. Male test groups were restricted to cells negative for *XIST*. Excluding *XIST*-negative female cells did not significantly affect the results (Supplementary Fig. S1b). Roughly 80–100% of female cells were positive for *XIST* depending on the cell type (Supplementary Fig. S2a). Unclassified Cluster 2 cells were always excluded; Platelets and Natural Killer T (NKT) cells were only included in Figs. 1a, 3 and Supplementary Fig. S2; 98.3% of NKT cells were positive for the platelet marker PPBP (Supplementary Fig. S2b). Antibody-dependent tags (ADTs) were excluded from analysis and did not count towards total transcript counts. Female cells tended to contain more transcripts per cell (Supplementary Fig. S2c), but there were no significant differences in the number of transcripts per positive cell (Supplementary Fig. S2d). In fact, the number of transcripts per positive cell did not fluctuate by more than 5.1% in Natural Killer cells and less than 2.8% in other cell types between females and males.

**Fig. 1.**
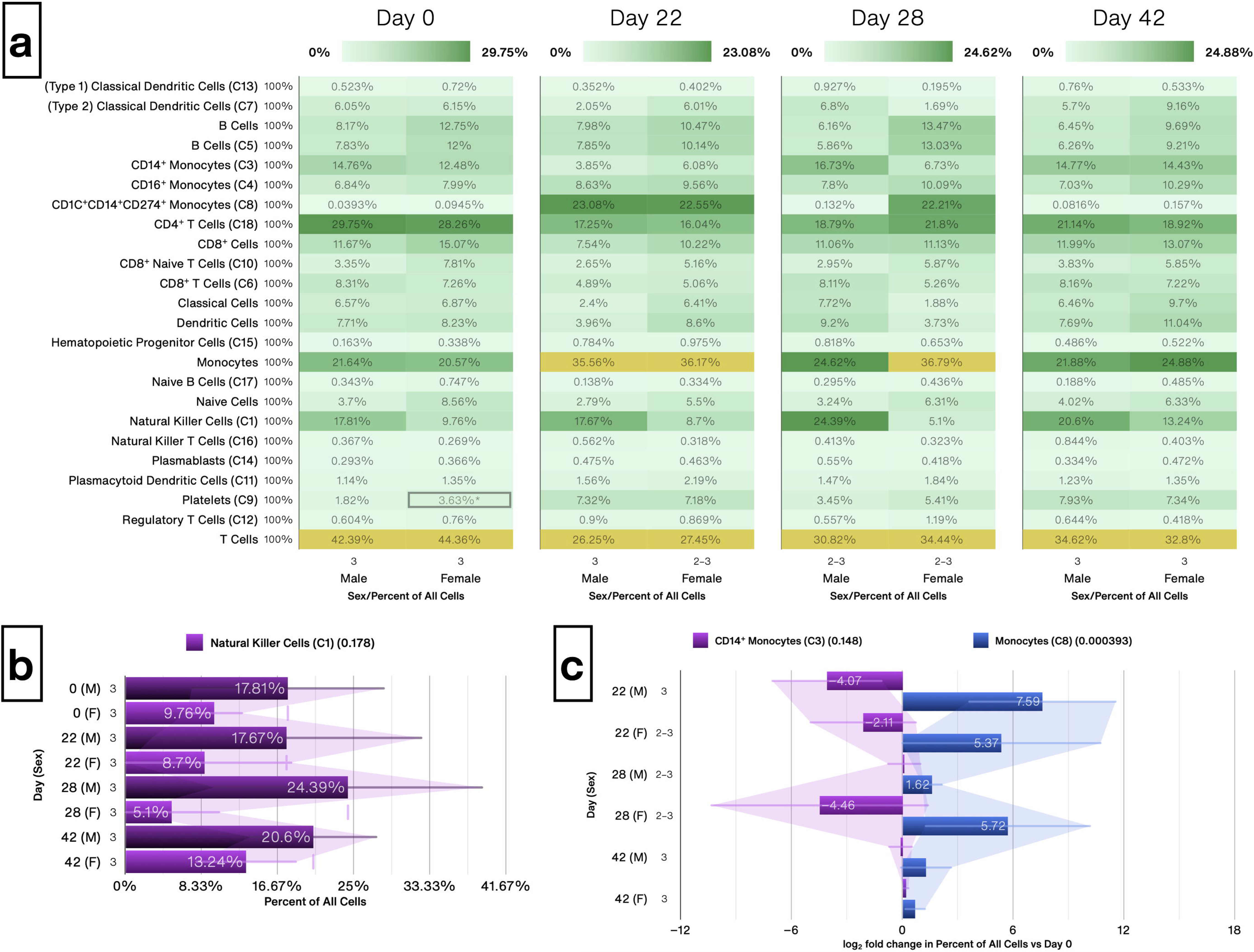
Immune cell subtype status before and after mRNA vaccination. (**a**) Heat map showing the absolute abundance of all included clusters and composite cell types, normalized to % of included cells, shown in males and females separately for each time point. Average across all samples shown; percentages next to symbols denote the percentage of samples in which cells of this type were detected; group sizes (n) are shown above group names. The maximum value of a heat map does not include outliers (highlighted in yellow) that are defined as less than Q1 − 1.5(IQR) or greater than Q3 + 1.5(IQR). IQRs were calculated by interpolating between data points to determine empirical quantiles. Composite cell types: B cells (naïve and mature B cells), CD8^+^ cells (naïve and mature CD8^+^ T cells), Classical cells (cDC1s and cDC2s), Dendritic cells (classical and plasmacytoid DCs), Monocytes (CD14^+^, CD16^+^ and Cluster 8), naïve cells (naïve B and CD8^+^ T cells) and T cells (CD4^+^, CD8^+^ and Regulatory T cells). (**b**) Natural Killer cells were increased non-significantly in males before (Day 0) and after the 2^nd^ BNT162b2 mRNA vaccine dose (Day 22, 28 and 42). Average across all samples is shown. Error bars: ± standard deviation. (**c**) Cluster 8 cells (blue bars) displaced CD14^+^ monocytes (purple bars) on Day 22 in males, and on Days 22 and 28 in females. log_2_ fold change was calculated for each sample versus its corresponding value on Day 0, with the average shown. Error bars: ± standard deviation. M: male; F: female. Numbers next to bars denote the number of included samples (n).

### Heat Maps

Heat maps were created to show several symbols in a single chart instead of multiple bar charts. Value labels show the group average or average log_2_ fold change vs the respective control group. Value labels are drawn for every symbol in absolute heat maps, and for values greater than 33% of the heat map’s maximum value in relative heat maps. Labels for values less than 16.67% of the maximum are drawn in black for legibility.

The maximum value of a heat map does not include outliers (highlighted in yellow) defined as less than Q1 − 1.5(IQR) or greater than Q3 + 1.5(IQR). IQRs were calculated by interpolating between data points to determine empirical quantiles. Degrees of freedom were calculated using the Welch– Satterthwaite equation, and exact p-values were calculated using the cumulative t-distribution functions as p = 2 * MIN(P(x), Q(x)). Labels next to gene symbols denote the percentage of cells across all test groups in which transcripts of the symbol were detected; percentages next to other symbols (e.g. cell types) denote the percentage of samples in which the symbol was detected. Numbers above test group labels denote the number of samples that were included (n); if not all symbols were present in the same number of samples a range is shown.

### Bar Charts

Values in bar charts are calculated in the same way as those in heat maps. Relative bar charts show average log_2_ fold change vs each test group’s control group, which is not shown. Absolute bar charts additionally include either all, the first or no control groups, which are drawn in darker colors than test groups. Bar chart legends may include the average normalized abundance of the first control group in parentheses next to symbols, and the sorted four cell types with greatest abundance across all test groups. Error bars represent the standard deviation. For female vs male comparisons the comparison mode Value-to-Average was used, in which the set of changes for each sample 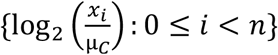 versus the normalized control average μ_*C*_ is averaged; for inter-time point comparisons the mode Value-to-Value was used, in which the set of changes for each sample versus its own control value 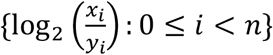 is averaged. Welch’s t-test was performed as in heat maps, with ^†, ††^ and ^†††^ denoting p < 0.05, 0.01 and 0.001 versus the previous test group (one bar above). Labels next to symbols denote the number of samples that were included (n). The filled fraction of a bar represents the percentage of cells in which transcripts of the symbol were detected; in charts showing changes in cell types, it represents the percentage of samples in which cells of that type were detected.

### Batch analysis

During batch analysis (used for volcano plots, Venn diagrams, Marker Genes, supplementary tables and to identify most changed genes), gene expression was calculated for every symbol in each test group, as calculated for bar charts, and symbols were sorted by ascending p-Value and descending absolute and relative change. For Impact rankings, symbols were sorted by the average rank across these 3 sorting methods (p-Value, relative change, and absolute change). Symbols were excluded using the criteria listed below. To identify symbols that were significantly changed across multiple test groups (related to Figs. 2 and 3), symbols were included if they were invalid in not more than 1 test group (per the exclusion criteria below), if they were significant in at least 66.7% of test groups and if they were changed in the same direction (all increased or all decreased) in every test group. The log_2_ fold change value 1 was imputed for symbols that were detected in a test group but not in its respective control group.

**Fig. 2.**
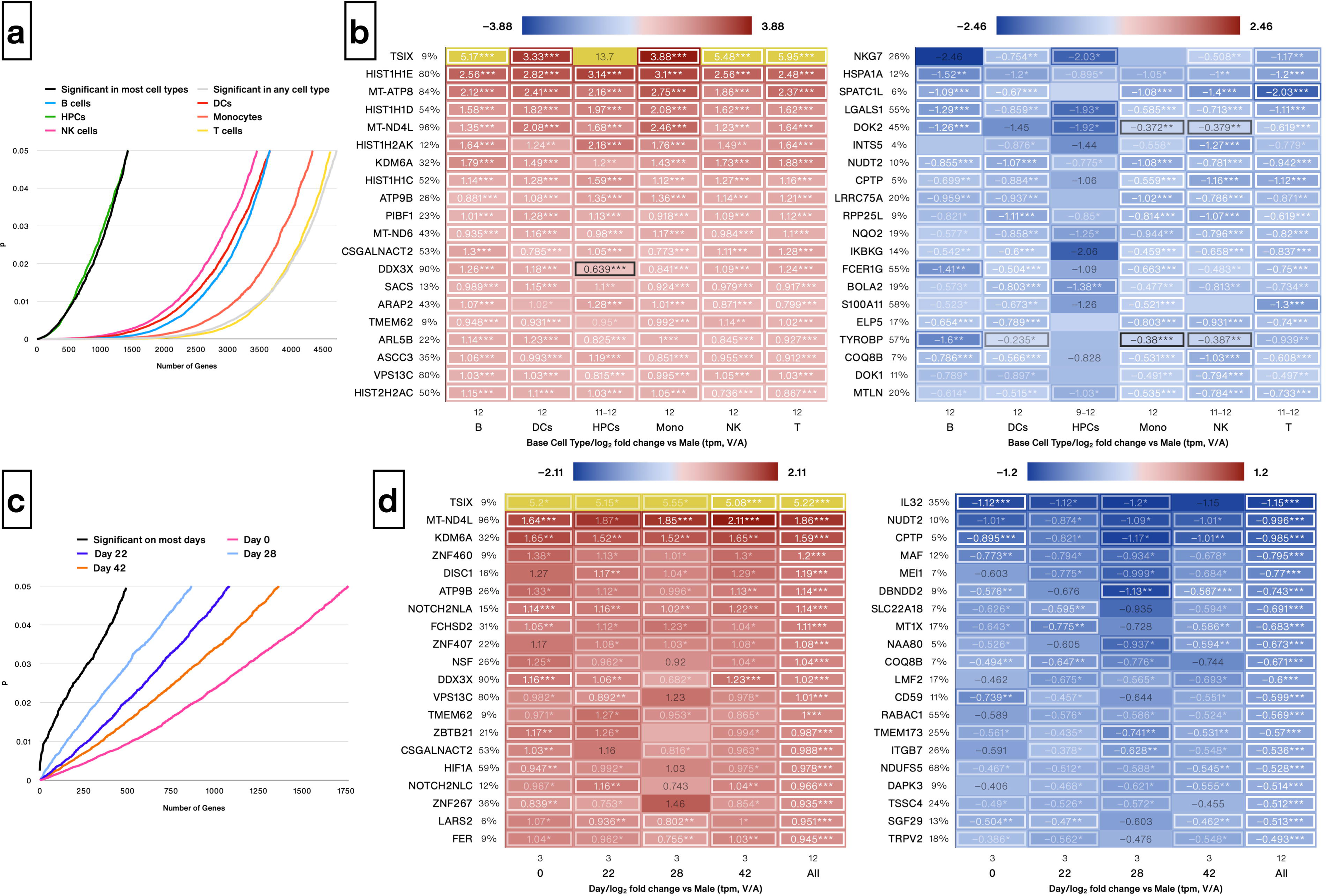
Genes with significant sex differences that held across most major immune cell base types and after the 2^nd^ vaccine dose. (**a**) Line chart visualizing the number of genes with significant (p < 0.05, Welch’s t-test) sex differences in each base immune cell type using pooled samples from Days 0, 22, 28 and 42. B cells include naïve and mature B cells; Dendritic cells (DCs) include types 1 and 2 classical (cDC1s/cDC2s) and plasmacytoid DCs; Monocytes include CD14^+^, CD16^+^ and Cluster 8; T cells include CD4^+^, CD8^+^ and Regulatory T cells. (**b**) Heat map of genes that were significantly increased (left) and decreased (right) in females versus males across most base cell types (see Methods), sorted by descending/ascending average log_2_ fold change vs male across all groups. Significantly changed values by Welch’s t-test are enclosed in a box and embellished with *, ** or *** denoting p < 0.05, 0.01 and 0.001, respectively. Average across all samples is shown; outliers are highlighted in yellow (see Methods). Labels next to symbols denote the percentage of cells in which transcripts were detected across all test groups; group sizes (n) are shown above group names. (**c**) Line chart visualizing the number of genes with significant (p < 0.05, Welch’s t-test) sex differences in any cell type on each sample day. (**d**) Heat map of genes that were significantly increased (left) and decreased (right) in females on most sample days (see Methods), sorted by descending/ascending average log_2_ fold change vs male across all groups.

**Fig. 3.**
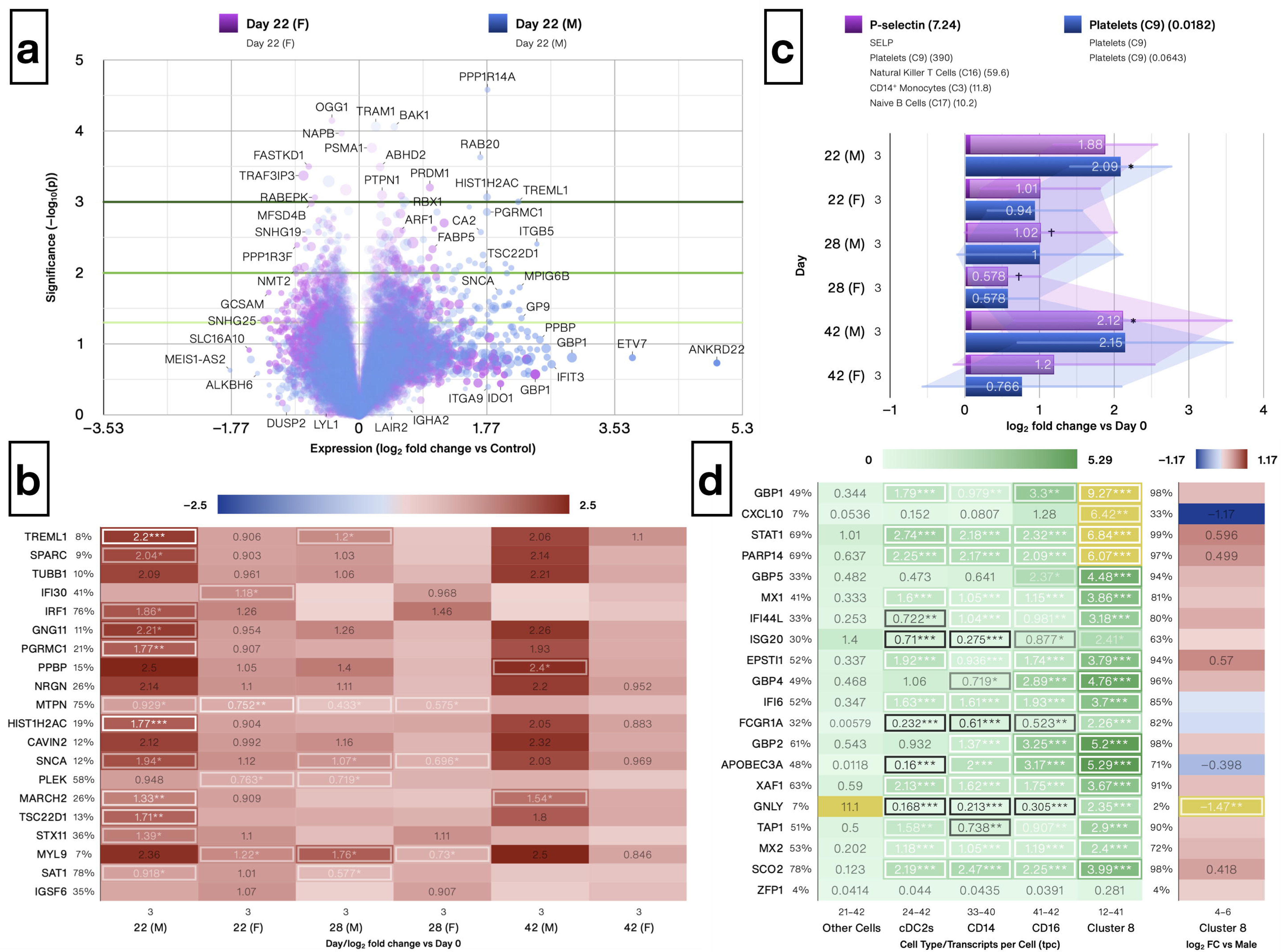
Platelet markers and counts increased more strongly in men after the 2^nd^ BNT162b2 dose, but Cluster 8 cells showed few sex differences. (**a**) Volcano plot showing changes in gene expression following the second vaccine dose (Day 22) in females (purple circles) and males (blue circles) separately, with platelets included. The size of a point represents its quantifiability, calculated as 0.375 + 0.625 * sqrt([percent of cells positive]) * [percent of samples positive] * [maximum point size]; the opacity of points represents expression. (**b**) Heat map of the 20 most-changed genes following the second vaccine dose (Day 22), with platelets included, sorted by impact score, as calculated for both sexes combined but with both sexes shown separately. log_2_ fold change was calculated for each sample versus its corresponding value on Day 0, with the average shown and *, ** and *** denoting p < 0.05, 0.01 and 0.001 by Welch’s t-test, respectively. Labels next to symbols denote the percentage of cells in which transcripts were detected across all groups; group sizes (n) are shown above group names. (**c**) Platelet counts and the platelet activation marker p-selectin (SELP) were consistently increased following the second vaccine dose, and this increase was greater in females than in males. log_2_ fold change was calculated for each sample versus its corresponding value on Day 0, with the average shown. Platelet counts were normalized to % of all cells; gene expression was normalized using TPM. Error bars: ± standard deviation; *, ** and *** denote p < 0.05, 0.01 and 0.001 vs Day 0 by Welch’s t-test, respectively. M: male; F: female. (**d**) Heat map of the top 20 markers for Cluster 8 cells relative to other monocytes and Type 2 Classical Dendritic Cells (cDC2s) across all 7 time points (left; transcripts per cell; *, ** and *** denote p < 0.05, 0.01 and 0.001 by Welch’s t-test vs Other Cells) and sex differences in their expression on Days 22 and 28 (right; log_2_ fold change female vs male). Labels next to symbols denote the percentage of cells in which transcripts were detected across all groups except Other Cells.

### Symbol exclusions

During batch analysis only symbols were included that met each of the following criteria in either the test or control group: a) the symbol is present in at least 4.1667% of cells (average across all samples), b) the group contains at least 32 cells total, and at least 4 cells that are positive for the symbol, and c) the symbol is present in at least 75% of samples.

### Venn diagrams

Venn diagrams show two test groups that were each compared to their respective control group (or the same control group), and highlight symbols that are: significantly changed only in the first (left) or only in the second test group (right); significantly increased (bottom left) or decreased (bottom right) in both test groups, or significantly increased in the first and significantly decreased in the second test group (top left) and vice versa (top right). All symbol lists are sorted by Impact ranking as described in the Batch analysis section, and values shown are log_2_ fold change versus control.

### Markers

Gene expression was calculated using the normalization modes % Positive, TPPC and TPM. TPC was not considered for marker identification because it is given by % Positive * TPPC. Only genes that had the highest or lowest abundance relative to the other groups using at least two normalization modes were considered as markers. Markers were sorted individually for each normalization mode, and the final markers were sorted by average rank across the 3 normalization modes.

## Results

### Women are overrepresented in COVID-19 vaccine adverse event reports

For COVID-19 vaccines, roughly ⅔ of AEs reported in surveillance databases were submitted for female patients, whereas ∼60% of serious AE reports were submitted for females. Databases that were used to search for AEs included vaccine adverse event reporting system (VAERS), VigiAcess, EudraVigilance (European Economic Area only), and Pfizer documents, covering a period between February 2021 and August 2022 (Table 2). Nearly 67.4% of all AEs and 60% of severe AEs for COVID-19 vaccines were reported by women, which is comparable to AE rates of other non-COVID-19 vaccines such as SHINGRIX (females: 67.2% of reports, 2017–2022) and quadrivalent influenza vaccines (70.4%, 2012–2022). We observed sex differences in the incidence of myocarditis (males: 0.91% of reports, females: 0.22%) for COVID-19 vaccines in VAERS. Given that COVID-19 symptoms, AEs, and outcomes that are reported by males and females differ, it is likely that COVID-19 mRNA vaccines result in both shared and distinct immunomodulation between the sexes.

**Table 2:** Adverse event report counts for BNT162b2, mRNA-1273, SHINGRIX (herpes zoster vaccine) and quadrivalent influenza vaccines, segregated by sex, using data from VAERS, VigiAccess, EudraVigilance and Pfizer document productions.

| Dataset | Details | All Adverse Events | Female | Male | Date |
| --- | --- | --- | --- | --- | --- |
| Vaccine Adverse Event Reporting System (VAERS) | COVID-19 Vaccines, US Only | 825,174 | 561,949 (68.1%) | 263,225 (31.9%) | August 19, 2022 |
| VigiAccess | COVID-19 Vaccines, World | 4,151,749 | 2,788,782 (67.17%) | 1,362,967 (32.83%) | August 28, 2022 |
| EudraVigilance | BNT162b2, Serious Adverse Events Only, European Economic Area | 234,766 | 140,683 (59.92%) | 94,083 (40.08%) | August 27, 2022 |
| EudraVigilance | mRNA-1273, Serious Adverse Events Only, European Economic Area | 41,693 | 25,268 (60.6%) | 16,425 (39.4%) | August 27, 2022 |
| Pfizer Document Productions | BNT162b2, Document 5.3.6 (Cumulative Analysis of Post-authorization Adverse Event Reports), World | 39,096 | 29,914 (76.51%) | 9,182 (23.49%) | February 28, 2021 |
| Reference (VAERS) | Quadrivalent Influenza Vaccines, US Only | 34,484 | 23,171 (67.19%) | 11,313 (32.81%) | 2012–2022 |
| Reference (VAERS) | SHINGRIX (Herpes zoster vaccine), US Only | 51,152 | 35,998 (70.37%) | 15,154 (29.63%) | 2017–2022 |

### Immune cell subtype status in women and men before and after 2 doses of Pfizer’s BNT162b2 mRNA vaccine

Before vaccination (Day 0), platelets were significantly increased in females compared to males (3.63% vs 1.82%, Fig. 1a, p = 0.0471). Although platelet counts are known to be higher in females,^11^ the difference was reported to be minor (7.4% more in females), and therefore these differences likely reflect individual variability in the study population rather than reproducible sex differences. B and T cells are known to be more abundant in females, whereas NK cells are known to be more abundant in males.^12^ In agreement with previous observations, similar trends were noted for mature B cells (12.72% vs 8.16%), naïve B cells (0.75% vs 0.34%) and NK cells (9.7% vs 17.8%) although they did not reach statistical significance likely due to low sample size and high age variability in the male group (Fig. 1b, Table 1). T cells counts were similar between sexes (44% vs 42%), with notably less naïve CD8^+^ T cells in males (3.35% vs 7.79%) but similar numbers of mature CD8^+^ T cells (8.3% vs 7.25%, Fig. 1a). In the original report, Arunachalam et al.^10^ had identified a novel monocyte population (Cluster 8 cells (C8)) that appeared transiently following the 2^nd^ vaccine dose (Days 22 and 28) and expressed the classical monocyte marker CD14, dendritic cell marker CD1C and high levels of programmed cell death 1 ligand 1 (PD-L1/CD274). We found that C8 cells also expressed high levels of DNA dC->dU-editing enzyme APOBEC-3A (Supplementary Fig. S3a). In males, these C8 cells displaced classical monocytes on Day 22 but mostly disappeared by Day 28, whereas C8 remained abundant in females through Day 28 (Fig. 1c).

### Baseline sex differences in single-cell transcriptome in immune cell subtypes of healthy subjects

Next, we identified sex differences in gene expression that held across most immune cell subtypes. To minimize false positive findings due to low sample size, we created subgroups for the six major base cell type (B cells, DCs, HPCs, Monocytes, NK cells and T cells) and filtered genes that were changed in the same direction in all groups, significant in 4 of 6 groups and passed exclusion criteria (see Methods). A total of 1,435 genes met these criteria (Supplementary Table S1), whereas 4,718 (of 18,383 total) differed significantly when all cells were aggregated (Fig. 2a). Differing genes were sorted by ascending (Fig. 2b) and descending (Fig. 2c) expression. The most increased gene in females by significance and relative change was *TSIX* (17.6x increase, p = 10^−8.1^), which is antisense to X-inactive specific transcript (*XIST*) and transcribed exclusively from the inactive X chromosome.^13^ A number of genes encoding histones were among the most increased genes including H1.4 (*HIST1H1E*) and H1.2 (*HIST1H1C*). Several subunits of the mitochondrial complex I NADH-ubiquinone oxidoreductase chain^14^ (5 of 7) were significantly increased in females, namely *MT-ND4L* (3.3x), *MT-ND6* (2.1x), *MT-ND5* (83.7%), *MT-ND2* (44.6%) and *MT-ND4* (31.8%). *KDM6A* expression was increased 3.3-fold in females.

The most decreased gene in females was *NKG7*, a marker for NK cells that is also highly expressed in mature (but not naïve) CD8^+^ T cells (Supplementary Figs. S3b, S4a). A significant negative correlation between *KDM6A* and several NK effector genes including *CST7, NKG7* and *PRF1* was present in female but not male cells (Supplementary Fig. S4b), with the strongest correlation seen with *CST7* (females: r = −0.935, p = 10^−5.1;^; males: −0.304, p = 0.337). We used a similar approach to identify genes that were significantly changed across most time points, yielding 496 genes (Fig. 2c, Supplementary Table S2). *IL32* was the most decreased gene in females across all time points (Fig. 2d).

### BNT162b2 mRNA vaccine-induced sex differences in single-cell transcriptome in immune cell subtypes of healthy subjects

We first compared responses to vaccination (Day 22 vs Day 0) in males and females separately and found marked increases in the expression of platelet markers (Fig. 3a). To confirm that platelets are strongly involved in the vaccine response, we identified the most impactfully changed genes on Day 22 in both sexes combined and found that most of the top genes were platelet markers (Supplementary Table S3). Furthermore, when inspecting the same gene set in a sex-segregated fashion, platelet markers increased much more in males than in females, e.g. *TREML1* increased 4.6-fold in males but only 1.87-fold in females (Fig. 3b). In addition to increases in platelet markers, the number of platelets (% of all cells) increased 4.3-fold in males but only 1.9-fold in females, and the platelet activation marker p-selectin (SELP)^15^ increased 3.6-fold in men but only 2-fold in women (Fig. 3c). The sex difference in platelets before vaccination (3.6% in females vs 1.8% in males) was no longer present on Day 22 (7.3% vs 7.2%) and notably absent again on Day 42 (7.9% vs 7.3%). Cluster 8 cells were characterized by high expression of interferon-stimulated genes including guanylate-binding proteins *GBP1, GBP2, GBP4* and *GBP5*, guanosine-5’-triphosphate (GTP)-binding proteins *MX1* and *MX2, IFI6, IFI44L* and *ISG20* (Fig. 3d). Interestingly, C8 cells expressed granulysin (*GNLY*) which is usually restricted to lymphocytes. There were few sex differences in C8 cell markers on Days 22 and 28; of the top 20 markers only *GNYL* was significantly decreased in females. Female C8 cells tended to contain more transcripts of *STAT1* and *EPSTI1* and less *CXCL10* and *APOBEC3A*, although these did not attain statistical significance.

To identify sex differences in other immune cell types besides those in platelets, we performed the same analysis without platelets, which resulted in fewer apparent sex differences (Fig. 4a). 1,709 genes differed significantly following vaccination during sex-aggregated analysis (Supplementary Table S4). There did not appear to be any notable sex differences in the 40 most increased and decreased genes by impact score (top 10 shown in Fig. 4b), although the vaccine-induced gene signature was still evident (non-significantly) only in females on Day 28. We next sought to identify sex differences in the response to vaccination and searched for genes with the greatest difference between the change in males and the change in females on Day 22 vs Day 0. Genes were sorted by differences in p-value, relative change and absolute change, and the average rank across the 3 sorting methods was calculated (Supplementary Table S5), with the 20 most-differing genes shown in Fig. 4c. *TSC22D3, CEBPD, DDIT4* and *CEBPB* increased in females but not in males on Days 22 and 42, whereas *KCNE3* increased in males but decreased in females on Days 22, 28 and 42.

**Fig. 4.**
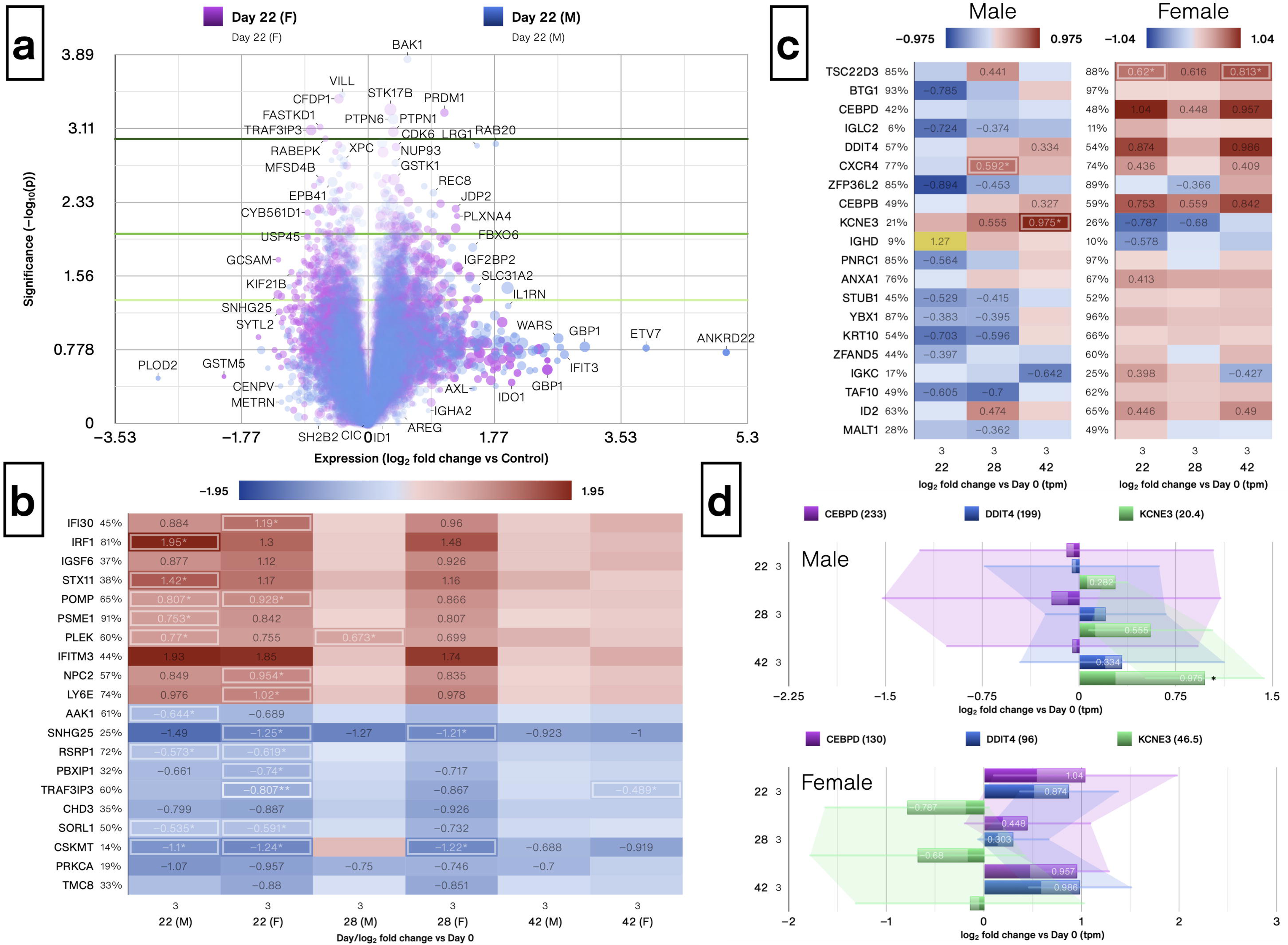
The gene signature induced by the 2^nd^ BNT162b2 dose did not differ substantially between males and females after platelets were excluded. (**a**) Volcano plot showing changes in gene expression following the second vaccine dose (Day 22) in females (purple) and males (blue) separately, with platelets excluded. The size of a point represents its quantifiability, calculated as 0.375 + 0.625 * sqrt([percent of cells positive]) * [percent of samples positive] * [maximum point size]; the opacity of points represents expression. (**b**) Heat map of the 10 most increased (top) and decreased (bottom) genes after administration of the 2^nd^ BNT162b2 dose (Day 22), with platelets excluded, sorted by impact score, as calculated for both sexes combined but with both sexes shown separately. log_2_ fold change was calculated for each sample versus its corresponding value on Day 0, with the average shown and *, ** and *** denoting p < 0.05, 0.01 and 0.001 by Welch’s t-test, respectively. Labels next to symbols denote the percentage of cells in which transcripts were detected across all groups; group sizes (n) are shown above group names. (**c**) Heat map of 20 genes exhibiting the greatest differences in response to vaccination (Day 22 vs Day 0) between males and females. Genes were ranked according to difference in p-value [((1−p_2_) * d_2_) − ((1−p_1_) * d_1_); d = 1 for increased and −1 for decreased genes], relative change (average log_2_ fold change of each sample vs its corresponding Day 0 value) and absolute change (difference in mean abundance) and sorted by average rank across the 3 methods. log_2_ fold change in transcripts per million (TPM) was calculated for each sample versus its corresponding value on Day 0, with the average shown; outliers are highlighted in yellow. (**d**) Bar charts for *CEBPD, DDIT4* and *KCNE3* in males (top) and females (bottom). log_2_ fold change was calculated for each sample versus its corresponding value on Day 0, with the average shown. Error bars: ± standard deviation, and *, ** and *** denote p < 0.05, 0.01 and 0.001 by Welch’s t-test, respectively. Average TPM on Day 0 is shown in parentheses next to symbols.

We selected *CEBPD, DDIT4* and *KCNE3* for further investigation and noted baseline sex differences in all 3 genes that disappeared following the 2^nd^ BNT162b2 dose. Before vaccination, *CEBPD* and *DDIT4* were ∼twice as abundant in males, but the 2-fold increase on Day 22 in females only resulted in similar levels between sexes (Fig. 4d). The opposite trend was seen with *KCNE3*, which was 50% less abundant in males on Day 0 but doubled by Day 42 in males only. Finally, BRCA1-associated ring domain 1 (*BARD1*) was the only gene that increased significantly in males and decreased significantly in females following vaccination (Fig. 5a), mainly in HPCs (Fig. 5b). Interestingly, even though most samples contained a very small number of Cluster 8 cells there was a strong, negative correlation between Cluster 8 cells and CD14^+^ monocytes (Fig. 5c) with highly similar r, R^2^ and p-values between sexes. We observed a significant negative correlation between monocytes and T cells (Fig. 5d).

**Fig. 5.**
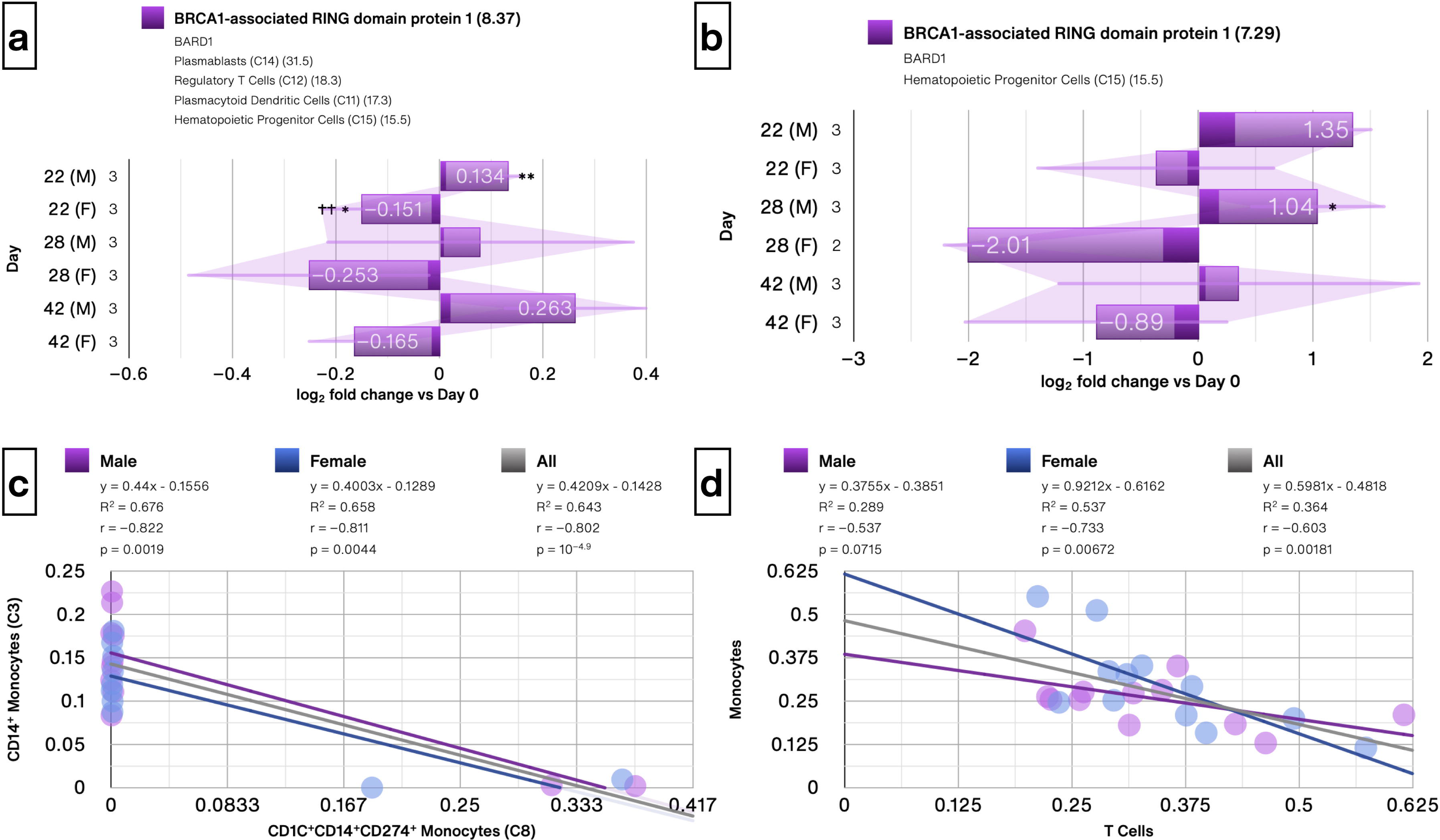
*BARD1* is a sexually dimorphic gene in HPCs and significant correlations between specific immune cell types. Bar charts shown for (**a**) all cells and (b) in hematopoietic progenitor cells showing *BARD1* was the only gene that was significantly increased in males and significantly decreased in females on Day 22. log_2_ fold change was calculated for each sample versus its corresponding value on Day 0, with the average shown. Error bars: ± standard deviation, and *, ** and *** denote p < 0.05, 0.01 and 0.001 by Welch’s t-test, respectively. (**c**) Cluster 8 cells showed a significant negative correlation with CD14^+^ monocytes in males and females independently. Samples from Days 0, 22, 28 and 42 were included. Linear trendlines, or an n^th^-degree polynomial trendline if its goodness-of-fit is either 50% greater than, or if it explains at least half of the variance not explained by the (n – 1)^th^-degree polynomial, were fit to the data. R^2^, r and p denote goodness-of-fit, Pearson’s correlation coefficient and significance of the correlation, respectively. (**d**) A strong negative correlation was noted between Monocytes and T cells that was significant in females but did not reach statistical significance in men.

## Discussion

To our knowledge, this is the first study to analyze baseline and BNT162b2 mRNA vaccine-induced changes in the single cell transcriptome of immune cells in healthy women and men who were also COVID-19 negative.^10^ During our analysis for presences of sex differences in the transcriptome after vaccination, we noted a discrepancy in the data reporting of the original study, and alerted Nature Editors and the authors. A correction has since been published.^16^ We have several novel key findings-T cell and monocytes displayed the greatest numbers of sexually dimorphic genes in single cell transcriptome analysis (4,619 and 4,346, respectively) while HPCs displayed the lowest (1,439), possibly due to low abundance. Baseline platelet counts were greater in females than in males, but this difference disappeared after mRNA vaccination. Increases in platelet counts and persisting platelet activation have also been reported in COVID-19 survivors.^17^ *BARD1*, encoding a tumor suppressor protein, was the only gene that increased significantly in males and decreased significantly in females following vaccination, driven by expression changes in HPCs.

Previously, BNT162b2 mRNA vaccination has been shown to stimulate innate immune responses after the second dose. Sex differences in immune cell and cytokines have been reported in COVID-19 patients.^18, 19^ Cluster 8 cells were identified by Arunachalam et al.^10^ as a novel monocyte population uniquely induced after the second mRNA vaccine dose. We report that C8 cells also expressed high levels of *APOBEC3A*, which promotes replication, propagation, and evolution of SARS-CoV-2.^20, 21^ In men, C8 cells displaced classical monocytes on Day 22 and mostly disappeared by Day 28, whereas they remained abundant in females through Day 28. The importance of C8 cells in vaccine-induced immunity and/or AEs remains to be fully elucidated.

Female NK cells were recently reported to exhibit decreased fitness but increased cell killing capacity and increased expression of cytolytic proteins compared to males, and it was suggested that these differences are mediated by increased expression of *KDM6A*.^22^ We also found that *KDM6A* expression was increased in females (3.3-fold), therefore the reason for decreased expression of NK effector genes is unclear. *KDM6A*, an X-linked gene, escapes inactivation and is shown to be a key player in autoimmune diseases, such as multiple sclerosis. A significant negative correlation between *KDM6A* and several hallmark NK cell genes including *CST7, NKG7* and *PRF1* was present in female but not male samples, with CST7 exhibiting the strongest correlation. However, it was also recently reported that NK cell gene expression is affected by age, with younger females expressing slightly less NKG7 than younger males whereas older males express significantly more NKG7 than older females^23^. Since the female group in this dataset was younger (average age females: 29, males: 40) our findings are consistent with these results. Moreover, the outlier of age 54 in the male group may contribute to the high variance in NK cell count in males. The observed negative relationship between KDM6A and NK effector genes warrants further investigation.

*IFI30*, the top changed gene following Dose 2 (after excluding platelets) was reported to be associated with COVID-19 severity and specifically induced during infection with the Delta strain but not the original Wuhan-like SARS-CoV-2 strain.^24^ *IFI30* induction was also observed in HIV-1 infection but not COVID-19, based on samples collected in China during a time period when the Wuhan-like SARS-CoV-2 strain was circulating.^25^ Other genes such as *IRF1*^26^, *PSME1*^27^ and *IFITM3*, whose expression was increased in our analysis, are known to correlate with markers of COVID-19 severity. *IFITM3* inhibits SARS-CoV-2 replication in HEK293T cells, but it was reported to be essential for infection of human lung cells and greatly promoted the production of infectious virus.^28^ On the other hand, vaccination also beneficially modulated genes known to restrict SARS-CoV-2 infection including *LY6E*^29^ and *AAK1* (#1 decreased).^30^

In our analysis of VAERS data, a 4-fold higher incidence in myocarditis AE in men was noted. In younger men, the risk of myocarditis after the 2^nd^ dose of the mRNA-1273 vaccine is reported.^31^ Voltage-gated potassium (K(V)) channel subunits are encoded by *KCNE* family of genes. In cardiomyopathic heart tissues, *KCNE3* expression is increased and a balanced expression of (K(V)) subunits is required for normal cardiac function.^32^ Gain-of-function mutations in KCNE3 are associated with Brugada syndrome,^33^ and forced overexpression of *KCNE3* in myocytes resulted in shortened QTc interval and accelerated cardiac repolarization.^34^ Brugada syndrome is a condition characterized by electrocardiographic changes including accentuated J waves and ST-segment elevation, resulting in increased risk of arrhythmias and sudden cardiac death, that can be unmasked by natural SARS-CoV-2 infection.^35^ Myocardial effects of COVID-19 mRNA vaccination in healthy individuals also include significant decreases in maximum corrected QT interval (QTc max)^36^ and QTc interval.^37^ ST-segment elevation was noted in several COVID-19 mRNA vaccine-induced myocarditis case reports,^38^ and prominent J waves were observed in one case report.^39^ Thus, vaccine-induced electrocardiographic abnormalities show at least a partial overlap with the reported effects of KCNE3 overexpression. The increased expression of *KCNE3* specifically in males after the 2^nd^ BNT162b2 dose may indicate its involvement in the pathogenesis of cardiovascular AEs in males and warrants further investigation.

DNA damage inducible transcript 4 (*DDIT4*) modulates cellular metabolic function in an mTOR-dependent manner. Elevated DDIT4 expression is associated with worse outcomes for many cancers.^40^ The glucocorticoid-responsive transcription factors CCAAT/enhancer binding proteins delta (CEBPD) and CEBPB regulate transcription of many cytokines and have been described as amplifiers of IL6 production in various cell types.^41^ Female-specific increases in these genes may contribute to higher AE rates in females. Limitations of this study are low sample size, incomplete data for the first vaccine dose, age differences between male and female participants and unavailability of other clinical data associated with these individuals such as adverse events.

Additional studies are needed to validate these findings. This sexual dimorphism in single cell transcriptome might be potential underlying reasons for adverse events such as fatigue, cardiovascular, autoimmune, and neurological symptoms. Finally, biological sex must be considered as a variable during analysis, as the aggregation of data from both sexes may obscure significant, sex-specific changes.

## Institutional Review Board Statement

The study did not require ethical approval.

## Data Availability Statement

All data are contained within the manuscripts or supplemental data. Original raw data can be downloaded at: https://www.nature.com/articles/s41586-021-03791-x

