## Supplemental Figure legends and Tables 1-5 for "BNT162b2 mRNA vaccine-induced sex differences in the single-cell transcriptome of peripheral blood mononuclear cells in healthy adults"

### Supplementary Table Legends

**Supplementary Table S1**. Genes that differed between males and females in most cell types, sorted by ascending average p-value across the tested cell types (B cells, Dendritic Cells (DCs), Hematopoietic Progenitor Cells (HPCs), Monocytes, Natural Killer (NK) cells and T cells), with the expression ($log_2$ fold change vs male) and p-value given for each cell type, in addition to the averages across all cell types.

**Supplementary Table S2**. Genes that differed between males and females on most sample days, sorted by ascending average p-value across the tested days (0, 22, 28 and 42), with the expression ($log_2$ fold change vs male, TPM) and p-value given for each day, in addition to the averages across all days.

**Supplementary Table S3**. Genes that differed between Day 22 and Day 0 (sex-aggregated), including platelets, sorted by impact score (average rank across significance [p], relative change and absolute change), with the normalized control abundance (Day 0, TPM), expression ($log_2$ fold change Day 22 vs Day 0) and p-value given.

**Supplementary Table S4**. Genes that differed between Day 22 and Day 0 (sex-aggregated), excluding platelets, sorted by impact score (average rank across significance [p], relative change and absolute change), with the normalized control abundance (Day 0, TPM), expression ($log_2$ fold change Day 22 vs Day 0) and p-value given.

**Supplementary Table S5**. 200 genes with the greatest differences in response to vaccination (Day 22 vs Day 0) between males and females. Genes were ranked according to difference in p-value [$((1-p_2) * d_2) - ((1-p_1) * d_1)$; d = 1 for increased and −1 for decreased genes], relative change (average $log_2$ fold change of each sample vs its corresponding Day 0 value) and absolute change (difference in mean abundance), with male and female values given for each gene, sorted by average rank across the 3 methods. Gene expression was normalized using TPM.

### Supplementary Figure Legends

**Fig. S1**. **Comparison of normalization modes**. (**a**) Volcano plots showing differences in gene expression for females versus males, in all cell types and all included time points (Day 0, 22, 28, 42), using different normalization modes, and (**b**) including or excluding *XIST*$^{-}$ female cells.

**Fig. S2**. **Prevalence of *XIST* and *PPBP* in each cell cluster, and sex differences in total transcript count**. (**a**) *XIST* transcripts were detected in roughly 80% of female cells: bar chart showing the percentage of female cells positive for *XIST* in each cluster. The cell types with greatest abundance are shown above the bar charts. The filled fraction of a bar represents the percentage of samples positive for *XIST*. (**b**) 98% of cells in the Natural Killer T cell cluster were positive for the platelet marker *PPBP*: bar chart showing the percentage of cells positive for *PPBP* in each cluster.

**Fig. S3**. **Cluster 8 cells expressed the highest levels of *APOBEC3A*, and prevalence of *NKG7* in each cluster**. (**a**) Bar chart showing transcripts per positive cell (TPPC) of *APOBEC3A* in type 2 classical Dendritic Cells (cDC2s), CD14$^{+}$ and CD16$^{+}$ monocytes, and cluster 8 cells; other cell types only expressed marginal levels of *APOBEC3A*. The filled fraction of a bar represents the percentage of cells positive for *APOBEC3A*. Labels next to bars indicate the number of samples (n) in each group. All 7 time points were included. *, **, and *** denote $p < 0.05$, 0.01 and 0.001 versus other immune cells by Welch's t-test; $^{\dagger}$, $^{\dagger\dagger}$ and $^{\dagger\dagger\dagger}$ denote $p < 0.05$, 0.01 and 0.001 versus the previous group (one bar above). Error bars represent the standard deviation. (**b**) *NKG7* transcripts were mainly detected in Natural Killer (NK) cells and mature CD8$^{+}$ T cells: bar chart showing the percentage of cells positive for *NKG7* in each cluster.

**Fig. S4**. **Natural Killer (NK) cell markers and their negative correlation with Lysine-specific demethylase 6A (UTX/*KDM6A*)**. (**a**) Top 10 NK cell markers relative to the other cell types from Fig. 2, with ranks shown. (**b**) Scatter plots showing negative correlations between *CST7*, *NKG7* and *PRF1* with *KDM6A* in male (purple) and female (blue) NK cells, separately and combined (gray trendline). Samples from Days 0, 22, 28 and 42 were included. Linear trendlines, or an $n^{th}$-degree polynomial trendline if its goodness-of-fit is either 50% greater than, or if it explains at least half of the variance not explained by the $(n – 1)^{th}$-degree polynomial, were fit to the data. $R^2$, r and p denote goodness-of-fit, Pearson's correlation coefficient and significance of the correlation, respectively.

Supplementary Table 1

| Position | Symbol | Average Xpr | Average p | Xpr (B) | p (B) | Xpr (DCs) | p (DCs) | Xpr (HPCs) | p (HPCs) | Xpr (Mono) | p (Mono) | Xpr (NK) | p (NK) | Xpr (T) | p (T) |
|---|---|---|---|---|---|---|---|---|---|---|---|---|---|---|---|
| #1 | XIST | 1 | 0 | 19.06569 | 0 | 19.36197 | 0 | 19.60673 | 0 | 19.22787 | 0 | 19.63304 | 0 | 19.09735 | 0 |
| #2 | TSIX | 4.137411 | 8.01715E-09 | 5.175411 | 4.05E-11 | 3.33357 | 1.57E-08 | 13.73554 | 0 | 3.878696 | 1.81E-08 | 5.482751 | 5.76E-09 | 5.954039 | 8.54E-09 |
| #3 | MT-ND4L | 1.738995 | 1.81955E-08 | 1.348642 | 2.32E-10 | 2.077913 | 7.39E-11 | 1.679174 | 4.37E-08 | 2.45522 | 1.57E-10 | 1.233398 | 0.000000065 | 1.639624 | 5.24E-11 |
| #4 | HNRNPU | 0.626732 | 1.93093E-06 | 0.5605818 | 1.56E-12 | 0.709305 | 0.000000185 | 0.5968718 | 0.0000057 | 0.6353956 | 0.000000285 | 0.6177456 | 0.00000542 | 0.6404926 | 3.31E-11 |
| #5 | SH3GLB1 | 0.4745818 | 3.76389E-06 | 0.5487656 | 1.77E-08 | 0.4055332 | 0.00000376 | 0.665487 | 0.0000164 | 0.3639225 | 1.66E-09 | 0.4563752 | 0.00000117 | 0.4074075 | 0.0000012 |
| #6 | ZC3H7A | 0.7768181 | 5.02481E-06 | 0.8546084 | 6.44E-08 | 0.6847191 | 0.0000014 | 1.059371 | 0.0000022 | 0.4844361 | 0.00000506 | 0.7051322 | 0.0000214 | 0.8726414 | 6.84E-09 |
| #7 | COPS2 | 0.7625681 | 7.18863E-06 | 0.8452247 | 0.00000528 | 0.6817959 | 0.000000069 | 0.7594181 | 0.0000358 | 0.6928197 | 9.52E-08 | 0.8291993 | 0.000000143 | 0.7669507 | 0.00000172 |
| #8 | NUFIP2 | 0.8196898 | 7.61317E-06 | 0.9497752 | 1.99E-09 | 0.9181516 | 0.000000026 | 0.7829479 | 0.000021 | 0.6646061 | 0.0000034 | 0.749646 | 0.0000212 | 0.8530117 | 8.6E-09 |
| #9 | ZMYM4 | 0.9479666 | 1.02448E-05 | 1.001876 | 2.11E-08 | 0.7900097 | 0.00000873 | 1.208485 | 0.00000332 | 0.800837 | 0.00000334 | 0.8276366 | 0.000046 | 1.058955 | 0.000000016 |
| #10 | SETD2 | 0.9739417 | 1.03898E-05 | 1.142814 | 3.23E-12 | 1.029402 | 2.37E-10 | 0.9146989 | 0.0000507 | 0.7510262 | 7.23E-11 | 1.002939 | 0.0000116 | 1.00277 | 3.13E-09 |
| #11 | CSGALNACT2 | 1.049669 | 1.51728E-05 | 1.298775 | 7.08E-08 | 0.7853774 | 0.0000413 | 1.054763 | 0.0000474 | 0.77321 | 0.00000164 | 1.105063 | 0.000000624 | 1.280826 | 1.81E-09 |
| #12 | SMC5 | 0.6685104 | 1.54997E-05 | 0.7099566 | 1.43E-10 | 0.7005845 | 8.05E-11 | 0.6481916 | 0.0000906 | 0.5909591 | 0.00000133 | 0.7442495 | 0.00000111 | 0.617121 | 2.52E-11 |
| #13 | CCNL1 | 0.8614123 | 1.9368E-05 | 0.9495694 | 8.71E-08 | 0.9477144 | 2.75E-08 | 0.7825439 | 0.000115 | 0.7502007 | 0.000000708 | 0.7462307 | 1.33E-08 | 0.9922147 | 7.21E-12 |
| #14 | NCL | 0.5537126 | 1.9505E-05 | 0.5057466 | 1.09E-09 | 0.5409257 | 0.000046 | 0.5468787 | 0.0000702 | 0.5352722 | 1.48E-08 | 0.618663 | 5.45E-08 | 0.5747896 | 0.000000754 |
| #15 | KAT7 | 0.7462729 | 2.07594E-05 | 0.7899 | 0.00000128 | 0.6842789 | 0.0000061 | 0.8697005 | 0.000112 | 0.6358342 | 0.000000153 | 0.7527731 | 0.00000413 | 0.7451507 | 0.000000702 |
| #16 | SHOC2 | 0.6366872 | 3.26617E-05 | 0.7720329 | 8.54E-13 | 0.6511041 | 0.0000265 | 0.7383315 | 0.000169 | 0.5427113 | 2.75E-10 | 0.5775787 | 0.00000039 | 0.5383649 | 9.01E-10 |
| #17 | ATP9B | 1.169691 | 3.27918E-05 | 0.8809133 | 0.00000373 | 1.076459 | 0.0000274 | 1.353762 | 0.00000468 | 1.355306 | 0.000000137 | 1.138865 | 0.000161 | 1.212842 | 6.79E-08 |
| #18 | HIST1H1E | 2.777198 | 3.28633E-05 | 2.55699 | 0.0000364 | 2.820762 | 0.0000147 | 3.136767 | 0.00000547 | 3.101974 | 0.00000688 | 2.563688 | 0.0000604 | 2.483005 | 0.0000733 |
| #19 | JPX | 0.8639147 | 3.31915E-05 | 0.9153743 | 3.39E-11 | 0.9390532 | 0.000169 | 0.8150815 | 0.0000304 | 0.5697781 | 0.000000071 | 0.8885147 | 4.11E-08 | 1.055686 | 4.11E-12 |
| #20 | NORAD | 0.74274 | 3.39338E-05 | 0.8216306 | 0.000017 | 0.8407976 | 0.0000107 | 0.6839065 | 0.000166 | 0.4911447 | 0.00000557 | 0.7908647 | 7.95E-08 | 0.8280956 | 0.00000475 |
| #21 | ARL5B | 0.9950359 | 3.49237E-05 | 1.1363 | 0.00000482 | 1.233978 | 0.000000997 | 0.8247769 | 0.0000656 | 1.003042 | 0.00000114 | 0.8448465 | 0.000137 | 0.9272713 | 0.000000401 |
| #22 | HIST2H2AC | 0.9896931 | 3.76814E-05 | 1.146124 | 4.74E-08 | 1.103887 | 7.97E-09 | 1.030252 | 0.0000778 | 1.055108 | 8.85E-09 | 0.7356486 | 0.000148 | 0.8671384 | 0.000000318 |
| #23 | PHF3 | 0.6872429 | 4.59071E-05 | 0.7371275 | 1.62E-10 | 0.7095309 | 0.00000181 | 0.6615888 | 0.0000903 | 0.4723368 | 0.0000921 | 0.7395299 | 0.0000912 | 0.8033435 | 1.24E-09 |
| #24 | PTPN11 | 0.73029 | 4.79986E-05 | 0.7952739 | 3.97E-08 | 0.7169584 | 0.00000148 | 0.7816716 | 0.000285 | 0.5506937 | 0.000000588 | 0.7694364 | 0.000000412 | 0.7677057 | 1.22E-10 |
| #25 | SFPQ | 0.6111602 | 4.93508E-05 | 0.6798289 | 1.96E-08 | 0.5548051 | 7.32E-08 | 0.5366966 | 0.000113 | 0.5520766 | 1.4E-09 | 0.7063917 | 0.000181 | 0.6371622 | 0.00000229 |
| #26 | MAP4 | 0.6624064 | 5.17056E-05 | 0.6679608 | 0.000000027 | 0.6787749 | 0.000254 | 0.9121447 | 0.0000385 | 0.6155561 | 1.28E-08 | 0.638797 | 0.000000679 | 0.4612049 | 0.0000168 |
| #27 | CAPRIN1 | 0.6708141 | 5.97237E-05 | 0.7365272 | 1.02E-08 | 0.588303 | 0.0000675 | 0.6229219 | 0.000287 | 0.5812108 | 0.00000108 | 0.7875672 | 0.00000304 | 0.7083546 | 7.71E-09 |
| #28 | ASCC3 | 0.9935634 | 6.02742E-05 | 1.056074 | 0.000000489 | 0.9923681 | 0.000137 | 1.194987 | 0.00014 | 0.8505241 | 0.0000496 | 0.9553954 | 0.0000348 | 0.9120311 | 6.43E-10 |
| #29 | ATXN7L3B | 0.7396785 | 6.4999E-05 | 0.8006827 | 0.000000784 | 0.8047903 | 0.000206 | 0.6039441 | 0.000181 | 0.4866651 | 0.00000189 | 0.8828272 | 0.000000127 | 0.8591618 | 0.000000651 |
| #30 | HIST1H1D | 1.781648 | 6.6715E-05 | 1.582044 | 0.0000156 | 1.816372 | 0.00000472 | 1.973247 | 0.000168 | 2.076264 | 0.00000101 | 1.618276 | 0.000204 | 1.623685 | 0.00000675 |
| #31 | VPS13C | 0.9907282 | 7.08618E-05 | 1.028602 | 0.000000571 | 1.028843 | 0.0000034 | 0.8148028 | 0.000418 | 0.995104 | 0.00000256 | 1.047064 | 0.000000428 | 1.029952 | 4.23E-14 |
| #32 | OPA1 | 0.874954 | 7.9269E-05 | 0.8374331 | 6.2E-11 | 0.790635 | 2.91E-10 | 0.9330432 | 0.000471 | 0.8418681 | 1.49E-09 | 0.9039529 | 0.00000452 | 0.9427918 | 8.45E-09 |
| #33 | TNKS2 | 0.7058949 | 8.42429E-05 | 0.7367636 | 0.000000182 | 0.5960616 | 0.00000313 | 0.9313398 | 0.000493 | 0.5402245 | 0.00000927 | 0.7952042 | 1.18E-09 | 0.6357758 | 2.87E-08 |
| #34 | HIST1H1C | 1.259207 | 8.48158E-05 | 1.138656 | 0.0000368 | 1.278559 | 0.000102 | 1.59409 | 0.0000147 | 1.115115 | 0.000000273 | 1.266473 | 0.000211 | 1.162348 | 0.000145 |
| #35 | YEATS2 | 0.8863804 | 8.51153E-05 | 0.7292559 | 0.00000713 | 0.779251 | 0.0000938 | 1.034891 | 0.000233 | 0.6473017 | 0.0000499 | 1.280821 | 0.000126 | 0.8467628 | 0.000000166 |
| #36 | CSNK1G3 | 0.7860781 | 9.00864E-05 | 0.6923007 | 6.58E-08 | 0.9539455 | 0.00037 | 0.8342343 | 0.000168 | 0.5781075 | 0.00000139 | 0.8549563 | 0.00000107 | 0.8029244 | 3.42E-08 |
| #37 | MYSM1 | 0.8126996 | 0.000102927 | 0.9349234 | 0.000000122 | 0.8411331 | 9.22E-08 | 0.9161348 | 0.000611 | 0.6721169 | 0.00000579 | 0.7209931 | 0.000000148 | 0.790896 | 2.88E-09 |
| #38 | LAMP2 | 0.5757272 | 0.000108117 | 0.4489792 | 0.00016 | 0.6372803 | 0.00000114 | 0.9864692 | 0.0000455 | 0.3500896 | 0.000305 | 0.6360206 | 0.000000854 | 0.3955242 | 0.000136 |
| #39 | KPNA4 | 0.4974007 | 0.000115421 | 0.4712185 | 0.00015 | 0.4451173 | 0.0000224 | 0.7795013 | 0.000108 | 0.3923908 | 0.0000964 | 0.4867723 | 0.0000356 | 0.409404 | 0.000281 |
| #40 | CHD1 | 0.8063545 | 0.000121242 | 0.8387514 | 3.36E-11 | 0.6272312 | 0.000526 | 0.9352122 | 0.0000144 | 0.8998399 | 8.07E-09 | 0.7335235 | 0.000178 | 0.8035689 | 0.00000862 |
| #41 | OAZ1 | -0.3389532 | 0.000126014 | -0.3061354 | 0.000000269 | -0.2404471 | 0.00000697 | -0.3595101 | 0.000634 | -0.3206997 | 2.39E-09 | -0.4037269 | 0.000115 | -0.4031997 | 3.41E-08 |
| #42 | TM9SF3 | 0.6916225 | 0.000134132 | 0.7066922 | 0.0000145 | 0.8299253 | 0.0000849 | 0.7663803 | 0.000667 | 0.5226242 | 0.000000141 | 0.6297941 | 0.000036 | 0.6943188 | 0.0000024 |
| #43 | MT-ND6 | 1.054522 | 0.000134711 | 0.9355355 | 0.00000267 | 1.157272 | 0.00026 | 0.9801259 | 0.000474 | 1.171001 | 6.27E-11 | 0.9838755 | 0.000071 | 1.099323 | 0.000000474 |
| #44 | WBP11 | 0.5768917 | 0.000137376 | 0.6564457 | 0.00000137 | 0.5694418 | 3.08E-08 | 0.6072953 | 0.000822 | 0.4730789 | 2.41E-12 | 0.5976464 | 0.000000871 | 0.5574421 | 3.45E-08 |
| #45 | HNRNPA3 | 0.4329304 | 0.000137805 | 0.3893276 | 1.82E-08 | 0.4264983 | 0.000137 | 0.4283307 | 0.00068 | 0.4599177 | 0.00000014 | 0.4591884 | 0.0000102 | 0.4343199 | 7.11E-11 |
| #46 | DDX3X | 1.041315 | 0.00015411 | 1.258553 | 6.39E-16 | 1.178321 | 4.75E-12 | 0.6394026 | 0.000925 | 0.8405129 | 0.000000023 | 1.093033 | 4.35E-11 | 1.238066 | 1.26E-13 |
| #47 | ZNF644 | 0.6262162 | 0.000157837 | 0.5992605 | 1.65E-08 | 0.671417 | 0.0000327 | 0.7216852 | 0.0004 | 0.5242076 | 0.000000385 | 0.5473162 | 0.000514 | 0.6934109 | 0.000000101 |
| #48 | NIN | 0.9051104 | 0.000172744 | 0.8392215 | 1.76E-08 | 1.032371 | 0.00000368 | 0.8729424 | 0.00103 | 0.7053468 | 8.81E-09 | 0.9864899 | 8.76E-08 | 0.9942906 | 6.79E-09 |
| #49 | TOP2B | 0.6828294 | 0.000177329 | 0.5216276 | 0.00039 | 0.7942043 | 0.0000599 | 0.5890897 | 0.00051 | 0.6940985 | 0.00000012 | 0.806603 | 0.000101 | 0.6913536 | 0.00000261 |
| #50 | PIBF1 | 1.092037 | 0.000180104 | 1.008083 | 1.13E-09 | 1.283582 | 0.000199 | 1.13359 | 0.000803 | 0.9182611 | 0.0000674 | 1.088149 | 0.0000108 | 1.120556 | 4.06E-08 |
| #51 | MT-ATP8 | 2.278959 | 0.000184437 | 2.118186 | 0.0000448 | 2.414868 | 0.0000481 | 2.16268 | 0.000372 | 2.751953 | 0.000305 | 1.860201 | 0.000256 | 2.365867 | 0.0000812 |
| #52 | COPA | 0.5020926 | 0.000189955 | 0.6054023 | 0.00000423 | 0.4508903 | 0.0000326 | 0.6881952 | 0.000605 | 0.2726679 | 0.00000468 | 0.5417145 | 0.000103 | 0.4536851 | 0.00039 |
| #53 | TRPM7 | 0.5909828 | 0.000208568 | 0.4896996 | 0.00000433 | 0.5169407 | 0.000322 | 0.7804221 | 0.000867 | 0.486757 | 0.00000235 | 0.6461639 | 0.0000556 | 0.6259138 | 9.33E-10 |
| #54 | SACS | 1.008705 | 0.000222186 | 0.9898931 | 0.0000113 | 1.145621 | 0.0000385 | 1.09681 | 0.00102 | 0.9239425 | 1.5E-09 | 0.9791488 | 0.000265 | 0.9168176 | 4.89E-09 |
| #55 | SMC1A | 0.7184093 | 0.000222988 | 0.6607568 | 0.00000935 | 0.6354704 | 0.00000182 | 0.7348503 | 0.00133 | 0.576176 | 2.36E-10 | 0.7953284 | 0.000000297 | 0.9078738 | 1.29E-09 |
| #56 | HNRNPA2B1 | 0.3700545 | 0.000223033 | 0.3054815 | 5.74E-08 | 0.4687866 | 0.00000263 | 0.2886043 | 0.00132 | 0.4375405 | 0.0000189 | 0.3601172 | 8.73E-09 | 0.3597968 | 6.09E-10 |
| #57 | STX16 | 0.6744871 | 0.00022371 | 0.6989229 | 1.73E-08 | 0.6692108 | 0.00000116 | 0.5972284 | 0.0013 | 0.4682717 | 0.000000658 | 0.7082726 | 0.0000406 | 0.9050164 | 2.09E-10 |
| #58 | CTDSPL2 | 0.8149147 | 0.000225494 | 1.011626 | 3.63E-10 | 0.8204538 | 4.05E-10 | 0.729056 | 0.00133 | 0.661474 | 2.27E-09 | 0.7364206 | 0.0000205 | 0.9304583 | 2.69E-11 |
| #59 | PAFAH1B1 | 0.4929215 | 0.00024094 | 0.5552712 | 4.05E-08 | 0.388546 | 0.00000262 | 0.733042 | 0.000823 | 0.3173343 | 0.000000437 | 0.4453276 | 0.000619 | 0.5180081 | 4.44E-08 |
| #60 | RANBP6 | 0.7570147 | 0.000246817 | 0.7576539 | 0.0000975 | 0.9036112 | 0.000022 | 0.9274195 | 0.000112 | 0.3679053 | 0.00102 | 0.7838327 | 0.000231 | 0.8016657 | 0.00000081 |
| #61 | BCLAF1 | 0.5175156 | 0.00025275 | 0.6301344 | 9.99E-09 | 0.5631589 | 0.000000618 | 0.3910497 | 0.00151 | 0.4489621 | 3.71E-11 | 0.5407473 | 0.00000217 | 0.5310412 | 0.000000158 |
| #62 | EIF1AX | 0.6235543 | 0.00026188 | 0.7211507 | 1.95E-09 | 0.595131 | 0.00000564 | 0.3283186 | 0.00156 | 0.5539895 | 0.00000122 | 0.7797683 | 4.12E-10 | 0.7629674 | 5.67E-10 |
| #63 | TLK2 | 0.7999503 | 0.000269278 | 0.7853539 | 0.00000235 | 0.7853361 | 0.00000669 | 0.8410854 | 0.000772 | 0.6761109 | 0.00000448 | 0.8544971 | 0.00083 | 0.8573184 | 0.000000162 |
| #64 | FUBP1 | 0.6108829 | 0.000274313 | 0.69375 | 0.000000476 | 0.5820965 | 0.000569 | 0.5514432 | 0.00107 | 0.5135343 | 0.00000357 | 0.6569128 | 0.000000857 | 0.6675609 | 3.76E-11 |
| #65 | RSRC2 | 0.4758865 | 0.000276082 | 0.6868969 | 0.000000244 | 0.372802 | 0.0000906 | 0.4618306 | 0.00145 | 0.4254324 | 0.00000174 | 0.3924677 | 0.000115 | 0.5158894 | 0.0000041 |
| #66 | ATE1 | 0.6964662 | 0.000277969 | 0.4699788 | 0.00116 | 0.6413707 | 0.000319 | 1.476268 | 0.00000665 | 0.3743197 | 0.000157 | 0.590014 | 0.0000238 | 0.6268457 | 5.25E-08 |
| #67 | DMTF1 | 0.767476 | 0.000288224 | 0.7495304 | 0.00000106 | 0.8646956 | 0.000997 | 0.9309948 | 0.00057 | 0.5904739 | 0.00000313 | 0.6520924 | 0.000158 | 0.8170691 | 3.68E-12 |
| #68 | SMARCC1 | 0.4947976 | 0.000288268 | 0.5386399 | 0.00000155 | 0.4268301 | 0.00000107 | 0.3742402 | 0.00153 | 0.5421657 | 5.04E-10 | 0.4848225 | 0.000192 | 0.602087 | 1.23E-08 |
| #69 | TMOD3 | 0.5053857 | 0.000298927 | 0.5795313 | 0.00000015 | 0.5261766 | 0.000000118 | 0.5705155 | 0.00179 | 0.4962426 | 1.1E-09 | 0.4440418 | 0.00000119 | 0.4158065 | 9.93E-09 |

| | | | | | | | | | | | | | | | |
|---|---|---|---|---|---|---|---|---|---|---|---|---|---|---|---|
| #70 | MARCH7 | 0.6345179 | 0.000303782 | 0.7220091 | 5.06E-08 | 0.653286 | 0.000162 | 0.7482087 | 0.00000291 | 0.3858071 | 0.00166 | 0.6703456 | 0.000000118 | 0.6274511 | 1.42E-08 |
| #71 | UHRF2 | 0.9486197 | 0.000303825 | 0.8359052 | 0.00000684 | 0.858457 | 0.00171 | 1.206249 | 0.0000694 | 0.898672 | 6.19E-08 | 0.885769 | 0.000034 | 1.006666 | 4E-10 |
| #72 | MON2 | 0.7761923 | 0.000307805 | 0.7464661 | 0.000000348 | 0.8882374 | 0.0000647 | 0.88245 | 0.00137 | 0.6206456 | 0.000118 | 0.6540319 | 0.000291 | 0.865323 | 0.00000012 |
| #73 | ZNF326 | 0.6944668 | 0.000322252 | 0.7319281 | 0.0000416 | 0.7537658 | 0.0000316 | 0.5632478 | 0.00143 | 0.6099782 | 0.00000121 | 0.6178475 | 0.000424 | 0.8900332 | 0.000000292 |
| #74 | MT-ND5 | 0.8772968 | 0.000341153 | 0.7733716 | 0.0000331 | 1.137427 | 6.45E-08 | 0.5341178 | 0.00193 | 1.387315 | 1.41E-10 | 0.6529291 | 0.0000439 | 0.7786209 | 0.0000403 |
| #75 | TPR | 0.3651893 | 0.000361539 | 0.3158038 | 0.00000046 | 0.4100041 | 0.00109 | 0.3870233 | 0.000684 | 0.3016603 | 0.00026 | 0.4267928 | 0.000125 | 0.3498516 | 0.00000501 |
| #76 | HP1BP3 | 0.4318256 | 0.000373371 | 0.519568 | 1.91E-10 | 0.414979 | 0.0000749 | 0.4779887 | 0.00115 | 0.3272076 | 0.000262 | 0.338826 | 0.000754 | 0.5123841 | 3.26E-12 |
| #77 | ACIN1 | 0.6252882 | 0.000373802 | 0.7399781 | 0.000000221 | 0.4725474 | 0.00000961 | 0.5768388 | 0.000319 | 0.4374509 | 0.00000793 | 0.7536506 | 0.00191 | 0.7712635 | 4.43E-08 |
| #78 | HIF1A | 0.841485 | 0.000393281 | 0.7394061 | 0.00000535 | 0.566313 | 0.00145 | 0.9611227 | 0.00084 | 0.7921265 | 0.0000118 | 0.909941 | 0.0000526 | 1.080001 | 4.52E-09 |
| #79 | ATP2C1 | 0.7253804 | 0.000394204 | 0.6212487 | 0.000103 | 0.7011162 | 0.00067 | 0.5053961 | 0.00159 | 0.560588 | 0.00000362 | 1.029604 | 0.000000248 | 0.9343296 | 2.69E-08 |
| #80 | KDM5C | 0.8357051 | 0.000403805 | 0.8543339 | 6.85E-08 | 0.6281734 | 0.0000234 | 1.237132 | 0.0000217 | 0.562731 | 0.000000445 | 0.693545 | 0.00238 | 1.038316 | 0.000000538 |
| #81 | CAPN1 | -0.5414057 | 0.000404019 | -0.483407 | 0.000181 | -0.3539281 | 0.00104 | -0.7984473 | 0.0000118 | -0.3818817 | 0.00113 | -0.5779873 | 0.0000614 | -0.6527826 | 0.00000123 |
| #82 | WAPL | 0.6259193 | 0.000421846 | 0.5093935 | 0.000926 | 0.5501719 | 0.0000293 | 0.5385085 | 0.00158 | 0.6935683 | 6.37E-09 | 0.8522461 | 2.44E-08 | 0.6116275 | 5.7E-09 |
| #83 | CD46 | 0.6034193 | 0.000429664 | 0.5727488 | 0.00000818 | 0.7552755 | 1.5E-10 | 0.5554514 | 0.00257 | 0.5398268 | 4.55E-08 | 0.6213585 | 0.000000384 | 0.575855 | 2.04E-08 |
| #84 | ADAM10 | 0.5729141 | 0.000429865 | 0.6363194 | 0.0000208 | 0.5794845 | 1.68E-08 | 0.8282458 | 0.000463 | 0.3489762 | 0.00187 | 0.4654309 | 0.00023 | 0.5790281 | 1.16E-08 |
| #85 | KDM6A | 1.586403 | 0.000438865 | 1.785404 | 5.68E-10 | 1.487586 | 6.26E-11 | 1.202806 | 0.00263 | 1.43056 | 3.25E-11 | 1.732722 | 0.00000126 | 1.87934 | 6.88E-11 |
| #86 | EPC2 | 0.7414187 | 0.000443411 | 0.8257653 | 0.000000158 | 0.8034968 | 0.0000816 | 0.7599509 | 0.00244 | 0.5887488 | 5.14E-08 | 0.5846388 | 0.000137 | 0.8859116 | 6.18E-09 |
| #87 | TMED5 | 0.5451737 | 0.000450206 | 0.7105465 | 2.16E-08 | 0.7841166 | 0.0000471 | 0.7137063 | 0.000967 | 0.4101798 | 5.29E-08 | 0.3669281 | 0.00147 | 0.2855649 | 0.000222 |
| #88 | ANKLE2 | 0.6879434 | 0.00045112 | 0.7152816 | 0.00000562 | 0.7551976 | 5.38E-08 | 0.7804877 | 0.00056 | 0.5270805 | 0.000129 | 0.6926718 | 0.002 | 0.6569412 | 0.00000894 |
| #89 | SH3BGRL3 | -0.5683991 | 0.000455006 | -0.4496395 | 0.000221 | -0.666402 | 0.0025 | -0.572495 | 0.00000592 | -0.3492457 | 5.98E-08 | -0.6986462 | 0.00000493 | -0.6739664 | 0.000000712 |
| #90 | DDHD1 | 0.7181468 | 0.000461566 | 0.6907853 | 0.000000607 | 0.8590387 | 0.000000727 | 0.6620431 | 0.00241 | 0.8014839 | 0.000000612 | 0.6288666 | 0.000353 | 0.6666631 | 5.66E-08 |
| #91 | CAPN7 | 0.7976757 | 0.000471799 | 0.8527848 | 0.000000104 | 0.7657635 | 0.00000733 | 1.145993 | 0.0027 | 0.6085632 | 0.0000049 | 0.6105237 | 0.000122 | 0.8024263 | 1.17E-09 |
| #92 | PDS5A | 0.5676511 | 0.000486552 | 0.4701795 | 0.0000273 | 0.6073756 | 0.000000557 | 0.6321021 | 0.00278 | 0.5715354 | 9.04E-10 | 0.651752 | 0.000131 | 0.4729616 | 0.000000513 |
| #93 | ATP5MG | -0.2586666 | 0.000486683 | -0.2171479 | 0.000192 | -0.2186316 | 0.000307 | -0.3015444 | 0.0000149 | -0.2780793 | 0.000000473 | -0.3657495 | 0.00208 | -0.1708471 | 0.000323 |
| #94 | PTBP2 | 0.9335063 | 0.000488742 | 0.8969557 | 0.000000426 | 1.104523 | 0.000000211 | 0.8536163 | 0.000854 | 0.8606247 | 0.000000904 | 0.7586654 | 0.00208 | 1.126653 | 0.000000368 |
| #95 | RABEP1 | 0.6208853 | 0.000489686 | 0.656542 | 4.78E-08 | 0.4340318 | 0.00276 | 0.80186 | 0.0000625 | 0.5595762 | 0.0000586 | 0.5561125 | 0.0000554 | 0.7171894 | 8.76E-08 |
| #96 | IMPA1 | 0.6364037 | 0.00051759 | 0.5584065 | 0.00049 | 0.8849456 | 0.0000463 | 0.7678868 | 0.00106 | 0.5449238 | 0.0000339 | 0.5455885 | 0.00138 | 0.5166708 | 0.0000972 |
| #97 | USP25 | 0.9853516 | 0.000518027 | 0.8720417 | 0.000284 | 0.8723691 | 0.00000311 | 1.322947 | 0.00266 | 0.9395528 | 0.000146 | 1.050129 | 0.00000971 | 0.8550702 | 0.00000642 |
| #98 | ITGA4 | 0.5082598 | 0.000521681 | 0.5379648 | 0.0000216 | 0.492323 | 0.0000368 | 0.4408444 | 0.00119 | 0.6208934 | 0.000103 | 0.5302062 | 0.00000584 | 0.4273268 | 0.00177 |
| #99 | MT-ND2 | 0.5319585 | 0.000529599 | 0.3454972 | 0.0000462 | 0.8279172 | 0.000000737 | 0.4953536 | 0.00012 | 0.7178213 | 0.0000703 | 0.4123981 | 0.00257 | 0.3927638 | 0.000369 |
| #100 | CPSF6 | 0.4783613 | 0.000535244 | 0.5768581 | 0.00000209 | 0.4927975 | 0.0000137 | 0.4592378 | 0.00311 | 0.3208404 | 0.0000876 | 0.4761581 | 0.0000021 | 0.544276 | 0.000000387 |
| #101 | TET2 | 0.8454929 | 0.000535708 | 1.099226 | 1.36E-08 | 0.8522701 | 0.00000162 | 0.7365959 | 0.00173 | 0.6788123 | 0.00000529 | 0.8066457 | 0.00148 | 0.8994075 | 0.000000087 |
| #102 | TRAPPC1 | -0.3929708 | 0.000546323 | -0.3995955 | 0.000463 | -0.3287215 | 0.0000202 | -0.4367622 | 0.0000888 | -0.3184677 | 0.0000408 | -0.4740761 | 0.00186 | -0.4002019 | 0.00000129 |
| #103 | SPECC1L | 0.8554278 | 0.000571639 | 1.067692 | 6.28E-10 | 0.816428 | 0.0000116 | 1.155178 | 0.0000342 | 0.5247237 | 0.00104 | 0.7967944 | 0.00234 | 0.7717505 | 2.88E-08 |
| #104 | FNDC3A | 0.8022904 | 0.000594218 | 0.9102683 | 1.09E-09 | 0.8165415 | 0.00184 | 0.7862517 | 0.00142 | 0.5567146 | 0.000309 | 0.9055621 | 7.27E-10 | 0.8384045 | 1.09E-08 |
| #105 | SHLD2 | 0.7757848 | 0.00059609 | 0.8857923 | 9.84E-08 | 0.6348291 | 0.000000123 | 1.145286 | 0.00356 | 0.4472513 | 2.39E-08 | 0.9309437 | 0.0000175 | 0.6106061 | 0.00000173 |
| #106 | HIST1H2AK | 1.659321 | 0.0006017 | 1.645049 | 0.000000736 | 1.243198 | 0.0015 | 2.177378 | 0.000959 | 1.764352 | 0.00000172 | 1.490567 | 0.00106 | 1.635383 | 0.0000875 |
| #107 | UBQLN1 | 0.5207334 | 0.000601789 | 0.5456751 | 0.0000533 | 0.4157797 | 0.0000192 | 0.546764 | 0.00354 | 0.5225382 | 9.71E-08 | 0.5218696 | 0.00000011 | 0.5717737 | 8.52E-11 |
| #108 | GPBP1 | 0.5553913 | 0.000607094 | 0.6798295 | 0.0000106 | 0.5103893 | 0.000165 | 0.424119 | 0.00335 | 0.4929171 | 0.0000679 | 0.5430053 | 0.00000121 | 0.6820879 | 0.0000469 |
| #109 | ZBTB1 | 0.8075016 | 0.000612666 | 0.8025892 | 0.000079 | 0.9164893 | 0.00000183 | 0.7291747 | 0.00225 | 0.8599716 | 0.0000134 | 0.8099803 | 0.000345 | 0.7268044 | 0.000985 |
| #110 | GLS | 0.8841522 | 0.000628752 | 0.9516207 | 0.000966 | 1.083356 | 0.000000762 | 0.6452037 | 0.00321 | 0.9923398 | 0.000444 | 0.8599193 | 0.00000831 | 0.7724742 | 0.00000377 |
| #111 | IDE | 0.5886994 | 0.0006351 | 0.5219633 | 0.00212 | 0.623138 | 0.0000207 | 0.7528973 | 0.000597 | 0.513939 | 0.0000115 | 0.5708461 | 0.00105 | 0.5494125 | 0.00000954 |
| #112 | MACF1 | 0.6587499 | 0.000640441 | 0.5703337 | 0.00000344 | 0.614734 | 0.0000153 | 0.6197866 | 0.000961 | 0.6820616 | 0.0000511 | 0.8634452 | 0.00271 | 0.6021383 | 0.0000984 |
| #113 | RIF1 | 0.6260737 | 0.000647647 | 0.5667071 | 0.00000429 | 0.6126332 | 0.00000265 | 0.5492844 | 0.00336 | 0.5706218 | 0.00000217 | 0.8346792 | 0.000515 | 0.6225162 | 0.000000513 |
| #114 | AGFG1 | 0.8766797 | 0.000648155 | 1.046797 | 0.000238 | 0.7242101 | 0.000576 | 0.9331181 | 0.00259 | 0.5953974 | 0.000224 | 0.9254702 | 0.000224 | 1.035085 | 0.0000395 |
| #115 | UHRF1BP1L | 0.7484246 | 0.000650621 | 0.7035676 | 0.000000546 | 0.7737164 | 0.000000179 | 0.8085626 | 0.00369 | 0.6322425 | 0.000000107 | 0.8258887 | 0.00208 | 0.7465699 | 9.91E-08 |
| #116 | PDCD6IP | 0.4118272 | 0.00066398 | 0.3864863 | 0.000128 | 0.4478048 | 0.0000125 | 0.5385016 | 0.000988 | 0.5232691 | 0.00000143 | 0.2866298 | 0.00222 | 0.2882715 | 0.00063 |
| #117 | CASD1 | 0.7433503 | 0.000678429 | 0.5792409 | 0.0015 | 0.6276641 | 0.00186 | 1.09118 | 0.00012 | 0.4910271 | 0.000374 | 0.9812855 | 0.000191 | 0.6897039 | 0.0000338 |
| #118 | UTRN | 0.8570439 | 0.000687856 | 0.7586108 | 7.1E-12 | 0.8896299 | 0.00000601 | 0.9282967 | 0.00036 | 0.9375669 | 6.44E-10 | 0.8619375 | 0.00376 | 0.7662219 | 0.00000062 |
| #119 | PRPF4B | 0.7719608 | 0.000689204 | 0.9206287 | 9.55E-12 | 0.7804739 | 0.000000673 | 0.4568305 | 0.00399 | 0.6281491 | 3.27E-09 | 0.9114679 | 0.000148 | 0.9342146 | 4.34E-09 |
| #120 | MED13 | 0.577624 | 0.000691446 | 0.5966027 | 0.00000757 | 0.5899085 | 0.00000802 | 0.7061997 | 0.00261 | 0.4694025 | 0.0000771 | 0.4340911 | 0.00145 | 0.6695396 | 5.84E-09 |
| #121 | KIF2A | 0.5713318 | 0.000693037 | 0.6791572 | 0.000000293 | 0.4336477 | 0.00383 | 0.6384961 | 0.000246 | 0.6505806 | 0.00000213 | 0.4553157 | 0.0000787 | 0.5707934 | 1.48E-08 |
| #122 | KIAA1586 | 0.8597659 | 0.000695541 | 0.8539686 | 5.02E-08 | 0.8137369 | 0.00000253 | 1.155414 | 0.00000128 | 0.4577783 | 0.00401 | 0.8086371 | 0.000155 | 1.06906 | 5.89E-09 |
| #123 | FYTTD1 | 0.5355209 | 0.000708135 | 0.4659151 | 0.000154 | 0.6641559 | 0.00145 | 0.6156447 | 0.00257 | 0.4922139 | 0.0000389 | 0.5347466 | 0.0000349 | 0.440449 | 9.77E-09 |
| #124 | ARFGEF1 | 0.7739582 | 0.000728558 | 0.726431 | 0.000000019 | 0.6418041 | 9.23E-09 | 0.8679211 | 0.000343 | 0.6398433 | 1.93E-08 | 0.8366528 | 0.00403 | 0.931097 | 2.24E-08 |
| #125 | SF3B1 | 0.5972026 | 0.000730077 | 0.707927 | 5.14E-11 | 0.5680785 | 0.000000118 | 0.4050827 | 0.00438 | 0.5557653 | 0.000000215 | 0.6554786 | 1.48E-08 | 0.6908832 | 1.88E-09 |
| #126 | ZNF254 | 0.6253298 | 0.000738667 | 0.7123597 | 0.000000117 | 0.5621265 | 0.00203 | 0.7135319 | 0.00115 | 0.6276575 | 0.0000124 | 0.440077 | 0.00124 | 0.696226 | 0.000000132 |
| #127 | PANK3 | 0.6762673 | 0.00074781 | 0.6183936 | 0.0000717 | 0.6319353 | 0.000186 | 0.8023233 | 0.00397 | 0.5919838 | 0.000000173 | 0.6353797 | 0.000259 | 0.7775881 | 4.27E-09 |
| #128 | ATP13A3 | 0.6057466 | 0.00075118 | 0.5539436 | 0.000966 | 0.6165489 | 0.0000914 | 0.8411529 | 0.00319 | 0.5320558 | 0.000209 | 0.6487985 | 0.00000183 | 0.4419797 | 0.0000519 |
| #129 | RANBP2 | 0.7815688 | 0.000751385 | 0.7987692 | 0.00000517 | 0.8901542 | 0.000000311 | 0.4922691 | 0.00446 | 0.6814489 | 0.00000126 | 1.070887 | 0.0000344 | 0.7558847 | 0.00000232 |
| #130 | HEATR1 | 0.7243256 | 0.00079504 | 0.6125789 | 0.0000361 | 0.6944389 | 0.000533 | 0.7700158 | 0.00352 | 0.7338661 | 0.000000215 | 0.7462407 | 0.000682 | 0.7888134 | 0.000000214 |
| #131 | BACH1 | 0.7591892 | 0.000812262 | 0.9614887 | 0.000132 | 0.5600752 | 0.00331 | 0.8441042 | 0.000869 | 0.5126341 | 0.000196 | 0.871347 | 0.000366 | 0.8054858 | 0.00000213 |
| #132 | EIF2AK1 | 0.3863528 | 0.000826318 | 0.3634206 | 0.000169 | 0.4035651 | 0.00000747 | 0.5582994 | 0.00017 | 0.3257723 | 0.0000023 | 0.3072005 | 0.0046 | 0.359859 | 0.00000773 |
| #133 | AZIN1 | 0.5771883 | 0.000839533 | 0.7282029 | 0.000000327 | 0.571447 | 0.000492 | 0.7911653 | 0.00294 | 0.4719284 | 0.000000112 | 0.508498 | 0.000223 | 0.391888 | 0.00138 |
| #134 | ELF1 | 0.4704108 | 0.000844324 | 0.5101645 | 0.00013 | 0.4514633 | 0.0000274 | 0.4794108 | 0.0036 | 0.4531582 | 0.00000053 | 0.4941801 | 0.000626 | 0.4340878 | 0.000681 |
| #135 | LCORL | 0.7134552 | 0.000849273 | 0.7495596 | 0.0000143 | 0.8280627 | 0.0000945 | 0.8117918 | 0.00122 | 0.4955006 | 0.0000139 | 0.5639745 | 0.00375 | 0.8318417 | 0.000000818 |
| #136 | SMARCA5 | 0.5674267 | 0.000854504 | 0.6578273 | 4.63E-12 | 0.5153647 | 0.000162 | 0.4860315 | 0.00495 | 0.5260677 | 0.00000109 | 0.6095043 | 0.00000915 | 0.6097649 | 4.16E-11 |
| #137 | ATP5ME | -0.3645571 | 0.000857721 | -0.3499712 | 0.00029 | -0.4238326 | 0.000105 | -0.3478994 | 0.00324 | -0.3174052 | 0.000994 | -0.4384022 | 0.0000165 | -0.309832 | 0.000503 |
| #138 | TPM3 | 0.3421169 | 0.000864117 | 0.3367168 | 0.0000235 | 0.2446256 | 0.000423 | 0.3554489 | 0.00328 | 0.481564 | 1.69E-08 | 0.3702172 | 0.000664 | 0.2641288 | 0.000793 |
| #139 | UQCR11 | -0.3543606 | 0.000865601 | -0.3244833 | 0.00012 | -0.3421776 | 0.00000992 | -0.3392751 | 0.00482 | -0.3254311 | 0.0000734 | -0.4190414 | 0.000174 | -0.3757552 | 0.000000131 |
| #140 | EIF4G2 | 0.5476361 | 0.000880177 | 0.4929659 | 6.54E-08 | 0.6640719 | 0.000185 | 0.4200969 | 0.00509 | 0.4363711 | 0.00000208 | 0.6824386 | 0.000000465 | 0.5898725 | 0.000000063 |

| | | | | | | | | | | | | | | | |
|---|---|---|---|---|---|---|---|---|---|---|---|---|---|---|---|
| **#141** | WDR26 | 0.621975 | 0.000880556 | 0.6407103 | 0.0000433 | 0.6807622 | 0.000000111 | 0.6582733 | 0.00502 | 0.4986169 | 0.0000261 | 0.6081401 | 0.000193 | 0.6453474 | 0.00000172 |
| **#142** | MAPKAP1 | 0.5967956 | 0.000896341 | 0.6724502 | 0.0000516 | 0.6181574 | 0.000371 | 0.7121688 | 0.00344 | 0.4439304 | 0.0000462 | 0.4958954 | 0.00146 | 0.6381715 | 0.0000115 |
| **#143** | CLINT1 | 0.37069 | 0.000910811 | 0.40527 | 0.0000104 | 0.3371608 | 0.00323 | 0.5274815 | 0.00153 | 0.296038 | 0.0000128 | 0.2860779 | 0.000684 | 0.3721116 | 0.00000285 |
| **#144** | UBR3 | 0.6570352 | 0.000913886 | 0.6542076 | 0.0000396 | 0.573566 | 0.000161 | 0.8369578 | 0.000936 | 0.5319575 | 0.0000087 | 0.684191 | 0.00434 | 0.6613315 | 0.0000018 |
| **#145** | ZFC3H1 | 0.6687879 | 0.000921379 | 0.5450646 | 7.09E-08 | 0.8046186 | 3.94E-11 | 0.5215147 | 0.00136 | 0.662599 | 0.00000177 | 0.8258606 | 0.00416 | 0.6530701 | 8.76E-10 |
| **#146** | ACSL4 | 0.6694606 | 0.000921777 | 0.5938925 | 0.00219 | 0.515846 | 0.0000341 | 0.869324 | 0.00329 | 0.5897784 | 0.00000244 | 0.8592813 | 0.00000662 | 0.5886413 | 5.68E-08 |
| **#147** | NR2C2 | 0.6869819 | 0.000925757 | 0.5768846 | 0.0000164 | 0.7748008 | 0.000384 | 1.208244 | 0.00297 | 0.4683839 | 0.000029 | 0.4463567 | 0.00213 | 0.6472213 | 0.000017 |
| **#148** | ZNF267 | 0.9486396 | 0.000930798 | 1.094988 | 0.00353 | 1.02181 | 0.0000119 | 1.062633 | 0.000404 | 0.8040168 | 0.000147 | 0.7436085 | 0.0003 | 0.964781 | 0.0012 |
| **#149** | ZCCHC8 | 0.5794105 | 0.000933457 | 0.5848739 | 0.00000643 | 0.5860561 | 0.00119 | 0.7910781 | 0.00257 | 0.5026502 | 0.0000123 | 0.5143725 | 0.00164 | 0.4974326 | 0.000176 |
| **#150** | RBM33 | 0.5618641 | 0.000958052 | 0.4495662 | 0.0000786 | 0.5707194 | 0.000000534 | 0.7141946 | 0.00523 | 0.4444949 | 0.0000324 | 0.5361939 | 0.000409 | 0.6560153 | 0.00000014 |
| **#151** | TCEA1 | 0.3877375 | 0.000966852 | 0.2801993 | 0.00000447 | 0.333068 | 0.0053 | 0.6685963 | 0.000445 | 0.3343704 | 4.97E-08 | 0.3689357 | 0.0000473 | 0.3412552 | 6.37E-08 |
| **#152** | MSL2 | 0.7921024 | 0.000979533 | 0.8356322 | 0.00000223 | 1.033762 | 0.000151 | 0.6612289 | 0.00543 | 0.519659 | 0.000000322 | 0.8229988 | 0.000295 | 0.879333 | 0.00000228 |
| **#153** | NDUFB4 | -0.3382881 | 0.000997639 | -0.3025117 | 0.00421 | -0.3018763 | 0.0000731 | -0.3537731 | 0.000134 | -0.3699414 | 0.000000203 | -0.4345055 | 0.00157 | -0.2671204 | 0.00000696 |
| **#154** | PTPRC | 0.4493377 | 0.001003896 | 0.3750303 | 0.00000364 | 0.4385609 | 0.00422 | 0.4981326 | 0.0018 | 0.4868187 | 0.00000174 | 0.3948692 | 0.000000747 | 0.5026147 | 2.36E-08 |
| **#155** | ANGEL2 | 0.6552871 | 0.001006022 | 0.6497773 | 0.00000339 | 0.6203164 | 0.000016 | 0.9628418 | 0.00462 | 0.4244086 | 0.0000303 | 0.6703207 | 0.00137 | 0.6040578 | 0.000000541 |
| **#156** | AFTPH | 0.5407271 | 0.001010317 | 0.6676549 | 0.000000571 | 0.4296692 | 0.00151 | 0.7307451 | 0.00329 | 0.3921102 | 0.000786 | 0.5907198 | 0.000082 | 0.4334637 | 0.000396 |
| **#157** | PRKX | 0.8461219 | 0.001022296 | 0.8799823 | 5.38E-10 | 0.9310595 | 7.45E-11 | 0.7464179 | 0.00612 | 0.8854383 | 3.31E-11 | 0.7386056 | 0.00000872 | 0.8952279 | 0.00000235 |
| **#158** | HNRNPK | 0.2216318 | 0.001025557 | 0.1888809 | 0.00000221 | 0.2478271 | 0.000000233 | 0.2233067 | 0.00591 | 0.2084219 | 0.0000396 | 0.2568648 | 0.000174 | 0.2044892 | 0.0000238 |
| **#159** | ZNF43 | 0.6230528 | 0.001036665 | 0.6748523 | 0.000012 | 0.8151081 | 0.00149 | 0.678643 | 0.00284 | 0.4438589 | 0.0000262 | 0.4655063 | 0.00185 | 0.6603484 | 2.34E-08 |
| **#160** | ARHGEF6 | 0.6142485 | 0.001087181 | 0.822902 | 0.000000337 | 0.3441525 | 0.00641 | 0.714053 | 0.0000533 | 0.4508779 | 0.000000923 | 0.6196241 | 0.000056 | 0.7338816 | 0.000000719 |
| **#161** | FNBP1 | 0.3984369 | 0.001103104 | 0.4627519 | 3.06E-09 | 0.2449935 | 0.00318 | 0.4236809 | 0.00324 | 0.3807561 | 0.000201 | 0.4585825 | 9.14E-09 | 0.4198563 | 2.86E-09 |
| **#162** | KAT6A | 0.5727649 | 0.00111237 | 0.5994396 | 0.000000103 | 0.4960652 | 0.0000119 | 0.4867025 | 0.00648 | 0.5596433 | 0.000000737 | 0.6709174 | 0.000179 | 0.6238212 | 2.26E-09 |
| **#163** | LSM14A | 0.3499777 | 0.001116978 | 0.3514875 | 0.000104 | 0.2953068 | 0.0000236 | 0.4805006 | 0.00634 | 0.2584939 | 0.0000223 | 0.3833844 | 0.00016 | 0.3306928 | 0.000051 |
| **#164** | IAH1 | -0.4875495 | 0.001129034 | -0.4405749 | 0.00595 | -0.3784703 | 0.000421 | -0.7002941 | 0.000395 | -0.4863318 | 3.15E-08 | -0.4606734 | 0.0000103 | -0.4589527 | 0.00000106 |
| **#165** | KANSL1L | 0.8629123 | 0.001131042 | 0.7217834 | 0.000108 | 0.7954149 | 0.000174 | 1.031342 | 0.00362 | 0.6872873 | 0.0000121 | 0.8040028 | 0.00285 | 1.137643 | 0.0000154 |
| **#166** | TM9SF2 | 0.5539685 | 0.00116549 | 0.6187418 | 0.00000159 | 0.6860965 | 0.00438 | 0.6311719 | 0.00134 | 0.3797192 | 0.0000268 | 0.4357039 | 0.00125 | 0.5723777 | 0.00000244 |
| **#167** | YWHAZ | 0.3149579 | 0.001169133 | 0.3053183 | 0.000000144 | 0.3620554 | 0.0000428 | 0.3816776 | 0.00508 | 0.3262684 | 3.46E-08 | 0.2925486 | 0.000781 | 0.2218793 | 0.00111 |
| **#168** | GOLGA4 | 0.5916931 | 0.001170449 | 0.4830208 | 0.0000227 | 0.6841574 | 0.000178 | 0.6804552 | 0.000164 | 0.538775 | 3.17E-08 | 0.7072262 | 0.0065 | 0.4565238 | 0.000155 |
| **#169** | TOMM70 | 0.4590163 | 0.001175502 | 0.3956482 | 0.00308 | 0.4751291 | 0.00229 | 0.6590362 | 0.000875 | 0.3259832 | 0.000504 | 0.4493421 | 0.000294 | 0.4489587 | 0.0000113 |
| **#170** | RBM27 | 0.296745 | 0.001185012 | 0.3029126 | 0.000452 | 0.2886502 | 0.00123 | 0.3812429 | 0.00533 | 0.1985007 | 0.0000857 | 0.2844799 | 0.000018 | 0.3246838 | 6.47E-08 |
| **#171** | SYAP1 | 0.6022013 | 0.001216839 | 0.8495017 | 3.59E-10 | 0.2759846 | 0.00676 | 0.9277761 | 0.000123 | 0.3719888 | 0.000161 | 0.4573495 | 0.000253 | 0.7306073 | 6.5E-09 |
| **#172** | SCYL2 | 0.538781 | 0.001221736 | 0.6294602 | 0.000765 | 0.410032 | 0.000505 | 0.807395 | 0.00147 | 0.4441696 | 0.00115 | 0.5142225 | 0.00331 | 0.427407 | 0.000137 |
| **#173** | ERC1 | 0.8141843 | 0.001248656 | 0.6446564 | 0.00000295 | 0.7340524 | 0.00000255 | 0.847434 | 0.00307 | 0.9414552 | 0.000000829 | 0.7121599 | 0.00442 | 1.005348 | 3.46E-10 |
| **#174** | DMXL1 | 0.6303346 | 0.001256882 | 0.670519 | 0.000000686 | 0.7285756 | 0.00468 | 0.638412 | 0.00286 | 0.56504 | 0.000000215 | 0.5995222 | 0.00000252 | 0.5799389 | 0.000000083 |
| **#175** | ZNF92 | 0.9044718 | 0.001264163 | 0.8929069 | 0.00000218 | 1.197353 | 8.54E-09 | 0.5877649 | 0.00751 | 0.979927 | 5.93E-08 | 0.901249 | 0.0000308 | 0.8676299 | 0.0000388 |
| **#176** | BLOC1S6 | 0.4144959 | 0.001269283 | 0.3491959 | 0.000115 | 0.3848774 | 0.000074 | 0.3545182 | 0.00709 | 0.5731664 | 0.00029 | 0.4341745 | 0.0000453 | 0.3910429 | 0.000000248 |
| **#177** | COQ10B | 0.493957 | 0.001280741 | 0.5125052 | 0.000134 | 0.4488846 | 0.00336 | 0.5802542 | 0.00407 | 0.5101657 | 0.00000774 | 0.4847681 | 0.0000668 | 0.4271644 | 0.0000489 |
| **#178** | ZNF708 | 0.7949031 | 0.001282613 | 0.8642687 | 9.48E-10 | 0.8487818 | 0.000618 | 0.8070975 | 0.00701 | 0.6573252 | 0.0000588 | 0.7064584 | 0.00000833 | 0.8854867 | 5.4E-10 |
| **#179** | PTBP3 | 0.4505708 | 0.00131686 | 0.4321842 | 0.000101 | 0.5577357 | 0.000759 | 0.5328132 | 0.00624 | 0.3373288 | 0.0000256 | 0.4384216 | 0.00075 | 0.4049413 | 0.0000254 |
| **#180** | CCDC6 | 0.8321356 | 0.001320115 | 1.004168 | 0.0000309 | 0.5314626 | 0.0049 | 0.773818 | 0.00289 | 0.9445729 | 0.000035 | 0.8494786 | 0.0000556 | 0.8893135 | 0.00000527 |
| **#181** | MYO5A | 0.7444821 | 0.00133154 | 0.6509709 | 0.0000105 | 0.9329912 | 6.38E-10 | 0.8338923 | 0.00559 | 0.695945 | 0.0000189 | 0.7926722 | 0.00233 | 0.5604209 | 0.0000426 |
| **#182** | ABI1 | 0.597773 | 0.001341912 | 0.6942582 | 4.96E-10 | 0.6739351 | 0.000123 | 0.6080038 | 0.0077 | 0.4017573 | 0.00000549 | 0.5663585 | 0.00022 | 0.6423248 | 0.000000046 |
| **#183** | CENPC | 0.7080303 | 0.001378303 | 0.6426894 | 0.00000404 | 0.906258 | 0.0000213 | 0.5636224 | 0.00214 | 0.793134 | 0.000000365 | 0.5399537 | 0.0061 | 0.8025242 | 0.00000116 |
| **#184** | ATP5MPL | -0.3401479 | 0.001379702 | -0.3561848 | 0.000203 | -0.2830014 | 0.0000546 | -0.2905005 | 0.008 | -0.3816645 | 0.00000187 | -0.4033245 | 0.0000192 | -0.3262119 | 0.00000182 |
| **#185** | USPL1 | 0.7650139 | 0.001381493 | 0.6123863 | 0.000131 | 0.8991923 | 0.0000263 | 0.9374409 | 0.00153 | 0.5479457 | 0.0000181 | 0.6522762 | 0.00658 | 0.940842 | 0.000000239 |
| **#186** | USP12 | 0.7057995 | 0.001393234 | 0.666363 | 0.00316 | 0.6565837 | 0.00146 | 0.7029891 | 0.00272 | 0.5920529 | 0.000928 | 0.8829248 | 0.000033 | 0.7338836 | 0.0000567 |
| **#187** | AP3B1 | 0.5851231 | 0.001394581 | 0.4810597 | 0.000583 | 0.6149174 | 1.73E-08 | 0.4701775 | 0.00774 | 0.5228922 | 4.04E-08 | 0.7643404 | 0.0000469 | 0.6573517 | 1.67E-11 |
| **#188** | AFF1 | 0.6527885 | 0.001396798 | 0.4192434 | 0.00124 | 0.7748649 | 0.000728 | 0.6926608 | 0.00263 | 0.7409364 | 0.00365 | 0.5597073 | 0.000065 | 0.729318 | 0.0000665 |
| **#189** | TBC1D5 | 0.4498667 | 0.001440609 | 0.5143408 | 5.14E-08 | 0.4068004 | 0.0000215 | 0.5188617 | 0.00847 | 0.3638581 | 0.0000413 | 0.5795321 | 0.0000545 | 0.3158072 | 0.0000603 |
| **#190** | GUF1 | 0.6743101 | 0.001456356 | 0.6164933 | 0.0000046 | 0.6378892 | 0.00112 | 0.7144555 | 0.00691 | 0.6303213 | 0.00000221 | 0.6835673 | 0.000697 | 0.7631338 | 0.00000778 |
| **#191** | CDKN1B | 0.5416351 | 0.001479666 | 0.503707 | 0.000162 | 0.5231667 | 0.00327 | 0.8342412 | 0.00122 | 0.5243152 | 0.000000497 | 0.4239509 | 0.00405 | 0.4404298 | 0.000176 |
| **#192** | RFX7 | 0.751555 | 0.001481234 | 0.9893826 | 0.000000595 | 0.6605496 | 0.000335 | 0.749544 | 0.00531 | 0.4771197 | 0.00306 | 0.8079532 | 0.000183 | 0.8247808 | 0.000000238 |
| **#193** | CDC16 | 0.598168 | 0.00149598 | 0.5544492 | 0.0000786 | 0.5147514 | 0.00134 | 0.7324789 | 0.00354 | 0.5478743 | 0.00000495 | 0.5795992 | 0.00401 | 0.6598552 | 0.00000262 |
| **#194** | ZFP91 | 0.4467952 | 0.001496566 | 0.4421484 | 0.000751 | 0.4170326 | 0.00122 | 0.425961 | 0.00612 | 0.4794426 | 0.0000828 | 0.435596 | 0.000803 | 0.4805906 | 0.00000346 |
| **#195** | TUBGCP2 | 0.5349727 | 0.001530639 | 0.5579331 | 0.000000293 | 0.3917373 | 0.00136 | 0.5566871 | 0.00782 | 0.5055154 | 4.92E-08 | 0.57519 | 0.000000275 | 0.6227731 | 6.24E-11 |
| **#196** | CLK1 | 0.617602 | 0.001534414 | 0.6268408 | 0.000000189 | 0.6806057 | 0.000000603 | 0.4611403 | 0.009 | 0.6696302 | 0.00000971 | 0.6446524 | 0.000189 | 0.6227424 | 0.00000495 |
| **#197** | PHC3 | 0.643436 | 0.001553805 | 0.5997152 | 9.15E-08 | 0.7664435 | 0.00214 | 0.6208752 | 0.00701 | 0.5647336 | 0.000000281 | 0.6670763 | 0.000173 | 0.6417724 | 4.08E-08 |
| **#198** | ANAPC4 | 0.5269725 | 0.001554515 | 0.4256387 | 0.0017 | 0.5159008 | 0.0000985 | 0.597149 | 0.00724 | 0.4880559 | 0.0000214 | 0.7025371 | 0.00027 | 0.4325535 | 0.00000429 |
| **#199** | KDM1A | 0.6149636 | 0.001624675 | 0.6833767 | 0.0000445 | 0.7996875 | 0.000229 | 0.4632741 | 0.00851 | 0.439612 | 0.000946 | 0.6547419 | 0.00000545 | 0.6490895 | 0.0000149 |
| **#200** | AHCTF1 | 0.9038561 | 0.001631169 | 0.9020663 | 0.000000309 | 1.033938 | 5.16E-09 | 0.8461745 | 0.00951 | 0.9009059 | 0.000000121 | 0.9516473 | 0.000276 | 0.7884044 | 0.000000417 |
| **#201** | UBR1 | 0.6719792 | 0.001633415 | 0.4361477 | 0.000285 | 0.4106639 | 0.00461 | 1.019489 | 0.00333 | 0.6535288 | 4.01E-08 | 0.7274983 | 0.00158 | 0.7845482 | 1.65E-09 |
| **#202** | LYSMD3 | 0.7490576 | 0.001657814 | 0.6963125 | 0.000182 | 0.8213933 | 0.000172 | 1.078264 | 0.00127 | 0.3376006 | 0.0083 | 0.8015332 | 0.000000871 | 0.7592421 | 0.0000161 |
| **#203** | CANX | 0.3749688 | 0.001660953 | 0.3966476 | 0.00000139 | 0.5278809 | 0.00139 | 0.4814341 | 0.000166 | 0.3279312 | 0.0000023 | 0.3477999 | 0.00262 | 0.1681192 | 0.00578 |
| **#204** | NDUFA3 | -0.5468008 | 0.001675318 | -0.5974826 | 0.00088 | -0.529365 | 0.000412 | -0.6583778 | 0.00833 | -0.5322269 | 0.0000577 | -0.4831564 | 0.000181 | -0.4801962 | 0.000195 |
| **#205** | HBS1L | 0.6870697 | 0.001708628 | 0.7182914 | 0.0000177 | 0.7529419 | 0.000656 | 0.5655532 | 0.00948 | 0.5349912 | 0.0000377 | 0.7857364 | 0.0000401 | 0.7649042 | 0.0000195 |
| **#206** | TRRAP | 0.6336955 | 0.001713878 | 0.3113981 | 0.00219 | 0.7423603 | 0.0000981 | 0.8029106 | 0.00689 | 0.6798343 | 0.00000129 | 0.6958446 | 0.00076 | 0.5698251 | 0.00035 |
| **#207** | MRPL43 | -0.4086556 | 0.00172311 | -0.3277344 | 0.0008 | -0.3625227 | 0.0000198 | -0.6053566 | 0.00714 | -0.4640918 | 0.00000131 | -0.3524561 | 0.00238 | -0.3397717 | 3.32E-10 |
| **#208** | VPS28 | -0.3985557 | 0.00173047 | -0.3336434 | 0.00122 | -0.4896404 | 0.0000212 | -0.3812068 | 0.00911 | -0.4026394 | 2.65E-11 | -0.4008094 | 0.000033 | -0.3833946 | 0.0000138 |
| **#209** | CUL4B | 0.6955134 | 0.001739563 | 0.544069 | 0.00081 | 0.7871042 | 0.00928 | 0.8872665 | 0.000312 | 0.573134 | 0.000018 | 0.6750996 | 0.0000213 | 0.7064072 | 8.8E-09 |
| **#210** | DCUN1D1 | 0.4114469 | 0.001745321 | 0.4118682 | 0.00056 | 0.3814973 | 0.0000584 | 0.4827253 | 0.00747 | 0.3773043 | 0.0000121 | 0.369475 | 0.00237 | 0.4458115 | 0.000000858 |
| **#211** | KMT2C | 0.5471663 | 0.001751884 | 0.4995519 | 0.00000488 | 0.5371095 | 0.00000423 | 0.5354599 | 0.00117 | 0.5863723 | 0.000013 | 0.5810509 | 0.00932 | 0.5434535 | 0.00000132 |

| | | | | | | | | | | | | | | | |
|---|---|---|---|---|---|---|---|---|---|---|---|---|---|---|---|
| #212 | EIF3F | -0.2902623 | 0.00175359 | -0.2453492 | 0.000109 | -0.258085 | 0.000446 | -0.2811539 | 0.00824 | -0.274285 | 0.000698 | -0.4484252 | 0.000809 | -0.2342756 | 0.000215 |
| #213 | PNN | 0.3847541 | 0.001759361 | 0.2873787 | 0.00288 | 0.475703 | 0.00189 | 0.3945531 | 0.00477 | 0.2991054 | 0.000367 | 0.421586 | 0.000608 | 0.4301987 | 0.0000406 |
| #214 | RPL41 | -0.3128841 | 0.001773222 | -0.3005845 | 0.0000004 | -0.2464558 | 0.000135 | -0.2127988 | 0.00966 | -0.2535955 | 0.000841 | -0.5097933 | 0.00000177 | -0.3540769 | 4.32E-08 |
| #215 | ATP5F1B | -0.3011913 | 0.001827836 | -0.3221875 | 0.000428 | -0.29381 | 0.0000273 | -0.3174448 | 0.00609 | -0.2156413 | 0.0018 | -0.3107768 | 0.00252 | -0.3472876 | 0.00011 |
| #216 | ZMYM6 | 0.756337 | 0.001859989 | 0.7253171 | 2.62E-08 | 0.8673064 | 0.00000021 | 0.7269148 | 0.0108 | 0.719847 | 0.0000052 | 0.8022271 | 0.000235 | 0.6964094 | 0.000126 |
| #217 | TTC37 | 0.5662858 | 0.001861842 | 0.6361385 | 0.0000043 | 0.5806197 | 0.00686 | 0.6133276 | 0.00316 | 0.4397331 | 0.000422 | 0.6147931 | 0.000673 | 0.5131031 | 0.0000189 |
| #218 | RPLP1 | -0.2960377 | 0.001885239 | -0.3193462 | 2.25E-08 | -0.2464221 | 0.0000105 | -0.1803479 | 0.00852 | -0.2150654 | 0.00132 | -0.4120379 | 0.00147 | -0.4030064 | 1.61E-09 |
| #219 | TBC1D1 | 0.5366789 | 0.001911904 | 0.3562487 | 0.000128 | 0.5613789 | 0.000297 | 0.8873234 | 0.00819 | 0.4182841 | 0.00000502 | 0.6530471 | 0.00001 | 0.3437914 | 0.00284 |
| #220 | INO80D | 0.5757267 | 0.001926384 | 0.460081 | 0.00000741 | 0.736986 | 0.000000119 | 0.5089502 | 0.0112 | 0.6522065 | 0.000000129 | 0.5413558 | 0.00035 | 0.5547805 | 0.000000049 |
| #221 | ZFR | 0.3656265 | 0.001945024 | 0.299719 | 0.00399 | 0.2370546 | 0.00233 | 0.677942 | 0.00147 | 0.2782053 | 0.000117 | 0.4465305 | 0.000793 | 0.2543075 | 0.00296 |
| #222 | MTRNR2L12 | 0.7365039 | 0.001956523 | 0.7424663 | 0.0000514 | 0.8129384 | 0.0000487 | 0.666068 | 0.00211 | 0.7441902 | 0.000172 | 0.6510763 | 0.00931 | 0.8022843 | 0.0000514 |
| #223 | SUZ12 | 0.5184506 | 0.001981486 | 0.6030905 | 0.00000113 | 0.5333495 | 0.000474 | 0.5588558 | 0.0114 | 0.407179 | 0.0000346 | 0.468448 | 0.0000122 | 0.5397808 | 0.00000287 |
| #224 | ZBTB11 | 0.7544617 | 0.002001975 | 0.844314 | 0.000000115 | 0.7924951 | 0.000000272 | 0.6031187 | 0.0115 | 0.532255 | 0.00056 | 0.8997048 | 3.63E-08 | 0.8548824 | 1.88E-08 |
| #225 | SUCO | 0.8482943 | 0.002007117 | 0.8379495 | 0.00000307 | 1.091942 | 0.00000367 | 0.5823397 | 0.012 | 0.8589414 | 1.44E-08 | 0.7986438 | 0.0000301 | 0.9199493 | 6.28E-11 |
| #226 | LIMD2 | -0.4206107 | 0.00208908 | -0.3462046 | 0.000419 | -0.5821332 | 0.00168 | -0.50027 | 0.00728 | -0.5235029 | 0.00000311 | -0.3177815 | 0.000247 | -0.2537724 | 0.0029 |
| #227 | PRRC2C | 0.4384989 | 0.002102636 | 0.4502196 | 0.00000129 | 0.3743326 | 0.00000937 | 0.4573476 | 0.0000395 | 0.4086686 | 1.89E-09 | 0.4805763 | 0.0126 | 0.4598484 | 0.000000227 |
| #228 | PCM1 | 0.5751576 | 0.00211775 | 0.6017036 | 3E-10 | 0.6370784 | 0.00115 | 0.6198511 | 0.000453 | 0.5392523 | 0.0000431 | 0.4734937 | 0.0111 | 0.5795663 | 0.00000146 |
| #229 | SECISBP2 | 0.4814653 | 0.002143471 | 0.3852062 | 0.0000496 | 0.5830355 | 0.00128 | 0.6274864 | 0.00236 | 0.3960677 | 0.000000113 | 0.3958666 | 0.00916 | 0.5011292 | 0.00000398 |
| #230 | RB1CC1 | 0.4245523 | 0.002150324 | 0.5583491 | 0.00000561 | 0.4484303 | 0.0000988 | 0.3662802 | 0.0102 | 0.2429804 | 0.000289 | 0.5184225 | 0.00214 | 0.4128515 | 0.000166 |
| #231 | GOLGB1 | 0.5663942 | 0.002156113 | 0.6160196 | 0.000011 | 0.6152002 | 0.00000164 | 0.5014576 | 0.00116 | 0.4926746 | 0.00000411 | 0.6105267 | 0.0117 | 0.5624867 | 0.0000228 |
| #232 | SPTLC2 | 0.5169686 | 0.002164031 | 0.4133144 | 0.00136 | 0.4149696 | 0.000019 | 0.7892849 | 0.000675 | 0.5288564 | 0.00778 | 0.5253407 | 0.00216 | 0.4300454 | 0.000987 |
| #233 | MDM2 | 0.6058257 | 0.002167161 | 0.5180817 | 0.0000323 | 0.8275884 | 0.000389 | 0.7152024 | 0.0117 | 0.5387232 | 0.000105 | 0.5594599 | 0.000762 | 0.4758987 | 0.0000381 |
| #234 | UXT | -0.2724133 | 0.002180401 | -0.3051868 | 0.00000125 | -0.2403531 | 0.0000164 | -0.2343703 | 0.0129 | -0.2569467 | 0.000000414 | -0.4117407 | 0.0000991 | -0.1858821 | 0.0000762 |
| #235 | NDUFB10 | -0.3044488 | 0.002187204 | -0.267918 | 0.000515 | -0.2522986 | 0.000386 | -0.2747453 | 0.0104 | -0.2996193 | 0.0000148 | -0.3845493 | 0.00175 | -0.3475625 | 0.0000407 |
| #236 | RNASEK | -0.3730507 | 0.002188466 | -0.4316144 | 0.00000778 | -0.2763826 | 0.00000432 | -0.3116211 | 0.0118 | -0.4026739 | 9.18E-08 | -0.3670209 | 0.0013 | -0.4489912 | 7.54E-08 |
| #237 | FCHSD2 | 0.9297305 | 0.002193047 | 0.8563297 | 3.25E-08 | 1.01645 | 0.011 | 1.040672 | 0.00203 | 0.8435274 | 0.000000458 | 0.8768518 | 0.000119 | 0.9445518 | 3.97E-09 |
| #238 | ELP4 | 0.6024038 | 0.002215533 | 0.4726858 | 0.00171 | 0.4339674 | 0.0104 | 0.8482914 | 0.000518 | 0.5327224 | 0.000592 | 0.7210332 | 0.0000251 | 0.6057228 | 0.000000136 |
| #239 | JOSD1 | 0.6148457 | 0.002251842 | 0.7569925 | 0.0000571 | 0.4976587 | 0.00333 | 0.7829672 | 0.00934 | 0.529501 | 0.0000578 | 0.5835562 | 0.000443 | 0.5383987 | 0.000278 |
| #240 | COX6B1 | -0.3290586 | 0.002257623 | -0.3080912 | 0.000548 | -0.2997514 | 0.0000142 | -0.265111 | 0.0108 | -0.3152419 | 0.00000886 | -0.4543718 | 0.00222 | -0.331784 | 4.37E-08 |
| #241 | UBR5 | 0.5476663 | 0.002326389 | 0.4895126 | 0.0000827 | 0.4217627 | 0.00283 | 0.697622 | 0.00316 | 0.6211669 | 0.00000371 | 0.5473207 | 0.00774 | 0.5086129 | 0.000136 |
| #242 | SPCS3 | 0.6157104 | 0.002365784 | 0.6203992 | 0.000000524 | 0.8059401 | 0.00424 | 0.5329003 | 0.00995 | 0.5670671 | 0.000001 | 0.6185407 | 0.000000412 | 0.5494152 | 5.69E-10 |
| #243 | TMEM30A | 0.693499 | 0.00238473 | 0.6417765 | 0.0000212 | 0.825302 | 0.000337 | 0.563253 | 0.0139 | 0.6388206 | 0.00000111 | 0.8844949 | 0.0000114 | 0.6073469 | 4.38E-08 |
| #244 | POLR2E | -0.3448049 | 0.002415517 | -0.3847865 | 0.00101 | -0.2556045 | 0.00147 | -0.4708417 | 0.01 | -0.2801819 | 0.0000197 | -0.4194742 | 0.00135 | -0.2579407 | 0.000596 |
| #245 | CSNK2B | -0.3396843 | 0.002438005 | -0.2911698 | 0.000184 | -0.384742 | 0.00000179 | -0.2857586 | 0.0125 | -0.38514 | 0.00000214 | -0.3229304 | 0.00197 | -0.368365 | 0.00000183 |
| #246 | ABHD14B | -0.4446329 | 0.002491085 | -0.4281375 | 0.0000494 | -0.4592077 | 0.00000445 | -0.3637437 | 0.0119 | -0.3497527 | 0.003 | -0.5404378 | 0.00000794 | -0.5265184 | 0.00000434 |
| #247 | PCNX1 | 0.5776504 | 0.002494195 | 0.6347761 | 0.00000314 | 0.5978201 | 0.00393 | 0.6037102 | 0.00922 | 0.4576438 | 0.00157 | 0.6910928 | 0.000196 | 0.4808592 | 0.0000453 |
| #248 | ARGLU1 | 0.4298226 | 0.002503543 | 0.4813916 | 2.14E-11 | 0.403504 | 0.000111 | 0.286288 | 0.0136 | 0.3725237 | 0.000124 | 0.4865003 | 0.00119 | 0.5487282 | 0.000002 |
| #249 | BDP1 | 0.5146047 | 0.002508418 | 0.5156704 | 0.00000012 | 0.5727825 | 0.000135 | 0.4678192 | 0.0114 | 0.4884661 | 0.0000195 | 0.5325696 | 0.00349 | 0.5103203 | 0.00000171 |
| #250 | NDUFA8 | -0.5778681 | 0.00252245 | -0.7133192 | 0.00603 | -0.4937612 | 0.00064 | -0.5769624 | 0.00684 | -0.5365164 | 0.0000874 | -0.5655484 | 0.00146 | -0.5811011 | 0.0000811 |
| #251 | SLC36A4 | 0.6225371 | 0.002525009 | 0.8236125 | 0.00000238 | 0.5379689 | 0.00187 | 0.8437471 | 0.00391 | 0.2436737 | 0.00855 | 0.6363452 | 0.000805 | 0.6498752 | 0.00000717 |
| #252 | NF1 | 0.5280165 | 0.002527414 | 0.4401243 | 0.0117 | 0.3895574 | 0.00153 | 0.7354274 | 0.00149 | 0.5211009 | 0.0000286 | 0.6186894 | 0.000113 | 0.4631999 | 0.000271 |
| #253 | SDHC | -0.4074612 | 0.002534325 | -0.4585943 | 0.00555 | -0.3555566 | 0.0000215 | -0.5751888 | 0.00324 | -0.2360387 | 0.00328 | -0.4441637 | 0.00296 | -0.3752251 | 0.000148 |
| #254 | XRN2 | 0.3224205 | 0.002578153 | 0.2939064 | 0.000698 | 0.2488015 | 0.00983 | 0.3863069 | 0.0043 | 0.1675581 | 0.000639 | 0.3802942 | 0.000000302 | 0.4576557 | 1.25E-08 |
| #255 | EPS15 | 0.4133977 | 0.00258739 | 0.3922377 | 0.00000136 | 0.483075 | 0.000249 | 0.4307525 | 0.0147 | 0.4289111 | 0.000000036 | 0.4085181 | 0.000017 | 0.3368916 | 0.000516 |
| #256 | UBR2 | 0.5521821 | 0.002589871 | 0.6087241 | 0.00000019 | 0.5849664 | 0.00638 | 0.58875 | 0.00173 | 0.459972 | 0.000000431 | 0.5065107 | 0.00742 | 0.5641692 | 0.00000652 |
| #257 | PHB2 | -0.3535869 | 0.00259065 | -0.3026681 | 0.0000386 | -0.2968456 | 0.0000224 | -0.3185631 | 0.0148 | -0.3093666 | 0.000671 | -0.5405694 | 0.0000426 | -0.3535085 | 0.000000219 |
| #258 | ETNK1 | 0.4212614 | 0.00260994 | 0.4098689 | 0.000356 | 0.5512385 | 0.00212 | 0.4863967 | 0.0101 | 0.3309892 | 0.0029 | 0.3961197 | 0.00000281 | 0.3529556 | 0.00022 |
| #259 | ZDHHC20 | 0.6307456 | 0.002646818 | 0.6528084 | 0.0000397 | 0.656977 | 0.0000035 | 0.5444208 | 0.0158 | 0.6956422 | 0.0000205 | 0.624505 | 0.0000109 | 0.61012 | 0.000000341 |
| #260 | BRK1 | -0.4266171 | 0.00264794 | -0.4889775 | 0.000068 | -0.3468061 | 0.000000332 | -0.3774047 | 0.0135 | -0.3834064 | 0.0000195 | -0.4903971 | 0.0023 | -0.472711 | 0.000000339 |
| #261 | MSH3 | 0.6023637 | 0.002680347 | 0.7044989 | 0.00000457 | 0.5031381 | 0.000508 | 0.725858 | 0.0103 | 0.4599838 | 0.00251 | 0.598043 | 0.00278 | 0.6226603 | 0.00000209 |
| #262 | ELOB | -0.3157403 | 0.002703949 | -0.3188911 | 0.00162 | -0.3192933 | 0.000506 | -0.2984233 | 0.0139 | -0.3105662 | 0.0000613 | -0.3470594 | 0.000156 | -0.3002082 | 0.000000155 |
| #263 | MATR3.1 | 0.9681903 | 0.002719695 | 1.022441 | 1.64E-12 | 1.092135 | 0.000263 | 0.9242606 | 0.0159 | 1.01929 | 0.00000047 | 0.6765446 | 0.000124 | 1.07447 | 5.88E-09 |
| #264 | EPM2AIP1 | 0.5267863 | 0.002739179 | 0.5131941 | 0.000101 | 0.592742 | 0.000441 | 0.4639782 | 0.0158 | 0.3654974 | 0.0000228 | 0.5841589 | 0.0000357 | 0.6411475 | 2.85E-08 |
| #265 | NUDT2 | -0.9178358 | 0.002761315 | -0.8599916 | 0.000774 | -1.0687 | 0.000386 | -0.7750298 | 0.0154 | -1.080475 | 0.00000709 | -0.7808774 | 0.0000142 | -0.941941 | 0.00000471 |
| #266 | ATP11B | 0.7669165 | 0.002790954 | 0.7738453 | 0.000000746 | 0.8390677 | 0.000287 | 0.7300721 | 0.0164 | 0.6337741 | 2.66E-09 | 0.7564831 | 0.0000138 | 0.8682567 | 1.96E-10 |
| #267 | EHMT1 | 0.4194139 | 0.002793379 | 0.2983767 | 0.00155 | 0.3059328 | 0.00123 | 0.6059951 | 0.000113 | 0.3136162 | 0.0000768 | 0.3699806 | 0.0138 | 0.6225821 | 7.88E-08 |
| #268 | KIAA1109 | 0.9076599 | 0.002825061 | 1.048318 | 2.87E-09 | 0.8446856 | 1.05E-10 | 0.8636843 | 0.000278 | 1.032873 | 2.45E-14 | 0.8214295 | 0.0167 | 0.8349695 | 1.58E-08 |
| #269 | MAPK6 | 0.5438289 | 0.002828629 | 0.5503254 | 0.000391 | 0.5655222 | 0.00036 | 0.527985 | 0.0142 | 0.5367322 | 0.000433 | 0.546294 | 0.0015 | 0.5361145 | 0.0000536 |
| #270 | CCNDBP1 | -0.4038652 | 0.002858136 | -0.2652496 | 0.0147 | -0.4609257 | 0.000213 | -0.4779508 | 0.00202 | -0.3229861 | 0.000076 | -0.4273675 | 0.0000319 | -0.4687116 | 0.0000637 |
| #271 | ZBTB21 | 0.9289152 | 0.002861958 | 0.8869139 | 0.0000297 | 1.204115 | 0.000000469 | 0.6184979 | 0.0171 | 0.7991031 | 0.00000538 | 0.9901336 | 0.0000601 | 1.074728 | 0.00002 |
| #272 | RPL39 | -0.3321137 | 0.002875386 | -0.3458562 | 0.00000182 | -0.2939977 | 0.000435 | -0.3092681 | 0.0146 | -0.2247598 | 0.00172 | -0.4660676 | 0.000449 | -0.3527328 | 0.00000378 |
| #273 | SREK1 | 0.5510726 | 0.002880182 | 0.6017376 | 0.000000626 | 0.6202687 | 0.00000475 | 0.3664115 | 0.0172 | 0.5546828 | 0.00000158 | 0.5566152 | 0.0000359 | 0.6067198 | 0.000000448 |
| #274 | BRWD1 | 0.6212605 | 0.002889629 | 0.7892729 | 5.77E-08 | 0.6273584 | 0.000168 | 0.5515306 | 0.00496 | 0.5059341 | 0.000235 | 0.5610979 | 0.012 | 0.6923691 | 2.76E-08 |
| #275 | LATS1 | 0.6054952 | 0.002892103 | 0.6052096 | 0.0000268 | 0.4827914 | 0.00833 | 0.8478675 | 0.00897 | 0.4744815 | 0.0000122 | 0.690961 | 0.0000171 | 0.5316602 | 0.000000217 |
| #276 | API5 | 0.4649966 | 0.002931196 | 0.3903481 | 0.000319 | 0.4684641 | 0.000383 | 0.5613295 | 0.0128 | 0.4138568 | 0.00182 | 0.4677263 | 0.00222 | 0.4882546 | 1.48E-08 |
| #277 | CUL3 | 0.4933783 | 0.002931792 | 0.530613 | 0.000193 | 0.4924639 | 0.000674 | 0.4510393 | 0.0138 | 0.4625846 | 0.000101 | 0.462164 | 0.00206 | 0.561405 | 0.000798 |
| #278 | IL6ST | 0.8734882 | 0.002938742 | 1.05806 | 0.000303 | 1.178458 | 0.00597 | 0.7309804 | 0.00531 | 0.552074 | 0.00424 | 0.6797674 | 0.0018 | 1.04159 | 0.00000303 |
| #279 | UQCR10 | -0.3328282 | 0.002955378 | -0.3929226 | 0.000962 | -0.2704536 | 0.00197 | -0.4247654 | 0.00327 | -0.2826251 | 0.000321 | -0.3497023 | 0.011 | -0.2765002 | 0.000165 |
| #280 | C1GALT1 | 0.5381568 | 0.002984843 | 0.5917932 | 0.0000178 | 0.6803965 | 0.00334 | 0.7536355 | 0.00492 | 0.3893481 | 0.00644 | 0.3533589 | 0.00318 | 0.4604085 | 0.00000134 |
| #281 | PSMD6-AS2 | 0.7842448 | 0.002991145 | 0.7763024 | 0.00371 | 0.8345976 | 0.000116 | 1.006041 | 0.00521 | 0.6793862 | 0.00628 | 0.6704505 | 0.00261 | 0.7386908 | 0.0000206 |
| #282 | STK26 | 0.4388622 | 0.002993656 | 0.4160345 | 0.00117 | 0.4206727 | 0.00239 | 0.7665989 | 0.000502 | 0.2805847 | 0.00609 | 0.3486465 | 0.0063 | 0.4006362 | 0.00152 |

| | | | | | | | | | | | | | | |
|---|---|---|---|---|---|---|---|---|---|---|---|---|---|---|
| #283 | NDUFB11 | -0.3205134 | 0.003020971 | -0.2989212 | 0.000591 | -0.258933 | 0.0000439 | -0.3068722 | 0.0167 | -0.3155283 | 0.000000946 | -0.4094024 | 0.000806 | -0.3334235 | 3.94E-08 |
| #284 | ARPC1B | -0.3778565 | 0.003023692 | -0.3383534 | 0.00179 | -0.3637971 | 0.000514 | -0.3781474 | 0.00667 | -0.3415318 | 0.000000613 | -0.3896738 | 0.00915 | -0.4556353 | 0.0000172 |
| #285 | RPS28 | -0.3277155 | 0.003039126 | -0.2942902 | 0.0000132 | -0.2383967 | 0.0169 | -0.3719676 | 0.000754 | -0.2984563 | 0.000242 | -0.4945558 | 0.0000758 | -0.2686265 | 0.000293 |
| #286 | PNPLA8 | 0.8588802 | 0.003039178 | 1.014642 | 0.00000477 | 0.7604314 | 0.0000127 | 0.582961 | 0.0182 | 0.558406 | 0.00000382 | 1.12747 | 0.0000245 | 1.10937 | 0.000000236 |
| #287 | NOP58 | 0.5357635 | 0.003040571 | 0.615684 | 0.000000317 | 0.5044925 | 0.00861 | 0.3202129 | 0.00932 | 0.413994 | 0.000304 | 0.78492 | 0.000000846 | 0.5752773 | 1.49E-09 |
| #288 | SPAG9 | 0.5845241 | 0.003062798 | 0.5201857 | 0.0000176 | 0.5689096 | 0.00000553 | 0.5134735 | 0.0173 | 0.5718194 | 6.94E-09 | 0.6123232 | 0.00105 | 0.7204333 | 9.49E-08 |
| #289 | PPIP5K2 | 0.5893941 | 0.003096217 | 0.6261659 | 0.00000277 | 0.58872 | 2.66E-09 | 0.4760758 | 0.0162 | 0.3831661 | 0.0022 | 0.7219459 | 0.000187 | 0.7402907 | 4.58E-08 |
| #290 | CEP95 | 0.650338 | 0.003153011 | 0.7636569 | 4.27E-08 | 0.7079441 | 0.000804 | 0.5331327 | 0.0129 | 0.5676803 | 0.000153 | 0.5417552 | 0.00503 | 0.7878588 | 3.17E-08 |
| #291 | PTAR1 | 0.4623182 | 0.003164004 | 0.3462674 | 0.000901 | 0.6035093 | 0.00000125 | 0.4866549 | 0.016 | 0.4508852 | 0.000208 | 0.3842362 | 0.00187 | 0.5023563 | 5.84E-08 |
| #292 | TMEM131 | 0.7461273 | 0.003185858 | 0.8041766 | 0.000000604 | 0.805696 | 0.0000218 | 0.6594082 | 0.015 | 0.6898141 | 0.00327 | 0.8049711 | 0.000778 | 0.712698 | 0.00000858 |
| #293 | NRIP1 | 0.6428682 | 0.003190453 | 0.8056672 | 0.000000514 | 0.6945699 | 0.0000018 | 0.4624286 | 0.0171 | 0.6771448 | 0.000419 | 0.5125589 | 0.00157 | 0.7048398 | 0.0000431 |
| #294 | PUM2 | 0.5269913 | 0.003198133 | 0.6319721 | 0.0000103 | 0.4658759 | 0.0000785 | 0.4512064 | 0.0187 | 0.4435902 | 0.000358 | 0.6078134 | 0.000012 | 0.5614895 | 0.00000173 |
| #295 | ARAP2 | 1.007263 | 0.003198974 | 1.066594 | 0.000000307 | 1.016849 | 0.0184 | 1.281763 | 0.00000028€ | 1.008628 | 6.37E-09 | 0.8714377 | 0.000754 | 0.7983065 | 2.74E-08 |
| #296 | CGGBP1 | 0.3285176 | 0.00320335 | 0.2311367 | 0.00104 | 0.3471319 | 0.000169 | 0.4993923 | 0.00206 | 0.1763133 | 0.0157 | 0.3719976 | 0.000289 | 0.3451339 | 3.08E-08 |
| #297 | MTMR12 | 0.6670562 | 0.003209484 | 0.5725251 | 0.0000281 | 0.7785573 | 0.0000319 | 0.701946 | 0.0191 | 0.5784485 | 0.0000258 | 0.6028531 | 0.0000376 | 0.7680073 | 0.00000308 |
| #298 | HSPA5 | 0.5968149 | 0.003225301 | 0.615944 | 0.00000898 | 0.6378411 | 0.00307 | 0.6443206 | 0.00336 | 0.7485627 | 0.000536 | 0.6123489 | 0.00206 | 0.3218723 | 0.0103 |
| #299 | TRIM44 | 0.5386897 | 0.003248471 | 0.5528172 | 0.00000435 | 0.5270153 | 0.00282 | 0.4644263 | 0.0108 | 0.3819577 | 0.0000164 | 0.5456801 | 0.00582 | 0.7602414 | 0.000000935 |
| #300 | TUT4 | 0.7320918 | 0.003292588 | 0.835641 | 1.54E-12 | 0.9073159 | 0.00984 | 0.6101869 | 0.00342 | 0.4203657 | 0.00293 | 0.7414493 | 0.00357 | 0.8775921 | 5.79E-09 |
| #301 | NIPBL | 0.4417109 | 0.003302098 | 0.3843729 | 0.000213 | 0.4104935 | 0.000313 | 0.4497064 | 0.0146 | 0.3347195 | 0.00112 | 0.5512383 | 0.00357 | 0.5197347 | 0.000000671 |
| #302 | PAK2 | 0.3189636 | 0.003314745 | 0.4000901 | 7E-12 | 0.2922642 | 0.0000615 | 0.2994164 | 0.018 | 0.3123143 | 0.00000218 | 0.2777996 | 0.0018 | 0.3318972 | 1.48E-08 |
| #303 | NDUFS5 | -0.4712715 | 0.003315209 | -0.3974712 | 0.0000787 | -0.5527278 | 1.18E-08 | -0.2882864 | 0.0197 | -0.7192948 | 1.91E-09 | -0.5028442 | 0.000117 | -0.3670043 | 7.55E-08 |
| #304 | SNHG29 | -0.3226369 | 0.003333336 | -0.3618872 | 0.000000371 | -0.2846371 | 0.000349 | -0.2441441 | 0.0194 | -0.3446585 | 0.0000179 | -0.4179103 | 0.0000277 | -0.2825844 | 0.000157 |
| #305 | UGGT1 | 0.4866535 | 0.00338811 | 0.4666976 | 0.00092 | 0.5193325 | 0.00569 | 0.6525017 | 0.0111 | 0.4976868 | 3.17E-08 | 0.4151269 | 0.00171 | 0.3685758 | 0.000907 |
| #306 | EXOC2 | 0.5186087 | 0.003458667 | 0.3943253 | 0.000269 | 0.5676207 | 0.00494 | 0.7944752 | 0.0139 | 0.4799939 | 0.00104 | 0.5319986 | 0.000371 | 0.3432388 | 0.000235 |
| #307 | ANKHD1 | 0.4398483 | 0.003460281 | 0.4868889 | 3.77E-09 | 0.3375102 | 0.000332 | 0.6300739 | 0.0000651 | 0.3065201 | 0.000581 | 0.4056305 | 0.0198 | 0.4724663 | 0.00000145 |
| #308 | ATF7IP | 0.6495965 | 0.003487889 | 0.7735178 | 0.00000001 | 0.732432 | 0.000026 | 0.5372402 | 0.00161 | 0.384883 | 0.0189 | 0.6928454 | 0.000397 | 0.7766607 | 0.00000036 |
| #309 | SUMO2 | -0.2827451 | 0.003492211 | -0.2475566 | 0.0000432 | -0.2386525 | 0.000208 | -0.193974 | 0.0154 | -0.3754624 | 0.000000788 | -0.3520713 | 0.00523 | -0.2887536 | 0.000113 |
| #310 | MFSD10 | -0.4669108 | 0.003508764 | -0.4407856 | 0.000285 | -0.4986626 | 0.0000517 | -0.6176344 | 0.0198 | -0.2617828 | 0.000662 | -0.4448299 | 0.0000058 | -0.5377699 | 0.000219 |
| #311 | SDE2 | 0.6163319 | 0.003513066 | 0.5209255 | 0.00256 | 0.6549476 | 0.00925 | 0.7726589 | 0.00569 | 0.5442215 | 0.000151 | 0.6806764 | 0.0000844 | 0.5245613 | 0.00334 |
| #312 | BTBD1 | 0.6407417 | 0.003518099 | 0.6202792 | 0.0000284 | 0.5413015 | 0.00015 | 0.706661 | 0.0207 | 0.5555761 | 0.000115 | 0.6707693 | 0.0000616 | 0.7498632 | 0.0000102 |
| #313 | CEP350 | 0.5883 | 0.003531182 | 0.5725971 | 0.000000368 | 0.5931006 | 2.62E-08 | 0.6731128 | 0.02 | 0.4590519 | 0.000148 | 0.6597365 | 0.00104 | 0.5722009 | 1.38E-10 |
| #314 | CASP8 | 0.4589961 | 0.00353188 | 0.3773249 | 0.000683 | 0.5346237 | 7.48E-08 | 0.5268956 | 0.00644 | 0.5710896 | 0.000000945 | 0.3975433 | 0.0101 | 0.3464995 | 0.00397 |
| #315 | TMEM123 | 0.4186804 | 0.003571766 | 0.3548555 | 0.00746 | 0.3642952 | 0.00596 | 0.4013167 | 0.00372 | 0.4564154 | 0.00319 | 0.6225784 | 0.000979 | 0.312621 | 0.000125 |
| #316 | YME1L1 | 0.376337 | 0.003617184 | 0.2960317 | 0.00126 | 0.4484098 | 0.00428 | 0.5614176 | 0.00978 | 0.3065844 | 0.00235 | 0.3658477 | 0.00274 | 0.2797307 | 0.0013 |
| #317 | NAA50 | 0.3640014 | 0.003619177 | 0.4451142 | 0.00172 | 0.3308657 | 0.00175 | 0.473745 | 0.014 | 0.2941435 | 0.0000243 | 0.428309 | 0.000318 | 0.2118306 | 0.00395 |
| #318 | CEBPZ | 0.4647578 | 0.003654017 | 0.5249319 | 3.77E-09 | 0.4417222 | 0.000000801 | 0.4132331 | 0.0215 | 0.2752528 | 0.000371 | 0.587337 | 0.0000214 | 0.5460701 | 2.04E-08 |
| #319 | KMT2A | 0.4243387 | 0.003664855 | 0.5010261 | 0.0000583 | 0.3943364 | 0.0107 | 0.3927138 | 0.00681 | 0.3556897 | 0.0000338 | 0.5553887 | 0.00411 | 0.3468777 | 0.000315 |
| #320 | SPTBN1 | 0.5388551 | 0.00366655 | 0.5977684 | 0.00000147 | 0.5911721 | 0.000379 | 0.570911 | 0.00375 | 0.522448 | 0.00128 | 0.2905166 | 0.0166 | 0.6603143 | 4.81E-08 |
| #321 | STARD7 | 0.3765238 | 0.003679673 | 0.3982833 | 0.00182 | 0.2205985 | 0.014 | 0.5047701 | 0.00166 | 0.3568631 | 0.000921 | 0.5077918 | 0.000392 | 0.270836 | 0.00328 |
| #322 | N4BP1 | 0.5888578 | 0.003713422 | 0.8147925 | 0.00000328 | 0.5619006 | 0.000553 | 0.6126518 | 0.0206 | 0.4589256 | 0.000369 | 0.5716438 | 0.000483 | 0.5132327 | 0.000265 |
| #323 | H3F3A | -0.2461708 | 0.003720992 | -0.3272392 | 0.00000895 | -0.2438284 | 0.0000888 | -0.1718509 | 0.00302 | -0.3136143 | 2.59E-08 | -0.2400539 | 0.0189 | -0.1804378 | 0.000331 |
| #324 | RAD23B | 0.4839581 | 0.003741838 | 0.5131412 | 0.0000762 | 0.3468792 | 0.0209 | 0.4731432 | 0.0013 | 0.4745301 | 0.0000176 | 0.5369058 | 0.0000773 | 0.5591492 | 0.0000803 |
| #325 | CSNK1A1 | 0.2494614 | 0.003770988 | 0.2108888 | 0.00169 | 0.2665613 | 0.0000105 | 0.3100013 | 0.0168 | 0.2187819 | 0.0000287 | 0.284418 | 0.00398 | 0.2061172 | 0.000135 |
| #326 | ARIH1 | 0.4337534 | 0.003782272 | 0.417672 | 0.00126 | 0.3701859 | 0.000202 | 0.6889726 | 0.0163 | 0.3622519 | 0.00000523 | 0.29126 | 0.00492 | 0.4721781 | 0.000013 |
| #327 | POLD4 | -0.517687 | 0.003793524 | -0.6399986 | 0.000000487 | -0.4252743 | 0.0000119 | -0.5764107 | 0.0221 | -0.5677202 | 0.000614 | -0.5276667 | 0.0000432 | -0.3690517 | 0.000000293 |
| #328 | MOB1A | 0.3579136 | 0.003806331 | 0.4223863 | 0.000213 | 0.394867 | 0.00255 | 0.4019683 | 0.0107 | 0.3310398 | 0.00485 | 0.3166545 | 0.00442 | 0.2805657 | 0.0000991 |
| #329 | DIS3 | 0.6119197 | 0.003810915 | 0.6369005 | 0.00000177 | 0.5735808 | 0.000183 | 0.5718263 | 0.0227 | 0.5150511 | 0.00000953 | 0.7495469 | 0.00000119 | 0.6246124 | 0.00000307 |
| #330 | SP100 | 0.4902712 | 0.003813754 | 0.4048605 | 1.8E-09 | 0.3331526 | 0.0111 | 0.8490712 | 0.000275 | 0.3559628 | 0.0108 | 0.5454563 | 0.0000846 | 0.4531238 | 0.000652 |
| #331 | VPS13A | 0.6877185 | 0.003843459 | 0.54714 | 0.00000142 | 0.8126778 | 0.00618 | 0.7556547 | 0.00176 | 0.7933433 | 0.00000202 | 0.4933757 | 0.0151 | 0.7241193 | 0.00000151 |
| #332 | RNF103 | 0.6590293 | 0.003864257 | 0.6751125 | 0.00115 | 0.8849479 | 0.00000572 | 0.8848563 | 0.00781 | 0.3163009 | 0.00824 | 0.4797326 | 0.00596 | 0.7132259 | 0.00000909 |
| #333 | KCTD20 | 0.4307428 | 0.003874697 | 0.4435788 | 0.0000466 | 0.3829909 | 0.000842 | 0.3890456 | 0.022 | 0.2951096 | 0.000192 | 0.5628113 | 0.0000973 | 0.5109207 | 0.0000441 |
| #334 | MIER1 | 0.3638677 | 0.003950072 | 0.361095 | 0.0000116 | 0.3930744 | 0.00000147 | 0.3610773 | 0.0237 | 0.2330992 | 0.00000055 | 0.3858242 | 0.0000159 | 0.4490361 | 0.000000156 |
| #335 | CHD8 | 0.4786333 | 0.003973211 | 0.4316771 | 0.000436 | 0.53341 | 0.00000156 | 0.4600305 | 0.0177 | 0.445084 | 0.000000341 | 0.4181186 | 0.00575 | 0.5834798 | 9.09E-09 |
| #336 | RMND5A | 0.5404996 | 0.003976584 | 0.3382717 | 0.0103 | 0.597972 | 0.000423 | 0.956674 | 0.0017 | 0.4348676 | 0.00118 | 0.3799695 | 0.0101 | 0.5352425 | 0.000211 |
| #337 | DNM1L | 0.4101283 | 0.003980911 | 0.2984766 | 0.00187 | 0.4186606 | 0.0000821 | 0.3498786 | 0.0117 | 0.5070805 | 2.57E-08 | 0.389594 | 0.0102 | 0.4970795 | 0.00000636 |
| #338 | TAOK1 | 0.3904507 | 0.004029504 | 0.4522537 | 0.00015 | 0.4243787 | 0.000107 | 0.4783567 | 0.00693 | 0.2855679 | 0.00368 | 0.3462824 | 0.0133 | 0.3558648 | 0.0000336 |
| #339 | ZNHIT1 | -0.2866528 | 0.004049492 | -0.3547816 | 0.000184 | -0.2052838 | 0.00539 | -0.2620188 | 0.0176 | -0.2885478 | 0.000016 | -0.3441684 | 0.000602 | -0.2651163 | 0.000502 |
| #340 | BAX | -0.3548247 | 0.00405606 | -0.328447 | 0.00000635 | -0.235969 | 0.0000278 | -0.481758 | 0.0213 | -0.3685013 | 0.0000119 | -0.3614439 | 0.00301 | -0.3528287 | 0.00001 |
| #341 | ZNF451 | 0.5257056 | 0.004058962 | 0.4325751 | 0.000000913 | 0.6480052 | 0.000109 | 0.4082567 | 0.0241 | 0.4390342 | 0.0000237 | 0.6205743 | 0.000144 | 0.6057879 | 6.29E-08 |
| #342 | CD164 | 0.4919791 | 0.004061913 | 0.5833798 | 0.00000024 | 0.5113922 | 0.0193 | 0.5831591 | 0.00498 | 0.3297322 | 0.0000816 | 0.5489201 | 0.000000076 | 0.3952913 | 0.00000133 |
| #343 | DGUOK | -0.3223095 | 0.004063892 | -0.2172157 | 0.003 | -0.2760277 | 0.00168 | -0.4722272 | 0.0185 | -0.3203397 | 0.0000402 | -0.4048833 | 0.00113 | -0.2431636 | 0.00000876 |
| #344 | ATRX | 0.3980213 | 0.004098839 | 0.4701968 | 0.000000117 | 0.3592934 | 0.000396 | 0.3563105 | 0.0202 | 0.2812578 | 0.000939 | 0.5152055 | 0.0031 | 0.405864 | 0.000000394 |
| #345 | USP48 | 0.5247634 | 0.004151796 | 0.582437 | 1.71E-08 | 0.5476838 | 0.0000183 | 0.3294858 | 0.0247 | 0.5337101 | 0.000122 | 0.6231635 | 0.0000902 | 0.5321001 | 0.000000142 |
| #346 | GABPA | 0.6233435 | 0.004167025 | 0.6743613 | 0.000187 | 0.6215162 | 0.0000456 | 0.6285094 | 0.0247 | 0.500036 | 0.0000102 | 0.6281538 | 0.0000495 | 0.6874844 | 1.23E-09 |
| #347 | UBR4 | 0.6540307 | 0.004187271 | 0.6622481 | 0.0000025 | 0.7479043 | 6.83E-08 | 0.5138059 | 0.0238 | 0.6138852 | 2.58E-09 | 0.7804229 | 0.00128 | 0.6059176 | 0.0000228 |
| #348 | CAT | -0.4879665 | 0.004203969 | -0.4855618 | 0.0000461 | -0.3929186 | 0.00197 | -0.6168859 | 0.00631 | -0.5609073 | 0.000341 | -0.4759507 | 0.0163 | -0.3955747 | 0.000238 |
| #349 | PSAP | 0.3737094 | 0.004303517 | 0.4541826 | 0.00000261 | 0.3702855 | 0.000000151 | 0.629801 | 0.016 | 0.2448702 | 0.00276 | 0.3108368 | 0.00593 | 0.2322803 | 0.00114 |
| #350 | GPATCH2L | 0.4413205 | 0.004309389 | 0.2852545 | 0.0128 | 0.5217043 | 0.00000265 | 0.5629677 | 0.0031 | 0.3874113 | 0.0000289 | 0.4308068 | 0.00991 | 0.4597785 | 0.00000964 |
| #351 | TFRC | 0.6423087 | 0.004322972 | 0.4866632 | 0.0135 | 0.7829615 | 0.00223 | 0.8013022 | 0.00376 | 0.6974705 | 0.0000882 | 0.6292254 | 0.000989 | 0.4562294 | 0.00534 |
| #352 | PICALM | 0.4994384 | 0.004348771 | 0.7016906 | 0.0000134 | 0.459281 | 0.00483 | 0.6725325 | 0.0141 | 0.2615458 | 0.00695 | 0.4739231 | 0.000152 | 0.4276575 | 0.0000306 |
| #353 | UBE2H | 0.4836133 | 0.004363561 | 0.5536832 | 0.0000243 | 0.5419447 | 0.00000792 | 0.6617659 | 0.00828 | 0.2538923 | 0.00166 | 0.3468942 | 0.0162 | 0.5434997 | 0.00000221 |

| | | | | | | | | | | | | | | | |
|---|---|---|---|---|---|---|---|---|---|---|---|---|---|---|---|
| **#354** | CHMP1A | -0.5355061 | 0.004383723 | -0.4066088 | 0.0116 | -0.5735847 | 0.0000565 | -1.097397 | 0.000953 | -0.2118795 | 0.0133 | -0.5376387 | 0.000166 | -0.385928 | 0.000237 |
| **#355** | CBX5 | 0.4469676 | 0.004424708 | 0.6080633 | 0.00000114 | 0.3868374 | 0.00336 | 0.4074999 | 0.00729 | 0.3472768 | 0.00304 | 0.360643 | 0.0129 | 0.5714853 | 0.00000223 |
| **#356** | TRIP12 | 0.8663257 | 0.004437479 | 1.08303 | 9.88E-11 | 0.7314936 | 1.63E-08 | 0.4276898 | 0.0266 | 0.7158107 | 3.26E-08 | 1.174159 | 0.0000649 | 1.065771 | 1.12E-12 |
| **#357** | ELK4 | 0.6747796 | 0.004447342 | 0.6068779 | 0.000022 | 0.7453707 | 0.000000313 | 0.6462384 | 0.0267 | 0.5955955 | 0.00000109 | 0.7230378 | 5.07E-08 | 0.731557 | 0.000000221 |
| **#358** | MAPK8 | 0.4898276 | 0.004525317 | 0.3589999 | 0.0161 | 0.6362222 | 0.0000851 | 0.548527 | 0.0104 | 0.3558059 | 0.000212 | 0.3811831 | 0.000358 | 0.6582276 | 0.00000601 |
| **#359** | USP14 | 0.3614667 | 0.004546127 | 0.3264784 | 0.00402 | 0.3013582 | 0.000417 | 0.4468056 | 0.0193 | 0.4091281 | 4.39E-08 | 0.4240132 | 0.000225 | 0.2610169 | 0.0033 |
| **#360** | ITPR2 | 0.7427645 | 0.004586505 | 0.7922755 | 1.44E-08 | 0.7883846 | 0.00373 | 0.430566 | 0.0202 | 0.7038727 | 6.48E-08 | 0.862857 | 0.00364 | 0.8786311 | 2.21E-08 |
| **#361** | MIA2 | 0.5015774 | 0.004645821 | 0.5061691 | 0.000767 | 0.4740945 | 0.0000663 | 0.4770225 | 0.027 | 0.5483569 | 0.000000683 | 0.5353798 | 0.0000464 | 0.4684414 | 0.000000194 |
| **#362** | LSM 7.00 | -0.2423403 | 0.004663234 | -0.3066456 | 0.0000758 | -0.2755029 | 0.0000159 | -0.3077415 | 0.0164 | -0.161762 | 0.0000527 | -0.2585305 | 0.0106 | -0.1438593 | 0.000822 |
| **#363** | ITPRID2 | 0.7481946 | 0.004668232 | 0.9086265 | 0.0000328 | 0.7573597 | 0.000000364 | 0.8986069 | 0.00434 | 0.7165703 | 0.00000722 | 0.3729333 | 0.0236 | 0.8350708 | 0.000000218 |
| **#364** | WHAMM | 0.4859832 | 0.004675935 | 0.4171825 | 0.00346 | 0.528822 | 0.000894 | 0.5961366 | 0.0156 | 0.2842701 | 0.00658 | 0.5584639 | 0.000619 | 0.5310238 | 0.000943 |
| **#365** | SMARCA2 | 0.4419134 | 0.004712687 | 0.3907296 | 0.0000505 | 0.3971663 | 0.00000325 | 0.4577173 | 0.0279 | 0.4536665 | 0.00000185 | 0.4786188 | 0.000285 | 0.4735817 | 0.00000197 |
| **#366** | ILF3 | 0.4791249 | 0.004726387 | 0.5171817 | 0.00000715 | 0.3587942 | 0.00000248 | 0.3768523 | 0.0282 | 0.4878363 | 1.61E-08 | 0.5574093 | 0.000166 | 0.5766759 | 0.000000251 |
| **#367** | RPL28 | -0.1952679 | 0.004771372 | -0.2259184 | 8.31E-08 | -0.1606238 | 0.00668 | -0.1214296 | 0.0211 | -0.1508364 | 0.000322 | -0.3216752 | 0.000366 | -0.1911241 | 0.000117 |
| **#368** | COX7A2 | -0.275286 | 0.004796253 | -0.2482909 | 0.0000243 | -0.2262761 | 0.0147 | -0.2220953 | 0.0123 | -0.3330293 | 0.000615 | -0.4175684 | 0.000795 | -0.2044562 | 0.000263 |
| **#369** | LGALS1 | -1.081276 | 0.00481304 | -1.287019 | 0.00000554 | -0.8588641 | 0.00393 | -1.933134 | 0.0245 | -0.5851183 | 0.000157 | -0.7129563 | 0.000264 | -1.110563 | 0.0000106 |
| **#370** | CAST | 0.5189092 | 0.004816714 | 0.544316 | 0.0000807 | 0.4313093 | 0.016 | 0.7454898 | 0.00189 | 0.5290131 | 0.000000356 | 0.5015675 | 0.00204 | 0.3617596 | 0.00887 |
| **#371** | ZCCHC2 | 0.6020209 | 0.004836113 | 0.7438363 | 0.000265 | 0.3809702 | 0.0138 | 0.9057439 | 0.00333 | 0.4788559 | 0.00104 | 0.7010536 | 0.000824 | 0.4016657 | 0.00975 |
| **#372** | VPS51 | -0.3614152 | 0.004872231 | -0.4054361 | 0.00185 | -0.3090814 | 0.00544 | -0.4014682 | 0.00811 | -0.2485944 | 0.0114 | -0.4806353 | 0.0000303 | -0.3232759 | 0.00236 |
| **#373** | KDM3A | 0.6916257 | 0.00494595 | 0.6935143 | 0.0000823 | 0.8796828 | 0.0000361 | 0.6281525 | 0.0181 | 0.7059408 | 0.000142 | 0.6437748 | 0.0112 | 0.5986891 | 0.0000969 |
| **#374** | TMEM62 | 0.9960805 | 0.004979518 | 0.9485924 | 0.0000149 | 0.9314213 | 0.00000077 | 0.9499826 | 0.028 | 0.9923878 | 0.000000155 | 1.138044 | 0.0018 | 1.016054 | 0.0000114 |
| **#375** | PCMTD2 | 0.4242053 | 0.004982935 | 0.5383216 | 0.0000918 | 0.41842 | 0.00171 | 0.3988634 | 0.0219 | 0.3574563 | 0.00267 | 0.3711367 | 0.00355 | 0.4610336 | 0.00000102 |
| **#376** | SIRT1 | 0.856163 | 0.00498865 | 0.9718657 | 0.00000308 | 0.6154531 | 0.014 | 0.8665597 | 0.0158 | 0.6772057 | 0.0000377 | 0.947361 | 0.0000155 | 1.058533 | 0.00000594 |
| **#377** | MCPH1 | 0.6587987 | 0.004990653 | 0.7454716 | 8.59E-09 | 0.625354 | 0.0014 | 0.5211352 | 0.0285 | 0.5805037 | 5.11E-08 | 0.8271776 | 0.000000111 | 0.6531498 | 5.66E-10 |
| **#378** | BOLA2 | -0.7963647 | 0.00500237 | -0.5730329 | 0.0182 | -0.8032224 | 0.0000919 | -1.378859 | 0.00182 | -0.4765133 | 0.00597 | -0.8125344 | 0.0014 | -0.7340261 | 0.00252 |
| **#379** | ZFX | 0.9271577 | 0.005009859 | 1.044233 | 4.22E-09 | 0.9878867 | 0.000000036 | 0.5846844 | 0.0301 | 0.8281432 | 3.25E-10 | 1.041525 | 4.59E-11 | 1.076474 | 4.06E-11 |
| **#380** | SENP7 | 0.7594856 | 0.005027252 | 0.7741125 | 0.000000116 | 0.7346953 | 0.0000351 | 0.5901434 | 0.0288 | 0.5308277 | 0.00117 | 0.8856413 | 0.000155 | 1.041493 | 4.58E-08 |
| **#381** | GALNT3 | 0.9071334 | 0.005043835 | 0.7847736 | 0.000646 | 1.117532 | 0.0000722 | 0.9619986 | 0.0224 | 0.9408318 | 0.0000108 | 1.006105 | 0.0000548 | 0.6315587 | 0.00707 |
| **#382** | MIEN1 | -0.3739374 | 0.005126458 | -0.2864845 | 0.00514 | -0.2134283 | 0.0186 | -0.4864614 | 0.00664 | -0.4017142 | 0.0000358 | -0.4769967 | 0.0000625 | -0.3785391 | 0.000298 |
| **#383** | N4BP2 | 0.6243471 | 0.005170559 | 0.5707932 | 0.00117 | 0.6057697 | 0.0157 | 0.4748168 | 0.0133 | 0.5432495 | 0.0000519 | 0.8355294 | 0.000762 | 0.7159239 | 2.62E-09 |
| **#384** | TSPYL4 | 0.5857017 | 0.005185165 | 0.5235238 | 0.000139 | 0.564393 | 0.0196 | 0.5894641 | 0.000954 | 0.4718979 | 0.000922 | 0.4699013 | 0.00949 | 0.8950303 | 0.00000531 |
| **#385** | YTHDC1 | 0.5075929 | 0.00522336 | 0.4674918 | 0.000000542 | 0.5494519 | 0.00000093 | 0.3467452 | 0.031 | 0.4905243 | 0.0000137 | 0.5521888 | 0.000315 | 0.6391553 | 0.000000013 |
| **#386** | ARF5 | -0.5557404 | 0.00526487 | -0.3792906 | 0.00296 | -0.745699 | 0.0000237 | -0.5273482 | 0.0282 | -0.5653106 | 0.000246 | -0.5684692 | 0.000071 | -0.5483248 | 0.0000657 |
| **#387** | OTULIN | 0.7259079 | 0.005280545 | 0.6823435 | 0.000322 | 0.8320436 | 0.0000233 | 0.6694902 | 0.0306 | 0.7479942 | 0.00000777 | 0.7204213 | 0.00067 | 0.7031545 | 0.0000267 |
| **#388** | RFC1 | 0.4124527 | 0.005293896 | 0.366233 | 0.0000214 | 0.4393645 | 0.00265 | 0.4704777 | 0.0291 | 0.3287705 | 0.000000616 | 0.439886 | 0.0000381 | 0.4299843 | 0.000000564 |
| **#389** | TAX1BP1 | 0.3892601 | 0.005304417 | 0.3943985 | 0.0000056 | 0.4508308 | 0.0000181 | 0.2852127 | 0.0318 | 0.3237023 | 0.00000915 | 0.4502386 | 0.00000364 | 0.4311777 | 0.000000788 |
| **#390** | BZW1 | 0.3510733 | 0.005343867 | 0.44619 | 0.00119 | 0.3452301 | 0.000272 | 0.3104721 | 0.0292 | 0.2452852 | 0.000194 | 0.4062654 | 0.000863 | 0.352997 | 0.000387 |
| **#391** | DHX9 | 0.5730226 | 0.005356717 | 0.5376018 | 3.62E-09 | 0.6429183 | 0.000000333 | 0.382022 | 0.0321 | 0.5150419 | 3.77E-10 | 0.6844277 | 2.89E-08 | 0.6761238 | 1.82E-13 |
| **#392** | VAV3 | 0.8426971 | 0.005398727 | 0.8136955 | 0.00015 | 0.9116254 | 0.000000994 | 0.5393886 | 0.0317 | 0.8710411 | 0.0000368 | 0.9464722 | 0.0000209 | 0.9739598 | 0.000507 |
| **#393** | PLCL2 | 0.678423 | 0.005418061 | 0.9206955 | 0.000000153 | 0.5317489 | 0.00183 | 0.4836434 | 0.0306 | 0.5474193 | 0.0000399 | 0.6964944 | 0.0000845 | 0.8905368 | 5.85E-09 |
| **#394** | TASOR | 0.5384927 | 0.005433124 | 0.5821505 | 0.000000829 | 0.5946783 | 0.0000119 | 0.4487732 | 0.0325 | 0.5620545 | 0.00000502 | 0.6211725 | 0.0000442 | 0.4221272 | 0.000000867 |
| **#395** | HIVEP1 | 0.8683123 | 0.005439307 | 0.9060318 | 0.00000012 | 0.6863771 | 0.0295 | 1.053833 | 0.00292 | 0.525631 | 0.00000403 | 1.078752 | 0.000229 | 0.9592482 | 7.71E-08 |
| **#396** | HTT | 0.5478714 | 0.005441527 | 0.3219536 | 0.000717 | 0.6606772 | 0.000000274 | 0.4413226 | 0.0318 | 0.6929732 | 0.00000074 | 0.5497436 | 0.000124 | 0.6205581 | 0.00000589 |
| **#397** | QKI | 0.4821113 | 0.0054476 | 0.4888274 | 0.00638 | 0.549103 | 0.00078 | 0.563495 | 0.00538 | 0.465749 | 0.000559 | 0.4117864 | 0.00588 | 0.413707 | 0.0137 |
| **#398** | DCAF5 | 0.396709 | 0.005448427 | 0.3946067 | 0.00394 | 0.4653851 | 0.0000397 | 0.441551 | 0.0279 | 0.3061915 | 0.000203 | 0.4383646 | 0.000435 | 0.3341549 | 0.000187 |
| **#399** | DHX40 | 0.4355635 | 0.005482491 | 0.443693 | 0.0000432 | 0.4646089 | 0.0000125 | 0.5055129 | 0.0216 | 0.3166338 | 0.000947 | 0.3579866 | 0.0102 | 0.5249455 | 1.58E-08 |
| **#400** | UBXN2A | 0.4660245 | 0.005485576 | 0.3932569 | 0.00115 | 0.6209313 | 0.000373 | 0.5217745 | 0.0294 | 0.4465286 | 0.000111 | 0.4435339 | 0.00188 | 0.3701217 | 0.0000283 |
| **#401** | HSPD1 | 0.3400984 | 0.005516213 | 0.2745285 | 0.0178 | 0.3005773 | 0.00449 | 0.3609203 | 0.0062 | 0.4809667 | 0.00045 | 0.372298 | 0.0000285 | 0.2512992 | 0.00414 |
| **#402** | NUP88 | 0.4496439 | 0.005559134 | 0.4327131 | 0.00000763 | 0.4740245 | 0.00803 | 0.8463326 | 0.0115 | 0.2086607 | 0.0102 | 0.3507446 | 0.00352 | 0.3853876 | 0.000117 |
| **#403** | U2SURP | 0.3112921 | 0.005562816 | 0.3157214 | 0.00000144 | 0.2649547 | 0.00218 | 0.2834841 | 0.0311 | 0.3012675 | 0.000000102 | 0.3487848 | 0.0000486 | 0.35354 | 1.25E-11 |
| **#404** | EWSR1 | 0.3754422 | 0.005570318 | 0.4037599 | 0.00000237 | 0.4801369 | 0.000361 | 0.2998249 | 0.0316 | 0.3390925 | 0.000000233 | 0.4432616 | 0.000796 | 0.2865774 | 0.000674 |
| **#405** | EIF3K | -0.2788809 | 0.005583712 | -0.1985746 | 0.000142 | -0.2966442 | 0.0203 | -0.2953933 | 0.0000629 | -0.2872248 | 0.000449 | -0.3951583 | 0.00807 | -0.2002902 | 0.00443 |
| **#406** | CORO1A | -0.4886561 | 0.005607375 | -0.4806451 | 0.00000476 | -0.4528164 | 0.000118 | -0.9853107 | 0.0249 | -0.3354462 | 0.00000462 | -0.316829 | 0.00861 | -0.3608894 | 0.00000327 |
| **#407** | USP1 | 0.3072852 | 0.005617088 | 0.3182901 | 0.000636 | 0.3156561 | 0.000627 | 0.4819014 | 0.00695 | 0.2512665 | 0.00256 | 0.3488141 | 0.00171 | 0.1277829 | 0.0212 |
| **#408** | TNPO1 | 0.4278007 | 0.00562265 | 0.2554104 | 0.0111 | 0.5223896 | 0.000866 | 0.4141611 | 0.021 | 0.4430044 | 0.000000272 | 0.4261387 | 0.000765 | 0.5057001 | 0.00000375 |
| **#409** | SRSF10 | 0.3543939 | 0.005629037 | 0.3662104 | 0.000448 | 0.4141839 | 0.0044 | 0.2380252 | 0.0287 | 0.3317322 | 0.000000328 | 0.4070811 | 0.000117 | 0.3691309 | 0.000103 |
| **#410** | SPEN | 0.6075294 | 0.005643003 | 0.7844113 | 9.52E-08 | 0.6246013 | 0.0000197 | 0.4676448 | 0.0258 | 0.5628594 | 0.0000103 | 0.6044773 | 0.00804 | 0.6011825 | 0.000000615 |
| **#411** | ZNF280D | 0.5253222 | 0.005656448 | 0.5529282 | 0.0000237 | 0.6470561 | 0.000678 | 0.7054845 | 0.000388 | 0.2346274 | 0.0306 | 0.3980517 | 0.00226 | 0.613785 | 0.00000595 |
| **#412** | RPP25L | -0.8817191 | 0.005695532 | -0.8208513 | 0.0195 | -1.11162 | 0.0000654 | -0.8498429 | 0.0142 | -0.8141483 | 0.000178 | -1.074703 | 0.000129 | -0.6191501 | 0.0000546 |
| **#413** | ACSL3 | 0.6651612 | 0.005741471 | 0.8519205 | 0.000000223 | 0.651699 | 0.00719 | 0.6517984 | 0.0268 | 0.4490288 | 0.000424 | 0.582895 | 0.0000222 | 0.8036255 | 2.83E-08 |
| **#414** | TRNT1 | 0.5428966 | 0.005765088 | 0.6833788 | 0.00000347 | 0.505008 | 0.0097 | 0.6547181 | 0.0145 | 0.3793696 | 0.00168 | 0.4987013 | 0.00858 | 0.5362037 | 0.0000859 |
| **#415** | STK17B | 0.5907353 | 0.005790958 | 0.6394011 | 1.5E-09 | 0.5954831 | 0.0000664 | 0.4256162 | 0.0347 | 0.5605396 | 0.000000113 | 0.6792906 | 0.000000149 | 0.6440811 | 3.01E-11 |
| **#416** | MED21 | 0.4846833 | 0.005814615 | 0.3605908 | 0.0141 | 0.6429762 | 0.00468 | 0.626394 | 0.00679 | 0.3391178 | 0.000303 | 0.3588563 | 0.00901 | 0.5801648 | 0.0000428 |
| **#417** | ATG2B | 0.684705 | 0.005815156 | 0.5749877 | 0.00141 | 0.7724755 | 0.00383 | 0.847176 | 0.0031 | 0.5640301 | 0.000913 | 0.5980351 | 0.0256 | 0.7515257 | 0.00000494 |
| **#418** | LONP2 | 0.3503659 | 0.005827777 | 0.3815541 | 0.000104 | 0.3207013 | 0.000244 | 0.4402618 | 0.0242 | 0.2574645 | 0.000224 | 0.3582223 | 0.00885 | 0.3439914 | 0.00136 |
| **#419** | RPL38 | -0.3375383 | 0.005848534 | -0.3035639 | 0.000928 | -0.2869239 | 0.00351 | -0.2791334 | 0.0295 | -0.276875 | 0.000681 | -0.5447695 | 0.00000571 | -0.333964 | 0.000431 |
| **#420** | PSENEN | -0.4911174 | 0.005854368 | -0.3417927 | 0.0126 | -0.3210863 | 0.0118 | -0.6974185 | 0.00976 | -0.5048825 | 0.000242 | -0.6070014 | 0.00066 | -0.4745227 | 0.0000425 |
| **#421** | PHF6 | 0.4372312 | 0.005891254 | 0.4300191 | 0.00158 | 0.3804002 | 0.0142 | 0.7557575 | 0.00354 | 0.3108745 | 0.0101 | 0.4726601 | 0.000166 | 0.2736754 | 0.00574 |
| **#422** | DCAF1 | 0.4662503 | 0.005903601 | 0.3876756 | 0.00092 | 0.3661673 | 0.00517 | 0.7865493 | 0.0192 | 0.4866761 | 0.00000784 | 0.4363674 | 0.00184 | 0.334066 | 0.00834 |
| **#423** | EEF1G | -0.2297492 | 0.005908184 | -0.2783541 | 0.000129 | -0.240586 | 0.000543 | -0.1629985 | 0.0235 | -0.1669606 | 0.00962 | -0.338615 | 0.00059 | -0.1909813 | 0.00105 |
| **#424** | EDRF1 | 0.6785137 | 0.005934882 | 0.6003569 | 0.00034 | 0.6340252 | 0.0284 | 0.8051631 | 0.00664 | 0.6892839 | 0.00000207 | 0.6852388 | 0.000253 | 0.6570143 | 0.00000616 |

| | | | | | | | | | | | | | | | |
|---|---|---|---|---|---|---|---|---|---|---|---|---|---|---|---|
| #425 | PAPOLA | 0.3045557 | 0.005980616 | 0.1361306 | 0.0152 | 0.3439639 | 0.000000762 | 0.2763635 | 0.0207 | 0.3519788 | 1.75E-08 | 0.3578559 | 0.00000103 | 0.3610416 | 1.15E-09 |
| #426 | AKAP8 | 0.3957093 | 0.005998718 | 0.2765309 | 0.00837 | 0.5005685 | 0.000338 | 0.4833024 | 0.0195 | 0.269576 | 0.0012 | 0.4334619 | 0.00458 | 0.4108158 | 0.00202 |
| #427 | ME2 | 0.5023265 | 0.006003065 | 0.448343 | 0.000445 | 0.2286699 | 0.0342 | 1.000109 | 0.0000144 | 0.3177505 | 0.000507 | 0.47294 | 0.000861 | 0.5461472 | 0.0000102 |
| #428 | SMG1 | 0.6045367 | 0.006058376 | 0.5437338 | 0.0000936 | 0.7465866 | 0.000000454 | 0.4364714 | 0.0307 | 0.7347293 | 0.000000503 | 0.6616621 | 0.00555 | 0.5040371 | 0.0000339 |
| #429 | RAD21 | 0.2616804 | 0.006084468 | 0.3268175 | 0.000000143 | 0.2022501 | 0.00261 | 0.3002115 | 0.0286 | 0.2732323 | 0.000132 | 0.2334125 | 0.00519 | 0.2341583 | 0.0000251 |
| #430 | WBP2 | -0.4387969 | 0.00611469 | -0.3498169 | 0.00321 | -0.2687384 | 0.00109 | -0.8463524 | 0.0324 | -0.314731 | 0.000000197 | -0.4439696 | 0.00000964 | -0.4091732 | 0.00000287 |
| #431 | PFDN5 | -0.2207521 | 0.006187915 | -0.1836489 | 0.000454 | -0.1945969 | 0.0268 | -0.2129091 | 0.00813 | -0.208713 | 0.00044 | -0.3328572 | 0.00123 | -0.1917873 | 0.0000684 |
| #432 | UBA52 | -0.2113105 | 0.006199666 | -0.1778472 | 7.42E-08 | -0.1973343 | 0.00032 | -0.1091384 | 0.0363 | -0.1987203 | 0.0000023 | -0.4263455 | 0.0000349 | -0.158477 | 0.000503 |
| #433 | CD37 | -0.3220842 | 0.006200299 | -0.292513 | 0.0000325 | -0.2171324 | 0.00199 | -0.260274 | 0.0313 | -0.3843768 | 0.00366 | -0.3936186 | 0.000204 | -0.3845905 | 0.000000264 |
| #434 | PDXDC1 | 0.573943 | 0.006216346 | 0.5519004 | 0.000754 | 0.3777205 | 0.00286 | 0.7947 | 0.0042 | 0.6410725 | 2.51E-08 | 0.3732046 | 0.0294 | 0.7050602 | 0.0000758 |
| #435 | MRPS21 | -0.333649 | 0.006265534 | -0.2689604 | 0.0032 | -0.2687776 | 0.00113 | -0.3028593 | 0.0331 | -0.3643906 | 0.00000482 | -0.4978624 | 0.000129 | -0.2990435 | 0.000000513 |
| #436 | SETX | 0.7268503 | 0.006283105 | 0.7852904 | 4.41E-09 | 0.8527795 | 0.000114 | 0.4965548 | 0.0375 | 0.6726475 | 0.00000306 | 0.8668176 | 0.0000479 | 0.6870122 | 4.69E-08 |
| #437 | EIF3L | -0.2594947 | 0.006292171 | -0.246121 | 0.0000599 | -0.2300711 | 0.00972 | -0.2716516 | 0.00764 | -0.2520962 | 0.0199 | -0.3316868 | 0.000408 | -0.2253416 | 0.00000565 |
| #438 | CNST | 0.7973925 | 0.006302214 | 1.020208 | 1.03E-08 | 0.7371399 | 0.00253 | 0.4838448 | 0.0353 | 0.6735605 | 0.00000731 | 0.8501319 | 0.00000126 | 1.01947 | 8.07E-08 |
| #439 | ERAP1 | 0.5178459 | 0.006370318 | 0.3978924 | 0.00928 | 0.7444141 | 0.000719 | 0.5898072 | 0.0142 | 0.6008609 | 0.00165 | 0.4424129 | 0.00193 | 0.3316879 | 0.0105 |
| #440 | AP2S1 | -0.2955742 | 0.006374934 | -0.279492 | 0.0216 | -0.3722866 | 0.00205 | -0.4274013 | 0.004 | -0.1799423 | 0.00353 | -0.2407732 | 0.00693 | -0.27355 | 0.000157 |
| #441 | NRAS | 0.5288577 | 0.006379015 | 0.6573178 | 0.0000137 | 0.538027 | 0.00279 | 0.5846838 | 0.00221 | 0.4915168 | 0.000000586 | 0.3532462 | 0.0333 | 0.5483547 | 0.0000056 |
| #442 | TOPBP1 | 0.4591321 | 0.006379612 | 0.5296671 | 0.0000178 | 0.4811847 | 0.000223 | 0.5216813 | 0.0379 | 0.4333915 | 0.0000165 | 0.4440057 | 0.000134 | 0.3448621 | 0.00000941 |
| #443 | PYCARD | -0.497037 | 0.006384039 | -0.4304469 | 0.00131 | -0.470428 | 0.000022 | -0.4745342 | 0.0303 | -0.3425602 | 0.000102 | -0.586945 | 0.00661 | -0.6773076 | 0.00000776 |
| #444 | CNPPD1 | -0.5197996 | 0.00643596 | -0.2811807 | 0.0292 | -0.3579195 | 0.00596 | -1.134539 | 0.00249 | -0.2605079 | 0.000701 | -0.6177013 | 0.000119 | -0.4669495 | 0.0000953 |
| #445 | PRPF40A | 0.2337475 | 0.006460966 | 0.2915928 | 0.00177 | 0.217202 | 0.00117 | 0.2496036 | 0.027 | 0.2277395 | 0.000467 | 0.2603171 | 0.000709 | 0.1560302 | 0.00768 |
| #446 | FMR1 | 0.5197291 | 0.006506385 | 0.548725 | 0.00551 | 0.3569291 | 0.0191 | 0.7668835 | 0.00329 | 0.4638901 | 0.000796 | 0.4527303 | 0.00995 | 0.5292165 | 0.000411 |
| #447 | RBX1 | -0.3951402 | 0.00653078 | -0.3658386 | 0.000779 | -0.3142007 | 0.0158 | -0.4551978 | 0.0216 | -0.3090446 | 0.000125 | -0.4707382 | 0.000808 | -0.4558214 | 0.0000371 |
| #448 | CSDE1 | 0.3222226 | 0.006574482 | 0.3757555 | 0.000000221 | 0.4107281 | 0.00000201 | 0.1679874 | 0.0388 | 0.3029986 | 0.00000709 | 0.3634969 | 0.000684 | 0.3123692 | 0.00000339 |
| #449 | LBR | 0.5343881 | 0.006683148 | 0.653634 | 3.76E-08 | 0.5313749 | 0.00000011 | 0.5616679 | 0.0213 | 0.2239916 | 0.0188 | 0.5647134 | 0.00000449 | 0.6709471 | 2.13E-08 |
| #450 | NDUFS8 | -0.3101772 | 0.006685797 | -0.2370674 | 0.00632 | -0.3459726 | 0.0000214 | -0.4167176 | 0.0331 | -0.301551 | 0.000618 | -0.3137801 | 0.0001 | -0.2459745 | 0.000000562 |
| #451 | GAPDH | -0.4606875 | 0.006779787 | -0.325928 | 0.000673 | -0.7687966 | 0.013 | -0.2837794 | 0.0232 | -0.4635069 | 0.0000643 | -0.3570957 | 0.00362 | -0.5650182 | 0.0000778 |
| #452 | SERF2 | -0.2555592 | 0.006785895 | -0.3090014 | 9.78E-08 | -0.224513 | 0.000323 | -0.1800371 | 0.0364 | -0.2369197 | 0.00333 | -0.3372405 | 0.000641 | -0.2456437 | 0.000000713 |
| #453 | PPP3R1 | 0.4597272 | 0.006791621 | 0.4410436 | 0.00261 | 0.519144 | 0.000495 | 0.5221881 | 0.0327 | 0.3599336 | 0.000463 | 0.5307137 | 0.000226 | 0.3853404 | 0.00426 |
| #454 | STAG2 | 0.322723 | 0.006800614 | 0.3014334 | 0.00527 | 0.307908 | 0.000415 | 0.4143709 | 0.00591 | 0.2423804 | 0.0194 | 0.2529895 | 0.00985 | 0.4172561 | 0.0000057 |
| #455 | PPP1R15B | 0.6434908 | 0.006824584 | 0.6384889 | 0.000119 | 0.711847 | 0.000204 | 0.4554716 | 0.0406 | 0.5912315 | 0.00000858 | 0.7596076 | 0.000000135 | 0.704298 | 0.00000145 |
| #456 | WDR83OS | -0.3155987 | 0.006846385 | -0.2239934 | 0.000902 | -0.3639044 | 0.00979 | -0.467082 | 0.0013 | -0.2887367 | 1.37E-09 | -0.2741912 | 0.029 | -0.2756845 | 0.000043 |
| #457 | NR1D2 | 0.5685209 | 0.006873103 | 0.5134769 | 0.0000567 | 0.799661 | 0.0000114 | 0.5320078 | 0.0361 | 0.6371323 | 0.000434 | 0.4803522 | 0.00407 | 0.4484954 | 0.000573 |
| #458 | CASC4 | 0.4073176 | 0.006896483 | 0.3494469 | 0.000593 | 0.4831887 | 0.000628 | 0.4556217 | 0.0339 | 0.2738713 | 0.00625 | 0.5359471 | 0.00000415 | 0.3458303 | 0.000000167 |
| #459 | TRANK1 | 0.6601753 | 0.006912136 | 0.9366062 | 0.000725 | 0.5024929 | 0.00609 | 0.6421029 | 0.00905 | 0.7720374 | 0.00497 | 0.5528385 | 0.0204 | 0.5549738 | 0.000222 |
| #460 | TRIM24 | 0.5829541 | 0.006960634 | 0.3926148 | 0.00836 | 0.5240908 | 0.000749 | 0.7519271 | 0.000419 | 0.3202915 | 0.0322 | 0.713691 | 0.0000766 | 0.7951094 | 1.46E-08 |
| #461 | GPSM3 | -0.4683189 | 0.006963856 | -0.5775252 | 0.000000411 | -0.4193664 | 0.00000128 | -0.5070933 | 0.0418 | -0.3770683 | 0.00000325 | -0.5505341 | 0.00000242 | -0.3783262 | 5.13E-09 |
| #462 | INTS11 | -0.4108989 | 0.006999688 | -0.2452478 | 0.0342 | -0.2921208 | 0.0000746 | -0.9888134 | 0.000384 | -0.1942282 | 0.00708 | -0.3339185 | 0.000277 | -0.4110647 | 0.0000119 |
| #463 | DOP1A | 0.8037368 | 0.007034707 | 0.5461273 | 0.00508 | 0.639059 | 0.00565 | 1.147474 | 0.0167 | 0.8052225 | 0.000037 | 0.9110271 | 0.0144 | 0.7735107 | 0.000318 |
| #464 | ZC3H14 | 0.4146988 | 0.00706865 | 0.3209053 | 0.000362 | 0.3535869 | 0.00692 | 0.431512 | 0.0248 | 0.2537376 | 0.0102 | 0.6626647 | 0.0000045 | 0.4657861 | 0.000116 |
| #465 | TOR2A | -0.6346828 | 0.007075023 | -0.4019623 | 0.014 | -0.7311565 | 0.0177 | -0.8403806 | 0.00879 | -0.3290376 | 0.000012 | -0.7888099 | 0.00196 | -0.7167501 | 0.0000103 |
| #466 | MORC3 | 0.6603585 | 0.007079997 | 0.8614047 | 3.49E-09 | 0.6329476 | 0.000125 | 0.4864149 | 0.0423 | 0.5361421 | 0.0000654 | 0.7018303 | 2.75E-08 | 0.7434113 | 5.15E-09 |
| #467 | GTF2F1 | 0.3793753 | 0.007098028 | 0.36958 | 0.000333 | 0.4664017 | 0.000062 | 0.2755507 | 0.0419 | 0.3800822 | 0.0000192 | 0.3895794 | 0.000279 | 0.3950581 | 0.0000031 |
| #468 | SSR1 | 0.2732385 | 0.007110313 | 0.2618201 | 0.00391 | 0.3416312 | 0.0000297 | 0.3640971 | 0.00977 | 0.238701 | 0.0000378 | 0.3057903 | 0.00186 | 0.1273915 | 0.0271 |
| #469 | SLMAP | 0.4370599 | 0.00712122 | 0.4654396 | 0.000371 | 0.5862213 | 0.000000156 | 0.3777321 | 0.0384 | 0.3031436 | 0.00048 | 0.4507362 | 0.00243 | 0.4390866 | 0.00105 |
| #470 | COX6A1 | -0.2456918 | 0.007173049 | -0.2147422 | 0.00882 | -0.1900954 | 0.000154 | -0.2666536 | 0.0276 | -0.2284239 | 0.0021 | -0.3631141 | 0.00379 | -0.2111215 | 0.000615 |
| #471 | GRB2 | 0.3313688 | 0.007185096 | 0.4507325 | 0.000000224 | 0.3975956 | 0.000352 | 0.3333469 | 0.0409 | 0.2687351 | 0.00111 | 0.1819967 | 0.000707 | 0.3558058 | 3.55E-08 |
| #472 | RPS27 | -0.2971707 | 0.007234528 | -0.2549242 | 0.00077 | -0.2363217 | 0.0217 | -0.2396523 | 0.0171 | -0.3113293 | 0.000913 | -0.48391 | 0.0000624 | -0.256887 | 0.00285 |
| #473 | BAZ2B | 0.5228803 | 0.007265885 | 0.5687316 | 0.00000862 | 0.5340883 | 1.06E-08 | 0.5221007 | 0.019 | 0.2917036 | 0.0185 | 0.7300993 | 0.00597 | 0.490558 | 0.0000347 |
| #474 | PARP8 | 0.6080451 | 0.007348828 | 0.5691061 | 0.00111 | 0.8012418 | 0.00000453 | 0.778324 | 0.0106 | 0.3692722 | 0.0303 | 0.5212846 | 0.00208 | 0.6090419 | 0.000000579 |
| #475 | RPRD2 | 0.4612252 | 0.007350178 | 0.4720669 | 0.0000735 | 0.4874012 | 0.00000912 | 0.5111016 | 0.0401 | 0.2763175 | 0.00324 | 0.4750798 | 0.000648 | 0.5453839 | 0.00000074 |
| #476 | USP33 | 0.5675855 | 0.007359666 | 0.644585 | 0.000000269 | 0.5841557 | 0.00000577 | 0.3607255 | 0.0442 | 0.6383506 | 0.000000721 | 0.5834736 | 0.000000953 | 0.5942226 | 6.53E-09 |
| #477 | PLCG2 | 0.9880019 | 0.007386993 | 0.8764754 | 0.000152 | 1.044775 | 0.000899 | 1.132514 | 0.00000728 | 0.9177256 | 0.000468 | 1.110275 | 0.0368 | 0.8462461 | 0.00604 |
| #478 | KTN1 | 0.2649016 | 0.007405403 | 0.2432327 | 0.00297 | 0.3087258 | 0.000111 | 0.3446927 | 0.0193 | 0.2145911 | 0.0000932 | 0.3277876 | 0.00107 | 0.1503798 | 0.0209 |
| #479 | ZNF91 | 0.6547111 | 0.007431383 | 0.6061735 | 0.00000105 | 0.8351252 | 0.000000474 | 0.4334561 | 0.0396 | 0.7865693 | 0.00000102 | 0.522156 | 0.00496 | 0.7447863 | 0.00000093 |
| #480 | HIPK3 | 0.6274856 | 0.007440899 | 0.8372177 | 0.0000113 | 0.6011551 | 0.000856 | 0.4474968 | 0.0412 | 0.4551322 | 0.00214 | 0.7342906 | 0.000449 | 0.6896209 | 1.98E-08 |
| #481 | TLK1 | 0.3424609 | 0.007450183 | 0.2205584 | 0.0115 | 0.3786192 | 0.00126 | 0.3246199 | 0.0319 | 0.4040102 | 0.0000242 | 0.3485599 | 0.0000014 | 0.3783977 | 0.0000233 |
| #482 | TBC1D15 | 0.3779389 | 0.007463422 | 0.3967335 | 0.00363 | 0.488185 | 0.000416 | 0.3438271 | 0.0402 | 0.260541 | 0.000346 | 0.3080962 | 0.00016 | 0.4702505 | 0.0000332 |
| #483 | RLF | 0.7715925 | 0.007485625 | 0.7177757 | 0.0000408 | 0.9055342 | 0.0000346 | 0.472065 | 0.0447 | 0.6298563 | 0.0000533 | 1.088888 | 0.000053 | 0.8154356 | 0.000015 |
| #484 | TRIR | -0.3161858 | 0.007500072 | -0.2413028 | 0.0000232 | -0.280605 | 0.0000159 | -0.2459323 | 0.0441 | -0.3033054 | 0.000774 | -0.4811027 | 0.000111 | -0.3448667 | 0.000000143 |
| #485 | ALG13 | 0.778597 | 0.007501377 | 0.9320595 | 2.87E-08 | 0.7987704 | 0.0000107 | 0.3703511 | 0.045 | 0.5189472 | 0.00000119 | 1.030404 | 6.8E-09 | 1.02105 | 4.58E-13 |
| #486 | RPL34 | -0.2305995 | 0.007533891 | -0.2279056 | 0.0000228 | -0.1481162 | 0.0106 | -0.2290549 | 0.0325 | -0.2152187 | 0.000735 | -0.3773969 | 0.000072 | -0.1859045 | 0.00127 |
| #487 | RPS5 | -0.2667092 | 0.007556392 | -0.3253866 | 5.68E-08 | -0.2030022 | 0.000537 | -0.1738884 | 0.0441 | -0.2403443 | 0.000284 | -0.4415897 | 0.000207 | -0.2160442 | 0.000178 |
| #488 | CCP110 | 0.4494478 | 0.007564851 | 0.5093106 | 0.00131 | 0.3352947 | 0.00191 | 0.5085087 | 0.0364 | 0.4534548 | 0.00273 | 0.5596655 | 0.00278 | 0.3304522 | 0.00024 |
| #489 | UBA5 | 0.3711311 | 0.007619335 | 0.4036472 | 0.00667 | 0.4538459 | 0.016 | 0.4335038 | 0.0191 | 0.2939906 | 0.00124 | 0.3268681 | 0.00271 | 0.314931 | 0.0000176 |
| #490 | MTRF1 | 0.5995442 | 0.007625235 | 0.6989924 | 0.0000513 | 0.6317474 | 0.00349 | 0.9141331 | 0.0217 | 0.3964974 | 0.00407 | 0.5582608 | 0.0164 | 0.3976338 | 0.0000153 |
| #491 | NONO | 0.2563431 | 0.007738849 | 0.2643841 | 0.000217 | 0.3021723 | 0.00000505 | 0.2876198 | 0.0361 | 0.1501647 | 0.0101 | 0.3478475 | 0.00000035 | 0.1858702 | 0.00000948 |
| #492 | PTEN | 0.4507208 | 0.007740092 | 0.3642087 | 0.00656 | 0.4885457 | 0.000181 | 0.7478852 | 0.000543 | 0.2258428 | 0.035 | 0.4259448 | 0.00185 | 0.4518974 | 0.0023 |
| #493 | GOLGA5 | 0.485456 | 0.007786343 | 0.5104899 | 0.00152 | 0.4189848 | 0.00125 | 0.6520406 | 0.041 | 0.475131 | 0.0000155 | 0.4622093 | 0.0029 | 0.3938808 | 0.0000382 |
| #494 | RLIM | 0.6663567 | 0.007811293 | 0.8301655 | 0.00000767 | 0.6992545 | 0.0000146 | 0.4692676 | 0.0461 | 0.6481148 | 6.17E-09 | 0.5881049 | 0.000701 | 0.7632331 | 2.86E-08 |
| #495 | RPL19 | -0.1968147 | 0.007813881 | -0.179133 | 0.0000355 | -0.1648991 | 0.0118 | -0.1336525 | 0.0322 | -0.1729756 | 0.00144 | -0.3647731 | 0.000739 | -0.165455 | 0.000695 |

| | | | | | | | | | | | | | | | |
|---|---|---|---|---|---|---|---|---|---|---|---|---|---|---|---|
| #496 | ETAA1 | 0.5175964 | 0.007830056 | 0.4759832 | 0.0000614 | 0.508416 | 0.0159 | 0.6658206 | 0.0143 | 0.2887118 | 0.0123 | 0.5551838 | 0.00439 | 0.6114632 | 0.00000177 |
| #497 | NQO2 | -0.8739463 | 0.007856335 | -0.5769539 | 0.0207 | -0.8585252 | 0.00533 | -1.246669 | 0.0197 | -0.9445294 | 0.0011 | -0.7956244 | 0.000232 | -0.8213765 | 0.0000294 |
| #498 | TRIM23 | 0.5539791 | 0.007871606 | 0.6767731 | 0.0000275 | 0.6861815 | 0.00399 | 0.6117347 | 0.0136 | 0.3016455 | 0.00983 | 0.3431001 | 0.0198 | 0.7044395 | 0.000000468 |
| #499 | RPL37 | -0.2778896 | 0.007896935 | -0.2808438 | 0.000265 | -0.2389539 | 0.00231 | -0.2225047 | 0.0424 | -0.2599429 | 0.000559 | -0.4051774 | 0.000407 | -0.2599147 | 0.0014 |
| #500 | ANKRD36C | 0.6838261 | 0.007903708 | 0.5320807 | 0.000875 | 0.8968403 | 0.000626 | 0.763384 | 0.000524 | 0.4665903 | 0.0127 | 0.7703587 | 0.032 | 0.6737025 | 0.000707 |
| #501 | POLR2L | -0.4042433 | 0.0079596 | -0.4741387 | 0.00126 | -0.3845196 | 0.00000984 | -0.3702857 | 0.0343 | -0.4291119 | 0.00000561 | -0.3470305 | 0.0122 | -0.420373 | 0.000085 |
| #502 | MTLN | -0.7025349 | 0.008014121 | -0.6133753 | 0.015 | -0.5148389 | 0.00599 | -1.034695 | 0.0271 | -0.5342757 | 0.0000146 | -0.7843456 | 0.0000185 | -0.7336784 | 0.0000314 |
| #503 | TP53BP2 | 0.8794009 | 0.008070228 | 1.026065 | 8.65E-08 | 1.082674 | 0.00000014 | 0.6321703 | 0.0483 | 0.7378671 | 0.0000276 | 0.8970523 | 0.0000616 | 0.9005768 | 0.00000459 |
| #504 | CFLAR | 0.3423298 | 0.008101731 | 0.571052 | 0.0000531 | 0.2813355 | 0.00128 | 0.5280489 | 0.00494 | 0.2581804 | 0.00408 | 0.2346462 | 0.0198 | 0.1807161 | 0.0184 |
| #505 | EXOC8 | 0.8161724 | 0.008110243 | 1.063852 | 0.0000125 | 0.9500384 | 0.00000346 | 0.4409387 | 0.0479 | 0.6402605 | 0.00000353 | 0.7942324 | 0.000788 | 1.007712 | 8.77E-09 |
| #506 | STAU1 | 0.2618375 | 0.008136138 | 0.3463878 | 0.00124 | 0.2091324 | 0.00341 | 0.5238557 | 0.00242 | 0.1582344 | 0.00366 | 0.1717774 | 0.0363 | 0.161637 | 0.00184 |
| #507 | WDR7 | 0.6172933 | 0.008145416 | 0.5650514 | 0.000626 | 0.7242061 | 0.0000217 | 0.7253723 | 0.0243 | 0.5689787 | 0.00000483 | 0.4356641 | 0.0239 | 0.6844872 | 0.000000014 |
| #508 | RUFY2 | 0.4293524 | 0.008168994 | 0.4824538 | 0.00101 | 0.360372 | 0.0206 | 0.5514177 | 0.0196 | 0.3983274 | 0.000522 | 0.4929426 | 0.00226 | 0.2906008 | 0.005 |
| #509 | PARG | 0.9555856 | 0.008188175 | 0.9654672 | 3.11E-08 | 1.051973 | 0.000000361 | 0.6175465 | 0.0486 | 0.8410658 | 0.0000488 | 1.128603 | 0.000431 | 1.128857 | 0.000000174 |
| #510 | TPP2 | 0.5462254 | 0.008189888 | 0.5811925 | 0.0000263 | 0.5558745 | 0.000014 | 0.3695789 | 0.0477 | 0.5123977 | 0.00000172 | 0.5882536 | 0.00134 | 0.670055 | 0.000011 |
| #511 | FRYL | 0.584082 | 0.008233888 | 0.5438394 | 0.0000669 | 0.8196973 | 0.00122 | 0.3787177 | 0.0447 | 0.611368 | 0.00000568 | 0.5756436 | 0.00345 | 0.5752262 | 0.00000286 |
| #512 | RESF1 | 0.6785439 | 0.008240306 | 0.8832552 | 0.00000118 | 0.7468303 | 0.0000311 | 0.7158939 | 0.000742 | 0.3272672 | 0.0484 | 0.6513339 | 0.000241 | 0.7466829 | 6.46E-08 |
| #513 | CDV3 | 0.5301206 | 0.008289309 | 0.6276224 | 0.00454 | 0.5111708 | 0.00622 | 0.5574626 | 0.00605 | 0.5181523 | 0.00579 | 0.5268245 | 0.00763 | 0.439491 | 0.0195 |
| #514 | REST | 0.3184302 | 0.008335068 | 0.185681 | 0.0157 | 0.4517293 | 0.00132 | 0.3667259 | 0.00255 | 0.417173 | 0.0000157 | 0.1931922 | 0.0305 | 0.29608 | 0.0000115 |
| #515 | ZNF207 | 0.3546619 | 0.008370473 | 0.3839349 | 0.00023 | 0.3733345 | 0.00108 | 0.3420025 | 0.0414 | 0.2676867 | 0.00017 | 0.302224 | 0.00731 | 0.4587885 | 0.00000632 |
| #516 | EIF6 | -0.3629974 | 0.008371082 | -0.2887315 | 0.000896 | -0.3759998 | 0.00000203 | -0.3810836 | 0.0436 | -0.2765008 | 0.00374 | -0.4189746 | 0.00196 | -0.436694 | 0.000021 |
| #517 | MRPL28 | -0.3823998 | 0.008376203 | -0.3369417 | 0.00515 | -0.2227573 | 0.00139 | -0.8517248 | 0.000992 | -0.1725049 | 0.0425 | -0.3704474 | 0.00000321 | -0.3400227 | 0.000224 |
| #518 | ATXN7 | 0.4781111 | 0.00839764 | 0.5198729 | 0.0000355 | 0.3391953 | 0.0014 | 0.8028702 | 0.000412 | 0.4033242 | 0.0158 | 0.3331878 | 0.0327 | 0.4702165 | 0.0000129 |
| #519 | CDK13 | 0.5161673 | 0.00841138 | 0.6012737 | 0.000132 | 0.687281 | 0.00000035 | 0.3412375 | 0.0456 | 0.392229 | 0.00024 | 0.4808255 | 0.00437 | 0.5941569 | 0.000138 |
| #520 | MAP4K5 | 0.4952415 | 0.008416878 | 0.5882273 | 0.00000602 | 0.4405264 | 0.0000411 | 0.3818167 | 0.0501 | 0.4931617 | 0.00000317 | 0.5168217 | 0.000304 | 0.5508955 | 0.00000183 |
| #521 | SAFB | 0.3392344 | 0.008419381 | 0.2551064 | 0.00257 | 0.2586567 | 0.02 | 0.3719575 | 0.0273 | 0.3849804 | 0.0000327 | 0.4457162 | 0.00014 | 0.3189891 | 0.000505 |
| #522 | ZNF805 | 0.764962 | 0.008457161 | 0.5801876 | 0.000326 | 1.047226 | 0.000843 | 0.526631 | 0.0492 | 0.6946222 | 0.0000372 | 0.9566011 | 0.000232 | 0.7845039 | 0.0000843 |
| #523 | TTC3 | 0.4212248 | 0.008488433 | 0.3106103 | 0.0000293 | 0.5402101 | 0.0000639 | 0.2866636 | 0.047 | 0.3880681 | 0.00366 | 0.4663837 | 0.000144 | 0.5354127 | 0.000000153 |
| #524 | RB1 | 0.4824178 | 0.008492283 | 0.5504458 | 0.000000877 | 0.5099146 | 0.0000285 | 0.4401333 | 0.0467 | 0.483744 | 0.00333 | 0.3723239 | 0.000914 | 0.5379453 | 0.00000264 |
| #525 | PPP4R3A | 0.6959603 | 0.008505493 | 0.8023487 | 1.72E-08 | 0.8031294 | 1.89E-08 | 0.3640041 | 0.051 | 0.7686216 | 7.59E-11 | 0.7793419 | 0.000000168 | 0.6583159 | 0.000000191 |
| #526 | JAK1 | 0.4580422 | 0.008565001 | 0.4669687 | 0.0000305 | 0.4666231 | 0.00000154 | 0.392981 | 0.0513 | 0.4087236 | 0.0000049 | 0.4997115 | 0.0000119 | 0.513245 | 0.00000712 |
| #527 | YOD1 | 0.6931318 | 0.008621867 | 0.438572 | 0.0384 | 0.6686933 | 0.00282 | 0.9003487 | 0.00616 | 0.640373 | 0.000214 | 0.7344026 | 0.00382 | 0.7764014 | 0.00033 |
| #528 | RAPGEF6 | 0.5010965 | 0.008715842 | 0.4192697 | 0.0000339 | 0.430377 | 0.0000725 | 0.5094177 | 0.0508 | 0.3772403 | 0.00109 | 0.565562 | 0.00000915 | 0.7047122 | 0.0000308 |
| #529 | NDUFA7 | -0.3203184 | 0.008727108 | -0.3847533 | 0.000734 | -0.2267716 | 0.0385 | -0.3744277 | 0.00567 | -0.3584529 | 0.00000123 | -0.3010221 | 0.00746 | -0.2764828 | 0.00000874 |
| #530 | CHORDC1 | 0.4848508 | 0.008755769 | 0.4501218 | 0.000139 | 0.4509923 | 0.0037 | 0.555445 | 0.0484 | 0.4216108 | 0.000176 | 0.6333826 | 0.00000491 | 0.3975525 | 0.0001 |
| #531 | CAPRIN2 | 0.671381 | 0.008771149 | 0.5171556 | 0.00431 | 0.877018 | 0.00277 | 0.8045537 | 0.0138 | 0.4335013 | 0.0136 | 0.6944655 | 0.0181 | 0.7015917 | 0.0000166 |
| #532 | PDIK1L | 0.670027 | 0.008814097 | 0.5019778 | 0.0204 | 0.7689132 | 0.000474 | 0.6371595 | 0.0297 | 0.5831217 | 0.000738 | 0.9805519 | 0.000254 | 0.5484377 | 0.00135 |
| #533 | DCUN1D4 | 0.6185188 | 0.008837633 | 0.6939843 | 0.000809 | 0.6653288 | 0.00176 | 0.5831618 | 0.0355 | 0.4692782 | 0.00963 | 0.6714105 | 0.00523 | 0.6279492 | 0.0000884 |
| #534 | ATF6 | 0.343371 | 0.008863106 | 0.2718867 | 0.0225 | 0.2785623 | 0.00439 | 0.4745956 | 0.0208 | 0.1909023 | 0.00333 | 0.3716168 | 0.00222 | 0.4726626 | 0.0000425 |
| #535 | DENND4C | 0.4978416 | 0.008954732 | 0.4891452 | 0.0005 | 0.6124365 | 0.000918 | 0.4861076 | 0.0284 | 0.5299197 | 0.00493 | 0.3762347 | 0.0189 | 0.4932059 | 0.0000924 |
| #536 | ANP32E | 0.2959626 | 0.008965145 | 0.1489594 | 0.00237 | 0.212121 | 0.000143 | 0.3868719 | 0.0505 | 0.5005479 | 0.0000453 | 0.2883496 | 0.000727 | 0.2389256 | 0.0000313 |
| #537 | ERCC6L2 | 0.6419302 | 0.009009229 | 0.6983249 | 0.000000207 | 0.9181279 | 0.000172 | 0.3627463 | 0.0536 | 0.4680435 | 0.000019 | 0.6933258 | 0.000247 | 0.7110128 | 8.65E-09 |
| #538 | DOCK11 | 0.5403976 | 0.009020653 | 0.5172464 | 0.0000623 | 0.4083516 | 0.00452 | 0.4132625 | 0.0494 | 0.6080753 | 1.29E-09 | 0.7109973 | 0.0000304 | 0.5844526 | 0.0000681 |
| #539 | RPL27 | -0.1671378 | 0.009021764 | -0.1601088 | 1.07E-08 | -0.1431064 | 0.00000141 | -0.1219195 | 0.0507 | -0.0958656 | 0.00253 | -0.3441201 | 0.000385 | -0.1377066 | 0.000495 |
| #540 | GNA13 | 0.6690706 | 0.009043032 | 0.8252595 | 0.00000342 | 0.7071581 | 0.0000864 | 0.3958257 | 0.0541 | 0.698925 | 0.000000167 | 0.8242647 | 0.0000836 | 0.5629905 | 0.0000283 |
| #541 | GPCPD1 | 0.4619692 | 0.009049093 | 0.5013816 | 0.0000151 | 0.522745 | 0.000384 | 0.3752804 | 0.0411 | 0.5117022 | 0.000176 | 0.3717757 | 0.0126 | 0.4889306 | 0.000000512 |
| #542 | VPS13B | 0.8267986 | 0.009063821 | 1.022257 | 4.36E-10 | 0.7364234 | 3.36E-09 | 0.5537317 | 0.045 | 0.9576665 | 0.000000019 | 0.9106888 | 0.00937 | 0.7800244 | 8.84E-08 |
| #543 | ATP5MF | -0.3372585 | 0.009079491 | -0.3344287 | 0.00638 | -0.3577911 | 0.000109 | -0.3213294 | 0.0238 | -0.2984262 | 0.00175 | -0.3432224 | 0.0224 | -0.3683534 | 0.0000492 |
| #544 | ERVK3-1 | 0.3285136 | 0.00909739 | 0.3150521 | 0.000352 | 0.3417137 | 0.00413 | 0.5554809 | 0.00889 | 0.2310422 | 0.00407 | 0.2046921 | 0.0367 | 0.3231004 | 0.00045 |
| #545 | R3HDM1 | 0.4489717 | 0.009101701 | 0.4724742 | 0.000256 | 0.3875265 | 0.0387 | 0.5024636 | 0.0152 | 0.3415503 | 0.000148 | 0.449296 | 0.000558 | 0.5405197 | 5.04E-09 |
| #546 | TIMM10 | -0.6650892 | 0.009194048 | -0.4007773 | 0.0343 | -0.5160515 | 0.0000945 | -1.195593 | 0.0168 | -0.5264452 | 0.000334 | -0.8553627 | 0.00033 | -0.4963052 | 0.00332 |
| #547 | RALY | -0.2299235 | 0.009206246 | -0.1777632 | 0.024 | -0.1197488 | 0.0118 | -0.4011944 | 0.00208 | -0.1360528 | 0.00918 | -0.3258297 | 0.00177 | -0.2189519 | 0.00636 |
| #548 | RBM25 | 0.3874514 | 0.009216209 | 0.4496718 | 0.0000163 | 0.3983421 | 0.00035 | 0.239485 | 0.0543 | 0.4509227 | 0.00000486 | 0.3443069 | 0.000664 | 0.4419799 | 0.00000274 |
| #549 | MOB1B | 0.5812038 | 0.009220394 | 0.6894263 | 0.000533 | 0.6775645 | 0.000201 | 0.737846 | 0.0127 | 0.3049843 | 0.0354 | 0.3738788 | 0.00645 | 0.703523 | 6.55E-08 |
| #550 | LIG3 | 0.5312193 | 0.009228706 | 0.400726 | 0.0367 | 0.6583322 | 0.000284 | 0.5260423 | 0.0129 | 0.4846651 | 0.000275 | 0.5215794 | 0.00519 | 0.595971 | 0.000000139 |
| #551 | TMEM134 | -0.4170783 | 0.009240792 | -0.2874198 | 0.00138 | -0.2743585 | 0.00384 | -0.6976197 | 0.0183 | -0.413768 | 0.0312 | -0.5027771 | 0.000137 | -0.3265267 | 0.000567 |
| #552 | SOS 1 | 0.5547 | 0.009324653 | 0.6559001 | 0.00246 | 0.4388182 | 0.0141 | 0.5547031 | 0.0338 | 0.4743249 | 0.0054 | 0.7097586 | 0.0000994 | 0.4946948 | 0.0000868 |
| #553 | SOCS4 | 0.70258 | 0.009328895 | 0.7851304 | 0.0000141 | 0.8221291 | 0.000019 | 0.5627086 | 0.0557 | 0.4745196 | 0.000239 | 0.7997406 | 0.000028 | 0.7712517 | 8.99E-08 |
| #554 | CRNKL1 | 0.6311304 | 0.009362836 | 0.768799 | 0.00000153 | 0.5423197 | 0.000274 | 0.5650325 | 0.0532 | 0.3907382 | 0.00272 | 0.7700367 | 0.0000137 | 0.7498564 | 0.000000764 |
| #555 | DBF4 | 0.5847036 | 0.009364039 | 0.7335053 | 0.000436 | 0.5644419 | 0.0184 | 0.5842627 | 0.0343 | 0.5466198 | 0.000798 | 0.6121743 | 0.00181 | 0.4672174 | 0.000361 |
| #556 | HIST1H2BE | 0.7556036 | 0.009364108 | 0.6237304 | 0.0153 | 0.853843 | 0.00111 | 0.7425791 | 0.0319 | 0.656218 | 0.000555 | 0.7760789 | 0.00721 | 0.8811721 | 0.0000595 |
| #557 | NKTR | 0.5242182 | 0.009413445 | 0.5761601 | 7.16E-08 | 0.6577533 | 1.68E-10 | 0.2960502 | 0.0236 | 0.5054633 | 0.000577 | 0.5657451 | 0.0322 | 0.5441371 | 0.000127 |
| #558 | TRAPPC10 | 0.6125462 | 0.009484144 | 0.641202 | 0.00152 | 0.7013123 | 0.0000874 | 0.4860986 | 0.0529 | 0.666859 | 0.0000992 | 0.5819038 | 0.000682 | 0.5979016 | 0.00163 |
| #559 | EFCAB2 | 0.7958393 | 0.009514062 | 1.120385 | 0.00000385 | 1.069364 | 2.05E-08 | 0.565353 | 0.0371 | 0.5454469 | 0.00189 | 0.5496181 | 0.0181 | 0.9248691 | 0.0000115 |
| #560 | PET100 | -0.3112346 | 0.009517475 | -0.1943871 | 0.0335 | -0.3245822 | 0.013 | -0.2880789 | 0.00988 | -0.4641359 | 0.000000644 | -0.3619779 | 0.0000972 | -0.2342453 | 0.000656 |
| #561 | FBXO11 | 0.4396541 | 0.009557336 | 0.4408796 | 0.000408 | 0.4250929 | 0.000348 | 0.3938358 | 0.0447 | 0.454118 | 0.00000404 | 0.4556523 | 0.0118 | 0.4683462 | 0.0000983 |
| #562 | HERC1 | 0.4870291 | 0.009586498 | 0.3449922 | 0.00317 | 0.4858668 | 0.000602 | 0.3939014 | 0.0454 | 0.5811156 | 0.0000888 | 0.5854032 | 0.00827 | 0.5308952 | 0.00000314 |
| #563 | ZRANB1 | 0.5080473 | 0.009635653 | 0.4171434 | 0.00949 | 0.5668108 | 0.0000337 | 0.4554546 | 0.0403 | 0.5627209 | 0.0000329 | 0.4519282 | 0.00793 | 0.5942258 | 0.0000215 |
| #564 | COMT | -0.4358927 | 0.00964456 | -0.4587655 | 0.00027 | -0.3623662 | 0.00202 | -0.3174177 | 0.0546 | -0.5502928 | 0.000464 | -0.4812433 | 0.000235 | -0.4452706 | 0.000317 |
| #565 | NFAT5 | 0.5542784 | 0.009700201 | 0.6763274 | 0.0000223 | 0.6376794 | 0.0192 | 0.5514838 | 0.021 | 0.6265821 | 0.0000234 | 0.4366622 | 0.0145 | 0.3969355 | 0.00344 |
| #566 | RPS4X | 0.3275186 | 0.009705346 | 0.3225126 | 0.0000088 | 0.4215703 | 0.0000268 | 0.487912 | 0.00000317 | 0.1890991 | 0.00858 | 0.193009 | 0.0496 | 0.3510086 | 0.000013 |

| | | | | | | | | | | | | | | | |
|---|---|---|---|---|---|---|---|---|---|---|---|---|---|---|---|
| #567 | LY75 | 0.6793943 | 0.009738188 | 0.8627264 | 0.00000138 | 0.5800816 | 0.0000261 | 0.4041533 | 0.0317 | 0.7033742 | 0.0264 | 0.7311052 | 0.000297 | 0.7949248 | 0.00000118 |
| #568 | TLN1 | 0.2358904 | 0.009745594 | 0.154045 | 0.0313 | 0.1708971 | 0.00371 | 0.2480604 | 0.0165 | 0.2780928 | 0.00000162 | 0.3372538 | 0.00326 | 0.2269936 | 0.00369 |
| #569 | ZNF107 | 0.8294713 | 0.009766733 | 0.6813151 | 0.0000128 | 0.9061764 | 0.0124 | 1.208531 | 0.00222 | 0.6950233 | 0.00107 | 0.5271814 | 0.0429 | 0.9586007 | 2.2E-11 |
| #570 | EDF1 | -0.212023 | 0.009780495 | -0.1580664 | 0.000814 | -0.1488654 | 0.00245 | -0.2171464 | 0.0542 | -0.2274655 | 0.00001 | -0.2728324 | 0.000987 | -0.2477618 | 0.000208 |
| #571 | MAGED2 | -0.5039252 | 0.009795395 | -0.5735718 | 0.007 | -0.6247659 | 0.0000479 | -0.5001705 | 0.0158 | -0.298828 | 0.0316 | -0.4350188 | 0.00433 | -0.5911962 | 2.45E-08 |
| #572 | VKORC1 | -0.4946888 | 0.009869283 | -0.5676723 | 0.0273 | -0.2625599 | 0.00463 | -0.5276757 | 0.021 | -0.3916118 | 0.0000255 | -0.5484148 | 0.00627 | -0.6701981 | 0.00000448 |
| #573 | APC | 0.5292018 | 0.009871877 | 0.4607042 | 0.0000995 | 0.4412948 | 0.00016 | 0.4407499 | 0.0586 | 0.5082285 | 0.000117 | 0.6754788 | 0.000268 | 0.6487546 | 1.23E-08 |
| #574 | VCPIP1 | 0.6024009 | 0.009886505 | 0.9276626 | 0.000176 | 0.6246148 | 0.000632 | 0.4533403 | 0.047 | 0.5496172 | 0.00145 | 0.5927765 | 0.000439 | 0.466394 | 0.00967 |
| #575 | DDX24 | 0.375161 | 0.009895064 | 0.4519965 | 0.000000295 | 0.5142849 | 0.00495 | 0.2834811 | 0.0542 | 0.2374917 | 0.0000242 | 0.384282 | 0.000176 | 0.3794297 | 4.04E-09 |
| #576 | CAPNS1 | -0.3917475 | 0.009927542 | -0.269533 | 0.0501 | -0.3725539 | 0.000228 | -0.4026434 | 0.00905 | -0.3266056 | 0.00000147 | -0.4895923 | 0.000128 | -0.4895569 | 0.0000276 |
| #577 | ARCN1 | 0.4830977 | 0.009941924 | 0.4176435 | 0.00181 | 0.5333129 | 0.00305 | 0.5141881 | 0.0419 | 0.2804817 | 0.0121 | 0.4890026 | 0.000789 | 0.6639574 | 0.000000056 |
| #578 | RPS2 | -0.2319781 | 0.009947665 | -0.2330984 | 0.00000526 | -0.1990566 | 0.00597 | -0.137986 | 0.0523 | -0.277651 | 0.0000677 | -0.3338831 | 0.0013 | -0.2101938 | 0.0000882 |
| #579 | FASTKD2 | 0.5125084 | 0.01007171 | 0.4359168 | 0.00142 | 0.3545621 | 0.015 | 0.4617894 | 0.0435 | 0.4951068 | 0.0000569 | 0.6210004 | 0.00046 | 0.7066748 | 9.92E-08 |
| #580 | EMB | 0.5750938 | 0.01021382 | 0.5560192 | 8.24E-09 | 0.6887764 | 0.000289 | 0.7415005 | 0.00447 | 0.2921056 | 0.0564 | 0.6571571 | 0.000142 | 0.5150043 | 1.87E-08 |
| #581 | CCDC124 | -0.2851624 | 0.01022463 | -0.2230473 | 0.029 | -0.2959599 | 0.0000057 | -0.4706223 | 0.0159 | -0.1572791 | 0.00184 | -0.3265193 | 0.0109 | -0.2375463 | 0.00371 |
| #582 | GABARAP | -0.3538499 | 0.01023653 | -0.2802578 | 0.000000686 | -0.3487141 | 0.0000236 | -0.4115453 | 0.0369 | -0.2384316 | 0.0245 | -0.4819255 | 0.0000149 | -0.3622254 | 1.05E-08 |
| #583 | USP36 | 0.5319285 | 0.01027696 | 0.6337423 | 0.000154 | 0.5720851 | 0.0000169 | 0.4197773 | 0.0551 | 0.483919 | 0.000219 | 0.648819 | 0.00132 | 0.4332282 | 0.00482 |
| #584 | B4GALT7 | -0.5183114 | 0.01027971 | -0.4234664 | 0.00094 | -0.47984 | 0.00278 | -0.8237735 | 0.0477 | -0.3410917 | 0.00000884 | -0.5643366 | 0.00987 | -0.47736 | 0.000358 |
| #585 | TP53BP1 | 0.6048792 | 0.01028927 | 0.6861211 | 0.0000264 | 0.6737945 | 0.00125 | 0.5176962 | 0.0537 | 0.6861221 | 0.000409 | 0.5355595 | 0.00583 | 0.5299816 | 0.00048 |
| #586 | RPS21 | -0.2917689 | 0.01030423 | -0.2665719 | 0.000969 | -0.2668448 | 0.00134 | -0.2523131 | 0.0526 | -0.2461918 | 0.000968 | -0.4806132 | 0.000163 | -0.2380784 | 0.00576 |
| #587 | VPS4A | 0.333415 | 0.01031681 | 0.2988884 | 0.00544 | 0.2599509 | 0.000174 | 0.3741626 | 0.0539 | 0.304226 | 0.0000383 | 0.5369922 | 0.0000531 | 0.2262701 | 0.00231 |
| #588 | CNTRL | 0.393466 | 0.01032243 | 0.5623842 | 0.00000254 | 0.2891524 | 0.0131 | 0.3840446 | 0.0139 | 0.3502085 | 0.00336 | 0.4187769 | 0.0296 | 0.3562297 | 0.00196 |
| #589 | LPGAT1 | 0.5445995 | 0.01032586 | 0.6323186 | 0.00013 | 0.5608931 | 0.0000391 | 0.4881211 | 0.0485 | 0.4826487 | 0.00958 | 0.6224467 | 0.00022 | 0.4811693 | 0.00346 |
| #590 | PPP2R5E | 0.2693673 | 0.01033432 | 0.2929817 | 0.00183 | 0.3197688 | 0.00232 | 0.3290504 | 0.0463 | 0.2605293 | 0.000525 | 0.2186138 | 0.00987 | 0.1952596 | 0.00117 |
| #591 | EIF3D | -0.2135784 | 0.0103433 | -0.1677873 | 0.000429 | -0.1621249 | 0.0406 | -0.244612 | 0.0073 | -0.1921713 | 0.0135 | -0.3288536 | 0.000102 | -0.1859213 | 0.000137 |
| #592 | LRRC75A | -0.8953168 | 0.01035295 | -0.9589594 | 0.00433 | -0.9369566 | 0.00352 | -0.7980386 | 0.05 | -1.021281 | 0.000452 | -0.7862015 | 0.000673 | -0.8704634 | 0.00314 |
| #593 | MTREX | 0.4665052 | 0.01036036 | 0.469286 | 0.00000701 | 0.5119128 | 0.00166 | 0.4814258 | 0.0549 | 0.394303 | 0.000474 | 0.5102529 | 0.00508 | 0.4318508 | 0.00000694 |
| #594 | MPHOSPH9 | 0.5372919 | 0.01036844 | 0.4247772 | 0.0000891 | 0.7054091 | 0.0000105 | 0.3361621 | 0.0615 | 0.5271533 | 0.00036 | 0.6106546 | 0.000241 | 0.6195954 | 0.0000409 |
| #595 | GPR108 | -0.4503623 | 0.01047164 | -0.3749535 | 0.0188 | -0.4334646 | 0.000912 | -0.638675 | 0.0399 | -0.4081651 | 0.00000318 | -0.3778555 | 0.00326 | -0.4690598 | 0.0000103 |
| #596 | LMBR1 | 0.6871434 | 0.0105226 | 0.5906195 | 0.00118 | 0.719844 | 0.00000352 | 0.5638324 | 0.0328 | 0.6461192 | 0.00000438 | 0.5014107 | 0.0292 | 1.101035 | 0.000000151 |
| #597 | USP34 | 0.5998221 | 0.01054578 | 0.6758096 | 9.57E-11 | 0.5531087 | 0.000000111 | 0.4418671 | 0.0625 | 0.5735102 | 9.67E-09 | 0.6954136 | 0.00074 | 0.6592234 | 1.34E-09 |
| #598 | EIF2A | 0.3039259 | 0.01054669 | 0.3410031 | 0.000000107 | 0.3569983 | 0.00000223 | 0.2728516 | 0.0535 | 0.2410039 | 0.00594 | 0.2773175 | 0.00386 | 0.3343813 | 0.000000108 |
| #599 | PDK1 | 0.7190713 | 0.01056667 | 0.3492401 | 0.0328 | 0.9235257 | 0.00184 | 0.9377309 | 0.0249 | 0.7085192 | 0.000463 | 0.5282592 | 0.00278 | 0.8671523 | 0.000542 |
| #600 | DCTN4 | 0.7927989 | 0.01059028 | 0.8374097 | 0.000000144 | 0.9677651 | 0.000000134 | 0.5049423 | 0.0635 | 0.5392058 | 0.000000161 | 0.9408521 | 0.0000395 | 0.9666181 | 4.11E-08 |
| #601 | NCOA1 | 0.3850596 | 0.01067586 | 0.3262133 | 0.00917 | 0.5079138 | 0.000144 | 0.3652385 | 0.0516 | 0.2215916 | 0.0000887 | 0.4324153 | 0.00296 | 0.4569851 | 0.0000868 |
| #602 | PITPNB | 0.4219256 | 0.01069222 | 0.4540963 | 0.0000221 | 0.4788655 | 0.00068 | 0.4415411 | 0.0631 | 0.4565707 | 0.00000119 | 0.3868577 | 0.000315 | 0.3136225 | 0.000000295 |
| #603 | PARP4 | 0.5151067 | 0.01070576 | 0.6808689 | 0.000000459 | 0.5520515 | 0.0000393 | 0.387906 | 0.064 | 0.5726358 | 0.000000861 | 0.5345001 | 0.0000551 | 0.3626777 | 0.000151 |
| #604 | NRDC | 0.5357808 | 0.01071419 | 0.5603434 | 0.000000217 | 0.604119 | 0.00117 | 0.3107733 | 0.0631 | 0.5208149 | 0.000000242 | 0.5221916 | 0.0000317 | 0.6964428 | 0.000000025 |
| #605 | SLC25A5 | -0.3240252 | 0.01071668 | -0.2651182 | 0.00794 | -0.3447478 | 0.0014 | -0.419982 | 0.0114 | -0.4171243 | 0.0000694 | -0.3329886 | 0.0153 | -0.1641906 | 0.0282 |
| #606 | B3GNT2 | 0.6251315 | 0.01073262 | 0.4981018 | 0.00551 | 0.7395637 | 0.00000194 | 0.5397622 | 0.0586 | 0.5496882 | 0.0000432 | 0.6734697 | 0.00024 | 0.7502035 | 0.0000429 |
| #607 | ACAP2 | 0.3488454 | 0.01074641 | 0.3220902 | 0.00000433 | 0.422189 | 5.8E-09 | 0.2850514 | 0.0616 | 0.266055 | 0.00287 | 0.379489 | 0.00000128 | 0.418198 | 3.04E-09 |
| #608 | ZC3H15 | 0.2887073 | 0.0107712 | 0.3275164 | 0.000128 | 0.2761301 | 0.000837 | 0.2317875 | 0.0636 | 0.2156079 | 0.0000191 | 0.3711616 | 0.000000113 | 0.31004 | 0.000000154 |
| #609 | DNAJC2 | 0.3118969 | 0.01081169 | 0.3124524 | 0.00366 | 0.2941322 | 0.00034 | 0.3750205 | 0.05 | 0.3353236 | 0.00000262 | 0.296197 | 0.0101 | 0.2582556 | 0.000767 |
| #610 | PJA2 | 0.443585 | 0.01082891 | 0.5107239 | 0.0000547 | 0.3840234 | 0.0000526 | 0.3071864 | 0.0649 | 0.3870008 | 0.000000909 | 0.5412811 | 0.00000149 | 0.5312943 | 1.58E-08 |
| #611 | MAP3K2 | 0.4197613 | 0.01085764 | 0.4671146 | 0.000217 | 0.5767399 | 0.000228 | 0.4562076 | 0.0581 | 0.2634756 | 0.00426 | 0.360355 | 0.00233 | 0.3946749 | 0.00000491 |
| #612 | OGT | 0.5810068 | 0.01087098 | 0.6388277 | 0.00000186 | 0.6953574 | 0.0154 | 0.3460156 | 0.0346 | 0.4855042 | 0.0114 | 0.6285929 | 0.00383 | 0.6917429 | 0.0000334 |
| #613 | MED13L | 0.5317478 | 0.01087684 | 0.5113697 | 0.000000283 | 0.6623357 | 0.0000234 | 0.410525 | 0.0629 | 0.557924 | 5.58E-10 | 0.4752222 | 0.00236 | 0.5731105 | 0.000000131 |
| #614 | LNPEP | 0.5557679 | 0.01091634 | 0.6689936 | 0.000000885 | 0.6495911 | 0.000269 | 0.3461839 | 0.0652 | 0.5464236 | 0.000000867 | 0.5402118 | 0.000000279 | 0.5832033 | 0.00000176 |
| #615 | UQCRQ | -0.4536478 | 0.01091864 | -0.4984597 | 0.000171 | -0.4769963 | 0.000486 | -0.3629527 | 0.0636 | -0.4986973 | 0.000932 | -0.4604998 | 0.000281 | -0.4242811 | 0.0000696 |
| #616 | ZNF510 | 0.8298788 | 0.0109196 | 0.9930265 | 0.000000612 | 0.5344103 | 0.0388 | 1.244899 | 0.000788 | 0.7165807 | 0.0000804 | 0.5181552 | 0.0259 | 0.9722011 | 9.95E-09 |
| #617 | DIAPH2 | 0.4875134 | 0.01095043 | 0.3827754 | 0.000717 | 0.593575 | 0.0000347 | 0.6149217 | 0.013 | 0.6827673 | 0.000000073 | 0.4417554 | 0.0227 | 0.2092859 | 0.0293 |
| #618 | AP2M1 | -0.3029488 | 0.01096398 | -0.3667712 | 0.00000238 | -0.2899963 | 0.0000376 | -0.226299 | 0.0613 | -0.2709175 | 0.000000503 | -0.3420552 | 0.00437 | -0.3216536 | 0.0000781 |
| #619 | PRKACB | 0.4558387 | 0.01102132 | 0.4498692 | 0.00021 | 0.4013315 | 0.0238 | 0.6833919 | 0.00275 | 0.2755934 | 0.039 | 0.4574161 | 0.00014 | 0.4674303 | 0.000198 |
| #620 | ZC3HAV1 | 0.5395022 | 0.0110217 | 0.5572749 | 0.00000289 | 0.7443376 | 0.00000525 | 0.2757064 | 0.0633 | 0.6611062 | 0.00285 | 0.4402615 | 0.00000111 | 0.5583268 | 3.54E-09 |
| #621 | SAMD9 | 0.6301919 | 0.01106863 | 0.5228475 | 0.00583 | 0.6184065 | 0.011 | 0.7127128 | 0.00871 | 0.5644003 | 0.04 | 0.7278061 | 0.00000804 | 0.6349784 | 0.000855 |
| #622 | RPS6KA1 | -0.5759723 | 0.01107942 | -0.5956512 | 0.000983 | -0.5091597 | 0.00000644 | -1.046735 | 0.0487 | -0.2338248 | 0.0132 | -0.4487139 | 0.0036 | -0.6217491 | 0.00000153 |
| #623 | CYLD | 0.5690266 | 0.01110632 | 0.6497774 | 0.0000132 | 0.6270926 | 0.000000279 | 0.3966873 | 0.0666 | 0.6208068 | 0.000000374 | 0.6617524 | 0.00000105 | 0.4580428 | 0.000000181 |
| #624 | BORCS5 | 0.500741 | 0.01111354 | 0.2823561 | 0.0341 | 0.4129238 | 0.0171 | 0.6803274 | 0.0102 | 0.7468076 | 0.000000104 | 0.3401856 | 0.00508 | 0.5418454 | 0.000109 |
| #625 | DOK2 | -1.000179 | 0.01111355 | -1.261386 | 0.000588 | -1.450932 | 0.051 | -1.919535 | 0.0109 | -0.3716898 | 0.00115 | -0.3787264 | 0.00306 | -0.6188016 | 0.0000377 |
| #626 | ASH1L | 0.5879892 | 0.01123266 | 0.6059131 | 1.14E-09 | 0.6313043 | 3.65E-08 | 0.4079171 | 0.0599 | 0.5805706 | 0.0000007 | 0.6558955 | 0.00752 | 0.6463344 | 0.000000157 |
| #627 | UBLCP1 | 0.4091722 | 0.01130942 | 0.4168441 | 0.000118 | 0.4339448 | 0.0404 | 0.5005929 | 0.00894 | 0.3555083 | 0.000192 | 0.3967568 | 0.0181 | 0.3513865 | 0.0000932 |
| #628 | TMEM33 | 0.3036616 | 0.01131389 | 0.4156425 | 0.000327 | 0.2347832 | 0.0061 | 0.4190492 | 0.0141 | 0.2161995 | 0.0172 | 0.2622693 | 0.0286 | 0.2740258 | 0.0015 |
| #629 | ZBTB10 | 0.8906559 | 0.01133213 | 0.9131476 | 0.000000278 | 1.031043 | 0.00208 | 0.6311127 | 0.0659 | 0.771216 | 0.0000159 | 1.001691 | 0.000000658 | 0.9957259 | 0.0000242 |
| #630 | FIP1L1 | 0.3541793 | 0.01135553 | 0.3670756 | 0.00216 | 0.3908235 | 0.0177 | 0.3537374 | 0.0481 | 0.3266716 | 0.0000152 | 0.3379975 | 0.000132 | 0.3487703 | 5.91E-08 |
| #631 | UBXN4 | 0.2585407 | 0.01143576 | 0.2926723 | 0.000885 | 0.2808533 | 0.012 | 0.1780491 | 0.0402 | 0.1882471 | 0.0154 | 0.3115354 | 0.00014 | 0.2998868 | 0.00000452 |
| #632 | NBN | 0.4124137 | 0.01145621 | 0.3790812 | 0.00334 | 0.3514075 | 0.00135 | 0.3261244 | 0.063 | 0.4916568 | 0.0000116 | 0.468103 | 0.000812 | 0.4581093 | 0.000202 |
| #633 | FKBP8 | -0.4512382 | 0.01149575 | -0.3299071 | 0.00111 | -0.5516098 | 0.0000438 | -0.4282284 | 0.0678 | -0.3860842 | 0.0000076 | -0.5582549 | 0.00000895 | -0.4533447 | 0.00000864 |
| #634 | CUEDC2 | -0.4120494 | 0.01154246 | -0.3390732 | 0.000858 | -0.4626191 | 9.82E-08 | -0.4352808 | 0.0672 | -0.3364802 | 0.000763 | -0.52726 | 0.000368 | -0.3715833 | 0.0000189 |
| #635 | NGLY1 | 0.4270506 | 0.01159386 | 0.5171332 | 8.35E-08 | 0.5509437 | 0.0435 | 0.3229749 | 0.0254 | 0.4754405 | 0.0000104 | 0.3675151 | 0.000322 | 0.3282964 | 0.000327 |
| #636 | RIOK2 | 0.5424801 | 0.01163887 | 0.4550443 | 0.000181 | 0.7350485 | 0.000558 | 0.3551932 | 0.0663 | 0.5297343 | 0.0000831 | 0.6086601 | 0.00262 | 0.5712 | 0.000127 |
| #637 | AEBP2 | 0.3982727 | 0.01167478 | 0.2555109 | 0.0125 | 0.5217353 | 0.000259 | 0.4259668 | 0.0347 | 0.4093085 | 0.000108 | 0.3233536 | 0.0225 | 0.4537609 | 0.000000166 |

| | | | | | | | | | | | | | | | |
|---|---|---|---|---|---|---|---|---|---|---|---|---|---|---|---|
| **#638** | CPTP | -0.912572 | 0.01167963 | -0.6989586 | 0.00357 | -0.8841045 | 0.00463 | -1.060829 | 0.0618 | -0.5581126 | 0.0000103 | -1.157721 | 0.0000215 | -1.115706 | 0.000000376 |
| **#639** | ATP5F1E | -0.2562206 | 0.0116939 | -0.2350871 | 0.000833 | -0.1783074 | 0.0546 | -0.3681569 | 0.000198 | -0.2004607 | 0.00329 | -0.2758003 | 0.0112 | -0.2795115 | 0.00000344 |
| **#640** | NOP14 | 0.3649841 | 0.01176928 | 0.2406555 | 0.00765 | 0.3549838 | 0.0027 | 0.4304934 | 0.0583 | 0.3752315 | 0.000324 | 0.519663 | 0.00162 | 0.2688772 | 0.00000757 |
| **#641** | PURB | 0.4767074 | 0.0117713 | 0.5534704 | 0.000568 | 0.4509793 | 0.000889 | 0.4127936 | 0.0355 | 0.3606108 | 0.000241 | 0.4008824 | 0.0334 | 0.681508 | 0.0000168 |
| **#642** | CHM | 0.488886 | 0.01179113 | 0.5216205 | 0.000114 | 0.3953443 | 0.0132 | 0.7651364 | 0.00635 | 0.2716265 | 0.0357 | 0.4587108 | 0.0154 | 0.5208776 | 0.0000155 |
| **#643** | FAU | -0.2081254 | 0.01185903 | -0.1655138 | 0.000378 | -0.1340825 | 0.00293 | -0.1419125 | 0.0663 | -0.2275946 | 0.000123 | -0.3910904 | 0.000919 | -0.1885588 | 0.000506 |
| **#644** | SF3A2 | -0.419539 | 0.01187347 | -0.4194355 | 0.00368 | -0.2974651 | 0.0252 | -0.6031369 | 0.0256 | -0.3428204 | 0.00761 | -0.3678637 | 0.009 | -0.4865128 | 0.00011 |
| **#645** | RNF38 | 0.4963879 | 0.01190654 | 0.8270429 | 0.0000002 | 0.4880162 | 0.0000919 | 0.3382947 | 0.0713 | 0.3475923 | 0.0000123 | 0.4672915 | 0.0000836 | 0.5100898 | 0.00000102 |
| **#646** | PSMD8 | -0.3205216 | 0.01192071 | -0.304511 | 0.000264 | -0.2953747 | 3.36E-08 | -0.2698436 | 0.0648 | -0.3282349 | 0.000996 | -0.3500626 | 0.00545 | -0.3751032 | 0.0000102 |
| **#647** | ABCE1 | 0.3551926 | 0.01192827 | 0.2617604 | 0.00247 | 0.4463654 | 0.003 | 0.3509933 | 0.0642 | 0.3585022 | 0.000303 | 0.3770489 | 0.00157 | 0.3364856 | 0.0000106 |
| **#648** | PIK3R1 | 0.4554637 | 0.01192947 | 0.2155714 | 0.0397 | 0.4840966 | 0.00514 | 0.680474 | 0.00817 | 0.5604551 | 0.00786 | 0.3921904 | 0.00416 | 0.3999944 | 0.00655 |
| **#649** | PPM1A | 0.3572144 | 0.01193048 | 0.3349843 | 0.000593 | 0.4615522 | 0.00176 | 0.2947077 | 0.0613 | 0.3621437 | 0.000601 | 0.4234203 | 0.000967 | 0.2664783 | 0.00638 |
| **#650** | TAF1 | 0.3954907 | 0.01195185 | 0.3415265 | 0.00221 | 0.4036202 | 0.00218 | 0.5380871 | 0.0327 | 0.3725859 | 0.000343 | 0.3675688 | 0.0328 | 0.3495558 | 0.00142 |
| **#651** | HBP1 | 0.4400132 | 0.01195968 | 0.6875356 | 0.0000701 | 0.3618606 | 0.00346 | 0.2485813 | 0.0681 | 0.2802235 | 0.0000507 | 0.5785875 | 0.00000383 | 0.4832905 | 0.0000548 |
| **#652** | ZNF117 | 0.7561319 | 0.01196018 | 0.7432169 | 0.00229 | 0.8916819 | 0.0012 | 0.6265333 | 0.0285 | 1.014232 | 3.28E-08 | 0.7229195 | 0.034 | 0.5382073 | 0.00578 |
| **#653** | XRN1 | 0.7948862 | 0.01196962 | 0.8751956 | 0.000387 | 0.8555881 | 0.00433 | 0.4495883 | 0.0478 | 0.7943365 | 0.0193 | 0.8823431 | 0.00000546 | 0.9122654 | 0.00000168 |
| **#654** | PEF1 | -0.45399 | 0.01197329 | -0.437434 | 0.0615 | -0.3667131 | 0.00185 | -0.7101349 | 0.00598 | -0.3559771 | 0.0000187 | -0.5107635 | 0.00218 | -0.3429177 | 0.000305 |
| **#655** | USP15 | 0.5660687 | 0.01199643 | 0.6444285 | 0.000284 | 0.6651278 | 0.00000756 | 0.3414872 | 0.0555 | 0.2904258 | 0.0162 | 0.6992578 | 0.0000271 | 0.7556851 | 0.000000323 |
| **#656** | FEM1C | 0.8972087 | 0.01200873 | 0.9997472 | 0.0000966 | 0.8691873 | 0.000603 | 0.6212399 | 0.0713 | 0.9501764 | 9.68E-09 | 1.141288 | 0.0000102 | 0.8016131 | 0.000000279 |
| **#657** | CHCHD2 | -0.3003548 | 0.01205907 | -0.299892 | 0.000901 | -0.2077877 | 0.0357 | -0.2043213 | 0.0239 | -0.348098 | 0.0000469 | -0.3804072 | 0.0115 | -0.3616227 | 0.000188 |
| **#658** | PGAM1 | -0.4954436 | 0.01207205 | -0.5347798 | 0.000386 | -0.4253044 | 0.00183 | -0.5458119 | 0.0619 | -0.2837373 | 0.00753 | -0.5204424 | 0.000783 | -0.6625858 | 0.0000184 |
| **#659** | COQ8B | -0.7247667 | 0.01218111 | -0.7859817 | 0.000658 | -0.5660334 | 0.000681 | -0.8284979 | 0.0712 | -0.5305671 | 0.000121 | -1.029361 | 0.000318 | -0.6081593 | 0.0000758 |
| **#660** | ACTR2 | 0.2799047 | 0.01219323 | 0.3139379 | 0.00000016 | 0.2652884 | 0.00000795 | 0.2673218 | 0.0725 | 0.2464146 | 0.000221 | 0.3793185 | 0.000000101 | 0.207147 | 0.000465 |
| **#661** | SMAD4 | 0.5655116 | 0.01228612 | 0.7042338 | 0.0000102 | 0.4107775 | 0.011 | 0.5063096 | 0.0329 | 0.6102158 | 0.00000283 | 0.3605091 | 0.0298 | 0.8010235 | 0.000000497 |
| **#662** | SEPTIN2 | 0.3869735 | 0.01232588 | 0.3303069 | 0.00885 | 0.4544263 | 0.0000146 | 0.2960558 | 0.0447 | 0.4667829 | 6.65E-09 | 0.4595802 | 0.000245 | 0.3146889 | 0.0202 |
| **#663** | DIP2B | 0.5497513 | 0.01233047 | 0.5332505 | 0.00000296 | 0.4244043 | 0.00126 | 0.5544884 | 0.00231 | 0.6205136 | 0.0000198 | 0.448949 | 0.0704 | 0.716902 | 0.00000954 |
| **#664** | CCSER2 | 0.4856102 | 0.01242009 | 0.4582356 | 0.0018 | 0.5035272 | 0.00068 | 0.4903223 | 0.0526 | 0.4864576 | 0.00000993 | 0.4579476 | 0.0188 | 0.5171708 | 0.000604 |
| **#665** | MAN1A2 | 0.4097304 | 0.01247095 | 0.5415656 | 0.000000254 | 0.3131746 | 0.000905 | 0.2875438 | 0.0721 | 0.2578433 | 0.00178 | 0.5945245 | 0.00000887 | 0.4637307 | 4.35E-08 |
| **#666** | LAMTOR5 | -0.3229853 | 0.01253311 | -0.3827992 | 0.0000161 | -0.2496567 | 0.0138 | -0.4473733 | 0.023 | -0.4317 | 0.000000591 | -0.1811033 | 0.0377 | -0.2452796 | 0.000633 |
| **#667** | PEPD | -0.4405816 | 0.01254667 | -0.4388368 | 0.00525 | -0.3272057 | 0.000458 | -0.7657123 | 0.0637 | -0.3239656 | 0.000163 | -0.4619471 | 0.00488 | -0.3258219 | 0.000857 |
| **#668** | SPOPL | 0.4885166 | 0.01255021 | 0.6157626 | 0.000264 | 0.438789 | 0.0103 | 0.5394182 | 0.0215 | 0.6093311 | 0.0000519 | 0.4247209 | 0.0284 | 0.303078 | 0.0148 |
| **#669** | CDC73 | 0.3809691 | 0.01258716 | 0.4010976 | 0.000413 | 0.4271487 | 0.00561 | 0.2676301 | 0.0682 | 0.4059328 | 0.0000163 | 0.3270482 | 0.00128 | 0.456957 | 0.000000968 |
| **#670** | SET | 0.2441937 | 0.01260701 | 0.2042111 | 0.0317 | 0.2208409 | 0.00295 | 0.2419039 | 0.0149 | 0.2955803 | 0.000106 | 0.269654 | 0.00943 | 0.2329718 | 0.0166 |
| **#671** | IDH3B | -0.294132 | 0.01263215 | -0.2107333 | 0.0266 | -0.2666371 | 0.000931 | -0.4996179 | 0.0398 | -0.2248017 | 5.72E-08 | -0.37949 | 0.00264 | -0.1835118 | 0.00579 |
| **#672** | RPLP2 | -0.1691681 | 0.01264685 | -0.1532011 | 0.00501 | -0.1424452 | 0.0131 | -0.1237953 | 0.0481 | -0.09655138 | 0.00678 | -0.3362317 | 0.000266 | -0.1627842 | 0.00263 |
| **#673** | NDUFB2 | -0.3056651 | 0.01265518 | -0.2915491 | 0.00298 | -0.3813666 | 0.0000973 | -0.226225 | 0.0666 | -0.2876353 | 0.000569 | -0.334196 | 0.00556 | -0.3130184 | 0.000109 |
| **#674** | GTF2I | 0.3573886 | 0.01285086 | 0.3995529 | 0.00103 | 0.4400363 | 0.000576 | 0.3549596 | 0.00567 | 0.1653974 | 0.0575 | 0.3018417 | 0.0122 | 0.4825435 | 0.0000973 |
| **#675** | MZT2A | -0.5173352 | 0.01285088 | -0.3901353 | 0.0395 | -0.4707699 | 0.0000842 | -0.3123332 | 0.0373 | -0.5448553 | 0.000029 | -0.871452 | 0.00000249 | -0.5144658 | 0.000115 |
| **#676** | G3BP2 | 0.4232045 | 0.01288278 | 0.5552791 | 0.0000133 | 0.4590198 | 2.48E-09 | 0.2150335 | 0.0773 | 0.4270636 | 0.0000019 | 0.4160705 | 0.00000014 | 0.4667602 | 0.00000613 |
| **#677** | SNU13 | -0.2856925 | 0.01292678 | -0.2097336 | 0.00161 | -0.161301 | 0.0304 | -0.3478188 | 0.0438 | -0.3334666 | 0.000000047 | -0.3428903 | 0.00176 | -0.3189444 | 0.0000015 |
| **#678** | DNAJB14 | 0.4643003 | 0.01296692 | 0.4334321 | 0.0000004 | 0.7062313 | 0.0000106 | 0.2805964 | 0.0778 | 0.4250078 | 0.000000576 | 0.5336248 | 0.0000249 | 0.4069096 | 0.00000412 |
| **#679** | RACK1 | -0.239092 | 0.01302527 | -0.2420514 | 0.0000683 | -0.1991082 | 0.0134 | -0.1195964 | 0.0642 | -0.3258869 | 0.000183 | -0.3439655 | 0.000227 | -0.2039433 | 0.000074 |
| **#680** | TMEM256 | -0.3312359 | 0.01303045 | -0.2159159 | 0.00973 | -0.3885632 | 0.000000337 | -0.3213115 | 0.0656 | -0.307131 | 0.000833 | -0.4761549 | 0.002 | -0.2783388 | 0.0000388 |
| **#681** | UBR7 | 0.5061432 | 0.01303732 | 0.6984847 | 0.0000201 | 0.5529399 | 0.00721 | 0.3698279 | 0.0702 | 0.4046601 | 0.000106 | 0.6500744 | 0.000235 | 0.3608725 | 0.000488 |
| **#682** | HIGD2A | -0.3910918 | 0.01306458 | -0.3615644 | 0.00849 | -0.4020164 | 0.00361 | -0.3955198 | 0.0524 | -0.3791041 | 0.00685 | -0.4732371 | 0.00134 | -0.3351088 | 0.00566 |
| **#683** | SENP2 | 0.5310156 | 0.01309782 | 0.5720614 | 0.0000731 | 0.6834059 | 0.00363 | 0.404058 | 0.0572 | 0.4978021 | 0.000000575 | 0.4065166 | 0.0176 | 0.6222494 | 0.0000121 |
| **#684** | EIF2S3 | 0.5190712 | 0.01315679 | 0.6801412 | 0.00000001 | 0.4175091 | 0.00000145 | 0.4316312 | 0.000968 | 0.1610245 | 0.078 | 0.7613627 | 5.94E-12 | 0.6627585 | 2.25E-14 |
| **#685** | ATF2 | 0.4323383 | 0.01318999 | 0.5363768 | 0.000584 | 0.4231633 | 0.0122 | 0.7244016 | 0.0033 | 0.3188178 | 0.0291 | 0.240015 | 0.0333 | 0.3512553 | 0.000732 |
| **#686** | RPL37A | -0.3432557 | 0.01319046 | -0.3575928 | 0.000342 | -0.2933511 | 0.00436 | -0.2957311 | 0.0631 | -0.3276292 | 0.00104 | -0.4940536 | 0.000121 | -0.2911766 | 0.0101 |
| **#687** | SPSB3 | -0.3839879 | 0.01319816 | -0.204388 | 0.0158 | -0.4203303 | 0.0000188 | -0.6155005 | 0.0398 | -0.2922631 | 0.0022 | -0.3365692 | 0.0212 | -0.4348762 | 0.000223 |
| **#688** | ARHGDIB | -0.2807507 | 0.01319982 | -0.2784321 | 0.0000228 | -0.4349465 | 0.000134 | -0.164696 | 0.0208 | -0.3491688 | 0.000123 | -0.2498218 | 0.055 | -0.2074389 | 0.0031 |
| **#689** | STX10 | -0.4066399 | 0.01322702 | -0.2578476 | 0.0146 | -0.6145789 | 0.0283 | -0.369294 | 0.0314 | -0.4578121 | 0.000292 | -0.4409325 | 0.00473 | -0.299374 | 0.0000543 |
| **#690** | WDFY1 | 0.6391667 | 0.01325506 | 1.081615 | 0.000637 | 0.7416364 | 0.000000208 | 0.4891424 | 0.0626 | 0.5064782 | 0.0155 | 0.4459717 | 0.000736 | 0.5701565 | 0.0000862 |
| **#691** | NPM1 | 0.2179203 | 0.01325557 | 0.2088402 | 0.0000851 | 0.2699363 | 0.000506 | 0.2700962 | 0.000803 | 0.1686444 | 0.01 | 0.2030676 | 0.0637 | 0.1869374 | 0.00443 |
| **#692** | SOS 2 | 0.7021013 | 0.01328593 | 0.7075101 | 0.00021 | 0.7713413 | 0.000000513 | 0.3448226 | 0.0789 | 0.6056527 | 0.000419 | 0.920022 | 0.00011 | 0.8632588 | 0.000119 |
| **#693** | MCL1 | 0.5065291 | 0.01329595 | 0.667876 | 0.00798 | 0.5106457 | 0.000834 | 0.3113747 | 0.0694 | 0.3762679 | 0.00141 | 0.6666857 | 0.000104 | 0.5063249 | 0.0000617 |
| **#694** | SKIL | 0.461243 | 0.01332396 | 0.2637146 | 0.0429 | 0.6568714 | 0.000567 | 0.5013836 | 0.0302 | 0.5105911 | 0.0000535 | 0.3841508 | 0.0023 | 0.4507464 | 0.0039 |
| **#695** | MIS18BP1 | 0.2863001 | 0.01334238 | 0.2802921 | 0.0000397 | 0.3077029 | 0.000192 | 0.3079567 | 0.0524 | 0.247866 | 0.00667 | 0.3656617 | 0.000143 | 0.2083213 | 0.0206 |
| **#696** | ZDHHC5 | 0.7070063 | 0.01334872 | 0.9321621 | 0.0000483 | 0.6032247 | 0.00576 | 0.4708262 | 0.074 | 0.5057926 | 0.0000525 | 0.8853061 | 0.000278 | 0.8447261 | 0.000000899 |
| **#697** | TCERG1 | 0.3771139 | 0.0133771 | 0.3456082 | 0.000126 | 0.3258504 | 0.000000255 | 0.3266026 | 0.0768 | 0.4011997 | 0.0000245 | 0.5100948 | 0.00333 | 0.3533277 | 0.0000101 |
| **#698** | PSMC5 | -0.2607689 | 0.01339065 | -0.2640597 | 0.000894 | -0.2850415 | 0.0000315 | -0.3983756 | 0.0132 | -0.1618929 | 0.0111 | -0.216597 | 0.0548 | -0.2386469 | 0.000322 |
| **#699** | WTAP | 0.2505344 | 0.01345712 | 0.2425647 | 0.00928 | 0.223312 | 0.000539 | 0.3626409 | 0.0192 | 0.3053296 | 0.0000264 | 0.2204574 | 0.00515 | 0.1489016 | 0.0465 |
| **#700** | ZNF518B | 0.6287831 | 0.01354437 | 0.7764493 | 0.00000211 | 0.4925327 | 0.000305 | 0.4965021 | 0.0739 | 0.635506 | 0.00000662 | 0.4433026 | 0.00703 | 0.9284056 | 1.77E-08 |
| **#701** | HPS3 | 0.4847278 | 0.01361274 | 0.372008 | 0.00813 | 0.5392616 | 0.0000552 | 0.406227 | 0.0729 | 0.553661 | 0.00000305 | 0.6098456 | 0.00000411 | 0.4273636 | 0.000616 |
| **#702** | PCNP | 0.2315222 | 0.01370212 | 0.2323168 | 0.0000734 | 0.2581005 | 0.00104 | 0.2038211 | 0.0541 | 0.2524289 | 0.00000435 | 0.2687025 | 0.0148 | 0.1737635 | 0.0122 |
| **#703** | SLK | 0.5709858 | 0.01374661 | 0.6693986 | 0.0000756 | 0.6029672 | 0.000000693 | 0.3859218 | 0.0813 | 0.601828 | 0.0000216 | 0.64237 | 0.000146 | 0.5234292 | 0.00093 |
| **#704** | CDK17 | 0.9394534 | 0.01375753 | 1.115882 | 3.57E-10 | 0.9276487 | 0.00000742 | 0.5056655 | 0.0825 | 0.9711297 | 7.43E-08 | 1.091503 | 0.00000883 | 1.024892 | 0.000000107 |
| **#705** | TMPO | 0.4720336 | 0.01376637 | 0.4584253 | 0.0000348 | 0.5449149 | 0.000286 | 0.3071306 | 0.0384 | 0.3220958 | 0.0438 | 0.5199404 | 0.00014 | 0.6796944 | 0.000000003 |
| **#706** | UBA6 | 0.4300569 | 0.01382536 | 0.3919953 | 0.00229 | 0.4785188 | 0.000626 | 0.5519441 | 0.00882 | 0.5459252 | 0.000492 | 0.380382 | 0.00864 | 0.2315757 | 0.0621 |
| **#707** | POLR3C | 0.389702 | 0.01386334 | 0.380035 | 0.00299 | 0.3762189 | 0.0273 | 0.3433989 | 0.0496 | 0.289825 | 0.00222 | 0.6595024 | 0.00000373 | 0.2892316 | 0.00107 |
| **#708** | NPIPB5 | 0.8231277 | 0.01387304 | 0.792924 | 0.000232 | 1.085703 | 0.00000168 | 0.527713 | 0.0403 | 0.937093 | 0.0000209 | 0.7532177 | 0.0422 | 0.8421152 | 0.000496 |

| | | | | | | | | | | | | | | | |
|---|---|---|---|---|---|---|---|---|---|---|---|---|---|---|---|
| #709 | SMG6 | 0.5562095 | 0.01389879 | 0.7021881 | 0.000000301 | 0.4550473 | 0.00193 | 0.474157 | 0.0697 | 0.507906 | 0.0000673 | 0.5824285 | 0.0117 | 0.6155304 | 0.00000743 |
| #710 | EGLN2 | -0.5878347 | 0.01397825 | -0.5086016 | 0.0124 | -0.554453 | 0.000843 | -1.191641 | 0.00111 | -0.3811472 | 0.00121 | -0.3455256 | 0.0669 | -0.5456401 | 0.00151 |
| #711 | LSM 2.00 | -0.3487198 | 0.01400159 | -0.2382032 | 0.00778 | -0.385548 | 0.0000284 | -0.2528736 | 0.0734 | -0.3581076 | 0.0000579 | -0.5158226 | 0.00234 | -0.3417639 | 0.000428 |
| #712 | EIF4G1 | 0.3349471 | 0.01408169 | 0.1981239 | 0.045 | 0.3769291 | 0.0000847 | 0.3935024 | 0.0101 | 0.361053 | 0.000395 | 0.3683162 | 0.0283 | 0.311758 | 0.000629 |
| #713 | BTF3 | -0.2127026 | 0.01409749 | -0.1341709 | 0.00728 | -0.2739747 | 0.0168 | -0.1643567 | 0.0293 | -0.2974817 | 0.0000775 | -0.255257 | 0.0266 | -0.1509744 | 0.00459 |
| #714 | CCDC115 | -0.4603189 | 0.01418159 | -0.3413757 | 0.000131 | -0.3410367 | 0.0000104 | -0.5723647 | 0.0805 | -0.3916468 | 0.00263 | -0.570522 | 0.00182 | -0.5449676 | 0.000000109 |
| #715 | TTC19 | 0.3048924 | 0.01420802 | 0.179707 | 0.0324 | 0.446477 | 0.0000528 | 0.3626063 | 0.0217 | 0.3857674 | 0.00194 | 0.1929984 | 0.0287 | 0.2617981 | 0.000449 |
| #716 | LRPPRC | 0.5686016 | 0.01426355 | 0.5022335 | 0.0000178 | 0.7138961 | 0.000000563 | 0.2895761 | 0.0856 | 0.6037754 | 0.000000809 | 0.6456558 | 0.000000758 | 0.6564726 | 1.26E-09 |
| #717 | ERBIN | 0.4902442 | 0.01426818 | 0.5112518 | 0.00000302 | 0.6342446 | 0.000000999 | 0.2990476 | 0.0856 | 0.4811876 | 0.0000271 | 0.5095736 | 0.0000111 | 0.5061601 | 0.00000148 |
| #718 | MANBA | 0.3756715 | 0.01432834 | 0.2206469 | 0.0284 | 0.4666973 | 0.0019 | 0.4565841 | 0.0108 | 0.3461459 | 0.00151 | 0.3715686 | 0.0433 | 0.392386 | 0.0001 |
| #719 | TWF2 | -0.3895504 | 0.01433997 | -0.4058119 | 0.037 | -0.3398153 | 0.000207 | -0.4038818 | 0.0408 | -0.2431519 | 0.00583 | -0.4774257 | 0.00205 | -0.4672161 | 0.0000896 |
| #720 | ELOF1 | -0.5180113 | 0.01441677 | -0.2838067 | 0.0194 | -0.2314226 | 0.0456 | -1.395004 | 0.00159 | -0.3222504 | 0.0000187 | -0.4965884 | 0.0199 | -0.3789961 | 0.00000626 |
| #721 | TRIM73 | 0.5918351 | 0.01443477 | 0.3988409 | 0.00428 | 0.7327184 | 0.0162 | 0.540398 | 0.0161 | 0.450891 | 0.0134 | 0.6012894 | 0.0365 | 0.8268731 | 0.0000864 |
| #722 | OXLD1 | -0.4039898 | 0.01446168 | -0.3565138 | 0.000493 | -0.3874374 | 0.000186 | -0.5255431 | 0.0623 | -0.4365759 | 0.000000672 | -0.2937677 | 0.0129 | -0.4241011 | 0.011 |
| #723 | ANKRD36 | 0.6983165 | 0.01447538 | 0.5474972 | 0.00288 | 0.5801402 | 0.0469 | 0.8794912 | 0.000588 | 0.491484 | 0.0192 | 0.7924148 | 0.0172 | 0.8988716 | 0.0000693 |
| #724 | ZNF669 | 0.4438247 | 0.01447885 | 0.5356884 | 0.00284 | 0.287922 | 0.0697 | 0.6729041 | 0.00931 | 0.2725452 | 0.000631 | 0.5736912 | 0.000574 | 0.3201977 | 0.00377 |
| #725 | RABGGTB | 0.3906039 | 0.01451751 | 0.3960856 | 0.0000738 | 0.4517007 | 0.000856 | 0.3514113 | 0.0725 | 0.3979474 | 0.000546 | 0.3842256 | 0.0122 | 0.362253 | 0.000884 |
| #726 | R3HDM2 | 0.3705114 | 0.01453927 | 0.4861385 | 0.000128 | 0.2903793 | 0.0126 | 0.4407102 | 0.0253 | 0.2593043 | 0.03 | 0.36629 | 0.0158 | 0.3802463 | 0.00347 |
| #727 | SRP72 | 0.2942178 | 0.01454104 | 0.2526653 | 0.00202 | 0.3704434 | 0.000258 | 0.2564456 | 0.0748 | 0.3800164 | 0.000000988 | 0.3287737 | 0.00237 | 0.1769623 | 0.00781 |
| #728 | MAN1A1 | 0.6053622 | 0.01462042 | 0.8717119 | 0.00000948 | 0.7516304 | 0.0249 | 0.5512025 | 0.0337 | 0.4729637 | 0.00386 | 0.6362569 | 0.00301 | 0.3484077 | 0.0223 |
| #729 | POU2F1 | 0.4071059 | 0.01471948 | 0.3484918 | 0.000816 | 0.5366029 | 0.000906 | 0.4670556 | 0.0394 | 0.2872283 | 0.0127 | 0.2971939 | 0.0344 | 0.5060629 | 0.0000932 |
| #730 | HNRNPD | 0.2965846 | 0.01476897 | 0.2017385 | 0.0703 | 0.2510291 | 0.00632 | 0.3071186 | 0.00233 | 0.3189523 | 0.00069 | 0.3885973 | 0.000189 | 0.3120716 | 0.00878 |
| #731 | MRE11 | 0.4733733 | 0.01477045 | 0.4961544 | 0.000083 | 0.4452821 | 0.000329 | 0.2671936 | 0.0856 | 0.3873403 | 0.00261 | 0.7042757 | 0.00000681 | 0.5399938 | 0.00000311 |
| #732 | NDEL1 | 0.659282 | 0.01480112 | 0.8445216 | 0.00000319 | 0.7695685 | 0.00000543 | 0.3852143 | 0.0888 | 0.672041 | 0.00000165 | 0.7918915 | 0.00000244 | 0.4924552 | 0.00000428 |
| #733 | MBNL1 | 0.3289765 | 0.01482425 | 0.2649127 | 0.000102 | 0.2977575 | 0.000706 | 0.2784221 | 0.088 | 0.2683751 | 0.000165 | 0.4226415 | 0.0000135 | 0.4417503 | 1.57E-09 |
| #734 | BPTF | 0.4304026 | 0.01482597 | 0.4429931 | 0.00000324 | 0.4786154 | 0.000154 | 0.251453 | 0.0719 | 0.3109082 | 0.0135 | 0.5870703 | 0.00343 | 0.5113757 | 0.000000816 |
| #735 | RBM39 | 0.2570235 | 0.01483567 | 0.283399 | 0.000652 | 0.3072842 | 0.00018 | 0.2124546 | 0.0309 | 0.2835519 | 0.000000549 | 0.1893088 | 0.0568 | 0.2661423 | 0.000465 |
| #736 | AKAP11 | 0.5762032 | 0.0148436 | 0.6891292 | 1.07E-08 | 0.4725088 | 0.00303 | 0.418723 | 0.0836 | 0.5207096 | 0.000206 | 0.7883643 | 0.00216 | 0.5677845 | 0.0000182 |
| #737 | AFF4 | 0.4040307 | 0.01489206 | 0.4734255 | 0.0000241 | 0.4839728 | 0.000456 | 0.4093377 | 0.05 | 0.3957826 | 0.0000249 | 0.3216249 | 0.0371 | 0.3400409 | 0.00182 |
| #738 | TUBGCP5 | 0.5942142 | 0.01489436 | 0.5818051 | 0.00145 | 0.744642 | 0.00227 | 0.3822566 | 0.0839 | 0.6635873 | 0.000282 | 0.6460176 | 0.00141 | 0.5469765 | 0.0000812 |
| #739 | SMIM26 | -0.316875 | 0.01490099 | -0.117212 | 0.0781 | -0.3891421 | 0.006 | -0.2946956 | 0.00333 | -0.4604038 | 0.00000067 | -0.3714588 | 0.000844 | -0.2683375 | 0.00115 |
| #740 | HNRNPR | 0.299626 | 0.01495597 | 0.3115066 | 0.0000178 | 0.3263003 | 0.000128 | 0.2522158 | 0.0858 | 0.3716259 | 0.000204 | 0.3245251 | 0.00246 | 0.2115825 | 0.00115 |
| #741 | POLI | 0.5701836 | 0.01497567 | 0.4623051 | 0.0193 | 0.5717875 | 0.00243 | 0.9116323 | 0.000551 | 0.2898633 | 0.0294 | 0.5617831 | 0.0381 | 0.6237303 | 0.00000181 |
| #742 | MYO9A | 0.5690732 | 0.01500354 | 0.685405 | 0.00000172 | 0.4631428 | 0.0703 | 0.7352148 | 0.00572 | 0.4834364 | 0.0000553 | 0.4578558 | 0.014 | 0.5893842 | 0.00000177 |
| #743 | COX5B | -0.2373426 | 0.01503479 | -0.2042589 | 0.00379 | -0.2990981 | 0.0000662 | -0.1887523 | 0.0827 | -0.2556354 | 0.0021 | -0.2911902 | 0.000849 | -0.1851209 | 0.000702 |
| #744 | ICAM3 | -0.4008995 | 0.01505884 | -0.4138265 | 0.00000194 | -0.5968401 | 0.0133 | -0.2784444 | 0.0559 | -0.4519256 | 0.0184 | -0.3161622 | 0.00271 | -0.3481983 | 0.00000349 |
| #745 | TXN2 | -0.3356577 | 0.01505992 | -0.2381048 | 0.0146 | -0.3653153 | 0.0000245 | -0.3070468 | 0.0723 | -0.2775911 | 0.000771 | -0.5438492 | 0.000277 | -0.2820392 | 0.00246 |
| #746 | ATP5MC3 | -0.2732086 | 0.01507581 | -0.2651974 | 0.0102 | -0.3245205 | 0.00000258 | -0.1708859 | 0.0662 | -0.3574317 | 0.000193 | -0.3354904 | 0.0124 | -0.1857255 | 0.00143 |
| #747 | RPL36A | -0.2923949 | 0.01510844 | -0.278995 | 0.0000136 | -0.2943869 | 0.000627 | -0.2415769 | 0.0875 | -0.2747021 | 0.000287 | -0.3488937 | 0.00219 | -0.3158151 | 0.0000295 |
| #748 | RPL36 | -0.2665678 | 0.01513572 | -0.247605 | 0.0000828 | -0.2744656 | 0.0111 | -0.1943918 | 0.077 | -0.1944062 | 0.00244 | -0.3554347 | 0.000198 | -0.3331034 | 0.0000167 |
| #749 | POLB | 0.5571884 | 0.01519727 | 0.394121 | 0.0177 | 0.7592566 | 0.0291 | 0.5908811 | 0.0406 | 0.6018742 | 0.00017 | 0.3901546 | 0.00359 | 0.6068429 | 0.0000022 |
| #750 | CNPY3 | -0.287063 | 0.01523606 | -0.1332802 | 0.0389 | -0.1824707 | 0.0116 | -0.5263054 | 0.0346 | -0.2108636 | 0.00591 | -0.4033313 | 0.000143 | -0.2661264 | 0.000213 |
| #751 | ARHGDIA | -0.3241673 | 0.01524529 | -0.252333 | 0.00877 | -0.2228462 | 0.0000177 | -0.5460331 | 0.0028 | -0.1156278 | 0.0555 | -0.3151171 | 0.0244 | -0.4930469 | 0.00000167 |
| #752 | SCAF11 | 0.3411556 | 0.01524768 | 0.372603 | 0.000000949 | 0.397976 | 0.0049 | 0.2007912 | 0.0856 | 0.3512097 | 0.00055 | 0.3712571 | 0.000463 | 0.3530966 | 0.000000309 |
| #753 | RAB5C | -0.328416 | 0.01532606 | -0.2897408 | 0.000328 | -0.3485927 | 0.000237 | -0.4599458 | 0.0784 | -0.2333189 | 0.000896 | -0.3454456 | 0.00993 | -0.2934526 | 0.00218 |
| #754 | TIMM17B | -0.3608479 | 0.01534521 | -0.2846677 | 0.0108 | -0.1485676 | 0.0397 | -0.7821111 | 0.0409 | -0.2289807 | 0.000262 | -0.4074733 | 0.000175 | -0.313287 | 0.000239 |
| #755 | RELCH | 0.716721 | 0.01537613 | 0.893267 | 3.91E-08 | 0.6498664 | 0.0000416 | 0.6278819 | 0.0904 | 0.4846224 | 0.000744 | 0.8678083 | 0.00109 | 0.7768803 | 1.27E-09 |
| #756 | KLHL15 | 0.6646458 | 0.01540377 | 0.8131072 | 0.0000186 | 0.8967387 | 0.000000662 | 0.4061919 | 0.0583 | 0.690578 | 0.00000631 | 0.4316137 | 0.0338 | 0.7496455 | 0.000301 |
| #757 | RBM19 | 0.4528195 | 0.01541443 | 0.4567806 | 0.00069 | 0.2442097 | 0.0649 | 0.4266935 | 0.0242 | 0.5856038 | 3.92E-11 | 0.4991596 | 0.00264 | 0.5044699 | 0.00000211 |
| #758 | MEX3C | 0.5850163 | 0.01544171 | 0.5348829 | 0.004 | 0.9079772 | 0.0168 | 0.5676899 | 0.0509 | 0.4286447 | 0.00318 | 0.5346043 | 0.00431 | 0.5362991 | 0.0134 |
| #759 | TTC21B | 0.7374067 | 0.01545401 | 0.7342723 | 0.000000358 | 0.8018719 | 0.000411 | 0.4615384 | 0.0906 | 0.6719097 | 0.0000565 | 0.7166051 | 0.00165 | 1.038243 | 1.68E-09 |
| #760 | DICER1 | 0.468737 | 0.01546195 | 0.5140223 | 0.0000535 | 0.3933636 | 0.000218 | 0.3727091 | 0.0854 | 0.3153578 | 0.000105 | 0.5526601 | 0.00699 | 0.6643089 | 0.00000143 |
| #761 | EIF2AK2 | 0.6324265 | 0.0154663 | 0.6295758 | 0.00995 | 0.8613573 | 0.00564 | 0.4051794 | 0.0529 | 0.6823242 | 0.0131 | 0.6530246 | 0.00312 | 0.5630978 | 0.00803 |
| #762 | RPS3A | -0.2245585 | 0.01552577 | -0.2472521 | 0.00000527 | -0.1260233 | 0.034 | -0.1760526 | 0.0545 | -0.2636103 | 0.00000139 | -0.3614232 | 0.000152 | -0.1729897 | 0.00456 |
| #763 | NDUFA1 | -0.2727616 | 0.0156029 | -0.3086749 | 0.00313 | -0.2832382 | 0.000877 | -0.2240797 | 0.0783 | -0.2592769 | 0.00488 | -0.2745346 | 0.00639 | -0.2867651 | 0.0000355 |
| #764 | EP300 | 0.5329983 | 0.01567139 | 0.6588142 | 0.0000293 | 0.5645123 | 0.00000333 | 0.3206451 | 0.0913 | 0.5173499 | 0.000151 | 0.5599321 | 0.00255 | 0.5767364 | 0.0000258 |
| #765 | CMTM3 | -0.4670263 | 0.0157127 | -0.9142448 | 0.0227 | -0.2642981 | 0.000484 | -0.6300776 | 0.057 | -0.2364297 | 0.00724 | -0.4368389 | 0.00254 | -0.3202688 | 0.00432 |
| #766 | MBIP | 0.4455572 | 0.01575332 | 0.5876225 | 0.00282 | 0.3653401 | 0.0441 | 0.5647133 | 0.0386 | 0.3109017 | 0.00135 | 0.4110265 | 0.00743 | 0.4337393 | 0.000194 |
| #767 | RALGPS2 | 0.5612495 | 0.01575483 | 0.6504544 | 1.04E-11 | 0.6109987 | 0.0386 | 0.3856277 | 0.0436 | 0.4939331 | 0.00294 | 0.437083 | 0.00947 | 0.7893999 | 7.41E-08 |
| #768 | MT-ND4 | 0.3979913 | 0.01583762 | 0.2986634 | 0.00483 | 0.4872277 | 0.000164 | 0.2306841 | 0.0732 | 0.6786844 | 0.0000301 | 0.3007947 | 0.0158 | 0.3918935 | 0.001 |
| #769 | DNAJC3 | 0.3697702 | 0.01600073 | 0.292115 | 0.0362 | 0.5159811 | 0.00609 | 0.6204465 | 0.0043 | 0.3274789 | 0.00125 | 0.3402229 | 0.0101 | 0.1223765 | 0.0381 |
| #770 | TMEM39A | 0.4730044 | 0.01602726 | 0.3744745 | 0.0188 | 0.6327609 | 0.00834 | 0.4773056 | 0.0595 | 0.2762565 | 0.00507 | 0.5172619 | 0.00432 | 0.5599668 | 0.0000854 |
| #771 | SETDB1 | 0.412669 | 0.01613462 | 0.4001986 | 0.000145 | 0.3946717 | 0.000321 | 0.3090749 | 0.0946 | 0.2587064 | 0.000119 | 0.5649257 | 0.00156 | 0.5484366 | 0.000109 |
| #772 | TMA7 | -0.2889251 | 0.01619032 | -0.2565099 | 0.000153 | -0.259747 | 0.0013 | -0.1678689 | 0.0955 | -0.3468632 | 0.0000973 | -0.4045254 | 0.000052 | -0.2980364 | 0.00000379 |
| #773 | NDUFS7 | -0.3551708 | 0.01625344 | -0.3317851 | 0.00023 | -0.3220407 | 0.000149 | -0.3019512 | 0.0971 | -0.2891611 | 0.0000125 | -0.5537308 | 1.14E-08 | -0.3323557 | 8.65E-08 |
| #774 | RPS29 | -0.2944045 | 0.01625746 | -0.2605583 | 0.00307 | -0.2825932 | 0.00175 | -0.2471855 | 0.072 | -0.1949249 | 0.0192 | -0.4510187 | 0.00092 | -0.3301464 | 0.000583 |
| #775 | GATAD2B | 0.3364091 | 0.01634501 | 0.2975944 | 0.0185 | 0.3018651 | 0.00891 | 0.3617499 | 0.0611 | 0.2733451 | 0.000218 | 0.3337855 | 0.00933 | 0.4501144 | 0.0000306 |
| #776 | SMCHD1 | 0.8116219 | 0.01635142 | 0.8060408 | 0.00000143 | 0.9962663 | 1.43E-09 | 0.4488722 | 0.0978 | 0.8425414 | 0.000281 | 0.8452533 | 0.000022 | 0.9307572 | 0.000000037 |
| #777 | RBM48 | 0.6175412 | 0.01636372 | 0.745953 | 0.0000106 | 0.5823808 | 0.00897 | 0.4248745 | 0.0891 | 0.4845726 | 0.0000944 | 0.7066154 | 0.00000318 | 0.7608509 | 0.00000023 |
| #778 | PDZD8 | 0.5497407 | 0.01641077 | 0.6020548 | 0.000513 | 0.6354026 | 0.000259 | 0.314214 | 0.097 | 0.6206763 | 0.000000203 | 0.6792599 | 0.000277 | 0.4468366 | 0.00037 |
| #779 | BIRC6 | 0.6254222 | 0.01643634 | 0.7563977 | 2.2E-09 | 0.645271 | 0.000000132 | 0.295695 | 0.0986 | 0.6499649 | 1.16E-10 | 0.7368954 | 0.0000637 | 0.6683092 | 1.45E-09 |

| | | | | | | | | | | | | | | |
|---|---|---|---|---|---|---|---|---|---|---|---|---|---|---|
| #780 | RTTN | 0.5941427 | 0.01645208 | 0.5240076 | 0.00031 | 0.6387526 | 0.0258 | 0.5239534 | 0.0393 | 0.6036321 | 0.00245 | 0.4440235 | 0.0309 | 0.8304871 | 0.000000817 |
| #781 | HSPA1A | -1.143242 | 0.01648099 | -1.514944 | 0.00385 | -1.198208 | 0.0208 | -0.895351 | 0.0482 | -1.047438 | 0.0214 | -1.003177 | 0.00463 | -1.200331 | 0.0000345 |
| #782 | STXBP3 | 0.3267237 | 0.01649141 | 0.4379249 | 0.000467 | 0.2459975 | 0.0227 | 0.3222855 | 0.067 | 0.2759644 | 0.00727 | 0.3712484 | 0.0014 | 0.3069216 | 0.0000521 |
| #783 | WDR36 | 0.5958822 | 0.01656336 | 0.566393 | 0.000497 | 0.8502263 | 0.000151 | 0.3496377 | 0.0982 | 0.4869919 | 0.000214 | 0.6058748 | 0.000334 | 0.7161697 | 2.46E-08 |
| #784 | WDR44 | 0.5655089 | 0.0165758 | 0.7362391 | 0.0000417 | 0.50648 | 0.0272 | 0.6536436 | 0.0461 | 0.5339107 | 0.000178 | 0.5054592 | 0.0259 | 0.4573208 | 0.0000327 |
| #785 | ADNP | 0.4809608 | 0.01658086 | 0.5047343 | 0.000749 | 0.5560625 | 0.00479 | 0.2959243 | 0.0791 | 0.3337583 | 0.0139 | 0.6066885 | 0.000588 | 0.5885966 | 0.000396 |
| #786 | ELK3 | 0.5682561 | 0.01662226 | 0.7905808 | 0.000246 | 0.3815698 | 0.0133 | 0.5104749 | 0.0534 | 0.593681 | 0.00193 | 0.9117837 | 0.0000119 | 0.2214467 | 0.0309 |
| #787 | CLN3 | -0.4479102 | 0.01668429 | -0.412279 | 0.0216 | -0.3021326 | 0.00497 | -0.7031357 | 0.0659 | -0.2639271 | 0.00472 | -0.485899 | 0.00198 | -0.5200878 | 0.000959 |
| #788 | PPP1R2 | 0.2629382 | 0.01682674 | 0.3012646 | 0.000934 | 0.375976 | 0.0225 | 0.3924273 | 0.0408 | 0.2133244 | 0.00256 | 0.1755463 | 0.0265 | 0.1190909 | 0.00771 |
| #789 | PRDX5 | -0.2550788 | 0.01686223 | -0.1679385 | 0.0056 | -0.2789083 | 0.00132 | -0.2825859 | 0.0516 | -0.2515028 | 0.0027 | -0.3498591 | 0.0325 | -0.1996781 | 0.00746 |
| #790 | NDUFS6 | -0.2449045 | 0.01692686 | -0.2494507 | 0.00211 | -0.2740598 | 0.000548 | -0.3205229 | 0.0027 | -0.1571045 | 0.0951 | -0.2633806 | 0.00104 | -0.2049086 | 0.0000863 |
| #791 | RABGEF1 | 0.4658766 | 0.01694242 | 0.5785178 | 0.0000046 | 0.5426826 | 0.00000685 | 0.5949073 | 0.0676 | 0.3237871 | 0.0085 | 0.3703915 | 0.0254 | 0.384973 | 0.0000515 |
| #792 | UBL5 | -0.3441679 | 0.01699766 | -0.3588028 | 0.00161 | -0.2996226 | 0.00657 | -0.1999539 | 0.0912 | -0.3876304 | 0.000267 | -0.4236219 | 0.00233 | -0.3953761 | 0.0000245 |
| #793 | SLC30A9 | 0.4038593 | 0.01704225 | 0.3661967 | 0.0266 | 0.3949391 | 0.0109 | 0.3831582 | 0.0345 | 0.3371855 | 0.0000754 | 0.396306 | 0.0301 | 0.54537 | 0.000000863 |
| #794 | FBXO3 | 0.4260798 | 0.01708034 | 0.2373328 | 0.0468 | 0.5593737 | 0.00745 | 0.5042518 | 0.0452 | 0.3863217 | 0.0000424 | 0.3123824 | 0.00293 | 0.5568165 | 0.0000321 |
| #795 | LAMTOR2 | -0.3895547 | 0.01718502 | -0.4622757 | 0.004 | -0.3521399 | 0.0212 | -0.3583646 | 0.0767 | -0.2436597 | 0.0011 | -0.4834636 | 0.000101 | -0.4374245 | 0.00000252 |
| #796 | UHMK1 | 0.2723767 | 0.01720456 | 0.2391368 | 0.00374 | 0.3694351 | 0.00638 | 0.3365208 | 0.0687 | 0.2860806 | 0.00000887 | 0.2799405 | 0.000451 | 0.1231464 | 0.024 |
| #797 | ARHGAP15 | 0.3592581 | 0.01721415 | 0.4729066 | 0.00000201 | 0.3451937 | 0.0131 | 0.272402 | 0.0356 | 0.2081592 | 0.0543 | 0.4706448 | 0.0000826 | 0.3862425 | 0.000166 |
| #798 | SEC63 | 0.2778648 | 0.01723728 | 0.1244999 | 0.0615 | 0.4343867 | 0.00255 | 0.3414129 | 0.0317 | 0.1802583 | 0.00477 | 0.360005 | 0.00268 | 0.2266258 | 0.000266 |
| #799 | HTATSF1 | 0.325008 | 0.01727178 | 0.2055676 | 0.0176 | 0.3067644 | 0.000854 | 0.3005212 | 0.0818 | 0.4485892 | 0.000000096 | 0.4419428 | 0.0000795 | 0.2466631 | 0.00337 |
| #800 | RAB5A | 0.5740207 | 0.01727285 | 0.8018707 | 0.00000795 | 0.5319166 | 0.00000984 | 0.4635098 | 0.104 | 0.4584744 | 0.00000213 | 0.5945563 | 0.0000262 | 0.5937962 | 0.00000221 |
| #801 | MAGT1 | 0.3829224 | 0.01728107 | 0.2798967 | 0.00637 | 0.4993454 | 0.0236 | 0.3613316 | 0.0734 | 0.3584849 | 0.0000382 | 0.4591399 | 0.000231 | 0.3393358 | 0.0000762 |
| #802 | SRRM2 | 0.4517489 | 0.01731513 | 0.46407 | 0.000000615 | 0.5048967 | 0.00000439 | 0.2104357 | 0.101 | 0.4610496 | 0.0000274 | 0.6082173 | 0.00283 | 0.4618239 | 0.00000288 |
| #803 | SEC24B | 0.4421073 | 0.01734697 | 0.4488693 | 0.00127 | 0.3741782 | 0.0017 | 0.3598356 | 0.0973 | 0.4249148 | 0.0000371 | 0.6238687 | 0.00161 | 0.420977 | 0.00213 |
| #804 | IKBKG | -0.8587526 | 0.01739019 | -0.5453662 | 0.00556 | -0.5991154 | 0.00004 | -2.055104 | 0.0985 | -0.45872 | 0.00000163 | -0.6576727 | 0.000274 | -0.8365372 | 0.000000022 |
| #805 | IMMT | 0.3071444 | 0.01739874 | 0.19341 | 0.0611 | 0.2442983 | 0.00659 | 0.3477031 | 0.0364 | 0.2483669 | 0.000257 | 0.514633 | 0.00000149 | 0.294455 | 0.00000721 |
| #806 | BTAF1 | 0.708241 | 0.01745352 | 0.6255163 | 0.000000957 | 0.9522453 | 0.0000267 | 0.4626061 | 0.0949 | 0.7107065 | 1.2E-09 | 0.7159147 | 0.00979 | 0.7824571 | 0.0000117 |
| #807 | MORF4L2 | 0.342058 | 0.01747407 | 0.4103392 | 0.000271 | 0.461839 | 0.00627 | 0.3371208 | 0.0244 | 0.3119886 | 0.00215 | 0.2017998 | 0.0715 | 0.3292605 | 0.000219 |
| #808 | HDAC8 | 0.3165786 | 0.01752368 | 0.3199912 | 0.0023 | 0.2332416 | 0.0334 | 0.3977826 | 0.0397 | 0.1965135 | 0.0165 | 0.3628679 | 0.0127 | 0.3890746 | 0.00048 |
| #809 | IPO7 | 0.5302107 | 0.01752832 | 0.657625 | 0.0000112 | 0.4925664 | 0.000087 | 0.3121577 | 0.105 | 0.4650104 | 0.0000736 | 0.717198 | 0.000169 | 0.5367069 | 0.00000415 |
| #810 | TRMT11 | 0.6248656 | 0.01757118 | 0.7861278 | 0.0000032 | 0.5761341 | 0.0116 | 0.4686362 | 0.0852 | 0.7165354 | 0.000117 | 0.5427011 | 0.00851 | 0.6590591 | 0.000000386 |
| #811 | ZNF678 | 0.5743524 | 0.0175862 | 0.3942534 | 0.0562 | 0.4949132 | 0.0238 | 0.7191383 | 0.00905 | 0.5143401 | 0.0125 | 0.5463745 | 0.00365 | 0.777095 | 0.000289 |
| #812 | ATP6V0E1 | -0.4003919 | 0.01760984 | -0.4060743 | 0.0241 | -0.2820861 | 0.0287 | -0.4395891 | 0.0458 | -0.2908784 | 0.00123 | -0.4841046 | 0.00577 | -0.4996188 | 0.0000159 |
| #813 | SP3 | 0.5421208 | 0.01763694 | 0.6238875 | 0.00000696 | 0.5646397 | 0.000059 | 0.3079479 | 0.106 | 0.5182506 | 0.0000288 | 0.5762255 | 0.000125 | 0.6617739 | 0.000000186 |
| #814 | FXR1 | 0.2292357 | 0.01765406 | 0.2519521 | 0.000943 | 0.2394525 | 0.000793 | 0.2499351 | 0.0281 | 0.1093637 | 0.0675 | 0.2418501 | 0.00853 | 0.2828604 | 0.0000017 |
| #815 | SEPTIN7 | 0.2212613 | 0.01767152 | 0.218418 | 0.000139 | 0.2595826 | 0.0000274 | 0.1824665 | 0.0826 | 0.2207425 | 0.0000209 | 0.2085337 | 0.0227 | 0.2378244 | 0.000572 |
| #816 | PSMA7 | -0.2696196 | 0.01768218 | -0.2793336 | 0.00424 | -0.2427264 | 0.000962 | -0.245219 | 0.0337 | -0.193523 | 0.0572 | -0.3219324 | 0.00998 | -0.3349833 | 0.00000707 |
| #817 | AKAP9 | 0.3989015 | 0.01769886 | 0.3227152 | 0.000139 | 0.5266408 | 0.00179 | 0.4999567 | 0.00533 | 0.3836106 | 0.0000664 | 0.3539709 | 0.094 | 0.3065147 | 0.00488 |
| #818 | MGA | 0.5057196 | 0.01773428 | 0.4870089 | 0.000539 | 0.4957874 | 0.00607 | 0.5074278 | 0.0624 | 0.4620696 | 0.00658 | 0.5818766 | 0.0307 | 0.5001475 | 0.0000403 |
| #819 | G6PD | -0.4250436 | 0.01779877 | -0.4587433 | 0.00547 | -0.4062867 | 0.00441 | -0.7168782 | 0.0169 | -0.2748534 | 0.00856 | -0.2454108 | 0.0714 | -0.4480891 | 0.0000731 |
| #820 | MKLN1 | 0.445967 | 0.01782184 | 0.5878571 | 0.00000655 | 0.4655072 | 0.0000463 | 0.3345969 | 0.106 | 0.3587053 | 0.0000501 | 0.4821742 | 0.000531 | 0.4469612 | 0.0000162 |
| #821 | SCAF4 | 0.3675819 | 0.01790975 | 0.3893468 | 0.000969 | 0.4414466 | 0.00341 | 0.2580246 | 0.0819 | 0.2967712 | 0.00696 | 0.3753656 | 0.014 | 0.4445367 | 0.000195 |
| #822 | ZNF844 | 0.5987259 | 0.0179359 | 0.5941688 | 0.0000175 | 0.3437138 | 0.0414 | 0.512595 | 0.05 | 0.5520217 | 0.00186 | 0.6559856 | 0.0144 | 0.9338704 | 0.00000795 |
| #823 | SHKBP1 | -0.4339374 | 0.01797082 | -0.3689652 | 0.000254 | -0.4026528 | 0.00538 | -0.6046725 | 0.0986 | -0.2842486 | 0.000339 | -0.4329846 | 0.00321 | -0.5101005 | 0.0000238 |
| #824 | MAP2K4 | 0.4802623 | 0.0180113 | 0.5247059 | 0.00062 | 0.3787366 | 0.0301 | 0.4653861 | 0.0633 | 0.3137078 | 0.0107 | 0.4705844 | 0.00329 | 0.7284527 | 0.000000675 |
| #825 | KANSL2 | 0.5594992 | 0.01803435 | 0.5314899 | 0.00018 | 0.6965038 | 0.00515 | 0.3925136 | 0.102 | 0.434824 | 0.000305 | 0.5606088 | 0.000323 | 0.7410552 | 0.00000915 |
| #826 | ARFGEF2 | 0.6985472 | 0.01803645 | 0.7578966 | 0.000000721 | 0.8134446 | 0.000721 | 0.3291984 | 0.102 | 0.9328936 | 8.64E-10 | 0.5571691 | 0.00574 | 0.8006811 | 2.04E-10 |
| #827 | SMARCE1 | 0.2333505 | 0.01804983 | 0.3468849 | 0.000144 | 0.1901735 | 0.077 | 0.2924214 | 0.0023 | 0.1831998 | 0.0000169 | 0.1458779 | 0.0288 | 0.2415455 | 0.0000426 |
| #828 | PDZD11 | -0.6320577 | 0.01815276 | -0.5225504 | 0.0756 | -0.5230696 | 0.00077 | -1.151353 | 0.0112 | -0.4719526 | 0.00294 | -0.5395396 | 0.0183 | -0.5838814 | 0.0000193 |
| #829 | ARPP19 | 0.3273593 | 0.01819342 | 0.4556206 | 0.000436 | 0.4082124 | 0.000949 | 0.2392742 | 0.107 | 0.2621269 | 0.0000536 | 0.3360195 | 0.00106 | 0.2629024 | 0.0000314 |
| #830 | NFKBIZ | 0.4862441 | 0.01819486 | 0.5032335 | 0.000995 | 0.4279813 | 0.00152 | 0.3849143 | 0.0676 | 0.2714576 | 0.0379 | 0.7347582 | 0.000216 | 0.5951197 | 0.000956 |
| #831 | PSMF1 | -0.320787 | 0.01823534 | -0.305341 | 0.00532 | -0.3474788 | 0.0000486 | -0.2958079 | 0.0992 | -0.2100687 | 0.0048 | -0.3529742 | 0.0000187 | -0.4130512 | 3.06E-08 |
| #832 | GNPDA2 | 0.4315124 | 0.01824561 | 0.4141508 | 0.00334 | 0.5334795 | 0.0229 | 0.5011859 | 0.0455 | 0.4617675 | 0.0000651 | 0.4673197 | 0.00038 | 0.2111712 | 0.0372 |
| #833 | CHD6 | 0.3525836 | 0.01830248 | 0.4611124 | 0.0000754 | 0.239447 | 0.00187 | 0.4037138 | 0.0338 | 0.420031 | 0.00212 | 0.2128916 | 0.0716 | 0.3783058 | 0.000359 |
| #834 | RPS23 | -0.1773974 | 0.01835549 | -0.1664011 | 0.00000944 | -0.1464055 | 0.000438 | -0.1258586 | 0.0859 | -0.204901 | 0.0000501 | -0.3235574 | 0.00182 | -0.09726095 | 0.022 |
| #835 | CALCOCO2 | 0.4963043 | 0.01842989 | 0.7065895 | 0.00000317 | 0.3862246 | 0.0000819 | 0.2216404 | 0.103 | 0.4678869 | 0.00649 | 0.6315113 | 0.000766 | 0.5639729 | 0.0000211 |
| #836 | STK24 | 0.4021428 | 0.01845228 | 0.4691563 | 0.0035 | 0.2969625 | 0.00717 | 0.7321323 | 0.0119 | 0.2541063 | 0.00417 | 0.4192698 | 0.00774 | 0.2412294 | 0.0762 |
| #837 | JMY | 0.4936399 | 0.01851057 | 0.3606135 | 0.0248 | 0.6628137 | 0.000417 | 0.3579288 | 0.0837 | 0.5394761 | 0.000712 | 0.5046623 | 0.000809 | 0.536345 | 0.000597 |
| #838 | OSBPL8 | 0.4034407 | 0.01851898 | 0.4446302 | 0.000153 | 0.3779368 | 0.00011 | 0.5142215 | 0.0266 | 0.1720275 | 0.0844 | 0.4813514 | 0.00000226 | 0.4304768 | 7.8E-09 |
| #839 | IPO5 | 0.410882 | 0.01856339 | 0.1911337 | 0.0886 | 0.5785049 | 0.0000125 | 0.5972556 | 0.00639 | 0.4751472 | 0.000013 | 0.2834391 | 0.0162 | 0.3398115 | 0.000191 |
| #840 | DNAJC13 | 0.586467 | 0.01856846 | 0.6330229 | 0.00000804 | 0.6794947 | 0.00000638 | 0.3606579 | 0.111 | 0.7168549 | 0.000176 | 0.6156883 | 0.000362 | 0.5130834 | 0.0000114 |
| #841 | MGAT1 | -0.4450697 | 0.01857491 | -0.3307686 | 0.046 | -0.2521589 | 0.00105 | -0.9390903 | 0.0355 | -0.214734 | 0.0123 | -0.5251988 | 0.00412 | -0.4084677 | 0.0125 |
| #842 | DDHD2 | 0.414778 | 0.01862347 | 0.1905362 | 0.0726 | 0.4786676 | 0.00625 | 0.6237189 | 0.0184 | 0.4795497 | 0.000337 | 0.4574411 | 0.0018 | 0.2587543 | 0.0123 |
| #843 | HIST1H3D | 0.7505399 | 0.01862914 | 0.8052351 | 0.000414 | 1.128304 | 0.00021 | 0.3659127 | 0.0774 | 1.122755 | 0.000000317 | 0.6593664 | 0.00138 | 0.4216667 | 0.0324 |
| #844 | TMX3 | 0.3792096 | 0.01864373 | 0.246454 | 0.017 | 0.4809349 | 0.00976 | 0.2968887 | 0.0842 | 0.3842995 | 0.000129 | 0.3614767 | 0.000756 | 0.5052037 | 0.000000205 |
| #845 | CDKAL1 | 0.5570963 | 0.01866244 | 0.4893037 | 0.014 | 0.5253656 | 0.00488 | 0.3730074 | 0.0929 | 0.6288849 | 0.00000103 | 0.6938526 | 0.000215 | 0.6321634 | 7.33E-09 |
| #846 | CGAS | 0.7703253 | 0.01875214 | 0.5333946 | 0.000689 | 1.139762 | 0.00355 | 0.7660424 | 0.0864 | 0.9075994 | 0.000125 | 0.9474685 | 0.0000656 | 0.3276846 | 0.0217 |
| #847 | PKN2 | 0.6891317 | 0.01880458 | 0.8038636 | 1.64E-09 | 0.7129497 | 4.95E-08 | 0.3197182 | 0.112 | 0.5757123 | 0.00000357 | 0.7751617 | 0.000456 | 0.9473848 | 5.37E-10 |
| #848 | RO60 | 0.3959819 | 0.01883095 | 0.447766 | 0.00000124 | 0.5202954 | 0.000000436 | 0.1762024 | 0.113 | 0.5266831 | 4.79E-10 | 0.3890895 | 0.000142 | 0.3158552 | 0.00000337 |
| #849 | SNHG6 | -0.2357473 | 0.01883212 | -0.2187763 | 0.00244 | -0.1565822 | 0.0165 | -0.1839494 | 0.0917 | -0.1928198 | 0.00229 | -0.4052647 | 0.0000563 | -0.2570911 | 0.00000334 |
| #850 | ZNF800 | 0.5412332 | 0.0188719 | 0.5658668 | 0.000148 | 0.7236919 | 0.000672 | 0.3421398 | 0.112 | 0.4558021 | 0.000000772 | 0.5820987 | 0.000181 | 0.5778002 | 0.000000373 |

| | | | | | | | | | | | | | | | |
|---|---|---|---|---|---|---|---|---|---|---|---|---|---|---|---|
| #851 | MRPL51 | -0.4240504 | 0.01889671 | -0.5333841 | 0.00621 | -0.3531977 | 0.00000272 | -0.3677132 | 0.102 | -0.3283584 | 0.00297 | -0.5676161 | 0.000479 | -0.3940331 | 0.00193 |
| #852 | RALGAPB | 0.5242157 | 0.01891058 | 0.5517249 | 0.0000608 | 0.5895836 | 0.0000403 | 0.5748118 | 0.0607 | 0.5999803 | 0.00000173 | 0.3703775 | 0.0526 | 0.4588158 | 0.0000851 |
| #853 | CEP57 | 0.3514502 | 0.01892701 | 0.3803875 | 0.0000298 | 0.3967041 | 0.0004 | 0.4526276 | 0.0134 | 0.1732538 | 0.0936 | 0.4120149 | 0.00571 | 0.2937132 | 0.000459 |
| #854 | LARS | 0.3928137 | 0.01897582 | 0.4136666 | 0.00000686 | 0.4064723 | 0.00268 | 0.1766669 | 0.11 | 0.3382408 | 0.00092 | 0.5020609 | 0.000132 | 0.5197749 | 0.0000349 |
| #855 | CCDC107 | -0.4078041 | 0.018977 | -0.3893326 | 0.0104 | -0.2284973 | 0.0177 | -0.7409285 | 0.0519 | -0.3754482 | 0.00000365 | -0.208429 | 0.0338 | -0.5041888 | 0.00000412 |
| #856 | ACO1 | 0.5036989 | 0.01898639 | 0.7082345 | 0.00127 | 0.6420569 | 0.00219 | 0.5077808 | 0.0725 | 0.2613863 | 0.00402 | 0.35359 | 0.0335 | 0.5491446 | 0.000403 |
| #857 | XPO1 | 0.5143466 | 0.01911043 | 0.6079504 | 0.00000298 | 0.5338542 | 0.000184 | 0.3383804 | 0.113 | 0.4850771 | 0.000601 | 0.5621289 | 0.00125 | 0.5586889 | 0.0000263 |
| #858 | SLC25A28 | 0.5676469 | 0.01911203 | 0.59183 | 0.00326 | 0.6679792 | 0.00469 | 0.8400797 | 0.00227 | 0.3835268 | 0.0619 | 0.3948365 | 0.0421 | 0.5276294 | 0.000475 |
| #859 | USF3 | 0.6765756 | 0.0191184 | 0.8018648 | 0.000000483 | 0.8663989 | 0.000149 | 0.3070071 | 0.115 | 0.6139834 | 3.59E-10 | 0.7759905 | 5.26E-08 | 0.6942091 | 0.00000444 |
| #860 | SMIM12 | -0.433184 | 0.01911863 | -0.2741803 | 0.0571 | -0.6610456 | 0.0000636 | -0.6077851 | 0.047 | -0.3363959 | 0.0000322 | -0.3793387 | 0.0106 | -0.3403582 | 0.00000482 |
| #861 | HNRNPAB | 0.3792809 | 0.01913538 | 0.4377705 | 0.0102 | 0.4769533 | 0.000432 | 0.3579419 | 0.0259 | 0.325991 | 0.0179 | 0.3248081 | 0.0363 | 0.3522207 | 0.0241 |
| #862 | FTH1 | -0.3163396 | 0.019195 | -0.2641605 | 0.00251 | -0.2793007 | 0.0375 | -0.1922427 | 0.0635 | -0.3816674 | 0.00329 | -0.4617339 | 0.00578 | -0.3189328 | 0.00255 |
| #863 | RPL8 | -0.310379 | 0.01919814 | -0.3623491 | 0.000874 | -0.3827245 | 0.0126 | -0.1531365 | 0.0948 | -0.2852134 | 0.00264 | -0.3793535 | 0.00212 | -0.2994967 | 0.00215 |
| #864 | RC3H1 | 0.3305631 | 0.01920375 | 0.3865204 | 0.000029 | 0.2147406 | 0.0701 | 0.3064548 | 0.0387 | 0.3756145 | 0.000206 | 0.299513 | 0.00613 | 0.4005352 | 0.0000277 |
| #865 | LUC7L | 0.4491012 | 0.01921266 | 0.5858259 | 0.000000346 | 0.5269196 | 0.0000819 | 0.2628013 | 0.106 | 0.3634597 | 0.000417 | 0.2939938 | 0.0083 | 0.6616066 | 0.000000297 |
| #866 | AURKAIP1 | -0.2593482 | 0.01923925 | -0.1520555 | 0.0389 | -0.1970911 | 0.000772 | -0.4213144 | 0.0357 | -0.1777337 | 0.0372 | -0.3405265 | 0.00182 | -0.2673681 | 0.00105 |
| #867 | TMF1 | 0.3354036 | 0.01924189 | 0.4654133 | 0.00000907 | 0.3731226 | 0.00143 | 0.2791917 | 0.105 | 0.2451118 | 0.00315 | 0.42354 | 0.000194 | 0.2260423 | 0.00555 |
| #868 | ZMYM2 | 0.5147073 | 0.01925899 | 0.5552604 | 0.0000198 | 0.7194012 | 0.000779 | 0.2813204 | 0.114 | 0.5626735 | 0.0000454 | 0.5229219 | 0.000107 | 0.4466666 | 0.000289 |
| #869 | TRIM25 | 0.5147051 | 0.0192794 | 0.5307989 | 0.00454 | 0.5303048 | 0.00474 | 0.7733267 | 0.0279 | 0.3595645 | 0.0712 | 0.4006662 | 0.00472 | 0.4935695 | 0.0026 |
| #870 | SPAST | 0.4418439 | 0.01930099 | 0.547721 | 0.0000143 | 0.3970204 | 0.012 | 0.2283196 | 0.102 | 0.4731784 | 0.0000329 | 0.4940692 | 0.00138 | 0.5107548 | 0.00000159 |
| #871 | UBE2Q2 | 0.4684778 | 0.01934038 | 0.6193291 | 0.00131 | 0.3992496 | 0.000814 | 0.3628418 | 0.109 | 0.4742674 | 0.0011 | 0.5962117 | 0.0000421 | 0.3589674 | 0.00343 |
| #872 | IMP3 | -0.3202482 | 0.01941536 | -0.2477805 | 0.0297 | -0.3419867 | 0.00135 | -0.4117919 | 0.0701 | -0.3209304 | 0.0000504 | -0.3782134 | 0.000915 | -0.2207863 | 0.0143 |
| #873 | CHD9 | 0.3997266 | 0.01946447 | 0.3580294 | 0.0000423 | 0.4340325 | 0.0647 | 0.2634331 | 0.0502 | 0.4284372 | 0.00000766 | 0.4687577 | 0.00163 | 0.4456697 | 0.000188 |
| #874 | MRPL57 | -0.257184 | 0.01948533 | -0.2859879 | 0.0224 | -0.2299276 | 0.0319 | -0.2639896 | 0.0605 | -0.2878089 | 0.000000357 | -0.303074 | 0.000988 | -0.172316 | 0.00109 |
| #875 | ZRANB2 | 0.3248642 | 0.01948935 | 0.244929 | 0.0301 | 0.4356283 | 0.0317 | 0.2874265 | 0.00573 | 0.2251309 | 0.0313 | 0.3503828 | 0.0176 | 0.4056881 | 0.000473 |
| #876 | ACTG1 | -0.4746664 | 0.01955449 | -0.5281484 | 0.000314 | -0.5870662 | 0.000386 | -0.3547367 | 0.0579 | -0.5035486 | 0.0000343 | -0.3988476 | 0.0584 | -0.4756512 | 0.000282 |
| #877 | ORMDL2 | -0.5889419 | 0.01955536 | -0.3414946 | 0.0744 | -0.6466425 | 0.0111 | -0.6638499 | 0.0303 | -0.5194757 | 0.00000665 | -0.5772858 | 0.00146 | -0.7849031 | 0.0000909 |
| #878 | ZNF638 | 0.4315743 | 0.01961864 | 0.4934232 | 0.0000162 | 0.419148 | 0.00808 | 0.3188326 | 0.0493 | 0.3375813 | 0.0163 | 0.4068602 | 0.0439 | 0.6136004 | 0.00000739 |
| #879 | WARS2 | 0.5858456 | 0.01964594 | 0.5979608 | 0.0429 | 0.4207962 | 0.00832 | 0.4726385 | 0.0598 | 0.3025112 | 0.00353 | 0.6884475 | 0.00331 | 1.03272 | 5.84E-09 |
| #880 | NEDD8 | -0.2485864 | 0.01964932 | -0.1922459 | 0.00114 | -0.15394 | 0.0408 | -0.2061467 | 0.0754 | -0.2707988 | 0.00000165 | -0.3710783 | 0.000529 | -0.2973091 | 0.00000011 |
| #881 | EPB41 | 0.4885059 | 0.01965644 | 0.49982 | 0.000000777 | 0.6139277 | 1.39E-10 | 0.2643428 | 0.109 | 0.4181542 | 0.0000115 | 0.4226757 | 0.00885 | 0.7121149 | 0.000000268 |
| #882 | TNKS | 0.7003517 | 0.01966542 | 0.7944952 | 0.000000475 | 0.9183833 | 0.00000104 | 0.3390232 | 0.116 | 0.5532379 | 0.000132 | 0.7152874 | 0.00147 | 0.8816831 | 2.26E-10 |
| #883 | OSBP | 0.3790019 | 0.01968673 | 0.5200453 | 0.000291 | 0.3785657 | 0.00115 | 0.39861 | 0.0706 | 0.3614467 | 0.000136 | 0.2912276 | 0.0283 | 0.3241164 | 0.0176 |
| #884 | IFIH1 | 0.6586746 | 0.01970767 | 0.7104936 | 0.000831 | 0.4776195 | 0.0336 | 0.7217193 | 0.0253 | 0.6374864 | 0.0382 | 0.7957587 | 0.00114 | 0.6089699 | 0.0192 |
| #885 | RAB11FIP2 | 0.572734 | 0.01971307 | 0.6053122 | 0.000768 | 0.644471 | 0.0000412 | 0.275651 | 0.117 | 0.3818724 | 0.0000763 | 0.6844891 | 0.000169 | 0.844608 | 0.0000032 |
| #886 | YIPF3 | -0.3937473 | 0.01973375 | -0.3052895 | 0.0203 | -0.4054864 | 0.0000394 | -0.285442 | 0.0979 | -0.3936691 | 0.000000382 | -0.4815737 | 0.000134 | -0.4910232 | 0.0000144 |
| #887 | ABHD13 | 0.3333186 | 0.01973463 | 0.3475405 | 0.00577 | 0.4321232 | 0.000157 | 0.4810706 | 0.0393 | 0.2570344 | 0.000566 | 0.2500912 | 0.0497 | 0.2320515 | 0.0229 |
| #888 | CCNL2 | 0.4629163 | 0.01981619 | 0.4183586 | 0.0000227 | 0.461712 | 0.000123 | 0.2630614 | 0.109 | 0.4072018 | 0.00203 | 0.5130906 | 0.00792 | 0.7140735 | 0.000000757 |
| #889 | PCGF5 | 0.4299957 | 0.01992091 | 0.5139901 | 0.000207 | 0.1088658 | 0.0971 | 0.6998495 | 0.0021 | 0.3387225 | 0.00356 | 0.4460626 | 0.0158 | 0.4724837 | 0.000813 |
| #890 | TUFM | -0.2627911 | 0.02006089 | -0.2010542 | 0.0878 | -0.2827996 | 0.00222 | -0.2852131 | 0.0215 | -0.2104205 | 0.00186 | -0.3750266 | 0.00215 | -0.2222324 | 0.00484 |
| #891 | TBC1D10C | -0.5806246 | 0.0201173 | -0.4660167 | 0.000265 | -0.5562398 | 0.0000506 | -0.770961 | 0.112 | -0.7414833 | 0.00843 | -0.4982471 | 0.0000104 | -0.4507999 | 0.0000238 |
| #892 | WASL | 0.3895856 | 0.02015253 | 0.2680233 | 0.0451 | 0.354567 | 0.0183 | 0.4491279 | 0.0413 | 0.3443771 | 0.00112 | 0.3941449 | 0.015 | 0.5272735 | 0.000046 |
| #893 | PNRC2 | 0.2611209 | 0.020247 | 0.2330636 | 0.00271 | 0.2743263 | 0.00313 | 0.3433431 | 0.0141 | 0.1109561 | 0.101 | 0.305536 | 0.000433 | 0.2995 | 0.000357 |
| #894 | ZNF26 | 0.4098741 | 0.02026449 | 0.3529721 | 0.0107 | 0.3129914 | 0.0639 | 0.6504263 | 0.0201 | 0.3007518 | 0.0003 | 0.4050484 | 0.0263 | 0.4370545 | 0.000327 |
| #895 | ABTB1 | -0.5133493 | 0.02026977 | -0.3869532 | 0.00256 | -0.4358433 | 0.0355 | -0.6509583 | 0.0753 | -0.6114101 | 0.00659 | -0.560172 | 0.00125 | -0.4347589 | 0.000445 |
| #896 | MARF1 | 0.465696 | 0.02033857 | 0.6027941 | 0.00000127 | 0.3522771 | 0.00388 | 0.5017348 | 0.0638 | 0.3332696 | 0.0184 | 0.405445 | 0.036 | 0.5986553 | 0.0000122 |
| #897 | PDS5B | 0.4322945 | 0.02038367 | 0.4930921 | 0.000126 | 0.4622679 | 0.0000216 | 0.2614619 | 0.118 | 0.4715685 | 0.000000254 | 0.4505123 | 0.0038 | 0.4548645 | 0.000000956 |
| #898 | PRDX3 | -0.5652792 | 0.02046415 | -0.7899319 | 0.0285 | -0.5118842 | 0.00000993 | -0.4666909 | 0.0216 | -0.3601354 | 0.00496 | -0.5634627 | 0.0664 | -0.6995702 | 0.0013 |
| #899 | FBXO28 | 0.4840468 | 0.02051695 | 0.2151617 | 0.104 | 0.5831245 | 0.00000408 | 0.6117066 | 0.0181 | 0.3568167 | 0.000375 | 0.5351755 | 0.000215 | 0.6022959 | 0.0000206 |
| #900 | STXBP5 | 0.6473202 | 0.02069459 | 0.3715518 | 0.0596 | 0.8833694 | 0.0000594 | 0.9449671 | 0.00419 | 0.2917392 | 0.0576 | 0.6613091 | 0.00272 | 0.7309846 | 0.00000752 |
| #901 | COPS6 | -0.2564829 | 0.02069572 | -0.2151474 | 0.0197 | -0.3044661 | 0.00172 | -0.3462697 | 0.08 | -0.2585507 | 0.00000161 | -0.1718376 | 0.0219 | -0.2426256 | 0.000932 |
| #902 | PPP2R2A | 0.3492149 | 0.02072266 | 0.4229776 | 0.0000135 | 0.3539478 | 0.0121 | 0.3177813 | 0.112 | 0.378989 | 0.0000736 | 0.3230932 | 0.000254 | 0.2985009 | 0.000139 |
| #903 | EIF4B | 0.3308361 | 0.02074702 | 0.2579799 | 0.00000657 | 0.3767067 | 0.0000234 | 0.5147859 | 0.000105 | 0.1641222 | 0.124 | 0.3470857 | 0.000052 | 0.3243362 | 3.05E-08 |
| #904 | POLDIP2 | -0.3043078 | 0.02078893 | -0.2466457 | 0.0156 | -0.2583663 | 0.00576 | -0.6078394 | 0.0173 | -0.197702 | 0.000367 | -0.3034767 | 0.0796 | -0.2118165 | 0.00609 |
| #905 | NUMB | 0.4442076 | 0.02078958 | 0.4328578 | 0.0000223 | 0.3897527 | 0.00237 | 0.4444026 | 0.121 | 0.2760471 | 0.000445 | 0.5940966 | 0.00125 | 0.5280887 | 0.00000936 |
| #906 | RNF111 | 0.5316998 | 0.02079619 | 0.7445817 | 0.000000385 | 0.2924696 | 0.0697 | 0.6509061 | 0.0124 | 0.4040518 | 0.00306 | 0.4081212 | 0.0396 | 0.6900682 | 0.000000161 |
| #907 | VIM | -0.4219123 | 0.02082047 | -0.384617 | 0.000779 | -0.3813858 | 0.0289 | -0.3900229 | 0.0532 | -0.2271592 | 0.042 | -0.5697379 | 0.0000365 | -0.578551 | 0.000000116 |
| #908 | COX7A2L | -0.3238757 | 0.02087045 | -0.2566306 | 0.00256 | -0.2444215 | 0.0106 | -0.332961 | 0.11 | -0.4192278 | 0.0000424 | -0.4695359 | 0.00152 | -0.2204775 | 0.000821 |
| #909 | PRRC2B | 0.4988918 | 0.02088674 | 0.4767507 | 0.0000959 | 0.5546932 | 0.00000151 | 0.4288922 | 0.122 | 0.3474602 | 0.000529 | 0.5486311 | 0.00254 | 0.6369237 | 0.00000132 |
| #910 | OGFRL1 | 0.6166879 | 0.02096342 | 0.7908615 | 0.000233 | 0.4832712 | 0.0233 | 0.5751403 | 0.0955 | 0.5054213 | 0.00163 | 0.8364131 | 0.000182 | 0.5090196 | 0.00492 |
| #911 | SENP6 | 0.4755775 | 0.0211924 | 0.5978588 | 0.000004 | 0.5436865 | 0.0000783 | 0.2682145 | 0.127 | 0.3635411 | 0.0000913 | 0.6100736 | 0.0000149 | 0.4700904 | 0.0000308 |
| #912 | MIA3 | 0.623377 | 0.02122598 | 0.6736087 | 0.00000969 | 0.9024171 | 0.00453 | 0.3181894 | 0.123 | 0.5631947 | 7.27E-09 | 0.5624947 | 0.000258 | 0.7203575 | 5.75E-09 |
| #913 | ZNF518A | 0.3778417 | 0.02125283 | 0.388839 | 0.000563 | 0.3732371 | 0.00198 | 0.5079551 | 0.0622 | 0.2054291 | 0.0415 | 0.3361393 | 0.0212 | 0.4554504 | 0.0000378 |
| #914 | YTHDF3 | 0.6578182 | 0.02130448 | 0.681791 | 0.0000865 | 0.7352293 | 0.000000641 | 0.3985794 | 0.128 | 0.5502779 | 0.000000211 | 0.9150507 | 0.00000255 | 0.6659808 | 0.000000607 |
| #915 | DYM | 0.3529946 | 0.02132482 | 0.3381862 | 0.014 | 0.1695733 | 0.073 | 0.4303069 | 0.0356 | 0.3334022 | 0.000712 | 0.4488886 | 0.00467 | 0.3976101 | 0.0000166 |
| #916 | ZNF654 | 0.4095819 | 0.02133621 | 0.3542883 | 0.0362 | 0.4225269 | 0.00147 | 0.2934866 | 0.0795 | 0.4527297 | 0.0000235 | 0.4360336 | 0.0108 | 0.4984259 | 0.0000115 |
| #917 | SGF29 | -0.4993766 | 0.0213947 | -0.5457485 | 0.000658 | -0.3569914 | 0.000488 | -0.5179507 | 0.127 | -0.5027189 | 0.0000962 | -0.6728612 | 0.000393 | -0.3999887 | 8.25E-08 |
| #918 | OST4 | -0.3198371 | 0.0214663 | -0.2786941 | 0.0000835 | -0.1771914 | 0.127 | -0.3262502 | 0.00141 | -0.3339156 | 0.000025 | -0.4943007 | 0.0000685 | -0.3086703 | 6.78E-08 |
| #919 | OSBPL3 | 0.6105659 | 0.02146666 | 0.8400387 | 0.0000597 | 0.1953446 | 0.099 | 0.7525487 | 0.0201 | 0.5234797 | 0.00648 | 0.936865 | 0.000584 | 0.4151187 | 0.00262 |
| #920 | SPATC1L | -1.103229 | 0.02148302 | -1.091196 | 0.0000398 | -0.6692665 | 0.000791 | -0.3484669 | 0.128 | -1.078954 | 0.000416 | -1.401807 | 0.0000711 | -2.029681 | 0.0000191 |
| #921 | DDX6 | 0.3377999 | 0.02155166 | 0.2854695 | 0.000962 | 0.493716 | 0.000486 | 0.2022177 | 0.12 | 0.3899051 | 0.000778 | 0.2706371 | 0.00718 | 0.3848539 | 0.000152 |

| | | | | | | | | | | | | | | | |
|---|---|---|---|---|---|---|---|---|---|---|---|---|---|---|---|
| #922 | UROD | -0.3253579 | 0.0215744 | -0.4039876 | 0.00536 | -0.2604224 | 0.00252 | -0.5831547 | 0.00313 | -0.2450929 | 0.000548 | -0.2432159 | 0.118 | -0.2162739 | 0.000325 |
| #923 | MAGOH | -0.2531236 | 0.02159875 | -0.21894 | 0.0507 | -0.2875868 | 0.00221 | -0.3400961 | 0.0115 | -0.297126 | 0.000193 | -0.2696611 | 0.0186 | -0.1053314 | 0.0463 |
| #924 | EML4 | 0.5485719 | 0.02161213 | 0.6003148 | 9.51E-08 | 0.6789309 | 0.00603 | 0.5009229 | 0.118 | 0.6651127 | 0.00000116 | 0.4705482 | 0.00472 | 0.3756017 | 0.000546 |
| #925 | PDCD6 | -0.2696248 | 0.0216305 | -0.1935666 | 0.0412 | -0.3299575 | 0.0000885 | -0.2659368 | 0.0536 | -0.2660562 | 0.000726 | -0.356325 | 0.00994 | -0.2059068 | 0.0242 |
| #926 | MAP3K4 | 0.5774618 | 0.02171465 | 0.6246613 | 0.0000638 | 0.6642948 | 0.0000709 | 0.2762272 | 0.111 | 0.4849675 | 0.00128 | 0.5647762 | 0.0176 | 0.8498438 | 6.27E-09 |
| #927 | LTN1 | 0.4052233 | 0.02172323 | 0.3115519 | 0.00162 | 0.2961511 | 0.0329 | 0.3803145 | 0.0929 | 0.4053688 | 0.000891 | 0.5871266 | 0.00201 | 0.4508268 | 0.0000704 |
| #928 | PPP3CA | 0.3768822 | 0.02179078 | 0.4984217 | 0.00000226 | 0.3392381 | 0.00000223 | 0.2790589 | 0.102 | 0.2499646 | 0.0205 | 0.5899839 | 0.000176 | 0.304626 | 0.00818 |
| #929 | CSNK1G1 | 0.3952298 | 0.02195015 | 0.4225733 | 0.00208 | 0.4642802 | 0.00656 | 0.3661806 | 0.1 | 0.350133 | 0.000802 | 0.3227862 | 0.0218 | 0.4454255 | 0.0000146 |
| #930 | SYNCRIP | 0.3630489 | 0.02197176 | 0.4038657 | 0.0000128 | 0.4265078 | 0.000000393 | 0.184357 | 0.132 | 0.3667076 | 0.00000376 | 0.4095104 | 0.000000889 | 0.3873446 | 0.000000844 |
| #931 | TARS | 0.4187549 | 0.0219786 | 0.5208307 | 0.00159 | 0.3064745 | 0.00682 | 0.2443237 | 0.117 | 0.4262314 | 0.00534 | 0.541924 | 0.000213 | 0.4727449 | 0.000649 |
| #932 | APRT | -0.2676727 | 0.0219791 | -0.3061551 | 0.000527 | -0.1964527 | 0.0127 | -0.281026 | 0.0434 | -0.2184546 | 0.00871 | -0.3150286 | 0.0643 | -0.2889192 | 0.00231 |
| #933 | RPL35A | -0.2102156 | 0.02199906 | -0.2226418 | 0.00000317 | -0.1665815 | 0.00273 | -0.135109 | 0.127 | -0.2058275 | 0.000483 | -0.3730976 | 0.0000968 | -0.1580365 | 0.00142 |
| #934 | PGRMC2 | 0.4114258 | 0.02200379 | 0.5562218 | 0.0000288 | 0.4088475 | 0.0012 | 0.2932563 | 0.122 | 0.2957248 | 0.0000582 | 0.4947705 | 0.00877 | 0.4197337 | 0.000133 |
| #935 | BROX | 0.2948478 | 0.02204473 | 0.2974677 | 0.00912 | 0.2254701 | 0.0244 | 0.5227217 | 0.0197 | 0.3585399 | 0.000000249 | 0.1436807 | 0.0789 | 0.2212064 | 0.000139 |
| #936 | ARPC3 | -0.2032636 | 0.02205291 | -0.1401282 | 0.00507 | -0.186918 | 0.00272 | -0.2331853 | 0.00474 | -0.2261249 | 0.00229 | -0.2488136 | 0.0932 | -0.1844116 | 0.0243 |
| #937 | ARID4B | 0.2481736 | 0.02210336 | 0.2349683 | 0.000162 | 0.2145302 | 0.0706 | 0.4405178 | 0.00049 | 0.2064901 | 0.000146 | 0.1543635 | 0.0609 | 0.2381718 | 0.000244 |
| #938 | DYNLL1 | -0.5466378 | 0.0221406 | -0.5599771 | 0.0363 | -0.4771921 | 0.0237 | -0.4867435 | 0.046 | -0.5035098 | 0.00513 | -0.4964466 | 0.0216 | -0.7559576 | 0.000155 |
| #939 | PIP4K2A | 0.2698136 | 0.02216901 | 0.2877987 | 0.0227 | 0.2780571 | 0.00426 | 0.2707143 | 0.0284 | 0.3107355 | 0.0000208 | 0.301303 | 0.00572 | 0.170273 | 0.0719 |
| #940 | CYBA | -0.3604108 | 0.02218315 | -0.352931 | 0.000000651 | -0.2026083 | 0.115 | -0.5082379 | 0.00543 | -0.2570783 | 0.0126 | -0.3636426 | 0.000162 | -0.477967 | 0.0000519 |
| #941 | ZNRD1 | -0.5760041 | 0.0222864 | -0.2307287 | 0.0354 | -0.6564513 | 0.00000548 | -0.4877774 | 0.0982 | -0.8085451 | 6.77E-08 | -0.7474112 | 0.000118 | -0.525111 | 0.0000333 |
| #942 | SENP5 | 0.6276532 | 0.02234754 | 0.6714449 | 0.000000391 | 0.8206091 | 0.000000253 | 0.3316734 | 0.128 | 0.7518212 | 2.46E-08 | 0.5420239 | 0.0064 | 0.6483465 | 0.00000973 |
| #943 | TADA3 | -0.4560024 | 0.02247176 | -0.4771826 | 0.0000848 | -0.4679635 | 0.00000118 | -0.5128232 | 0.121 | -0.2854122 | 0.0138 | -0.4685707 | 0.000158 | -0.5240622 | 0.00000577 |
| #944 | RPP21 | -0.468189 | 0.02261379 | -0.4134611 | 0.000466 | -0.5620543 | 0.0291 | -0.6244756 | 0.102 | -0.5052688 | 0.0000382 | -0.4600161 | 0.00212 | -0.2438579 | 0.0026 |
| #945 | DNAJC11 | 0.4276218 | 0.0226143 | 0.2653614 | 0.0541 | 0.218158 | 0.0527 | 0.6780835 | 0.0223 | 0.3251795 | 0.000257 | 0.6320794 | 0.00626 | 0.4468688 | 0.0000854 |
| #946 | AKT3 | 0.5034694 | 0.02266986 | 0.6096887 | 0.000103 | 0.3669185 | 0.0127 | 0.5498249 | 0.0192 | 0.295311 | 0.103 | 0.6714235 | 0.000818 | 0.5276501 | 0.000188 |
| #947 | DYRK1A | 0.560511 | 0.02269016 | 0.5187066 | 0.000365 | 0.6516991 | 8.41E-08 | 0.2565879 | 0.136 | 0.5640509 | 7.65E-08 | 0.7132606 | 0.000151 | 0.658761 | 0.000000126 |
| #948 | GPS1 | -0.3744608 | 0.02273613 | -0.3645811 | 0.0148 | -0.2167701 | 0.0568 | -0.9768392 | 0.000393 | -0.107885 | 0.0232 | -0.3333809 | 0.0114 | -0.2473085 | 0.0298 |
| #949 | CAB39 | 0.4292017 | 0.02273764 | 0.4819486 | 0.00283 | 0.2888068 | 0.0636 | 0.6058712 | 0.0152 | 0.2268246 | 0.0444 | 0.6102037 | 0.0000599 | 0.3615552 | 0.0103 |
| #950 | NPAT | 0.589747 | 0.02281857 | 0.662522 | 0.00000999 | 0.4127579 | 0.0244 | 1.019311 | 0.00375 | 0.2679749 | 0.0751 | 0.3607282 | 0.0337 | 0.8151877 | 0.00000974 |
| #951 | ECD | 0.5185211 | 0.02285327 | 0.4259436 | 0.00294 | 0.6378522 | 0.0000175 | 0.373798 | 0.134 | 0.4384865 | 0.000124 | 0.5898541 | 0.0000122 | 0.645192 | 0.00000362 |
| #952 | RBM34 | 0.6804393 | 0.0229735 | 0.6846981 | 0.000759 | 0.971277 | 0.0123 | 0.379015 | 0.119 | 0.549859 | 0.002 | 0.5294204 | 0.00334 | 0.9683662 | 0.00000218 |
| #953 | PHIP | 0.3226586 | 0.02298478 | 0.3228533 | 0.00177 | 0.3500287 | 0.000306 | 0.2606828 | 0.0956 | 0.2972978 | 0.000796 | 0.3506595 | 0.0383 | 0.3544297 | 0.00117 |
| #954 | MALT1 | 0.6792779 | 0.02310976 | 0.5875765 | 0.0000327 | 0.9356351 | 1.87E-09 | 0.3522322 | 0.139 | 0.6998308 | 6.79E-10 | 0.7513752 | 0.000000684 | 0.7490178 | 0.000000583 |
| #955 | GMPR2 | -0.386516 | 0.02321525 | -0.2765808 | 0.0108 | -0.3034687 | 0.0034 | -0.287339 | 0.125 | -0.4103433 | 0.000000798 | -0.6365489 | 0.000409 | -0.4048154 | 0.0000478 |
| #956 | COX8A | -0.2693969 | 0.02324473 | -0.2447551 | 0.000763 | -0.2835179 | 0.000106 | -0.2035947 | 0.131 | -0.2313772 | 0.00182 | -0.3831225 | 0.00618 | -0.270014 | 0.0000134 |
| #957 | SEM1 | -0.3541024 | 0.02331864 | -0.388892 | 0.0223 | -0.3989718 | 0.000436 | -0.2300037 | 0.0611 | -0.2630897 | 0.0199 | -0.3458869 | 0.0355 | -0.4977706 | 0.000728 |
| #958 | TSR1 | 0.3613032 | 0.02335191 | 0.3519122 | 0.00333 | 0.4079055 | 0.000168 | 0.2385268 | 0.116 | 0.2505116 | 0.000721 | 0.4376571 | 0.02 | 0.4813058 | 0.0000221 |
| #959 | TULP4 | 0.6295639 | 0.02337094 | 0.6847477 | 0.0000227 | 0.8779065 | 0.00992 | 0.2366073 | 0.13 | 0.5745865 | 0.000337 | 0.6498109 | 0.0000262 | 0.7537244 | 9.77E-09 |
| #960 | VAMP8 | -0.3670824 | 0.02339561 | -0.4389722 | 0.000039 | -0.2968145 | 0.0341 | -0.4005089 | 0.00112 | -0.2564619 | 0.0734 | -0.3293322 | 0.0317 | -0.4804046 | 0.000000287 |
| #961 | CCDC186 | 0.4567398 | 0.02354706 | 0.5020285 | 0.000000213 | 0.5289551 | 0.0804 | 0.3627858 | 0.0593 | 0.4833047 | 0.0000284 | 0.3769183 | 0.00153 | 0.4864465 | 0.000000284 |
| #962 | RC3H2 | 0.3982428 | 0.02354915 | 0.5215334 | 0.00133 | 0.4414389 | 0.00393 | 0.310967 | 0.121 | 0.3443263 | 0.000022 | 0.2621434 | 0.0152 | 0.5090481 | 0.00000106 |
| #963 | HMGCL | -0.5620882 | 0.02354992 | -0.5258637 | 0.0345 | -0.4642956 | 0.0288 | -0.7257408 | 0.0585 | -0.5314358 | 0.000142 | -0.7623332 | 0.00305 | -0.3628599 | 0.0163 |
| #964 | ARHGEF12 | 0.547077 | 0.02355268 | 0.6518358 | 0.00305 | 0.6086665 | 0.00716 | 0.2994475 | 0.116 | 0.6703975 | 0.000641 | 0.6078058 | 0.00424 | 0.444309 | 0.0103 |
| #965 | RNF168 | 0.4250356 | 0.02355354 | 0.474345 | 0.000479 | 0.6378344 | 0.00000054 | 0.3060988 | 0.115 | 0.4824849 | 0.000546 | 0.3209606 | 0.0199 | 0.3284898 | 0.00522 |
| #966 | MFSD6 | 0.6033785 | 0.02356387 | 0.5989205 | 0.00144 | 0.3198217 | 0.126 | 0.8584724 | 0.00278 | 0.4509926 | 0.00211 | 0.6263004 | 0.00914 | 0.7657637 | 0.00000149 |
| #967 | CCDC88A | 0.4774962 | 0.02356395 | 0.8209615 | 0.000415 | 0.4514662 | 0.0000789 | 0.3757982 | 0.0596 | 0.3448734 | 0.00139 | 0.4344699 | 0.0705 | 0.437408 | 0.0094 |
| #968 | DENND4A | 0.5232848 | 0.02368262 | 0.5251277 | 0.0000425 | 0.4653375 | 0.00309 | 0.2501235 | 0.135 | 0.5075806 | 0.000179 | 0.6827479 | 0.00359 | 0.7087913 | 0.00000452 |
| #969 | DENND1B | 0.4382454 | 0.02374831 | 0.5499214 | 0.000484 | 0.3059072 | 0.00739 | 0.4684605 | 0.0912 | 0.5847183 | 0.0146 | 0.4992401 | 0.000728 | 0.221225 | 0.0281 |
| #970 | TIAL1 | 0.2205557 | 0.02374985 | 0.2786288 | 0.00103 | 0.1601871 | 0.0103 | 0.2750945 | 0.0939 | 0.2134704 | 0.00194 | 0.2249483 | 0.00848 | 0.1710053 | 0.0269 |
| #971 | SNX1 | 0.3723796 | 0.02378579 | 0.3286332 | 0.00307 | 0.3515258 | 0.001 | 0.2768438 | 0.131 | 0.2834842 | 0.00592 | 0.4900761 | 0.00164 | 0.5037146 | 0.00000377 |
| #972 | RAB2B | 0.5380479 | 0.02384197 | 0.4537793 | 0.00975 | 0.7361645 | 0.00275 | 0.3166057 | 0.13 | 0.5431013 | 0.000000977 | 0.5014736 | 0.0000845 | 0.6771632 | 0.000000308 |
| #973 | PKM | -0.3884055 | 0.02388416 | -0.2910465 | 0.00527 | -0.457836 | 0.0104 | -0.8621093 | 0.00222 | -0.2114468 | 0.038 | -0.2421174 | 0.0873 | -0.2658772 | 0.0000794 |
| #974 | NDUFB1 | -0.4894607 | 0.02391241 | -0.4480431 | 0.0113 | -0.5305473 | 0.000055 | -0.3743761 | 0.131 | -0.6234891 | 0.00000123 | -0.5358009 | 0.000604 | -0.4245079 | 0.000766 |
| #975 | KCTD18 | 0.3151396 | 0.02398018 | 0.2709552 | 0.0151 | 0.3293944 | 0.0868 | 0.5318438 | 0.019 | 0.1950775 | 0.00628 | 0.2653877 | 0.0154 | 0.2981792 | 0.00118 |
| #976 | UQCRB | -0.2273452 | 0.0239977 | -0.1812377 | 0.000738 | -0.1780495 | 0.00267 | -0.2522402 | 0.00407 | -0.3414214 | 0.00348 | -0.342438 | 0.0000611 | -0.06868448 | 0.133 |
| #977 | ATP5IF1 | -0.2329186 | 0.02403098 | -0.2340244 | 0.0286 | -0.1845043 | 0.00918 | -0.2741109 | 0.096 | -0.2276084 | 0.00744 | -0.2662652 | 0.000859 | -0.2109987 | 0.00209 |
| #978 | SIGIRR | -0.5864833 | 0.02403282 | -0.5772133 | 0.000656 | -0.7926425 | 0.0107 | -0.8971047 | 0.0203 | -0.5428637 | 0.00155 | -0.3122942 | 0.11 | -0.3967816 | 0.000871 |
| #979 | MTRF1L | 0.3237219 | 0.02403467 | 0.490673 | 0.0000879 | 0.329818 | 0.0019 | 0.3601888 | 0.0931 | 0.1277889 | 0.0335 | 0.3054445 | 0.0145 | 0.3284184 | 0.00112 |
| #980 | CWC15 | 0.2387587 | 0.02407125 | 0.3206436 | 0.00245 | 0.1784025 | 0.0168 | 0.1793788 | 0.117 | 0.2017558 | 0.000287 | 0.3141169 | 0.00142 | 0.2382549 | 0.00612 |
| #981 | ANKRD36B | 0.7902628 | 0.02418457 | 0.9789428 | 0.0000151 | 0.7142269 | 0.0309 | 1.031231 | 0.000498 | 0.6006341 | 0.0521 | 0.5238157 | 0.0615 | 0.8927259 | 0.0000277 |
| #982 | DNAJC10 | 0.5805772 | 0.02422785 | 0.4606027 | 0.000129 | 0.7214135 | 2.15E-09 | 0.1870862 | 0.145 | 0.5263218 | 0.000129 | 0.7746388 | 0.0000102 | 0.8134002 | 2.48E-10 |
| #983 | PBRM1 | 0.3473959 | 0.02432232 | 0.3783447 | 0.00016 | 0.4658525 | 0.00043 | 0.2216295 | 0.141 | 0.2895927 | 0.00444 | 0.3119155 | 0.0000406 | 0.4170406 | 0.0000183 |
| #984 | MICOS13 | -0.2961121 | 0.02433852 | -0.3901716 | 0.0000544 | -0.1937064 | 0.013 | -0.4772149 | 0.109 | -0.2059625 | 0.00945 | -0.3172819 | 0.00866 | -0.1923356 | 0.00549 |
| #985 | CRCP | 0.4822937 | 0.0244642 | 0.5109185 | 0.000483 | 0.5541694 | 0.00000242 | 0.2953457 | 0.146 | 0.5650298 | 0.00000605 | 0.3834258 | 0.000383 | 0.5848729 | 4.31E-08 |
| #986 | HSP90B1 | 0.4060189 | 0.02447764 | 0.4195068 | 0.000000369 | 0.5926892 | 0.0522 | 0.5092567 | 0.00229 | 0.5057661 | 0.000463 | 0.3401046 | 0.00394 | 0.06878974 | 0.0879 |
| #987 | ATP5MC2 | -0.2495568 | 0.0245131 | -0.1886074 | 0.00234 | -0.2965041 | 0.0146 | -0.1634838 | 0.128 | -0.2665427 | 0.000271 | -0.3644207 | 0.00121 | -0.2177822 | 0.000348 |
| #988 | RPL21 | -0.2128587 | 0.02452499 | -0.2039011 | 0.000000249 | -0.1993616 | 0.0000655 | -0.1409509 | 0.0534 | -0.2372295 | 0.000263 | -0.3996328 | 0.000194 | -0.09607641 | 0.0933 |
| #989 | HSF2 | 0.5469413 | 0.02453403 | 0.6871964 | 0.000153 | 0.4903001 | 0.00207 | 0.2831581 | 0.14 | 0.4908231 | 0.00168 | 0.6413654 | 0.00312 | 0.6888046 | 0.00000535 |
| #990 | ARHGAP21 | 0.6419916 | 0.02455633 | 1.017213 | 0.00117 | 0.5192555 | 0.00182 | 0.4446055 | 0.128 | 0.6055415 | 0.000288 | 0.6239941 | 0.0149 | 0.64134 | 0.00145 |
| #991 | CYB5R3 | -0.3410311 | 0.0246258 | -0.3983401 | 0.00106 | -0.3314396 | 0.000502 | -0.5648775 | 0.0321 | -0.1160485 | 0.0373 | -0.1921184 | 0.0768 | -0.4433623 | 0.00000737 |
| #992 | PSPC1 | 0.3605225 | 0.02464824 | 0.3842236 | 0.00729 | 0.3695012 | 0.00493 | 0.2758057 | 0.135 | 0.2919672 | 0.000262 | 0.4223267 | 0.000756 | 0.4193105 | 0.000139 |

| | | | | | | | | | | | | | | | |
|---|---|---|---|---|---|---|---|---|---|---|---|---|---|---|---|
| #993 | CLPP | -0.3399696 | 0.02470453 | -0.3669604 | 0.0000876 | -0.2180418 | 0.0896 | -0.4304166 | 0.0283 | -0.2770658 | 0.0068 | -0.3643821 | 0.0235 | -0.3829508 | 0.000000535 |
| #994 | BNIP2 | 0.3101951 | 0.02472285 | 0.3440369 | 0.000387 | 0.3455807 | 0.0000366 | 0.2265524 | 0.141 | 0.3099592 | 0.0000187 | 0.2750961 | 0.00712 | 0.3599455 | 0.0000486 |
| #995 | LCP1 | 0.658348 | 0.02472369 | 0.6925209 | 0.000000103 | 0.704685 | 6.02E-10 | 0.4004058 | 0.148 | 0.7203939 | 0.0000289 | 0.8393384 | 0.000118 | 0.5927442 | 0.000000308 |
| #996 | CD53 | -0.414613 | 0.02474863 | -0.3554338 | 0.00486 | -0.2831882 | 0.0134 | -0.7525681 | 0.00429 | -0.1922739 | 0.117 | -0.4178872 | 0.00843 | -0.486327 | 0.0000651 |
| #997 | PPP2CA | 0.3570348 | 0.02477974 | 0.3639396 | 0.00938 | 0.3407785 | 0.000141 | 0.2180747 | 0.131 | 0.431293 | 4.99E-08 | 0.4934732 | 0.0000149 | 0.2946499 | 0.00767 |
| #998 | SNX14 | 0.320325 | 0.02478566 | 0.2665078 | 0.0209 | 0.2022783 | 0.0623 | 0.5828314 | 0.000665 | 0.369791 | 0.000000284 | 0.3040604 | 0.00266 | 0.1964809 | 0.0622 |
| #999 | ATP5MD | -0.3751169 | 0.02481524 | -0.4178041 | 0.0013 | -0.2871124 | 0.0152 | -0.3274515 | 0.132 | -0.3489439 | 0.0000748 | -0.4839097 | 0.000367 | -0.3854797 | 0.00000305 |
| #1000 | JMJD1C | 0.5967006 | 0.02483858 | 0.6438336 | 2.06E-08 | 0.7540919 | 0.000000142 | 0.2711724 | 0.145 | 0.5748463 | 0.0000263 | 0.6097523 | 0.00357 | 0.726507 | 1.01E-09 |
| #1001 | ZHX1 | 0.557986 | 0.0248439 | 0.698405 | 0.000307 | 0.5560234 | 0.0000478 | 0.2588223 | 0.136 | 0.6206193 | 0.0000014 | 0.5295701 | 0.0123 | 0.6844759 | 0.0000596 |
| #1002 | CHP1 | -0.4007215 | 0.02488241 | -0.3159309 | 0.0483 | -0.3041122 | 0.054 | -0.6948543 | 0.0165 | -0.3196176 | 0.00389 | -0.4205752 | 0.0242 | -0.3492389 | 0.00247 |
| #1003 | RPL18A | -0.1981829 | 0.02494293 | -0.289957 | 0.0000205 | -0.1775838 | 0.0168 | -0.1021408 | 0.121 | -0.1203495 | 0.00973 | -0.3323536 | 0.00148 | -0.1667126 | 0.000776 |
| #1004 | SSR2 | -0.2716142 | 0.02499654 | -0.2309112 | 0.000689 | -0.1667687 | 0.00211 | -0.1562292 | 0.145 | -0.3058863 | 0.000348 | -0.4410845 | 0.00135 | -0.3288052 | 0.00000176 |
| #1005 | TERF2 | 0.5177956 | 0.02501705 | 0.5491488 | 0.00172 | 0.2349283 | 0.106 | 0.5461879 | 0.0379 | 0.3909967 | 0.00276 | 0.6204525 | 0.00149 | 0.7650593 | 0.0000026 |
| #1006 | WDR75 | 0.6261947 | 0.02509435 | 0.7540635 | 0.00000155 | 0.6461273 | 0.00033 | 0.3872637 | 0.149 | 0.5173315 | 0.000418 | 0.6780089 | 0.000702 | 0.7743731 | 4.12E-08 |
| #1007 | CEP290 | 0.4306851 | 0.02512561 | 0.443935 | 0.00104 | 0.3330249 | 0.0692 | 0.4796511 | 0.0235 | 0.2707814 | 0.0521 | 0.6275084 | 0.00424 | 0.4292099 | 0.000723 |
| #1008 | ING4 | -0.4025595 | 0.02514019 | -0.3706498 | 0.0731 | -0.3172713 | 0.00462 | -0.5127574 | 0.0551 | -0.3452552 | 0.00818 | -0.4467452 | 0.00687 | -0.4226781 | 0.003 |
| #1009 | ATP5PD | -0.2313321 | 0.02520955 | -0.2549934 | 0.00757 | -0.1951561 | 0.0076 | -0.2308101 | 0.107 | -0.1219951 | 0.0202 | -0.3677068 | 0.00885 | -0.2173312 | 0.00000083 |
| #1010 | GFPT1 | 0.4383205 | 0.02521655 | 0.4450936 | 0.00737 | 0.5231931 | 0.000907 | 0.3899879 | 0.133 | 0.4966083 | 0.000376 | 0.4854089 | 0.00219 | 0.2896312 | 0.00773 |
| #1011 | EIF4EBP1 | -0.3873938 | 0.02521941 | -0.390285 | 0.00212 | -0.285052 | 0.0254 | -0.3108265 | 0.101 | -0.3254877 | 0.00098 | -0.6809637 | 0.000287 | -0.3317479 | 0.0213 |
| #1012 | RBBP7 | 0.344091 | 0.0253417 | 0.3599653 | 0.0000242 | 0.3122453 | 0.0532 | 0.2661111 | 0.0984 | 0.4182915 | 0.000127 | 0.4706659 | 0.00000591 | 0.2372669 | 0.00034 |
| #1013 | FXYD5 | -0.4634458 | 0.02537506 | -0.4136281 | 0.0345 | -0.6224603 | 0.011 | -0.4434339 | 0.023 | -0.5418992 | 0.000452 | -0.1843144 | 0.0833 | -0.5749389 | 0.00000533 |
| #1014 | ESCO1 | 0.6344563 | 0.02547132 | 0.6419107 | 0.000000812 | 0.8516294 | 0.00000191 | 0.2617473 | 0.153 | 0.6956254 | 0.000000487 | 0.7691782 | 0.0000482 | 0.5866466 | 0.000000641 |
| #1015 | NDUFA13 | -0.2291499 | 0.02548882 | -0.2307601 | 0.00134 | -0.2674937 | 0.000397 | -0.2027325 | 0.141 | -0.2106049 | 0.00472 | -0.259242 | 0.00408 | -0.2040665 | 0.000936 |
| #1016 | TEC | 0.7389369 | 0.02558275 | 0.4190725 | 0.00829 | 0.3841489 | 0.0349 | 0.2289367 | 0.11 | 0.685236 | 0.000231 | 1.650318 | 0.00019 | 1.065909 | 0.0000445 |
| #1017 | OCEL1 | -0.6285751 | 0.02573404 | -0.7500302 | 0.00185 | -0.6303472 | 0.0333 | -0.8544871 | 0.103 | -0.2982966 | 0.0151 | -0.6117976 | 0.000928 | -0.626492 | 0.00000737 |
| #1018 | TRAK2 | 0.5027032 | 0.02590734 | 0.4162533 | 0.0159 | 0.5413122 | 0.00319 | 0.5170509 | 0.135 | 0.4418264 | 0.0000242 | 0.6775825 | 0.00127 | 0.4221938 | 0.00000108 |
| #1019 | NBAS | 0.4367487 | 0.02594559 | 0.4609676 | 0.000259 | 0.4104247 | 0.0135 | 0.4009068 | 0.141 | 0.4752401 | 3.84E-09 | 0.4900303 | 0.000649 | 0.3829229 | 0.000129 |
| #1020 | PRMT3 | 0.614054 | 0.02598048 | 0.5999094 | 0.00022 | 0.6641478 | 0.00234 | 0.2391973 | 0.151 | 0.7727823 | 0.000000851 | 0.7768094 | 0.00266 | 0.631478 | 0.00000329 |
| #1021 | RTL8A | -0.5744696 | 0.02601578 | -0.4418773 | 0.0908 | -0.6806215 | 0.000238 | -0.8346381 | 0.0151 | -0.4847526 | 0.0000048 | -0.346146 | 0.0498 | -0.6587821 | 0.0000322 |
| #1022 | ZNF431 | 0.445665 | 0.0260449 | 0.3697315 | 0.0102 | 0.5032911 | 0.00358 | 0.9228372 | 0.000189 | 0.1687689 | 0.0985 | 0.262447 | 0.0437 | 0.4469141 | 0.0000816 |
| #1023 | PAIP2 | 0.2431984 | 0.02610815 | 0.1692517 | 0.0042 | 0.3480993 | 0.000614 | 0.3525269 | 0.000475 | 0.1738302 | 0.00692 | 0.1092187 | 0.144 | 0.3062635 | 0.0000739 |
| #1024 | ZBTB37 | 0.4198538 | 0.02611166 | 0.3929058 | 0.00215 | 0.4603611 | 0.00142 | 0.3088909 | 0.0847 | 0.3029725 | 0.0102 | 0.4854315 | 0.0581 | 0.5685608 | 0.0000952 |
| #1025 | HNRNPM | 0.1757815 | 0.02612809 | 0.1156224 | 0.0609 | 0.1889087 | 0.0184 | 0.1669828 | 0.0607 | 0.2414544 | 0.00156 | 0.188604 | 0.0121 | 0.1531167 | 0.00312 |
| #1026 | POLR3K | -0.6564888 | 0.02613098 | -0.5970336 | 0.00969 | -0.574947 | 0.0406 | -1.180427 | 0.00207 | -0.2569878 | 0.0951 | -0.7773584 | 0.00884 | -0.5521795 | 0.000503 |
| #1027 | RPL29 | -0.2265632 | 0.02614191 | -0.2368487 | 0.0000703 | -0.1678228 | 0.00772 | -0.1058415 | 0.148 | -0.2686607 | 0.0000299 | -0.3881078 | 0.000188 | -0.1920974 | 0.000494 |
| #1028 | MDH2 | -0.2961499 | 0.02622755 | -0.251657 | 0.0327 | -0.2673047 | 0.000254 | -0.2857351 | 0.0449 | -0.2701405 | 0.000126 | -0.3668082 | 0.0784 | -0.3352539 | 0.00105 |
| #1029 | MSL3 | 0.3792755 | 0.02624317 | 0.3702447 | 0.0000479 | 0.307048 | 0.0196 | 0.2564595 | 0.121 | 0.2212663 | 0.00832 | 0.3950277 | 0.00843 | 0.7256067 | 8.52E-08 |
| #1030 | ATAD5 | 0.6393489 | 0.02626285 | 0.5872634 | 0.000488 | 0.9156145 | 0.00845 | 0.3176253 | 0.148 | 0.5614151 | 0.000227 | 0.6889921 | 0.000776 | 0.7651832 | 2.13E-08 |
| #1031 | PAXBP1 | 0.8673376 | 0.02629281 | 0.9367761 | 0.00000262 | 1.128035 | 0.000000709 | 0.3125119 | 0.158 | 0.9719982 | 0.0000161 | 0.874448 | 0.0000253 | 0.9802567 | 0.000000578 |
| #1032 | NEPRO | 0.3359117 | 0.02629546 | 0.3780253 | 0.0123 | 0.4954062 | 0.000192 | 0.363064 | 0.0668 | 0.3279524 | 0.0032 | 0.1796369 | 0.0749 | 0.2713858 | 0.000399 |
| #1033 | NSD2 | 0.4736476 | 0.02635167 | 0.6770995 | 0.000109 | 0.4914401 | 0.000829 | 0.286036 | 0.0883 | 0.5112422 | 0.0000879 | 0.4570714 | 0.0586 | 0.4189965 | 0.0102 |
| #1034 | HSPA13 | 0.4744224 | 0.02638052 | 0.7300686 | 0.000847 | 0.5291832 | 0.0687 | 0.3444262 | 0.0839 | 0.3149012 | 0.00119 | 0.5248387 | 0.00269 | 0.4031165 | 0.000888 |
| #1035 | DYNLL2 | 0.34777 | 0.02638573 | 0.3308502 | 0.016 | 0.3403406 | 0.0166 | 0.4155168 | 0.102 | 0.2938784 | 0.00265 | 0.3310868 | 0.0205 | 0.3749469 | 0.000961 |
| #1036 | EEF1D | -0.2266772 | 0.02640462 | -0.2515904 | 0.000028 | -0.1093379 | 0.0988 | -0.1599576 | 0.0549 | -0.188857 | 0.00456 | -0.4292386 | 4.94E-08 | -0.2210819 | 0.0000987 |
| #1037 | HEIH | -0.5532986 | 0.02643985 | -0.3958743 | 0.0312 | -0.7013899 | 0.00263 | -0.7362601 | 0.107 | -0.5736669 | 0.00013 | -0.5539974 | 0.0173 | -0.3586033 | 0.000737 |
| #1038 | TOLLIP | -0.4029655 | 0.02648842 | -0.4543576 | 0.0154 | -0.2720434 | 0.00148 | -0.5700628 | 0.142 | -0.2450722 | 0.000206 | -0.4772575 | 0.000169 | -0.3989994 | 0.000035 |
| #1039 | SMG7 | 0.5410437 | 0.02657474 | 0.6851393 | 0.000000185 | 0.6399507 | 0.000131 | 0.293885 | 0.159 | 0.3819772 | 0.000119 | 0.7137834 | 0.00000832 | 0.5315266 | 0.000000166 |
| #1040 | EIF3G | -0.259634 | 0.02660098 | -0.2716828 | 0.0000666 | -0.2860479 | 0.00093 | -0.1959927 | 0.0753 | -0.2743695 | 0.000461 | -0.2617937 | 0.0828 | -0.2679171 | 0.0000324 |
| #1041 | BTBD7 | 0.3636931 | 0.02663855 | 0.2359923 | 0.0584 | 0.2482745 | 0.0235 | 0.6634614 | 0.0245 | 0.2161435 | 0.0353 | 0.4340468 | 0.0166 | 0.3842404 | 0.00148 |
| #1042 | ARIH2 | 0.3437077 | 0.02665521 | 0.403273 | 0.0000337 | 0.3456562 | 0.000235 | 0.2256689 | 0.16 | 0.3631076 | 0.000000148 | 0.3395324 | 0.0000502 | 0.3850081 | 7.34E-08 |
| #1043 | SLC52A2 | -0.4546726 | 0.02673529 | -0.2838125 | 0.153 | -0.4420163 | 0.00102 | -0.8679868 | 0.00182 | -0.3548765 | 0.000444 | -0.3209065 | 0.00419 | -0.458437 | 0.0000225 |
| #1044 | TRAF5 | 0.4464098 | 0.02681078 | 0.2653852 | 0.00105 | 0.2606344 | 0.13 | 0.666253 | 0.00642 | 0.5155612 | 0.0128 | 0.6312537 | 0.00247 | 0.339371 | 0.00815 |
| #1045 | RPS25 | -0.2020042 | 0.02682986 | -0.1589294 | 0.0000124 | -0.1489397 | 0.0000236 | -0.1054715 | 0.161 | -0.1927671 | 0.0000341 | -0.4197111 | 0.00000448 | -0.186206 | 0.000165 |
| #1046 | ZBTB18 | 0.63512 | 0.02692018 | 0.4597611 | 0.0147 | 0.8624866 | 0.000554 | 0.4051446 | 0.136 | 0.5495967 | 0.000118 | 0.6676366 | 0.0103 | 0.8660947 | 0.0000301 |
| #1047 | ANKRD13A | 0.3920351 | 0.02696889 | 0.6158407 | 0.000182 | 0.3678184 | 0.00135 | 0.3271322 | 0.139 | 0.4318296 | 0.00196 | 0.2963959 | 0.0178 | 0.3131939 | 0.00144 |
| #1048 | NUB1 | 0.4448132 | 0.02703745 | 0.4965907 | 0.0262 | 0.4043893 | 0.0907 | 0.4002851 | 0.0161 | 0.5188311 | 0.0286 | 0.4150011 | 0.000377 | 0.4337819 | 0.000322 |
| #1049 | EIF3I | -0.2300155 | 0.02708905 | -0.1860731 | 0.0179 | -0.2511939 | 0.000593 | -0.2530306 | 0.112 | -0.1843249 | 0.00504 | -0.2538256 | 0.0255 | -0.251645 | 0.0015 |
| #1050 | RPLP0 | -0.2537567 | 0.02709685 | -0.3029056 | 0.00000973 | -0.1599692 | 0.0554 | -0.2067046 | 0.00949 | -0.1227339 | 0.0938 | -0.4315368 | 0.0029 | -0.2986904 | 0.00101 |
| #1051 | DHX38 | 0.272637 | 0.02717215 | 0.2532549 | 0.0116 | 0.2094229 | 0.0269 | 0.3558623 | 0.0717 | 0.2614473 | 0.000481 | 0.2901949 | 0.0466 | 0.26564 | 0.00585 |
| #1052 | ASXL2 | 0.436221 | 0.02726775 | 0.5648803 | 0.00217 | 0.4916281 | 0.00654 | 0.3921696 | 0.141 | 0.3314898 | 0.00466 | 0.4417459 | 0.0051 | 0.3954124 | 0.0044 |
| #1053 | PDK3 | 0.3311605 | 0.02731325 | 0.3380596 | 0.00721 | 0.2367464 | 0.0197 | 0.4710826 | 0.0242 | 0.2910824 | 0.000703 | 0.1762971 | 0.112 | 0.473695 | 0.000000303 |
| #1054 | CLIP1 | 0.4304748 | 0.02738042 | 0.673524 | 0.0000263 | 0.3887092 | 0.000177 | 0.2589063 | 0.142 | 0.4421347 | 5.7E-09 | 0.4382454 | 0.000272 | 0.3813291 | 0.0218 |
| #1055 | MAML2 | 0.5022751 | 0.02745058 | 0.3692514 | 0.000641 | 0.3257608 | 0.00858 | 0.6794173 | 0.0279 | 0.1794159 | 0.127 | 0.7590121 | 0.0000655 | 0.7007933 | 0.000114 |
| #1056 | SIMC1 | 0.5938457 | 0.02748801 | 0.4725621 | 0.00595 | 0.7747604 | 0.00155 | 0.7709992 | 0.106 | 0.6324587 | 0.000031 | 0.4973826 | 0.0278 | 0.4149111 | 0.0233 |
| #1057 | FKBP1A | -0.4093181 | 0.02752865 | -0.4668876 | 0.000417 | -0.4320538 | 0.000119 | -0.2642567 | 0.162 | -0.3176496 | 0.000256 | -0.5537965 | 0.0022 | -0.4212644 | 0.0000229 |
| #1058 | SEPTIN6 | 0.2928061 | 0.02754032 | 0.2355699 | 0.00385 | 0.2944891 | 0.0432 | 0.1613123 | 0.11 | 0.3263054 | 0.00803 | 0.1992991 | 0.000403 | 0.539861 | 7.91E-13 |
| #1059 | CAMSAP1 | 0.5471474 | 0.02755125 | 0.7443732 | 0.00116 | 0.4873024 | 0.0205 | 0.295076 | 0.136 | 0.6611659 | 0.000216 | 0.5395446 | 0.00624 | 0.5554223 | 0.00143 |
| #1060 | BIRC2 | 0.4453782 | 0.0276245 | 0.6732122 | 0.0000021 | 0.2546685 | 0.00894 | 0.2409364 | 0.157 | 0.2325018 | 0.0000768 | 0.6711817 | 0.00000882 | 0.5997685 | 0.000000461 |
| #1061 | TMEM205 | -0.3115297 | 0.02764668 | -0.2825866 | 0.0046 | -0.415163 | 0.032 | -0.326071 | 0.0801 | -0.3626441 | 0.000203 | -0.3024679 | 0.0192 | -0.1802456 | 0.0297 |
| #1062 | RAD50 | 0.2714867 | 0.0277399 | 0.3380614 | 0.00404 | 0.2594565 | 0.00388 | 0.264669 | 0.0768 | 0.2149855 | 0.0286 | 0.3329505 | 0.0361 | 0.2187975 | 0.017 |
| #1063 | INO80 | 0.4884185 | 0.02774905 | 0.5919413 | 0.0000104 | 0.5052762 | 0.0000174 | 0.2487703 | 0.166 | 0.4150338 | 0.00000197 | 0.6216836 | 0.0000458 | 0.5478056 | 0.0000119 |

| | | | | | | | | | | | | | | | |
|---|---|---|---|---|---|---|---|---|---|---|---|---|---|---|---|
| #1064 | SNX19 | 0.3486483 | 0.02782743 | 0.3734222 | 0.000454 | 0.2906294 | 0.00999 | 0.3476407 | 0.12 | 0.3887244 | 0.00234 | 0.264954 | 0.0341 | 0.4265189 | 0.0000137 |
| #1065 | ZNF83 | 0.7789773 | 0.02787412 | 0.7330574 | 0.00000502 | 0.9756517 | 0.000000387 | 0.3019239 | 0.165 | 0.9034621 | 0.0000538 | 0.9179553 | 0.00217 | 0.8418136 | 0.00000813 |
| #1066 | FBXO34 | 0.5193654 | 0.02804428 | 0.619279 | 0.0000525 | 0.5850658 | 0.000375 | 0.2958241 | 0.167 | 0.4750003 | 0.000177 | 0.6163937 | 0.0000968 | 0.5246295 | 0.0000745 |
| #1067 | IBTK | 0.3883061 | 0.02817213 | 0.3189118 | 0.00722 | 0.4312037 | 0.00000316 | 0.3210156 | 0.118 | 0.5293048 | 0.000000176 | 0.5311261 | 0.00117 | 0.1982744 | 0.0426 |
| #1068 | PSME4 | 0.401911 | 0.0282839 | 0.2518227 | 0.0714 | 0.4530243 | 0.000148 | 0.3097932 | 0.0927 | 0.4193136 | 0.00000263 | 0.427553 | 0.00379 | 0.5499593 | 0.00164 |
| #1069 | CUL5 | 0.2993695 | 0.02839302 | 0.180285 | 0.017 | 0.3020407 | 0.00676 | 0.3093185 | 0.128 | 0.3526193 | 0.000308 | 0.2704809 | 0.0187 | 0.3814723 | 0.000000217 |
| #1070 | DNTTIP2 | 0.3908888 | 0.02839398 | 0.3806497 | 0.00225 | 0.4443664 | 0.0000476 | 0.2220268 | 0.167 | 0.4285461 | 0.00000111 | 0.4192336 | 0.000997 | 0.4505102 | 0.000061 |
| #1071 | RBBP4 | 0.2237919 | 0.02843306 | 0.1444416 | 0.0147 | 0.2509634 | 0.106 | 0.229453 | 0.0486 | 0.242198 | 0.000512 | 0.2488091 | 0.000746 | 0.226886 | 0.0000811 |
| #1072 | HIST4H4 | 0.7753207 | 0.02847722 | 0.8863386 | 0.0000173 | 0.5147109 | 0.0218 | 0.5795454 | 0.148 | 1.066245 | 0.0000116 | 0.9166463 | 0.000566 | 0.6884383 | 0.000141 |
| #1073 | KAT6B | 0.403345 | 0.02851937 | 0.3912088 | 0.00251 | 0.5791778 | 0.0000128 | 0.2232188 | 0.138 | 0.2990452 | 0.0108 | 0.4360058 | 0.02 | 0.4914133 | 0.000065 |
| #1074 | NKG7 | -1.288391 | 0.02857138 | -2.457956 | 0.0637 | -0.7534487 | 0.00382 | -2.02816 | 0.0291 | -0.808876 | 0.0707 | -0.5075144 | 0.0014 | -1.174389 | 0.00272 |
| #1075 | NAA80 | -0.6428687 | 0.02857279 | -0.5418871 | 0.00579 | -0.7857691 | 0.0157 | -0.6457143 | 0.119 | -0.5339548 | 0.0121 | -0.501651 | 0.019 | -0.8482359 | 0.0000064 |
| #1076 | PPP1R18 | -0.3963998 | 0.02868411 | -0.3909594 | 0.0754 | -0.2264917 | 0.067 | -0.6155674 | 0.0167 | -0.1294885 | 0.00289 | -0.4189829 | 0.0101 | -0.5969092 | 0.0000512 |
| #1077 | RARS2 | 0.3761214 | 0.02868869 | 0.5750169 | 0.0000514 | 0.4395466 | 0.0122 | 0.3066118 | 0.131 | 0.2716876 | 0.000124 | 0.3076187 | 0.0284 | 0.3562467 | 0.000113 |
| #1078 | TRAM1 | 0.2988631 | 0.02872698 | 0.1557284 | 0.0198 | 0.4267528 | 0.0469 | 0.3755245 | 0.0132 | 0.3766694 | 0.0000306 | 0.3703074 | 0.000886 | 0.08819585 | 0.0915 |
| #1079 | ST8SIA4 | 0.5249673 | 0.02878503 | 0.7730633 | 0.00000021 | 0.5378093 | 0.0000727 | 0.363156 | 0.163 | 0.4207329 | 0.00806 | 0.4836503 | 0.00155 | 0.5713918 | 0.00000886 |
| #1080 | CHML | 0.6140686 | 0.02879341 | 0.5224384 | 0.000601 | 0.8213416 | 0.000513 | 0.5178355 | 0.0158 | 0.4551677 | 0.00691 | 0.236739 | 0.149 | 1.13089 | 7.53E-08 |
| #1081 | RHOG | -0.4806189 | 0.02887853 | -0.3596756 | 0.0119 | -0.6903067 | 0.00681 | -0.3236135 | 0.154 | -0.3779431 | 0.0000408 | -0.559941 | 0.000616 | -0.5722334 | 0.000152 |
| #1082 | DHX37 | 0.543965 | 0.02900691 | 0.4660557 | 0.00488 | 0.6169839 | 0.000554 | 0.3481243 | 0.166 | 0.6191433 | 0.000000935 | 0.6264427 | 0.00241 | 0.5870404 | 0.0000524 |
| #1083 | NCOR1 | 0.2352716 | 0.02907293 | 0.1822528 | 0.0121 | 0.2693892 | 0.000644 | 0.1820695 | 0.15 | 0.2518743 | 0.00116 | 0.289524 | 0.00787 | 0.23652 | 0.00222 |
| #1084 | KANSL3 | 0.3952736 | 0.02913633 | 0.5494769 | 0.00229 | 0.2526593 | 0.0558 | 0.524014 | 0.0259 | 0.3110742 | 0.00046 | 0.4022181 | 0.0739 | 0.3321991 | 0.0164 |
| #1085 | LSP1 | -0.3886168 | 0.02914448 | -0.2889241 | 0.00602 | -0.4105644 | 0.000175 | -0.7814263 | 0.0471 | -0.2417578 | 0.111 | -0.3031341 | 0.0102 | -0.3058945 | 0.000211 |
| #1086 | CHD4 | 0.3042649 | 0.02924723 | 0.3380317 | 0.000256 | 0.3377815 | 0.000184 | 0.1763351 | 0.164 | 0.3438979 | 0.00013 | 0.3318153 | 0.00757 | 0.2977277 | 0.00364 |
| #1087 | BRD1 | 0.4653694 | 0.02931438 | 0.5351395 | 0.0131 | 0.5436557 | 0.00585 | 0.180695 | 0.111 | 0.3639512 | 0.0243 | 0.5410592 | 0.0205 | 0.6277159 | 0.00163 |
| #1088 | CEP170 | 0.4745689 | 0.02935267 | 0.5632853 | 0.00000167 | 0.2216271 | 0.0779 | 0.66181 | 0.0024 | 0.3605895 | 0.000807 | 0.375506 | 0.095 | 0.6645956 | 3.96E-09 |
| #1089 | KIAA0232 | 0.4129296 | 0.02937237 | 0.2891283 | 0.0702 | 0.341944 | 0.0117 | 0.4024792 | 0.0882 | 0.4106066 | 0.00153 | 0.4935692 | 0.00433 | 0.5398504 | 0.00031 |
| #1090 | SSX2IP | 0.5556515 | 0.02937802 | 0.4110217 | 0.0378 | 0.7640761 | 0.00682 | 0.3921848 | 0.13 | 0.5862774 | 0.00184 | 0.6107429 | 0.0000275 | 0.5696063 | 0.00000317 |
| #1091 | TYROBP | -0.7255997 | 0.02939765 | -1.597906 | 0.0092 | -0.2343527 | 0.0185 | -0.8166483 | 0.136 | -0.3794327 | 0.00000105 | -0.386649 | 0.00844 | -0.9386092 | 0.00404 |
| #1092 | KIF3B | 0.3836848 | 0.02954308 | 0.4514265 | 0.0077 | 0.3138013 | 0.0304 | 0.4539438 | 0.0537 | 0.3143581 | 0.0177 | 0.4665127 | 0.066 | 0.3020665 | 0.00174 |
| #1093 | TSC22D4 | -0.4552764 | 0.02957799 | -0.4290826 | 0.00738 | -0.3749176 | 0.0000942 | -0.5093551 | 0.167 | -0.3239243 | 0.000231 | -0.4705881 | 0.00234 | -0.6237904 | 0.000208 |
| #1094 | ANKRD44 | 0.3205379 | 0.02958314 | 0.2487353 | 0.0061 | 0.2806329 | 0.00498 | 0.2681746 | 0.161 | 0.3799407 | 0.000399 | 0.3499698 | 0.00439 | 0.3957739 | 0.000829 |
| #1095 | ADIPOR2 | 0.3103105 | 0.02958362 | 0.3170351 | 0.00891 | 0.3028088 | 0.00464 | 0.3137098 | 0.139 | 0.3168786 | 0.00000711 | 0.2774113 | 0.0244 | 0.3340196 | 0.000192 |
| #1096 | TECR | -0.3625367 | 0.02962767 | -0.4076285 | 0.000961 | -0.3456782 | 0.0005 | -0.2455459 | 0.141 | -0.4450918 | 0.0152 | -0.4840014 | 0.000334 | -0.2472742 | 0.02 |
| #1097 | PPP4R2 | 0.2260693 | 0.02963643 | 0.2227511 | 0.00764 | 0.2106426 | 0.00684 | 0.2129059 | 0.155 | 0.2009737 | 0.00303 | 0.2637991 | 0.00419 | 0.2453432 | 0.00127 |
| #1098 | PNRC1 | 0.5192099 | 0.02967497 | 0.547406 | 0.000673 | 0.7132779 | 0.000127 | 0.6186864 | 0.00129 | 0.1801766 | 0.172 | 0.4923209 | 0.00391 | 0.5633916 | 0.000526 |
| #1099 | VWA8 | 0.6155764 | 0.02975249 | 0.3567571 | 0.0437 | 0.7063836 | 0.00000308 | 0.5153156 | 0.134 | 0.533283 | 0.00117 | 0.8596532 | 0.0000864 | 0.7220661 | 0.00000041 |
| #1100 | ETHE1 | -0.463742 | 0.02978607 | -0.2282971 | 0.096 | -0.6320891 | 0.0499 | -0.7107217 | 0.0188 | -0.4128033 | 0.000541 | -0.4434554 | 0.00774 | -0.3550853 | 0.00569 |
| #1101 | COX7C | -0.3348161 | 0.02979174 | -0.317105 | 0.000132 | -0.3457151 | 0.0000113 | -0.1430329 | 0.179 | -0.4007602 | 0.00000159 | -0.5274711 | 0.0000221 | -0.2748122 | 0.00000265 |
| #1102 | SERBP1 | 0.203633 | 0.0299468 | 0.1862598 | 0.000366 | 0.2006464 | 0.000274 | 0.1065619 | 0.179 | 0.1739595 | 0.0004 | 0.3157933 | 0.00000641 | 0.2385773 | 0.00000251 |
| #1103 | AAMP | -0.3358943 | 0.03004559 | -0.3129978 | 0.026 | -0.3327691 | 0.00103 | -0.3760663 | 0.114 | -0.278129 | 0.0184 | -0.2559161 | 0.0207 | -0.4594879 | 0.000139 |
| #1104 | RRM2B | 0.3481996 | 0.03007287 | 0.4114614 | 0.00566 | 0.4363714 | 0.000359 | 0.4325186 | 0.0939 | 0.3222251 | 0.000534 | 0.2467198 | 0.0415 | 0.2399016 | 0.0384 |
| #1105 | PRDM2 | 0.4402466 | 0.03007719 | 0.4216512 | 0.00104 | 0.5674814 | 0.0000203 | 0.2263121 | 0.176 | 0.3927857 | 0.000114 | 0.5358744 | 0.00344 | 0.4973746 | 0.00000258 |
| #1106 | NSD1 | 0.4765341 | 0.03008612 | 0.4741749 | 0.00000244 | 0.5433654 | 1.28E-09 | 0.2502857 | 0.178 | 0.5714308 | 4.25E-09 | 0.5100254 | 0.00282 | 0.5099227 | 3.13E-09 |
| #1107 | NOSIP | -0.3548323 | 0.03015225 | -0.4594006 | 0.0115 | -0.3945248 | 0.000108 | -0.3162047 | 0.127 | -0.3710752 | 0.000000368 | -0.4076708 | 0.000419 | -0.1801176 | 0.0417 |
| #1108 | DHX32 | 0.7574189 | 0.03023677 | 0.7341885 | 0.00208 | 0.2776221 | 0.175 | 1.308399 | 0.000048 | 0.3673781 | 0.00393 | 0.9857016 | 0.0000143 | 0.8712237 | 0.0000101 |
| #1109 | GOLGA1 | 0.500171 | 0.03028346 | 0.4942042 | 0.000811 | 0.5067248 | 0.00601 | 0.7222772 | 0.0209 | 0.4804405 | 0.00242 | 0.1866946 | 0.152 | 0.6106848 | 0.0000322 |
| #1110 | ITSN2 | 0.4176769 | 0.03039659 | 0.4615372 | 0.0000063 | 0.3937558 | 0.00091 | 0.1850551 | 0.153 | 0.510955 | 0.0000248 | 0.4417131 | 0.0287 | 0.5130449 | 0.00000281 |
| #1111 | PREPL | 0.4694022 | 0.03039983 | 0.5473136 | 0.000344 | 0.4879817 | 0.00994 | 0.5346539 | 0.0148 | 0.2105881 | 0.115 | 0.3837708 | 0.0418 | 0.6521055 | 0.0000137 |
| #1112 | CFL1 | -0.2867205 | 0.03042484 | -0.2541025 | 0.00000254 | -0.3343369 | 0.00000107 | -0.393671 | 0.000292 | -0.1818888 | 0.00199 | -0.2451979 | 0.18 | -0.3111256 | 0.0000587 |
| #1113 | VRK2 | 0.5685 | 0.03064858 | 0.7155029 | 0.00145 | 0.6805962 | 0.00148 | 0.2822518 | 0.14 | 0.5407167 | 0.0408 | 0.6976411 | 0.000262 | 0.4942911 | 0.000000508 |
| #1114 | ALG11 | 0.4927678 | 0.03066666 | 0.3644434 | 0.00346 | 0.2819849 | 0.163 | 1.224332 | 0.00186 | 0.2514808 | 0.0106 | 0.4775025 | 0.0036 | 0.3568634 | 0.00106 |
| #1115 | TRPV2 | -0.4703579 | 0.03070303 | -0.509929 | 0.00159 | -0.3312211 | 0.0069 | -0.8920264 | 0.000107 | -0.1988164 | 0.105 | -0.3594288 | 0.0702 | -0.5307257 | 0.000093 |
| #1116 | MYCBP2 | 0.3513642 | 0.03075735 | 0.3113126 | 0.000579 | 0.288764 | 0.000733 | 0.1623137 | 0.181 | 0.4816839 | 5.75E-08 | 0.4582442 | 0.00207 | 0.4058668 | 0.0000145 |
| #1117 | KIF5B | 0.2859049 | 0.03077325 | 0.2172286 | 0.00191 | 0.397269 | 0.00308 | 0.2672127 | 0.173 | 0.3557652 | 0.00000469 | 0.2328298 | 0.00544 | 0.2451242 | 0.00125 |
| #1118 | MBNL3 | 0.4934536 | 0.0309622 | 0.5510775 | 0.0003 | 0.5593941 | 0.00071 | 0.2962774 | 0.18 | 0.5161137 | 0.00105 | 0.4998269 | 0.00333 | 0.538032 | 0.000305 |
| #1119 | UBE3A | 0.5080993 | 0.03107633 | 0.4876703 | 0.000000173 | 0.5611673 | 0.0000275 | 0.2569889 | 0.186 | 0.478233 | 0.000000277 | 0.6445418 | 0.000139 | 0.6199942 | 1.07E-08 |
| #1120 | SMIM19 | -0.3553025 | 0.03108566 | -0.2409999 | 0.000903 | -0.3863447 | 0.00639 | -0.3614385 | 0.178 | -0.3854868 | 0.00000627 | -0.4118263 | 0.000943 | -0.3457187 | 0.0000441 |
| #1121 | RHOQ | 0.4662093 | 0.03118481 | 0.38021 | 0.00226 | 0.1918909 | 0.115 | 1.270408 | 0.00149 | 0.3331941 | 3.05E-08 | 0.3945007 | 0.0203 | 0.2270524 | 0.0482 |
| #1122 | BORCS6 | -0.6826862 | 0.03120273 | -0.468939 | 0.0363 | -0.8190825 | 0.00105 | -0.7189999 | 0.149 | -0.5623475 | 0.000279 | -0.7821676 | 0.00000487 | -0.7445806 | 0.000339 |
| #1123 | CDK6 | 0.365886 | 0.0312188 | 0.307879 | 0.0148 | 0.4342682 | 0.0217 | 0.3403536 | 0.0131 | 0.528931 | 0.00145 | 0.2588376 | 0.0905 | 0.3250467 | 0.0456 |
| #1124 | PNPT1 | 0.4153767 | 0.0312259 | 0.4213525 | 0.00631 | 0.5146294 | 0.00164 | 0.2357907 | 0.158 | 0.5635105 | 0.00105 | 0.3465616 | 0.0199 | 0.4104155 | 0.000185 |
| #1125 | CTDSP1 | -0.3308877 | 0.03122609 | -0.2629898 | 0.00528 | -0.2982148 | 0.0117 | -0.5434711 | 0.161 | -0.2854235 | 0.00095 | -0.3094174 | 0.00604 | -0.2858094 | 0.00216 |
| #1126 | CCDC93 | 0.4668834 | 0.03127241 | 0.4506319 | 0.00026 | 0.4548818 | 0.002 | 0.384721 | 0.182 | 0.4393878 | 0.0000463 | 0.4711543 | 0.00292 | 0.6005233 | 0.00000249 |
| #1127 | ELP5 | -0.7261516 | 0.03127354 | -0.6539169 | 0.000308 | -0.789284 | 0.0000143 | -0.4406569 | 0.187 | -0.8026925 | 0.0000102 | -0.930826 | 0.0000564 | -0.739533 | 0.0000658 |
| #1128 | PEAK1 | 0.7103265 | 0.03128426 | 0.7611157 | 0.000000344 | 0.7842856 | 0.0000787 | 0.4872733 | 0.182 | 0.5001673 | 0.00415 | 1.019591 | 0.00128 | 0.7095264 | 0.0000424 |
| #1129 | CDC14A | 0.4550022 | 0.03129859 | 0.2571697 | 0.0561 | 0.5927801 | 0.0462 | 0.3825339 | 0.0664 | 0.5035511 | 0.00134 | 0.4787727 | 0.0175 | 0.5152059 | 0.000201 |
| #1130 | OTUB1 | -0.3025949 | 0.03137519 | -0.1965923 | 0.0965 | -0.3748604 | 0.000022 | -0.4694979 | 0.0865 | -0.2203549 | 0.00000136 | -0.307435 | 0.00475 | -0.2468289 | 0.000515 |
| #1131 | PPM1B | 0.3751336 | 0.03138559 | 0.3680126 | 0.000716 | 0.4272287 | 0.00116 | 0.2240918 | 0.173 | 0.3855433 | 0.0000122 | 0.4412874 | 0.0132 | 0.4046377 | 0.000017 |
| #1132 | ENTPD4 | 0.4890609 | 0.03147246 | 0.5852558 | 0.0000138 | 0.6311848 | 0.0116 | 0.3777281 | 0.0882 | 0.468856 | 0.00000897 | 0.4628825 | 0.0877 | 0.408458 | 0.00132 |
| #1133 | NUP54 | 0.320894 | 0.0315138 | 0.4309981 | 0.00164 | 0.2880033 | 0.0153 | 0.2938213 | 0.13 | 0.284174 | 0.000785 | 0.2627846 | 0.041 | 0.3655828 | 0.0000347 |
| #1134 | ETF1 | 0.5274283 | 0.03158165 | 0.6218365 | 0.0000327 | 0.593855 | 0.0000533 | 0.3090186 | 0.188 | 0.4755659 | 0.000000912 | 0.6744214 | 0.000202 | 0.4898724 | 0.00128 |

| | | | | | | | | | | | | | | | |
|---|---|---|---|---|---|---|---|---|---|---|---|---|---|---|---|
| **#1135** | BCCIP | 0.2600229 | 0.03164435 | 0.1893673 | 0.0315 | 0.2321691 | 0.0419 | 0.2558555 | 0.112 | 0.2306304 | 0.00251 | 0.4156708 | 0.00228 | 0.2364442 | 0.000141 |
| **#1136** | EIF3A | 0.2539799 | 0.03170578 | 0.3483258 | 0.000133 | 0.1807133 | 0.0262 | 0.1868103 | 0.119 | 0.2521859 | 0.00417 | 0.3891058 | 0.00521 | 0.1667383 | 0.0352 |
| **#1137** | DAD1 | -0.3963349 | 0.03185015 | -0.4512979 | 0.0000388 | -0.2703438 | 0.174 | -0.3892664 | 0.00338 | -0.4582656 | 6.11E-08 | -0.4112626 | 0.0134 | -0.3975731 | 0.0000844 |
| **#1138** | ZNF33A | 0.4778156 | 0.0318682 | 0.6399914 | 7.5E-09 | 0.5846277 | 0.0000988 | 0.4833067 | 0.0236 | 0.132038 | 0.167 | 0.3090515 | 0.000305 | 0.7178781 | 3.73E-09 |
| **#1139** | GSPT1 | 0.3881369 | 0.03190852 | 0.3653523 | 0.0509 | 0.2936093 | 0.0442 | 0.2586248 | 0.0951 | 0.4333641 | 0.0000675 | 0.4751131 | 0.000953 | 0.5027578 | 0.000222 |
| **#1140** | SIK3 | 0.3959122 | 0.03191379 | 0.4747059 | 0.00177 | 0.3159054 | 0.0317 | 0.303944 | 0.149 | 0.2828661 | 0.00105 | 0.5453608 | 0.00219 | 0.452691 | 0.00597 |
| **#1141** | ADAR | 0.5132194 | 0.03199497 | 0.6113768 | 0.00117 | 0.6848137 | 0.00299 | 0.2685287 | 0.184 | 0.5774097 | 0.00268 | 0.4806867 | 0.0000767 | 0.4565007 | 0.000768 |
| **#1142** | THOC1 | 0.554236 | 0.0320011 | 0.6731056 | 0.00000563 | 0.6789152 | 0.0000264 | 0.2077853 | 0.189 | 0.5719429 | 5.77E-08 | 0.5037585 | 0.00298 | 0.6899086 | 0.000000331 |
| **#1143** | TRAPPC8 | 0.5025858 | 0.0320149 | 0.6718733 | 0.0000307 | 0.3431547 | 0.0491 | 0.4413673 | 0.129 | 0.3352186 | 0.0133 | 0.7005476 | 0.000771 | 0.523353 | 0.0000515 |
| **#1144** | EXOC4 | 0.340526 | 0.03203928 | 0.1263599 | 0.102 | 0.5188908 | 0.00242 | 0.4483899 | 0.044 | 0.3746631 | 0.0000496 | 0.2234302 | 0.0435 | 0.3514224 | 0.00000609 |
| **#1145** | BSCL2 | -0.4474522 | 0.03204008 | -0.3500126 | 0.0303 | -0.4441927 | 0.000447 | -0.5536603 | 0.154 | -0.5193629 | 0.000956 | -0.4773948 | 0.0059 | -0.3400902 | 0.00023 |
| **#1146** | PLEKHM1 | 0.3837706 | 0.03213007 | 0.3828294 | 0.0154 | 0.5290026 | 0.0000393 | 0.3650279 | 0.0854 | 0.371098 | 0.00249 | 0.1717724 | 0.0893 | 0.4828932 | 0.000209 |
| **#1147** | CHMP4B | 0.3427706 | 0.03225777 | 0.3404593 | 0.00577 | 0.3961979 | 0.000249 | 0.2379557 | 0.162 | 0.4461349 | 0.0000764 | 0.3402371 | 0.00538 | 0.2956386 | 0.0204 |
| **#1148** | PPP4R3B | 0.5285737 | 0.03228829 | 0.5311116 | 0.000000521 | 0.6479136 | 0.000016 | 0.1896388 | 0.194 | 0.5729325 | 0.0000717 | 0.6226526 | 0.0000273 | 0.6071932 | 2.15E-08 |
| **#1149** | ZFAND5 | 0.4977724 | 0.03230633 | 0.6509644 | 0.000000896 | 0.2707057 | 0.08 | 0.6527837 | 0.00241 | 0.2382896 | 0.111 | 0.680012 | 0.0000172 | 0.4938789 | 0.0000708 |
| **#1150** | TRIM37 | 0.5135615 | 0.03231827 | 0.1720353 | 0.126 | 0.7286971 | 0.00063 | 0.6730018 | 0.0205 | 0.4588816 | 0.00551 | 0.415051 | 0.0412 | 0.6337024 | 0.0000218 |
| **#1151** | ZNF148 | 0.3432763 | 0.0323338 | 0.4019728 | 0.000015 | 0.3908025 | 0.00322 | 0.1791389 | 0.174 | 0.2010799 | 0.0164 | 0.4475147 | 0.0000066 | 0.439149 | 9.72E-09 |
| **#1152** | NAA30 | 0.5258666 | 0.03243205 | 0.8615705 | 0.0000193 | 0.5942499 | 0.000392 | 0.1582731 | 0.194 | 0.4771908 | 0.00000002 | 0.5415496 | 0.00000864 | 0.5223655 | 0.00000102 |
| **#1153** | CNN2 | -0.3798986 | 0.03243546 | -0.3698974 | 0.00185 | -0.5468474 | 0.00755 | -0.6872035 | 0.0701 | -0.275676 | 0.0279 | -0.2071251 | 0.026 | -0.192642 | 0.0612 |
| **#1154** | UQCRH | -0.2252974 | 0.0324668 | -0.2336218 | 0.000971 | -0.3072508 | 0.0416 | -0.1328008 | 0.141 | -0.2505145 | 0.000229 | -0.2946593 | 0.00296 | -0.1329372 | 0.00798 |
| **#1155** | CTSD | -0.4713667 | 0.03249098 | -0.6518151 | 0.000719 | -0.3204793 | 0.0000488 | -0.4755486 | 0.188 | -0.6144397 | 0.00197 | -0.3937961 | 0.00196 | -0.3721212 | 0.00221 |
| **#1156** | CLTC | 0.5086814 | 0.03249413 | 0.5707192 | 0.00035 | 0.5848382 | 0.000119 | 0.2539905 | 0.194 | 0.479849 | 0.000000252 | 0.6204513 | 0.0000105 | 0.5422401 | 1.84E-08 |
| **#1157** | HCST | -0.5031187 | 0.03260205 | -0.7304181 | 0.0423 | -0.3367081 | 0.00728 | -0.7526565 | 0.112 | -0.5474356 | 0.00000171 | -0.3440326 | 0.0341 | -0.3074613 | 0.0000667 |
| **#1158** | SBDS | 0.4200796 | 0.03260475 | 0.5609687 | 0.00424 | 0.6043746 | 0.0103 | 0.3153147 | 0.115 | 0.3567706 | 0.000472 | 0.3601789 | 0.0299 | 0.3228701 | 0.0354 |
| **#1159** | LIN54 | 0.5413804 | 0.03270887 | 0.5833478 | 0.000303 | 0.3400308 | 0.124 | 0.6335118 | 0.071 | 0.5300157 | 0.000323 | 0.5850035 | 0.000809 | 0.5763727 | 0.0000024 |
| **#1160** | GPBP1L1 | 0.2657237 | 0.03271482 | 0.1815043 | 0.00542 | 0.2865656 | 0.000331 | 0.1961232 | 0.189 | 0.295794 | 0.00000454 | 0.340437 | 0.00103 | 0.2939179 | 0.00001 |
| **#1161** | POM121 | 0.3115988 | 0.03273189 | 0.18534 | 0.0501 | 0.2311855 | 0.00711 | 0.6488952 | 0.0107 | 0.1062509 | 0.126 | 0.464774 | 0.000272 | 0.2331469 | 0.00218 |
| **#1162** | NAA25 | 0.3902085 | 0.03289489 | 0.2703995 | 0.0422 | 0.6375799 | 0.00105 | 0.3344098 | 0.127 | 0.3337905 | 0.000297 | 0.3562562 | 0.0262 | 0.4088151 | 0.000129 |
| **#1163** | SERINC1 | 0.4335504 | 0.03303211 | 0.6837766 | 0.0000864 | 0.5135555 | 0.00148 | 0.3092311 | 0.186 | 0.2127965 | 0.00731 | 0.3859153 | 0.00312 | 0.4960275 | 0.0000356 |
| **#1164** | GOLGA3 | 0.4530744 | 0.03308737 | 0.3844918 | 0.00252 | 0.5667224 | 0.0000955 | 0.2464016 | 0.189 | 0.4928185 | 0.00000317 | 0.4810954 | 0.00721 | 0.5469169 | 0.00000109 |
| **#1165** | TPM4 | 0.3873068 | 0.0331138 | 0.2952593 | 0.00584 | 0.3734888 | 0.0000876 | 0.2967606 | 0.147 | 0.5637615 | 0.000949 | 0.5808119 | 0.00026 | 0.2137585 | 0.0445 |
| **#1166** | PSMD3 | -0.2696579 | 0.03320886 | -0.3082651 | 0.0308 | -0.2592793 | 0.0000185 | -0.2978224 | 0.0805 | -0.2369802 | 0.00595 | -0.2166928 | 0.0816 | -0.2989078 | 0.000363 |
| **#1167** | DIPK2A | 0.6676113 | 0.03322036 | 0.6213058 | 0.000266 | 0.9495605 | 0.0000469 | 0.2234672 | 0.196 | 0.7952338 | 0.0000103 | 0.66281 | 0.00297 | 0.7532902 | 0.000169 |
| **#1168** | GNG5 | -0.334787 | 0.03331775 | -0.3873166 | 0.00287 | -0.2931942 | 0.0000176 | -0.206978 | 0.188 | -0.3487977 | 0.0000269 | -0.3650251 | 0.00887 | -0.4074103 | 0.0000104 |
| **#1169** | ATRAID | -0.3247934 | 0.03334316 | -0.2899644 | 0.000184 | -0.2983119 | 0.00183 | -0.2824298 | 0.19 | -0.3336447 | 0.00000808 | -0.4424462 | 0.00799 | -0.3019631 | 0.0000117 |
| **#1170** | PSMD6 | 0.2938303 | 0.03343842 | 0.4269816 | 0.0000901 | 0.1575332 | 0.0219 | 0.2042646 | 0.177 | 0.1994929 | 0.00188 | 0.4434559 | 0.0000149 | 0.3312533 | 0.000000447 |
| **#1171** | DOK1 | -0.711591 | 0.03344987 | -0.7888955 | 0.0172 | -0.8967205 | 0.0333 | -0.8013356 | 0.147 | -0.4913038 | 0.00126 | -0.7941669 | 0.0000829 | -0.4971238 | 0.00184 |
| **#1172** | UBL4A | -0.6044521 | 0.03345949 | -0.672809 | 0.0492 | -0.3125167 | 0.112 | -1.125683 | 0.0246 | -0.2628004 | 0.0105 | -0.7342584 | 0.00481 | -0.518645 | 0.0000514 |
| **#1173** | LARS2 | 0.8397329 | 0.03354759 | 1.155278 | 0.0000246 | 0.5053212 | 0.0152 | 0.2455487 | 0.183 | 0.8910004 | 3.57E-08 | 1.212892 | 0.00264 | 1.028357 | 0.000000293 |
| **#1174** | LDB1 | 0.3956642 | 0.03359482 | 0.2309412 | 0.0645 | 0.4325353 | 0.00532 | 0.8059557 | 0.0197 | 0.3791331 | 0.00622 | 0.2753007 | 0.0971 | 0.2501189 | 0.00877 |
| **#1175** | PPDPF | -0.611777 | 0.03374 | -0.5599443 | 0.0192 | -0.5544199 | 0.0191 | -0.688789 | 0.00175 | -0.4564948 | 0.155 | -0.7121066 | 0.00287 | -0.6989076 | 0.00432 |
| **#1176** | TRIM22 | 0.653314 | 0.03375149 | 0.7367087 | 0.0191 | 0.6362178 | 0.0316 | 0.607271 | 0.0628 | 0.5739666 | 0.0617 | 0.6850582 | 0.00372 | 0.6806616 | 0.0236 |
| **#1177** | ACBD5 | 0.3348286 | 0.03378534 | 0.2988117 | 0.0273 | 0.4529178 | 0.00117 | 0.305809 | 0.148 | 0.4207581 | 0.00000216 | 0.2341042 | 0.0249 | 0.2965709 | 0.00165 |
| **#1178** | GPR107 | 0.3866862 | 0.03385931 | 0.3717151 | 0.0296 | 0.3731373 | 0.00206 | 0.2148741 | 0.167 | 0.3578275 | 0.00000603 | 0.457606 | 0.00384 | 0.544957 | 0.000124 |
| **#1179** | YPEL3 | -0.4116006 | 0.03387249 | -0.2761851 | 0.0391 | -0.1784104 | 0.0746 | -0.4710454 | 0.0716 | -0.6339849 | 0.000626 | -0.5288378 | 0.00215 | -0.3811401 | 0.0152 |
| **#1180** | SOD2 | 0.28907 | 0.03391645 | 0.2996994 | 0.00478 | 0.3167476 | 0.0107 | 0.3831397 | 0.0335 | 0.248045 | 0.136 | 0.233352 | 0.0116 | 0.2534363 | 0.00721 |
| **#1181** | LSM 12.00 | 0.2183789 | 0.03394445 | 0.2892724 | 0.0107 | 0.2543407 | 0.00161 | 0.3350094 | 0.0443 | 0.1806649 | 0.000678 | 0.1282101 | 0.123 | 0.1227757 | 0.0236 |
| **#1182** | CCNI | -0.1925279 | 0.03414126 | -0.1386821 | 0.0481 | -0.1639422 | 0.0117 | -0.1672213 | 0.0913 | -0.2421072 | 0.0386 | -0.1852474 | 0.015 | -0.2579674 | 0.0000842 |
| **#1183** | ANKRD11 | 0.3649931 | 0.03428822 | 0.4516089 | 0.00000554 | 0.3873109 | 0.0154 | 0.338262 | 0.0862 | 0.2814926 | 0.00148 | 0.3225705 | 0.0996 | 0.4087138 | 0.00309 |
| **#1184** | FCER1G | -0.8157022 | 0.0343966 | -1.40835 | 0.00216 | -0.5042248 | 0.00022 | -1.086843 | 0.194 | -0.66265 | 0.000283 | -0.4827902 | 0.0097 | -0.7493551 | 0.000456 |
| **#1185** | PNKD | -0.3306752 | 0.03443362 | -0.295859 | 0.026 | -0.3440242 | 0.00265 | -0.3611162 | 0.142 | -0.4423387 | 0.00511 | -0.3286063 | 0.0244 | -0.2121067 | 0.00601 |
| **#1186** | ATR | 0.4066717 | 0.0345199 | 0.3487583 | 0.000134 | 0.5534638 | 0.000103 | 0.2727359 | 0.189 | 0.558854 | 0.00000025 | 0.3219718 | 0.0182 | 0.3842466 | 0.000186 |
| **#1187** | RNF20 | 0.3702798 | 0.03452252 | 0.2477562 | 0.0519 | 0.267545 | 0.0322 | 0.3258731 | 0.121 | 0.2852746 | 0.00036 | 0.6605523 | 0.00185 | 0.4346774 | 0.000106 |
| **#1188** | KLF13 | 0.4015138 | 0.03452544 | 0.4900299 | 0.00244 | 0.533063 | 0.000422 | 0.465469 | 0.0477 | 0.2474118 | 0.0713 | 0.2899283 | 0.0694 | 0.3831805 | 0.0158 |
| **#1189** | CPEB2 | 0.6229001 | 0.03453514 | 0.682543 | 0.00156 | 0.5367639 | 0.0121 | 1.433912 | 0.0363 | 0.4856377 | 0.000247 | 0.2784668 | 0.113 | 0.3200769 | 0.0438 |
| **#1190** | UFM1 | 0.2294392 | 0.03471057 | 0.2974608 | 0.000059 | 0.2952036 | 0.039 | 0.3723585 | 0.0459 | 0.1311314 | 0.00797 | 0.1202377 | 0.115 | 0.1602433 | 0.000187 |
| **#1191** | RHOT1 | 0.4009161 | 0.03475317 | 0.3758711 | 0.0147 | 0.1606232 | 0.157 | 0.7458775 | 0.00797 | 0.4303113 | 0.000436 | 0.2994814 | 0.0286 | 0.3933322 | 0.00004 |
| **#1192** | SH3BGRL | 0.2018433 | 0.03481137 | 0.1699251 | 0.00554 | 0.2282036 | 0.00118 | 0.2936143 | 0.0221 | 0.2817597 | 0.0000334 | 0.1341922 | 0.133 | 0.103365 | 0.0474 |
| **#1193** | UBXN1 | -0.1972117 | 0.03486011 | -0.1269312 | 0.00206 | -0.1797388 | 0.0194 | -0.2773122 | 0.106 | -0.1892983 | 0.0745 | -0.2801675 | 0.00563 | -0.1298223 | 0.00173 |
| **#1194** | LRBA | 0.663072 | 0.03496361 | 0.7615695 | 0.0000105 | 0.6571781 | 0.00413 | 0.1738501 | 0.205 | 0.8784882 | 0.00000342 | 0.777523 | 0.000553 | 0.7298233 | 3.74E-09 |
| **#1195** | MBNL2 | 0.4156328 | 0.03503289 | 0.656039 | 0.0000208 | 0.4835303 | 0.00118 | 0.2385107 | 0.172 | 0.3645426 | 0.0174 | 0.3704556 | 0.0193 | 0.3807188 | 0.000476 |
| **#1196** | ATP6V0B | -0.2842032 | 0.03505907 | -0.3103481 | 0.00175 | -0.1634495 | 0.0194 | -0.180519 | 0.187 | -0.223302 | 0.000156 | -0.3935332 | 0.00191 | -0.4340677 | 0.0000004 |
| **#1197** | TSPAN32 | -0.654487 | 0.03506092 | -0.5475167 | 0.097 | -1.230701 | 0.00753 | -1.199442 | 0.000438 | -0.4037991 | 0.0079 | -0.2959229 | 0.0544 | -0.2495404 | 0.0431 |
| **#1198** | WDR1 | -0.3895931 | 0.03522104 | -0.353655 | 0.0000367 | -0.2296875 | 0.0254 | -1.105968 | 0.00413 | -0.1250678 | 0.0296 | -0.1575905 | 0.152 | -0.3655896 | 0.0000654 |
| **#1199** | NAA38 | -0.3653396 | 0.03523641 | -0.3352139 | 0.00022 | -0.3900444 | 0.00188 | -0.3352516 | 0.208 | -0.2937598 | 0.00000022 | -0.3662775 | 0.000953 | -0.4714907 | 5.69E-09 |
| **#1200** | ORAI3 | -0.55283 | 0.03538586 | -0.5734907 | 0.00126 | -0.5134832 | 0.0000422 | -0.331057 | 0.206 | -0.5527007 | 0.0000686 | -0.7255434 | 0.00449 | -0.6207053 | 0.000946 |
| **#1201** | ECH1 | -0.4290009 | 0.03558272 | -0.3676714 | 0.0068 | -0.5065735 | 0.00012 | -0.1496894 | 0.204 | -0.6106381 | 1.85E-09 | -0.4855102 | 0.00293 | -0.453923 | 0.0000689 |
| **#1202** | PRKD3 | 0.5154102 | 0.03569799 | 0.7534343 | 4.93E-09 | 0.5630747 | 0.00000719 | 0.2247844 | 0.214 | 0.5409085 | 0.0000277 | 0.4270112 | 0.0000451 | 0.5832484 | 0.000000224 |
| **#1203** | TAF15 | 0.2911359 | 0.03574011 | 0.394707 | 0.0000737 | 0.2983545 | 0.00109 | 0.1927404 | 0.213 | 0.2807916 | 0.000485 | 0.2343407 | 0.000244 | 0.3458811 | 0.00000117 |
| **#1204** | APEH | -0.405317 | 0.03574283 | -0.3846938 | 0.0477 | -0.2053217 | 0.101 | -0.9962668 | 0.0185 | -0.1572319 | 0.044 | -0.3924507 | 0.00178 | -0.2959369 | 0.00119 |
| **#1205** | CNOT10 | 0.3428747 | 0.03578976 | 0.3551965 | 0.012 | 0.3664785 | 0.00138 | 0.1994145 | 0.199 | 0.2921787 | 0.00146 | 0.4397159 | 0.000652 | 0.4042639 | 0.0000823 |

| | | | | | | | | | | | | | | | |
|---|---|---|---|---|---|---|---|---|---|---|---|---|---|---|---|
| #1206 | RPS14 | -0.1909292 | 0.03579477 | -0.155845 | 0.0026 | -0.1647557 | 0.0114 | -0.09244209 | 0.192 | -0.1932659 | 0.00352 | -0.3820325 | 0.00118 | -0.1572342 | 0.0036 |
| #1207 | TMBIM1 | -0.5165345 | 0.03581924 | -0.630762 | 0.0195 | -0.3901109 | 0.0654 | -0.7137276 | 0.0947 | -0.3520448 | 0.00643 | -0.4588697 | 0.027 | -0.5536922 | 0.00197 |
| #1208 | MIER3 | 0.6706406 | 0.0358976 | 0.7151193 | 0.00037 | 1.031576 | 0.0000578 | 0.4080724 | 0.215 | 0.4925022 | 0.0000442 | 0.622818 | 0.00007 | 0.7537555 | 0.00000255 |
| #1209 | THAP12 | 0.3996937 | 0.03590769 | 0.306642 | 0.000427 | 0.3921775 | 0.00457 | 0.2161078 | 0.21 | 0.3945091 | 0.000124 | 0.6466066 | 0.00000121 | 0.4421195 | 0.000106 |
| #1210 | HDGFL2 | -0.3766631 | 0.03608282 | -0.4668669 | 0.0077 | -0.3263977 | 0.0252 | -0.503872 | 0.0558 | -0.1556166 | 0.118 | -0.3929602 | 0.0097 | -0.4142652 | 0.0000448 |
| #1211 | DCAF17 | 0.4671158 | 0.03611872 | 0.4899036 | 0.00596 | 0.53153 | 0.00926 | 0.08037499 | 0.195 | 0.3943038 | 0.0051 | 0.5033215 | 0.00117 | 0.8032607 | 1.38E-08 |
| #1212 | AFG3L2 | 0.3640405 | 0.03612852 | 0.2324206 | 0.107 | 0.3437202 | 0.00121 | 0.2950699 | 0.091 | 0.5483717 | 0.00000542 | 0.3559855 | 0.014 | 0.4086749 | 0.00374 |
| #1213 | ZNF639 | 0.4010699 | 0.03621926 | 0.5255114 | 0.00109 | 0.3499048 | 0.0178 | 0.2827683 | 0.173 | 0.3666551 | 0.00423 | 0.4338385 | 0.0181 | 0.4477416 | 0.00302 |
| #1214 | LGALS8 | 0.318321 | 0.03621947 | 0.2948076 | 0.0399 | 0.4765635 | 0.000000638 | 0.2530588 | 0.174 | 0.2240716 | 0.000282 | 0.3420367 | 0.00335 | 0.319388 | 0.000224 |
| #1215 | RNF213 | 0.4678822 | 0.03625907 | 0.3705501 | 0.0453 | 0.5928342 | 0.0172 | 0.3632523 | 0.117 | 0.6621861 | 0.0284 | 0.44911 | 0.00547 | 0.3693604 | 0.00433 |
| #1216 | SUN2 | -0.3923846 | 0.03626272 | -0.3506159 | 0.00961 | -0.5639648 | 0.0157 | -0.5617792 | 0.11 | -0.2134563 | 0.0553 | -0.3060751 | 0.00877 | -0.3584163 | 0.0182 |
| #1217 | MRPS31 | -0.5268026 | 0.0363443 | -0.5383346 | 0.00952 | -0.6361911 | 0.0231 | -0.6547078 | 0.0253 | -0.6216658 | 0.00652 | -0.3528441 | 0.0603 | -0.3570722 | 0.0932 |
| #1218 | SESN3 | 0.4872932 | 0.03635306 | 0.6655193 | 9.8E-09 | 0.5653384 | 0.0052 | 0.5423749 | 0.0212 | 0.3634346 | 0.0445 | 0.5230037 | 0.0848 | 0.2640883 | 0.0625 |
| #1219 | TMEM179B | -0.3950302 | 0.03637821 | -0.2402818 | 0.123 | -0.3705807 | 0.00769 | -0.6444688 | 0.0693 | -0.3402322 | 0.00911 | -0.3867692 | 0.00856 | -0.3878483 | 0.000816 |
| #1220 | KDM4A | 0.62221 | 0.03642511 | 0.9226089 | 0.0000368 | 0.5602134 | 0.000221 | 0.172976 | 0.213 | 0.5633937 | 0.000031 | 0.7705135 | 0.00499 | 0.7435545 | 0.000018 |
| #1221 | BMT2 | 0.3889265 | 0.03651103 | 0.4736202 | 0.00408 | 0.3529102 | 0.00951 | 0.2980478 | 0.164 | 0.4145977 | 0.000346 | 0.3165037 | 0.0412 | 0.4778791 | 0.0000617 |
| #1222 | FAR1 | 0.5888473 | 0.0365511 | 0.5244244 | 0.00014 | 0.7362591 | 0.00000826 | 0.3407459 | 0.218 | 0.660387 | 0.00000952 | 0.5831264 | 0.000753 | 0.688141 | 0.00000951 |
| #1223 | CLPX | 0.3146731 | 0.03664787 | 0.1986193 | 0.0266 | 0.3748165 | 0.0000854 | 0.357805 | 0.187 | 0.2859504 | 0.00532 | 0.3580784 | 0.000306 | 0.3127692 | 0.00106 |
| #1224 | HNRNPUL2 | 0.2552729 | 0.03669687 | 0.2227665 | 0.00885 | 0.1305613 | 0.171 | 0.3638725 | 0.0284 | 0.2098886 | 0.000173 | 0.3349234 | 0.00878 | 0.2696249 | 0.00343 |
| #1225 | TMEM87B | 0.5966997 | 0.03672613 | 0.8660131 | 0.00012 | 0.8091645 | 0.000357 | 0.4273135 | 0.194 | 0.2788965 | 0.0244 | 0.5807463 | 0.00124 | 0.6180642 | 0.000726 |
| #1226 | SUPT16H | 0.3061619 | 0.03703934 | 0.3025344 | 0.00126 | 0.3627566 | 0.00336 | 0.1799576 | 0.217 | 0.2850005 | 0.0000974 | 0.3361591 | 0.000597 | 0.3705631 | 1.02E-08 |
| #1227 | TCP1 | 0.1846382 | 0.0371955 | 0.2218351 | 0.00971 | 0.1650689 | 0.0367 | 0.2269583 | 0.111 | 0.228491 | 0.000423 | 0.1501909 | 0.0429 | 0.1152848 | 0.0223 |
| #1228 | HINT1 | -0.2458445 | 0.03725256 | -0.2345165 | 0.00333 | -0.1554287 | 0.093 | -0.2087796 | 0.0909 | -0.25453 | 0.00111 | -0.2739356 | 0.0352 | -0.3478767 | 0.0000354 |
| #1229 | MFNG | -0.4789322 | 0.03737241 | -0.4364417 | 0.00112 | -0.6283883 | 0.0445 | -0.4469364 | 0.178 | -0.4004762 | 0.000522 | -0.590317 | 0.000393 | -0.3710338 | 0.0000117 |
| #1230 | SRFBP1 | 0.2970241 | 0.03737633 | 0.3543987 | 0.00478 | 0.181154 | 0.116 | 0.3961922 | 0.0499 | 0.2804597 | 0.00206 | 0.2275904 | 0.0508 | 0.3423498 | 0.000268 |
| #1231 | RPS16 | -0.1775262 | 0.03742652 | -0.1471644 | 0.0000676 | -0.1659175 | 0.00702 | -0.07829728 | 0.193 | -0.1978459 | 0.0163 | -0.3582008 | 0.00223 | -0.1177317 | 0.00599 |
| #1232 | NACA | -0.1776434 | 0.03744588 | -0.1491411 | 0.000488 | -0.1132822 | 0.0647 | -0.1150582 | 0.126 | -0.2830964 | 0.00511 | -0.2747154 | 0.0127 | -0.1305672 | 0.0156 |
| #1233 | KIF13B | 0.4088941 | 0.03750693 | 0.4197691 | 0.0000349 | 0.5176126 | 0.000315 | 0.2222472 | 0.222 | 0.4720943 | 0.0000959 | 0.4392051 | 0.000828 | 0.3824361 | 0.00154 |
| #1234 | MBTD1 | 0.5142491 | 0.03753386 | 0.6515633 | 0.000264 | 0.631634 | 0.0002 | 0.3386824 | 0.183 | 0.4573693 | 0.000584 | 0.3409343 | 0.0413 | 0.665311 | 0.00000499 |
| #1235 | PPP1R10 | 0.5025618 | 0.0375686 | 0.5858074 | 0.000915 | 0.603068 | 0.0000446 | 0.1370844 | 0.202 | 0.3762734 | 0.0208 | 0.5597831 | 0.00203 | 0.7533545 | 0.00000177 |
| #1236 | MYL12A | -0.3386583 | 0.03767207 | -0.3942451 | 0.00484 | -0.2587956 | 0.0461 | -0.3814626 | 0.0504 | -0.2299779 | 0.101 | -0.4293883 | 0.0234 | -0.3380802 | 0.000186 |
| #1237 | NUP205 | 0.4537607 | 0.03768949 | 0.549756 | 0.0041 | 0.5417562 | 0.00263 | 0.2198565 | 0.183 | 0.4930026 | 0.00167 | 0.4337191 | 0.0339 | 0.4844735 | 0.000835 |
| #1238 | CIAO2B | -0.3635245 | 0.03771641 | -0.2550233 | 0.0248 | -0.1863239 | 0.178 | -0.6730183 | 0.0212 | -0.2530751 | 0.00238 | -0.4048922 | 0.000214 | -0.4088143 | 0.0000247 |
| #1239 | ELMO1 | 0.3789062 | 0.03774993 | 0.5505159 | 0.0000103 | 0.325289 | 0.0114 | 0.1787977 | 0.203 | 0.3574437 | 0.00755 | 0.3704791 | 0.00476 | 0.4909121 | 0.00000564 |
| #1240 | NMD3 | 0.3042321 | 0.03779949 | 0.1991237 | 0.0515 | 0.262053 | 0.00435 | 0.698546 | 0.0186 | 0.2854158 | 0.0000722 | 0.1489437 | 0.147 | 0.2313106 | 0.00522 |
| #1241 | KDM1B | 0.4396907 | 0.0379052 | 0.4895273 | 0.00405 | 0.2377561 | 0.12 | 0.3917347 | 0.0771 | 0.4132926 | 0.00389 | 0.7544356 | 0.0036 | 0.3513979 | 0.0187 |
| #1242 | MEA1 | -0.363798 | 0.03790678 | -0.5463887 | 0.000547 | -0.2359369 | 0.0197 | -0.3245486 | 0.2 | -0.3222304 | 0.0000642 | -0.4430502 | 0.00753 | -0.3106331 | 0.0000141 |
| #1243 | APH1A | -0.2850743 | 0.03793843 | -0.2427798 | 0.00066 | -0.2516472 | 0.000513 | -0.4046153 | 0.0648 | -0.2693183 | 0.0039 | -0.261667 | 0.156 | -0.2804185 | 0.00161 |
| #1244 | RSL1D1 | 0.2943601 | 0.03802463 | 0.3443935 | 0.00000225 | 0.3451961 | 0.000228 | 0.1141429 | 0.207 | 0.2195945 | 0.0204 | 0.3993479 | 0.00000605 | 0.3434856 | 3.11E-09 |
| #1245 | TGOLN2 | 0.3210705 | 0.03806686 | 0.414616 | 1.86E-10 | 0.2307427 | 0.00238 | 0.1792654 | 0.226 | 0.2483046 | 0.000154 | 0.4186547 | 0.000077 | 0.4348397 | 1.5E-09 |
| #1246 | TEP1 | 0.4571912 | 0.03807994 | 0.4163975 | 0.0137 | 0.5604432 | 0.0113 | 0.4427335 | 0.118 | 0.5785051 | 0.000144 | 0.4268696 | 0.0281 | 0.3181985 | 0.0572 |
| #1247 | SMARCAD1 | 0.7464324 | 0.03808119 | 0.9893098 | 0.000000708 | 0.9285895 | 0.000546 | 0.2076135 | 0.228 | 0.70579 | 0.0000315 | 0.8019538 | 0.0000128 | 0.8453378 | 2.05E-09 |
| #1248 | GXYLT1 | 0.4437658 | 0.03808403 | 0.5172706 | 0.000685 | 0.2034547 | 0.144 | 0.5010772 | 0.0821 | 0.4072527 | 0.000343 | 0.4738836 | 0.0017 | 0.5596562 | 0.0000146 |
| #1249 | TMEM167B | 0.256231 | 0.03834489 | 0.295881 | 0.017 | 0.2366118 | 0.0127 | 0.2603188 | 0.196 | 0.1730465 | 0.00282 | 0.3130734 | 0.000304 | 0.2584545 | 0.0016 |
| #1250 | ZNF675 | 0.4802214 | 0.03837549 | 0.6750434 | 0.0000061 | 0.4023654 | 0.0321 | 0.3638065 | 0.177 | 0.3554015 | 0.0189 | 0.4667508 | 0.00226 | 0.617961 | 0.00000167 |
| #1251 | FASTKD5 | 0.4342216 | 0.03840256 | 0.3492792 | 0.0461 | 0.4310039 | 0.00644 | 0.4350211 | 0.126 | 0.4706905 | 0.000274 | 0.6086181 | 0.000954 | 0.3107168 | 0.0507 |
| #1252 | PPM1D | 0.4603708 | 0.03843106 | 0.8659289 | 0.000668 | 0.3594974 | 0.014 | 0.2388448 | 0.17 | 0.3092559 | 0.0197 | 0.5718968 | 0.00939 | 0.4168011 | 0.0169 |
| #1253 | GOLGA8A | 0.7683498 | 0.03844272 | 0.8869551 | 0.0023 | 0.5543069 | 0.0262 | 0.8003493 | 0.00667 | 0.2499169 | 0.164 | 1.18773 | 0.0289 | 0.9308405 | 0.0021 |
| #1254 | RASA2 | 0.5114127 | 0.03846539 | 0.5020025 | 0.0000495 | 0.4203274 | 0.000533 | 0.2888732 | 0.23 | 0.4410668 | 0.00000789 | 0.7055395 | 0.000000415 | 0.7106669 | 2.03E-10 |
| #1255 | CYP51A1 | 0.4193784 | 0.0384997 | 0.3262327 | 0.0527 | 0.4585472 | 0.00146 | 0.3571803 | 0.156 | 0.486005 | 0.00215 | 0.5559846 | 0.00886 | 0.3323206 | 0.0102 |
| #1256 | USP47 | 0.3575585 | 0.03862023 | 0.3858753 | 0.000329 | 0.3361589 | 0.0299 | 0.2219472 | 0.195 | 0.3372044 | 0.000245 | 0.3849173 | 0.00574 | 0.4792479 | 0.000000466 |
| #1257 | TARDBP | 0.3446872 | 0.03862239 | 0.3938894 | 0.0000188 | 0.3908881 | 0.00136 | 0.1681552 | 0.194 | 0.3990985 | 0.000239 | 0.2861005 | 0.036 | 0.4299917 | 0.0000786 |
| #1258 | INTS5 | -0.9338919 | 0.03870425 | -0.6754076 | 0.0646 | -0.8795672 | 0.0163 | -1.443267 | 0.122 | -0.5574968 | 0.0167 | -1.268498 | 0.000443 | -0.7791147 | 0.0118 |
| #1259 | USP24 | 0.6718943 | 0.03900253 | 0.517904 | 0.000208 | 0.9339926 | 0.0109 | 0.3108256 | 0.2 | 0.7991602 | 0.000000028 | 0.6858867 | 0.0226 | 0.7835965 | 0.0000246 |
| #1260 | RSF1 | 0.2430462 | 0.03904045 | 0.2483394 | 0.000515 | 0.2618188 | 0.00406 | 0.2787401 | 0.083 | 0.2200839 | 0.00069 | 0.1553479 | 0.146 | 0.2939474 | 0.0000882 |
| #1261 | ATP2A2 | 0.550411 | 0.03921818 | 0.6472886 | 0.0000109 | 0.455399 | 0.0000374 | 0.2408571 | 0.235 | 0.5950908 | 0.000123 | 0.7324323 | 0.0000472 | 0.6313982 | 0.000103 |
| #1262 | USE1 | -0.3117274 | 0.03932508 | -0.3136016 | 0.00284 | -0.2975912 | 0.00717 | -0.4113146 | 0.181 | -0.3148171 | 0.00224 | -0.4025756 | 0.016 | -0.1304644 | 0.0266 |
| #1263 | USP3 | 0.4238575 | 0.0394406 | 0.4438221 | 0.000000383 | 0.1685354 | 0.144 | 0.4579863 | 0.0915 | 0.3952218 | 0.0000123 | 0.5790451 | 0.00122 | 0.4985342 | 0.00000665 |
| #1264 | RNF19A | 0.564141 | 0.03947975 | 0.943164 | 0.0000205 | 0.729486 | 0.00266 | 0.2364844 | 0.228 | 0.5890053 | 4.88E-09 | 0.3513998 | 0.00531 | 0.5353067 | 0.00112 |
| #1265 | NDUFS3 | -0.2979989 | 0.03948334 | -0.4050722 | 0.0012 | -0.2959155 | 0.000363 | -0.2582548 | 0.2 | -0.3338998 | 0.0000149 | -0.23089 | 0.035 | -0.2639609 | 0.0000124 |
| #1266 | JPT1 | -0.3919299 | 0.03958893 | -0.4529353 | 0.0181 | -0.301127 | 0.0129 | -0.5013049 | 0.105 | -0.2752437 | 0.0486 | -0.3069106 | 0.0526 | -0.5140576 | 0.0000666 |
| #1267 | RPAP3 | 0.4010975 | 0.03964345 | 0.3962177 | 0.000354 | 0.5571354 | 0.000806 | 0.2506278 | 0.22 | 0.371933 | 0.000184 | 0.3520709 | 0.016 | 0.4786 | 0.000101 |
| #1268 | NOL8 | 0.3606229 | 0.03965317 | 0.344206 | 0.000139 | 0.1649261 | 0.194 | 0.4360315 | 0.0258 | 0.315544 | 0.00258 | 0.4958596 | 0.0155 | 0.4071702 | 0.0000435 |
| #1269 | ATL2 | 0.4227587 | 0.03966832 | 0.4758128 | 0.00272 | 0.4565237 | 0.0000162 | 0.1715701 | 0.228 | 0.3602854 | 0.00628 | 0.5582787 | 0.00127 | 0.5140812 | 0.0000749 |
| #1270 | RALGAPA1 | 0.4225844 | 0.03967811 | 0.4507404 | 0.00216 | 0.4268145 | 0.000733 | 0.2837404 | 0.147 | 0.5398358 | 0.00103 | 0.348886 | 0.087 | 0.4854894 | 0.0000623 |
| #1271 | RPL10 | -0.1969855 | 0.03969369 | -0.1673864 | 0.000686 | -0.1063022 | 0.0417 | -0.09650676 | 0.176 | -0.2254745 | 0.0156 | -0.4042104 | 0.000613 | -0.1820328 | 0.0033 |
| #1272 | RPS15A | -0.1799829 | 0.03985731 | -0.1582303 | 0.000201 | -0.1142673 | 0.0114 | -0.0841094 | 0.226 | -0.2126201 | 0.000441 | -0.3463337 | 0.00088 | -0.1643367 | 0.000138 |
| #1273 | ARRDC1 | -0.5024258 | 0.03989126 | -0.4101502 | 0.0216 | -0.4914989 | 0.000832 | -1.035588 | 0.0448 | -0.2490294 | 0.00237 | -0.3214973 | 0.169 | -0.5067907 | 0.000767 |
| #1274 | HSPA14 | 0.5274593 | 0.03990005 | 0.2168115 | 0.163 | 0.601551 | 0.0122 | 0.6403449 | 0.0627 | 0.419742 | 0.0000944 | 0.5291023 | 0.00141 | 0.7572038 | 0.00000179 |
| #1275 | RNF169 | 0.3870787 | 0.03990699 | 0.5260097 | 0.0112 | 0.4405783 | 0.0252 | 0.1972878 | 0.0851 | 0.4412226 | 0.00512 | 0.2850601 | 0.0896 | 0.4323134 | 0.0232 |
| #1276 | PIK3C3 | 0.5207594 | 0.04008003 | 0.6220744 | 0.00000312 | 0.5680914 | 0.000038 | 0.2815871 | 0.238 | 0.4690285 | 0.000649 | 0.556042 | 0.00182 | 0.6277332 | 0.00000195 |

| | | | | | | | | | | | | | | | |
|---|---|---|---|---|---|---|---|---|---|---|---|---|---|---|---|
| **#1277** | NUS1 | 0.6145444 | 0.04012172 | 0.7226077 | 0.00000464 | 0.6758477 | 0.00705 | 0.1742096 | 0.233 | 0.6343784 | 1.24E-08 | 0.79582 | 0.000238 | 0.6844033 | 5.16E-09 |
| **#1278** | PPP2R2D | 0.3654857 | 0.0401878 | 0.5087881 | 0.0032 | 0.3457114 | 0.0332 | 0.4684569 | 0.04 | 0.3425094 | 0.00651 | 0.2074696 | 0.122 | 0.3199788 | 0.0362 |
| **#1279** | FIS1 | -0.2462932 | 0.04025049 | -0.1644131 | 0.0264 | -0.2400063 | 0.00138 | -0.1349415 | 0.211 | -0.2732061 | 0.000126 | -0.4354808 | 0.000554 | -0.2297117 | 0.00221 |
| **#1280** | GABARAPL2 | -0.3317848 | 0.04035038 | -0.3476346 | 0.0000665 | -0.2332143 | 0.052 | -0.2694137 | 0.188 | -0.33474 | 0.0000915 | -0.5690313 | 0.000238 | -0.2366747 | 0.00162 |
| **#1281** | TXNRD1 | 0.3403269 | 0.04047886 | 0.4261043 | 0.000556 | 0.3017367 | 0.00674 | 0.2505293 | 0.216 | 0.2841841 | 0.0177 | 0.4589805 | 0.00129 | 0.3204269 | 0.00106 |
| **#1282** | CCT7 | -0.2857027 | 0.04051583 | -0.2069865 | 0.0718 | -0.242585 | 0.000477 | -0.316811 | 0.117 | -0.2138226 | 0.0104 | -0.3486029 | 0.0437 | -0.3854081 | 5.84E-08 |
| **#1283** | JARID2 | 0.3802063 | 0.04059549 | 0.1772662 | 0.0812 | 0.4256485 | 0.000229 | 0.2761234 | 0.149 | 0.2261844 | 0.0128 | 0.6863083 | 0.0000415 | 0.4897073 | 0.000504 |
| **#1284** | ZNF354B | 0.6669299 | 0.04062994 | 0.8277093 | 0.00023 | 0.7631583 | 0.000567 | 0.609607 | 0.189 | 0.5238008 | 0.00827 | 0.2943835 | 0.046 | 0.9829205 | 0.000000856 |
| **#1285** | EFCAB14 | 0.3339817 | 0.04063746 | 0.2856016 | 0.00194 | 0.4033214 | 0.000557 | 0.2950867 | 0.241 | 0.2934595 | 0.0000386 | 0.4144686 | 0.00023 | 0.3119523 | 0.0000762 |
| **#1286** | PAIP1 | 0.3061003 | 0.04063806 | 0.2432735 | 0.0399 | 0.3876105 | 0.0673 | 0.3534478 | 0.0855 | 0.3481762 | 0.0000466 | 0.2672641 | 0.0197 | 0.23683 | 0.0315 |
| **#1287** | TRIM33 | 0.411278 | 0.04065013 | 0.6094835 | 0.00000117 | 0.3662243 | 0.0000877 | 0.1979239 | 0.235 | 0.3809792 | 0.00000567 | 0.4292796 | 0.00845 | 0.4837778 | 0.0000216 |
| **#1288** | ZMYM5 | 0.3218077 | 0.04066752 | 0.4041439 | 0.00241 | 0.281808 | 0.0071 | 0.3499221 | 0.121 | 0.2071105 | 0.102 | 0.3714128 | 0.0106 | 0.3164489 | 0.00109 |
| **#1289** | CKAP5 | 0.5569608 | 0.04071453 | 0.1454402 | 0.223 | 0.6567449 | 0.000704 | 0.7730915 | 0.0113 | 0.4654264 | 0.00205 | 0.8221037 | 0.00611 | 0.4789584 | 0.00108 |
| **#1290** | PGK 1.00 | -0.262256 | 0.04085239 | -0.3481335 | 0.0000167 | -0.2227846 | 0.0000502 | -0.4232499 | 0.0236 | -0.1501146 | 0.0674 | -0.2467177 | 0.139 | -0.1825358 | 0.0155 |
| **#1291** | HNRNPDL | 0.2016473 | 0.04092291 | 0.1995104 | 0.0000272 | 0.3338157 | 0.000141 | 0.1370755 | 0.159 | 0.2141625 | 0.00257 | 0.1009068 | 0.0833 | 0.2244132 | 0.0000132 |
| **#1292** | PRKRIP1 | 0.3909941 | 0.04095487 | 0.2732235 | 0.00279 | 0.2639774 | 0.0679 | 1.204847 | 0.000761 | 0.1276575 | 0.0201 | 0.1768783 | 0.153 | 0.2993811 | 0.000863 |
| **#1293** | NDUFAF3 | -0.4344484 | 0.04103413 | -0.5629323 | 0.0193 | -0.2842085 | 0.156 | -0.4052066 | 0.0617 | -0.395523 | 0.0000405 | -0.4995125 | 0.0075 | -0.4593075 | 0.00158 |
| **#1294** | HERC3 | 0.3925705 | 0.04103988 | 0.2788703 | 0.0606 | 0.2282152 | 0.121 | 0.5562704 | 0.036 | 0.5050347 | 0.000154 | 0.2866196 | 0.0284 | 0.5004128 | 0.0000442 |
| **#1295** | SRSF4 | 0.2572218 | 0.04109403 | 0.2747023 | 0.0000538 | 0.2527575 | 0.00134 | 0.1723492 | 0.238 | 0.3025578 | 5.54E-08 | 0.3121667 | 0.0029 | 0.2287974 | 0.0041 |
| **#1296** | PTPN1 | 0.4721504 | 0.04112525 | 0.6379059 | 0.000135 | 0.4806617 | 0.00192 | 0.1915579 | 0.245 | 0.5129517 | 0.0000366 | 0.5059158 | 0.00000147 | 0.5039094 | 0.000000223 |
| **#1297** | IRAK4 | 0.3003642 | 0.04113549 | 0.4324373 | 0.00000122 | 0.4023729 | 0.0279 | 0.3976162 | 0.0907 | 0.2541118 | 0.00504 | 0.1530872 | 0.118 | 0.1625596 | 0.00559 |
| **#1298** | DCAF16 | 0.4970578 | 0.04121258 | 0.4621668 | 0.0000174 | 0.6541592 | 0.00845 | 0.3089842 | 0.239 | 0.4129663 | 0.00000007 | 0.5898156 | 0.0000122 | 0.5542546 | 0.000000587 |
| **#1299** | S100A11 | -0.7932621 | 0.04121577 | -0.5226264 | 0.0174 | -0.6727035 | 0.00285 | -1.258018 | 0.113 | -0.5211655 | 0.000000401 | -0.4805835 | 0.114 | -1.304475 | 0.00000717 |
| **#1300** | TOMM5 | -0.2506169 | 0.04122586 | -0.2071752 | 0.00735 | -0.1774464 | 0.0155 | -0.2140976 | 0.214 | -0.2436307 | 0.0000551 | -0.3671094 | 0.0103 | -0.2942423 | 0.000000809 |
| **#1301** | RUNX1 | 0.3494442 | 0.04123523 | 0.2203397 | 0.107 | 0.4394451 | 0.0000474 | 0.367742 | 0.0719 | 0.4557587 | 0.0223 | 0.2625679 | 0.0449 | 0.3508119 | 0.00127 |
| **#1302** | MIF4GD | -0.4634156 | 0.0412985 | -0.4162529 | 0.00328 | -0.3213045 | 0.178 | -0.6794511 | 0.0354 | -0.4816489 | 0.0000234 | -0.4402673 | 0.0312 | -0.4415689 | 0.000156 |
| **#1303** | CYBC1 | -0.3718896 | 0.04134042 | -0.3349427 | 0.0165 | -0.3693599 | 0.00477 | -0.5683906 | 0.189 | -0.3050514 | 0.00156 | -0.3344557 | 0.0339 | -0.3191373 | 0.0027 |
| **#1304** | WDR82 | 0.3109328 | 0.04134608 | 0.379001 | 0.000357 | 0.279689 | 0.00089 | 0.2113674 | 0.243 | 0.2719305 | 0.00269 | 0.370179 | 0.0000793 | 0.3534302 | 0.000625 |
| **#1305** | WAC | 0.3286553 | 0.04136801 | 0.4274079 | 0.00000055 | 0.32094 | 0.00041 | 0.1933136 | 0.245 | 0.2959936 | 0.00135 | 0.4177499 | 0.0000935 | 0.3165267 | 0.00103 |
| **#1306** | IKZF5 | 0.4806168 | 0.04150884 | 0.4671208 | 0.00645 | 0.6069217 | 0.000101 | 0.2243683 | 0.24 | 0.3771013 | 0.00162 | 0.5462854 | 0.000697 | 0.661903 | 0.000000733 |
| **#1307** | TMEM87A | 0.2856302 | 0.04153578 | 0.3447398 | 0.000981 | 0.3054497 | 0.0013 | 0.156629 | 0.246 | 0.3364216 | 0.000000239 | 0.3041867 | 0.000126 | 0.2663546 | 0.00106 |
| **#1308** | STIM2 | 0.5321946 | 0.04154133 | 0.5328297 | 0.000193 | 0.6997037 | 0.0000247 | 0.2363289 | 0.249 | 0.6516375 | 3.17E-10 | 0.5519329 | 0.0000505 | 0.520735 | 0.00000034 |
| **#1309** | TBL1XR1 | 0.2842339 | 0.04165444 | 0.1879081 | 0.0279 | 0.3967394 | 0.00576 | 0.3343871 | 0.139 | 0.2991085 | 0.00707 | 0.1797089 | 0.0696 | 0.3075513 | 0.0000424 |
| **#1310** | KXD1 | -0.3242402 | 0.04167109 | -0.2252307 | 0.0583 | -0.2687381 | 0.000217 | -0.4343643 | 0.19 | -0.2702025 | 0.0000675 | -0.3945472 | 0.000951 | -0.3523585 | 2.15E-08 |
| **#1311** | DOCK10 | 0.7965025 | 0.04175392 | 0.9448145 | 7.95E-09 | 0.7147282 | 0.00000803 | 0.3326077 | 0.25 | 1.054205 | 0.0000165 | 0.7984102 | 0.00000213 | 0.9342488 | 2.14E-13 |
| **#1312** | JAK2 | 0.8085883 | 0.04180254 | 1.085339 | 0.000536 | 0.7472286 | 0.0000404 | 0.09587332 | 0.233 | 0.8886938 | 0.0169 | 1.189017 | 0.000000545 | 0.8453783 | 0.000166 |
| **#1313** | FAAP20 | -0.2214427 | 0.04188057 | -0.1847885 | 0.0539 | -0.1881217 | 0.00635 | -0.2783067 | 0.166 | -0.2055445 | 0.000518 | -0.2863413 | 0.0231 | -0.1855534 | 0.00193 |
| **#1314** | RRAGB | 0.4163024 | 0.04190001 | 0.3430541 | 0.00502 | 0.4318861 | 0.00955 | 0.7066623 | 0.00792 | 0.4036672 | 0.000437 | 0.1091276 | 0.228 | 0.5034168 | 0.0000627 |
| **#1315** | NCKAP1L | 0.2362717 | 0.04200949 | 0.1991816 | 0.036 | 0.305438 | 0.00316 | 0.2343482 | 0.112 | 0.2258053 | 0.000102 | 0.2935267 | 0.0173 | 0.1593303 | 0.0834 |
| **#1316** | ATG14 | 0.5787396 | 0.04206107 | 0.6949122 | 0.0000625 | 0.7234385 | 0.0000207 | 0.2037276 | 0.25 | 0.4530674 | 0.000524 | 0.6032991 | 0.00217 | 0.793993 | 0.000000247 |
| **#1317** | RCOR3 | 0.4512585 | 0.04207502 | 0.5619454 | 0.00000235 | 0.7235063 | 0.00000468 | 0.2124167 | 0.24 | 0.334661 | 0.0033 | 0.3917681 | 0.00889 | 0.4832537 | 0.00000458 |
| **#1318** | ATAD2 | 0.4548698 | 0.04207654 | 0.4150962 | 0.0065 | 0.6181333 | 0.000815 | 0.2965812 | 0.235 | 0.4981697 | 0.00000729 | 0.4398899 | 0.00983 | 0.4613482 | 0.000143 |
| **#1319** | CELF2 | 0.2260543 | 0.04208763 | 0.241417 | 0.014 | 0.2398683 | 0.000672 | 0.1671648 | 0.219 | 0.2682762 | 0.000023 | 0.1977518 | 0.0149 | 0.2418473 | 0.00383 |
| **#1320** | RAB8B | 0.4345385 | 0.04212557 | 0.5997877 | 0.0000089 | 0.2833748 | 0.044 | 0.1625488 | 0.208 | 0.3340387 | 0.000852 | 0.6690209 | 0.00000104 | 0.5584599 | 0.0000437 |
| **#1321** | TEX10 | 0.5599533 | 0.04213468 | 0.5659874 | 0.00025 | 0.6251258 | 0.00225 | 0.3215504 | 0.226 | 0.4056231 | 0.0175 | 0.7080434 | 0.00663 | 0.7333899 | 0.0000177 |
| **#1322** | MSI2 | 0.3428246 | 0.04213568 | 0.449932 | 0.000116 | 0.3887428 | 0.0277 | 0.2700218 | 0.0596 | 0.1440284 | 0.145 | 0.3772953 | 0.0207 | 0.4269276 | 0.0000116 |
| **#1323** | USP38 | 0.5395447 | 0.04221162 | 0.7363726 | 0.000179 | 0.5444862 | 0.000979 | 0.06320915 | 0.252 | 0.5268487 | 0.0000219 | 0.6944405 | 0.000204 | 0.6719112 | 0.0000727 |
| **#1324** | COX6C | -0.169651 | 0.04226675 | -0.1895033 | 0.000961 | -0.1582893 | 0.0308 | -0.150961 | 0.0981 | -0.1911502 | 0.0115 | -0.2227352 | 0.0628 | -0.1052668 | 0.0494 |
| **#1325** | TBL3 | -0.5545244 | 0.04232583 | -0.9018447 | 0.00951 | -0.3993112 | 0.101 | -0.8238378 | 0.0996 | -0.2256028 | 0.00772 | -0.6359961 | 0.00454 | -0.340554 | 0.0315 |
| **#1326** | PPP4R1 | 0.3734067 | 0.04232803 | 0.2089673 | 0.197 | 0.4921689 | 0.0000443 | 0.3259082 | 0.0535 | 0.4428739 | 0.0000169 | 0.4199848 | 0.00137 | 0.3505373 | 0.0024 |
| **#1327** | NAA16 | 0.5618366 | 0.04269614 | 0.5763341 | 0.0001 | 0.645899 | 0.00135 | 0.1982434 | 0.251 | 0.4038342 | 0.00378 | 0.6418983 | 0.000193 | 0.9048103 | 6.81E-08 |
| **#1328** | IFIT5 | 0.6868888 | 0.04273302 | 0.8607661 | 0.000197 | 0.9309939 | 0.000342 | 0.1284426 | 0.24 | 0.6517312 | 0.0128 | 0.7058134 | 0.000732 | 0.8435854 | 0.00193 |
| **#1329** | LRCH1 | 0.3348124 | 0.04275122 | 0.5032246 | 0.0000171 | 0.3759033 | 0.0212 | 0.3493205 | 0.139 | 0.3154879 | 0.00337 | 0.1886165 | 0.085 | 0.2763218 | 0.00822 |
| **#1330** | PDIA3 | 0.365008 | 0.04276068 | 0.350166 | 0.00538 | 0.5302195 | 0.00856 | 0.2892127 | 0.0349 | 0.5888015 | 0.0131 | 0.2991256 | 0.0386 | 0.1325229 | 0.156 |
| **#1331** | TMEM258 | -0.3799077 | 0.04277388 | -0.4625083 | 0.000306 | -0.2387433 | 0.197 | -0.2595426 | 0.0593 | -0.3886147 | 0.000000772 | -0.5576835 | 0.00000255 | -0.3723536 | 0.00000472 |
| **#1332** | BCL10 | 0.4274776 | 0.04278455 | 0.5529242 | 0.000123 | 0.6057669 | 0.000000316 | 0.2260738 | 0.234 | 0.3559205 | 0.00000271 | 0.3433675 | 0.0213 | 0.4808124 | 0.00132 |
| **#1333** | NEK1 | 0.4494212 | 0.04279639 | 0.4054498 | 0.0134 | 0.4914396 | 0.000835 | 0.3923574 | 0.167 | 0.3989602 | 0.00768 | 0.3702264 | 0.0673 | 0.6380939 | 0.0000016 |
| **#1334** | GSS | -0.3184601 | 0.04285229 | -0.389953 | 0.129 | -0.2493905 | 0.0165 | -0.4602163 | 0.0868 | -0.157061 | 0.000686 | -0.4054741 | 0.00565 | -0.2486654 | 0.0179 |
| **#1335** | SCO1 | 0.2901369 | 0.04289354 | 0.3231483 | 0.0068 | 0.2571013 | 0.0299 | 0.4061891 | 0.0623 | 0.2020381 | 0.0301 | 0.2185787 | 0.128 | 0.3337658 | 0.000475 |
| **#1336** | ALDH16A1 | -0.4959002 | 0.04290611 | -0.4801675 | 0.00763 | -0.4586459 | 0.000701 | -0.3966373 | 0.247 | -0.4336112 | 0.00156 | -0.664823 | 0.000666 | -0.5415161 | 0.0000469 |
| **#1337** | ZNRF2 | 0.4882724 | 0.04290774 | 0.4884187 | 0.0197 | 0.5853243 | 0.0673 | 0.732592 | 0.0226 | 0.4184204 | 0.0329 | 0.2936852 | 0.0827 | 0.4111941 | 0.0323 |
| **#1338** | MPLKIP | -0.388667 | 0.04297139 | -0.2157035 | 0.186 | -0.4363883 | 0.00246 | -0.6302111 | 0.0347 | -0.3023937 | 0.000588 | -0.4021878 | 0.0298 | -0.3451178 | 0.00451 |
| **#1339** | EFR3A | 0.3514027 | 0.04299391 | 0.2532486 | 0.0898 | 0.3296066 | 0.00633 | 0.4782147 | 0.116 | 0.4472587 | 0.00114 | 0.3498123 | 0.0293 | 0.2502754 | 0.015 |
| **#1340** | TAF12 | -0.398525 | 0.04316711 | -0.3628861 | 0.0182 | -0.4285927 | 0.000312 | -0.6425842 | 0.227 | -0.2936046 | 0.0018 | -0.2868194 | 0.0111 | -0.3766628 | 0.000339 |
| **#1341** | ZNF688 | -0.4335748 | 0.04316882 | -0.2764451 | 0.0329 | -0.2879228 | 0.0267 | -1.16772 | 0.0538 | -0.2576567 | 0.00638 | -0.3110358 | 0.139 | -0.3006676 | 0.00000703 |
| **#1342** | ATAD1 | 0.4079744 | 0.04319133 | 0.6704327 | 0.00000917 | 0.469383 | 0.0068 | 0.3460131 | 0.135 | 0.1598358 | 0.109 | 0.4424674 | 0.00133 | 0.3597146 | 0.00707 |
| **#1343** | RMDN1 | 0.3982462 | 0.04328015 | 0.4278929 | 0.000738 | 0.4067017 | 0.00499 | 0.142938 | 0.253 | 0.2964996 | 0.000544 | 0.5723883 | 0.000256 | 0.5430566 | 0.0000047 |
| **#1344** | AGPS | 0.408719 | 0.04334336 | 0.3945239 | 0.00528 | 0.09890618 | 0.252 | 0.538082 | 0.0015 | 0.4048025 | 0.0000727 | 0.500245 | 0.000998 | 0.5157544 | 0.00001 |
| **#1345** | NVL | 0.4059941 | 0.04338206 | 0.5034479 | 0.000897 | 0.5005146 | 0.00237 | 0.1955713 | 0.149 | 0.2233465 | 0.0621 | 0.36071 | 0.0455 | 0.6523742 | 0.00000213 |
| **#1346** | SNTB1 | 0.618662 | 0.04343805 | 0.8652299 | 0.00279 | 0.7520581 | 0.00288 | 0.2813101 | 0.248 | 0.5057273 | 0.00542 | 0.5791986 | 0.000487 | 0.7284482 | 0.00115 |
| **#1347** | SLAIN2 | 0.4257868 | 0.04356375 | 0.5390996 | 0.0000183 | 0.4593775 | 0.0000512 | 0.1836762 | 0.261 | 0.4242041 | 0.000216 | 0.4785019 | 0.000184 | 0.4698615 | 0.000000381 |

| | | | | | | | | | | | | | | | |
|---|---|---|---|---|---|---|---|---|---|---|---|---|---|---|---|
| **#1348** | CD2AP | 0.5730266 | 0.04356698 | 0.5513941 | 0.0013 | 0.6776496 | 0.0486 | 0.2640821 | 0.185 | 0.7348455 | 0.0224 | 0.5159994 | 0.00364 | 0.6941887 | 0.000011 |
| **#1349** | ZNF506 | 0.525882 | 0.04363185 | 0.6772165 | 1.09E-08 | 0.6766356 | 0.0159 | 0.2473227 | 0.221 | 0.4099336 | 0.00638 | 0.3892067 | 0.0181 | 0.7549769 | 2.63E-11 |
| **#1350** | SMC4 | 0.3785339 | 0.04363432 | 0.4367294 | 0.00411 | 0.6148676 | 0.000171 | 0.232858 | 0.241 | 0.2877755 | 0.00137 | 0.3455417 | 0.014 | 0.353431 | 0.0015 |
| **#1351** | GNPTAB | 0.3975243 | 0.04372782 | 0.2961354 | 0.0173 | 0.5097647 | 0.000951 | 0.208448 | 0.233 | 0.4918976 | 1.29E-08 | 0.62711 | 0.00000431 | 0.2517899 | 0.0109 |
| **#1352** | MLLT10 | 0.241817 | 0.04380748 | 0.3080621 | 0.00745 | 0.1788677 | 0.0259 | 0.3102428 | 0.116 | 0.1595384 | 0.0904 | 0.348198 | 0.0101 | 0.1459932 | 0.0134 |
| **#1353** | TAB2 | 0.3473883 | 0.04389186 | 0.5844629 | 0.0000343 | 0.2508671 | 0.0459 | 0.0944146 | 0.216 | 0.3553728 | 0.000127 | 0.4781724 | 0.00133 | 0.3210402 | 0.0000588 |
| **#1354** | PTTG1IP | 0.3460819 | 0.04396832 | 0.1589386 | 0.201 | 0.3966228 | 0.00192 | 0.5155346 | 0.00716 | 0.4839762 | 0.000367 | 0.2688776 | 0.037 | 0.2525419 | 0.0164 |
| **#1355** | PPP1R11 | -0.4102826 | 0.04398226 | -0.3393068 | 0.00653 | -0.2671672 | 0.174 | -0.7539399 | 0.0563 | -0.2267706 | 0.0253 | -0.4455506 | 0.00188 | -0.4289606 | 0.000000943 |
| **#1356** | DDX21 | 0.3284659 | 0.04400368 | 0.4442166 | 0.000534 | 0.3361233 | 0.00000215 | 0.1788075 | 0.262 | 0.2727888 | 0.000000196 | 0.3781432 | 0.000362 | 0.3607161 | 0.00101 |
| **#1357** | CIB1 | -0.2801282 | 0.04410379 | -0.1056606 | 0.13 | -0.3854241 | 0.025 | -0.3236458 | 0.106 | -0.2891636 | 0.0000946 | -0.2826491 | 0.00303 | -0.2942263 | 0.00035 |
| **#1358** | ZNF253 | 0.5413093 | 0.04411458 | 0.5611229 | 0.000997 | 0.6235004 | 0.021 | 0.3766724 | 0.145 | 0.3950466 | 0.0669 | 0.3822195 | 0.0312 | 0.9092937 | 0.000000217 |
| **#1359** | SNRPE | -0.1729692 | 0.04411947 | -0.140245 | 0.125 | -0.2093017 | 0.000697 | -0.1590541 | 0.045 | -0.2124928 | 0.000596 | -0.2118276 | 0.0378 | -0.1048939 | 0.0556 |
| **#1360** | MOSPD3 | -0.5436276 | 0.04423378 | -0.5556369 | 0.0846 | -0.5478611 | 0.00351 | -0.5847865 | 0.169 | -0.5850771 | 0.000275 | -0.5616327 | 0.0078 | -0.426771 | 0.0000891 |
| **#1361** | GALNT4 | 0.575689 | 0.04449424 | 0.5788589 | 0.00155 | 0.3050595 | 0.198 | 0.8771714 | 0.0126 | 0.305582 | 0.0497 | 0.8384134 | 0.00554 | 0.5490485 | 0.00000159 |
| **#1362** | DIAPH1 | 0.2642045 | 0.04474024 | 0.4193926 | 0.0000617 | 0.1569529 | 0.0817 | 0.2223738 | 0.181 | 0.2180363 | 0.000815 | 0.3419693 | 0.0000902 | 0.2265021 | 0.00437 |
| **#1363** | DTX3L | 0.6480097 | 0.04480672 | 0.7693494 | 0.0314 | 0.4374038 | 0.0398 | 0.8445996 | 0.0406 | 0.5973399 | 0.0427 | 0.7683952 | 0.0137 | 0.4709701 | 0.101 |
| **#1364** | FGFR1OP2 | 0.3283104 | 0.04519236 | 0.2307829 | 0.0308 | 0.4098643 | 0.00103 | 0.6340884 | 0.000666 | 0.3075005 | 0.00173 | 0.2452696 | 0.0632 | 0.1423565 | 0.174 |
| **#1365** | MTR | 0.3832736 | 0.04520235 | 0.3798046 | 0.0189 | 0.5705346 | 0.00248 | 0.5629249 | 0.0353 | 0.2835851 | 0.0156 | 0.1746254 | 0.191 | 0.3281672 | 0.00775 |
| **#1366** | MED16 | -0.4466388 | 0.04529703 | -0.5355425 | 0.00247 | -0.3317913 | 0.00775 | -0.4510924 | 0.188 | -0.2069037 | 0.0708 | -0.5399783 | 0.0023 | -0.6145244 | 0.0000191 |
| **#1367** | CUL2 | 0.5237094 | 0.04551925 | 0.7298608 | 0.0000238 | 0.5821333 | 0.00000319 | 0.1194836 | 0.273 | 0.5765949 | 0.00000853 | 0.5106902 | 0.000323 | 0.6234935 | 4.05E-09 |
| **#1368** | RNMT | 0.3550777 | 0.04561597 | 0.3445076 | 0.00456 | 0.5055481 | 0.00747 | 0.1968439 | 0.224 | 0.32821 | 0.0000137 | 0.249423 | 0.0374 | 0.5059336 | 0.00000576 |
| **#1369** | TARBP1 | 0.4993098 | 0.04566108 | 0.4368427 | 0.04 | 0.7758392 | 0.00618 | 0.2380661 | 0.166 | 0.4832531 | 0.0112 | 0.4432512 | 0.0505 | 0.6186064 | 0.0000224 |
| **#1370** | PDE4B | 0.4642142 | 0.04568117 | 0.4641926 | 0.0106 | 0.6540174 | 0.016 | 0.1589884 | 0.15 | 0.6879427 | 0.00926 | 0.3932686 | 0.0849 | 0.4268758 | 0.00292 |
| **#1371** | S100A4 | -0.5800695 | 0.04573358 | -0.5934824 | 0.0107 | -0.589985 | 0.0934 | -0.2834043 | 0.17 | -0.43571 | 0.0000828 | -0.6607806 | 0.00000785 | -0.9170547 | 1.16E-08 |
| **#1372** | GTPBP4 | 0.2744058 | 0.04580637 | 0.1550278 | 0.145 | 0.3164539 | 0.000269 | 0.2789136 | 0.0845 | 0.3616065 | 0.0000565 | 0.2972587 | 0.0443 | 0.2371745 | 0.000355 |
| **#1373** | RABAC1 | -0.4443809 | 0.04589791 | -0.5799759 | 0.00000677 | -0.3257229 | 0.191 | -0.3932574 | 0.0785 | -0.4893903 | 0.00000442 | -0.4098734 | 0.00549 | -0.4680658 | 0.0000196 |
| **#1374** | MED4 | 0.2416978 | 0.04596014 | 0.3094996 | 0.00114 | 0.2649071 | 0.00124 | 0.1328589 | 0.271 | 0.2386794 | 0.000117 | 0.2512667 | 0.0019 | 0.2529749 | 0.00011 |
| **#1375** | ANKRD17 | 0.3722474 | 0.04602955 | 0.5443284 | 0.0000119 | 0.3724136 | 0.000436 | 0.3034134 | 0.159 | 0.2759855 | 0.0205 | 0.2424439 | 0.0967 | 0.4948998 | 0.00000257 |
| **#1376** | PIAS2 | 0.322733 | 0.04602966 | 0.3515737 | 0.0162 | 0.3908876 | 0.0355 | 0.4472391 | 0.0219 | 0.1832517 | 0.0748 | 0.1690547 | 0.127 | 0.3943913 | 0.000372 |
| **#1377** | DR1 | 0.256236 | 0.04603135 | 0.3068139 | 0.0111 | 0.2219713 | 0.0512 | 0.2372615 | 0.204 | 0.2317449 | 0.000079 | 0.3189516 | 0.00147 | 0.2206726 | 0.00823 |
| **#1378** | STAT3 | 0.4040116 | 0.04609275 | 0.4624793 | 0.00572 | 0.531975 | 0.000123 | 0.1974389 | 0.246 | 0.3444043 | 0.0217 | 0.4470049 | 0.00203 | 0.4407671 | 0.00113 |
| **#1379** | FRS2 | 0.5045559 | 0.0461568 | 0.6307448 | 0.000116 | 0.477897 | 0.00491 | 0.132238 | 0.271 | 0.622946 | 0.0000188 | 0.5146609 | 0.000786 | 0.6488484 | 2.19E-08 |
| **#1380** | ZBED5 | 0.368874 | 0.04638755 | 0.3192918 | 0.000567 | 0.4543303 | 0.000621 | 0.2715145 | 0.206 | 0.2521772 | 0.0408 | 0.4521856 | 0.0297 | 0.4637446 | 0.000113 |
| **#1381** | MBD2 | 0.3509723 | 0.046446 | 0.3620493 | 0.0544 | 0.3264212 | 0.0293 | 0.3551324 | 0.0996 | 0.3120921 | 0.0242 | 0.3631119 | 0.0343 | 0.3870271 | 0.0368 |
| **#1382** | EPG5 | 0.4563042 | 0.04652732 | 0.4913199 | 0.000809 | 0.5656395 | 0.0000592 | 0.2601392 | 0.239 | 0.5994806 | 0.0136 | 0.4508225 | 0.0231 | 0.3704234 | 0.00224 |
| **#1383** | ATF7 | 0.2678611 | 0.0468352 | 0.5336263 | 0.00143 | 0.1440051 | 0.0687 | 0.4914117 | 0.0256 | 0.1021906 | 0.038 | 0.1326766 | 0.14 | 0.2032563 | 0.00692 |
| **#1384** | PLXNC1 | 0.486454 | 0.04687693 | 0.8308993 | 0.0000554 | 0.1986037 | 0.174 | 0.3335122 | 0.0898 | 0.380928 | 0.000743 | 0.5378535 | 0.0165 | 0.6369276 | 0.0000236 |
| **#1385** | CYB561D2 | -0.4543063 | 0.04691463 | -0.3541639 | 0.0293 | -0.1877221 | 0.0849 | -0.6613917 | 0.167 | -0.4650425 | 0.000012 | -0.5118416 | 0.000657 | -0.5456762 | 0.000000435 |
| **#1386** | CDK5RAP2 | 0.40404 | 0.04705809 | 0.4433515 | 0.00141 | 0.4838592 | 0.000339 | 0.3152052 | 0.214 | 0.3682723 | 0.00665 | 0.3873326 | 0.0591 | 0.4262195 | 0.000405 |
| **#1387** | PRR14L | 0.492901 | 0.04711938 | 0.5561255 | 0.0000456 | 0.5398346 | 0.0000289 | 0.2995794 | 0.222 | 0.4856841 | 0.000909 | 0.4024036 | 0.0593 | 0.673779 | 0.000000178 |
| **#1388** | CSTB | -0.5633284 | 0.04717652 | -0.6953777 | 0.000599 | -0.7625593 | 0.0182 | -0.3714736 | 0.259 | -0.3765014 | 0.00398 | -0.4881898 | 0.00146 | -0.6858687 | 2.71E-09 |
| **#1389** | PA2G4 | 0.1563147 | 0.04723216 | 0.07823055 | 0.173 | 0.1379385 | 0.0142 | 0.3269098 | 0.00675 | 0.1742165 | 0.00155 | 0.1295879 | 0.0356 | 0.09100493 | 0.0519 |
| **#1390** | ACBD3 | 0.3573319 | 0.04734103 | 0.3524227 | 0.0399 | 0.3554182 | 0.0228 | 0.2963804 | 0.152 | 0.231991 | 0.0573 | 0.4514294 | 0.00451 | 0.4563499 | 0.00781 |
| **#1391** | COX4I1 | -0.1706636 | 0.04745174 | -0.1727496 | 0.00576 | -0.1262275 | 0.129 | -0.2605456 | 0.0162 | -0.1161195 | 0.0675 | -0.2307571 | 0.032 | -0.1175826 | 0.0338 |
| **#1392** | NFE2L1 | 0.3292056 | 0.04752588 | 0.2455169 | 0.0328 | 0.2276559 | 0.116 | 0.7593593 | 0.0282 | 0.2985834 | 0.00317 | 0.237723 | 0.0852 | 0.206395 | 0.0197 |
| **#1393** | ALKBH8 | 0.5454511 | 0.04756013 | 0.448842 | 0.0129 | 0.9296711 | 0.00179 | 0.3499216 | 0.228 | 0.6657502 | 0.000603 | 0.4108958 | 0.0416 | 0.467626 | 0.000765 |
| **#1394** | DNAJB6 | 0.2140248 | 0.04767136 | 0.2025554 | 0.0062 | 0.2316348 | 0.00311 | 0.1373131 | 0.243 | 0.2016883 | 0.00257 | 0.2171063 | 0.0308 | 0.2938506 | 0.000647 |
| **#1395** | PRPF3 | 0.4685962 | 0.04770806 | 0.4270524 | 0.00528 | 0.3833991 | 0.0274 | 0.1821653 | 0.252 | 0.5706098 | 0.000337 | 0.6240549 | 0.00157 | 0.6242956 | 0.000000416 |
| **#1396** | SMARCC2 | 0.2218999 | 0.04778626 | 0.2134116 | 0.00544 | 0.23844 | 0.000663 | 0.4346121 | 0.00269 | 0.077805 | 0.198 | 0.2237166 | 0.0437 | 0.1434143 | 0.0361 |
| **#1397** | ZNF37A | 0.3286665 | 0.04784605 | 0.2664797 | 0.0445 | 0.4015124 | 0.0615 | 0.2295007 | 0.148 | 0.2316725 | 0.0173 | 0.4083979 | 0.0161 | 0.434436 | 0.000116 |
| **#1398** | PHF11 | 0.3143099 | 0.0479217 | 0.3368294 | 0.0158 | 0.439692 | 0.0135 | 0.32577 | 0.089 | 0.2283528 | 0.109 | 0.2545714 | 0.0344 | 0.3006437 | 0.0258 |
| **#1399** | ENPP4 | 0.5736485 | 0.04794028 | 0.5483255 | 0.0375 | 0.3399368 | 0.112 | 0.9432036 | 0.00367 | 0.3403051 | 0.132 | 0.8229046 | 0.000000199 | 0.4472153 | 0.0029 |
| **#1400** | WWP1 | 0.4005251 | 0.04794138 | 0.5942703 | 0.000101 | 0.3306324 | 0.00416 | 0.2198583 | 0.267 | 0.3151955 | 0.00611 | 0.4285803 | 0.0101 | 0.5146136 | 0.0000195 |
| **#1401** | THOC6 | -0.4395015 | 0.04797693 | -0.3339101 | 0.1 | -0.4300973 | 0.0000212 | -0.8283395 | 0.00947 | -0.3268771 | 0.00365 | -0.3789998 | 0.173 | -0.3387853 | 0.00119 |
| **#1402** | UCP2 | -0.5025507 | 0.04799638 | -0.5552462 | 0.000664 | -0.3495644 | 0.223 | -0.5493054 | 0.0184 | -0.3047729 | 0.0458 | -0.5660764 | 0.000283 | -0.6903387 | 0.0000051 |
| **#1403** | RNF4 | 0.3047649 | 0.04807175 | 0.4878156 | 0.000083 | 0.2222471 | 0.0108 | 0.1542619 | 0.271 | 0.2988196 | 0.000000753 | 0.3696228 | 0.00497 | 0.2958222 | 0.00175 |
| **#1404** | METTL16 | 0.4107015 | 0.04808038 | 0.4538114 | 0.000787 | 0.4357381 | 0.00205 | 0.2301611 | 0.283 | 0.2928022 | 0.00234 | 0.5178545 | 0.000523 | 0.5338417 | 1.55E-08 |
| **#1405** | MLKL | 0.4698481 | 0.04815047 | 0.5741152 | 0.0214 | 0.2362043 | 0.104 | 0.4022633 | 0.148 | 0.4529001 | 0.0146 | 0.5985041 | 0.00162 | 0.5551019 | 0.0000671 |
| **#1406** | SLC66A3 | -0.3940462 | 0.048175 | -0.3671585 | 0.00771 | -0.3535531 | 0.0466 | -0.4008688 | 0.221 | -0.1919409 | 0.0133 | -0.4544089 | 0.000734 | -0.596347 | 1.58E-08 |
| **#1407** | CAND1 | 0.5211701 | 0.0482027 | 0.5442606 | 0.00000822 | 0.7199753 | 0.000168 | 0.2122518 | 0.289 | 0.4652922 | 0.000000684 | 0.5056702 | 6.11E-08 | 0.6795705 | 9.06E-12 |
| **#1408** | STAG1 | 0.4404966 | 0.04824254 | 0.4117257 | 0.0000244 | 0.5523967 | 0.00191 | 0.2106849 | 0.285 | 0.4221262 | 0.0000554 | 0.5263031 | 0.00223 | 0.5197427 | 2.23E-08 |
| **#1409** | ZNF776 | 0.4725693 | 0.04842576 | 0.7168165 | 0.000355 | 0.4132092 | 0.0227 | 0.3983282 | 0.124 | 0.2837468 | 0.0475 | 0.2924482 | 0.0954 | 0.7308667 | 0.000149 |
| **#1410** | SZT2 | 0.3439595 | 0.04849414 | 0.2296514 | 0.0626 | 0.4346654 | 0.000987 | 0.2663224 | 0.187 | 0.2357766 | 0.0146 | 0.5629667 | 0.000991 | 0.3343745 | 0.025 |
| **#1411** | ZMAT1 | 0.4853732 | 0.04855537 | 0.4320736 | 0.00535 | 0.6634476 | 0.0135 | 0.310081 | 0.178 | 0.4317086 | 0.0172 | 0.6183756 | 0.0638 | 0.4565528 | 0.0132 |
| **#1412** | PGGT1B | 0.2887145 | 0.04859278 | 0.3817959 | 0.00000796 | 0.3924691 | 0.0168 | 0.1003828 | 0.266 | 0.2010915 | 0.00494 | 0.3292667 | 0.00408 | 0.3272807 | 0.00000448 |
| **#1413** | KDM3B | 0.346073 | 0.04866257 | 0.2836127 | 0.0239 | 0.3684799 | 0.00874 | 0.2350658 | 0.248 | 0.3025689 | 0.00283 | 0.4308978 | 0.00823 | 0.4558127 | 0.000559 |
| **#1414** | KLHL5 | 0.3722879 | 0.04868062 | 0.3178519 | 0.0144 | 0.1710623 | 0.186 | 0.4662209 | 0.0543 | 0.3386573 | 0.00152 | 0.4911062 | 0.0279 | 0.4488286 | 0.00828 |
| **#1415** | MARK3 | 0.4198732 | 0.04872729 | 0.4305159 | 0.0000137 | 0.4556004 | 0.0000194 | 0.2043425 | 0.283 | 0.6412293 | 0.000000112 | 0.3827133 | 0.00818 | 0.4048378 | 0.00136 |
| **#1416** | APOOL | 0.277296 | 0.04877657 | 0.3452543 | 0.0398 | 0.3002388 | 0.0204 | 0.2518324 | 0.2 | 0.2321707 | 0.00445 | 0.2959631 | 0.00989 | 0.2383166 | 0.018 |
| **#1417** | YTHDC2 | 0.6058224 | 0.04889677 | 0.5151982 | 8.92E-08 | 0.8072723 | 0.000184 | 0.2107974 | 0.291 | 0.6689332 | 4.43E-08 | 0.6986897 | 0.00179 | 0.7340435 | 0.000000206 |
| **#1418** | ASXL1 | 0.4069296 | 0.04894441 | 0.5992548 | 0.00243 | 0.3647206 | 0.0328 | 0.3000464 | 0.149 | 0.2272421 | 0.0767 | 0.424251 | 0.03 | 0.5260631 | 0.00263 |

| | | | | | | | | | | | | | | | |
|---|---|---|---|---|---|---|---|---|---|---|---|---|---|---|---|
| **#1419** | NME3 | -0.2434038 | 0.04899345 | -0.2562924 | 0.0329 | -0.1436937 | 0.00637 | -0.4512955 | 0.101 | -0.3672431 | 0.00353 | -0.1197465 | 0.118 | -0.1221514 | 0.0329 |
| **#1420** | ALPK1 | 0.6964785 | 0.04900837 | 0.8342013 | 0.0109 | 0.6482996 | 0.00096 | 0.4219786 | 0.0714 | 0.7639897 | 0.00303 | 0.4407696 | 0.2 | 1.069632 | 0.00815 |
| **#1421** | WDR48 | 0.3388925 | 0.04909689 | 0.2784654 | 0.027 | 0.3323306 | 0.00254 | 0.1583153 | 0.264 | 0.3256073 | 0.0000111 | 0.4438949 | 0.000912 | 0.4947415 | 0.00000944 |
| **#1422** | PIKFYVE | 0.5094244 | 0.04913458 | 0.6860543 | 0.0000119 | 0.5879567 | 0.0000629 | 0.2248072 | 0.292 | 0.5783604 | 6.31E-09 | 0.5046157 | 0.00238 | 0.4747522 | 0.0000736 |
| **#1423** | BSG | -0.3032587 | 0.04917838 | -0.3520387 | 0.00131 | -0.2416658 | 0.000715 | -0.2661826 | 0.11 | -0.09307363 | 0.183 | -0.4113809 | 0.00000175 | -0.4552105 | 0.00000273 |
| **#1424** | SNAPC2 | -0.5365501 | 0.04917855 | -0.5628323 | 0.00182 | -0.3657235 | 0.00446 | -0.5278595 | 0.284 | -0.4320647 | 0.00289 | -0.6707042 | 0.00156 | -0.6601163 | 0.00000647 |
| **#1425** | PFN1 | -0.2973292 | 0.04929215 | -0.2633268 | 0.000854 | -0.3882798 | 0.000312 | -0.3584365 | 0.000503 | -0.2328373 | 0.00242 | -0.2226281 | 0.291 | -0.3184668 | 0.00102 |
| **#1426** | SENP1 | 0.5194369 | 0.04931324 | 0.4545583 | 0.0174 | 0.6315555 | 0.00107 | 0.213374 | 0.277 | 0.4441023 | 0.00037 | 0.7171835 | 0.000361 | 0.6558475 | 0.000000856 |
| **#1427** | UBE2V1 | 0.1771977 | 0.04953964 | 0.265143 | 0.000195 | 0.1399061 | 0.00838 | 0.1823723 | 0.179 | 0.06642599 | 0.107 | 0.2390064 | 0.00249 | 0.1703326 | 0.00023 |
| **#1428** | ZEB1 | 0.5300332 | 0.0497476 | 0.5759325 | 0.0000241 | 0.4857413 | 0.0331 | 0.4047827 | 0.0801 | 0.4013006 | 0.157 | 0.6163718 | 0.0284 | 0.6960702 | 8.94E-09 |
| **#1429** | TASOR2 | 0.6544946 | 0.04978837 | 0.7665185 | 6.88E-08 | 0.7994272 | 8.42E-08 | 0.2625642 | 0.298 | 0.7614233 | 0.000000106 | 0.6635434 | 0.000283 | 0.6734909 | 3.74E-09 |
| **#1430** | NOP10 | -0.4769585 | 0.0498151 | -0.4211451 | 0.034 | -0.3550154 | 0.000196 | -0.189943 | 0.263 | -0.7427325 | 0.000000229 | -0.6446758 | 0.000323 | -0.5082395 | 0.00114 |
| **#1431** | ATRN | 0.4822968 | 0.0498769 | 0.7667804 | 0.00336 | 0.3070838 | 0.0349 | 0.390128 | 0.239 | 0.3639913 | 0.00163 | 0.599814 | 0.0171 | 0.4659832 | 0.00364 |
| **#1432** | TTF2 | 0.2976609 | 0.04988693 | 0.3024374 | 0.00798 | 0.295034 | 0.0121 | 0.2345528 | 0.206 | 0.2672438 | 0.000691 | 0.3066656 | 0.0721 | 0.3800315 | 0.000772 |
| **#1433** | RGS14 | -0.599642 | 0.04989527 | -0.3726391 | 0.235 | -0.5759166 | 0.00386 | -0.9748189 | 0.00794 | -0.6325641 | 0.0072 | -0.6007515 | 0.0284 | -0.4411616 | 0.0166 |
| **#1434** | IKZF1 | 0.3371912 | 0.04990898 | 0.3470445 | 0.00109 | 0.4658725 | 0.00758 | 0.1968707 | 0.284 | 0.1928554 | 0.000317 | 0.3754872 | 0.0066 | 0.4450168 | 0.0000748 |
| **#1435** | NABP1 | 0.443896 | 0.04997548 | 0.731822 | 0.000054 | 0.4423385 | 0.0466 | 0.3814262 | 0.181 | 0.4413239 | 0.00475 | 0.3725374 | 0.0616 | 0.2939283 | 0.00608 |

Supplementary Table 2

| Position | Symbol | Average Xpr | Average p | Xpr (Day 0 (F)) | p (Day 0 (F)) | Xpr (Day 22 (F)) | p (Day 22 (F)) | Xpr (Day 28 (F)) | p (Day 28 (F)) | Xpr (Day 42 (F)) | p (Day 42 (F)) |
|---|---|---|---|---|---|---|---|---|---|---|---|
| #1 | XIST | 1 | 0 | 19.42448 | 0 | 19.08499 | 0 | 19.04847 | 0 | 19.31931 | 0 |
| #2 | NOTCH2NLA | 1.136245 | 0.002471211 | 1.144378 | 0.000319 | 1.156901 | 0.00234 | 1.020438 | 0.00458 | 1.223266 | 0.00265 |
| #3 | DHX9 | 0.5713194 | 0.004020883 | 0.5443665 | 0.000149 | 0.5837978 | 0.00111 | 0.5223831 | 0.00717 | 0.6347301 | 0.00765 |
| #4 | PKN2 | 0.7330221 | 0.004204099 | 0.8707205 | 0.00615 | 0.7054915 | 0.0034 | 0.6282621 | 0.00472 | 0.7276142 | 0.00254 |
| #5 | KDM6A | 1.587672 | 0.00472242 | 1.650321 | 0.0043 | 1.522509 | 0.00614 | 1.524022 | 0.0055 | 1.653835 | 0.00296 |
| #6 | STK17B | 0.6914698 | 0.004867262 | 0.6246672 | 0.00749 | 0.8025925 | 0.0000239 | 0.6450944 | 0.00535 | 0.693525 | 0.0066 |
| #7 | EXOC6B | 0.9072001 | 0.005254112 | 1.186969 | 0.00266 | 0.9404669 | 0.00563 | 0.609544 | 0.0063 | 0.89182 | 0.00642 |
| #8 | GUF1 | 0.6822029 | 0.005577845 | 0.7543898 | 0.00473 | 0.6624761 | 0.00098 | 0.7067023 | 0.00718 | 0.6052435 | 0.00942 |
| #9 | CLASP2 | 0.5862472 | 0.005628475 | 0.707457 | 0.00801 | 0.6953752 | 0.00885 | 0.2625034 | 0.00322 | 0.6796534 | 0.00244 |
| #10 | DDX3X | 1.029803 | 0.006122316 | 1.155521 | 0.0000355 | 1.056468 | 0.00387 | 0.6819304 | 0.0198 | 1.225294 | 0.000813 |
| #11 | G3BP2 | 0.4080018 | 0.006658441 | 0.4325773 | 0.0073 | 0.4893561 | 0.00299 | 0.3575719 | 0.00193 | 0.352502 | 0.0144 |
| #12 | USP9X | 0.7222853 | 0.006846451 | 0.7710205 | 0.0195 | 0.7311175 | 0.0022 | 0.6628995 | 0.00294 | 0.7241037 | 0.00272 |
| #13 | MEF2A | 0.7691779 | 0.007114503 | 0.7612256 | 0.00265 | 0.8284429 | 0.0035 | 0.7728141 | 0.0183 | 0.7142291 | 0.00397 |
| #14 | ZFX | 0.9021359 | 0.00712985 | 0.8623953 | 0.00026 | 0.965111 | 0.00678 | 0.8034174 | 0.000497 | 0.9776198 | 0.021 |
| #15 | NIN | 0.9232088 | 0.007187902 | 0.8805847 | 0.00165 | 0.9103675 | 0.00377 | 0.8918708 | 0.0221 | 1.010012 | 0.00123 |
| #16 | ASPH | 0.7044835 | 0.00722311 | 0.5424245 | 0.00378 | 0.8002053 | 0.00566 | 0.6708635 | 0.0169 | 0.8044408 | 0.00257 |
| #17 | HNRNPU | 0.5984384 | 0.00740621 | 0.5504128 | 0.00353 | 0.68098 | 0.00397 | 0.5228652 | 0.0142 | 0.6394954 | 0.00789 |
| #18 | ARFGEF1 | 0.7222837 | 0.007550908 | 0.9040513 | 0.00844 | 0.7174097 | 0.011 | 0.5729352 | 0.00644 | 0.6947386 | 0.00433 |
| #19 | SECISBP2L | 0.5260233 | 0.008162985 | 0.4804926 | 0.0184 | 0.4562437 | 0.0073 | 0.6496445 | 0.00455 | 0.5177125 | 0.00239 |
| #20 | ARFGEF2 | 0.8284713 | 0.008590317 | 0.7527047 | 0.0000716 | 0.9412665 | 0.00198 | 0.940917 | 0.0138 | 0.6789971 | 0.0185 |
| #21 | N4BP2 | 0.6329123 | 0.008711493 | 0.7265211 | 0.00264 | 0.6759859 | 0.00439 | 0.6360151 | 0.00674 | 0.4931269 | 0.0211 |
| #22 | LCOR | 0.5531743 | 0.008950259 | 0.5424225 | 0.0141 | 0.5801661 | 0.0044 | 0.4812909 | 0.0108 | 0.6088179 | 0.00652 |
| #23 | VPS13B | 0.8210373 | 0.009368321 | 0.8798064 | 0.0189 | 0.7857275 | 0.012 | 0.7033827 | 0.00441 | 0.9152328 | 0.00215 |
| #24 | SMC1A | 0.6683213 | 0.00945243 | 0.7294062 | 0.000614 | 0.6398563 | 0.0101 | 0.6550419 | 0.00897 | 0.6489809 | 0.0181 |
| #25 | MAN2A1 | 0.7940228 | 0.009481538 | 0.7653211 | 0.000591 | 0.7064467 | 0.00852 | 0.9261173 | 0.0232 | 0.7782061 | 0.00562 |
| #26 | SHOC2 | 0.6201056 | 0.009553228 | 0.6958944 | 0.000549 | 0.6367221 | 0.0146 | 0.5817164 | 0.00416 | 0.5660895 | 0.0189 |
| #27 | JMJD1C | 0.7350239 | 0.009574669 | 0.7958617 | 0.00405 | 0.7995268 | 0.00546 | 0.6162612 | 0.0285 | 0.7284459 | 0.000268 |
| #28 | KIAA1109 | 0.885126 | 0.009752549 | 0.9307159 | 0.0256 | 0.8479262 | 0.0000914 | 0.8988905 | 0.0112 | 0.8629714 | 0.0021 |
| #29 | SMG7 | 0.4729096 | 0.009777015 | 0.4775841 | 0.0146 | 0.4896175 | 0.00866 | 0.3949579 | 0.000733 | 0.5294788 | 0.0151 |
| #30 | HIVEP1 | 0.8224698 | 0.009883247 | 0.851256 | 0.00445 | 0.7298902 | 0.00531 | 0.9004394 | 0.0293 | 0.8082937 | 0.000512 |
| #31 | MED13 | 0.5188122 | 0.009906129 | 0.5490306 | 0.00373 | 0.4928914 | 0.00185 | 0.4325316 | 0.034 | 0.6007952 | 0.0000892 |
| #32 | BIRC6 | 0.5954039 | 0.009917207 | 0.6734339 | 0.00209 | 0.669153 | 0.00955 | 0.4292864 | 0.0244 | 0.6097422 | 0.00365 |
| #33 | INO80D | 0.4981123 | 0.009960864 | 0.523811 | 0.0364 | 0.5293979 | 0.000151 | 0.459013 | 0.00305 | 0.4802275 | 0.000287 |
| #34 | LARS2 | 0.9527316 | 0.01015461 | 1.068141 | 0.0215 | 0.9363311 | 0.00532 | 0.8018949 | 0.00147 | 1.004559 | 0.0123 |
| #35 | ERC1 | 0.9374578 | 0.01018748 | 1.052353 | 0.0186 | 1.035376 | 0.00279 | 0.8428649 | 0.0149 | 0.8192367 | 0.00446 |
| #36 | AHCTF1 | 0.8666476 | 0.0102519 | 0.9398245 | 0.0271 | 0.9696719 | 0.00154 | 0.7490216 | 0.0106 | 0.8080723 | 0.00177 |
| #37 | UBR4 | 0.6074512 | 0.01037659 | 0.7459333 | 0.0164 | 0.6282696 | 0.00533 | 0.4639753 | 0.0077 | 0.5916266 | 0.0121 |
| #38 | RANBP2 | 0.7813215 | 0.01049776 | 0.8706855 | 0.0129 | 0.8978609 | 0.00645 | 0.5551112 | 0.0171 | 0.8016284 | 0.00552 |
| #39 | PPP4R3A | 0.703079 | 0.01053663 | 0.7347229 | 0.00181 | 0.7870437 | 0.00961 | 0.6396544 | 0.0151 | 0.650895 | 0.0157 |
| #40 | DOCK10 | 0.8201369 | 0.01063223 | 0.8980195 | 0.0177 | 0.8868367 | 0.0135 | 0.7803097 | 0.00658 | 0.7153818 | 0.00475 |
| #41 | RBM19 | 0.447356 | 0.01070123 | 0.5213097 | 0.0195 | 0.2989485 | 0.0155 | 0.4328015 | 0.0000776 | 0.5363645 | 0.00774 |
| #42 | USP48 | 0.553089 | 0.01079481 | 0.6185375 | 0.000984 | 0.5904947 | 0.0339 | 0.4078794 | 0.00394 | 0.5954444 | 0.00433 |
| #43 | RLIM | 0.7270363 | 0.01084289 | 0.8740485 | 0.00148 | 0.8045146 | 0.00933 | 0.5673754 | 0.0109 | 0.6622069 | 0.0216 |
| #44 | LRPPRC | 0.6383047 | 0.01087918 | 0.6335985 | 0.00843 | 0.7017869 | 0.00505 | 0.5088103 | 0.0243 | 0.7090232 | 0.00573 |
| #45 | MT-ND4L | 1.868037 | 0.010897 | 1.638331 | 0.000633 | 1.867839 | 0.0422 | 1.852987 | 0.000116 | 2.11299 | 0.000633 |
| #46 | WARS2 | 0.6741437 | 0.01090896 | 0.7441708 | 0.0109 | 0.6941582 | 0.0115 | 0.4231308 | 0.00975 | 0.8351152 | 0.0115 |
| #47 | UBE3A | 0.5167995 | 0.01097657 | 0.6206397 | 0.00313 | 0.6160818 | 0.00955 | 0.4164574 | 0.0285 | 0.4140192 | 0.00268 |
| #48 | PPFIBP2 | 0.8236225 | 0.01100236 | 0.754735 | 0.021 | 0.876927 | 0.00362 | 0.6576594 | 0.0157 | 1.005169 | 0.00366 |
| #49 | FNBP1 | 0.3456333 | 0.01138707 | 0.3832007 | 0.000163 | 0.3425683 | 0.011 | 0.3013791 | 0.0278 | 0.3553852 | 0.0066 |
| #50 | USP34 | 0.5214733 | 0.01140884 | 0.6612035 | 0.0131 | 0.5935749 | 0.0151 | 0.3313179 | 0.0165 | 0.499797 | 0.0009 |
| #51 | TRIP12 | 0.8692235 | 0.01173119 | 0.975967 | 0.0000714 | 0.8343771 | 0.00861 | 0.847449 | 0.0286 | 0.8191009 | 0.00967 |
| #52 | WBP11 | 0.4826569 | 0.01184201 | 0.5006439 | 0.000165 | 0.4567961 | 0.0137 | 0.4075944 | 0.0000937 | 0.5655929 | 0.0334 |
| #53 | PANK3 | 0.6510844 | 0.01205887 | 0.611931 | 0.00333 | 0.6222692 | 0.00646 | 0.6661351 | 0.0102 | 0.7040022 | 0.0283 |
| #54 | NUP98 | 0.4507565 | 0.012275 | 0.5226792 | 0.0116 | 0.4776946 | 0.0147 | 0.3824258 | 0.00291 | 0.4202263 | 0.0199 |
| #55 | FNDC3A | 0.6675154 | 0.0123169 | 0.8214109 | 0.0178 | 0.5953403 | 0.0276 | 0.6638862 | 0.000882 | 0.5894243 | 0.00298 |
| #56 | CAPN7 | 0.7211366 | 0.01248466 | 0.80385 | 0.00503 | 0.7099316 | 0.000683 | 0.6872133 | 0.031 | 0.6835516 | 0.0132 |

| | | | | | | | | | | |
|---|---|---|---|---|---|---|---|---|---|---|
| #57 | ATP2C1 | 0.755905 | 0.01255465 | 0.8767742 | 0.00878 | 0.709667 | 0.0356 | 0.6741368 | 0.00101 | 0.763042 | 0.00483 |
| #58 | ZMYM6 | 0.8025775 | 0.01260694 | 0.8844613 | 0.00229 | 0.8318789 | 0.019 | 0.6587636 | 0.0205 | 0.8352062 | 0.0086 |
| #59 | FNBP4 | 0.3701415 | 0.01275434 | 0.4563622 | 0.0153 | 0.4533277 | 0.00517 | 0.2375294 | 0.0111 | 0.3333465 | 0.0194 |
| #60 | MRPL43 | -0.4417892 | 0.01282575 | -0.4020261 | 0.017 | -0.4432784 | 0.003 | -0.4953984 | 0.00667 | -0.426454 | 0.0246 |
| #61 | CCNL1 | 0.8516011 | 0.01282703 | 0.8469793 | 0.00214 | 0.8084187 | 0.00118 | 0.750872 | 0.041 | 1.000135 | 0.00698 |
| #62 | UHRF1BP1 | 0.7106231 | 0.01304431 | 0.7090006 | 0.00236 | 0.7180429 | 0.0205 | 0.5970102 | 0.0131 | 0.8184388 | 0.0161 |
| #63 | PRKDC | 0.4188863 | 0.0130492 | 0.4444733 | 0.00832 | 0.4672059 | 0.00896 | 0.3312565 | 0.023 | 0.4326096 | 0.012 |
| #64 | YTHDC2 | 0.6754016 | 0.01310857 | 0.8273874 | 0.00991 | 0.6470856 | 0.022 | 0.6019688 | 0.00952 | 0.6251648 | 0.011 |
| #65 | DNAJC10 | 0.7416302 | 0.01311702 | 0.75377 | 0.00164 | 0.7325407 | 0.0321 | 0.624758 | 0.0184 | 0.8554519 | 0.000277 |
| #66 | TBC1D5 | 0.4486398 | 0.01324915 | 0.4402632 | 0.00727 | 0.4379669 | 0.00198 | 0.3454673 | 0.0321 | 0.5708618 | 0.0117 |
| #67 | PAPOLA | 0.3458716 | 0.01341913 | 0.3568701 | 0.00515 | 0.3431718 | 0.0227 | 0.2993531 | 0.0117 | 0.3840914 | 0.0141 |
| #68 | MED14 | 0.8559488 | 0.01349145 | 0.9522348 | 0.00265 | 0.9546194 | 0.00797 | 0.7560415 | 0.0349 | 0.7608994 | 0.00842 |
| #69 | ZDHHC17 | 0.5678104 | 0.01358298 | 0.7218499 | 0.00349 | 0.5806541 | 0.0143 | 0.4516988 | 0.0314 | 0.5170389 | 0.00521 |
| #70 | BTAF1 | 0.7534817 | 0.01365572 | 0.9656293 | 0.0108 | 0.7833148 | 0.014 | 0.5735481 | 0.0235 | 0.6914345 | 0.00632 |
| #71 | RBAK | 0.6459604 | 0.01366249 | 0.6312414 | 0.00295 | 0.6221202 | 0.0144 | 0.5087727 | 0.00647 | 0.8217071 | 0.0308 |
| #72 | NAB1 | 0.5195494 | 0.01367225 | 0.5073632 | 0.00548 | 0.5140963 | 0.00162 | 0.4901136 | 0.0235 | 0.5666248 | 0.0241 |
| #73 | CHD8 | 0.3932808 | 0.01377443 | 0.4694883 | 0.0168 | 0.3919865 | 0.0202 | 0.2736985 | 0.0129 | 0.4379499 | 0.00514 |
| #74 | EIF2S3 | 0.5579646 | 0.01387024 | 0.56661 | 0.00276 | 0.6184527 | 0.00131 | 0.451571 | 0.0494 | 0.5952245 | 0.00201 |
| #75 | MATR3.1 | 0.8686266 | 0.01405783 | 1.06078 | 0.0154 | 0.8743681 | 0.0038 | 0.598406 | 0.0363 | 0.9409524 | 0.000714 |
| #76 | IDE | 0.648105 | 0.01423029 | 0.702633 | 0.00653 | 0.7899036 | 0.00774 | 0.5784299 | 0.0232 | 0.5214535 | 0.0195 |
| #77 | USP33 | 0.5158085 | 0.01434634 | 0.609518 | 0.00214 | 0.4013147 | 0.0402 | 0.5263004 | 0.00205 | 0.5261008 | 0.013 |
| #78 | SNX13 | 0.6134711 | 0.01441162 | 0.7080192 | 0.013 | 0.530022 | 0.00127 | 0.5033151 | 0.0221 | 0.7125283 | 0.0213 |
| #79 | NDUFS5 | -0.5279851 | 0.01458556 | -0.4665346 | 0.0125 | -0.5124729 | 0.0263 | -0.5881778 | 0.0143 | -0.5447553 | 0.00527 |
| #80 | HNRNPA3 | 0.3638518 | 0.01463873 | 0.3949127 | 0.00544 | 0.3111501 | 0.0118 | 0.3837091 | 0.0363 | 0.3656354 | 0.00504 |
| #81 | KDM5C | 0.7535294 | 0.01468136 | 0.8909333 | 0.0229 | 0.6375173 | 0.00558 | 0.6087169 | 0.000999 | 0.8769503 | 0.0293 |
| #82 | WAPL | 0.6360418 | 0.01473879 | 0.6972819 | 0.0203 | 0.7194014 | 0.0254 | 0.5527676 | 0.00963 | 0.5747164 | 0.00371 |
| #83 | ZC3H7A | 0.7088086 | 0.01496796 | 0.6398616 | 0.000929 | 0.8381172 | 0.0105 | 0.6867792 | 0.0478 | 0.6704762 | 0.000627 |
| #84 | OTUD6B | 0.5052961 | 0.01503741 | 0.7322139 | 0.00154 | 0.4484557 | 0.0267 | 0.3763925 | 0.00678 | 0.4641224 | 0.0251 |
| #85 | ZNF117 | 0.8713899 | 0.01514751 | 1.093244 | 0.00149 | 0.8159497 | 0.0136 | 0.9635801 | 0.0152 | 0.6127853 | 0.0303 |
| #86 | EIF1AX | 0.6698175 | 0.01520428 | 0.6746941 | 0.00199 | 0.6637667 | 0.0434 | 0.6505952 | 0.00835 | 0.6902139 | 0.00705 |
| #87 | YEATS2 | 0.7848911 | 0.01554255 | 0.8763928 | 0.00104 | 0.8873521 | 0.00782 | 0.6255584 | 0.0433 | 0.7502611 | 0.00998 |
| #88 | SF3B1 | 0.5547123 | 0.01554798 | 0.6671123 | 0.00511 | 0.5573246 | 0.0225 | 0.4185652 | 0.0111 | 0.5758471 | 0.0235 |
| #89 | ZNF451 | 0.4864665 | 0.01555804 | 0.5638431 | 0.00784 | 0.4859921 | 0.00505 | 0.383578 | 0.0432 | 0.5124529 | 0.00612 |
| #90 | TRPM7 | 0.495697 | 0.01571597 | 0.5906907 | 0.00849 | 0.4632048 | 0.0309 | 0.3756263 | 0.0229 | 0.553266 | 0.000623 |
| #91 | KRTCAP2 | -0.3807723 | 0.01573302 | -0.4439984 | 0.00795 | -0.3723201 | 0.0222 | -0.3122798 | 0.00673 | -0.394491 | 0.026 |
| #92 | ZMYM4 | 0.9234587 | 0.01584298 | 0.9804538 | 0.00461 | 0.9353786 | 0.0466 | 0.9047699 | 0.00528 | 0.8732325 | 0.00692 |
| #93 | ARFIP2 | -0.2814698 | 0.01598153 | -0.3105988 | 0.00298 | -0.303284 | 0.0154 | -0.3039257 | 0.00885 | -0.2080709 | 0.0367 |
| #94 | PRKX | 0.7411333 | 0.01605607 | 0.6712684 | 0.031 | 0.7770656 | 0.000472 | 0.8201213 | 0.0301 | 0.6960777 | 0.00268 |
| #95 | KLHL15 | 0.8013366 | 0.01614865 | 0.7622688 | 0.0306 | 0.9863087 | 0.00988 | 0.469316 | 0.0217 | 0.9874527 | 0.00241 |
| #96 | OPA1 | 0.8748752 | 0.0161619 | 0.8863352 | 0.000163 | 0.9631096 | 0.000487 | 0.7624302 | 0.0236 | 0.8876257 | 0.0404 |
| #97 | GPBP1L1 | 0.2711146 | 0.01620445 | 0.3390233 | 0.0263 | 0.2237517 | 0.00598 | 0.2422943 | 0.0123 | 0.279389 | 0.0202 |
| #98 | STX16 | 0.6681846 | 0.01629309 | 0.758345 | 0.0432 | 0.6318459 | 0.0118 | 0.596435 | 0.00984 | 0.6861124 | 0.000311 |
| #99 | PDS5A | 0.5299098 | 0.01635082 | 0.5158697 | 0.00133 | 0.5360281 | 0.0122 | 0.4818043 | 0.0517 | 0.585937 | 0.000173 |
| #100 | VPS13D | 0.6958257 | 0.01636954 | 0.7608988 | 0.0492 | 0.7070154 | 0.00378 | 0.5039094 | 0.0124 | 0.8114792 | 0.0000942 |
| #101 | SETD2 | 0.8511889 | 0.01670255 | 0.9186778 | 0.00417 | 0.9116269 | 0.0451 | 0.7356191 | 0.00308 | 0.838832 | 0.0144 |
| #102 | HECA | 0.6314139 | 0.01675524 | 0.696051 | 0.00665 | 0.7432294 | 0.0127 | 0.4474161 | 0.0326 | 0.638959 | 0.0151 |
| #103 | MYSM1 | 0.6378324 | 0.01678752 | 0.8290377 | 0.0104 | 0.6449804 | 0.0385 | 0.4132595 | 0.00843 | 0.6640518 | 0.00979 |
| #104 | CDC27 | 0.3073062 | 0.016833 | 0.3113982 | 0.00659 | 0.2887635 | 0.0327 | 0.3181702 | 0.00629 | 0.310893 | 0.0218 |
| #105 | SUCO | 0.8223468 | 0.01699394 | 0.8534 | 0.00361 | 0.8141786 | 0.0375 | 0.7237239 | 0.00692 | 0.8980848 | 0.0199 |
| #106 | ACSL3 | 0.5760641 | 0.01740843 | 0.6552913 | 0.00766 | 0.5002107 | 0.0211 | 0.6339282 | 0.000541 | 0.5148262 | 0.0403 |
| #107 | ZFC3H1 | 0.6557162 | 0.01767093 | 0.8118346 | 0.0103 | 0.6026098 | 0.0447 | 0.5982622 | 0.000536 | 0.6101582 | 0.0151 |
| #108 | ATP11B | 0.658277 | 0.01783823 | 0.6812443 | 0.000713 | 0.7119649 | 0.0121 | 0.5074571 | 0.0509 | 0.7324415 | 0.00767 |
| #109 | CEP350 | 0.5301601 | 0.01786411 | 0.6300366 | 0.0026 | 0.5570756 | 0.0308 | 0.4326078 | 0.0341 | 0.5009204 | 0.00389 |
| #110 | SMARCA5 | 0.516744 | 0.01795177 | 0.6284801 | 0.000273 | 0.5213186 | 0.0391 | 0.4954039 | 0.00284 | 0.4217733 | 0.0296 |
| #111 | CHD1 | 0.8461189 | 0.01806481 | 0.8180662 | 0.00828 | 0.9476504 | 0.00402 | 0.7869678 | 0.0557 | 0.8317911 | 0.00425 |
| #112 | SNX16 | 0.7043125 | 0.01816315 | 0.6388859 | 0.0333 | 0.736123 | 0.0201 | 0.5272873 | 0.00334 | 0.9149538 | 0.0158 |
| #113 | GABPA | 0.6055721 | 0.01829353 | 0.6071413 | 0.0341 | 0.5853736 | 0.0187 | 0.6569768 | 0.00466 | 0.5727968 | 0.0157 |
| #114 | ALMS1 | 0.8673279 | 0.01831984 | 1.106385 | 0.0332 | 0.7877089 | 0.01 | 0.7892492 | 0.0264 | 0.7859685 | 0.00357 |

| | | | | | | | | | | | |
|---|---|---|---|---|---|---|---|---|---|---|---|
| #115 | TAF2 | 0.5506945 | 0.0183282 | 0.7062725 | 0.000787 | 0.4872907 | 0.0208 | 0.4230207 | 0.0227 | 0.5861939 | 0.029 |
| #116 | TUSC2 | -0.4919952 | 0.01845065 | -0.5254242 | 0.0329 | -0.5641273 | 0.00164 | -0.3936551 | 0.0281 | -0.4847743 | 0.0111 |
| #117 | ITPR2 | 0.7788111 | 0.01850549 | 0.844884 | 0.000146 | 0.7625321 | 0.000799 | 0.7672312 | 0.071 | 0.7405972 | 0.00204 |
| #118 | CTDSPL2 | 0.6845389 | 0.018585 | 0.7670645 | 0.0118 | 0.6770916 | 0.0281 | 0.5195008 | 0.00668 | 0.7744987 | 0.0277 |
| #119 | TUBGCP3 | 0.558149 | 0.01868453 | 0.5870949 | 0.0174 | 0.5903233 | 0.000617 | 0.3556355 | 0.0388 | 0.6995425 | 0.018 |
| #120 | LIG3 | 0.4282826 | 0.0187565 | 0.608687 | 0.00487 | 0.3463698 | 0.00378 | 0.3927885 | 0.0425 | 0.3652852 | 0.0238 |
| #121 | PTPN1 | 0.5382643 | 0.01883538 | 0.4790373 | 0.0311 | 0.5212336 | 0.0101 | 0.740052 | 0.0104 | 0.4127341 | 0.0238 |
| #122 | BRWD3 | 0.6221287 | 0.01892178 | 0.7500026 | 0.0352 | 0.4428976 | 0.0116 | 0.5782713 | 0.0151 | 0.7173432 | 0.0138 |
| #123 | WDR75 | 0.667785 | 0.01895864 | 0.8557241 | 0.0418 | 0.6015889 | 0.0154 | 0.5662827 | 0.00531 | 0.6475443 | 0.0133 |
| #124 | CPTP | -0.9741671 | 0.01907458 | -0.8946526 | 0.000999 | -0.8207664 | 0.0302 | -1.170923 | 0.0387 | -1.010327 | 0.00642 |
| #125 | AP3B1 | 0.6210766 | 0.01924941 | 0.7105244 | 0.000599 | 0.5269373 | 0.0181 | 0.6216949 | 0.0469 | 0.6251498 | 0.0114 |
| #126 | MAN1A2 | 0.3491699 | 0.0192885 | 0.4072909 | 0.0185 | 0.3698008 | 0.0206 | 0.299681 | 0.0354 | 0.3199071 | 0.00272 |
| #127 | AP1G1 | 0.6537377 | 0.01929246 | 0.6995882 | 0.00401 | 0.6988669 | 0.000414 | 0.5003327 | 0.0306 | 0.7161629 | 0.0421 |
| #128 | CHML | 0.8867125 | 0.01931934 | 0.8079463 | 0.00719 | 0.9270412 | 0.0105 | 0.9600441 | 0.0414 | 0.8518183 | 0.0182 |
| #129 | MON2 | 0.6416803 | 0.01939319 | 0.7458155 | 0.0434 | 0.6013913 | 0.0313 | 0.4915771 | 0.00228 | 0.7279373 | 0.000547 |
| #130 | IPO7 | 0.5826069 | 0.01947336 | 0.5609661 | 0.00636 | 0.6432621 | 0.0125 | 0.4590268 | 0.0453 | 0.6671725 | 0.0138 |
| #131 | NMD3 | 0.265665 | 0.01954132 | 0.3445422 | 0.00675 | 0.2709826 | 0.00285 | 0.200841 | 0.0306 | 0.2462943 | 0.038 |
| #132 | WDR7 | 0.5085725 | 0.01959914 | 0.5458223 | 0.0419 | 0.5196218 | 0.0106 | 0.446082 | 0.0234 | 0.5227638 | 0.00251 |
| #133 | SPEN | 0.6183386 | 0.01964384 | 0.6649626 | 0.00488 | 0.6845151 | 0.0183 | 0.506523 | 0.0467 | 0.6173537 | 0.00871 |
| #134 | RNF38 | 0.4411125 | 0.02007223 | 0.3820915 | 0.00135 | 0.4834605 | 0.000208 | 0.4028643 | 0.0737 | 0.4960338 | 0.00502 |
| #135 | GFM1 | 0.3754029 | 0.02021023 | 0.4491449 | 0.0151 | 0.4206174 | 0.00833 | 0.3594902 | 0.00833 | 0.2723592 | 0.0491 |
| #136 | MCPH1 | 0.5702376 | 0.02025863 | 0.8126542 | 0.00419 | 0.5885503 | 0.0277 | 0.3347124 | 0.0432 | 0.5450335 | 0.00593 |
| #137 | USF3 | 0.6556665 | 0.02029184 | 0.6221618 | 0.0018 | 0.6494551 | 0.00553 | 0.6962684 | 0.00678 | 0.6547807 | 0.0671 |
| #138 | SHLD2 | 0.6033548 | 0.02034644 | 0.7338864 | 0.00196 | 0.5846383 | 0.00402 | 0.5895705 | 0.0304 | 0.5053239 | 0.045 |
| #139 | DDHD1 | 0.6267102 | 0.02035695 | 0.6024131 | 0.0534 | 0.7473648 | 0.00104 | 0.5807339 | 0.00269 | 0.5763291 | 0.0243 |
| #140 | SYNCRIP | 0.396365 | 0.02040869 | 0.3970761 | 0.0064 | 0.4088271 | 0.00554 | 0.385924 | 0.0638 | 0.3936328 | 0.00593 |
| #141 | SETX | 0.7024399 | 0.0205138 | 0.7871204 | 0.00802 | 0.5589172 | 0.043 | 0.8027826 | 0.0249 | 0.6609396 | 0.00616 |
| #142 | DMXL1 | 0.6869961 | 0.02054669 | 0.7321547 | 0.00137 | 0.6464312 | 0.0352 | 0.7319516 | 0.0367 | 0.637447 | 0.009 |
| #143 | SCAMP3 | -0.3619187 | 0.02077334 | -0.4860863 | 0.00556 | -0.3040643 | 0.00617 | -0.4277911 | 0.0712 | -0.2297333 | 0.000157 |
| #144 | RAB3GAP1 | 0.7274323 | 0.02078978 | 0.7149858 | 0.00778 | 0.5917215 | 0.0637 | 0.8674513 | 0.0106 | 0.7355707 | 0.00112 |
| #145 | ABI1 | 0.5140332 | 0.02080265 | 0.6392417 | 0.00859 | 0.525783 | 0.0195 | 0.3578163 | 0.0458 | 0.5332917 | 0.00934 |
| #146 | U2SURP | 0.2720151 | 0.0208324 | 0.3101194 | 0.000872 | 0.219295 | 0.0473 | 0.2460964 | 0.0219 | 0.3125498 | 0.0132 |
| #147 | PRRC2C | 0.3453424 | 0.0208894 | 0.4711445 | 0.0728 | 0.3158282 | 0.000983 | 0.2938027 | 0.00644 | 0.3005944 | 0.00336 |
| #148 | TMEM131L | 0.7487734 | 0.02092189 | 0.8928196 | 0.0147 | 0.600189 | 0.0584 | 0.812841 | 0.00899 | 0.6892441 | 0.00163 |
| #149 | PIK3CB | 0.6493721 | 0.02094217 | 0.5983619 | 0.00316 | 0.6860979 | 0.00833 | 0.5433112 | 0.063 | 0.7697173 | 0.00923 |
| #150 | ILF3 | 0.4864504 | 0.02097095 | 0.5814594 | 0.0435 | 0.4933347 | 0.00935 | 0.3817761 | 0.0237 | 0.4892315 | 0.00734 |
| #151 | POLA1 | 0.7599782 | 0.02097965 | 0.9480376 | 0.00523 | 0.7593696 | 0.0178 | 0.5104772 | 0.0371 | 0.8220284 | 0.0238 |
| #152 | PRPF4B | 0.6802223 | 0.02104174 | 0.8769685 | 0.0056 | 0.6766335 | 0.0334 | 0.4686553 | 0.0123 | 0.6986319 | 0.0328 |
| #153 | FER | 0.9470931 | 0.02115148 | 1.037789 | 0.0276 | 0.9620843 | 0.046 | 0.7554652 | 0.00957 | 1.033034 | 0.00149 |
| #154 | LCORL | 0.6077586 | 0.02117285 | 0.6580439 | 0.00321 | 0.5942808 | 0.051 | 0.5731708 | 0.000856 | 0.6055388 | 0.0297 |
| #155 | LATS1 | 0.4495929 | 0.02122295 | 0.4414809 | 0.0173 | 0.4572421 | 0.0266 | 0.3553209 | 0.0276 | 0.5443279 | 0.0134 |
| #156 | TSIX | 5.247547 | 0.02135783 | 5.200455 | 0.0205 | 5.151108 | 0.0366 | 5.554743 | 0.0279 | 5.083883 | 0.000407 |
| #157 | SGF29 | -0.5097298 | 0.02142746 | -0.5035125 | 0.00567 | -0.4704227 | 0.00767 | -0.6027896 | 0.0695 | -0.4621945 | 0.00283 |
| #158 | RELCH | 0.6440084 | 0.02146322 | 0.7573743 | 0.000559 | 0.5589165 | 0.0639 | 0.5755867 | 0.0176 | 0.6841563 | 0.00385 |
| #159 | DPP8 | 0.3666165 | 0.02147967 | 0.4417705 | 0.00378 | 0.3751925 | 0.00855 | 0.2132969 | 0.048 | 0.4362062 | 0.0256 |
| #160 | ALG13 | 0.8163902 | 0.02172828 | 0.9684436 | 0.0226 | 0.7678612 | 0.0264 | 0.726079 | 0.0264 | 0.8031771 | 0.0114 |
| #161 | CEP170 | 0.6645538 | 0.02188997 | 0.6826254 | 0.00207 | 0.787021 | 0.00349 | 0.5240806 | 0.0809 | 0.6644881 | 0.00107 |
| #162 | CDC16 | 0.5460657 | 0.02198659 | 0.652484 | 0.0000496 | 0.5544387 | 0.0194 | 0.4772477 | 0.0137 | 0.5000923 | 0.0548 |
| #163 | SGMS1 | 0.7421105 | 0.02225474 | 0.8814422 | 0.000036 | 0.6652851 | 0.00465 | 0.7075249 | 0.0744 | 0.7141897 | 0.00995 |
| #164 | MTMR10 | 0.8470828 | 0.02226783 | 0.8823407 | 0.000337 | 0.8418138 | 0.00108 | 0.7920524 | 0.0827 | 0.8721243 | 0.00493 |
| #165 | NSD1 | 0.4760705 | 0.02228235 | 0.5222849 | 0.000486 | 0.5255214 | 0.0371 | 0.3896771 | 0.0167 | 0.4667988 | 0.0349 |
| #166 | RCOR3 | 0.4342799 | 0.02245603 | 0.5036879 | 0.0117 | 0.4582633 | 0.0525 | 0.3181056 | 0.019 | 0.4570629 | 0.00658 |
| #167 | TMEM62 | 1.014216 | 0.02251626 | 0.9710025 | 0.0139 | 1.267628 | 0.043 | 0.953365 | 0.0165 | 0.8648674 | 0.0167 |
| #168 | CCNT2 | 0.4551685 | 0.02253611 | 0.4998422 | 0.0313 | 0.4001487 | 0.0133 | 0.3242728 | 0.000405 | 0.5964105 | 0.0452 |
| #169 | MIER1 | 0.2636689 | 0.02268803 | 0.2895516 | 0.00215 | 0.2328972 | 0.00132 | 0.2773624 | 0.0435 | 0.2548644 | 0.0438 |
| #170 | PPP1R15B | 0.6773049 | 0.02296697 | 0.6828982 | 0.00197 | 0.731878 | 0.00409 | 0.5495091 | 0.0719 | 0.7449342 | 0.014 |
| #171 | ICE1 | 0.7138069 | 0.02300454 | 0.5380493 | 0.0158 | 0.9075374 | 0.0346 | 0.8374472 | 0.0262 | 0.572194 | 0.0155 |
| #172 | HBP1 | 0.4211408 | 0.02304514 | 0.4275564 | 0.0251 | 0.3773261 | 0.00747 | 0.3459426 | 0.0295 | 0.5337379 | 0.0301 |

| | | | | | | | | | | | |
|---|---|---|---|---|---|---|---|---|---|---|---|
| #173 | MIOS | 0.5826493 | 0.02327252 | 0.369503 | 0.0017 | 0.8245707 | 0.0237 | 0.6309804 | 0.0221 | 0.5055432 | 0.0456 |
| #174 | PIKFYVE | 0.6108119 | 0.02329231 | 0.6763375 | 0.00956 | 0.5902746 | 0.0361 | 0.5476967 | 0.0333 | 0.6289389 | 0.0142 |
| #175 | VPS13C | 1.021289 | 0.02330887 | 0.9818775 | 0.0114 | 0.892187 | 0.00606 | 1.233453 | 0.0649 | 0.9776382 | 0.0109 |
| #176 | PHC3 | 0.5765899 | 0.02346409 | 0.6427994 | 0.0487 | 0.5493428 | 0.0105 | 0.4622513 | 0.0179 | 0.6519662 | 0.0167 |
| #177 | RBM25 | 0.3580878 | 0.02351167 | 0.3742838 | 0.0435 | 0.3365075 | 0.0368 | 0.3912839 | 0.00208 | 0.3302761 | 0.0117 |
| #178 | CAPRIN1 | 0.6730285 | 0.02358856 | 0.6805895 | 0.0111 | 0.7034466 | 0.0112 | 0.6323386 | 0.0628 | 0.6757391 | 0.00928 |
| #179 | UHRF2 | 0.8320118 | 0.02362896 | 0.8475549 | 0.00693 | 0.8789819 | 0.0171 | 0.7882751 | 0.05 | 0.8132354 | 0.0206 |
| #180 | SLC30A9 | 0.4656347 | 0.02363406 | 0.5523386 | 0.00291 | 0.47214 | 0.0317 | 0.3997107 | 0.019 | 0.4383497 | 0.0409 |
| #181 | UBR3 | 0.6055435 | 0.02376228 | 0.603676 | 0.00773 | 0.6954831 | 0.00176 | 0.383649 | 0.0843 | 0.7393657 | 0.00124 |
| #182 | TUBGCP2 | 0.5680013 | 0.02376741 | 0.5817134 | 0.00375 | 0.623418 | 0.0302 | 0.4875143 | 0.0521 | 0.5793596 | 0.00896 |
| #183 | PLEKHA8 | 0.7207649 | 0.02386383 | 0.6379107 | 0.0237 | 0.7670831 | 0.0196 | 0.6718285 | 0.0232 | 0.8062373 | 0.0289 |
| #184 | MTM1 | 0.7329349 | 0.02389362 | 0.6870407 | 0.0142 | 0.6300958 | 0.0456 | 0.8204311 | 0.0358 | 0.7941719 | 0.0000378 |
| #185 | NRDC | 0.525546 | 0.02402478 | 0.5803185 | 0.0026 | 0.4562498 | 0.0232 | 0.5212819 | 0.0629 | 0.544334 | 0.00733 |
| #186 | ATP9B | 1.143066 | 0.02407479 | 1.32517 | 0.0438 | 1.118482 | 0.0103 | 0.9962045 | 0.0336 | 1.132405 | 0.00863 |
| #187 | PDS5B | 0.4152854 | 0.02422463 | 0.5404398 | 0.00925 | 0.3804232 | 0.00586 | 0.3116915 | 0.0728 | 0.4285872 | 0.00898 |
| #188 | OSBPL9 | 0.2942891 | 0.02424273 | 0.4626863 | 0.000548 | 0.3355559 | 0.00362 | 0.1161216 | 0.067 | 0.2627925 | 0.0258 |
| #189 | CD46 | 0.5734433 | 0.02429332 | 0.5818517 | 0.00196 | 0.4753759 | 0.00944 | 0.7098829 | 0.0743 | 0.5266629 | 0.0115 |
| #190 | KAT6A | 0.4945804 | 0.02433219 | 0.6298668 | 0.0404 | 0.5392921 | 0.0165 | 0.3322362 | 0.0289 | 0.4769266 | 0.0115 |
| #191 | NOP14 | 0.3084959 | 0.02453303 | 0.4302316 | 0.00781 | 0.2462012 | 0.0254 | 0.2511381 | 0.0444 | 0.3064129 | 0.0205 |
| #192 | SNU13 | -0.294274 | 0.02473746 | -0.3342294 | 0.0172 | -0.229118 | 0.0249 | -0.3712664 | 0.00384 | -0.2424822 | 0.0531 |
| #193 | PPIP5K2 | 0.5773136 | 0.0247689 | 0.724844 | 0.0076 | 0.6142306 | 0.0286 | 0.3933032 | 0.0338 | 0.5768766 | 0.0291 |
| #194 | CUL4B | 0.6303268 | 0.02501426 | 0.6847922 | 0.0129 | 0.6379533 | 0.081 | 0.573195 | 0.002 | 0.6253669 | 0.00414 |
| #195 | COPS2 | 0.715069 | 0.02503062 | 0.7093625 | 0.00171 | 0.7361448 | 0.0408 | 0.6777216 | 0.00582 | 0.7370471 | 0.0518 |
| #196 | SPTLC1 | 0.3166831 | 0.02503104 | 0.3289671 | 0.0055 | 0.2963047 | 0.00525 | 0.3090842 | 0.0566 | 0.3323764 | 0.0327 |
| #197 | SEH1L | 0.4910649 | 0.02506461 | 0.4895629 | 0.0137 | 0.4317208 | 0.00786 | 0.601391 | 0.0617 | 0.4415848 | 0.017 |
| #198 | DENND4A | 0.6046004 | 0.02506606 | 0.72155 | 0.00139 | 0.4866443 | 0.0673 | 0.5421064 | 0.0288 | 0.6681009 | 0.00281 |
| #199 | RNF111 | 0.5682621 | 0.02509776 | 0.6673992 | 0.0132 | 0.585864 | 0.0308 | 0.388754 | 0.0469 | 0.631031 | 0.00954 |
| #200 | MALT1 | 0.7500634 | 0.02513138 | 0.5645551 | 0.0157 | 0.9256925 | 0.0322 | 0.8523861 | 0.0211 | 0.6576199 | 0.0314 |
| #201 | PIK3CA | 0.6175206 | 0.02519881 | 0.5897897 | 0.00453 | 0.6226453 | 0.00648 | 0.6110099 | 0.0507 | 0.6466375 | 0.0391 |
| #202 | POLR3C | 0.3218698 | 0.02524512 | 0.3409404 | 0.0118 | 0.3053098 | 0.0135 | 0.2901305 | 0.06 | 0.3510985 | 0.0157 |
| #203 | SLC25A46 | 0.3124853 | 0.02526875 | 0.3155341 | 0.0283 | 0.2917739 | 0.0533 | 0.2619796 | 0.0132 | 0.3806538 | 0.00628 |
| #204 | HNRNPH3 | 0.2908884 | 0.02528394 | 0.2710598 | 0.0267 | 0.264524 | 0.00168 | 0.2503353 | 0.0667 | 0.3776346 | 0.00599 |
| #205 | SENP2 | 0.5023893 | 0.02536373 | 0.577893 | 0.0106 | 0.4877924 | 0.0193 | 0.3351778 | 0.0298 | 0.6086942 | 0.0418 |
| #206 | NF1 | 0.4528618 | 0.02540955 | 0.37851 | 0.012 | 0.5387584 | 0.00484 | 0.4800335 | 0.0602 | 0.4141454 | 0.0247 |
| #207 | EPC2 | 0.6808109 | 0.02542051 | 0.7802732 | 0.00217 | 0.5794354 | 0.0425 | 0.7005845 | 0.0127 | 0.6629506 | 0.0443 |
| #208 | FCHSD2 | 1.109325 | 0.02555074 | 1.053886 | 0.0012 | 1.11832 | 0.0392 | 1.226241 | 0.0485 | 1.038854 | 0.0133 |
| #209 | ACIN1 | 0.5736003 | 0.02558742 | 0.7756571 | 0.0154 | 0.5428075 | 0.0754 | 0.3697392 | 0.0112 | 0.6061974 | 0.000313 |
| #210 | SLC22A18 | -0.6879001 | 0.02568515 | -0.6263377 | 0.0119 | -0.5954149 | 0.00226 | -0.9354732 | 0.0566 | -0.5943747 | 0.032 |
| #211 | TAB2 | 0.4218307 | 0.02570331 | 0.3611994 | 0.0414 | 0.5631329 | 0.00773 | 0.3431143 | 0.0246 | 0.4198761 | 0.0291 |
| #212 | UBQLN1 | 0.5352445 | 0.02581986 | 0.5692734 | 0.00291 | 0.5485612 | 0.00763 | 0.4861235 | 0.0812 | 0.5370197 | 0.0116 |
| #213 | RPS6KB1 | 0.4857096 | 0.02592395 | 0.5081477 | 0.0399 | 0.4294522 | 0.0332 | 0.4414241 | 0.00853 | 0.5638143 | 0.022 |
| #214 | NUS1 | 0.7018698 | 0.02593742 | 0.5529621 | 0.00391 | 0.7960768 | 0.00214 | 0.7362043 | 0.0965 | 0.7222361 | 0.00115 |
| #215 | SMARCC1 | 0.5169561 | 0.02598432 | 0.5153882 | 0.0425 | 0.5296323 | 0.0336 | 0.5001839 | 0.00478 | 0.5226202 | 0.023 |
| #216 | CLPP | -0.3993577 | 0.02599002 | -0.4780251 | 0.0248 | -0.3620146 | 0.00538 | -0.3764884 | 0.0693 | -0.3809026 | 0.00457 |
| #217 | TASOR2 | 0.6329015 | 0.02605192 | 0.631269 | 0.0482 | 0.7645366 | 0.00998 | 0.6457726 | 0.00631 | 0.4900278 | 0.0397 |
| #218 | EPM2AIP1 | 0.4954469 | 0.02606615 | 0.4390767 | 0.0563 | 0.3685306 | 0.00544 | 0.5840536 | 0.0163 | 0.5901267 | 0.0262 |
| #219 | MT-ND6 | 0.9038064 | 0.02607742 | 1.031633 | 0.0461 | 0.9154293 | 0.00949 | 0.7781648 | 0.0349 | 0.8899984 | 0.0138 |
| #220 | GOLT1B | 0.3515862 | 0.02614196 | 0.3665895 | 0.0175 | 0.2933546 | 0.0417 | 0.4262404 | 0.0127 | 0.3201604 | 0.0326 |
| #221 | TCF12 | 0.6383558 | 0.02617734 | 0.6832444 | 0.00809 | 0.6810654 | 0.0327 | 0.5120208 | 0.0397 | 0.6770927 | 0.0241 |
| #222 | GPATCH2L | 0.399477 | 0.02625349 | 0.451888 | 0.037 | 0.3559071 | 0.0194 | 0.2720934 | 0.0386 | 0.5180194 | 0.01 |
| #223 | PRMT3 | 0.6825043 | 0.0263854 | 0.875544 | 0.0104 | 0.6103427 | 0.0529 | 0.6685528 | 0.029 | 0.5755775 | 0.0132 |
| #224 | PEX16 | -0.3061473 | 0.02647551 | -0.2805797 | 0.00503 | -0.3568118 | 0.0116 | -0.2770916 | 0.0238 | -0.310106 | 0.0654 |
| #225 | PHF3 | 0.6345278 | 0.02654901 | 0.7602677 | 0.0206 | 0.6825192 | 0.0281 | 0.4353745 | 0.05 | 0.6599498 | 0.00749 |
| #226 | DISC1 | 1.192481 | 0.02655677 | 1.27264 | 0.0545 | 1.169633 | 0.00276 | 1.036333 | 0.0331 | 1.291316 | 0.0158 |
| #227 | RIF1 | 0.5968663 | 0.0265738 | 0.7258232 | 0.0363 | 0.5313767 | 0.0437 | 0.6237897 | 0.0122 | 0.5064755 | 0.0141 |
| #228 | TNKS2 | 0.6232032 | 0.02667706 | 0.5072648 | 0.0361 | 0.6168254 | 0.00453 | 0.7592082 | 0.0508 | 0.6095141 | 0.0153 |
| #229 | BCLAF1 | 0.4648095 | 0.02672635 | 0.5467144 | 0.00834 | 0.5370776 | 0.022 | 0.2914144 | 0.0654 | 0.4840318 | 0.0111 |
| #230 | TMEM30A | 0.6820243 | 0.02684047 | 0.5008141 | 0.0233 | 0.6546157 | 0.00462 | 0.8696939 | 0.0655 | 0.7029736 | 0.014 |

| | | | | | | | | | | | |
|---|---|---|---|---|---|---|---|---|---|---|---|
| **#231** | ARAP2 | 0.6781082 | 0.02684079 | 0.7873606 | 0.0202 | 0.6499812 | 0.0102 | 0.6559431 | 0.0515 | 0.619148 | 0.0254 |
| **#232** | LMBR1 | 0.8159947 | 0.02703605 | 0.7388551 | 0.0584 | 0.7507863 | 0.0177 | 0.8748264 | 0.0273 | 0.899511 | 0.00477 |
| **#233** | CERS6 | 0.7220227 | 0.02708261 | 0.6232767 | 0.0085 | 0.7913345 | 0.0035 | 0.82095 | 0.0137 | 0.6525296 | 0.0827 |
| **#234** | FRYL | 0.4509073 | 0.02715675 | 0.6537279 | 0.0124 | 0.4504611 | 0.0139 | 0.2662204 | 0.046 | 0.4332199 | 0.0363 |
| **#235** | CEP192 | 0.7131693 | 0.02729493 | 0.9164939 | 0.0397 | 0.6835834 | 0.0256 | 0.5117406 | 0.0131 | 0.7408592 | 0.0307 |
| **#236** | SFPQ | 0.5953003 | 0.02734275 | 0.5541039 | 0.0259 | 0.7158502 | 0.00668 | 0.5587001 | 0.058 | 0.552547 | 0.0188 |
| **#237** | NCBP1 | 0.4964538 | 0.02739226 | 0.5394582 | 0.0251 | 0.5040608 | 0.053 | 0.3918577 | 0.0109 | 0.5504387 | 0.0206 |
| **#238** | TMEM173 | -0.5670976 | 0.02741323 | -0.5611807 | 0.0485 | -0.4352656 | 0.0452 | -0.7409383 | 0.00687 | -0.5310058 | 0.0091 |
| **#239** | ERCC6L2 | 0.5956989 | 0.02744191 | 0.6870968 | 0.0196 | 0.590957 | 0.0731 | 0.5039666 | 0.0107 | 0.6007751 | 0.00633 |
| **#240** | CUL5 | 0.3117953 | 0.02754361 | 0.2968418 | 0.0158 | 0.3412877 | 0.00412 | 0.2994058 | 0.0222 | 0.309646 | 0.068 |
| **#241** | MAP4 | 0.5268429 | 0.02762883 | 0.6000099 | 0.0156 | 0.5061198 | 0.0157 | 0.4578643 | 0.0647 | 0.5433775 | 0.0146 |
| **#242** | MORC3 | 0.6255566 | 0.02763045 | 0.7554513 | 0.0266 | 0.5667992 | 0.0459 | 0.6018214 | 0.013 | 0.5781543 | 0.025 |
| **#243** | ANGEL2 | 0.4927349 | 0.02764182 | 0.6935815 | 0.00519 | 0.4925615 | 0.0422 | 0.347816 | 0.0285 | 0.4369808 | 0.0346 |
| **#244** | SCAF8 | 0.4586621 | 0.02782079 | 0.5057117 | 0.0184 | 0.5754151 | 0.0242 | 0.3035503 | 0.0415 | 0.4499711 | 0.0272 |
| **#245** | MAF | -0.7945272 | 0.0278305 | -0.7725582 | 0.00121 | -0.793679 | 0.0272 | -0.9342471 | 0.0477 | -0.6776244 | 0.0352 |
| **#246** | RFX3 | 0.6268749 | 0.02789704 | 0.7592827 | 0.047 | 0.6990203 | 0.0194 | 0.4755021 | 0.0348 | 0.5736944 | 0.0105 |
| **#247** | NGLY1 | 0.3844939 | 0.02791102 | 0.5168044 | 0.0128 | 0.3605638 | 0.0612 | 0.3935402 | 0.0117 | 0.2670672 | 0.026 |
| **#248** | KANSL1L | 0.9470033 | 0.02797552 | 1.05595 | 0.00429 | 0.7796074 | 0.0779 | 1.205189 | 0.00625 | 0.7472675 | 0.0234 |
| **#249** | UBE2E2 | 0.7356972 | 0.02801202 | 0.6704551 | 0.0286 | 0.7695296 | 0.0194 | 0.8022536 | 0.0467 | 0.7005504 | 0.0174 |
| **#250** | HUWE1 | 0.3685634 | 0.02808086 | 0.3989114 | 0.0233 | 0.3391833 | 0.078 | 0.331972 | 0.00232 | 0.4041869 | 0.00866 |
| **#251** | SETDB1 | 0.4075693 | 0.02820239 | 0.4585725 | 0.00561 | 0.2947438 | 0.0528 | 0.3821792 | 0.0381 | 0.4947816 | 0.0163 |
| **#252** | XRN2 | 0.308068 | 0.02830418 | 0.3244549 | 0.0304 | 0.2223943 | 0.000566 | 0.3815491 | 0.0226 | 0.3038737 | 0.0597 |
| **#253** | SMG1 | 0.5431062 | 0.02855091 | 0.6143029 | 0.077 | 0.6013189 | 0.000609 | 0.3798421 | 0.00936 | 0.5769608 | 0.0273 |
| **#254** | SMC5 | 0.5522359 | 0.02862668 | 0.6523111 | 0.00014 | 0.5448782 | 0.0641 | 0.377473 | 0.0345 | 0.6342811 | 0.0157 |
| **#255** | FBXO11 | 0.4939753 | 0.02865818 | 0.4209066 | 0.0059 | 0.5046767 | 0.0195 | 0.4330162 | 0.0688 | 0.6173019 | 0.0204 |
| **#256** | PRPF8 | 0.320345 | 0.02869708 | 0.2659535 | 0.0609 | 0.2741193 | 0.0159 | 0.2973421 | 0.0293 | 0.443965 | 0.00864 |
| **#257** | PTBP2 | 0.869162 | 0.02889074 | 0.9912239 | 0.027 | 0.8292225 | 0.0168 | 0.7588235 | 0.0353 | 0.8973779 | 0.0364 |
| **#258** | NAA80 | -0.6652194 | 0.02893946 | -0.5260413 | 0.0232 | -0.6046055 | 0.0508 | -0.9366316 | 0.0325 | -0.5935993 | 0.0092 |
| **#259** | GLYR1 | 0.4206958 | 0.02895313 | 0.4227714 | 0.00149 | 0.3725767 | 0.0401 | 0.4104632 | 0.0601 | 0.4769718 | 0.0141 |
| **#260** | STOML2 | -0.2794448 | 0.02897672 | -0.3061128 | 0.0194 | -0.2577356 | 0.0393 | -0.2832321 | 0.0562 | -0.2706986 | 0.00106 |
| **#261** | FBXL3 | 0.5498109 | 0.02898204 | 0.5516474 | 0.00291 | 0.5274993 | 0.0415 | 0.5731124 | 0.00421 | 0.5469843 | 0.0673 |
| **#262** | MIER3 | 0.598011 | 0.02898528 | 0.6376915 | 0.0201 | 0.7013495 | 0.0121 | 0.5307647 | 0.00693 | 0.5222381 | 0.0768 |
| **#263** | GPSM3 | -0.407375 | 0.0291275 | -0.4004785 | 0.00734 | -0.4317477 | 0.0254 | -0.3719522 | 0.0812 | -0.4253218 | 0.00256 |
| **#264** | WDR36 | 0.569359 | 0.02916169 | 0.6399694 | 0.0237 | 0.5749771 | 0.0368 | 0.5770127 | 0.016 | 0.4854768 | 0.0402 |
| **#265** | SLC35F5 | 0.421008 | 0.02943014 | 0.5027019 | 0.00314 | 0.3388128 | 0.0495 | 0.452591 | 0.0134 | 0.3899261 | 0.0517 |
| **#266** | COQ8B | -0.6651162 | 0.02947716 | -0.4941708 | 0.00208 | -0.6466354 | 0.00171 | -0.7757169 | 0.0228 | -0.7439417 | 0.0914 |
| **#267** | USP24 | 0.6741972 | 0.02954599 | 0.7779578 | 0.0261 | 0.6459355 | 0.0514 | 0.5240864 | 0.025 | 0.7488092 | 0.0157 |
| **#268** | SOCS4 | 0.5935421 | 0.02960778 | 0.8187843 | 0.000991 | 0.5888955 | 0.0196 | 0.4477183 | 0.0365 | 0.5187701 | 0.0613 |
| **#269** | ELF1 | 0.4382442 | 0.02960948 | 0.2543862 | 0.0349 | 0.5837027 | 0.0198 | 0.3984382 | 0.0119 | 0.5164497 | 0.0518 |
| **#270** | TRAF3 | 0.5076969 | 0.02967819 | 0.2860114 | 0.0648 | 0.5136741 | 0.0245 | 0.7944689 | 0.0252 | 0.4366334 | 0.00413 |
| **#271** | PTPN11 | 0.6736694 | 0.02967895 | 0.6600234 | 0.00337 | 0.6562422 | 0.056 | 0.6185883 | 0.0475 | 0.7598236 | 0.0118 |
| **#272** | RPS27L | -0.3383749 | 0.02985265 | -0.3490071 | 0.0104 | -0.3468462 | 0.00454 | -0.3139151 | 0.0432 | -0.3437312 | 0.0613 |
| **#273** | ANXA11 | -0.4499557 | 0.02989107 | -0.4324183 | 0.016 | -0.4721115 | 0.0226 | -0.4992961 | 0.0447 | -0.3959969 | 0.0363 |
| **#274** | MED13L | 0.6520776 | 0.02994255 | 0.6085678 | 0.0135 | 0.6466978 | 0.00905 | 0.5764474 | 0.0767 | 0.7765972 | 0.0205 |
| **#275** | UNC45A | -0.4706879 | 0.02997376 | -0.3354157 | 0.0166 | -0.5344016 | 0.0132 | -0.6446599 | 0.0285 | -0.3682745 | 0.0616 |
| **#276** | IAH1 | -0.4521254 | 0.02998805 | -0.3696271 | 0.015 | -0.4781189 | 0.0089 | -0.419166 | 0.0473 | -0.5415897 | 0.0488 |
| **#277** | PRPF39 | 0.5926062 | 0.03003758 | 0.7797868 | 0.033 | 0.54296 | 0.0143 | 0.3720314 | 0.0587 | 0.6756466 | 0.0142 |
| **#278** | SACS | 0.8194589 | 0.03015527 | 0.8568949 | 0.0661 | 0.9416905 | 0.0239 | 0.7760082 | 0.0295 | 0.7032421 | 0.0012 |
| **#279** | EPS15 | 0.3898574 | 0.03019878 | 0.3346662 | 0.0356 | 0.3931172 | 0.0458 | 0.4782253 | 0.0374 | 0.3534207 | 0.0021 |
| **#280** | TCF20 | 0.4737786 | 0.03024105 | 0.5956424 | 0.0292 | 0.5343256 | 0.0191 | 0.2744558 | 0.069 | 0.4906904 | 0.00366 |
| **#281** | ATAD2B | 0.6111732 | 0.03024234 | 0.7159667 | 0.00754 | 0.6512789 | 0.0236 | 0.4570501 | 0.0856 | 0.620397 | 0.00429 |
| **#282** | EXOSC10 | 0.4871619 | 0.03024997 | 0.597136 | 0.00118 | 0.4945721 | 0.0434 | 0.332982 | 0.0423 | 0.5239575 | 0.0341 |
| **#283** | DHX15 | 0.4602639 | 0.03026595 | 0.5016078 | 0.0221 | 0.4789645 | 0.0152 | 0.3542552 | 0.0716 | 0.5062281 | 0.0122 |
| **#284** | YTHDF3 | 0.6607415 | 0.03032551 | 0.6697123 | 0.0117 | 0.7579046 | 0.0261 | 0.4925013 | 0.0597 | 0.7228478 | 0.0238 |
| **#285** | CPSF6 | 0.4198887 | 0.03038532 | 0.5176199 | 0.00987 | 0.3868946 | 0.0211 | 0.2596751 | 0.0738 | 0.5153651 | 0.0168 |
| **#286** | TTBK2 | 0.4881592 | 0.03042309 | 0.4876712 | 0.0201 | 0.5289439 | 0.00284 | 0.4064087 | 0.076 | 0.5296128 | 0.0228 |
| **#287** | DBNDD2 | -0.7383037 | 0.0304701 | -0.5764515 | 0.00612 | -0.6761549 | 0.109 | -1.133521 | 0.00615 | -0.5670876 | 0.000583 |
| **#288** | DHX37 | 0.5390429 | 0.03052912 | 0.6320075 | 0.00518 | 0.4313969 | 0.0348 | 0.3964721 | 0.0489 | 0.6962949 | 0.0332 |

| | | | | | | | | | | | |
|---|---|---|---|---|---|---|---|---|---|---|---|
| **#289** | CD59 | -0.5976566 | 0.03059466 | -0.7391012 | 0.00796 | -0.4565584 | 0.0327 | -0.6435628 | 0.0595 | -0.5514042 | 0.0222 |
| **#290** | TNKS | 0.6243226 | 0.03069713 | 0.7168432 | 0.0125 | 0.5939187 | 0.072 | 0.5422701 | 0.00188 | 0.6442583 | 0.0364 |
| **#291** | ASH1L | 0.5740471 | 0.03074644 | 0.7034725 | 0.0791 | 0.5924027 | 0.0145 | 0.397713 | 0.0189 | 0.6026003 | 0.0105 |
| **#292** | GOLGB1 | 0.5078378 | 0.03079272 | 0.7169352 | 0.0147 | 0.5929466 | 0.065 | 0.3207053 | 0.0422 | 0.4007641 | 0.00125 |
| **#293** | UTP18 | 0.3719127 | 0.0308379 | 0.4441764 | 0.0561 | 0.2852331 | 0.0416 | 0.3320681 | 0.0252 | 0.4261734 | 0.000437 |
| **#294** | DCTN4 | 0.7358362 | 0.03088499 | 0.8121261 | 0.00911 | 0.8986535 | 0.0139 | 0.4239414 | 0.0881 | 0.8086238 | 0.0124 |
| **#295** | ZNF207 | 0.2853451 | 0.03092996 | 0.2912526 | 0.0188 | 0.2380053 | 0.0324 | 0.2460174 | 0.0157 | 0.366105 | 0.0569 |
| **#296** | TSR1 | 0.3720204 | 0.03101711 | 0.3780421 | 0.0341 | 0.3101834 | 0.00557 | 0.3533125 | 0.0138 | 0.4465436 | 0.0706 |
| **#297** | ZBTB21 | 1.006294 | 0.03106322 | 1.17279 | 0.00467 | 1.262722 | 0.0413 | 0.5961018 | 0.0624 | 0.9935605 | 0.0158 |
| **#298** | IL32 | -1.145894 | 0.03107553 | -1.121147 | 0.000808 | -1.116701 | 0.0333 | -1.197567 | 0.0109 | -1.148161 | 0.0794 |
| **#299** | OTUD4 | 0.6302248 | 0.03118112 | 0.5471915 | 0.000205 | 0.5656435 | 0.0759 | 0.7601138 | 0.0394 | 0.6479505 | 0.0092 |
| **#300** | NBAS | 0.4010873 | 0.03125284 | 0.4291727 | 0.00842 | 0.3155934 | 0.0723 | 0.3292974 | 0.0435 | 0.5302858 | 0.000779 |
| **#301** | THOC1 | 0.4874529 | 0.03125642 | 0.5867171 | 0.0818 | 0.3993028 | 0.0177 | 0.4678515 | 0.02 | 0.4959401 | 0.00553 |
| **#302** | TTC17 | 0.3911461 | 0.03126625 | 0.483063 | 0.016 | 0.4030203 | 0.00721 | 0.2031091 | 0.0856 | 0.475392 | 0.0162 |
| **#303** | CSDE1 | 0.3748055 | 0.0312999 | 0.2970986 | 0.0256 | 0.4038592 | 0.0419 | 0.4152117 | 0.0184 | 0.3830526 | 0.0394 |
| **#304** | SS18L2 | -0.451576 | 0.03132096 | -0.4351908 | 0.0126 | -0.4922899 | 0.0406 | -0.4569024 | 0.0141 | -0.4219209 | 0.058 |
| **#305** | SMAP2 | 0.4218526 | 0.031331 | 0.2889491 | 0.0126 | 0.4328408 | 0.0792 | 0.5094319 | 0.0264 | 0.4561885 | 0.00719 |
| **#306** | PRMT1 | -0.3692439 | 0.0313967 | -0.375543 | 0.00657 | -0.3013 | 0.0558 | -0.4085311 | 0.0199 | -0.3916016 | 0.0434 |
| **#307** | ITGB7 | -0.5361961 | 0.03151605 | -0.5909179 | 0.064 | -0.3776984 | 0.0268 | -0.6284755 | 0.00129 | -0.5476925 | 0.034 |
| **#308** | KDM4C | 0.4278161 | 0.03158083 | 0.5339286 | 0.00618 | 0.4292282 | 0.0255 | 0.2664465 | 0.07 | 0.4816612 | 0.0247 |
| **#309** | SPCS3 | 0.5147675 | 0.03172019 | 0.4633272 | 0.035 | 0.5036144 | 0.0257 | 0.5753686 | 0.0503 | 0.5167599 | 0.0159 |
| **#310** | KMT2C | 0.5781404 | 0.03197894 | 0.6401915 | 0.00113 | 0.6282606 | 0.0294 | 0.3978848 | 0.0915 | 0.6462247 | 0.00592 |
| **#311** | ZMYM2 | 0.5238638 | 0.03200923 | 0.5499781 | 0.0574 | 0.5709131 | 0.0349 | 0.3193807 | 0.0241 | 0.6551835 | 0.0116 |
| **#312** | MYO5A | 0.7786593 | 0.03226531 | 0.7451768 | 0.00147 | 0.8214966 | 0.0329 | 0.6951459 | 0.0796 | 0.8528179 | 0.0151 |
| **#313** | EIF2AK3 | 0.808546 | 0.03239277 | 0.8457129 | 0.0129 | 0.7457244 | 0.0558 | 0.8767648 | 0.0185 | 0.765982 | 0.0423 |
| **#314** | MRPS7 | -0.2618638 | 0.0325172 | -0.2733926 | 0.0142 | -0.3205468 | 0.00216 | -0.1911732 | 0.0977 | -0.2623424 | 0.016 |
| **#315** | NUDT2 | -0.9952737 | 0.03253825 | -1.006188 | 0.039 | -0.8742993 | 0.0241 | -1.092651 | 0.0439 | -1.007957 | 0.0233 |
| **#316** | MEI1 | -0.7651084 | 0.03254171 | -0.6027621 | 0.0585 | -0.7746898 | 0.0195 | -0.9989943 | 0.0186 | -0.6839874 | 0.0335 |
| **#317** | CRYBG3 | 0.8423991 | 0.03256774 | 0.9188775 | 0.0119 | 0.8399031 | 0.0403 | 0.8292765 | 0.0184 | 0.7815392 | 0.0597 |
| **#318** | FRS2 | 0.5956873 | 0.03268586 | 0.6676325 | 0.0489 | 0.5405046 | 0.0134 | 0.5597619 | 0.0204 | 0.6148503 | 0.048 |
| **#319** | PSMD5 | 0.4272523 | 0.03287894 | 0.4747518 | 0.00783 | 0.4550012 | 0.00311 | 0.494159 | 0.0332 | 0.2850972 | 0.0874 |
| **#320** | USPL1 | 0.7558102 | 0.0329468 | 0.6964253 | 0.0106 | 0.7777987 | 0.00259 | 0.6465954 | 0.0826 | 0.9024213 | 0.036 |
| **#321** | BABAM1 | -0.3010109 | 0.03305963 | -0.3423965 | 0.0508 | -0.2840971 | 0.00924 | -0.3475047 | 0.0381 | -0.2300454 | 0.0341 |
| **#322** | EML4 | 0.4206968 | 0.03306904 | 0.488735 | 0.0721 | 0.3723082 | 0.0151 | 0.4394271 | 0.0131 | 0.3823168 | 0.0319 |
| **#323** | SLC36A4 | 0.563915 | 0.03314163 | 0.6076662 | 0.00311 | 0.4257141 | 0.108 | 0.557935 | 0.0162 | 0.6643449 | 0.0056 |
| **#324** | ANAPC4 | 0.3983183 | 0.03317928 | 0.5534506 | 0.0103 | 0.350914 | 0.0779 | 0.2761363 | 0.0208 | 0.4127724 | 0.0237 |
| **#325** | VPS28 | -0.41007 | 0.03349187 | -0.4584575 | 0.00754 | -0.4239036 | 0.0434 | -0.3592525 | 0.06 | -0.3986664 | 0.023 |
| **#326** | TOPBP1 | 0.3359941 | 0.03361602 | 0.3577614 | 0.000616 | 0.2934283 | 0.105 | 0.2384801 | 0.0108 | 0.4543067 | 0.018 |
| **#327** | DIS3 | 0.5518072 | 0.03370744 | 0.6070724 | 0.00128 | 0.4869067 | 0.0418 | 0.5698873 | 0.0185 | 0.5433622 | 0.0733 |
| **#328** | TOR1AIP1 | 0.4631334 | 0.0337077 | 0.5610302 | 0.0391 | 0.4690031 | 0.0228 | 0.3871244 | 0.0371 | 0.4353758 | 0.0359 |
| **#329** | CUEDC2 | -0.358846 | 0.03372241 | -0.4951122 | 0.0252 | -0.3709064 | 0.00331 | -0.2600869 | 0.0991 | -0.3092785 | 0.00735 |
| **#330** | ARHGEF6 | 0.5781713 | 0.03373433 | 0.5585819 | 0.034 | 0.6498316 | 0.0742 | 0.5497746 | 0.00911 | 0.5544972 | 0.0176 |
| **#331** | DMTF1 | 0.6203333 | 0.03377905 | 0.6845182 | 0.0381 | 0.5631929 | 0.0759 | 0.6443019 | 0.0196 | 0.5893203 | 0.00148 |
| **#332** | KIAA1586 | 0.8311998 | 0.03380843 | 0.9452188 | 0.0123 | 0.7123752 | 0.0412 | 0.8994457 | 0.0693 | 0.7677594 | 0.0124 |
| **#333** | NPEPPS | 0.4546868 | 0.03388629 | 0.5256772 | 0.00924 | 0.5150439 | 0.0026 | 0.2673115 | 0.103 | 0.5107146 | 0.0205 |
| **#334** | SEC24B | 0.4424909 | 0.03393417 | 0.4315526 | 0.009 | 0.4694332 | 0.00601 | 0.2944225 | 0.11 | 0.5745552 | 0.0112 |
| **#335** | SAMD8 | 0.6931735 | 0.03411036 | 0.6984782 | 0.00576 | 0.7020304 | 0.05 | 0.6444353 | 0.0182 | 0.7277501 | 0.0625 |
| **#336** | SPECC1L | 0.6415617 | 0.03419134 | 0.8097857 | 0.0076 | 0.7567791 | 0.0663 | 0.3874736 | 0.0282 | 0.6122082 | 0.0347 |
| **#337** | FYTTD1 | 0.4441282 | 0.03427275 | 0.362449 | 0.0304 | 0.4728017 | 0.00647 | 0.5165357 | 0.0177 | 0.4247264 | 0.0826 |
| **#338** | ZKSCAN8 | 0.6057918 | 0.0343108 | 0.5675784 | 0.0438 | 0.6339469 | 0.0132 | 0.6114416 | 0.0506 | 0.6102004 | 0.0297 |
| **#339** | GOLGA3 | 0.4457319 | 0.03441144 | 0.5558343 | 0.0302 | 0.4466787 | 0.0134 | 0.3214087 | 0.0804 | 0.4590059 | 0.0136 |
| **#340** | ZNF407 | 1.089648 | 0.03447123 | 1.169724 | 0.0645 | 1.075853 | 0.0203 | 1.032305 | 0.0391 | 1.080712 | 0.014 |
| **#341** | SENP6 | 0.4752442 | 0.03464989 | 0.5362886 | 0.00536 | 0.4966589 | 0.041 | 0.3900541 | 0.0838 | 0.4779751 | 0.00847 |
| **#342** | ZNF644 | 0.5514002 | 0.03485019 | 0.603544 | 0.0461 | 0.5840816 | 0.0141 | 0.4612062 | 0.00724 | 0.5567691 | 0.072 |
| **#343** | SHISA5 | -0.3387989 | 0.03496671 | -0.4574413 | 0.00111 | -0.4290305 | 0.0409 | -0.1817537 | 0.0436 | -0.2869701 | 0.0543 |
| **#344** | TBC1D1 | 0.5316523 | 0.03500765 | 0.428835 | 0.0651 | 0.4905163 | 0.0297 | 0.6163819 | 0.0267 | 0.590876 | 0.0185 |
| **#345** | ZGPAT | -0.3929436 | 0.03508753 | -0.3344067 | 0.0237 | -0.3584875 | 0.0613 | -0.4447769 | 0.0148 | -0.4341034 | 0.0405 |
| **#346** | POGK | 0.4497076 | 0.03508799 | 0.470883 | 0.00356 | 0.2947204 | 0.0311 | 0.462762 | 0.0815 | 0.5704652 | 0.0241 |

| | | | | | | | | | | | |
|---|---|---|---|---|---|---|---|---|---|---|---|
| #347 | PRR14L | 0.5692572 | 0.03511433 | 0.7202757 | 0.00809 | 0.5814873 | 0.101 | 0.4332288 | 0.0126 | 0.5420369 | 0.0184 |
| #348 | BLOC1S6 | 0.543503 | 0.03522231 | 0.5138843 | 0.0145 | 0.533284 | 0.0294 | 0.57483 | 0.0933 | 0.5520137 | 0.00365 |
| #349 | TAX1BP1 | 0.3668528 | 0.03523959 | 0.3244851 | 0.0775 | 0.3178389 | 0.00529 | 0.4449796 | 0.046 | 0.3801078 | 0.0121 |
| #350 | SREK1 | 0.5193543 | 0.0352566 | 0.5544996 | 0.0465 | 0.4977378 | 0.0565 | 0.4052461 | 0.0135 | 0.6199336 | 0.0245 |
| #351 | MAP4K3 | 0.8134856 | 0.03553388 | 0.7198848 | 0.0101 | 0.8216264 | 0.0125 | 0.8401293 | 0.107 | 0.8723017 | 0.0126 |
| #352 | EIF4G2 | 0.5646715 | 0.03553755 | 0.5661977 | 0.00671 | 0.6522836 | 0.0671 | 0.3976807 | 0.0497 | 0.642524 | 0.0186 |
| #353 | SERINC1 | 0.3754363 | 0.03565744 | 0.3255963 | 0.0138 | 0.4255595 | 0.0348 | 0.3064641 | 0.0938 | 0.4441252 | 0.000152 |
| #354 | TRAK2 | 0.4615341 | 0.03568375 | 0.5022084 | 0.0148 | 0.4760234 | 0.0923 | 0.5431497 | 0.000579 | 0.3247549 | 0.035 |
| #355 | CSGALNACT2 | 0.9913643 | 0.03586611 | 1.029579 | 0.00449 | 1.15677 | 0.0604 | 0.8163311 | 0.0399 | 0.9627772 | 0.0387 |
| #356 | PARG | 0.9467627 | 0.03587519 | 1.255285 | 0.0235 | 0.913432 | 0.0937 | 0.7282288 | 0.004 | 0.8901052 | 0.0223 |
| #357 | ELP4 | 0.5016733 | 0.03587583 | 0.6465427 | 0.00952 | 0.3633467 | 0.0465 | 0.3560933 | 0.086 | 0.6407106 | 0.00146 |
| #358 | RABAC1 | -0.5686814 | 0.03595048 | -0.5891038 | 0.0582 | -0.5756683 | 0.0363 | -0.5858036 | 0.0244 | -0.5241499 | 0.025 |
| #359 | ARIH1 | 0.3288227 | 0.03596838 | 0.357957 | 0.0286 | 0.3969016 | 0.0321 | 0.2144269 | 0.0821 | 0.3460053 | 0.000971 |
| #360 | COP 1 | 0.6158004 | 0.03604837 | 0.551435 | 0.00661 | 0.5778052 | 0.0388 | 0.7075758 | 0.072 | 0.6263856 | 0.0268 |
| #361 | ZNF460 | 1.203402 | 0.03631857 | 1.379055 | 0.0283 | 1.125892 | 0.038 | 1.013319 | 0.0356 | 1.295342 | 0.0434 |
| #362 | G2E3 | 0.6073417 | 0.03635936 | 0.7237287 | 0.0468 | 0.6364892 | 0.011 | 0.4277507 | 0.0709 | 0.6413982 | 0.0167 |
| #363 | NDUFB11 | -0.3282486 | 0.03643266 | -0.3720861 | 0.0373 | -0.2920297 | 0.0409 | -0.3241987 | 0.00552 | -0.3246799 | 0.062 |
| #364 | CLN3 | -0.434573 | 0.03659933 | -0.3550169 | 0.0132 | -0.4300556 | 0.0297 | -0.6541392 | 0.0583 | -0.2990804 | 0.0451 |
| #365 | OGA | 0.3973394 | 0.03666842 | 0.5625629 | 0.0415 | 0.315325 | 0.0126 | 0.2620795 | 0.0633 | 0.4493903 | 0.0292 |
| #366 | MT1X | -0.6828351 | 0.03670405 | -0.6425602 | 0.0244 | -0.7751233 | 0.00842 | -0.7278507 | 0.112 | -0.5858061 | 0.00187 |
| #367 | TMX3 | 0.33489 | 0.03674818 | 0.3742844 | 0.014 | 0.3254599 | 0.0492 | 0.2426235 | 0.0395 | 0.3971921 | 0.0443 |
| #368 | PNPLA8 | 0.8969623 | 0.03675768 | 0.986853 | 0.00923 | 0.9576797 | 0.0432 | 0.8197918 | 0.0455 | 0.8235247 | 0.0491 |
| #369 | COPS6 | -0.3456552 | 0.03678423 | -0.3123678 | 0.0258 | -0.3839911 | 0.0763 | -0.4093063 | 0.0441 | -0.2769556 | 0.000885 |
| #370 | KAT7 | 0.597724 | 0.0369345 | 0.7750642 | 0.000223 | 0.6183843 | 0.0891 | 0.3802472 | 0.0287 | 0.6172005 | 0.0297 |
| #371 | MYO9A | 0.4929354 | 0.03695715 | 0.61989 | 0.00451 | 0.5742437 | 0.011 | 0.2585793 | 0.13 | 0.5190287 | 0.00232 |
| #372 | DAPK3 | -0.5124372 | 0.03705605 | -0.405662 | 0.0779 | -0.4681588 | 0.0205 | -0.6209376 | 0.0457 | -0.5549903 | 0.00415 |
| #373 | RAB11FIP2 | 0.5822485 | 0.03718199 | 0.4327597 | 0.0138 | 0.6678434 | 0.0118 | 0.6846384 | 0.014 | 0.5437523 | 0.109 |
| #374 | CUL2 | 0.4927335 | 0.03733988 | 0.566889 | 0.00554 | 0.4735104 | 0.0474 | 0.4784304 | 0.0637 | 0.4521042 | 0.0326 |
| #375 | ZRSR2 | 0.4532748 | 0.03737074 | 0.5123616 | 0.0047 | 0.3352249 | 0.123 | 0.4448504 | 0.00252 | 0.5206624 | 0.0191 |
| #376 | ANO6 | 0.5859178 | 0.03738512 | 0.5964932 | 0.023 | 0.6240692 | 0.00263 | 0.5033893 | 0.109 | 0.6197195 | 0.0153 |
| #377 | SFSWAP | 0.3294474 | 0.03738874 | 0.3705913 | 0.013 | 0.3109577 | 0.0926 | 0.2574287 | 0.0021 | 0.3788122 | 0.0419 |
| #378 | OXSR1 | 0.4881577 | 0.03745085 | 0.5301808 | 0.00629 | 0.6045024 | 0.00118 | 0.4141363 | 0.141 | 0.4038116 | 0.00182 |
| #379 | TAB3 | 0.5837183 | 0.03754588 | 0.5245174 | 0.0255 | 0.6489402 | 0.0225 | 0.4685062 | 0.101 | 0.6929095 | 0.000977 |
| #380 | DHX32 | 0.6155475 | 0.03757017 | 0.4951491 | 0.047 | 0.8634656 | 0.0263 | 0.6166901 | 0.0653 | 0.486885 | 0.0117 |
| #381 | HS2ST1 | 0.371525 | 0.03757802 | 0.358931 | 0.0111 | 0.4026502 | 0.0603 | 0.3338508 | 0.0489 | 0.390668 | 0.03 |
| #382 | YTHDC1 | 0.4392661 | 0.03769775 | 0.6395134 | 0.025 | 0.460245 | 0.0608 | 0.2134726 | 0.0355 | 0.4438334 | 0.0294 |
| #383 | CEP95 | 0.6251417 | 0.03805938 | 0.6930457 | 0.0428 | 0.5990151 | 0.0149 | 0.5859455 | 0.0719 | 0.6225607 | 0.0227 |
| #384 | ASCC3 | 0.8702296 | 0.03806529 | 0.9932762 | 0.00803 | 0.6942282 | 0.0922 | 1.005006 | 0.0484 | 0.7884084 | 0.0037 |
| #385 | HPS3 | 0.5167995 | 0.03811485 | 0.4569517 | 0.0922 | 0.5982229 | 0.0103 | 0.429568 | 0.00331 | 0.5824553 | 0.0467 |
| #386 | UBR5 | 0.5188739 | 0.038276 | 0.5879271 | 0.0244 | 0.5359523 | 0.0342 | 0.4690895 | 0.072 | 0.4825266 | 0.0225 |
| #387 | SIAH1 | 0.4359265 | 0.03845968 | 0.5250689 | 0.00388 | 0.4105875 | 0.0298 | 0.3278985 | 0.0404 | 0.4801511 | 0.0798 |
| #388 | RB1CC1 | 0.3250161 | 0.03861535 | 0.4212394 | 0.0411 | 0.2730216 | 0.057 | 0.28116 | 0.0201 | 0.3246435 | 0.0363 |
| #389 | DGUOK | -0.402928 | 0.03863414 | -0.3595902 | 0.0149 | -0.4105232 | 0.0436 | -0.4093964 | 0.0512 | -0.4322022 | 0.0448 |
| #390 | MSL3 | 0.4915708 | 0.03864265 | 0.5989624 | 0.00378 | 0.3887871 | 0.129 | 0.5903656 | 0.0147 | 0.388168 | 0.00675 |
| #391 | UBR2 | 0.4013409 | 0.03885785 | 0.4579934 | 0.0428 | 0.3343435 | 0.061 | 0.406543 | 0.00667 | 0.4064836 | 0.045 |
| #392 | ECPAS | 0.6128106 | 0.03887341 | 0.5262189 | 0.00457 | 0.5351178 | 0.00894 | 0.7277146 | 0.133 | 0.6621909 | 0.0094 |
| #393 | BCL2L12 | -0.4685953 | 0.03902701 | -0.4308237 | 0.00882 | -0.5054243 | 0.107 | -0.5651336 | 0.0128 | -0.3729996 | 0.0273 |
| #394 | FUBP1 | 0.5148566 | 0.03906652 | 0.5284136 | 0.00782 | 0.4934853 | 0.11 | 0.5688902 | 0.0151 | 0.4686374 | 0.0236 |
| #395 | SLC9A9 | 0.5573007 | 0.03914803 | 0.7391543 | 0.0179 | 0.4202512 | 0.0879 | 0.5113271 | 0.0249 | 0.5584703 | 0.026 |
| #396 | NAA20 | -0.2428101 | 0.03945105 | -0.2995585 | 0.00604 | -0.2205058 | 0.0173 | -0.1656417 | 0.0325 | -0.2855344 | 0.102 |
| #397 | RSL1D1 | 0.3609239 | 0.03955141 | 0.4011915 | 0.00297 | 0.3899082 | 0.0773 | 0.2556112 | 0.0436 | 0.3969848 | 0.0343 |
| #398 | CNST | 0.8301404 | 0.03959381 | 1.056363 | 0.0697 | 0.8646703 | 0.0451 | 0.5984071 | 0.0356 | 0.8011211 | 0.00798 |
| #399 | RMDN1 | 0.4526818 | 0.03961833 | 0.6016749 | 0.0315 | 0.5104294 | 0.0469 | 0.4037194 | 0.0305 | 0.2949035 | 0.0495 |
| #400 | UBAP2L | 0.3578143 | 0.03975128 | 0.4013979 | 0.00529 | 0.3224014 | 0.107 | 0.3087071 | 0.0347 | 0.3987509 | 0.0118 |
| #401 | NCOA2 | 0.4962281 | 0.03982297 | 0.5248339 | 0.0309 | 0.4545359 | 0.00482 | 0.4668863 | 0.117 | 0.5386564 | 0.00706 |
| #402 | FDPS | -0.4048237 | 0.03991915 | -0.3292057 | 0.0239 | -0.4366426 | 0.0766 | -0.5440647 | 0.0254 | -0.3093818 | 0.0337 |
| #403 | ZNF700 | 0.6321857 | 0.03994864 | 0.7252644 | 0.0867 | 0.6770769 | 0.0103 | 0.4315482 | 0.0413 | 0.6948534 | 0.0215 |
| #404 | HIF1A | 0.9861243 | 0.04006976 | 0.946658 | 0.00831 | 0.9922515 | 0.0339 | 1.030637 | 0.104 | 0.9749505 | 0.0139 |

| | | | | | | | | | | | |
|---|---|---|---|---|---|---|---|---|---|---|---|
| **#405** | SLC10A7 | 0.5366208 | 0.04010874 | 0.5934172 | 0.0223 | 0.5348306 | 0.00497 | 0.4153607 | 0.0882 | 0.6028747 | 0.045 |
| **#406** | CSNK1G3 | 0.6766855 | 0.04015337 | 0.7210143 | 0.0057 | 0.7658758 | 0.0648 | 0.6523699 | 0.0407 | 0.5674819 | 0.0494 |
| **#407** | ITSN2 | 0.5392035 | 0.04018891 | 0.5921829 | 0.011 | 0.5659157 | 0.0148 | 0.4038937 | 0.122 | 0.5948217 | 0.0125 |
| **#408** | HNRNPA2B1 | 0.3328804 | 0.04021838 | 0.3361989 | 0.000839 | 0.3474564 | 0.0435 | 0.2801409 | 0.0856 | 0.3677254 | 0.031 |
| **#409** | TTC21B | 0.6643527 | 0.04028198 | 0.9824675 | 0.00242 | 0.5200865 | 0.0929 | 0.4551906 | 0.0262 | 0.6996659 | 0.0396 |
| **#410** | PRPF31 | -0.3533524 | 0.04029969 | -0.4536183 | 0.00425 | -0.2679098 | 0.115 | -0.3891016 | 0.0156 | -0.3027798 | 0.0265 |
| **#411** | RIC1 | 0.8704753 | 0.04033179 | 0.6951148 | 0.00629 | 0.8161273 | 0.0143 | 1.06416 | 0.102 | 0.9064991 | 0.0383 |
| **#412** | UBXN2A | 0.3761408 | 0.04038665 | 0.2892627 | 0.000909 | 0.4757519 | 0.0306 | 0.3510977 | 0.0935 | 0.3884512 | 0.0366 |
| **#413** | SMARCAD1 | 0.6989594 | 0.04041637 | 0.8219066 | 0.0454 | 0.7602741 | 0.0608 | 0.5770093 | 0.01 | 0.6366477 | 0.0454 |
| **#414** | APC | 0.5443947 | 0.04042824 | 0.6314346 | 0.000103 | 0.5399684 | 0.0268 | 0.3320145 | 0.128 | 0.6741613 | 0.00678 |
| **#415** | MCM9 | 0.6957215 | 0.04043138 | 0.7832623 | 0.00753 | 0.7979298 | 0.0216 | 0.5554774 | 0.0413 | 0.6462167 | 0.0913 |
| **#416** | AKAP11 | 0.6227701 | 0.04044849 | 0.6757691 | 0.0281 | 0.6110646 | 0.0507 | 0.5923678 | 0.0374 | 0.611879 | 0.0456 |
| **#417** | NDUFB4 | -0.3118597 | 0.04060095 | -0.3353279 | 0.0159 | -0.2876559 | 0.0167 | -0.2708045 | 0.00433 | -0.3536505 | 0.125 |
| **#418** | RNF138 | 0.5382307 | 0.04077824 | 0.4816759 | 0.0236 | 0.5014312 | 0.0193 | 0.6287321 | 0.098 | 0.5410837 | 0.0222 |
| **#419** | ABCE1 | 0.3520996 | 0.04088648 | 0.3421941 | 0.0299 | 0.34225 | 0.0691 | 0.4562771 | 0.0375 | 0.2676771 | 0.027 |
| **#420** | TET2 | 0.8472441 | 0.04095027 | 0.8666317 | 0.00183 | 0.9367723 | 0.0236 | 0.6343924 | 0.133 | 0.9511801 | 0.00484 |
| **#421** | STIM2 | 0.4296583 | 0.04125907 | 0.3606254 | 0.0359 | 0.5148406 | 0.0677 | 0.4570447 | 0.0407 | 0.3861226 | 0.0208 |
| **#422** | UTRN | 0.667637 | 0.04125965 | 0.8935839 | 0.00758 | 0.6323708 | 0.0346 | 0.4707589 | 0.116 | 0.6738344 | 0.00711 |
| **#423** | QSER1 | 0.665294 | 0.0413179 | 0.646291 | 0.0415 | 0.6975235 | 0.00727 | 0.6124024 | 0.108 | 0.7049593 | 0.00899 |
| **#424** | NOTCH2 | 0.6403583 | 0.04152436 | 0.5459098 | 0.012 | 0.4856611 | 0.079 | 0.7634933 | 0.04 | 0.766369 | 0.035 |
| **#425** | AQR | 0.4991263 | 0.04163202 | 0.6981777 | 0.0229 | 0.41945 | 0.0869 | 0.4398478 | 0.0285 | 0.4390298 | 0.0282 |
| **#426** | MIA3 | 0.6182692 | 0.04185159 | 0.7193946 | 0.0181 | 0.529291 | 0.115 | 0.6330855 | 0.0256 | 0.5913056 | 0.00884 |
| **#427** | LMF2 | -0.5986234 | 0.04186341 | -0.4619363 | 0.0907 | -0.6750702 | 0.0115 | -0.5646526 | 0.0244 | -0.6928344 | 0.0409 |
| **#428** | GMPR2 | -0.3924768 | 0.04187348 | -0.4190492 | 0.0218 | -0.3745975 | 0.00549 | -0.3649186 | 0.11 | -0.4113418 | 0.03 |
| **#429** | TRPV2 | -0.4930071 | 0.0419194 | -0.3861863 | 0.0197 | -0.5618239 | 0.0204 | -0.4762652 | 0.106 | -0.5477532 | 0.0217 |
| **#430** | SLC52A2 | -0.4869127 | 0.04202405 | -0.4630344 | 0.0253 | -0.4604016 | 0.0242 | -0.5768936 | 0.0828 | -0.4473214 | 0.0358 |
| **#431** | GTPBP4 | 0.2606875 | 0.04229422 | 0.2520644 | 0.0103 | 0.2571589 | 0.115 | 0.2667641 | 0.0224 | 0.2667627 | 0.0219 |
| **#432** | NRIP1 | 0.7711604 | 0.04233945 | 0.6394617 | 0.0157 | 0.713316 | 0.0165 | 1.016932 | 0.101 | 0.7149316 | 0.036 |
| **#433** | DCAF17 | 0.542909 | 0.0424095 | 0.5817577 | 0.0353 | 0.545869 | 0.0336 | 0.494291 | 0.0678 | 0.5497184 | 0.033 |
| **#434** | CDK12 | 0.4073219 | 0.04243606 | 0.5266141 | 0.0788 | 0.4193498 | 0.0479 | 0.2747431 | 0.0111 | 0.4085804 | 0.032 |
| **#435** | MPP5 | 0.4790773 | 0.04263903 | 0.4323713 | 0.104 | 0.5417757 | 0.037 | 0.3204505 | 0.021 | 0.6217117 | 0.00812 |
| **#436** | PPP6R3 | 0.3655662 | 0.04268753 | 0.4185683 | 0.0315 | 0.3570532 | 0.0416 | 0.2378809 | 0.0972 | 0.4487625 | 0.000408 |
| **#437** | PDCD6IP | 0.4852757 | 0.04273882 | 0.4720621 | 0.00345 | 0.5133911 | 0.0228 | 0.4559837 | 0.11 | 0.499666 | 0.0348 |
| **#438** | TRNAU1AP | -0.3780199 | 0.04284553 | -0.1710028 | 0.0235 | -0.4002905 | 0.0271 | -0.4572423 | 0.0789 | -0.4835441 | 0.0418 |
| **#439** | GNA13 | 0.7152257 | 0.04309789 | 0.5385797 | 0.0487 | 0.9097276 | 0.00685 | 0.6927244 | 0.0689 | 0.7198711 | 0.0479 |
| **#440** | DICER1 | 0.5275527 | 0.04311309 | 0.44713 | 0.00353 | 0.4184332 | 0.0448 | 0.6985381 | 0.0361 | 0.5461097 | 0.088 |
| **#441** | TCERG1 | 0.4110767 | 0.0431353 | 0.5023562 | 0.00694 | 0.4703804 | 0.00173 | 0.2624 | 0.161 | 0.4091703 | 0.00312 |
| **#442** | CREBRF | 0.310258 | 0.04335952 | 0.2723895 | 0.0476 | 0.1882475 | 0.0531 | 0.4053499 | 0.0334 | 0.3750449 | 0.0394 |
| **#443** | CASC4 | 0.3260579 | 0.04347959 | 0.3267385 | 0.0167 | 0.3204224 | 0.104 | 0.203506 | 0.018 | 0.4535648 | 0.0351 |
| **#444** | ZDHHC20 | 0.6508881 | 0.04355276 | 0.4976429 | 0.00756 | 0.6148981 | 0.0173 | 0.7167766 | 0.132 | 0.7742345 | 0.0169 |
| **#445** | HTT | 0.5328628 | 0.04379895 | 0.5766897 | 0.0388 | 0.5732097 | 0.0411 | 0.4131965 | 0.0563 | 0.5683555 | 0.039 |
| **#446** | KDM7A | 0.7948666 | 0.04392005 | 0.7842218 | 0.000111 | 0.8157137 | 0.00668 | 0.7481286 | 0.161 | 0.8314022 | 0.00759 |
| **#447** | ZNF267 | 0.9751639 | 0.04393371 | 0.8386209 | 0.00917 | 0.7525731 | 0.0189 | 1.455498 | 0.102 | 0.8539634 | 0.0455 |
| **#448** | AFTPH | 0.5603661 | 0.04419116 | 0.6063794 | 0.034 | 0.6310899 | 0.0318 | 0.4362466 | 0.0836 | 0.5677485 | 0.0273 |
| **#449** | WDR70 | 0.4303018 | 0.04435744 | 0.6329439 | 0.0021 | 0.3478327 | 0.0978 | 0.3597051 | 0.0399 | 0.3807256 | 0.0376 |
| **#450** | SERGEF | -0.3896624 | 0.04465497 | -0.3572805 | 0.000525 | -0.4116977 | 0.0305 | -0.4074244 | 0.128 | -0.3822469 | 0.0198 |
| **#451** | CYP2U1 | 0.6271442 | 0.04467241 | 0.3936911 | 0.0781 | 0.681176 | 0.0389 | 0.8609957 | 0.0274 | 0.5727139 | 0.0342 |
| **#452** | BIRC2 | 0.3838777 | 0.04469741 | 0.4916985 | 0.00444 | 0.4050349 | 0.027 | 0.2940711 | 0.0269 | 0.3447063 | 0.12 |
| **#453** | USP45 | 0.8135277 | 0.04469789 | 1.090446 | 0.0093 | 0.5763091 | 0.108 | 0.8947702 | 0.0469 | 0.6925853 | 0.0143 |
| **#454** | UTP20 | 0.5675238 | 0.04517774 | 0.6788707 | 0.0138 | 0.4355857 | 0.116 | 0.7489039 | 0.00844 | 0.4067349 | 0.0426 |
| **#455** | PAICS | 0.4650224 | 0.04527453 | 0.4834251 | 0.00363 | 0.3817266 | 0.119 | 0.6643431 | 0.0413 | 0.3305948 | 0.0167 |
| **#456** | MBNL1 | 0.3168042 | 0.0453489 | 0.3909326 | 0.00923 | 0.3203179 | 0.0177 | 0.1970753 | 0.136 | 0.3588908 | 0.0189 |
| **#457** | RERE | 0.6888355 | 0.04545302 | 0.6693156 | 0.0193 | 0.5692537 | 0.0988 | 0.876941 | 0.0345 | 0.6398318 | 0.0292 |
| **#458** | PAXBP1 | 0.8857861 | 0.04547191 | 0.9227774 | 0.00636 | 1.062898 | 0.0459 | 0.7436786 | 0.123 | 0.81379 | 0.00683 |
| **#459** | TSSC4 | -0.5109273 | 0.04550239 | -0.4903252 | 0.0136 | -0.5263562 | 0.0412 | -0.5724416 | 0.0426 | -0.4545863 | 0.0846 |
| **#460** | NOM1 | 0.3015023 | 0.04557261 | 0.4208532 | 0.0387 | 0.1926782 | 0.0198 | 0.2645163 | 0.101 | 0.3279614 | 0.0229 |
| **#461** | PALB2 | 0.4785109 | 0.04575904 | 0.675874 | 0.0337 | 0.4305176 | 0.0484 | 0.4625427 | 0.00013 | 0.3451091 | 0.101 |
| **#462** | CCT7 | -0.3461669 | 0.045975 | -0.3941571 | 0.00357 | -0.3529037 | 0.0363 | -0.323002 | 0.0386 | -0.3146047 | 0.105 |

| | | | | | | | | | | | |
|---|---|---|---|---|---|---|---|---|---|---|---|
| **#463** | GGA2 | 0.4398575 | 0.04604548 | 0.4734928 | 0.0355 | 0.5235138 | 0.029 | 0.3613882 | 0.00931 | 0.4010352 | 0.11 |
| **#464** | TM9SF3 | 0.5929303 | 0.04613703 | 0.4266906 | 0.0392 | 0.6936029 | 0.0403 | 0.713531 | 0.0614 | 0.5378968 | 0.0437 |
| **#465** | BDP1 | 0.391957 | 0.04621749 | 0.6226084 | 0.0408 | 0.3514324 | 0.123 | 0.197904 | 0.0113 | 0.3958833 | 0.0095 |
| **#466** | ATF6 | 0.3741168 | 0.04624631 | 0.3149231 | 0.0231 | 0.3309751 | 0.0873 | 0.4076197 | 0.0489 | 0.4429493 | 0.0256 |
| **#467** | MLH1 | 0.2758086 | 0.04634284 | 0.4774223 | 0.039 | 0.1670025 | 0.00963 | 0.1539193 | 0.0932 | 0.3048902 | 0.0435 |
| **#468** | EWSR1 | 0.3278929 | 0.04634349 | 0.3622054 | 0.0433 | 0.3309069 | 0.0192 | 0.2312 | 0.123 | 0.3872594 | 0.000336 |
| **#469** | MYCBP2 | 0.4300107 | 0.04636062 | 0.362676 | 0.0135 | 0.4347778 | 0.136 | 0.5946917 | 0.0282 | 0.3278974 | 0.00812 |
| **#470** | ZBTB33 | 0.5802201 | 0.04641519 | 0.6165459 | 0.0163 | 0.5762275 | 0.0371 | 0.6185024 | 0.02 | 0.5096048 | 0.112 |
| **#471** | NEDD8 | -0.3142307 | 0.04653418 | -0.3564697 | 0.0432 | -0.3087278 | 0.0173 | -0.3038779 | 0.00306 | -0.2878474 | 0.123 |
| **#472** | NOTCH2NLC | 0.9767526 | 0.04654306 | 0.9666807 | 0.019 | 1.15741 | 0.00969 | 0.7430987 | 0.15 | 1.03982 | 0.00764 |
| **#473** | CYLD | 0.4999662 | 0.04672745 | 0.4450108 | 0.1 | 0.4534204 | 0.00774 | 0.6758913 | 0.037 | 0.4255421 | 0.0423 |
| **#474** | HAX1 | -0.3175667 | 0.04692613 | -0.309685 | 0.0737 | -0.3764235 | 0.0453 | -0.2223933 | 0.0242 | -0.3617649 | 0.0445 |
| **#475** | UHRF1BP1L | 0.7870639 | 0.04702878 | 0.7932375 | 0.0218 | 0.8035645 | 0.03 | 0.7112189 | 0.12 | 0.8402347 | 0.0163 |
| **#476** | ATP5MPL | -0.3174274 | 0.04706353 | -0.3018505 | 0.0234 | -0.2986846 | 0.0263 | -0.3590568 | 0.00755 | -0.3101175 | 0.131 |
| **#477** | FASTKD2 | 0.5127081 | 0.04710869 | 0.6073147 | 0.0174 | 0.523566 | 0.0201 | 0.4232735 | 0.128 | 0.4966782 | 0.023 |
| **#478** | RALGAPB | 0.441281 | 0.04721435 | 0.5371727 | 0.016 | 0.5134317 | 0.0147 | 0.3444423 | 0.136 | 0.3700772 | 0.0221 |
| **#479** | DOCK11 | 0.5477638 | 0.04743595 | 0.6365605 | 0.00406 | 0.5836371 | 0.037 | 0.4596367 | 0.14 | 0.5112207 | 0.00845 |
| **#480** | TOMM70 | 0.3865008 | 0.04763589 | 0.3154229 | 0.0344 | 0.5824186 | 0.0315 | 0.2750719 | 0.0756 | 0.3730899 | 0.0491 |
| **#481** | ZNF688 | -0.3527818 | 0.04792486 | -0.3080874 | 0.0396 | -0.3647087 | 0.00145 | -0.2212288 | 0.133 | -0.5171025 | 0.0178 |
| **#482** | WDR83 | -0.4290789 | 0.04816326 | -0.2833403 | 0.022 | -0.440586 | 0.0411 | -0.6396297 | 0.0429 | -0.3527595 | 0.0867 |
| **#483** | PPP4R3B | 0.5141426 | 0.0482783 | 0.609787 | 0.0156 | 0.4337036 | 0.133 | 0.5296738 | 0.0175 | 0.483406 | 0.0268 |
| **#484** | TNPO1 | 0.4151006 | 0.04845354 | 0.4188687 | 0.0297 | 0.4992954 | 0.0347 | 0.2691539 | 0.0818 | 0.4730846 | 0.0476 |
| **#485** | UVRAG | 0.6301903 | 0.04858392 | 0.5840615 | 0.0334 | 0.802891 | 0.0314 | 0.5258996 | 0.0837 | 0.6079093 | 0.0458 |
| **#486** | TRAPPC9 | 0.4495056 | 0.04864279 | 0.4515203 | 0.0206 | 0.477664 | 0.0333 | 0.3183265 | 0.104 | 0.5505115 | 0.0364 |
| **#487** | PPM1B | 0.3137943 | 0.04864743 | 0.3217481 | 0.135 | 0.3195285 | 0.00366 | 0.2123622 | 0.0243 | 0.4015386 | 0.0322 |
| **#488** | DNAJB14 | 0.3236494 | 0.04873853 | 0.4352221 | 0.00153 | 0.3624673 | 0.0339 | 0.1414175 | 0.132 | 0.3554906 | 0.0279 |
| **#489** | HNRNPK | 0.1848543 | 0.04880033 | 0.1959941 | 0.0453 | 0.1923453 | 0.00595 | 0.145826 | 0.129 | 0.2052517 | 0.0149 |
| **#490** | NAA30 | 0.4040812 | 0.04885254 | 0.3905616 | 0.0188 | 0.3752417 | 0.115 | 0.4147212 | 0.041 | 0.4358002 | 0.0211 |
| **#491** | NSF | 1.042605 | 0.0489267 | 1.24941 | 0.0359 | 0.9619108 | 0.0468 | 0.9196442 | 0.0714 | 1.039454 | 0.0416 |
| **#492** | FAR1 | 0.6826946 | 0.04915251 | 0.625664 | 0.0191 | 0.6181048 | 0.0355 | 0.7697515 | 0.12 | 0.7172582 | 0.0219 |
| **#493** | PCM1 | 0.4733845 | 0.04920345 | 0.6253697 | 0.0407 | 0.4854049 | 0.017 | 0.3324533 | 0.136 | 0.4503101 | 0.00287 |
| **#494** | WASHC1 | -0.47878 | 0.04922225 | -0.3585352 | 0.183 | -0.5282383 | 0.00892 | -0.654079 | 0.00169 | -0.3742672 | 0.00374 |
| **#495** | ZBTB10 | 0.8804541 | 0.04961772 | 1.028964 | 0.0104 | 0.948194 | 0.0427 | 0.6518073 | 0.112 | 0.8928514 | 0.033 |

Supplementary Table 3

| Rank | Symbol | Control Abundance | Expression | Significance (p) |
|---|---|---|---|---|
| #1 | TREML1 | 85.26282 | 1.552503 | 7.28784E-05 |
| #2 | SPARC | 79.08122 | 1.472755 | 0.000424952 |
| #3 | TUBB1 | 246.586 | 1.528078 | 0.002863439 |
| #4 | IFI30 | 374.5127 | 0.9925443 | 0.000990654 |
| #5 | IRF1 | 267.7736 | 1.562205 | 0.003189242 |
| #6 | GNG11 | 271.4535 | 1.583858 | 0.003413497 |
| #7 | PGRMC1 | 78.07159 | 1.33693 | 0.00103541 |
| #8 | PPBP | 1,269 | 1.77364 | 0.004593598 |
| #9 | NRGN | 541.1837 | 1.618094 | 0.004349181 |
| #10 | MTPN | 273.9814 | 0.8408795 | 9.33724E-05 |
| #11 | HIST1H2AC | 168.1354 | 1.336154 | 0.003289903 |
| #12 | CAVIN2 | 280.5133 | 1.555458 | 0.005147299 |
| #13 | SNCA | 32.20159 | 1.528557 | 0.000574943 |
| #14 | PLEK | 192.6149 | 0.855626 | 0.000691498 |
| #15 | MARCH2 | 69.68294 | 1.117741 | 0.000947394 |
| #16 | TSC22D1 | 106.1591 | 1.238208 | 0.002946624 |
| #17 | STX11 | 61.12124 | 1.24664 | 0.002061361 |
| #18 | MYL9 | 74.04531 | 1.790189 | 0.00494366 |
| #19 | SAT1 | 869.5761 | 0.9636897 | 0.004396921 |
| #20 | IGSF6 | 81.28348 | 0.941251 | 0.000630948 |
| #21 | BEX3 | 51.70968 | 1.159336 | 0.001476404 |
| #22 | MPP1 | 68.7961 | 1.23127 | 0.003505503 |
| #23 | LIMS1 | 200.7433 | 0.9848492 | 0.003905488 |
| #24 | HIST1H3H | 23.20346 | 1.695273 | 0.001890449 |
| #25 | ACRBP | 87.82995 | 1.469392 | 0.005682866 |
| #26 | FERMT3 | 158.8563 | 0.9185932 | 0.00312829 |
| #27 | POMP | 116.4362 | 0.799266 | 0.000695229 |
| #28 | IFITM3 | 128.616 | 1.827356 | 0.007724409 |
| #29 | MPIG6B | 37.20661 | 1.446975 | 0.003097915 |

| | | | | |
|---|---|---|---|---|
| **#30** | ACTB | 4,208 | 0.7007108 | 0.002813101 |
| **#31** | SOD2 | 176.5926 | 1.181239 | 0.00641125 |
| **#32** | PSME1 | 350.1364 | 0.7303608 | 0.00181629 |
| **#33** | SH3BGRL2 | 25.76241 | 1.453209 | 0.001409269 |
| **#34** | OAZ1 | 1,135 | 0.7101355 | 0.003096598 |
| **#35** | TLN1 | 426.2464 | 0.8644187 | 0.004954949 |
| **#36** | RAB31 | 68.62104 | 0.9241658 | 0.001612984 |
| **#37** | CALM3 | 203.5743 | 0.7534051 | 0.001778485 |
| **#38** | CTSA | 108.0967 | 1.197917 | 0.006147388 |
| **#39** | PF4 | 548.1379 | 1.540699 | 0.01063242 |
| **#40** | STOM | 51.67487 | 1.010017 | 0.001307027 |
| **#41** | NPC2 | 168.6719 | 0.8425448 | 0.003134795 |
| **#42** | LAMTOR1 | 176.8414 | 0.786132 | 0.002450775 |
| **#43** | SNN | 46.27349 | 1.101308 | 0.002030029 |
| **#44** | MMD | 56.42601 | 1.177986 | 0.003798577 |
| **#45** | AAK1 | 243.2914 | -0.7050016 | 0.000615042 |
| **#46** | FTH1 | 3,635 | 0.7188019 | 0.004124819 |
| **#47** | RGS18 | 229.5385 | 1.157158 | 0.007687986 |
| **#48** | ITGA2B | 52.56335 | 1.745756 | 0.007647537 |
| **#49** | SCO2 | 52.75488 | 1.573637 | 0.007176235 |
| **#50** | LY6E | 188.3516 | 0.9262981 | 0.00549401 |
| **#51** | SLC31A2 | 23.8802 | 1.446208 | 0.002115846 |
| **#52** | CLU | 183.0197 | 1.572422 | 0.01088413 |
| **#53** | PRDX6 | 147.1202 | 0.7663065 | 0.002564753 |
| **#54** | ARPC1B | 636.6273 | 0.6829581 | 0.003043442 |
| **#55** | H3F3A | 1,438 | 0.5927746 | 0.001165273 |
| **#56** | SRGN | 656.436 | 0.6875014 | 0.00346426 |
| **#57** | PDIA3 | 189.803 | 0.6229984 | 0.000120241 |
| **#58** | RSRP1 | 290.9315 | -0.6514141 | 0.001235739 |
| **#59** | SERPINB1 | 170.6303 | 0.6202152 | 0.000196614 |
| **#60** | RAB32 | 76.7119 | 0.8081275 | 0.001873669 |
| **#61** | RGS10 | 288.537 | 0.7254717 | 0.004174583 |

| | | | | |
|---|---|---|---|---|
| **#62** | WARS | 71.46555 | 2.450773 | 0.01394277 |
| **#63** | SNX10 | 65.10069 | 1.185343 | 0.007532101 |
| **#64** | PSMB9 | 197.9112 | 1.0246 | 0.009454648 |
| **#65** | F13A1 | 86.82678 | 1.46251 | 0.01149096 |
| **#66** | RNF11 | 51.24854 | 0.9186639 | 0.002547247 |
| **#67** | BLVRA | 37.65 | 0.9996893 | 0.001765115 |
| **#68** | CAPZA2 | 115.0265 | 0.6438666 | 0.000357015 |
| **#69** | ARF1 | 208.5302 | 0.5980173 | 0.00038831 |
| **#70** | CLEC1B | 36.35673 | 1.36764 | 0.006530241 |
| **#71** | MYL6 | 955.0634 | 0.5694011 | 0.001197342 |
| **#72** | PSME2 | 198.4488 | 1.537332 | 0.01470126 |
| **#73** | ARPC5 | 339.4519 | 0.5685621 | 0.000235682 |
| **#74** | MAFB | 57.13901 | 1.019821 | 0.005354769 |
| **#75** | CASP1 | 86.1545 | 0.8004241 | 0.003685053 |
| **#76** | FGL2 | 223.7214 | 1.07369 | 0.01211102 |
| **#77** | DAB2 | 38.87307 | 0.9679863 | 0.002183069 |
| **#78** | TGFB1 | 278.7403 | 0.6688764 | 0.003725286 |
| **#79** | VCL | 134.0362 | 1.092331 | 0.01083599 |
| **#80** | FBXO6 | 16.2687 | 1.400636 | 0.001126607 |
| **#81** | NAPA | 54.23557 | 0.8477231 | 0.002140858 |
| **#82** | GUCY1B1 | 14.36251 | 1.446188 | 0.00076453 |
| **#83** | PTGS1 | 28.97316 | 1.541266 | 0.007509216 |
| **#84** | PSMA4 | 100.4525 | 0.899605 | 0.007097641 |
| **#85** | PBXIP1 | 92.88151 | -0.7455351 | 0.000191493 |
| **#86** | UBE2L6 | 53.8472 | 1.567998 | 0.0124027 |
| **#87** | PDLIM1 | 63.53642 | 1.233167 | 0.009901992 |
| **#88** | AIF1 | 567.3137 | 0.7702925 | 0.00896633 |
| **#89** | SNHG25 | 98.91588 | -1.403959 | 0.008793474 |
| **#90** | ISG15 | 87.68257 | 1.392099 | 0.01396905 |
| **#91** | PKM | 216.5686 | 0.7696564 | 0.007449629 |
| **#92** | MYL12A | 762.5443 | 0.5715006 | 0.002443395 |
| **#93** | CD151 | 35.92985 | 1.031048 | 0.003655321 |

| | | | | |
|---|---|---|---|---|
| **#94** | NCOA4 | 186.0093 | 1.12203 | 0.01451419 |
| **#95** | LAP3 | 57.6571 | 2.20567 | 0.01810281 |
| **#96** | GAS2L1 | 25.30492 | 1.176548 | 0.003602347 |
| **#97** | PSMB10 | 158.8855 | 0.8353463 | 0.009303865 |
| **#98** | TUBA1B | 230.2193 | 0.5533347 | 0.000480459 |
| **#99** | PSMB8 | 156.954 | 0.7051543 | 0.00552566 |
| **#100** | PSTPIP2 | 35.37772 | 1.407344 | 0.01038153 |
| **#101** | BST2 | 143.4693 | 0.7171624 | 0.00608315 |
| **#102** | PLAC8 | 263.7675 | 0.6402878 | 0.005279271 |
| **#103** | IRF9 | 74.49532 | 0.8371239 | 0.006515788 |
| **#104** | GBP2 | 72.72225 | 1.638615 | 0.01786994 |
| **#105** | SNAP23 | 89.02329 | 0.7749897 | 0.006091786 |
| **#106** | RIPK2 | 24.39093 | 1.067087 | 0.002531851 |
| **#107** | NT5C3A | 83.98188 | 0.8826505 | 0.008821249 |
| **#108** | CHD3 | 106.7856 | -0.8758146 | 0.00619297 |
| **#109** | MTHFD2 | 19.95848 | 1.24364 | 0.003451202 |
| **#110** | MT2A | 138.9037 | 1.413852 | 0.01967868 |
| **#111** | HIST1H2BJ | 22.24434 | 1.393688 | 0.007309703 |
| **#112** | TAGLN2 | 426.9276 | 0.8069053 | 0.01408131 |
| **#113** | GLUL | 66.88105 | 0.7771799 | 0.004690735 |
| **#114** | ABCC3 | 17.83985 | 1.124283 | 0.0010314 |
| **#115** | NDUFAF3 | 65.97709 | 0.7404139 | 0.003615384 |
| **#116** | TMC8 | 85.68586 | -0.7959606 | 0.0032099 |
| **#117** | MFSD1 | 56.59911 | 0.7699583 | 0.003632305 |
| **#118** | FKBP1A | 178.4957 | 0.7212158 | 0.008792929 |
| **#119** | TMEM219 | 103.6238 | 0.5849305 | 0.000900956 |
| **#120** | DMTN | 18.48956 | 1.30223 | 0.00426662 |
| **#121** | APOL6 | 90.04748 | 1.087857 | 0.01569065 |
| **#122** | TALDO1 | 188.8779 | 0.6994689 | 0.008221061 |
| **#123** | BCL2A1 | 40.21757 | 1.014834 | 0.007662982 |
| **#124** | MAP3K7CL | 95.21595 | 1.297298 | 0.01901896 |
| **#125** | VMP1 | 86.60871 | 0.6353739 | 0.002284546 |

| | | | | |
|---|---|---|---|---|
| **#126** | GSTO1 | 127.4499 | 0.9060048 | 0.01427813 |
| **#127** | SH3BGRL3 | 1,384 | 0.6046945 | 0.008153714 |
| **#128** | TPM4 | 190.0267 | 0.9965476 | 0.01791927 |
| **#129** | SMIM5 | 12.51845 | 1.501917 | 0.003681389 |
| **#130** | MAX | 116.2007 | 0.8314045 | 0.01219535 |
| **#131** | FHL1 | 18.81557 | 0.9991001 | 0.000195433 |
| **#132** | SORL1 | 154.4806 | -0.570802 | 0.000526104 |
| **#133** | TRAF3IP3 | 220.9556 | -0.732279 | 0.009250088 |
| **#134** | VAMP5 | 80.04888 | 1.887429 | 0.02364802 |
| **#135** | MYD88 | 41.97574 | 0.7015947 | 0.000155344 |
| **#136** | SH3BGRL | 229.4835 | 0.5438657 | 0.002077024 |
| **#137** | PLSCR1 | 42.86829 | 1.137773 | 0.01235394 |
| **#138** | PLXDC2 | 47.43367 | 0.7912103 | 0.00398902 |
| **#139** | TUBA1C | 45.66631 | 1.172888 | 0.01374345 |
| **#140** | GRAP2 | 68.89452 | 1.096423 | 0.01569656 |
| **#141** | MOB1A | 179.4483 | 0.5369848 | 0.001107275 |
| **#142** | PSMA2 | 101.7928 | 0.5750256 | 0.001016785 |
| **#143** | PSMB3 | 131.9938 | 0.6121633 | 0.004449328 |
| **#144** | CD38 | 23.93994 | 0.9955934 | 0.003090926 |
| **#145** | ARHGAP18 | 43.23259 | 1.03754 | 0.01048744 |
| **#146** | LPCAT2 | 23.5803 | 1.221877 | 0.007533847 |
| **#147** | ENDOD1 | 21.66576 | 0.9689312 | 0.001655888 |
| **#148** | RAB8A | 87.70065 | 0.6061287 | 0.002274628 |
| **#149** | ATP1B3 | 67.22903 | 0.7257985 | 0.005727912 |
| **#150** | MLH3 | 35.51513 | 0.8577878 | 0.003936883 |
| **#151** | HIST1H4C | 215.4396 | -0.5249735 | 0.000751059 |
| **#152** | ILK | 66.43945 | 0.9973036 | 0.01556055 |
| **#153** | LEPROT | 58.65942 | 0.62536 | 0.000971188 |
| **#154** | CSKMT | 37.01425 | -1.195215 | 0.003443743 |
| **#155** | SLAMF7 | 30.2573 | 0.8083012 | 0.0017028 |
| **#156** | PTPN18 | 103.1893 | 0.617533 | 0.004174424 |
| **#157** | CYB5R3 | 44.94577 | 0.6945233 | 0.001525188 |

| | | | | |
|---|---|---|---|---|
| **#158** | STAT1 | 89.907 | 1.886564 | 0.02740865 |
| **#159** | ATG3 | 80.98414 | 0.5853333 | 0.001530175 |
| **#160** | ARF3 | 45.26316 | 0.6842558 | 0.001339459 |
| **#161** | CYBB | 203.2226 | 0.7678038 | 0.01564913 |
| **#162** | RAP1B | 395.8446 | 0.5170961 | 0.003119577 |
| **#163** | TIMP1 | 193.8146 | 0.6734755 | 0.01144659 |
| **#164** | CD9 | 35.87822 | 1.621229 | 0.02166026 |
| **#165** | TRAPPC1 | 159.6504 | 0.5439323 | 0.002586929 |
| **#166** | ETFA | 52.47802 | 0.6342019 | 0.001023901 |
| **#167** | GBP5 | 49.99532 | 2.052828 | 0.02705444 |
| **#168** | SMOX | 29.83004 | 1.088948 | 0.009839977 |
| **#169** | CD52 | 916.8304 | -0.497457 | 0.002893084 |
| **#170** | DYNLT1 | 60.30873 | 0.7371095 | 0.007013882 |
| **#171** | TMEM140 | 20.53386 | 1.444054 | 0.01276047 |
| **#172** | CTDSPL | 16.91608 | 1.329593 | 0.007658785 |
| **#173** | IFI35 | 32.59613 | 1.32232 | 0.01650662 |
| **#174** | SUPT4H1 | 79.30531 | 0.5574812 | 0.00033414 |
| **#175** | MARCKS | 112.2883 | 0.7981257 | 0.01527466 |
| **#176** | RORA | 117.2705 | -0.5815102 | 0.001645898 |
| **#177** | MPHOSPH8 | 199.2864 | -0.5212366 | 0.001175345 |
| **#178** | MTURN | 54.65477 | 1.01543 | 0.01645613 |
| **#179** | SERF2 | 1,108 | 0.4855637 | 0.003016262 |
| **#180** | THBS1 | 37.49991 | 1.137175 | 0.01546818 |
| **#181** | CDK2AP1 | 30.87776 | 0.9053864 | 0.006443155 |
| **#182** | ODF3B | 21.98973 | 1.139063 | 0.008044976 |
| **#183** | TNFAIP2 | 87.66051 | 0.811239 | 0.01535987 |
| **#184** | ACAP1 | 170.4188 | -0.5504126 | 0.002630124 |
| **#185** | TXN | 110.6441 | 0.5426534 | 0.001848397 |
| **#186** | PARVB | 41.46619 | 1.175033 | 0.01862982 |
| **#187** | PRKCA | 54.47474 | -0.9513305 | 0.007425294 |
| **#188** | TUBA4A | 162.0367 | 1.045195 | 0.02642224 |
| **#189** | IRF7 | 47.05295 | 1.03544 | 0.01701031 |

| | | | | |
|---|---|---|---|---|
| **#190** | PSMA6 | 102.1946 | 0.5287285 | 0.000961372 |
| **#191** | GBP4 | 50.01679 | 1.837063 | 0.02964097 |
| **#192** | TNFSF13B | 74.12083 | 1.165944 | 0.02498842 |
| **#193** | HLA-E | 1,346 | 0.4342604 | 0.000254717 |
| **#194** | FPR1 | 45.61882 | 1.16808 | 0.02006427 |
| **#195** | LUC7L3 | 205.7465 | -0.526571 | 0.002923791 |
| **#196** | GNG10 | 48.94649 | 0.636181 | 0.002413668 |
| **#197** | SKAP2 | 94.21613 | 0.5376006 | 0.001516436 |
| **#198** | DIP2A | 55.21024 | -0.8960165 | 0.007234998 |
| **#199** | MXD1 | 42.0605 | 0.7853724 | 0.007484894 |
| **#200** | MCUR1 | 34.65887 | 0.8063532 | 0.006163917 |
| **#201** | AKIRIN2 | 69.92097 | 0.5623434 | 0.001266506 |
| **#202** | GNB4 | 28.62868 | 0.9658225 | 0.008944235 |
| **#203** | FCGR2A | 61.01531 | 0.8618387 | 0.01568242 |
| **#204** | RAB20 | 8.067361 | 1.755049 | 0.007329876 |
| **#205** | RHOA | 335.1426 | 0.4623031 | 0.001299517 |
| **#206** | WDFY1 | 30.77316 | 0.838439 | 0.006227405 |
| **#207** | MYLK | 22.12326 | 1.323014 | 0.01587872 |
| **#208** | ZNF185 | 12.32575 | 1.212931 | 0.003222483 |
| **#209** | PLEKHO1 | 156.2719 | 0.7586936 | 0.02056245 |
| **#210** | GCH1 | 21.68344 | 1.198688 | 0.0131004 |
| **#211** | SLC8A1 | 30.67236 | 0.9839579 | 0.01202045 |
| **#212** | BCL9L | 52.46569 | -0.8601843 | 0.006901146 |
| **#213** | IL16 | 88.00106 | -0.606231 | 0.003311051 |
| **#214** | GABARAP | 447.7017 | 0.6080109 | 0.01616779 |
| **#215** | EPSTI1 | 39.11403 | 1.859859 | 0.03270598 |
| **#216** | AP2S1 | 142.2597 | 0.5554519 | 0.00680019 |
| **#217** | MYOF | 18.47413 | 1.867229 | 0.02460376 |
| **#218** | PLA2G12A | 40.03085 | 0.8991337 | 0.01376658 |
| **#219** | HLA-DMA | 144.2753 | 0.6480004 | 0.01487687 |
| **#220** | TMSB4X | 7,684 | 0.4234276 | 0.001265912 |
| **#221** | SPOCK2 | 122.5282 | -0.7187433 | 0.01342851 |

| | | | | |
|---|---|---|---|---|
| **#222** | LBH | 118.033 | -0.7497861 | 0.01441665 |
| **#223** | CD68 | 129.5169 | 0.6720034 | 0.01576295 |
| **#224** | HLA-DPA1 | 767.8102 | 0.7053928 | 0.02516231 |
| **#225** | TAP1 | 55.03341 | 1.159367 | 0.0265095 |
| **#226** | PCGF5 | 72.70684 | 0.5641451 | 0.003192383 |
| **#227** | PARP9 | 27.13306 | 1.529494 | 0.02594163 |
| **#228** | EPB41L3 | 22.45356 | 1.155293 | 0.01412061 |
| **#229** | BRD9 | 51.72163 | -0.7074692 | 0.002905013 |
| **#230** | CSF2RB | 16.90728 | 1.026812 | 0.004997302 |
| **#231** | RALB | 36.53144 | 0.6957921 | 0.003898157 |
| **#232** | BCL3 | 18.69603 | 1.082242 | 0.008368537 |
| **#233** | DAPP1 | 72.65127 | 0.8732628 | 0.02205958 |
| **#234** | RPS27 | 9,985 | -0.4482122 | 0.004078161 |
| **#235** | SERPINA1 | 192.4528 | 0.7085674 | 0.0218256 |
| **#236** | PAK1 | 53.12142 | 0.6704831 | 0.007358419 |
| **#237** | ATOX1 | 55.02361 | 0.6472922 | 0.006711595 |
| **#238** | GBP1 | 33.12629 | 2.688345 | 0.03955143 |
| **#239** | LILRB1 | 48.87227 | 0.9208508 | 0.01977271 |
| **#240** | APOBEC3A | 37.95317 | 1.744308 | 0.03493848 |
| **#241** | SORT1 | 16.60214 | 1.066653 | 0.006882561 |
| **#242** | GMPR | 22.82847 | 1.086441 | 0.01363819 |
| **#243** | UBE2D1 | 39.57919 | 0.636871 | 0.002971996 |
| **#244** | PYGL | 30.96921 | 0.8778252 | 0.01142576 |
| **#245** | GSN | 46.48734 | 0.7974313 | 0.01409043 |
| **#246** | RPS29 | 5,941 | -0.4609816 | 0.00681024 |
| **#247** | IFIT2 | 13.94526 | 2.066956 | 0.02669777 |
| **#248** | MALAT1 | 55,568 | -0.4982592 | 0.01137541 |
| **#249** | CLIC1 | 346.5417 | 0.5773873 | 0.01597065 |
| **#250** | CWF19L2 | 84.09774 | -0.6062573 | 0.004543617 |
| **#251** | TNFSF10 | 74.72967 | 1.522053 | 0.03844334 |
| **#252** | PLEKHA1 | 61.48795 | -0.6579646 | 0.003941605 |
| **#253** | HSBP1 | 74.14542 | 0.604966 | 0.007877475 |

| | | | | |
|---|---|---|---|---|
| #254 | CMTM6 | 125.8772 | 0.5610243 | 0.00893816 |
| #255 | S100A9 | 2,291 | 0.8969671 | 0.0395316 |
| #256 | CDKN1A | 51.32635 | 1.102344 | 0.02790789 |
| #257 | SCN1B | 16.28486 | 1.486016 | 0.01954213 |
| #258 | STAT2 | 50.64758 | 1.039247 | 0.02641776 |
| #259 | CNDP2 | 52.94057 | 0.7985594 | 0.01771098 |
| #260 | XKR8 | 11.07685 | 1.306351 | 0.007051714 |
| #261 | SQOR | 32.58362 | 0.9119594 | 0.0152438 |
| #262 | MLLT6 | 90.95438 | -0.5178247 | 0.000562922 |
| #263 | SMCO4 | 18.68924 | 1.095534 | 0.01233492 |
| #264 | TAX1BP3 | 24.88407 | 0.8683335 | 0.008783142 |
| #265 | ADIPOR1 | 40.53328 | 0.6399537 | 0.00464636 |
| #266 | BCL11B | 272.959 | -0.9804726 | 0.0358113 |
| #267 | SUMO3 | 73.52146 | 0.4955427 | 0.000264442 |
| #268 | PLBD1 | 51.92275 | 0.7832591 | 0.01746142 |
| #269 | NAMPT | 200.6389 | 0.8603874 | 0.03504233 |
| #270 | GLIPR2 | 77.25479 | 0.6055166 | 0.01084656 |
| #271 | NOP53 | 507.9641 | -0.6086622 | 0.02258153 |
| #272 | RIT1 | 35.42559 | 0.5893776 | 0.000739291 |
| #273 | CCND3 | 174.7214 | 0.4759332 | 0.004722021 |
| #274 | CARD16 | 123.509 | 0.5910062 | 0.01570235 |
| #275 | ATF5 | 15.60676 | 1.169526 | 0.01325784 |
| #276 | PLAAT4 | 176.4711 | 0.7078666 | 0.02779628 |
| #277 | TCF7 | 185.7125 | -0.8111375 | 0.03094064 |
| #278 | EIF2AK2 | 64.692 | 0.7717913 | 0.02231352 |
| #279 | P2RY13 | 21.89639 | 0.8997528 | 0.01010323 |
| #280 | CCL5 | 765.3792 | 0.6867127 | 0.03372901 |
| #281 | LILRA5 | 50.5441 | 0.7459862 | 0.01716031 |
| #282 | RBX1 | 137.1822 | 0.5122712 | 0.007788798 |
| #283 | PF4V1 | 52.90605 | 1.676891 | 0.04645206 |
| #284 | LIME1 | 157.7014 | -0.574965 | 0.01338737 |
| #285 | SLC39A3 | 24.92654 | 0.6430952 | 9.78812E-05 |

| | | | | |
|---|---|---|---|---|
| #286 | MCL1 | 233.7217 | 0.4993282 | 0.01034858 |
| #287 | PYCARD | 148.5479 | 0.6129427 | 0.02109064 |
| #288 | CLEC2D | 158.005 | -0.68407 | 0.02284112 |
| #289 | TNFRSF25 | 41.40611 | -0.9455685 | 0.01308667 |
| #290 | IFI6 | 50.01329 | 1.242913 | 0.03950027 |
| #291 | CARD19 | 51.41137 | 0.7468303 | 0.01883272 |
| #292 | OAS3 | 12.57072 | 1.506044 | 0.02096317 |
| #293 | TMEM50A | 129.1314 | 0.4662609 | 0.003330009 |
| #294 | SFT2D2 | 65.86748 | 0.5792758 | 0.008811847 |
| #295 | LYN | 173.5683 | 0.6304798 | 0.02507612 |
| #296 | GNS | 41.38806 | 0.5857852 | 0.00369565 |
| #297 | APP | 54.96778 | 0.5595049 | 0.004958433 |
| #298 | NMT2 | 28.64445 | -0.7763832 | 0.00066921 |
| #299 | WDR1 | 110.6317 | 0.5418216 | 0.01069745 |
| #300 | MARCH9 | 33.73771 | -0.9690071 | 0.009092025 |
| #301 | STMP1 | 97.9632 | 0.4736164 | 0.002809364 |
| #302 | FGD2 | 36.14675 | 0.7190948 | 0.01216478 |
| #303 | NLRP1 | 59.64371 | -0.6564836 | 0.007986717 |
| #304 | CD6 | 53.91928 | -0.7492688 | 0.011797 |
| #305 | HLA-B | 2,854 | 0.3775159 | 0.001318479 |
| #306 | PFKFB3 | 16.65154 | 1.212694 | 0.02021346 |
| #307 | ICAM1 | 14.92712 | 1.129484 | 0.01394083 |
| #308 | UBL4A | 25.75718 | 0.8153358 | 0.01229265 |
| #309 | IFIT3 | 11.28859 | 2.283272 | 0.0384755 |
| #310 | NCF1 | 151.6696 | 1.0843 | 0.04920712 |
| #311 | CALHM6 | 84.81405 | 1.183154 | 0.04780677 |
| #312 | FABP5 | 19.23942 | 0.7355913 | 0.001714778 |
| #313 | SOCS3 | 55.84749 | 0.9093368 | 0.03284633 |
| #314 | INAFM2 | 14.69761 | 0.9307394 | 0.005506011 |
| #315 | ITPKB | 43.82715 | -0.7698641 | 0.009022086 |
| #316 | B2M | 9,777 | 0.4071316 | 0.005764011 |
| #317 | MKRN1 | 90.09228 | 0.476912 | 0.003189385 |

| | | | | |
|---|---|---|---|---|
| **#318** | RPS21 | 3,528 | -0.4214315 | 0.007317794 |
| **#319** | OAS1 | 32.75495 | 0.9348449 | 0.02490737 |
| **#320** | GM2A | 16.97748 | 1.188184 | 0.02099202 |
| **#322** | ARPC3 | 627.1222 | 0.430472 | 0.007148129 |
| **#323** | SMIM3 | 22.75598 | 1.025456 | 0.02170226 |
| **#324** | FCER1G | 487.8559 | 0.7754708 | 0.04452778 |
| **#325** | CAP1 | 272.457 | 0.4323579 | 0.005052216 |
| **#326** | LACTB | 44.7433 | 0.6250178 | 0.01012605 |
| **#327** | BNIP2 | 99.46681 | 0.4841563 | 0.004936149 |
| **#328** | EMC3 | 67.72621 | 0.5675747 | 0.01077561 |
| **#329** | IL7R | 464.8571 | -0.9063263 | 0.04958369 |
| **#330** | SDCBP | 116.9788 | 0.4531241 | 0.003319799 |
| **#331** | NLRC3 | 28.01044 | -0.9092369 | 0.005829222 |
| **#332** | CORO1C | 35.42359 | 0.6449872 | 0.008249031 |
| **#333** | DNAJB6 | 126.9578 | 0.4542997 | 0.003847824 |
| **#334** | VDAC3 | 65.72218 | 0.4904277 | 0.002393215 |
| **#335** | OST4 | 481.9119 | 0.4908281 | 0.01614439 |
| **#336** | LRRK2 | 58.07528 | 1.011721 | 0.0411401 |
| **#337** | TPM1 | 51.17588 | 1.280236 | 0.04843565 |
| **#338** | SNHG7 | 110.7004 | -0.6343721 | 0.02032405 |
| **#339** | CASP4 | 103.773 | 0.5479106 | 0.01474253 |
| **#340** | HLA-DRA | 1,564 | 0.5934625 | 0.03558934 |
| **#341** | PPP1R14A | 29.23303 | 0.9972217 | 0.02669359 |
| **#342** | TST | 15.97626 | 1.098784 | 0.01722103 |
| **#343** | CMIP | 70.02094 | 0.5826804 | 0.01476457 |
| **#344** | TTC9 | 17.87434 | -1.132516 | 0.001442229 |
| **#345** | GPBAR1 | 22.99524 | 0.9104809 | 0.01838127 |
| **#346** | GASK1B | 18.41867 | 0.8172363 | 0.007004033 |
| **#347** | NUTF2 | 50.61725 | 0.4886801 | 0.000518913 |
| **#348** | TYROBP | 714.0618 | 0.6724409 | 0.04268223 |
| **#349** | FGD3 | 75.10953 | -0.5515207 | 0.006917309 |
| **#350** | MGLL | 21.3708 | 1.328762 | 0.03512716 |

| | | | | |
|---|---|---|---|---|
| **#351** | ABCA7 | 26.03673 | -0.7585754 | 0.000578365 |
| **#352** | ENKUR | 25.21841 | 1.326061 | 0.03916343 |
| **#353** | HACD4 | 68.19389 | 0.503647 | 0.005040202 |
| **#354** | GPX4 | 267.9593 | 0.3800099 | 0.000468342 |
| **#355** | S100A11 | 468.0866 | 0.6788525 | 0.04226242 |
| **#356** | ACSL1 | 18.19055 | 1.116483 | 0.02338926 |
| **#357** | ANKH | 25.10602 | -0.7578573 | 0.00025718 |
| **#358** | SYTL2 | 32.98976 | -0.9164402 | 0.01202702 |
| **#359** | LTB | 814.9799 | -0.5853953 | 0.03443149 |
| **#360** | H2AFJ | 71.86243 | 0.6053715 | 0.01953513 |
| **#361** | PDE5A | 17.94235 | 1.463599 | 0.03672127 |
| **#362** | RPL37 | 5,267 | -0.3743991 | 0.003648785 |
| **#363** | OGT | 110.8372 | -0.6464358 | 0.02408419 |
| **#365** | RBM47 | 24.37079 | 0.8586609 | 0.01853186 |
| **#366** | PURA | 49.28111 | -0.6273991 | 0.007166678 |
| **#367** | CA2 | 30.05045 | 1.276063 | 0.04352627 |
| **#368** | PILRA | 63.09992 | 0.6524393 | 0.02284699 |
| **#369** | JAK2 | 33.74229 | 1.101146 | 0.0394415 |
| **#370** | ATM | 297.765 | -0.5801686 | 0.02886658 |
| **#371** | YWHAQ | 101.6018 | 0.4362421 | 0.0022862 |
| **#372** | CALCOCO2 | 69.32134 | 0.5775778 | 0.01632816 |
| **#373** | LRRC25 | 56.0806 | 0.6566746 | 0.02170603 |
| **#374** | RCAN3 | 120.0915 | -0.8098676 | 0.03838632 |
| **#375** | ZNF267 | 29.22029 | 0.7922501 | 0.0194715 |
| **#376** | RPL36 | 2,814 | -0.3757455 | 0.003987655 |
| **#377** | GNG5 | 176.9147 | 0.4740996 | 0.01264515 |
| **#378** | ATXN2 | 42.0083 | -0.5848298 | 0.002054891 |
| **#379** | LILRA2 | 28.41263 | 0.7956814 | 0.01927861 |
| **#380** | NUDT16 | 24.59181 | 0.6152953 | 0.002034963 |
| **#381** | CBX7 | 24.09121 | -0.8033632 | 0.001644793 |
| **#382** | SPINT2 | 70.67748 | 0.4960771 | 0.006066586 |
| **#383** | HDAC7 | 37.7211 | -0.606126 | 0.001524512 |

| | | | | |
|---|---|---|---|---|
| #384 | SLC4A7 | 42.26664 | -0.73834 | 0.01178644 |
| #385 | AIG1 | 19.09251 | 0.6507846 | 0.000274522 |
| #386 | THEM4 | 41.82702 | -0.845203 | 0.01795275 |
| #387 | LCP1 | 509.1719 | 0.546874 | 0.03076999 |
| #388 | AP2M1 | 119.8807 | 0.4661317 | 0.008154752 |
| #389 | BST1 | 24.81472 | 0.6998211 | 0.008063466 |
| #390 | CD63 | 183.6964 | 0.6668564 | 0.04105972 |
| #391 | ITM2B | 772.473 | 0.3783764 | 0.003690453 |
| #392 | LMNB1 | 18.6786 | 1.049043 | 0.02485275 |
| #393 | RAB7A | 94.24534 | 0.4133851 | 0.000228447 |
| #394 | CDC25B | 38.64422 | -0.7290884 | 0.01002147 |
| #395 | CTNNA1 | 34.70408 | 0.5836944 | 0.006795953 |
| #396 | MICU1 | 38.28014 | 0.5287269 | 0.002566705 |
| #397 | TNRC6C | 49.25852 | -0.7718995 | 0.01980314 |
| #398 | MAGED2 | 40.2109 | 0.5822623 | 0.009683418 |
| #399 | RBCK1 | 64.67243 | 0.5189497 | 0.009193562 |
| #400 | PDE7A | 62.86163 | -0.8484656 | 0.03033779 |
| #401 | ZRANB2 | 154.9591 | -0.505599 | 0.01494853 |
| #402 | MS4A6A | 120.4732 | 0.5615459 | 0.02506501 |
| #403 | ATF3 | 13.02271 | 1.278653 | 0.0257969 |
| #404 | SAMD4A | 7.109568 | 1.776486 | 0.02511213 |
| #405 | RILPL2 | 77.49411 | 0.580414 | 0.02195588 |
| #406 | SSX2IP | 19.19062 | 0.9082685 | 0.01842 |
| #407 | SAP25 | 26.13638 | -0.7475381 | 0.002604335 |
| #408 | LILRA1 | 29.92986 | 0.8902712 | 0.03034195 |
| #409 | CERK | 40.28819 | -0.6098056 | 0.0042033 |
| #410 | KANSL1 | 131.7404 | -0.426284 | 0.002594909 |
| #411 | HLA-C | 2,251 | 0.4342826 | 0.01738659 |
| #412 | ENO1 | 269.238 | 0.3800043 | 0.002894323 |
| #413 | PRELID1 | 221.1261 | 0.5957662 | 0.03796963 |
| #414 | RIOK3 | 116.6736 | 0.5115065 | 0.01783739 |
| #415 | AKNA | 132.7873 | -0.5002777 | 0.01353585 |

| | | | | |
|---|---|---|---|---|
| **#416** | PGK 1.00 | 195.0155 | 0.4345247 | 0.00957631 |
| **#417** | OAS2 | 26.50794 | 1.081586 | 0.03849057 |
| **#418** | LGALS9 | 59.96524 | 0.6972658 | 0.03152261 |
| **#419** | FUOM | 20.01347 | 0.6274941 | 0.001012076 |
| **#420** | ATP8B2 | 26.56311 | -1.033078 | 0.01515834 |
| **#421** | ATP6V0C | 235.4855 | 0.422208 | 0.008524925 |
| **#424** | UBE2L3 | 101.0502 | 0.399549 | 0.000387782 |
| **#425** | CDKN2D | 180.0198 | 0.5576503 | 0.03146342 |
| **#426** | TLR8 | 17.23445 | 0.862381 | 0.0142694 |
| **#427** | PRKAR1A | 92.28045 | 0.4174219 | 0.001615858 |
| **#430** | TRIB1 | 11.18684 | 0.9071967 | 0.004307349 |
| **#431** | EIF5A | 197.4762 | 0.442802 | 0.01289395 |
| **#432** | MCTP1 | 24.66125 | 0.7425969 | 0.01600203 |
| **#433** | RAB11A | 71.69516 | 0.4870417 | 0.008476983 |
| **#434** | B4GALT5 | 12.7943 | 0.8073081 | 0.003043688 |
| **#436** | CTSB | 104.7269 | 0.5338157 | 0.02271713 |
| **#437** | DYNLL1 | 171.458 | 0.4515354 | 0.0143252 |
| **#438** | RAB5C | 97.24549 | 0.4971088 | 0.01632503 |
| **#439** | NCF2 | 119.8885 | 0.4844337 | 0.01638824 |
| **#441** | NEXN | 24.90576 | 1.197443 | 0.04875963 |
| **#442** | TAP2 | 42.03097 | 0.6905238 | 0.02718557 |
| **#443** | GCC2 | 207.2801 | -0.4263605 | 0.00890817 |
| **#444** | SPTLC2 | 46.81576 | 0.6293181 | 0.02377428 |
| **#445** | MVP | 53.28336 | 0.6789568 | 0.03299177 |
| **#446** | CD1D | 23.44554 | 0.5642986 | 0.001182316 |
| **#447** | DDX17 | 349.5649 | -0.4482314 | 0.01866617 |
| **#448** | RAB13 | 7.973399 | 1.215114 | 0.01177129 |
| **#449** | HK1 | 56.1201 | 0.4544604 | 0.003336684 |
| **#450** | TPMT | 23.03722 | 0.5591221 | 0.000532683 |
| **#451** | LILRB3 | 57.41656 | 0.6117839 | 0.02745442 |
| **#452** | PMAIP1 | 47.84103 | 0.6400113 | 0.02712144 |
| **#453** | MIAT | 16.58282 | -1.014857 | 0.003336433 |

| | | | | |
|---|---|---|---|---|
| #454 | SIGIRR | 107.7381 | -0.6006923 | 0.03100332 |
| #455 | NAP1L1 | 554.7086 | 0.333417 | 0.001252823 |
| #456 | HNRNPH2 | 46.60456 | 0.4444717 | 0.000140916 |
| #457 | ZNF503 | 10.41458 | 1.033274 | 0.01239073 |
| #458 | MRI1 | 23.96315 | -0.7150705 | 0.002664664 |
| #459 | NMI | 45.95154 | 0.7727884 | 0.04184828 |
| #460 | SNHG8 | 236.0252 | -0.3619454 | 0.001610007 |
| #461 | CPVL | 128.6328 | 0.4707885 | 0.0181585 |
| #462 | UBE2J1 | 60.5399 | 0.4742389 | 0.007824378 |
| #463 | REC8 | 10.92873 | 0.7815389 | 0.000394431 |
| #464 | NUCB1 | 70.30514 | 0.5037612 | 0.0153263 |
| #465 | PPIF | 23.47084 | 0.7685018 | 0.02167191 |
| #466 | DEK | 232.3872 | 0.3499076 | 0.001563948 |
| #467 | GRINA | 49.97576 | 0.6734593 | 0.03449516 |
| #468 | GRAMD1A | 39.82402 | -0.545587 | 0.003069784 |
| #469 | DGKZ | 92.12355 | -0.509656 | 0.01549006 |
| #470 | FTL | 4,134 | 0.4744083 | 0.03552899 |
| #471 | HSP90AA1 | 557.5443 | 0.3670692 | 0.007468555 |
| #473 | ZMYND8 | 49.58492 | -0.5620769 | 0.009374211 |
| #475 | CREBZF | 54.02566 | -0.5644402 | 0.01242088 |
| #476 | ANKZF1 | 31.31 | -0.5787342 | 0.001458058 |
| #477 | SPPL2A | 45.96303 | 0.4443287 | 0.00056545 |
| #478 | GUK1 | 203.144 | 0.4340637 | 0.01707385 |
| #479 | RPL38 | 1,983 | -0.4078027 | 0.01933446 |
| #480 | DUSP3 | 14.41317 | 0.8287533 | 0.01198332 |
| #481 | TMEM63A | 40.91283 | -0.7165525 | 0.01992979 |
| #482 | HLA-DMB | 119.1748 | 0.593537 | 0.04356614 |
| #483 | GAS5 | 384.489 | -0.3972117 | 0.01173021 |
| #484 | NLRP3 | 22.82666 | 0.7885107 | 0.02465049 |
| #485 | CMPK2 | 6.666507 | 1.566498 | 0.02516382 |
| #486 | RASGRP1 | 69.08911 | -0.5301063 | 0.01436078 |
| #487 | RNF7 | 86.66172 | 0.4254537 | 0.006373932 |

| | | | | |
|---|---|---|---|---|
| #488 | CUL1 | 33.46076 | 0.6604712 | 0.02411799 |
| #489 | CTSS | 1,151 | 0.4881597 | 0.03793699 |
| #490 | DUSP6 | 40.16407 | 0.7854454 | 0.04295753 |
| #491 | GLG1 | 111.2249 | -0.3814344 | 0.000401181 |
| #493 | EIF2AK1 | 77.76748 | 0.5053981 | 0.0198331 |
| #494 | XPNPEP1 | 22.93262 | 0.8120205 | 0.02767529 |
| #495 | OASL | 5.993366 | 1.431438 | 0.01488845 |
| #496 | LMO2 | 53.27656 | 0.6088544 | 0.03038362 |
| #497 | HERC2 | 50.41593 | -0.6042136 | 0.0170243 |
| #498 | NUB1 | 96.8978 | 0.5083632 | 0.0250733 |
| #499 | SENCR | 7.980117 | 1.151612 | 0.0120333 |
| #500 | DUSP5 | 14.13871 | 0.7128778 | 0.003500231 |
| #501 | KIF21B | 17.11091 | -0.8589283 | 0.002058308 |
| #503 | CLEC2B | 139.3719 | 0.4253023 | 0.01289383 |
| #504 | LILRB2 | 80.41848 | 0.5993227 | 0.03981213 |
| #505 | PIP4P2 | 22.0178 | 0.562895 | 0.002056005 |
| #506 | ZFP14 | 23.13739 | -0.6418743 | 0.000510889 |
| #507 | CBFA2T2 | 22.55892 | -0.6663268 | 0.001076943 |
| #508 | DSE | 26.94387 | 0.7124425 | 0.02509757 |
| #509 | ITGB1 | 215.2717 | 0.3307019 | 0.000277436 |
| #510 | STX2 | 23.65051 | -0.7067688 | 0.004242669 |
| #511 | ARID1B | 191.9373 | -0.3819887 | 0.006148652 |
| #512 | SYNRG | 89.68843 | -0.4626581 | 0.01137584 |
| #513 | UBE2F | 52.81129 | 0.5440347 | 0.02171879 |
| #514 | ODF2L | 70.65196 | -0.4807916 | 0.009464937 |
| #515 | TPM3 | 587.9751 | 0.4235659 | 0.02398677 |
| #516 | EPB41 | 195.7914 | -0.5199811 | 0.03587643 |
| #517 | ZCCHC17 | 45.7636 | 0.4783831 | 0.007443252 |
| #518 | SAMD1 | 21.6512 | -0.6642241 | 0.000956017 |
| #519 | PDCD7 | 47.59771 | -0.8678031 | 0.04319084 |
| #520 | FTX | 81.92166 | -0.5482509 | 0.0250882 |
| #521 | GNA15 | 15.82395 | 0.8349932 | 0.02006622 |

| | | | | |
|---|---|---|---|---|
| #522 | LYPLAL1 | 22.24921 | 0.6154331 | 0.009581391 |
| #523 | NDUFA6 | 82.05165 | 0.4370984 | 0.01068956 |
| #524 | IL15 | 11.18073 | 0.9855434 | 0.01862964 |
| #525 | SEC14L1 | 65.6424 | 0.5083096 | 0.02145996 |
| #527 | SNHG12 | 22.70659 | -0.6654939 | 0.00208324 |
| #528 | IFI27L2 | 75.1539 | 0.430391 | 0.008380906 |
| #529 | UBE2E3 | 52.73978 | 0.4234669 | 0.002336737 |
| #530 | CLTA | 87.49219 | 0.3857437 | 0.002953226 |
| #531 | INPP4A | 47.53225 | -0.4741532 | 0.002418777 |
| #532 | SLC40A1 | 36.3648 | 0.7071422 | 0.0388962 |
| #533 | TRIM25 | 22.33685 | 0.6044492 | 0.008743817 |
| #534 | TRIM73 | 35.68572 | -0.8759768 | 0.03297357 |
| #536 | NUP210 | 54.03763 | -0.4610913 | 0.003110854 |
| #538 | PIK3AP1 | 55.88359 | 0.5796147 | 0.03250738 |
| #539 | CYTH1 | 165.3565 | -0.3952714 | 0.008365163 |
| #540 | ASXL1 | 65.64826 | -0.6548167 | 0.03687147 |
| #541 | VRK2 | 17.85328 | 0.7759292 | 0.02082199 |
| #542 | IRF3 | 80.27183 | -0.4841878 | 0.01490773 |
| #543 | DNASE1 | 19.05973 | -0.9343852 | 0.01115429 |
| #544 | TMSB10 | 4,840 | 0.3289082 | 0.007674264 |
| #545 | NUMA1 | 66.79415 | -0.4623497 | 0.007626016 |
| #546 | RAB4A | 57.83648 | 0.3964755 | 0.000530672 |
| #547 | DYSF | 5.658208 | 1.67955 | 0.03096319 |
| #550 | NPHP3 | 26.09505 | -0.6099743 | 0.003023217 |
| #551 | ZBTB25 | 35.98382 | -0.6509382 | 0.01692056 |
| #553 | EIF4A2 | 278.2845 | -0.3446673 | 0.00409727 |
| #554 | KHDC4 | 39.34555 | -0.5174981 | 0.004330039 |
| #555 | SLC35E2B | 17.93927 | -0.7233688 | 0.000917714 |
| #556 | ARMCX6 | 25.79008 | 0.5089554 | 0.00227409 |
| #557 | CDC14A | 56.7321 | -0.5893771 | 0.02615925 |
| #558 | KIAA0355 | 42.28795 | -0.5835047 | 0.01560485 |
| #559 | ZNF276 | 51.26214 | -0.4371554 | 0.000882589 |

| | | | | |
|---|---|---|---|---|
| **#560** | HBEGF | 19.91995 | 0.9471469 | 0.04449296 |
| **#561** | HOTAIRM1 | 39.6499 | 0.5614856 | 0.02245297 |
| **#562** | SELENOT | 73.42013 | 0.4332865 | 0.01099727 |
| **#564** | CAMTA1 | 43.90912 | 0.4619117 | 0.006802998 |
| **#565** | XPC | 54.69584 | -0.4513649 | 0.003345836 |
| **#567** | DRAP1 | 178.4818 | 0.4254139 | 0.02242562 |
| **#568** | PRKAR1B | 12.41648 | 0.9227999 | 0.02207691 |
| **#569** | KATNBL1 | 41.06704 | -0.4881103 | 0.002743921 |
| **#570** | TMEM179B | 44.68407 | 0.4989166 | 0.01464824 |
| **#571** | BISPR | 15.23675 | 0.7113651 | 0.01026853 |
| **#572** | FOXP1 | 356.1775 | -0.412609 | 0.02309793 |
| **#573** | VAMP3 | 32.01752 | 0.4972747 | 0.006272037 |
| **#574** | CYTOR | 49.38432 | 0.5116108 | 0.02060847 |
| **#575** | CFAP36 | 25.15606 | -0.670609 | 0.007667737 |
| **#576** | SNHG6 | 408.9613 | -0.3225267 | 0.003559455 |
| **#578** | RBL2 | 114.5912 | -0.5138959 | 0.03261456 |
| **#581** | SFI1 | 40.95725 | -0.6187364 | 0.02102394 |
| **#582** | ATP6V0E1 | 201.1162 | 0.4074974 | 0.02054456 |
| **#583** | CREG1 | 31.59404 | 0.5439335 | 0.01408604 |
| **#584** | PIM1 | 48.85114 | 0.6029301 | 0.03895112 |
| **#585** | ARL8B | 31.03461 | 0.4855782 | 0.004006346 |
| **#587** | LTBP3 | 23.15866 | -0.8550643 | 0.01896308 |
| **#588** | NFIA | 23.73371 | -0.7204665 | 0.01034383 |
| **#589** | PTTG1IP | 59.9103 | 0.4655668 | 0.01619133 |
| **#590** | PSMB1 | 162.2829 | 0.3624616 | 0.007606992 |
| **#591** | VASP | 162.6036 | 0.4971661 | 0.0424303 |
| **#592** | UQCRH | 293.1955 | 0.3066223 | 0.001041777 |
| **#593** | BCL2L1 | 17.61727 | 0.8019908 | 0.02768865 |
| **#594** | ATP6V0D1 | 71.69791 | 0.506038 | 0.02931355 |
| **#595** | LY96 | 26.61228 | 0.583287 | 0.01645289 |
| **#596** | CARD11 | 31.81636 | -0.5567137 | 0.005377532 |
| **#597** | PSAP | 629.1832 | 0.4347083 | 0.03706855 |

| | | | | |
|---|---|---|---|---|
| **#599** | ANXA5 | 179.8846 | 0.5092714 | 0.04809307 |
| **#600** | SNHG19 | 15.35142 | -0.7616234 | 5.49659E-05 |
| **#601** | EMILIN2 | 47.2294 | 0.576957 | 0.03502517 |
| **#602** | PPP1R3E | 19.25681 | -0.834831 | 0.01036506 |
| **#603** | BRD8 | 20.52695 | -0.6560594 | 0.002411401 |
| **#605** | CDK13 | 132.1884 | -0.5050162 | 0.03591997 |
| **#606** | TRAFD1 | 14.6784 | 0.9671837 | 0.03794173 |
| **#607** | ANKRD36C | 43.95244 | -0.7917693 | 0.04523167 |
| **#609** | RPL36A | 2,081 | -0.3044457 | 0.004788347 |
| **#610** | AP5B1 | 21.77633 | 0.6914834 | 0.02572681 |
| **#611** | CCDC66 | 54.07354 | -0.5349351 | 0.01931258 |
| **#612** | UBA7 | 27.55347 | 0.5520094 | 0.01333539 |
| **#613** | INTS6 | 66.93065 | -0.4687256 | 0.01397636 |
| **#614** | STAT5B | 44.5843 | -0.5134215 | 0.01104319 |
| **#615** | ATP6V1A | 26.62881 | 0.5813381 | 0.01809547 |
| **#617** | DERA | 23.11423 | 0.4880964 | 0.00022063 |
| **#618** | CD36 | 72.85743 | 0.5531915 | 0.04449665 |
| **#619** | PTGS2 | 14.07019 | 0.9143216 | 0.0332234 |
| **#620** | CNIH4 | 28.72744 | 0.4539048 | 0.000471405 |
| **#621** | PSMB2 | 59.51916 | 0.4194761 | 0.008308769 |
| **#623** | CEP290 | 31.58469 | -0.6342881 | 0.01700801 |
| **#624** | NFIL3 | 23.04597 | 0.7913013 | 0.04325072 |
| **#625** | SUN2 | 142.3581 | -0.4968921 | 0.03927005 |
| **#626** | KAT6B | 106.4579 | -0.4813008 | 0.02853339 |
| **#627** | WBP2 | 58.19528 | 0.5012252 | 0.02710145 |
| **#628** | S100PBP | 24.92146 | -0.6038594 | 0.005307633 |
| **#629** | SCAI | 17.64192 | -0.6715521 | 0.000884265 |
| **#631** | SP110 | 139.1494 | 0.471123 | 0.03924044 |
| **#632** | RSBN1L | 89.78621 | -0.4249802 | 0.0134562 |
| **#634** | GKAP1 | 15.46629 | -0.7258914 | 0.000401867 |
| **#635** | CBLB | 53.31122 | -0.5224189 | 0.01984422 |
| **#636** | ZFAS1 | 454.6416 | -0.2835322 | 2.77489E-05 |

| | | | | |
|---|---|---|---|---|
| **#637** | GPAT3 | 13.4464 | 0.6879296 | 0.008096011 |
| **#638** | TMEM71 | 33.82814 | -0.5592886 | 0.01129211 |
| **#640** | PHF1 | 40.04828 | -0.4997096 | 0.007967973 |
| **#641** | PCMT1 | 59.54088 | 0.4034664 | 0.00713367 |
| **#642** | MAP2K6 | 16.63501 | -0.7016052 | 0.001434274 |
| **#643** | RPL34 | 5,702 | -0.3223725 | 0.0144322 |
| **#644** | MSRB2 | 18.24775 | 0.7195304 | 0.02669775 |
| **#645** | CFL1 | 917.3552 | 0.3831423 | 0.02964992 |
| **#646** | DOK3 | 18.31702 | 0.6180477 | 0.01307744 |
| **#648** | FAS | 11.89899 | 0.8256538 | 0.0191895 |
| **#649** | F11R | 20.18233 | 0.5332011 | 0.003699658 |
| **#650** | NDUFA9 | 37.61566 | 0.4076426 | 0.000436908 |
| **#652** | GNAQ | 58.08702 | 0.4078245 | 0.00797168 |
| **#653** | CNTLN | 11.94909 | 0.7263169 | 0.00885901 |
| **#654** | CABIN1 | 32.48984 | -0.4863562 | 0.001521446 |
| **#655** | ZFYVE21 | 10.20036 | 0.7726799 | 0.007537538 |
| **#656** | TNS3 | 12.1563 | 0.739963 | 0.01149561 |
| **#658** | PPP6R2 | 55.38053 | -0.5462151 | 0.02781268 |
| **#659** | ATP6V1B2 | 35.12081 | 0.5364132 | 0.02321096 |
| **#661** | FGFR1OP | 14.66217 | -0.7348496 | 0.000429865 |
| **#662** | PCNX4 | 32.66202 | -0.4845636 | 0.001662069 |
| **#663** | CHMP2A | 91.72536 | 0.4159621 | 0.01955674 |
| **#664** | RHEB | 39.2517 | 0.4078942 | 0.001486596 |
| **#665** | MSRB1 | 24.00685 | 0.7505894 | 0.04570504 |
| **#667** | HIST2H2BE | 12.89115 | 0.7950762 | 0.02113134 |
| **#668** | PSMA3 | 55.16922 | 0.3934501 | 0.004735095 |
| **#669** | ADAR | 137.8645 | 0.4255603 | 0.02954849 |
| **#673** | PLEKHB2 | 44.43113 | 0.418024 | 0.005450046 |
| **#675** | TRGC2 | 33.94825 | -0.6674508 | 0.02889835 |
| **#676** | DNAJA1 | 120.8405 | 0.4671595 | 0.04190579 |
| **#680** | F2R | 19.52752 | 0.7594753 | 0.03900087 |
| **#681** | NFATC3 | 41.89768 | -0.4712365 | 0.007084354 |

| | | | | |
|---|---|---|---|---|
| #682 | NSUN5 | 33.00278 | -0.4657499 | 0.000848924 |
| #683 | TNRC6A | 60.37616 | -0.4122706 | 0.005237888 |
| #684 | STRN4 | 20.54669 | 0.544841 | 0.007592414 |
| #685 | RGS1 | 19.60353 | 0.8421575 | 0.04981834 |
| #687 | HMG20B | 38.93436 | 0.4342145 | 0.00699842 |
| #688 | RNF19B | 15.56975 | 0.5816526 | 0.004429816 |
| #689 | CYFIP2 | 94.1194 | -0.372881 | 0.006812609 |
| #691 | TRAF1 | 11.67182 | -0.9353653 | 0.002276247 |
| #692 | TENT5A | 43.74 | 0.5629837 | 0.04195131 |
| #693 | TECPR1 | 31.65127 | -0.4980436 | 0.003736204 |
| #694 | RBM14 | 27.71395 | -0.4985945 | 0.001098598 |
| #696 | RGS14 | 32.90908 | -0.5930131 | 0.01992888 |
| #698 | AGAP2 | 13.93404 | -0.7665516 | 0.001226976 |
| #699 | TICAM2 | 7.216082 | 0.9846807 | 0.01330748 |
| #701 | ISOC2 | 13.69827 | 0.6161607 | 0.005230052 |
| #702 | MAPRE1 | 57.50595 | 0.4130403 | 0.01239675 |
| #703 | PCSK7 | 156.8031 | -0.3872912 | 0.01966902 |
| #704 | TTC39C | 69.63882 | -0.5592042 | 0.04527107 |
| #705 | GTF3C6 | 61.3168 | 0.4031853 | 0.01124603 |
| #706 | RHOU | 9.303525 | 0.7931728 | 0.008217159 |
| #707 | AGPAT1 | 25.75998 | 0.5516717 | 0.01965437 |
| #708 | ENOSF1 | 18.23765 | -0.6781322 | 0.005438097 |
| #709 | TOLLIP | 24.88427 | 0.5350577 | 0.01471255 |
| #710 | EHMT2 | 27.76567 | -0.4998442 | 0.0016758 |
| #712 | DNAJC15 | 69.54076 | 0.4351652 | 0.02357082 |
| #713 | HERC4 | 53.58119 | -0.5177943 | 0.02606741 |
| #714 | PRDX3 | 31.43543 | 0.5103215 | 0.01911632 |
| #715 | HNRNPH1 | 322.2129 | -0.3494173 | 0.0186497 |
| #716 | PGD | 51.80844 | 0.4931985 | 0.03154875 |
| #717 | P2RX7 | 12.06248 | 0.8854269 | 0.03241423 |
| #718 | PTPRJ | 52.98186 | 0.4984686 | 0.03442599 |
| #719 | PRDM1 | 29.95488 | 0.6081655 | 0.03850091 |

| | | | | |
|---|---|---|---|---|
| **#720** | ATP6V0B | 139.2925 | 0.42013 | 0.0331237 |
| **#721** | SNRNP70 | 103.8358 | -0.3745167 | 0.01039187 |
| **#722** | SEPTIN4 | 4.058672 | 1.590051 | 0.02650783 |
| **#723** | CAPZB | 226.6661 | 0.3319146 | 0.0127217 |
| **#725** | HIGD1A | 28.55285 | 0.4552117 | 0.00473694 |
| **#727** | CGAS | 14.49601 | 0.7525521 | 0.02654251 |
| **#728** | SLC5A3 | 26.68502 | -0.5067003 | 0.002195773 |
| **#729** | TJP2 | 9.855168 | 0.7203085 | 0.005071814 |
| **#731** | VAMP7 | 20.15623 | 0.4810984 | 0.000820728 |
| **#732** | ABCA5 | 15.05987 | -0.7378831 | 0.00362174 |
| **#734** | SPTAN1 | 89.67875 | -0.5193626 | 0.04750947 |
| **#735** | ADAP2 | 22.36549 | 0.6183031 | 0.02940183 |
| **#736** | IRAK3 | 46.69538 | 0.5240777 | 0.03966482 |
| **#737** | APOL1 | 14.40647 | 0.8684795 | 0.04427042 |
| **#738** | TOB2 | 23.35537 | -0.6697716 | 0.01769393 |
| **#740** | YWHAE | 109.9141 | 0.4103178 | 0.02828429 |
| **#741** | NSUN6 | 24.00774 | -0.5659587 | 0.006780377 |
| **#742** | PRH1 | 26.44896 | -0.4933927 | 0.00119432 |
| **#745** | VAMP8 | 196.0954 | 0.3931457 | 0.0333099 |
| **#747** | GADD45A | 33.48274 | 0.5897349 | 0.0440078 |
| **#748** | SCFD2 | 8.471621 | 1.003179 | 0.02720247 |
| **#749** | DNMT3A | 26.01293 | -0.5521895 | 0.007626293 |
| **#751** | PASK | 23.54683 | -0.7184978 | 0.02389552 |
| **#753** | KYNU | 23.79314 | 0.575535 | 0.02524083 |
| **#755** | ATP6V1F | 124.1755 | 0.385428 | 0.02474841 |
| **#756** | TM2D2 | 13.03222 | 0.6913146 | 0.01570492 |
| **#757** | STAT3 | 130.6241 | 0.3733133 | 0.02163116 |
| **#758** | PRDX5 | 132.5697 | 0.3655697 | 0.01976268 |
| **#759** | CFDP1 | 77.65998 | -0.3388209 | 0.00085013 |
| **#760** | PTPN4 | 74.31008 | -0.5077374 | 0.04207678 |
| **#761** | CALR | 219.7795 | 0.380175 | 0.03241728 |
| **#762** | MFSD4B | 17.88236 | -0.5962316 | 0.001310698 |

| | | | | |
|---|---|---|---|---|
| #763 | UPF2 | 88.71243 | -0.3567459 | 0.00648275 |
| #764 | RPS28 | 4,876 | -0.3067086 | 0.01836311 |
| #765 | TRAM1 | 119.1427 | 0.3245678 | 0.00629792 |
| #766 | GON4L | 60.29715 | -0.3901024 | 0.005808301 |
| #767 | RPL37A | 3,331 | -0.3173418 | 0.0216876 |
| #768 | PDIA6 | 92.29741 | 0.3562302 | 0.01122074 |
| #769 | ZNF721 | 48.09193 | -0.506201 | 0.0253635 |
| #770 | LAMP2 | 54.27128 | 0.4209896 | 0.0180061 |
| #771 | AKT3 | 32.73801 | -0.5815266 | 0.02416892 |
| #772 | LRRC59 | 21.73066 | 0.4618815 | 0.001554149 |
| #773 | LSM 1.00 | 43.32106 | 0.4020507 | 0.007119543 |
| #774 | SNX3 | 190.8579 | 0.3372594 | 0.01645136 |
| #775 | ABHD16A | 27.4245 | 0.4829635 | 0.01192668 |
| #776 | CD96 | 64.06588 | -0.5018942 | 0.03669826 |
| #778 | GLRX | 113.6568 | 0.4444534 | 0.04542742 |
| #779 | SLC1A5 | 16.81545 | 0.6173651 | 0.01878706 |
| #780 | RAB10 | 62.67834 | 0.3770967 | 0.009970346 |
| #781 | CTSO | 16.16528 | 0.5618972 | 0.007523355 |
| #783 | PWWP3A | 24.24161 | -0.503448 | 0.001560298 |
| #784 | RPL31 | 1,208 | -0.3069946 | 0.0165474 |
| #785 | RNF213 | 364.0129 | 0.4097759 | 0.04965275 |
| #786 | TPI1 | 187.3003 | 0.4064686 | 0.0411724 |
| #787 | SRA1 | 36.09132 | 0.4188645 | 0.007006365 |
| #788 | NOD2 | 8.786818 | 0.7560559 | 0.006807285 |
| #791 | CDC42 | 530.631 | 0.2790292 | 0.006479531 |
| #792 | TMEM259 | 79.67009 | -0.4897203 | 0.0419348 |
| #794 | R3HDM2 | 85.75457 | -0.4720612 | 0.03962604 |
| #795 | ARRDC2 | 40.94895 | -0.4796832 | 0.01485544 |
| #796 | AP2A2 | 32.4339 | -0.436387 | 0.000850439 |
| #797 | MSN | 335.6302 | 0.3439444 | 0.02544979 |
| #798 | RBM6 | 87.87487 | -0.3275506 | 0.00148767 |
| #799 | ZC3H8 | 34.07008 | -0.454691 | 0.004010379 |

| | | | | |
|---|---|---|---|---|
| **#800** | EAF1 | 15.38628 | 0.5461826 | 0.004370708 |
| **#802** | ASGR2 | 11.86382 | 0.950677 | 0.04878399 |
| **#806** | SLC25A45 | 39.9381 | -0.449633 | 0.007893448 |
| **#808** | RBP7 | 32.47567 | 0.5647956 | 0.04213292 |
| **#809** | BLOC1S1 | 152.5478 | 0.3777696 | 0.02888007 |
| **#811** | TSEN34 | 22.77102 | 0.480735 | 0.006806222 |
| **#812** | SLC10A3 | 11.64679 | 0.6142571 | 0.003909058 |
| **#813** | COQ10A | 14.12233 | -0.8495888 | 0.01227907 |
| **#816** | RCN1 | 11.89829 | 0.7028434 | 0.01681453 |
| **#817** | PDE4DIP | 20.62028 | -0.577221 | 0.005752577 |
| **#818** | TRIM52 | 37.3892 | -0.4648641 | 0.009790176 |
| **#819** | ARL6IP5 | 261.8628 | 0.3026094 | 0.01042474 |
| **#820** | ANKRD13A | 50.17964 | 0.4255655 | 0.01994946 |
| **#821** | CAPZA1 | 128.3425 | 0.2979464 | 0.00283775 |
| **#822** | ELF4 | 14.13961 | 0.5520234 | 0.003381353 |
| **#823** | POLR2E | 62.64715 | 0.4184891 | 0.0250566 |
| **#824** | CA5B | 34.59767 | -0.4855625 | 0.01119256 |
| **#825** | UBE2D3 | 280.532 | 0.2691846 | 0.002485802 |
| **#826** | ARHGAP15 | 88.66517 | -0.4364602 | 0.03364219 |
| **#827** | SPG7 | 71.59754 | -0.3936448 | 0.01373793 |
| **#828** | IKBKG | 24.04615 | 0.5346686 | 0.02181365 |
| **#829** | ACPP | 9.946032 | 0.9310707 | 0.03774985 |
| **#830** | AGPAT3 | 34.73572 | 0.4581901 | 0.01822282 |
| **#831** | KLHDC7B | 4.624617 | 1.059039 | 0.004834188 |
| **#832** | TMEM116 | 11.08941 | -0.8177103 | 0.001467764 |
| **#833** | GYG1 | 24.35245 | 0.4894858 | 0.01207377 |
| **#834** | LRCH3 | 32.21663 | -0.4676426 | 0.006611014 |
| **#837** | PML | 19.84474 | 0.6422194 | 0.037726 |
| **#840** | DUS1L | 48.29714 | -0.4914659 | 0.02701839 |
| **#841** | CEP250 | 15.37472 | -0.6179968 | 0.000937276 |
| **#842** | MAP1LC3B | 122.7306 | 0.3696833 | 0.02564365 |
| **#843** | RNF149 | 108.8974 | 0.4161633 | 0.04013555 |

| | | | | |
|---|---|---|---|---|
| **#844** | VCPIP1 | 35.54875 | 0.4249088 | 0.01100013 |
| **#846** | MPST | 24.10652 | 0.5143382 | 0.01930004 |
| **#847** | MRNIP | 20.06968 | -0.6585258 | 0.01668413 |
| **#848** | RPL39 | 4,957 | -0.3097384 | 0.02597985 |
| **#849** | HELZ2 | 16.90689 | 0.7057122 | 0.04068304 |
| **#850** | SLC25A42 | 18.15319 | -0.5790188 | 0.003091095 |
| **#851** | HPGD | 12.69923 | 0.6852977 | 0.02061403 |
| **#852** | PNISR | 459.6703 | -0.3101031 | 0.01917125 |
| **#853** | ECHS1 | 49.49323 | 0.359698 | 0.004521437 |
| **#854** | TECR | 77.78839 | -0.4121024 | 0.02390644 |
| **#855** | KIF22 | 28.24657 | -0.5469433 | 0.01702614 |
| **#856** | DDX5 | 1,210 | -0.2658147 | 0.007620356 |
| **#857** | STAC3 | 9.800049 | 0.7285674 | 0.01358746 |
| **#858** | MCM7 | 19.93174 | -0.5343013 | 0.001841819 |
| **#860** | SWI5 | 18.9082 | 0.6155906 | 0.03224242 |
| **#861** | RTN3 | 79.68736 | 0.3504511 | 0.01327474 |
| **#862** | DECR1 | 58.40961 | 0.3542397 | 0.007210118 |
| **#863** | RPL35 | 1,979 | -0.2383042 | 0.002296969 |
| **#864** | EPHB6 | 21.48318 | -0.5631188 | 0.008533088 |
| **#865** | ACOT9 | 22.95039 | 0.5217439 | 0.02064039 |
| **#866** | PARK7 | 202.9912 | 0.3304937 | 0.02185143 |
| **#867** | MINDY2 | 25.88841 | -0.4550515 | 0.000746869 |
| **#868** | RALGAPA1 | 38.18357 | -0.5570024 | 0.03615257 |
| **#869** | ZKSCAN1 | 51.05741 | -0.3768945 | 0.004120248 |
| **#870** | CEP126 | 12.57391 | -0.7875416 | 0.005852601 |
| **#872** | ORMDL1 | 117.6194 | -0.2927756 | 0.001135081 |
| **#873** | DYRK2 | 55.69929 | -0.4885521 | 0.03584638 |
| **#875** | RPL22 | 2,120 | -0.2851635 | 0.01688684 |
| **#876** | USP53 | 16.94594 | -0.8136974 | 0.02595243 |
| **#877** | FYB1 | 483.6165 | 0.3673031 | 0.04678518 |
| **#879** | HLA-A | 1,941 | 0.2801483 | 0.01558282 |
| **#880** | ZNF335 | 21.50238 | -0.5048328 | 0.001585345 |

| | | | | |
|---|---|---|---|---|
| #881 | WIPI1 | 11.40713 | 0.8546075 | 0.04258463 |
| #882 | PTOV1 | 30.28008 | -0.4555735 | 0.004886285 |
| #883 | CASP7 | 10.10416 | 0.8045982 | 0.02702976 |
| #884 | KMT2B | 34.02928 | -0.5645341 | 0.03283768 |
| #885 | PSD4 | 34.34877 | -0.4941377 | 0.0171049 |
| #886 | SRSF11 | 194.076 | -0.319493 | 0.01521661 |
| #888 | LPCAT4 | 11.4672 | -0.7641722 | 0.001924822 |
| #889 | ACOT7 | 5.402862 | 0.8528923 | 0.001281137 |
| #890 | ZNF439 | 18.64353 | -0.6171529 | 0.01055377 |
| #891 | TP53I13 | 28.4324 | -0.6608987 | 0.04171828 |
| #892 | ZNF559 | 13.02137 | -0.7097402 | 0.003587125 |
| #894 | ATXN7L1 | 20.08571 | -0.5216167 | 0.002220734 |
| #895 | FBXL20 | 33.17701 | -0.4571332 | 0.008764175 |
| #896 | CAMK2G | 27.55914 | -0.4376522 | 0.001013333 |
| #899 | PTPRE | 121.7057 | 0.4014274 | 0.04420025 |
| #900 | RPL23A | 3,236 | -0.2528385 | 0.007966896 |
| #901 | AP1S2 | 200.4336 | 0.3794774 | 0.04552694 |
| #902 | KIAA2026 | 67.6791 | -0.4208465 | 0.02613082 |
| #903 | PPWD1 | 32.45684 | -0.5256705 | 0.02385112 |
| #904 | WDR37 | 22.90549 | -0.4725232 | 0.000793655 |
| #905 | PLIN3 | 32.85872 | 0.4801336 | 0.02858374 |
| #906 | SHISAL2A | 18.77361 | -0.7557268 | 0.02826469 |
| #908 | ANKS3 | 10.59302 | -0.775002 | 0.001239202 |
| #909 | DENND5A | 24.66416 | 0.4983069 | 0.02163867 |
| #910 | ZBTB44 | 62.61894 | -0.3563621 | 0.00673121 |
| #911 | GABARAPL2 | 112.6963 | 0.2819534 | 0.002123176 |
| #913 | MTRF1L | 21.20996 | 0.4418476 | 0.002946785 |
| #914 | HHEX | 37.05824 | 0.4619225 | 0.0298668 |
| #916 | GRAMD1B | 9.415022 | 0.6473026 | 0.004583896 |
| #918 | SETD5 | 47.93754 | -0.3882593 | 0.007445098 |
| #921 | CD5 | 25.76979 | -0.6322886 | 0.03406816 |
| #922 | RASA3 | 66.82351 | -0.400388 | 0.0207115 |

| | | | | |
|---|---|---|---|---|
| #924 | HSPA6 | 8.139328 | 1.023379 | 0.04550506 |
| #925 | CKAP4 | 24.88447 | 0.500257 | 0.02365584 |
| #926 | TAPBP | 215.961 | 0.3455873 | 0.03443968 |
| #927 | PDP1 | 26.08194 | 0.4686563 | 0.01740005 |
| #928 | TMEM167A | 61.38092 | 0.3202125 | 0.002952445 |
| #930 | EZH1 | 36.20297 | -0.5260451 | 0.0323911 |
| #931 | ZNF44 | 28.04183 | -0.4766225 | 0.008487865 |
| #932 | HEATR5B | 28.42963 | -0.6107674 | 0.037464 |
| #933 | RPL32 | 5,767 | -0.2976092 | 0.02827454 |
| #934 | AKAP17A | 63.66388 | -0.4069551 | 0.02241418 |
| #935 | CERS2 | 40.21406 | 0.3443361 | 0.001004387 |
| #937 | ZNF827 | 14.63333 | -0.6642676 | 0.006669199 |
| #939 | DNPEP | 31.41071 | 0.538642 | 0.04923691 |
| #941 | BCKDK | 19.04802 | 0.5710426 | 0.02931736 |
| #944 | VAMP1 | 13.5268 | -0.6838286 | 0.00527422 |
| #945 | WWP1 | 49.40219 | -0.4811115 | 0.03661836 |
| #946 | PPM1M | 30.20744 | -0.472702 | 0.01155624 |
| #948 | PLXND1 | 22.03342 | -0.5793182 | 0.01811801 |
| #949 | NPTN | 42.75711 | 0.4593831 | 0.03770874 |
| #951 | RGCC | 39.54234 | -0.5625676 | 0.04877221 |
| #953 | TSC2 | 29.04281 | -0.4521719 | 0.006314618 |
| #955 | EIF5B | 132.7835 | -0.2753125 | 0.00173135 |
| #956 | FLYWCH1 | 9.911101 | -0.8194143 | 0.002646564 |
| #957 | RUFY3 | 38.292 | 0.4885635 | 0.04329033 |
| #958 | MED17 | 46.49661 | -0.4080058 | 0.01360609 |
| #959 | R3HDM4 | 56.18854 | 0.4090624 | 0.03095122 |
| #960 | TRIM65 | 11.04618 | -0.7077423 | 0.000924709 |
| #967 | WASHC2C | 32.82325 | 0.4136295 | 0.0131689 |
| #968 | RPL5 | 2,644 | -0.2686466 | 0.0178174 |
| #971 | PCNX2 | 14.11021 | -0.7916269 | 0.01884227 |
| #972 | THOC3 | 31.32765 | -0.5320934 | 0.02828048 |
| #973 | FARP2 | 8.551023 | 0.8870791 | 0.03702691 |

| | | | | |
|---|---|---|---|---|
| **#974** | TPCN1 | 18.12343 | -0.5257387 | 0.002010514 |
| **#977** | TARSL2 | 34.40759 | -0.507053 | 0.02814792 |
| **#978** | SYNGAP1 | 11.63919 | -0.84253 | 0.01186991 |
| **#980** | GHITM | 109.155 | 0.2703096 | 0.001366718 |
| **#983** | CYB561D1 | 11.41432 | -0.6847416 | 0.001321607 |
| **#984** | RHOQ | 41.71724 | 0.4291459 | 0.02918238 |
| **#985** | MTHFD2L | 20.81393 | 0.435769 | 0.004097114 |
| **#986** | PRPF38B | 162.2685 | -0.270708 | 0.003556009 |
| **#988** | ATP6V0E2 | 31.4762 | -0.511989 | 0.02593383 |
| **#990** | DIDO1 | 47.21831 | -0.3898763 | 0.01104475 |
| **#991** | RBM3 | 283.3812 | 0.3117471 | 0.02827893 |
| **#992** | RPS15A | 4,829 | -0.271585 | 0.02196994 |
| **#993** | P4HTM | 15.14104 | -0.6677196 | 0.0115867 |
| **#994** | PHF14 | 59.7721 | -0.387063 | 0.01922183 |
| **#995** | WDR6 | 25.56744 | -0.4608434 | 0.005816074 |
| **#996** | ANXA7 | 73.13504 | 0.2841037 | 0.000105588 |
| **#997** | RABL2B | 15.95156 | -0.5371489 | 0.0004682 |
| **#999** | CTC1 | 18.22467 | -0.7008239 | 0.02826095 |
| **#1001** | OCIAD2 | 67.76815 | -0.3962658 | 0.02617846 |
| **#1002** | HRH2 | 28.26139 | 0.4789676 | 0.03006983 |
| **#1003** | BAZ1A | 102.423 | 0.386719 | 0.04556499 |
| **#1005** | PHC1 | 14.05416 | -0.5820934 | 0.001229894 |
| **#1008** | TMC6 | 93.41519 | -0.3673997 | 0.02741303 |
| **#1011** | N4BP2L1 | 50.39046 | 0.3592942 | 0.01383553 |
| **#1013** | ABCA2 | 14.2264 | -0.852662 | 0.0281695 |
| **#1014** | PRRC2B | 71.33877 | -0.4315289 | 0.04370454 |
| **#1016** | AKR1A1 | 52.11528 | 0.4466093 | 0.04957503 |
| **#1017** | GRB2 | 190.9079 | 0.3359422 | 0.03698988 |
| **#1018** | EVL | 298.505 | -0.3254864 | 0.03555611 |
| **#1019** | SURF6 | 23.80033 | -0.4953329 | 0.01002643 |
| **#1021** | ADARB1 | 15.48749 | -0.6388474 | 0.01104129 |
| **#1024** | PSMA7 | 232.2417 | 0.308188 | 0.02649141 |

| | | | | |
|---|---|---|---|---|
| #1025 | PCNX1 | 63.25939 | -0.4444335 | 0.04342363 |
| #1026 | RNF44 | 42.3469 | -0.3634715 | 0.003490808 |
| #1027 | IL11RA | 9.405665 | -0.8709919 | 0.005730253 |
| #1028 | SH2B1 | 18.85147 | -0.4902832 | 0.00070063 |
| #1029 | SUB1 | 297.782 | 0.24012 | 0.003656691 |
| #1030 | IKBKB | 38.55927 | -0.3859228 | 0.005480042 |
| #1031 | TNRC6B | 224.194 | -0.3072539 | 0.02358021 |
| #1033 | SFMBT1 | 14.04463 | -0.6708992 | 0.009833958 |
| #1034 | PAN3 | 52.67272 | -0.3436355 | 0.004397103 |
| #1035 | SF1 | 370.1282 | -0.2823143 | 0.01959763 |
| #1036 | GABPB2 | 31.78508 | -0.4559838 | 0.01558082 |
| #1037 | RPL35A | 3,279 | -0.252817 | 0.01628112 |
| #1039 | BSCL2 | 16.30925 | 0.4801229 | 0.006170983 |
| #1040 | KIAA0513 | 18.59744 | 0.5049968 | 0.01830747 |
| #1041 | CEP164 | 27.87519 | -0.4174604 | 0.002620729 |
| #1042 | DAZAP2 | 234.4832 | 0.2662045 | 0.0100876 |
| #1043 | RPAP2 | 44.26896 | -0.4951198 | 0.04500683 |
| #1045 | CBR1 | 22.68566 | 0.532092 | 0.03846275 |
| #1047 | AMMECR1 | 15.33074 | -0.5353923 | 0.000521612 |
| #1049 | HEBP1 | 22.41472 | 0.4320051 | 0.008034564 |
| #1050 | VPS29 | 81.02858 | 0.2725578 | 0.00035627 |
| #1051 | DENND11 | 21.1559 | -0.5540012 | 0.01731064 |
| #1052 | PCED1A | 16.93581 | -0.5636826 | 0.006904919 |
| #1053 | PSMB6 | 85.3373 | 0.3766975 | 0.04011095 |
| #1054 | TRAPPC6B | 19.79492 | -0.5049037 | 0.005062639 |
| #1055 | ZMYND11 | 30.50574 | -0.482254 | 0.02117961 |
| #1057 | ERP44 | 60.28149 | 0.3011418 | 0.002534851 |
| #1059 | SPICE1 | 10.84636 | -0.8871486 | 0.01567217 |
| #1061 | PPP1R3F | 8.061954 | -0.9833781 | 0.005172234 |
| #1063 | TAZ | 35.40181 | -0.3734994 | 0.002148541 |
| #1064 | MRPL30 | 19.76027 | -0.4832219 | 0.002442139 |
| #1065 | APBB1 | 13.81517 | -0.7622463 | 0.02066481 |

| | | | | |
|---|---|---|---|---|
| **#1066** | RPL30 | 5,653 | -0.298 | 0.04150463 |
| **#1067** | ITPRIPL2 | 11.20774 | 0.7165351 | 0.03843625 |
| **#1068** | LTBP4 | 14.51689 | -0.659347 | 0.01325364 |
| **#1069** | KCTD7 | 12.68652 | -0.6250283 | 0.003529536 |
| **#1070** | PILRB | 36.50284 | -0.5069641 | 0.04099802 |
| **#1071** | PKD1 | 15.64164 | -0.6341254 | 0.01429409 |
| **#1072** | SUPT20H | 41.43061 | -0.3986432 | 0.01358074 |
| **#1073** | MUS81 | 17.00917 | -0.5728061 | 0.01001495 |
| **#1074** | RPS12 | 7,270 | -0.2892687 | 0.03815899 |
| **#1076** | ST3GAL1 | 50.39346 | -0.3530972 | 0.007768225 |
| **#1077** | NANS | 32.11983 | 0.4295014 | 0.02589778 |
| **#1082** | DRAM1 | 6.975008 | 0.8108571 | 0.01929832 |
| **#1083** | NOP56 | 69.95414 | -0.3082183 | 0.004562969 |
| **#1084** | COX5B | 262.5793 | 0.2942635 | 0.02663939 |
| **#1085** | LTBR | 15.6627 | 0.5668554 | 0.02885685 |
| **#1086** | KLHDC2 | 40.02662 | -0.4507951 | 0.02885477 |
| **#1087** | MEA1 | 29.82934 | 0.3845141 | 0.008106426 |
| **#1089** | RGL2 | 19.99121 | -0.4622016 | 0.00126175 |
| **#1090** | MTA1 | 21.63606 | -0.6272744 | 0.03806344 |
| **#1091** | DERL1 | 42.21927 | 0.3366947 | 0.006177781 |
| **#1092** | ACAP3 | 16.12552 | -0.5226759 | 0.002428331 |
| **#1093** | WAS | 125.5835 | 0.2565531 | 0.003418226 |
| **#1094** | GCSAM | 10.0366 | -0.93091 | 0.01720675 |
| **#1095** | TRMT13 | 24.10502 | -0.5955154 | 0.04020488 |
| **#1096** | DENND1A | 17.61527 | 0.5508592 | 0.03348142 |
| **#1099** | PYCR2 | 37.53322 | 0.3853783 | 0.01891401 |
| **#1100** | SLC35E2A | 10.87645 | -0.7209046 | 0.00668452 |
| **#1101** | SCARB2 | 38.45487 | 0.4512181 | 0.04687995 |
| **#1103** | ULK3 | 16.61475 | -0.5585307 | 0.008116749 |
| **#1104** | WDR60 | 28.91763 | -0.571261 | 0.04896935 |
| **#1105** | ZNF397 | 18.71116 | -0.6281825 | 0.0285996 |
| **#1106** | ZC3H6 | 28.86948 | -0.5131772 | 0.03240869 |

| | | | | |
|---|---|---|---|---|
| **#1107** | TOM1L2 | 12.94611 | -0.5697056 | 0.001077871 |
| **#1108** | NME3 | 79.37034 | -0.3469842 | 0.02277876 |
| **#1110** | COX8A | 218.7985 | 0.33183 | 0.04764128 |
| **#1113** | PSMA5 | 80.16164 | 0.3647194 | 0.04145769 |
| **#1115** | RP2 | 19.27778 | 0.4553343 | 0.01321556 |
| **#1116** | CNPY4 | 10.22059 | -0.8141557 | 0.01158957 |
| **#1117** | PCYOX1 | 15.03659 | -0.515208 | 0.000548522 |
| **#1119** | CDK10 | 20.96556 | -0.4537353 | 0.00245145 |
| **#1121** | ATP6V1G1 | 258.5813 | 0.2468556 | 0.009027427 |
| **#1122** | DANCR | 48.13031 | -0.4107491 | 0.02798124 |
| **#1124** | CCDC14 | 26.31825 | -0.4612204 | 0.01453691 |
| **#1125** | IDH3A | 18.80752 | 0.4456165 | 0.008745169 |
| **#1126** | HTATIP2 | 36.83634 | 0.3925425 | 0.02244823 |
| **#1127** | PSMD7 | 81.73737 | 0.2962396 | 0.01053009 |
| **#1132** | TTC14 | 111.9672 | -0.3519753 | 0.03873015 |
| **#1133** | TRA2A | 124.021 | -0.2979655 | 0.01683719 |
| **#1134** | SLC31A1 | 13.31258 | 0.581567 | 0.02647918 |
| **#1135** | RPL21 | 4,484 | -0.2544986 | 0.02560808 |
| **#1137** | N4BP2L2 | 285.5952 | -0.2977024 | 0.03226934 |
| **#1139** | CRYL1 | 19.40293 | 0.5236478 | 0.03573027 |
| **#1140** | CBR4 | 19.07536 | -0.5543905 | 0.01815844 |
| **#1143** | CSAD | 24.6936 | -0.4356313 | 0.006420286 |
| **#1144** | SYK | 74.90895 | 0.3304829 | 0.0250264 |
| **#1146** | ARHGEF1 | 154.3086 | -0.3056819 | 0.02533893 |
| **#1147** | CD33 | 32.44255 | 0.4425555 | 0.03817778 |
| **#1149** | POLG2 | 15.16148 | -0.5765131 | 0.009026435 |
| **#1150** | MTHFR | 26.53702 | 0.4352012 | 0.02450777 |
| **#1152** | DNAJC3 | 52.35265 | 0.362777 | 0.026549 |
| **#1153** | NUDCD3 | 33.0955 | -0.3989982 | 0.009324058 |
| **#1154** | TMEM41B | 34.68505 | -0.3716573 | 0.004913069 |
| **#1156** | P3H1 | 12.19018 | -0.588433 | 0.002544646 |
| **#1157** | SNTB1 | 12.99784 | 0.5544419 | 0.01947615 |

| | | | | |
|---|---|---|---|---|
| #1159 | SMURF2 | 23.56162 | -0.4571861 | 0.009441624 |
| #1160 | RAP1A | 116.972 | 0.2502401 | 0.003420873 |
| #1161 | PRMT2 | 186.7654 | -0.3248045 | 0.04116564 |
| #1162 | GPS2 | 102.7393 | -0.300295 | 0.01499044 |
| #1164 | METAP2 | 96.22856 | -0.2564177 | 0.000382825 |
| #1165 | TMEM164 | 9.65359 | 0.5897612 | 0.01018212 |
| #1168 | RNF13 | 60.32841 | 0.3265509 | 0.01720783 |
| #1170 | MAPKAPK5 | 30.38837 | -0.4331077 | 0.01672315 |
| #1171 | RPS25 | 2,977 | -0.2748422 | 0.03893468 |
| #1172 | AP3S1 | 42.24138 | 0.3560265 | 0.01763738 |
| #1173 | ETV6 | 43.09076 | 0.410025 | 0.04134171 |
| #1176 | MEPCE | 27.84946 | 0.4391697 | 0.03027957 |
| #1177 | PRKAA1 | 42.15943 | -0.3419203 | 0.004257409 |
| #1178 | RINL | 23.15982 | -0.544547 | 0.03201211 |
| #1179 | KLF6 | 289.3331 | 0.2985177 | 0.03938623 |
| #1181 | CFAP97 | 63.55283 | -0.3789074 | 0.03323627 |
| #1182 | BTN3A1 | 51.57838 | -0.4275013 | 0.04413264 |
| #1185 | RPS6 | 4,097 | -0.2559687 | 0.02970289 |
| #1186 | ARF4 | 43.97012 | 0.3042873 | 0.001832654 |
| #1189 | NAA40 | 11.54215 | -0.6747882 | 0.009212881 |
| #1190 | METTL17 | 27.88573 | -0.4296395 | 0.01311403 |
| #1192 | SCPEP1 | 57.97497 | 0.3841109 | 0.04453518 |
| #1193 | ATP6AP2 | 120.3243 | 0.2834287 | 0.01747723 |
| #1194 | TRIM28 | 53.08976 | -0.4011462 | 0.03500534 |
| #1195 | SKI | 23.29947 | -0.4535058 | 0.009701698 |
| #1199 | DEF6 | 54.16389 | -0.4086385 | 0.04027618 |
| #1200 | PADI4 | 18.36412 | 0.511919 | 0.03312643 |
| #1202 | FRA10AC1 | 30.35107 | -0.4447676 | 0.02161883 |
| #1204 | LIMK2 | 6.052576 | 0.9299487 | 0.03476185 |
| #1206 | USP28 | 30.72507 | -0.4671227 | 0.02982958 |
| #1207 | PLCG1 | 13.28249 | -0.917548 | 0.04853794 |
| #1208 | UBTF | 79.97102 | -0.3785803 | 0.04603253 |

| | | | | |
|---|---|---|---|---|
| #1209 | EML3 | 22.62037 | -0.4606448 | 0.01048887 |
| #1210 | ABRAXAS1 | 28.64817 | -0.403364 | 0.007520261 |
| #1213 | ACSL6 | 10.49529 | -0.8753966 | 0.02383204 |
| #1214 | RALGDS | 28.16796 | -0.4859073 | 0.03057667 |
| #1215 | SIN3B | 22.97216 | -0.4969596 | 0.02089153 |
| #1216 | SGSM3 | 34.72231 | -0.3669961 | 0.006024725 |
| #1217 | PRUNE1 | 9.273195 | 0.6953316 | 0.03021595 |
| #1218 | ELMOD3 | 16.94838 | -0.4692108 | 0.001205137 |
| #1220 | VPS51 | 76.86641 | -0.3586187 | 0.03507743 |
| #1221 | AGO2 | 39.65662 | -0.3375828 | 0.003218591 |
| #1222 | ZBTB4 | 23.8563 | -0.3993776 | 0.00092796 |
| #1223 | RFX1 | 20.96098 | -0.4707963 | 0.009656336 |
| #1225 | KDM1B | 27.02288 | 0.4571106 | 0.03997578 |
| #1227 | RPL13 | 7,663 | -0.2249791 | 0.01825006 |
| #1229 | LRRC14 | 10.08985 | -0.8087016 | 0.01638289 |
| #1231 | CLDND1 | 56.71923 | -0.2997468 | 0.002879645 |
| #1232 | RRAGC | 24.54177 | 0.354856 | 0.001251497 |
| #1233 | DGKE | 23.30554 | -0.5232379 | 0.03128538 |
| #1235 | LIAS | 17.05975 | -0.4544079 | 0.000114873 |
| #1236 | BICDL1 | 10.44755 | -0.8116282 | 0.01940488 |
| #1237 | DZIP3 | 15.73248 | -0.5783696 | 0.01630413 |
| #1238 | TPP1 | 78.54139 | 0.3058964 | 0.02038991 |
| #1239 | GARS-DT | 53.16037 | -0.4143294 | 0.04549775 |
| #1241 | UBE3C | 27.84906 | -0.4174771 | 0.01177036 |
| #1242 | RBM5 | 88.37602 | -0.3095003 | 0.01958847 |
| #1243 | ZC3H12D | 13.65123 | -0.670931 | 0.02214604 |
| #1245 | AKTIP | 17.2917 | -0.6403423 | 0.03739682 |
| #1247 | RPL14 | 2,803 | -0.2402504 | 0.02629263 |
| #1251 | FASTKD1 | 9.514834 | -0.6231038 | 0.000364058 |
| #1252 | CHMP2B | 41.13064 | 0.3398538 | 0.01509543 |
| #1253 | RPL41 | 7,856 | -0.255913 | 0.03779023 |
| #1255 | WDR59 | 16.57248 | -0.5366732 | 0.01312469 |

| | | | | |
|---|---|---|---|---|
| **#1256** | EEPD1 | 7.618866 | -0.7799701 | 0.001853243 |
| **#1257** | SPTSSA | 22.82442 | 0.3885969 | 0.008154037 |
| **#1260** | ZNF236 | 18.10473 | -0.4361798 | 0.000294597 |
| **#1261** | COX7B | 172.6064 | 0.2709954 | 0.02318039 |
| **#1262** | CHSY1 | 11.81058 | 0.5660715 | 0.02263425 |
| **#1263** | BCLAF3 | 14.04475 | -0.5652533 | 0.009412675 |
| **#1264** | SGSM2 | 13.25079 | -0.567682 | 0.00735663 |
| **#1265** | TTC13 | 13.41207 | -0.5846923 | 0.01040584 |
| **#1266** | CARMIL2 | 9.973127 | -0.9057988 | 0.02663936 |
| **#1267** | NAPEPLD | 7.679961 | -0.8504252 | 0.006564204 |
| **#1268** | RPL6 | 3,085 | -0.2199771 | 0.01819312 |
| **#1269** | MAML3 | 13.00862 | 0.5102666 | 0.01479941 |
| **#1270** | RAB2A | 97.76124 | 0.2945459 | 0.02378373 |
| **#1271** | HIBCH | 18.53176 | -0.5302621 | 0.01941364 |
| **#1274** | SORBS3 | 14.03152 | -0.5417891 | 0.006633613 |
| **#1275** | CDK11B | 39.41507 | -0.3396779 | 0.006137228 |
| **#1276** | KLF12 | 38.79952 | -0.3676357 | 0.0147002 |
| **#1277** | SZRD1 | 52.5696 | 0.2795077 | 0.002643064 |
| **#1280** | MAD2L1BP | 8.134919 | 0.538331 | 0.000624435 |
| **#1281** | SLC7A6 | 24.03077 | -0.4969631 | 0.03095702 |
| **#1283** | ASPHD2 | 5.444991 | 0.7881982 | 0.01406923 |
| **#1284** | ACTR2 | 339.7433 | 0.282908 | 0.04394112 |
| **#1285** | TXNL4B | 7.436187 | 0.5815645 | 0.003001593 |
| **#1286** | HIP1R | 15.98627 | -0.5751934 | 0.02087271 |
| **#1287** | UQCRB | 624.9097 | -0.2143302 | 0.01136063 |
| **#1288** | RPS27A | 5,393 | -0.2676809 | 0.04813443 |
| **#1289** | FBXW4 | 23.80071 | -0.4115322 | 0.006050109 |
| **#1290** | ALKBH6 | 9.451089 | -1.023684 | 0.03475772 |
| **#1291** | UBAC2 | 53.66019 | 0.3473905 | 0.03241868 |
| **#1292** | NFAM1 | 22.4307 | 0.4283366 | 0.02314301 |
| **#1293** | CIR1 | 87.87349 | 0.257199 | 0.006103182 |
| **#1294** | SNX6 | 114.5079 | 0.2635046 | 0.01451504 |

| | | | | |
|---|---|---|---|---|
| #1295 | VPS8 | 23.91611 | -0.4285274 | 0.01045411 |
| #1296 | SRSF1 | 62.63998 | -0.3472828 | 0.0292538 |
| #1297 | CYB5R1 | 21.87612 | 0.455221 | 0.0332283 |
| #1298 | RNF114 | 53.58469 | 0.3163349 | 0.01789675 |
| #1299 | CYCS | 111.9485 | 0.2920199 | 0.02773927 |
| #1302 | POLE4 | 72.30538 | 0.3073406 | 0.02396377 |
| #1304 | RGP1 | 13.99355 | -0.5192523 | 0.004585306 |
| #1307 | KIAA0930 | 35.67119 | 0.3988416 | 0.03920444 |
| #1308 | PRCP | 38.34323 | 0.3835113 | 0.03615665 |
| #1309 | BCL6 | 26.87952 | 0.4487568 | 0.04549063 |
| #1310 | PRR12 | 25.00352 | -0.4108314 | 0.008550815 |
| #1311 | DBNL | 113.8634 | 0.3038484 | 0.03691793 |
| #1313 | ZNF699 | 15.5475 | -0.4640762 | 0.001344424 |
| #1314 | BCAP31 | 93.61048 | 0.2942965 | 0.02621273 |
| #1316 | SMYD4 | 15.48184 | -0.5224357 | 0.01045855 |
| #1317 | FOXN2 | 34.30114 | 0.3841789 | 0.031299 |
| #1319 | RPS19 | 4,542 | -0.2043835 | 0.01557542 |
| #1320 | LENG8 | 88.73089 | -0.3251545 | 0.03506072 |
| #1321 | ERCC5 | 55.50604 | -0.3798714 | 0.04162104 |
| #1322 | TCP11L2 | 36.62229 | -0.4431952 | 0.04431062 |
| #1323 | CCDC130 | 30.26367 | -0.3493157 | 0.00241437 |
| #1326 | NSD3 | 149.859 | -0.3098154 | 0.04429546 |
| #1328 | ZNF431 | 45.48599 | -0.3811339 | 0.0315584 |
| #1331 | UCHL3 | 20.31818 | 0.3622981 | 0.000875835 |
| #1334 | MAP9 | 11.3419 | -0.5816818 | 0.005666308 |
| #1339 | RILP | 8.945911 | 0.6017752 | 0.01923039 |
| #1340 | CRLF3 | 52.33701 | -0.3568839 | 0.02888099 |
| #1341 | TIMM8B | 55.30241 | 0.3479301 | 0.03923745 |
| #1342 | INO80E | 27.75642 | -0.3681202 | 0.004202994 |
| #1343 | ZNF585A | 22.4115 | -0.4253846 | 0.008416609 |
| #1344 | SLC35B1 | 19.57042 | 0.4241596 | 0.0174573 |
| #1345 | OARD1 | 43.94064 | -0.3143631 | 0.005166493 |

| | | | | |
|---|---|---|---|---|
| **#1347** | EIF4E2 | 47.55164 | 0.3406852 | 0.02812187 |
| **#1349** | CARD8 | 81.06246 | -0.3203614 | 0.03042748 |
| **#1350** | WDR73 | 25.13892 | -0.4608228 | 0.02728455 |
| **#1351** | SSR4 | 308.1715 | 0.2633351 | 0.03658497 |
| **#1353** | STAU1 | 48.87189 | 0.2815672 | 0.003902319 |
| **#1356** | FUNDC2 | 46.81512 | 0.2956972 | 0.007296096 |
| **#1357** | GTF2B | 32.91893 | 0.3703285 | 0.02530626 |
| **#1361** | BAK1 | 14.90939 | 0.4626278 | 0.0152112 |
| **#1362** | DAP | 34.25815 | 0.4101982 | 0.04857917 |
| **#1367** | EID1 | 206.4581 | 0.2564373 | 0.02734725 |
| **#1368** | ERGIC2 | 40.97482 | -0.3697993 | 0.02425623 |
| **#1369** | HDGF | 64.91009 | 0.3040885 | 0.02322947 |
| **#1373** | KIF3A | 13.7779 | -0.4834722 | 0.001530713 |
| **#1374** | DDX39B | 131.9804 | -0.2695586 | 0.02222917 |
| **#1375** | NDE1 | 38.27905 | -0.337379 | 0.009101919 |
| **#1377** | SMARCD3 | 11.45728 | 0.5860535 | 0.03760711 |
| **#1378** | MFSD14A | 23.23273 | 0.348392 | 0.003373703 |
| **#1380** | RPS3 | 4,075 | -0.2339609 | 0.03742959 |
| **#1382** | ZNF609 | 31.68147 | -0.4441337 | 0.04122681 |
| **#1383** | NFKBIE | 9.806319 | 0.6280096 | 0.03700651 |
| **#1384** | RMND1 | 13.97963 | -0.491243 | 0.003757315 |
| **#1385** | ARHGEF9 | 21.11373 | -0.4791147 | 0.02475052 |
| **#1386** | EEF1B2 | 1,460 | -0.2369856 | 0.0358773 |
| **#1388** | GATA3 | 39.48105 | -0.4112635 | 0.04328903 |
| **#1389** | SLC12A6 | 41.286 | -0.3951098 | 0.03860298 |
| **#1390** | PIEZO1 | 35.82924 | -0.3374758 | 0.007999972 |
| **#1391** | PPARGC1B | 9.198717 | 0.600801 | 0.02533258 |
| **#1393** | LCLAT1 | 10.70074 | -0.6627902 | 0.01635679 |
| **#1394** | MRPL16 | 28.86323 | 0.3239003 | 0.004676225 |
| **#1397** | ZNF692 | 8.533766 | -0.6584452 | 0.00387868 |
| **#1400** | PHPT1 | 65.41028 | 0.3056361 | 0.02741829 |
| **#1404** | RPS23 | 4,939 | -0.2078274 | 0.02372411 |

| | | | | |
|---|---|---|---|---|
| #1407 | UNKL | 18.20799 | -0.4652414 | 0.01266411 |
| #1408 | TMX1 | 40.54517 | 0.2875702 | 0.003546035 |
| #1409 | TNNI2 | 8.69469 | 0.5780733 | 0.01743183 |
| #1410 | GATD1 | 13.33397 | -0.4973865 | 0.003933207 |
| #1411 | ESYT1 | 37.3657 | -0.367985 | 0.02257725 |
| #1412 | SAP18 | 182.4244 | 0.2437761 | 0.02304963 |
| #1413 | ZNF66 | 10.72514 | -0.8970671 | 0.04936761 |
| #1414 | DPH5 | 22.31037 | -0.4761708 | 0.03132413 |
| #1415 | RNPC3 | 41.83751 | -0.3484834 | 0.02152662 |
| #1416 | CROCC | 8.561149 | -0.6979011 | 0.008248283 |
| #1418 | EHMT1 | 54.4425 | -0.3602436 | 0.04260988 |
| #1419 | GRPEL1 | 31.56835 | 0.3899378 | 0.04069788 |
| #1420 | VDAC1 | 75.56703 | 0.3242179 | 0.04983672 |
| #1421 | HCG18 | 36.51806 | -0.3900043 | 0.03270965 |
| #1423 | ZNF720 | 22.33213 | -0.4929359 | 0.03788928 |
| #1424 | RPL3 | 3,792 | -0.2070953 | 0.02509905 |
| #1425 | S1PR4 | 67.52812 | -0.3468045 | 0.04720693 |
| #1426 | ARHGAP30 | 173.0907 | -0.2236884 | 0.01023875 |
| #1427 | UBN2 | 24.80982 | -0.483693 | 0.04395889 |
| #1428 | CUL9 | 11.27942 | -0.5652401 | 0.008001863 |
| #1429 | DOCK9 | 13.59838 | -0.6165196 | 0.03021533 |
| #1430 | ST6GALNAC | 17.21868 | -0.5859407 | 0.04441005 |
| #1431 | ITPR3 | 10.79774 | -0.717331 | 0.02772565 |
| #1432 | MECP2 | 85.07636 | -0.2918268 | 0.02555085 |
| #1433 | DNAJC24 | 16.81182 | -0.4715869 | 0.01187514 |
| #1434 | ATP6V0A2 | 24.83592 | -0.4248465 | 0.02197149 |
| #1435 | ZNF33B | 19.44433 | -0.5132884 | 0.03430891 |
| #1437 | MAPK1IP1L | 57.30576 | 0.2465771 | 0.001085017 |
| #1440 | MZF1 | 15.22569 | -0.4789878 | 0.008044104 |
| #1443 | PRPS2 | 13.7042 | 0.5091471 | 0.03357308 |
| #1444 | ZNF783 | 9.723145 | -0.5881001 | 0.004870703 |
| #1445 | APOLD1 | 9.880563 | -0.5716649 | 0.003609513 |

| | | | | |
|---|---|---|---|---|
| **#1447** | WSB2 | 14.99873 | 0.4268464 | 0.01051688 |
| **#1448** | SMARCC2 | 67.39715 | -0.2592204 | 0.004684218 |
| **#1452** | MAFG | 18.48012 | 0.4597747 | 0.03965238 |
| **#1453** | TMEM245 | 28.71116 | -0.4575598 | 0.04832687 |
| **#1457** | ZNF76 | 13.53692 | -0.5307151 | 0.01371495 |
| **#1458** | GLB1 | 20.03275 | 0.373628 | 0.008662924 |
| **#1459** | FBRSL1 | 25.16382 | -0.4351346 | 0.02865086 |
| **#1460** | AP1G2 | 27.07916 | -0.4046661 | 0.02194402 |
| **#1463** | BBS2 | 12.48493 | -0.6502435 | 0.03253683 |
| **#1465** | ANKRD26 | 21.23439 | -0.4948692 | 0.03808803 |
| **#1466** | PRR13 | 180.4496 | 0.253605 | 0.03327457 |
| **#1467** | PPP2R3B | 7.964576 | -0.6812247 | 0.005752042 |
| **#1468** | SH2D3A | 12.19902 | -0.7001361 | 0.03976848 |
| **#1469** | AGER | 9.372298 | -0.6243866 | 0.008254318 |
| **#1471** | MAGT1 | 32.29031 | 0.318292 | 0.01045597 |
| **#1472** | ZNF605 | 11.71549 | -0.6298604 | 0.02492261 |
| **#1473** | NDUFA5 | 81.89962 | 0.228522 | 0.003622602 |
| **#1475** | ANP32A | 129.4597 | 0.2081232 | 0.004594437 |
| **#1478** | PRIM2 | 12.61807 | -0.4761089 | 0.001960435 |
| **#1481** | PI4KA | 32.32891 | -0.3118793 | 0.001746197 |
| **#1482** | BDH1 | 11.13723 | -0.626742 | 0.02087615 |
| **#1485** | ZNF302 | 25.85226 | -0.363571 | 0.007258059 |
| **#1486** | HGSNAT | 32.52962 | -0.3784892 | 0.02597287 |
| **#1487** | DNAJB11 | 36.71923 | 0.3144295 | 0.0156643 |
| **#1489** | TXNDC17 | 42.71468 | 0.341669 | 0.03937699 |
| **#1491** | COMMD7 | 72.81072 | 0.2861936 | 0.02884143 |
| **#1492** | ZNF224 | 27.29671 | -0.3750741 | 0.01430451 |
| **#1493** | DESI1 | 23.91686 | 0.4196609 | 0.04416298 |
| **#1495** | TUBGCP6 | 16.83598 | -0.4755172 | 0.01653682 |
| **#1497** | SNF8 | 64.7276 | 0.3203724 | 0.04775983 |
| **#1498** | CNOT6 | 21.20649 | -0.4481654 | 0.02353417 |
| **#1500** | NDUFV2 | 82.17565 | 0.2349231 | 0.007241662 |

| | | | | |
|---|---|---|---|---|
| #1501 | TANC2 | 13.48137 | -0.508832 | 0.01153327 |
| #1503 | KRIT1 | 25.96298 | -0.3715223 | 0.01099609 |
| #1504 | KIZ | 20.59027 | -0.4757098 | 0.03197134 |
| #1505 | RXRB | 16.40482 | -0.4255662 | 0.003933508 |
| #1506 | ISYNA1 | 8.297925 | -0.6393624 | 0.005669327 |
| #1508 | ZSCAN30 | 8.972114 | -0.7304885 | 0.0212325 |
| #1510 | CLK2 | 12.58641 | -0.5110928 | 0.008553573 |
| #1511 | NFIX | 4.758337 | 0.8634033 | 0.03432591 |
| #1513 | NISCH | 24.73264 | -0.3593832 | 0.005038425 |
| #1515 | CNNM3 | 19.0268 | -0.3763539 | 0.000660478 |
| #1518 | ENSA | 106.8722 | 0.2433719 | 0.01949444 |
| #1519 | CNP | 16.01239 | 0.3981275 | 0.008523775 |
| #1520 | EXOSC4 | 7.043158 | 0.5861603 | 0.0132658 |
| #1522 | TIAL1 | 85.28592 | -0.2549242 | 0.01354239 |
| #1523 | GLMN | 14.62351 | -0.4469863 | 0.00376407 |
| #1525 | KNOP1 | 25.76488 | -0.3846674 | 0.01617679 |
| #1528 | KRI1 | 27.74186 | -0.3956826 | 0.02551902 |
| #1529 | NSMF | 17.03789 | -0.5458377 | 0.0410409 |
| #1532 | PRKCB | 171.5928 | 0.2662576 | 0.04747244 |
| #1533 | LYPLA1 | 66.42824 | 0.2404829 | 0.00572949 |
| #1536 | HOOK2 | 13.05031 | -0.5452406 | 0.0193363 |
| #1537 | LEMD2 | 24.08257 | -0.3470654 | 0.002280635 |
| #1539 | VTI1B | 52.66469 | 0.3057798 | 0.03149082 |
| #1540 | PPOX | 8.715965 | -0.5772995 | 0.002744912 |
| #1541 | AUH | 13.56856 | -0.5364848 | 0.01998278 |
| #1543 | BICD2 | 37.17851 | 0.3031014 | 0.01360353 |
| #1545 | CCNY | 61.22915 | 0.317291 | 0.04685611 |
| #1546 | FBXO3 | 38.3872 | -0.3636538 | 0.03420117 |
| #1547 | ZNF618 | 4.58051 | 0.6394546 | 0.001294992 |
| #1549 | TPT1 | 7,019 | -0.1835104 | 0.02299523 |
| #1550 | MZT2B | 249.8341 | -0.2495242 | 0.04198789 |
| #1553 | MAPRE2 | 48.36184 | 0.2731537 | 0.0106759 |

| | | | | |
|---|---|---|---|---|
| #1555 | NME1 | 24.84863 | 0.36883 | 0.02613199 |
| #1559 | SNHG29 | 474.6543 | -0.2413321 | 0.04835658 |
| #1564 | TM7SF2 | 8.427932 | -0.5538931 | 0.000385905 |
| #1565 | IMPAD1 | 22.11056 | 0.3584924 | 0.01446294 |
| #1567 | ZNF557 | 12.54246 | -0.5039555 | 0.009622589 |
| #1570 | SNX17 | 70.76521 | 0.301923 | 0.04509333 |
| #1571 | WDR54 | 22.37059 | -0.4382235 | 0.030261 |
| #1575 | PGAP1 | 4.329054 | 0.9633348 | 0.0487415 |
| #1576 | MOSMO | 11.26981 | -0.6383159 | 0.03096318 |
| #1578 | PATL1 | 22.48122 | 0.3764052 | 0.02410601 |
| #1581 | ATP5F1C | 119.7701 | 0.2449724 | 0.02727372 |
| #1582 | TJAP1 | 9.983021 | -0.5237057 | 0.002883818 |
| #1584 | ITFG2 | 31.69913 | -0.339233 | 0.01418302 |
| #1585 | TMEM129 | 13.29059 | -0.5438863 | 0.02369092 |
| #1586 | GCN1 | 15.25714 | -0.416591 | 0.002381798 |
| #1591 | RIC8B | 7.82577 | -0.6185471 | 0.004635714 |
| #1593 | TSGA10 | 14.01656 | -0.5715224 | 0.03683224 |
| #1594 | NCAPD3 | 14.71509 | -0.4665539 | 0.01221494 |
| #1595 | FBL | 114.2732 | -0.2112623 | 0.005281224 |
| #1600 | PRMT7 | 14.58513 | -0.4621765 | 0.01049121 |
| #1601 | URGCP | 9.479054 | -0.554876 | 0.005248575 |
| #1602 | ZNF248 | 9.353366 | -0.7275124 | 0.03159007 |
| #1603 | ZNF333 | 12.7753 | -0.4330924 | 0.000238388 |
| #1605 | CLSTN1 | 37.9766 | -0.3329161 | 0.02242279 |
| #1606 | GPR82 | 8.877265 | -0.6401052 | 0.01477484 |
| #1607 | BRF1 | 12.50334 | -0.5204713 | 0.01540577 |
| #1609 | PELP1 | 18.04039 | -0.4121839 | 0.009038047 |
| #1613 | FXR2 | 8.743381 | -0.5837521 | 0.006587082 |
| #1615 | NAP1L4 | 111.1731 | -0.2438422 | 0.0219241 |
| #1617 | TMEM181 | 25.20679 | -0.38466 | 0.021027 |
| #1618 | ATG2B | 15.08585 | -0.5591737 | 0.04228619 |
| #1622 | BNIP3L | 90.44752 | 0.272539 | 0.04096382 |

| | | | | |
|---|---|---|---|---|
| **#1623** | FBXO7 | 65.55275 | 0.2412594 | 0.009603042 |
| **#1626** | CFAP298 | 43.42068 | -0.3460234 | 0.03966553 |
| **#1627** | BECN1 | 44.42722 | 0.2427162 | 0.000490019 |
| **#1629** | RTCB | 32.79338 | 0.3063044 | 0.01471784 |
| **#1630** | ATP6V1E1 | 46.45743 | 0.2705469 | 0.01208394 |
| **#1632** | TSC1 | 20.34652 | -0.4282126 | 0.02312976 |
| **#1633** | AKAP10 | 24.39466 | -0.4062925 | 0.02745008 |
| **#1635** | PELI2 | 16.97611 | 0.3800626 | 0.01201021 |
| **#1636** | LIG1 | 15.10991 | -0.4706945 | 0.01715443 |
| **#1641** | ACP1 | 79.84964 | 0.2004124 | 0.000323943 |
| **#1643** | EEF2 | 661.1456 | -0.2219686 | 0.04688171 |
| **#1645** | IFFO1 | 13.57155 | -0.5447622 | 0.03060306 |
| **#1646** | PCTP | 5.886551 | 0.5619138 | 0.004713097 |
| **#1647** | DDHD2 | 16.59836 | -0.4413836 | 0.01535918 |
| **#1648** | AGO3 | 51.02115 | -0.2921144 | 0.0196694 |
| **#1649** | MRPL44 | 26.37579 | 0.3357665 | 0.01992738 |
| **#1651** | ST3GAL5 | 38.30971 | 0.3351185 | 0.04343858 |
| **#1653** | GTPBP3 | 10.85967 | -0.5089672 | 0.007380836 |
| **#1655** | OAF | 10.1355 | 0.5559343 | 0.04374601 |
| **#1657** | DTNBP1 | 31.06284 | 0.3422412 | 0.03382691 |
| **#1659** | ATP7A | 13.60629 | -0.4545555 | 0.007827421 |
| **#1660** | ABCB8 | 14.60023 | -0.4069716 | 0.001522267 |
| **#1661** | DDX51 | 17.45937 | -0.4598921 | 0.02562471 |
| **#1663** | CCDC152 | 9.960397 | -0.5214343 | 0.005219321 |
| **#1664** | STK4 | 301.5028 | -0.1816995 | 0.01286559 |
| **#1665** | MCEE | 11.22004 | -0.4883737 | 0.00508509 |
| **#1668** | ACSS1 | 24.39706 | -0.4270176 | 0.04171261 |
| **#1670** | SRPK2 | 68.34304 | -0.3085208 | 0.04907447 |
| **#1671** | TMA16 | 26.02417 | -0.3855997 | 0.02711701 |
| **#1672** | PTPN6 | 128.5856 | 0.2565634 | 0.04649724 |
| **#1675** | HEMK1 | 14.4345 | -0.4795221 | 0.01927357 |
| **#1676** | EEF1A1 | 9,128 | -0.1826146 | 0.03355719 |

| | | | | |
|---|---|---|---|---|
| **#1677** | TMEM208 | 38.37217 | 0.3237946 | 0.0374474 |
| **#1679** | PTGES3 | 211.4088 | 0.1975194 | 0.0185228 |
| **#1680** | OSER1-DT | 9.962125 | -0.647437 | 0.03063722 |
| **#1681** | SIDT1 | 16.65067 | -0.4986672 | 0.03741179 |
| **#1683** | SEC31B | 10.38497 | -0.4727312 | 0.001159427 |
| **#1685** | PSMC6 | 64.95858 | -0.2495062 | 0.01055088 |
| **#1686** | CACTIN | 8.809367 | -0.5087809 | 0.000487442 |
| **#1692** | TCHP | 27.11351 | -0.3892518 | 0.03441081 |
| **#1693** | SLF1 | 19.39498 | -0.4663034 | 0.03966979 |
| **#1694** | NIPSNAP3A | 17.06482 | 0.4250516 | 0.037108 |
| **#1695** | ATP1B1 | 6.67942 | 0.6572938 | 0.03863756 |
| **#1696** | COPZ1 | 51.73959 | 0.3003246 | 0.04119112 |
| **#1697** | ADAMTSL4 | 9.672876 | 0.5425476 | 0.03795529 |
| **#1698** | ZSWIM9 | 8.182982 | -0.5759165 | 0.006024651 |
| **#1699** | YIF1B | 24.81325 | 0.3835974 | 0.04710535 |
| **#1700** | RPL17 | 3,228 | -0.1733926 | 0.02697391 |
| **#1702** | KCTD13 | 11.11398 | -0.4786716 | 0.004828416 |
| **#1704** | SLC37A1 | 11.67938 | 0.4620041 | 0.02390745 |
| **#1705** | RAB1A | 43.26773 | 0.2840117 | 0.02109433 |
| **#1707** | PAAF1 | 11.58992 | -0.5095269 | 0.01421943 |
| **#1708** | ACTR1B | 27.11144 | -0.3732966 | 0.02667756 |
| **#1709** | HMCES | 22.30388 | -0.4394451 | 0.04305209 |
| **#1710** | ZNF514 | 8.709215 | -0.5443896 | 0.004827049 |
| **#1711** | KIFC2 | 7.541026 | -0.5566345 | 0.00067748 |
| **#1712** | WRN | 18.05912 | -0.4894789 | 0.0438942 |
| **#1714** | SLC27A1 | 10.43448 | -0.602257 | 0.0272317 |
| **#1715** | APPL2 | 14.86828 | -0.5025644 | 0.03183385 |
| **#1716** | ARHGAP45 | 147.1999 | -0.2441147 | 0.04058799 |
| **#1717** | ADPRHL2 | 10.66595 | 0.4102587 | 0.002932953 |
| **#1720** | MOGS | 24.68094 | -0.3277727 | 0.004462506 |
| **#1722** | RAB3GAP2 | 38.2108 | -0.3401592 | 0.03639047 |
| **#1725** | PYGO2 | 10.42289 | -0.5372352 | 0.01409213 |

| | | | | |
|---|---|---|---|---|
| #1726 | WDR41 | 14.58517 | 0.3992733 | 0.01540268 |
| #1727 | SNRNP200 | 90.48385 | -0.2710834 | 0.04115765 |
| #1729 | TMEM14A | 8.495578 | -0.6008986 | 0.01367429 |
| #1730 | ZNF593 | 17.50335 | 0.4055481 | 0.03177606 |
| #1734 | CYHR1 | 29.76066 | -0.3750351 | 0.03883293 |
| #1737 | TRMT10B | 21.50239 | -0.4180457 | 0.03162005 |
| #1740 | ZC3H13 | 91.63266 | -0.274321 | 0.04599467 |
| #1741 | SNHG10 | 11.49834 | -0.510582 | 0.01646688 |
| #1744 | SLC35E3 | 25.00958 | -0.3642096 | 0.02078582 |
| #1745 | LZTR1 | 9.386458 | -0.4894478 | 0.001901141 |
| #1748 | ARMCX3 | 32.57531 | 0.2870628 | 0.01236267 |
| #1754 | PCSK5 | 15.82818 | -0.4756625 | 0.03154252 |
| #1756 | SNHG3 | 66.2498 | -0.2319002 | 0.007443753 |
| #1757 | IL27RA | 50.37062 | -0.3083601 | 0.03875156 |
| #1758 | CEP68 | 18.69116 | -0.4473971 | 0.03501793 |
| #1763 | SLC26A11 | 9.487273 | -0.5685007 | 0.01689272 |
| #1764 | CAPN10 | 14.69463 | -0.4689561 | 0.02385525 |
| #1766 | ECHDC2 | 22.28261 | -0.4194457 | 0.0382049 |
| #1767 | EYA3 | 12.21661 | -0.4224495 | 0.001461024 |
| #1768 | GPR137 | 8.467417 | 0.4547461 | 0.005186579 |
| #1769 | GDF11 | 9.80854 | -0.6148289 | 0.03007289 |
| #1770 | WDR27 | 8.051074 | -0.6387301 | 0.01868097 |
| #1771 | CBX1 | 35.48043 | -0.3383049 | 0.03553378 |
| #1773 | DUSP10 | 7.379239 | 0.4990619 | 0.008728707 |
| #1774 | IFT80 | 12.36452 | -0.5121787 | 0.02485539 |
| #1777 | EEFSEC | 19.91447 | -0.3388735 | 0.002538313 |
| #1778 | ORC2 | 17.29922 | -0.4934908 | 0.04930428 |
| #1779 | GDE1 | 25.13002 | 0.3259923 | 0.02071981 |
| #1785 | WNK1 | 116.0719 | -0.2393175 | 0.03478913 |
| #1786 | ZNF445 | 17.65019 | -0.3866923 | 0.01042918 |
| #1787 | PRKCQ | 27.61271 | -0.3682976 | 0.03425402 |
| #1791 | KLHL36 | 32.66908 | -0.313632 | 0.01729887 |

| | | | | |
|---|---|---|---|---|
| **#1794** | STAT5A | 27.02725 | 0.2963101 | 0.009693544 |
| **#1795** | ZNF568 | 9.289742 | -0.5562159 | 0.01495796 |
| **#1796** | AAMP | 32.72488 | 0.3330822 | 0.04586943 |
| **#1797** | SDHA | 54.86661 | 0.246645 | 0.01648502 |
| **#1798** | CD163 | 23.26508 | 0.3567968 | 0.03506328 |
| **#1801** | RTL8C | 13.00073 | 0.3841293 | 0.007965947 |
| **#1804** | RPL18A | 3,995 | -0.1822677 | 0.04479346 |
| **#1806** | GAB2 | 9.381749 | 0.5454521 | 0.04807602 |
| **#1807** | TCF3 | 24.5543 | -0.3795077 | 0.03102033 |
| **#1810** | ACTR10 | 37.22638 | 0.2596658 | 0.00785255 |
| **#1811** | RPL4 | 1,203 | -0.1489807 | 0.01727409 |
| **#1812** | TTF1 | 32.74202 | -0.270038 | 0.001057592 |
| **#1813** | SIRT5 | 9.51982 | -0.4904749 | 0.005014489 |
| **#1818** | KBTBD3 | 11.6384 | -0.480413 | 0.01445591 |
| **#1823** | VKORC1 | 32.04672 | 0.3280142 | 0.04341411 |
| **#1825** | UQCRFS1 | 94.66095 | 0.248466 | 0.04472149 |
| **#1829** | SON | 432.5429 | -0.1386395 | 0.005500842 |
| **#1830** | HAGHL | 10.33392 | -0.6113287 | 0.04059231 |
| **#1833** | COG8 | 9.493621 | -0.4582753 | 0.001050747 |
| **#1837** | MMS22L | 14.31929 | -0.423583 | 0.01254772 |
| **#1840** | ATP6V1E2 | 10.13919 | -0.4553685 | 0.003142328 |
| **#1843** | ADIPOR2 | 23.93663 | 0.33059 | 0.02475771 |
| **#1845** | VAT1 | 7.647923 | -0.5172427 | 0.001316037 |
| **#1846** | USE1 | 33.18738 | -0.2959769 | 0.01234011 |
| **#1847** | MPC1 | 48.59175 | 0.2908327 | 0.04460679 |
| **#1848** | BLOC1S2 | 53.92514 | 0.2146383 | 0.002864889 |
| **#1850** | TMEM268 | 10.49331 | 0.4530364 | 0.02330602 |
| **#1852** | EMSY | 19.11326 | -0.4214823 | 0.03473359 |
| **#1853** | ZDHHC3 | 27.19815 | 0.296042 | 0.01369883 |
| **#1854** | POLRMT | 11.99447 | -0.5275058 | 0.03402187 |
| **#1855** | KIF21A | 12.35333 | -0.5146234 | 0.03258237 |
| **#1858** | ABT1 | 31.10386 | 0.3233748 | 0.04126845 |

| | | | | |
|---|---|---|---|---|
| **#1859** | ST3GAL2 | 20.35574 | -0.3417238 | 0.007347428 |
| **#1860** | CYP2R1 | 8.98936 | -0.4789861 | 0.002858141 |
| **#1863** | TRAPPC12 | 26.10932 | -0.3511397 | 0.02641551 |
| **#1866** | ZNF90 | 8.161995 | -0.6220829 | 0.02432435 |
| **#1867** | ADAT2 | 11.59261 | -0.5002192 | 0.02350816 |
| **#1870** | COMMD6 | 455.3492 | -0.1956301 | 0.04626695 |
| **#1871** | PIGL | 11.70619 | -0.5054475 | 0.0262306 |
| **#1872** | GIGYF1 | 31.3459 | -0.3566703 | 0.0482085 |
| **#1873** | GNPAT | 29.39461 | -0.2841102 | 0.003807289 |
| **#1875** | GATAD1 | 41.0158 | -0.2694629 | 0.0133577 |
| **#1876** | DGAT1 | 26.59489 | -0.3545893 | 0.03165927 |
| **#1877** | RABEPK | 9.619619 | -0.4605626 | 0.003430249 |
| **#1879** | GALNT11 | 29.21224 | -0.2761458 | 0.001972993 |
| **#1882** | ATG4C | 12.28717 | -0.4991398 | 0.02962683 |
| **#1883** | IL21R | 9.607376 | -0.5610481 | 0.02574356 |
| **#1885** | ABCC5 | 11.63815 | -0.4374084 | 0.007381655 |
| **#1888** | RNF216 | 63.78719 | -0.2692887 | 0.03836828 |
| **#1893** | LAX1 | 11.88081 | -0.5247299 | 0.03592451 |
| **#1894** | CASP3 | 21.79525 | 0.3522018 | 0.03582764 |
| **#1895** | DFFB | 8.736253 | -0.4889927 | 0.004090392 |
| **#1898** | SIL1 | 18.12385 | 0.3680261 | 0.02988886 |
| **#1905** | ABHD18 | 17.23133 | -0.4123257 | 0.02656978 |
| **#1906** | PIP4K2B | 15.09343 | -0.3850136 | 0.007852242 |
| **#1907** | FNIP2 | 19.06029 | 0.3781698 | 0.04178451 |
| **#1911** | SP4 | 21.93588 | -0.3920033 | 0.03906667 |
| **#1912** | CRTC2 | 22.44297 | -0.3511067 | 0.02048938 |
| **#1913** | PSMB4 | 83.39999 | 0.2282939 | 0.03240479 |
| **#1914** | CREB1 | 56.50407 | -0.2738906 | 0.03754577 |
| **#1915** | SLC50A1 | 23.67343 | 0.2759282 | 0.002586665 |
| **#1919** | RAD54L2 | 16.90036 | -0.3652104 | 0.008820016 |
| **#1921** | VMA21 | 48.88254 | 0.2223662 | 0.006546951 |
| **#1927** | TTC9C | 12.64554 | -0.5142308 | 0.04201552 |

| | | | | |
|---|---|---|---|---|
| #1930 | RPAIN | 45.99836 | -0.2208065 | 0.000243281 |
| #1933 | FBXO25 | 26.25043 | -0.2817715 | 0.001820456 |
| #1934 | MFSD8 | 19.66506 | -0.3867378 | 0.02795856 |
| #1935 | GIGYF2 | 37.94377 | -0.3219678 | 0.04677548 |
| #1937 | HNRNPR | 158.0172 | 0.18867 | 0.0263518 |
| #1942 | IL17RC | 2.942691 | 0.6280744 | 8.96252E-05 |
| #1943 | UBE3D | 7.863179 | -0.5841393 | 0.01968665 |
| #1945 | MMS19 | 11.83316 | -0.3847015 | 0.000303315 |
| #1946 | RPRD1A | 27.20297 | -0.3246992 | 0.02144815 |
| #1949 | PGAM5 | 13.44723 | -0.4908433 | 0.04033168 |
| #1951 | APBB1IP | 133.6467 | -0.1982053 | 0.02457064 |
| #1954 | NVL | 16.03667 | -0.4655555 | 0.04925976 |
| #1958 | TNK2 | 28.44488 | -0.3269617 | 0.02674242 |
| #1959 | E4F1 | 13.25912 | -0.3882159 | 0.004982247 |
| #1961 | PMS2 | 21.77199 | -0.3263215 | 0.009994432 |
| #1965 | RETREG3 | 32.84821 | -0.2813773 | 0.01215641 |
| #1967 | RUFY2 | 22.02163 | -0.3574589 | 0.02550965 |
| #1969 | MRFAP1 | 74.91616 | 0.225924 | 0.02939343 |
| #1970 | CTNS | 4.998267 | 0.5860176 | 0.01862168 |
| #1973 | RAB14 | 83.95709 | 0.1725718 | 0.002515819 |
| #1975 | METAP1 | 18.41067 | -0.3628212 | 0.01639397 |
| #1980 | PPP3CC | 34.53645 | -0.3111401 | 0.03568405 |
| #1981 | NBEAL1 | 26.88406 | -0.3332019 | 0.02822914 |
| #1982 | TARS2 | 10.59764 | -0.4209738 | 0.003631597 |
| #1983 | AP1M1 | 41.11195 | -0.2448587 | 0.00699307 |
| #1984 | PGM2L1 | 13.41367 | -0.4290364 | 0.01988844 |
| #1986 | CC2D1B | 11.34952 | -0.4178162 | 0.005897344 |
| #1987 | BCL2L2 | 5.37118 | 0.5248321 | 0.007937724 |
| #1988 | LTB4R | 21.39707 | -0.3936878 | 0.04433274 |
| #1994 | ZNF544 | 13.40786 | -0.4921556 | 0.04511059 |
| #1995 | PNKP | 38.573 | -0.3054991 | 0.04167711 |
| #1996 | ZNF567 | 13.34095 | -0.4623707 | 0.03357429 |

| | | | | |
|---|---|---|---|---|
| **#2003** | RBM28 | 30.70464 | -0.3193462 | 0.03292168 |
| **#2004** | ATAD3B | 10.31348 | -0.4607906 | 0.01280744 |
| **#2009** | TAMM41 | 11.31813 | -0.4017703 | 0.003227806 |
| **#2017** | RMC1 | 12.43521 | 0.3453126 | 0.003840756 |
| **#2018** | ZNF383 | 9.466991 | -0.4889789 | 0.01526291 |
| **#2019** | PSMA1 | 114.878 | 0.1510081 | 0.001571468 |
| **#2021** | ARID2 | 42.80441 | -0.2993599 | 0.04726685 |
| **#2023** | GNB1 | 119.4874 | 0.1836297 | 0.02062441 |
| **#2027** | PINK1 | 14.68623 | -0.4400694 | 0.03508203 |
| **#2030** | ZNF264 | 22.76436 | -0.346566 | 0.02698775 |
| **#2033** | TTC31 | 16.88163 | -0.3181496 | 0.000487778 |
| **#2034** | RPL26 | 4,140 | -0.1392496 | 0.03379032 |
| **#2042** | RTF2 | 74.85104 | 0.2252235 | 0.0350266 |
| **#2043** | ZNF286A | 9.795708 | -0.4183609 | 0.002139585 |
| **#2053** | PCNP | 98.70351 | 0.1925571 | 0.02249721 |
| **#2054** | TIMM44 | 19.71803 | -0.3125695 | 0.004828941 |
| **#2055** | EXOC7 | 39.48971 | -0.2385891 | 0.005300499 |
| **#2056** | TRNAU1AP | 26.15019 | -0.3030481 | 0.01526097 |
| **#2060** | NPLOC4 | 30.98153 | -0.2588574 | 0.004076448 |
| **#2063** | XPA | 56.36072 | -0.2404659 | 0.02490824 |
| **#2064** | LTO1 | 12.27636 | -0.3752817 | 0.002992748 |
| **#2069** | BUD13 | 9.197913 | -0.4564376 | 0.007674981 |
| **#2074** | CCAR1 | 63.94328 | -0.2520939 | 0.04221416 |
| **#2076** | PHF21A | 39.67364 | -0.2379582 | 0.006312056 |
| **#2078** | NFRKB | 16.05697 | -0.3627502 | 0.0134851 |
| **#2080** | CRYZL1 | 24.17562 | -0.3030111 | 0.01193233 |
| **#2083** | DLG1 | 51.33703 | -0.2762061 | 0.04961841 |
| **#2085** | HMBOX1 | 28.97238 | -0.3215131 | 0.03676965 |
| **#2086** | ADPRH | 4.128534 | 0.6804555 | 0.03346637 |
| **#2091** | NDUFAF7 | 13.37733 | -0.3865746 | 0.01130367 |
| **#2094** | PIGP | 21.67025 | -0.3694912 | 0.04227066 |
| **#2095** | SMPD4 | 19.0961 | -0.3438696 | 0.01781733 |

| | | | | |
|---|---|---|---|---|
| **#2097** | MAP7D3 | 20.13909 | -0.3073495 | 0.005513025 |
| **#2098** | IRF2BP1 | 10.40069 | -0.5385744 | 0.04598121 |
| **#2099** | SLC43A3 | 10.21792 | 0.4038019 | 0.01832909 |
| **#2103** | PDE8A | 13.4087 | -0.4558631 | 0.04093054 |
| **#2105** | NUP93 | 16.35481 | 0.299233 | 0.003134367 |
| **#2107** | PLEKHO2 | 14.16793 | 0.4048334 | 0.0489012 |
| **#2109** | TTI1 | 9.160158 | -0.4667183 | 0.01251126 |
| **#2110** | TYSND1 | 12.78268 | -0.4392247 | 0.02851385 |
| **#2111** | PSMD2 | 57.48597 | 0.2490564 | 0.04764256 |
| **#2112** | CSE1L | 13.73931 | -0.3842502 | 0.01328093 |
| **#2116** | CAPS | 9.896557 | -0.4144496 | 0.003729607 |
| **#2118** | POLG | 29.7707 | -0.2918649 | 0.02026683 |
| **#2119** | RPL27 | 1,627 | -0.146393 | 0.04299003 |
| **#2122** | SLC39A4 | 12.3469 | 0.3638208 | 0.01531623 |
| **#2125** | COQ8A | 16.33097 | -0.3487537 | 0.01088174 |
| **#2128** | REXO4 | 32.76882 | -0.3099469 | 0.04333686 |
| **#2129** | PPP1R37 | 7.741042 | -0.5393112 | 0.01897185 |
| **#2130** | BBS9 | 8.561377 | -0.5492315 | 0.0315609 |
| **#2131** | TDRD3 | 22.51628 | -0.368153 | 0.0480785 |
| **#2139** | SEC62 | 191.5605 | -0.1849358 | 0.04255931 |
| **#2140** | ZNF395 | 14.89899 | -0.3384392 | 0.003573405 |
| **#2142** | DYNLRB1 | 97.5744 | 0.1930647 | 0.02715267 |
| **#2143** | AP3M2 | 25.35725 | -0.3170292 | 0.02575482 |
| **#2145** | SDR42E2 | 9.625249 | -0.5583968 | 0.04694524 |
| **#2148** | SH2D3C | 21.40498 | -0.3491853 | 0.03228989 |
| **#2151** | DIS3L | 19.26775 | -0.3318879 | 0.01521471 |
| **#2152** | MR1 | 21.91098 | 0.2991365 | 0.02014835 |
| **#2154** | SERPINB8 | 13.95672 | 0.3579676 | 0.02251653 |
| **#2156** | UBE2Z | 40.41809 | 0.2676036 | 0.04336613 |
| **#2164** | SNRNP48 | 21.39207 | -0.3065811 | 0.01106728 |
| **#2166** | SLC38A6 | 9.76142 | -0.460039 | 0.01837399 |
| **#2171** | CYB561D2.1 | 8.285206 | -0.6002667 | 0.04711903 |

| | | | | |
|---|---|---|---|---|
| **#2174** | CCT5 | 77.17997 | 0.2051361 | 0.02918992 |
| **#2177** | DDX55 | 21.62757 | -0.3082437 | 0.01339709 |
| **#2178** | DDX50 | 45.34756 | -0.2643708 | 0.03739257 |
| **#2183** | ACAD11 | 7.498966 | -0.5973443 | 0.03570113 |
| **#2190** | CALCOCO1 | 36.62384 | -0.268337 | 0.02502751 |
| **#2192** | RFFL | 23.6401 | -0.3236198 | 0.02758491 |
| **#2194** | MLYCD | 11.42725 | -0.416952 | 0.01595852 |
| **#2195** | NHP2 | 70.24896 | 0.1847021 | 0.01165906 |
| **#2199** | SCARF1 | 3.107019 | 0.7806175 | 0.03621125 |
| **#2201** | CA11 | 9.973158 | -0.4539644 | 0.02000105 |
| **#2204** | ZNF738 | 10.73108 | -0.4076566 | 0.009893377 |
| **#2206** | PIK3CD | 41.74534 | -0.2699684 | 0.03888301 |
| **#2208** | NAE1 | 41.06399 | -0.2438807 | 0.01800326 |
| **#2210** | QRICH1 | 28.05367 | -0.2861307 | 0.01961434 |
| **#2211** | AK9 | 12.96516 | -0.4230199 | 0.03033395 |
| **#2212** | TMEM189 | 19.62466 | 0.3186521 | 0.02967844 |
| **#2213** | FNTA | 50.24987 | -0.2653776 | 0.0492964 |
| **#2219** | RBM41 | 17.62575 | -0.3583541 | 0.02626094 |
| **#2225** | RANBP10 | 8.970994 | -0.4230033 | 0.004932091 |
| **#2227** | PYGB | 16.8222 | 0.3584861 | 0.04609106 |
| **#2229** | TMEM265 | 7.176773 | -0.5203815 | 0.01560658 |
| **#2230** | DBT | 13.72572 | -0.4347015 | 0.04351475 |
| **#2232** | DROSHA | 9.547539 | -0.445395 | 0.01512137 |
| **#2233** | TMEM94 | 10.29833 | -0.3878895 | 0.003505791 |
| **#2234** | LMBRD1 | 56.38806 | -0.208442 | 0.01328359 |
| **#2236** | ZNF529 | 12.02493 | -0.394728 | 0.01439893 |
| **#2237** | MRPL14 | 44.85443 | 0.246653 | 0.03848174 |
| **#2241** | METTL3 | 14.10927 | -0.3593921 | 0.01189661 |
| **#2243** | TOP3B | 10.45428 | -0.3650959 | 0.000365045 |
| **#2247** | TAOK2 | 10.62836 | -0.4238571 | 0.01706318 |
| **#2248** | ZNF701 | 13.68069 | -0.3649103 | 0.01255179 |
| **#2249** | ZNF641 | 17.17582 | -0.3347526 | 0.01467283 |

| | | | | |
|---|---|---|---|---|
| #2253 | SLC26A2 | 10.0457 | -0.450484 | 0.02210517 |
| #2254 | APOBEC3D | 11.9709 | -0.4414257 | 0.03410112 |
| #2255 | UBE4B | 20.54072 | -0.3097309 | 0.01462372 |
| #2257 | TRMU | 15.86609 | -0.3218206 | 0.005241376 |
| #2263 | ACP2 | 5.94006 | 0.4667224 | 0.01265844 |
| #2264 | KLHL22 | 6.72186 | -0.5375935 | 0.01684023 |
| #2266 | ANKRD46 | 8.082682 | -0.470021 | 0.01244573 |
| #2278 | USP37 | 12.37977 | -0.4515599 | 0.04545112 |
| #2281 | PSME3 | 21.33819 | 0.2702644 | 0.009551174 |
| #2282 | ENY2 | 108.2861 | 0.1895953 | 0.04101609 |
| #2283 | ITGA5 | 18.72625 | -0.3453934 | 0.02861074 |
| #2285 | DXO | 9.743128 | -0.4450107 | 0.02018431 |
| #2291 | SRCAP | 22.67002 | -0.3252286 | 0.03417909 |
| #2294 | GPATCH1 | 11.42396 | -0.4034894 | 0.01772653 |
| #2296 | SNX8 | 7.241757 | 0.4282402 | 0.01484981 |
| #2297 | SEC22C | 22.49208 | -0.2775556 | 0.00699803 |
| #2300 | XRCC6 | 128.4972 | 0.1448118 | 0.01144837 |
| #2307 | ZNF789 | 9.940505 | -0.4166026 | 0.01314501 |
| #2311 | TAB1 | 8.595717 | -0.4588981 | 0.01679107 |
| #2318 | CEP89 | 9.232084 | -0.3892578 | 0.003068749 |
| #2320 | RPUSD4 | 14.12672 | -0.3252697 | 0.003945037 |
| #2329 | APBA3 | 9.014147 | -0.4444216 | 0.01591293 |
| #2330 | OXCT1 | 19.07364 | -0.2846078 | 0.003805411 |
| #2332 | COG4 | 22.33082 | -0.276443 | 0.007615434 |
| #2338 | CASP2 | 21.55624 | -0.2937689 | 0.01386977 |
| #2339 | INTS2 | 9.700839 | -0.4758863 | 0.03577199 |
| #2343 | ZNF337 | 8.628916 | -0.4722273 | 0.02249463 |
| #2346 | ANKMY1 | 7.167763 | -0.4246707 | 0.001920904 |
| #2348 | ZNF281 | 33.53917 | 0.2635011 | 0.04240799 |
| #2349 | NUP188 | 13.50337 | -0.3703129 | 0.01991624 |
| #2354 | OTUD5 | 40.96548 | -0.2409261 | 0.0252521 |
| #2356 | GBA2 | 12.69756 | -0.3365819 | 0.004114802 |

| | | | | |
|---|---|---|---|---|
| **#2358** | MAEA | 25.67996 | -0.3049935 | 0.03611193 |
| **#2359** | TNPO2 | 8.552814 | -0.4449429 | 0.01460772 |
| **#2361** | NUP42 | 28.56557 | -0.2715847 | 0.02153658 |
| **#2364** | ZNF444 | 8.794229 | -0.5107078 | 0.04144007 |
| **#2365** | MICB | 9.118977 | 0.4237924 | 0.03729305 |
| **#2371** | ABHD13 | 21.37276 | 0.2543819 | 0.005969706 |
| **#2372** | PPP6C | 30.68487 | 0.224573 | 0.007225764 |
| **#2375** | MTAP | 17.37625 | -0.3753302 | 0.04817359 |
| **#2378** | CAMKMT | 9.818687 | -0.4635277 | 0.03564431 |
| **#2383** | ARL 3.00 | 14.4086 | -0.3617861 | 0.02289064 |
| **#2384** | SDHB | 51.09485 | 0.2210913 | 0.03647682 |
| **#2387** | PRPSAP2 | 24.80642 | -0.2850813 | 0.02187305 |
| **#2388** | TBPL1 | 23.16742 | 0.231688 | 0.001052886 |
| **#2389** | UBAP2 | 23.22307 | -0.2844875 | 0.01734778 |
| **#2390** | ZNF621 | 13.46344 | -0.3399309 | 0.009783555 |
| **#2392** | ELP1 | 10.56019 | -0.4557527 | 0.04108679 |
| **#2395** | CNBP | 237.4304 | 0.1555336 | 0.04998294 |
| **#2403** | ASPSCR1 | 27.30542 | -0.2836109 | 0.02878851 |
| **#2405** | GAMT | 14.18368 | -0.368628 | 0.02665058 |
| **#2407** | FASTKD5 | 12.59968 | 0.3503617 | 0.02774414 |
| **#2409** | COG1 | 14.77554 | -0.3241371 | 0.008721269 |
| **#2410** | MBD1 | 19.54704 | -0.2921892 | 0.0110156 |
| **#2411** | ANKRD49 | 48.74878 | -0.2141981 | 0.0196214 |
| **#2414** | ANKRD54 | 9.121897 | -0.3671895 | 0.001170494 |
| **#2418** | LARP1B | 17.70035 | -0.3096409 | 0.01466099 |
| **#2421** | CDC37L1 | 18.28622 | -0.3528783 | 0.04304324 |
| **#2423** | ENGASE | 13.37783 | -0.4031768 | 0.04018661 |
| **#2426** | ZNF324 | 9.142314 | -0.3921276 | 0.006635264 |
| **#2430** | IFT88 | 14.28086 | -0.3929907 | 0.04263823 |
| **#2433** | MOCS2 | 20.43833 | -0.3052025 | 0.02193514 |
| **#2436** | EIF4E | 32.44952 | 0.2264058 | 0.01364671 |
| **#2443** | HMGB1 | 761.6482 | 0.1302931 | 0.04938332 |

| | | | | |
|---|---|---|---|---|
| **#2444** | KLHL42 | 12.47795 | -0.3831888 | 0.02502791 |
| **#2446** | ARFGAP1 | 13.07564 | -0.3343376 | 0.007812452 |
| **#2449** | UBE2E1 | 33.3442 | 0.2092048 | 0.005555144 |
| **#2450** | COQ6 | 11.87525 | -0.3342391 | 0.003371769 |
| **#2452** | ZNF426 | 9.656377 | -0.3823969 | 0.007248166 |
| **#2453** | IGF1R | 17.08862 | -0.3041732 | 0.01035295 |
| **#2457** | CCDC57 | 43.1714 | -0.2417756 | 0.03727814 |
| **#2458** | POLM | 17.83541 | -0.3580084 | 0.04671109 |
| **#2462** | FEZ2 | 26.40229 | 0.2774893 | 0.04264717 |
| **#2463** | SLC33A1 | 19.08697 | -0.2968791 | 0.0142156 |
| **#2465** | AP1S1 | 13.47315 | 0.328068 | 0.02353179 |
| **#2468** | SUDS3 | 33.5995 | -0.2677801 | 0.03811197 |
| **#2469** | RSBN1 | 64.3102 | -0.1925035 | 0.02254658 |
| **#2475** | CRAMP1 | 9.577722 | -0.4157473 | 0.01819984 |
| **#2481** | AASDH | 15.31076 | -0.3634583 | 0.03680204 |
| **#2482** | TAF6 | 16.32453 | -0.298786 | 0.006849241 |
| **#2489** | COPS7B | 16.57671 | -0.2965818 | 0.006993468 |
| **#2497** | HADHB | 52.07707 | 0.1925215 | 0.02054102 |
| **#2498** | HSD17B7 | 11.11422 | -0.4119573 | 0.03268131 |
| **#2500** | TMED2 | 80.75093 | 0.191269 | 0.04634349 |
| **#2505** | ACADSB | 13.35475 | -0.3903828 | 0.04087395 |
| **#2507** | AMPD3 | 6.741992 | -0.5072053 | 0.02530046 |
| **#2508** | NUP85 | 10.87399 | -0.3346896 | 0.001875087 |
| **#2509** | C1D | 47.71936 | 0.2213542 | 0.04191079 |
| **#2510** | ZNF821 | 10.67144 | -0.370726 | 0.01163097 |
| **#2511** | EIF1B | 87.61151 | 0.144078 | 0.007709938 |
| **#2514** | ZCCHC9 | 23.54534 | -0.3151191 | 0.04921819 |
| **#2519** | PLAGL1 | 11.73452 | 0.3300909 | 0.01693223 |
| **#2527** | TUBE1 | 9.069591 | -0.4582579 | 0.03516602 |
| **#2529** | ATG16L2 | 47.27616 | -0.2215438 | 0.03130703 |
| **#2530** | MRPL21 | 45.03086 | 0.1942461 | 0.01523418 |
| **#2536** | CERS5 | 22.17511 | -0.2695775 | 0.01304981 |

| | | | | |
|---|---|---|---|---|
| **#2538** | TAF1C | 9.51738 | -0.4401274 | 0.0326807 |
| **#2542** | ZNF346 | 10.96788 | -0.4117869 | 0.03445336 |
| **#2543** | MBD3 | 21.10053 | -0.3033177 | 0.03054366 |
| **#2549** | NCEH1 | 7.013177 | 0.4027131 | 0.01546217 |
| **#2551** | NAA80 | 7.983982 | -0.4938156 | 0.0379674 |
| **#2553** | SLC39A13 | 10.684 | -0.4291683 | 0.04190224 |
| **#2556** | NMNAT1 | 7.39555 | -0.4161352 | 0.007627247 |
| **#2563** | MRTFA | 21.80214 | -0.2395953 | 0.000659288 |
| **#2564** | NEK9 | 19.13006 | -0.2942354 | 0.01806199 |
| **#2574** | HAUS5 | 7.534509 | -0.4614159 | 0.02347738 |
| **#2578** | TBRG4 | 17.79421 | -0.3432562 | 0.04735242 |
| **#2579** | KAT2A | 9.545256 | -0.4202154 | 0.0262147 |
| **#2581** | OGFOD2 | 6.966826 | -0.4018814 | 0.003639592 |
| **#2584** | VILL | 8.158993 | -0.4230369 | 0.01658265 |
| **#2588** | CCDC28A | 23.65125 | -0.2375554 | 0.003370908 |
| **#2597** | TRIM39 | 8.293048 | -0.3963326 | 0.009766176 |
| **#2600** | MAP4K1 | 22.60606 | -0.2481246 | 0.006024855 |
| **#2603** | DDX19A | 22.97318 | -0.2700515 | 0.0189994 |
| **#2607** | ELMOD2 | 13.50178 | -0.3275939 | 0.01493795 |
| **#2613** | PIGQ | 7.203415 | -0.3986796 | 0.005270717 |
| **#2616** | TMEM80 | 20.12216 | -0.3215718 | 0.04728982 |
| **#2620** | CDC40 | 77.23525 | 0.178272 | 0.04050424 |
| **#2623** | ZDHHC8 | 9.432742 | -0.4280031 | 0.03281663 |
| **#2626** | PRPF38A | 43.9206 | -0.2052353 | 0.01863725 |
| **#2629** | MRPL47 | 33.96501 | 0.2427195 | 0.04695227 |
| **#2630** | TSG101 | 24.14298 | 0.2420557 | 0.01801315 |
| **#2632** | FCHO1 | 16.22916 | -0.2794529 | 0.005215267 |
| **#2633** | CFAP20 | 12.20689 | -0.4046199 | 0.04931271 |
| **#2634** | CTRL | 4.17594 | 0.5680161 | 0.0469333 |
| **#2635** | BICRAL | 21.42454 | -0.2906903 | 0.02939087 |
| **#2636** | TFG | 29.41808 | 0.2239394 | 0.0172632 |
| **#2638** | UPF3B | 30.95973 | -0.2483966 | 0.0285704 |

| | | | | |
|---|---|---|---|---|
| **#2640** | ADCK5 | 6.739964 | -0.4222823 | 0.009144264 |
| **#2641** | TBL1X | 35.54351 | -0.230799 | 0.02528448 |
| **#2642** | EEF2K | 7.573332 | -0.4525211 | 0.02557679 |
| **#2645** | NOL11 | 30.06819 | -0.2550866 | 0.03222651 |
| **#2646** | ZNF12 | 17.26852 | -0.3263153 | 0.03720871 |
| **#2651** | MLLT10 | 32.4491 | -0.2463548 | 0.0318123 |
| **#2653** | UFSP2 | 17.04481 | -0.3413892 | 0.0478006 |
| **#2654** | UBIAD1 | 14.27787 | -0.309813 | 0.01301399 |
| **#2663** | FLCN | 9.744487 | -0.3635023 | 0.01163036 |
| **#2664** | SLC35B3 | 9.055099 | -0.4299345 | 0.03348616 |
| **#2665** | COMMD1 | 30.12358 | 0.2250898 | 0.02161182 |
| **#2671** | ZSWIM8 | 23.08144 | -0.2442947 | 0.008150603 |
| **#2673** | SRPK1 | 28.03935 | -0.2538095 | 0.02695179 |
| **#2674** | IMPDH2 | 40.60577 | -0.2128538 | 0.0218311 |
| **#2685** | LNPK | 21.49688 | -0.3061185 | 0.04789472 |
| **#2686** | PSMG2 | 53.79208 | 0.1877518 | 0.03139114 |
| **#2688** | SCAP | 14.91123 | -0.3349302 | 0.03163488 |
| **#2692** | DENND4B | 32.75096 | -0.219225 | 0.01412466 |
| **#2697** | DCAF8 | 36.32967 | -0.2363474 | 0.03550404 |
| **#2698** | NSMCE2 | 19.39315 | -0.2637317 | 0.009445565 |
| **#2703** | STK3 | 8.10865 | 0.4140598 | 0.0489992 |
| **#2706** | DGCR8 | 12.16446 | -0.3442543 | 0.02117373 |
| **#2710** | TATDN1 | 31.5114 | -0.2436251 | 0.03193154 |
| **#2714** | STN 1.00 | 27.78809 | 0.2441474 | 0.03742327 |
| **#2716** | TEFM | 11.48742 | -0.3658636 | 0.02872139 |
| **#2719** | TTPAL | 11.31363 | -0.3595897 | 0.02391828 |
| **#2722** | CETN2 | 17.45498 | 0.289171 | 0.03550863 |
| **#2726** | EXOC4 | 37.76489 | -0.2396486 | 0.04623847 |
| **#2727** | CMAS | 18.83314 | 0.2628058 | 0.02110719 |
| **#2735** | TIMM21 | 7.880299 | -0.4523952 | 0.03605421 |
| **#2743** | SIK2 | 8.299364 | -0.4271815 | 0.03061603 |
| **#2747** | ATXN10 | 40.08952 | 0.2184753 | 0.04547332 |

| | | | | |
|---|---|---|---|---|
| **#2754** | CHERP | 18.03277 | -0.3057104 | 0.03550277 |
| **#2757** | RAI1 | 9.03867 | -0.4069519 | 0.02848765 |
| **#2763** | EIF3M | 147.7112 | -0.1370278 | 0.03676069 |
| **#2767** | TSTD2 | 11.34539 | -0.3900703 | 0.04611031 |
| **#2773** | CLYBL | 7.508366 | -0.4521121 | 0.03473796 |
| **#2774** | SPATA33 | 8.765103 | -0.4450715 | 0.04661918 |
| **#2785** | FUT4 | 6.099232 | 0.4273306 | 0.03318784 |
| **#2789** | ELP3 | 15.28084 | -0.3068775 | 0.0234984 |
| **#2793** | THOC7 | 67.95157 | 0.1771172 | 0.04571255 |
| **#2796** | ZMAT2 | 73.12128 | 0.1433193 | 0.01417639 |
| **#2799** | ABCD4 | 20.82759 | -0.2726686 | 0.02638683 |
| **#2801** | AMDHD2 | 12.14803 | -0.3357765 | 0.02233692 |
| **#2804** | DALRD3 | 19.15143 | -0.3076014 | 0.04789311 |
| **#2805** | ZNF518A | 41.25139 | -0.221428 | 0.04207837 |
| **#2809** | LDB1 | 15.4273 | -0.3332891 | 0.04498151 |
| **#2810** | RBFA | 18.78807 | -0.2757788 | 0.02128556 |
| **#2818** | SLC25A12 | 10.38538 | -0.338723 | 0.01377164 |
| **#2819** | VPS33A | 22.61911 | -0.225678 | 0.004162266 |
| **#2821** | RFC4 | 8.899809 | -0.3713841 | 0.01793982 |
| **#2823** | LETMD1 | 29.85371 | -0.2382142 | 0.03090697 |
| **#2829** | PRIMPOL | 8.704258 | -0.3865004 | 0.02266871 |
| **#2832** | SLC2A11 | 10.33896 | -0.3812429 | 0.03468048 |
| **#2836** | PIDD1 | 8.354459 | -0.3449065 | 0.006446627 |
| **#2837** | ASB8 | 37.68417 | 0.2121563 | 0.04066605 |
| **#2839** | TMEM43 | 23.85699 | -0.2551569 | 0.0267444 |
| **#2842** | PMPCA | 27.43067 | -0.2374849 | 0.02535124 |
| **#2844** | FBXL17 | 17.88985 | -0.3040903 | 0.04141218 |
| **#2847** | NCBP2 | 52.0512 | -0.1769902 | 0.02198762 |
| **#2852** | CIZ1 | 23.58552 | -0.2484989 | 0.02334006 |
| **#2854** | LONP1 | 15.72055 | -0.2635016 | 0.005856135 |
| **#2857** | TIGAR | 14.47046 | 0.3034983 | 0.0418676 |
| **#2858** | PPME1 | 9.260265 | -0.3185744 | 0.003417918 |

| | | | | |
|---|---|---|---|---|
| **#2863** | EFHC1 | 8.825477 | -0.3541563 | 0.01358453 |
| **#2874** | PPARA | 10.56275 | -0.3457756 | 0.02161257 |
| **#2886** | TRMT6 | 20.24324 | 0.221725 | 0.00721257 |
| **#2889** | DPF2 | 28.0572 | -0.2389236 | 0.03186106 |
| **#2904** | PMPCB | 54.21622 | -0.1880416 | 0.04036343 |
| **#2906** | SCRIB | 9.043518 | -0.3665584 | 0.02232525 |
| **#2910** | RBM7 | 33.48075 | 0.2209431 | 0.04545729 |
| **#2913** | UVSSA | 13.58504 | -0.3360974 | 0.04347602 |
| **#2915** | EDC4 | 13.34529 | -0.289492 | 0.01062844 |
| **#2923** | MPHOSPH6 | 23.93502 | -0.2700557 | 0.0492493 |
| **#2927** | MAP11 | 8.619957 | -0.4051981 | 0.03850731 |
| **#2936** | BTN2A1 | 23.25549 | -0.2403473 | 0.02112903 |
| **#2944** | USP19 | 11.36553 | -0.296241 | 0.007080181 |
| **#2949** | PACS2 | 15.5963 | -0.2831749 | 0.02117486 |
| **#2954** | TRIM26 | 21.62666 | 0.2510059 | 0.04053864 |
| **#2957** | ARSB | 7.787642 | 0.3806065 | 0.04379611 |
| **#2964** | SRD5A1 | 7.794769 | 0.3524203 | 0.0261807 |
| **#2970** | MTFMT | 10.93384 | -0.3332035 | 0.02310049 |
| **#2973** | VPS36 | 64.04362 | -0.1624022 | 0.03094992 |
| **#2977** | ANAPC7 | 20.38016 | -0.2637301 | 0.03092197 |
| **#2990** | DPH3 | 19.90197 | 0.2600942 | 0.04389114 |
| **#2991** | MYO19 | 13.44339 | -0.312001 | 0.0311591 |
| **#2998** | TBC1D19 | 9.099923 | -0.352983 | 0.02294231 |
| **#2999** | PRPF6 | 63.16245 | -0.1522323 | 0.02158359 |
| **#3019** | CTDP1 | 19.5954 | -0.2811555 | 0.04712533 |
| **#3029** | MROH1 | 13.83371 | -0.3011427 | 0.02769589 |
| **#3031** | FBXW2 | 30.19342 | -0.1896659 | 0.007845895 |
| **#3034** | KPTN | 3.435821 | 0.505641 | 0.04292789 |
| **#3038** | FCF1 | 19.16821 | -0.254404 | 0.02223073 |
| **#3040** | TWSG1 | 4.896651 | 0.447112 | 0.0469212 |
| **#3041** | STRIP1 | 15.9729 | -0.2727495 | 0.02196665 |
| **#3049** | STRADA | 25.36901 | -0.2030866 | 0.007497237 |

| | | | | |
|---|---|---|---|---|
| **#3054** | SEC24D | 13.94645 | 0.2584548 | 0.01602682 |
| **#3059** | CRIPT | 34.88738 | -0.2174276 | 0.04380319 |
| **#3061** | EIF2B4 | 17.4742 | -0.2426497 | 0.009778049 |
| **#3069** | NELFA | 16.75888 | -0.2507081 | 0.0124768 |
| **#3075** | OSBPL1A | 6.515534 | 0.3941297 | 0.04437531 |
| **#3078** | POLD1 | 6.582743 | -0.4215217 | 0.03590978 |
| **#3080** | VPS13B-DT | 7.094459 | -0.3870659 | 0.02650495 |
| **#3083** | SETD4 | 10.04018 | -0.3700633 | 0.04743616 |
| **#3086** | ATP13A2 | 14.11406 | -0.2744601 | 0.01540454 |
| **#3097** | FLOT2 | 53.44964 | -0.1453149 | 0.009273175 |
| **#3098** | ABCC10 | 8.520743 | -0.375832 | 0.03597462 |
| **#3109** | ZNF480 | 26.04248 | -0.2429312 | 0.04703453 |
| **#3112** | FXN | 20.65114 | -0.2505703 | 0.03123765 |
| **#3118** | TBP | 12.94065 | -0.2760132 | 0.0126112 |
| **#3137** | SLC20A2 | 12.08874 | -0.3277613 | 0.04373083 |
| **#3141** | EIF3J | 77.71726 | -0.1434412 | 0.03616972 |
| **#3146** | ALAD | 13.50102 | -0.2622217 | 0.008438872 |
| **#3149** | SART1 | 33.61914 | -0.2157689 | 0.04770333 |
| **#3151** | FECH | 7.727046 | 0.337811 | 0.03126186 |
| **#3152** | MUL1 | 7.714821 | 0.3618167 | 0.04773264 |
| **#3153** | NCAPH2 | 13.57809 | -0.2813014 | 0.02108257 |
| **#3157** | COG7 | 8.873123 | -0.2788528 | 0.001236616 |
| **#3158** | SMIM15 | 22.20304 | 0.2370507 | 0.04689857 |
| **#3159** | POGLUT1 | 16.73738 | -0.2621525 | 0.02623334 |
| **#3160** | ZMIZ2 | 19.11099 | -0.2382713 | 0.01914062 |
| **#3161** | SASS6 | 8.967936 | -0.3368616 | 0.02516249 |
| **#3167** | GPAT4 | 20.25325 | -0.2369422 | 0.02344954 |
| **#3184** | SLC7A6OS | 27.29916 | -0.2270023 | 0.04355216 |
| **#3185** | NDUFAF6 | 10.53312 | -0.3405691 | 0.04371755 |
| **#3194** | ACAD10 | 8.60465 | -0.3626027 | 0.03994829 |
| **#3196** | IL18BP | 9.08579 | 0.3066271 | 0.02680479 |
| **#3201** | ZFYVE19 | 9.462181 | -0.3446897 | 0.03785437 |

| | | | | |
|---|---|---|---|---|
| **#3210** | GPN2 | 11.33812 | -0.2979683 | 0.02288466 |
| **#3217** | CLCC1 | 13.09146 | -0.2780709 | 0.02180487 |
| **#3227** | BICRA | 8.380786 | -0.3308731 | 0.02278988 |
| **#3231** | CXXC1 | 22.19203 | -0.2237243 | 0.02420859 |
| **#3236** | MRS2 | 15.76698 | -0.24254 | 0.01301401 |
| **#3247** | POT1 | 7.007181 | -0.4022286 | 0.04858831 |
| **#3249** | CAD | 6.683756 | -0.3406994 | 0.0160078 |
| **#3254** | SLC39A11 | 8.212776 | 0.3191232 | 0.03237936 |
| **#3260** | RBBP5 | 15.29237 | -0.2416679 | 0.01142388 |
| **#3263** | IFNAR2 | 52.71785 | 0.15228 | 0.03613646 |
| **#3278** | SMG5 | 12.49741 | -0.2937446 | 0.03284973 |
| **#3290** | TEPSIN | 13.73351 | -0.2755966 | 0.0276019 |
| **#3291** | RNF214 | 21.4933 | -0.2411106 | 0.04316956 |
| **#3293** | MTHFSD | 6.831578 | -0.353333 | 0.02574915 |
| **#3317** | PHRF1 | 17.96468 | -0.2573334 | 0.04136538 |
| **#3318** | DEPDC5 | 13.4827 | -0.2677162 | 0.02212893 |
| **#3320** | THRAP3 | 131.8822 | 0.1113301 | 0.04926513 |
| **#3322** | SCCPDH | 10.68928 | 0.2792396 | 0.02765872 |
| **#3343** | UBE2K | 64.13104 | 0.1317041 | 0.02749568 |
| **#3345** | CENPW | 3.931531 | 0.4272555 | 0.04645433 |
| **#3354** | ACTR5 | 7.041069 | -0.3783792 | 0.04549181 |
| **#3356** | EIF2D | 24.42878 | -0.2021962 | 0.02143479 |
| **#3357** | TRIT1 | 8.388592 | -0.3293443 | 0.03048931 |
| **#3360** | TASP1 | 9.886648 | -0.2910442 | 0.01864882 |
| **#3367** | SLC35B4 | 8.461779 | -0.2858697 | 0.008424301 |
| **#3376** | WDTC1 | 21.71731 | -0.1829787 | 0.00298174 |
| **#3386** | STK19 | 16.07598 | 0.2479756 | 0.04375005 |
| **#3392** | SMARCAL1 | 9.539099 | -0.3201417 | 0.03592696 |
| **#3420** | CIAO3 | 10.62426 | -0.3180245 | 0.04777096 |
| **#3422** | NCAPD2 | 14.30609 | -0.2679755 | 0.03486614 |
| **#3446** | MAN2B2 | 20.75679 | -0.1865142 | 0.004520223 |
| **#3449** | SLC25A51 | 6.753965 | -0.3473338 | 0.03215262 |

| | | | | |
|---|---|---|---|---|
| **#3458** | ST7L | 9.741938 | -0.3233424 | 0.0467761 |
| **#3467** | DCTN6 | 19.82429 | 0.2052766 | 0.02475349 |
| **#3482** | CNOT3 | 20.17933 | -0.2255081 | 0.03458808 |
| **#3491** | PAXIP1 | 7.444148 | -0.3286395 | 0.03152701 |
| **#3497** | ZNF24 | 58.9229 | 0.1336649 | 0.03446406 |
| **#3514** | NUBP1 | 18.39556 | 0.2259828 | 0.04422271 |
| **#3524** | ANKRD52 | 8.50599 | -0.2971695 | 0.0224942 |
| **#3532** | ZNF740 | 11.31629 | -0.2998992 | 0.04778247 |
| **#3534** | POLL | 12.34929 | -0.2747703 | 0.03522896 |
| **#3541** | CCDC58 | 9.703533 | -0.3003042 | 0.03462802 |
| **#3546** | BBS7 | 6.646273 | -0.3156593 | 0.0197984 |
| **#3547** | RRP1 | 11.66798 | -0.2780137 | 0.03417403 |
| **#3550** | ZNF561 | 9.25503 | -0.3235231 | 0.04903955 |
| **#3557** | SND1 | 44.58776 | 0.1557764 | 0.04908471 |
| **#3559** | AARSD1 | 22.64879 | -0.1796455 | 0.009060795 |
| **#3576** | RIOX2 | 14.22035 | -0.2689087 | 0.0477287 |
| **#3601** | MCFD2 | 25.21806 | 0.1773952 | 0.02626364 |
| **#3604** | EME2 | 6.378089 | -0.2910847 | 0.009837569 |
| **#3628** | C1RL | 11.66848 | -0.2461172 | 0.01682401 |
| **#3631** | TMTC4 | 7.314699 | -0.3270211 | 0.03996423 |
| **#3637** | NRF1 | 11.91582 | -0.2605889 | 0.02832291 |
| **#3660** | AP4B1 | 7.284764 | -0.2998866 | 0.02425423 |
| **#3667** | SOCS7 | 6.939492 | -0.2789498 | 0.01143061 |
| **#3669** | RBM4B | 7.192727 | -0.2710724 | 0.009016 |
| **#3674** | SKIV2L | 11.90592 | -0.2729387 | 0.04266638 |
| **#3678** | COMMD10 | 22.04179 | 0.1898015 | 0.0334054 |
| **#3690** | CRK | 21.51344 | -0.2154233 | 0.04671932 |
| **#3698** | PCBD2 | 12.0095 | -0.2093313 | 0.002916611 |
| **#3722** | ZFYVE27 | 14.91854 | -0.2148063 | 0.01540428 |
| **#3732** | MRPS5 | 57.51543 | -0.135034 | 0.04610375 |
| **#3733** | PXMP4 | 7.557633 | -0.2499351 | 0.004915927 |
| **#3806** | DHX33 | 9.575573 | -0.2337899 | 0.01018365 |

| | | | | |
|---|---|---|---|---|
| **#3833** | TIMM23 | 10.05926 | 0.2287949 | 0.01966802 |
| **#3842** | RBMX2 | 27.33491 | -0.178833 | 0.04475248 |
| **#3853** | RPL26L1 | 24.27407 | 0.178403 | 0.0458415 |
| **#3917** | TMEM68 | 8.69628 | -0.2774833 | 0.04313587 |
| **#3936** | SLC38A9 | 9.245958 | -0.2580864 | 0.03303379 |
| **#3947** | RNF123 | 8.637579 | -0.2637442 | 0.03330244 |
| **#3954** | ZSCAN29 | 7.435408 | -0.2825826 | 0.03805382 |
| **#3961** | UTP4 | 9.338112 | 0.2033743 | 0.006973657 |
| **#3985** | PSMC2 | 25.52655 | 0.1521148 | 0.02646912 |
| **#4005** | CUL4A | 32.29538 | -0.148662 | 0.0358018 |
| **#4013** | ABRAXAS2 | 12.37589 | 0.2239719 | 0.04580264 |
| **#4029** | TACO1 | 9.587418 | 0.2382986 | 0.03986361 |
| **#4046** | FAHD2A | 12.12695 | -0.23863 | 0.0473919 |
| **#4059** | MRPL45 | 16.48537 | -0.1820657 | 0.01970758 |
| **#4066** | DEDD | 12.33468 | 0.2096283 | 0.03480911 |
| **#4068** | IMPACT | 14.98975 | 0.2060518 | 0.04918737 |
| **#4082** | XRCC1 | 14.67924 | -0.2044716 | 0.03266801 |
| **#4104** | WDR5 | 10.03274 | -0.2149938 | 0.01837427 |
| **#4150** | HCCS | 10.76088 | 0.207716 | 0.02862174 |
| **#4158** | MED24 | 13.23197 | -0.2074328 | 0.03482729 |
| **#4182** | SS18L1 | 7.243402 | -0.2330392 | 0.02057033 |
| **#4185** | SLC46A3 | 14.91645 | -0.2059614 | 0.04588618 |
| **#4192** | MRPS27 | 16.29747 | -0.1792618 | 0.02616605 |
| **#4239** | ALG2 | 12.26985 | 0.1999604 | 0.0399823 |
| **#4242** | PPP2R5D | 9.115909 | -0.2233766 | 0.03027215 |
| **#4304** | BRCC3 | 21.52842 | -0.1602068 | 0.0390648 |
| **#4336** | HECTD3 | 7.633177 | -0.1973508 | 0.01111522 |
| **#4351** | TM9SF4 | 21.4139 | 0.1509397 | 0.03926778 |
| **#4361** | TNPO3 | 25.6646 | -0.1464783 | 0.04247226 |
| **#4403** | ACLY | 25.64904 | 0.1320208 | 0.03158602 |
| **#4420** | SCLY | 6.989563 | -0.2332518 | 0.03826605 |
| **#4445** | CRTC1 | 10.23957 | -0.2112693 | 0.04669976 |

| | | | | |
|---|---|---|---|---|
| **#4450** | FARS2 | 12.59844 | 0.1746152 | 0.03422553 |
| **#4487** | ESS2 | 8.551648 | -0.2175071 | 0.04194881 |
| **#4490** | FBXO41 | 6.491824 | -0.2354527 | 0.04227126 |
| **#4520** | CAMTA2 | 14.24897 | -0.1692812 | 0.03388838 |
| **#4555** | DHDDS | 12.72862 | -0.1744394 | 0.03437961 |
| **#4628** | BCAT2 | 8.031031 | -0.2158146 | 0.04933868 |
| **#4787** | MLLT1 | 15.15672 | -0.1536513 | 0.0453876 |
| **#4791** | POC1B | 15.77313 | -0.1509124 | 0.04439962 |
| **#4812** | TCEAL3 | 15.58251 | 0.1297418 | 0.02502908 |
| **#4835** | TAF9 | 42.7939 | 0.08915955 | 0.04821046 |
| **#4968** | RNF121 | 5.747076 | -0.1639927 | 0.02261339 |
| **#5373** | PRKRA | 26.81787 | -0.08101684 | 0.03921349 |

Supplementary Table 4

| Rank | Symbol | Control Abundance | Expression | Significance (p) |
|---|---|---|---|---|
| #1 | IFI30 | 383.8093 | 1.064125 | 0.001057611 |
| #2 | IRF1 | 274.255 | 1.634085 | 0.003683898 |
| #3 | IGSF6 | 81.93609 | 1.020526 | 0.000368063 |
| #4 | STX11 | 56.63322 | 1.307442 | 0.001860728 |
| #5 | POMP | 117.8748 | 0.861872 | 0.000768375 |
| #6 | PSME1 | 358.2633 | 0.7975082 | 0.001858127 |
| #7 | PLEK | 173.0954 | 0.772246 | 0.000501262 |
| #8 | IFITM3 | 130.9113 | 1.90424 | 0.007537191 |
| #9 | NPC2 | 170.3921 | 0.909367 | 0.002817396 |
| #10 | LY6E | 192.5391 | 0.9956662 | 0.004908389 |
| #11 | SOD2 | 152.1733 | 1.214855 | 0.00746386 |
| #12 | SCO2 | 53.39916 | 1.659564 | 0.007273667 |
| #13 | PDIA3 | 191.4194 | 0.6695036 | 8.82645E-05 |
| #14 | AIF1 | 582.2145 | 0.8430264 | 0.006690409 |
| #15 | SLC31A2 | 24.26421 | 1.451713 | 0.001772667 |
| #16 | SAT1 | 809.3303 | 0.9536916 | 0.009026804 |
| #17 | CASP1 | 87.23621 | 0.8896548 | 0.00331111 |
| #18 | PSMB9 | 201.1459 | 1.081344 | 0.008698217 |
| #19 | BLVRA | 38.01017 | 1.0655 | 0.001806379 |
| #20 | PLAC8 | 270.4911 | 0.7098253 | 0.003296943 |
| #21 | MAFB | 57.83697 | 1.065289 | 0.004992742 |
| #22 | PSMA4 | 102.2819 | 0.9824251 | 0.007136935 |
| #23 | SNX10 | 66.34343 | 1.251423 | 0.008358826 |
| #24 | AAK1 | 249.4722 | -0.6613641 | 0.000916433 |
| #25 | FGL2 | 229.8065 | 1.14242 | 0.01164973 |
| #26 | RAB31 | 61.27285 | 0.9822807 | 0.005034739 |
| #27 | PSME2 | 202.914 | 1.61118 | 0.01480666 |
| #28 | WARS | 72.641 | 2.52139 | 0.01514413 |
| #29 | UBE2L6 | 54.38438 | 1.648584 | 0.01181487 |

| | | | | |
|---|---|---|---|---|
| **#30** | PSMB10 | 162.061 | 0.9036299 | 0.00907711 |
| **#31** | SERPINB1 | 159.8107 | 0.6054407 | 0.000262208 |
| **#32** | PSMB3 | 134.0157 | 0.6885632 | 0.002853126 |
| **#33** | MTPN | 214.952 | 0.5889395 | 0.000223543 |
| **#34** | FBXO6 | 16.28818 | 1.420632 | 0.001040734 |
| **#35** | PSMB8 | 157.2297 | 0.744258 | 0.005427581 |
| **#36** | IRF9 | 76.07531 | 0.8797513 | 0.005966076 |
| **#37** | ISG15 | 88.93194 | 1.476219 | 0.01387061 |
| **#38** | NAPA | 52.46777 | 0.8513312 | 0.002756861 |
| **#39** | ATP1B3 | 68.79183 | 0.7964792 | 0.003315166 |
| **#40** | BST2 | 146.5561 | 0.7813435 | 0.007375698 |
| **#41** | VMP1 | 88.2269 | 0.7100339 | 0.002110541 |
| **#42** | SNHG25 | 98.04268 | -1.376238 | 0.009249 |
| **#43** | MYD88 | 42.26308 | 0.7962985 | 9.75606E-05 |
| **#44** | RIPK2 | 24.81752 | 1.137336 | 0.002502036 |
| **#45** | PSTPIP2 | 28.59104 | 1.51307 | 0.00949458 |
| **#46** | ATG3 | 82.7728 | 0.6623572 | 0.000885241 |
| **#47** | PSMA2 | 103.1842 | 0.6282695 | 0.0009749 |
| **#48** | APOL6 | 91.63408 | 1.169253 | 0.01490142 |
| **#49** | LAP3 | 58.87503 | 2.283333 | 0.01878292 |
| **#50** | MARCKS | 114.6997 | 0.881333 | 0.01186633 |
| **#51** | MTHFD2 | 18.87297 | 1.285328 | 0.003297826 |
| **#52** | MT2A | 141.5404 | 1.489074 | 0.01913034 |
| **#53** | RSRP1 | 297.338 | -0.5844798 | 0.002179921 |
| **#54** | PBXIP1 | 94.57342 | -0.6877246 | 0.000392243 |
| **#55** | BCL2A1 | 40.7084 | 1.076513 | 0.008235589 |
| **#56** | GBP2 | 74.3969 | 1.702053 | 0.01836326 |
| **#57** | TRAF3IP3 | 225.4667 | -0.7160796 | 0.007725674 |
| **#58** | DYNLT1 | 61.22646 | 0.8047768 | 0.005237146 |
| **#59** | LPCAT2 | 23.62358 | 1.29705 | 0.006655451 |
| **#60** | HLA-DPA1 | 788.6352 | 0.7762951 | 0.01500907 |
| **#61** | PLSCR1 | 43.66707 | 1.207355 | 0.01286556 |

| | | | | |
|---|---|---|---|---|
| **#62** | CYBB | 208.5261 | 0.8353769 | 0.01443038 |
| **#63** | TXN | 111.4964 | 0.5881394 | 0.001118742 |
| **#64** | PLXDC2 | 45.42862 | 0.8126889 | 0.003766654 |
| **#65** | GSTO1 | 90.46233 | 0.7382364 | 0.007461116 |
| **#66** | CD38 | 24.1976 | 1.001172 | 0.002477258 |
| **#67** | ODF3B | 22.19166 | 1.204747 | 0.005789534 |
| **#68** | GLUL | 61.74834 | 0.7448171 | 0.004915195 |
| **#69** | CHD3 | 109.0066 | -0.8354072 | 0.008472509 |
| **#70** | SRGN | 618.1732 | 0.6422699 | 0.01150745 |
| **#71** | VAMP5 | 80.81437 | 1.979802 | 0.02412642 |
| **#72** | TNFAIP2 | 89.54648 | 0.864845 | 0.01404018 |
| **#73** | HLA-DMA | 147.575 | 0.722511 | 0.01160827 |
| **#74** | PSMA6 | 102.7678 | 0.5636318 | 0.000485218 |
| **#75** | MOB1A | 169.5137 | 0.5275524 | 0.000939246 |
| **#76** | GNG10 | 49.5884 | 0.7141534 | 0.002403197 |
| **#77** | SLAMF7 | 30.23277 | 0.8658504 | 0.002869589 |
| **#78** | SERPINA1 | 197.0563 | 0.7809628 | 0.01611892 |
| **#79** | IFI35 | 32.83117 | 1.381279 | 0.01611282 |
| **#80** | PYCARD | 150.1559 | 0.6838912 | 0.01105375 |
| **#81** | GLIPR2 | 78.19475 | 0.6883042 | 0.006422617 |
| **#82** | SORL1 | 158.106 | -0.5508051 | 0.000414733 |
| **#83** | IRF7 | 47.86457 | 1.113486 | 0.01581988 |
| **#84** | GBP5 | 50.38056 | 2.145067 | 0.025992 |
| **#85** | MCL1 | 239.2267 | 0.5673027 | 0.006123276 |
| **#86** | STAT1 | 91.89872 | 1.950308 | 0.02810676 |
| **#87** | GNB4 | 29.13478 | 1.039204 | 0.008821535 |
| **#88** | GCH1 | 21.82121 | 1.290323 | 0.01079564 |
| **#89** | SLC8A1 | 31.09864 | 1.05462 | 0.01046871 |
| **#90** | UBE2D1 | 39.88276 | 0.7306141 | 0.002402074 |
| **#91** | MXD1 | 40.28558 | 0.8479041 | 0.007629721 |
| **#92** | ACTB | 3,855 | 0.5741197 | 0.01199192 |
| **#93** | GABARAP | 451.647 | 0.6505976 | 0.01521945 |

| | | | | |
|---|---|---|---|---|
| **#94** | CSKMT | 37.44724 | -1.155314 | 0.00435012 |
| **#95** | PRKCA | 54.98711 | -0.9985678 | 0.008475287 |
| **#96** | TMC8 | 86.53219 | -0.7358427 | 0.006491393 |
| **#97** | TNFSF13B | 74.60338 | 1.244499 | 0.02466644 |
| **#98** | WDFY1 | 31.28444 | 0.8833754 | 0.006541272 |
| **#99** | FPR1 | 46.14713 | 1.240002 | 0.02120822 |
| **#100** | STOM | 36.20871 | 0.747142 | 0.003236762 |
| **#101** | DIP2A | 55.97285 | -0.8751868 | 0.006976583 |
| **#102** | GBP4 | 50.94555 | 1.915966 | 0.02953115 |
| **#103** | CARD16 | 125.044 | 0.6439035 | 0.01177343 |
| **#104** | ATOX1 | 55.58746 | 0.6956001 | 0.006775507 |
| **#105** | RAB20 | 8.046477 | 1.865663 | 0.00786654 |
| **#106** | PAK1 | 53.90599 | 0.6871846 | 0.006387026 |
| **#107** | GNS | 42.02302 | 0.6667083 | 0.002232994 |
| **#108** | ARPC3 | 635.0614 | 0.4786797 | 0.003594244 |
| **#109** | TAP1 | 55.80463 | 1.198683 | 0.02515214 |
| **#110** | PLEKHO1 | 127.2043 | 0.7558893 | 0.01925141 |
| **#111** | LYN | 173.1349 | 0.691589 | 0.01750989 |
| **#112** | CSF2RB | 16.92264 | 1.078215 | 0.004865574 |
| **#113** | S100A9 | 2,341 | 0.9693623 | 0.03335107 |
| **#114** | HLA-DRA | 1,608 | 0.6659029 | 0.02227294 |
| **#115** | NAMPT | 205.4292 | 0.9339365 | 0.02811568 |
| **#116** | BCL3 | 19.02403 | 1.143479 | 0.009451681 |
| **#117** | SFT2D2 | 66.93682 | 0.6546747 | 0.007975931 |
| **#118** | JDP2 | 17.27703 | 0.9737724 | 0.002485966 |
| **#119** | PLAAT4 | 179.9457 | 0.7691398 | 0.02267644 |
| **#120** | CNDP2 | 53.94765 | 0.8703895 | 0.01660523 |
| **#121** | TYROBP | 731.3584 | 0.7440518 | 0.02615405 |
| **#122** | FCGR2A | 59.54453 | 0.8811585 | 0.0184875 |
| **#123** | PILRA | 64.10478 | 0.711026 | 0.01134413 |
| **#124** | SPOCK2 | 125.1761 | -0.7112618 | 0.01324389 |
| **#125** | CPVL | 131.575 | 0.538817 | 0.005341546 |

| | | | | |
|---|---|---|---|---|
| #126 | STAT2 | 51.56906 | 1.120163 | 0.0245255 |
| #127 | SMCO4 | 18.72841 | 1.187817 | 0.01141135 |
| #128 | GNG5 | 178.9186 | 0.5279992 | 0.007002486 |
| #129 | PARP9 | 27.42306 | 1.603686 | 0.02594565 |
| #130 | RASA3 | 62.86212 | -0.6753003 | 0.003499522 |
| #131 | PRELID1 | 226.4883 | 0.6673992 | 0.02006014 |
| #132 | EPSTI1 | 39.84049 | 1.915125 | 0.03343816 |
| #133 | LCP1 | 523.9148 | 0.6145602 | 0.01914786 |
| #134 | CLEC2B | 140.6489 | 0.5158296 | 0.004714212 |
| #135 | LILRB1 | 49.45867 | 0.9763976 | 0.02161637 |
| #136 | HIST1H4C | 220.1684 | -0.4735615 | 0.001392293 |
| #137 | TNFSF10 | 75.73957 | 1.610244 | 0.03539857 |
| #138 | SQOR | 33.18346 | 0.9710474 | 0.01615142 |
| #139 | EPB41L3 | 21.74417 | 1.203202 | 0.01616219 |
| #140 | PLBD1 | 52.25088 | 0.8437258 | 0.01793887 |
| #141 | IFIT2 | 14.16542 | 2.103347 | 0.02663101 |
| #142 | APOBEC3A | 38.47575 | 1.824536 | 0.03489606 |
| #143 | ACAP1 | 174.2857 | -0.499244 | 0.003172439 |
| #144 | CASP4 | 104.4601 | 0.5847468 | 0.01034383 |
| #145 | SDCBP | 116.8721 | 0.4972493 | 0.002669969 |
| #146 | SKAP2 | 89.44363 | 0.5172624 | 0.002696972 |
| #147 | NCF2 | 122.0723 | 0.5701733 | 0.01071398 |
| #148 | EIF5A | 200.6135 | 0.4988911 | 0.006977178 |
| #149 | S100A11 | 478.8858 | 0.7524658 | 0.03031516 |
| #150 | TALDO1 | 164.2536 | 0.5536056 | 0.01157346 |
| #151 | GBP1 | 33.35953 | 2.765205 | 0.04047018 |
| #152 | FGD2 | 36.49806 | 0.7666076 | 0.01031714 |
| #153 | EIF2AK2 | 66.01941 | 0.8238418 | 0.02230124 |
| #154 | MYOF | 18.61774 | 1.915361 | 0.02969794 |
| #155 | BCL9L | 52.63021 | -0.7945592 | 0.009073739 |
| #156 | XKR8 | 11.0492 | 1.314083 | 0.007018772 |
| #157 | CLIC1 | 334.7764 | 0.571762 | 0.01668077 |

| | | | | |
|---|---|---|---|---|
| **#158** | RALB | 35.11818 | 0.6673455 | 0.003890243 |
| **#159** | RORA | 118.9641 | -0.5274609 | 0.003749386 |
| **#160** | MS4A6A | 122.9915 | 0.6405676 | 0.01918051 |
| **#161** | SORT1 | 15.56175 | 1.111016 | 0.009582169 |
| **#162** | IFI6 | 51.05087 | 1.315594 | 0.03620506 |
| **#163** | LACTB | 45.38494 | 0.684032 | 0.01070874 |
| **#164** | RHOA | 323.8111 | 0.4388001 | 0.002887811 |
| **#165** | LBH | 119.7913 | -0.7016282 | 0.01861679 |
| **#166** | SOCS3 | 56.72092 | 0.9542796 | 0.02888575 |
| **#167** | PMAIP1 | 48.2654 | 0.7279435 | 0.01462979 |
| **#168** | S100A8 | 1,389 | 1.019304 | 0.04681731 |
| **#169** | CTSB | 106.4246 | 0.6138325 | 0.01718541 |
| **#170** | CMTM6 | 120.2964 | 0.5431736 | 0.01131722 |
| **#171** | TNFRSF25 | 41.95217 | -0.9601901 | 0.01290987 |
| **#172** | RBCK1 | 65.52158 | 0.5802439 | 0.008223512 |
| **#173** | CALCOCO2 | 70.51741 | 0.6361305 | 0.01486008 |
| **#174** | LILRA5 | 50.99636 | 0.7847615 | 0.02073985 |
| **#175** | OAS3 | 12.78403 | 1.543418 | 0.02084159 |
| **#176** | ICAM1 | 14.96771 | 1.153759 | 0.01269932 |
| **#177** | ARPC5 | 306.6443 | 0.4306461 | 0.002989301 |
| **#178** | CD52 | 940.6831 | -0.4293451 | 0.004633249 |
| **#179** | IL16 | 89.09415 | -0.5551367 | 0.005505946 |
| **#180** | CTSS | 1,182 | 0.5609987 | 0.02382521 |
| **#181** | P2RY13 | 22.02729 | 0.966047 | 0.0148531 |
| **#182** | NCF1 | 155.2654 | 1.157925 | 0.04489176 |
| **#183** | MPHOSPH8 | 203.9169 | -0.4587899 | 0.003397205 |
| **#184** | CALHM6 | 86.54161 | 1.2598 | 0.04444754 |
| **#185** | LRRK2 | 58.82678 | 1.096435 | 0.03745223 |
| **#186** | ZNF267 | 29.649 | 0.8584792 | 0.01660976 |
| **#187** | GPBAR1 | 23.11723 | 0.9546603 | 0.01643083 |
| **#188** | RNF7 | 88.07746 | 0.4821415 | 0.002777844 |
| **#189** | PFKFB3 | 16.57569 | 1.28145 | 0.02125247 |

| | | | | |
|---|---|---|---|---|
| **#190** | GASK1B | 18.55594 | 0.8668825 | 0.006406882 |
| **#191** | BRD9 | 52.73255 | -0.6490696 | 0.005814429 |
| **#192** | PCGF5 | 65.09297 | 0.5004804 | 0.002067943 |
| **#193** | ATF5 | 15.69374 | 1.188831 | 0.01670789 |
| **#194** | IFIT3 | 11.40186 | 2.360722 | 0.03680483 |
| **#195** | OAS1 | 33.17188 | 0.9760653 | 0.02599384 |
| **#196** | ARF3 | 40.30766 | 0.5581671 | 0.001593198 |
| **#197** | LRRC25 | 56.86761 | 0.6823547 | 0.018461 |
| **#198** | CD68 | 119.7759 | 0.6094048 | 0.0223814 |
| **#199** | MARCH9 | 34.13854 | -0.9690898 | 0.0113985 |
| **#200** | TAP2 | 42.18626 | 0.7864118 | 0.0213911 |
| **#201** | FABP5 | 19.31981 | 0.7460613 | 0.001171759 |
| **#202** | LGALS9 | 60.91918 | 0.7704459 | 0.02698321 |
| **#203** | RAB8A | 78.1034 | 0.4798476 | 0.002589479 |
| **#204** | LUC7L3 | 210.5651 | -0.4607429 | 0.006344723 |
| **#205** | BCL11B | 279.5518 | -0.9233418 | 0.04489158 |
| **#206** | PLEKHA1 | 61.54461 | -0.5946063 | 0.006309495 |
| **#207** | SNHG7 | 111.0881 | -0.6217935 | 0.01831874 |
| **#208** | TCF7 | 189.1935 | -0.7770421 | 0.03627084 |
| **#209** | SPPL2A | 46.9002 | 0.5074389 | 0.000196874 |
| **#210** | DEK | 235.1446 | 0.4052546 | 0.001695519 |
| **#211** | FTH1 | 3,239 | 0.5746054 | 0.0324142 |
| **#212** | DRAP1 | 181.3957 | 0.475891 | 0.01127269 |
| **#213** | CD6 | 54.53776 | -0.7186936 | 0.01354441 |
| **#214** | GM2A | 17.10651 | 1.219933 | 0.02414238 |
| **#215** | TLR8 | 17.61432 | 0.9138176 | 0.01057456 |
| **#216** | ACSL1 | 18.32522 | 1.187158 | 0.02504064 |
| **#217** | RBM47 | 24.42817 | 0.8714818 | 0.01724394 |
| **#218** | NMT2 | 28.58133 | -0.7600297 | 0.00079204 |
| **#220** | HSP90AA1 | 569.0151 | 0.4262614 | 0.008117064 |
| **#221** | PSMB1 | 166.0609 | 0.4232719 | 0.002762993 |
| **#222** | TMSB10 | 4,959 | 0.3985808 | 0.006399041 |

| | | | | |
|---|---|---|---|---|
| #223 | NUDT16 | 24.91633 | 0.6436178 | 0.002012257 |
| #224 | BST1 | 24.86138 | 0.7363672 | 0.009099385 |
| #225 | FUOM | 20.02075 | 0.6896538 | 0.000759834 |
| #226 | LILRA2 | 28.81575 | 0.807295 | 0.01885076 |
| #227 | LILRB3 | 57.84639 | 0.6932188 | 0.02459032 |
| #228 | ANXA2 | 207.1029 | 0.60177 | 0.03123049 |
| #229 | JAK2 | 33.21346 | 1.172457 | 0.0392153 |
| #230 | LILRB2 | 81.78898 | 0.663401 | 0.02748469 |
| #231 | SYTL2 | 33.39764 | -0.937107 | 0.01321098 |
| #232 | HLA-B | 2,833 | 0.3683536 | 0.001786752 |
| #233 | PGK 1.00 | 194.703 | 0.4514686 | 0.009208394 |
| #234 | NLRP1 | 60.86117 | -0.6240447 | 0.01164976 |
| #235 | UBE2L3 | 100.2815 | 0.4263505 | 0.000482521 |
| #236 | ATF3 | 13.06442 | 1.349742 | 0.02391216 |
| #237 | SPTLC2 | 47.81363 | 0.6909844 | 0.02201974 |
| #238 | ITPKB | 44.63231 | -0.7382123 | 0.01240099 |
| #239 | LMO2 | 51.52841 | 0.6595581 | 0.02070347 |
| #240 | CTNNA1 | 34.14955 | 0.60532 | 0.006675587 |
| #241 | ABCA7 | 26.17458 | -0.7696077 | 0.000606233 |
| #242 | NUB1 | 99.30553 | 0.5757428 | 0.02213096 |
| #243 | CWF19L2 | 84.92154 | -0.5329442 | 0.00898338 |
| #244 | NLRP3 | 22.94638 | 0.8483496 | 0.0167755 |
| #245 | CAPZA2 | 97.08627 | 0.4384515 | 0.002226465 |
| #246 | OAS2 | 26.81481 | 1.16358 | 0.03684859 |
| #247 | CDC25B | 39.29063 | -0.7211369 | 0.008755609 |
| #248 | LMNB1 | 18.81195 | 1.113094 | 0.02731722 |
| #249 | ANXA5 | 183.6402 | 0.5787211 | 0.03032801 |
| #250 | TMEM179B | 45.32869 | 0.5649464 | 0.009048129 |
| #251 | CUL1 | 33.80161 | 0.7360095 | 0.0203676 |
| #252 | PIM1 | 49.14111 | 0.6698356 | 0.02257465 |
| #253 | CD1D | 23.6073 | 0.6297315 | 0.001647126 |
| #254 | PIK3AP1 | 56.96686 | 0.6491621 | 0.02349875 |

| | | | | |
|---|---|---|---|---|
| #255 | CALR | 223.7956 | 0.4538573 | 0.01317902 |
| #256 | ENO1 | 265.4821 | 0.3966118 | 0.003810751 |
| #257 | NLRC3 | 28.32631 | -0.8950067 | 0.008807065 |
| #258 | PSMB2 | 60.492 | 0.4763337 | 0.00359832 |
| #261 | CLEC2D | 162.0493 | -0.6361166 | 0.03385928 |
| #262 | MVP | 54.00115 | 0.7457051 | 0.03297325 |
| #263 | RAB7A | 94.25272 | 0.4156806 | 0.000374856 |
| #264 | PDIA6 | 93.81204 | 0.4336257 | 0.002907969 |
| #265 | TTC9 | 17.97848 | -1.079646 | 0.002169161 |
| #266 | SAMD4A | 7.006671 | 1.845619 | 0.02670025 |
| #267 | NMI | 46.50674 | 0.835554 | 0.03744537 |
| #268 | IGF2BP2 | 8.094321 | 1.117539 | 0.002548343 |
| #269 | ATP6V0B | 141.4401 | 0.4798383 | 0.01628838 |
| #270 | ANKH | 25.20019 | -0.7303083 | 0.000216938 |
| #271 | CBX7 | 24.4786 | -0.779214 | 0.001945043 |
| #272 | SP110 | 141.2066 | 0.556013 | 0.02883036 |
| #273 | PSAP | 635.8332 | 0.4831293 | 0.02632337 |
| #274 | MLLT6 | 91.46207 | -0.4394192 | 0.001195626 |
| #276 | SLC4A7 | 42.85759 | -0.7189214 | 0.01489073 |
| #277 | HLA-DMB | 121.3104 | 0.6413317 | 0.03927431 |
| #278 | B4GALT5 | 12.86694 | 0.8520038 | 0.003665426 |
| #279 | LILRA1 | 30.19311 | 0.8952145 | 0.03339408 |
| #280 | FCN1 | 507.3753 | 0.5905221 | 0.04272426 |
| #281 | EPB41 | 196.9539 | -0.5559634 | 0.02909283 |
| #282 | GUK1 | 203.3304 | 0.4496259 | 0.01517507 |
| #283 | TRIB1 | 11.23481 | 0.9187293 | 0.003761314 |
| #285 | THEM4 | 41.72111 | -0.8006646 | 0.02046948 |
| #287 | TRIM25 | 22.55461 | 0.6858133 | 0.008247279 |
| #288 | PURA | 49.48675 | -0.5937609 | 0.009667328 |
| #289 | PDE7A | 63.8749 | -0.8386705 | 0.03562291 |
| #290 | CD74 | 2,055 | 0.4829778 | 0.03129275 |
| #291 | GRINA | 50.56682 | 0.7344757 | 0.03504583 |

| | | | | |
|---|---|---|---|---|
| #292 | HNRNPH2 | 45.99532 | 0.460706 | 1.49855E-05 |
| #293 | AKNA | 135.7932 | -0.4838697 | 0.01537069 |
| #294 | RAB5C | 96.09468 | 0.5125715 | 0.02062162 |
| #295 | VAMP3 | 31.78636 | 0.5509351 | 0.003538232 |
| #296 | MALAT1 | 57,102 | -0.4328207 | 0.0217485 |
| #297 | ADAR | 140.7865 | 0.4953882 | 0.0226584 |
| #298 | NOP53 | 520.4771 | -0.5423119 | 0.03707565 |
| #299 | FGD3 | 76.39799 | -0.503642 | 0.01026657 |
| #300 | DUSP5 | 14.3002 | 0.7618023 | 0.002527604 |
| #301 | RCAN3 | 122.7982 | -0.7630417 | 0.04678607 |
| #302 | LIME1 | 160.0097 | -0.4982614 | 0.02075448 |
| #303 | LILRB4 | 21.4429 | 0.9669919 | 0.0318223 |
| #304 | PRKAR1A | 88.9195 | 0.4107871 | 0.001941113 |
| #305 | GAS6 | 6.165117 | 1.287028 | 0.003818917 |
| #306 | CHMP2A | 91.85306 | 0.4632453 | 0.01163515 |
| #307 | TPMT | 23.23076 | 0.577799 | 0.000608826 |
| #308 | HDAC7 | 38.1977 | -0.5683551 | 0.001712462 |
| #309 | RPS27 | 10,228 | -0.3846126 | 0.01211068 |
| #310 | DUSP6 | 40.72615 | 0.7925619 | 0.03821713 |
| #312 | OASL | 5.96631 | 1.509603 | 0.01282762 |
| #313 | TNRC6C | 49.44785 | -0.7251845 | 0.02372579 |
| #314 | RILPL2 | 74.40701 | 0.5787014 | 0.02876949 |
| #315 | CERK | 40.99389 | -0.5735307 | 0.004576408 |
| #316 | RPS29 | 6,091 | -0.3956023 | 0.01546076 |
| #317 | B2M | 9,491 | 0.3712562 | 0.01103284 |
| #318 | GRAMD1A | 40.03293 | -0.5531036 | 0.0023015 |
| #319 | HOTAIRM1 | 40.15361 | 0.604698 | 0.02083009 |
| #320 | SAP25 | 26.44426 | -0.7028675 | 0.003765756 |
| #322 | GLRX | 115.5644 | 0.5098284 | 0.02715843 |
| #323 | TMEM63A | 40.65264 | -0.726631 | 0.01988763 |
| #324 | ATP8B2 | 26.87494 | -0.9962726 | 0.01942064 |
| #325 | PYGL | 23.8184 | 0.5984768 | 0.004884322 |

| | | | | |
|---|---|---|---|---|
| **#326** | CMPK2 | 6.602384 | 1.660466 | 0.02618555 |
| **#327** | BNIP2 | 94.34854 | 0.4246971 | 0.007393293 |
| **#328** | SUMO3 | 68.52382 | 0.4162164 | 0.00201498 |
| **#329** | PTPRE | 124.6451 | 0.4746292 | 0.02187435 |
| **#330** | MIAT | 16.54283 | -1.013412 | 0.003295605 |
| **#331** | CAP1 | 259.9441 | 0.385109 | 0.009042134 |
| **#332** | PLEKHB2 | 43.61235 | 0.469309 | 0.00332458 |
| **#333** | PSMA3 | 56.08227 | 0.4467028 | 0.003885692 |
| **#334** | TUBA1B | 209.9756 | 0.3953951 | 0.0104301 |
| **#335** | HLA-C | 2,202 | 0.4115594 | 0.02232081 |
| **#336** | ATP6V0D1 | 72.03541 | 0.5397026 | 0.02642391 |
| **#337** | DNAJA1 | 123.5896 | 0.5357629 | 0.03465021 |
| **#338** | RNF149 | 111.3722 | 0.4909259 | 0.02500764 |
| **#339** | MRI1 | 24.2336 | -0.7156664 | 0.003350622 |
| **#340** | PRDX3 | 31.9102 | 0.5748441 | 0.01215615 |
| **#341** | REC8 | 11.02687 | 0.7906111 | 0.000245456 |
| **#342** | PSMB6 | 86.59923 | 0.4555792 | 0.01378963 |
| **#343** | CNIH4 | 28.94365 | 0.5021289 | 0.000208283 |
| **#344** | CTSZ | 171.1646 | 0.4936864 | 0.03200521 |
| **#345** | GRN | 156.78 | 0.5751783 | 0.04633571 |
| **#346** | ATP6V1A | 26.79805 | 0.6302658 | 0.01478047 |
| **#347** | KLF4 | 80.58967 | 0.5399539 | 0.02960934 |
| **#348** | CLEC7A | 92.59193 | 0.5868358 | 0.03942518 |
| **#349** | ATP6V1B2 | 35.75306 | 0.5892723 | 0.01803008 |
| **#350** | ATXN2 | 42.764 | -0.5325771 | 0.004268735 |
| **#351** | DUSP3 | 14.00745 | 0.8011962 | 0.009741046 |
| **#352** | RBM3 | 290.4851 | 0.3796719 | 0.01023596 |
| **#354** | IL15 | 11.15465 | 1.061734 | 0.01972736 |
| **#355** | CD226 | 48.77099 | -0.5695102 | 0.01342782 |
| **#356** | ATP2A3 | 46.9068 | -0.4914515 | 0.002379107 |
| **#357** | VRK2 | 17.83338 | 0.8365578 | 0.02243449 |
| **#358** | FKBP1A | 153.276 | 0.522635 | 0.03850066 |

| | | | | |
|---|---|---|---|---|
| **#359** | DECR1 | 59.22708 | 0.4301854 | 0.003924926 |
| **#360** | SRA1 | 36.59085 | 0.4768202 | 0.002597624 |
| **#361** | RGS2 | 179.652 | 0.5459015 | 0.04532497 |
| **#362** | TMEM219 | 93.18585 | 0.409116 | 0.007597456 |
| **#363** | ZNF503 | 10.42829 | 1.091699 | 0.02087975 |
| **#366** | OGT | 112.9201 | -0.5803807 | 0.03781309 |
| **#367** | LTB | 836.3367 | -0.5211701 | 0.04936029 |
| **#369** | ANKZF1 | 31.43522 | -0.563011 | 0.001610362 |
| **#370** | SH3BGRL | 205.1689 | 0.3919719 | 0.01264258 |
| **#371** | BID | 84.51529 | 0.5959645 | 0.04379122 |
| **#372** | HERPUD1 | 155.5204 | 0.483377 | 0.03252077 |
| **#373** | RGCC | 36.83564 | -0.8137008 | 0.02917824 |
| **#374** | BACH1 | 65.89884 | 0.5614031 | 0.0323947 |
| **#376** | VAMP8 | 196.7711 | 0.4339619 | 0.02298804 |
| **#377** | RNF213 | 372.9126 | 0.4800396 | 0.03887158 |
| **#378** | HHEX | 36.38078 | 0.5266085 | 0.0118668 |
| **#379** | APOL3 | 32.33678 | 0.8197198 | 0.04812659 |
| **#381** | MSRB1 | 24.31224 | 0.8147228 | 0.03757401 |
| **#382** | HERC2 | 50.60141 | -0.5907204 | 0.02147754 |
| **#383** | MSRB2 | 18.3352 | 0.7833684 | 0.02260952 |
| **#384** | STX2 | 23.79465 | -0.6995207 | 0.005515566 |
| **#385** | CREBZF | 55.16502 | -0.5300846 | 0.01586346 |
| **#386** | PRDM1 | 29.91548 | 0.6511944 | 0.02734215 |
| **#387** | SCIMP | 39.21707 | 0.6966581 | 0.04242723 |
| **#388** | NFIL3 | 23.18795 | 0.8341975 | 0.03861344 |
| **#389** | SNN | 26.24571 | 0.5845774 | 0.01222398 |
| **#391** | PDCD7 | 48.17577 | -0.8505292 | 0.04702668 |
| **#392** | EMILIN2 | 47.52071 | 0.6185227 | 0.03779099 |
| **#393** | ATM | 305.5287 | -0.5119857 | 0.04719374 |
| **#394** | VCPIP1 | 36.22511 | 0.4849578 | 0.00656275 |
| **#395** | LTBP3 | 23.32619 | -0.8652372 | 0.01650801 |
| **#396** | DNASE1 | 19.07366 | -0.929106 | 0.01130943 |

| | | | | |
|---|---|---|---|---|
| #397 | KIF21B | 17.18279 | -0.8462837 | 0.002992913 |
| #398 | GYG1 | 24.56249 | 0.56586 | 0.008181777 |
| #399 | PDP1 | 26.16251 | 0.5520723 | 0.008134373 |
| #400 | BISPR | 15.2807 | 0.7389525 | 0.01295212 |
| #401 | DYSF | 5.702668 | 1.690751 | 0.03139298 |
| #403 | ANKRD13A | 51.04406 | 0.4734695 | 0.01456645 |
| #405 | DGKZ | 93.14635 | -0.4866731 | 0.02394694 |
| #407 | ISOC2 | 13.68839 | 0.6709245 | 0.002184379 |
| #408 | ZFP14 | 23.18991 | -0.6249926 | 0.001117007 |
| #409 | COX5B | 267.5533 | 0.3526528 | 0.008909562 |
| #410 | TRIM73 | 36.05186 | -0.8720196 | 0.03923031 |
| #411 | KANSL1 | 134.4481 | -0.3732262 | 0.005125721 |
| #412 | AKT3 | 33.007 | -0.6513687 | 0.01623399 |
| #414 | MAP4K2 | 29.00379 | -0.5736292 | 0.003327657 |
| #415 | RNF19B | 15.7364 | 0.6380286 | 0.004048457 |
| #416 | AP5B1 | 21.92079 | 0.720061 | 0.02649806 |
| #417 | CREG1 | 30.60183 | 0.5543047 | 0.01517032 |
| #419 | NDUFA9 | 37.757 | 0.4385118 | 0.000933916 |
| #420 | AGPAT3 | 35.24877 | 0.5194285 | 0.01475121 |
| #421 | PSMD7 | 83.03316 | 0.3696193 | 0.00233927 |
| #423 | BLOC1S1 | 154.0669 | 0.4088004 | 0.01985867 |
| #425 | TNS3 | 11.19836 | 0.7830609 | 0.005785725 |
| #426 | HSBP1 | 68.05094 | 0.4978716 | 0.02871716 |
| #427 | LY96 | 26.73805 | 0.5920088 | 0.01712957 |
| #428 | CD300E | 44.24331 | 0.624058 | 0.04218756 |
| #429 | NCOA4 | 90.19239 | 0.3978096 | 0.009972895 |
| #431 | CBFA2T2 | 22.62762 | -0.6283165 | 0.00154259 |
| #434 | UBE2J1 | 58.73463 | 0.4576484 | 0.01652426 |
| #435 | CNTLN | 12.16884 | 0.7571201 | 0.00749597 |
| #436 | LDLRAP1 | 39.00427 | -0.8400451 | 0.0428187 |
| #437 | LAMP2 | 54.87585 | 0.4619781 | 0.01604143 |
| #438 | TSEN34 | 22.89659 | 0.5361055 | 0.004651931 |

| | | | | |
|---|---|---|---|---|
| **#439** | ZNF276 | 52.20319 | -0.4297289 | 0.000717198 |
| **#440** | OAZ1 | 870.6047 | 0.4003482 | 0.02951272 |
| **#441** | UBE2D3 | 283.723 | 0.3119375 | 0.001504127 |
| **#442** | RPS21 | 3,617 | -0.3540521 | 0.01776066 |
| **#443** | NFIA | 23.75545 | -0.7156553 | 0.01056668 |
| **#444** | ZMYND8 | 50.53788 | -0.5123871 | 0.0149732 |
| **#445** | KHDC4 | 39.68756 | -0.4942571 | 0.004340224 |
| **#446** | SIGIRR | 109.5637 | -0.5512359 | 0.04345246 |
| **#447** | TENT5A | 44.36905 | 0.5723311 | 0.03397011 |
| **#448** | RASGRP1 | 69.7576 | -0.485425 | 0.01994482 |
| **#450** | CARD11 | 31.96712 | -0.55002 | 0.005511901 |
| **#451** | RHOQ | 42.5059 | 0.4971417 | 0.01784489 |
| **#452** | TAPBP | 220.5985 | 0.4117797 | 0.02741201 |
| **#453** | PTGS2 | 14.17996 | 0.9424414 | 0.0339417 |
| **#454** | TBC1D10C | 139.1536 | -0.5445456 | 0.04959324 |
| **#456** | IRAK3 | 47.4818 | 0.5564769 | 0.03515296 |
| **#457** | FAS | 11.96035 | 0.8474458 | 0.01741093 |
| **#458** | PSMA7 | 234.9672 | 0.3518438 | 0.01149215 |
| **#459** | SFI1 | 41.45899 | -0.615597 | 0.0251585 |
| **#460** | DNAJC15 | 67.72073 | 0.4661614 | 0.02493189 |
| **#461** | PSMA5 | 81.45468 | 0.4410423 | 0.02207304 |
| **#462** | H3F3A | 1,210 | 0.3496701 | 0.01755558 |
| **#463** | AP2S1 | 132.8661 | 0.4415056 | 0.03055149 |
| **#464** | SAMD1 | 21.98165 | -0.6194496 | 0.001547277 |
| **#465** | INPP4A | 48.06731 | -0.4566862 | 0.004694202 |
| **#466** | PHPT1 | 66.07265 | 0.3958433 | 0.006776512 |
| **#467** | RNF144B | 22.76586 | 0.8335358 | 0.04871814 |
| **#469** | SNHG12 | 23.13412 | -0.6215435 | 0.003561423 |
| **#470** | LRRC59 | 21.93676 | 0.5014913 | 0.000593059 |
| **#471** | ADAP2 | 22.57168 | 0.643837 | 0.02286427 |
| **#472** | STAT5B | 45.01535 | -0.5159332 | 0.01390202 |
| **#474** | COX8A | 222.0889 | 0.384586 | 0.02159732 |

| | | | | |
|---|---|---|---|---|
| **#477** | MPP1 | 23.33004 | 0.5347861 | 0.0076988 |
| **#478** | ATP6V0C | 229.2737 | 0.3718791 | 0.01853205 |
| **#479** | HLA-E | 1,235 | 0.2864394 | 0.001345926 |
| **#480** | XPC | 55.14504 | -0.4384126 | 0.006034788 |
| **#482** | FERMT3 | 92.77556 | 0.3575475 | 0.004519747 |
| **#483** | ZRANB2 | 158.1027 | -0.4359144 | 0.02967169 |
| **#484** | CEP290 | 31.75683 | -0.6186467 | 0.01713833 |
| **#486** | CLTA | 85.74493 | 0.3630135 | 0.005024533 |
| **#488** | RGS1 | 19.63101 | 0.8597658 | 0.04785403 |
| **#490** | DNAJC3 | 53.34299 | 0.4241988 | 0.01087497 |
| **#491** | DDX17 | 356.577 | -0.4017701 | 0.03111995 |
| **#492** | CFAP36 | 25.33699 | -0.6256899 | 0.00917938 |
| **#493** | NPHP3 | 26.34945 | -0.5704958 | 0.004170474 |
| **#494** | SNHG19 | 15.42873 | -0.7516639 | 6.07616E-05 |
| **#495** | DGKD | 68.98437 | -0.5238155 | 0.03317474 |
| **#496** | RHOU | 9.159791 | 0.8526114 | 0.008686763 |
| **#497** | TPI1 | 185.7968 | 0.4308091 | 0.03634845 |
| **#498** | TRAFD1 | 14.60498 | 0.9468201 | 0.04196083 |
| **#499** | KIAA0355 | 42.81673 | -0.5606858 | 0.02168576 |
| **#500** | P2RX7 | 12.05227 | 0.9307756 | 0.03051755 |
| **#501** | SLC35E2B | 17.9638 | -0.6854493 | 0.001504619 |
| **#502** | BLOC1S6 | 70.11354 | 0.452996 | 0.02648907 |
| **#503** | NSUN5 | 33.50995 | -0.4738677 | 0.000706842 |
| **#505** | SLC1A5 | 16.80334 | 0.6420157 | 0.01359271 |
| **#508** | KYNU | 23.99987 | 0.6092412 | 0.02380264 |
| **#509** | WAS | 128.5484 | 0.3186541 | 0.000796095 |
| **#510** | BAZ1A | 104.8145 | 0.4526224 | 0.03491634 |
| **#512** | MBD2 | 71.39823 | 0.4654534 | 0.03102557 |
| **#513** | RUFY3 | 38.90902 | 0.5544712 | 0.03327848 |
| **#514** | BRD8 | 20.49936 | -0.6380069 | 0.003307158 |
| **#515** | RPL36 | 2,884 | -0.3095846 | 0.01086368 |
| **#516** | CPPED1 | 66.2166 | 0.4937978 | 0.03681009 |

| | | | | |
|---|---|---|---|---|
| **#517** | ZBTB25 | 36.05574 | -0.5945974 | 0.02154023 |
| **#518** | SNHG6 | 411.8845 | -0.3029313 | 0.004225157 |
| **#519** | GPAT3 | 13.61854 | 0.6718612 | 0.008657612 |
| **#520** | RPL37 | 5,399 | -0.3076897 | 0.01143102 |
| **#521** | ODF2L | 72.00995 | -0.438658 | 0.01646507 |
| **#522** | GKAP1 | 15.50889 | -0.7344853 | 0.000294391 |
| **#523** | MAP2K6 | 16.65387 | -0.7079397 | 0.001357495 |
| **#524** | COX7B | 174.8216 | 0.331073 | 0.007700313 |
| **#526** | ELF4 | 14.1038 | 0.595759 | 0.001798733 |
| **#527** | DNPEP | 31.54606 | 0.5951137 | 0.03589723 |
| **#528** | CAPZA1 | 129.3501 | 0.3173645 | 0.001658172 |
| **#529** | VDAC1 | 76.70572 | 0.396311 | 0.01513374 |
| **#531** | PRRG4 | 6.619518 | 1.2386 | 0.02679746 |
| **#532** | NOD2 | 8.77353 | 0.8121163 | 0.005507424 |
| **#533** | EAF1 | 15.02686 | 0.5870299 | 0.003122389 |
| **#537** | GCC2 | 212.401 | -0.3685145 | 0.01952944 |
| **#538** | DOK3 | 17.56266 | 0.6214845 | 0.01528036 |
| **#539** | TRGC2 | 33.8966 | -0.6613734 | 0.03058577 |
| **#540** | SCPEP1 | 59.27453 | 0.4502025 | 0.02559833 |
| **#541** | NUCB1 | 67.62161 | 0.4504881 | 0.02896126 |
| **#542** | KLF6 | 296.3571 | 0.3661654 | 0.02580493 |
| **#543** | TICAM2 | 7.197909 | 0.9927139 | 0.01275456 |
| **#544** | LAT | 86.79185 | -0.5244744 | 0.0468914 |
| **#545** | CGAS | 14.4719 | 0.7575842 | 0.02595776 |
| **#546** | ADA2 | 80.67871 | 0.4450503 | 0.03207368 |
| **#547** | PPP1R3E | 19.43828 | -0.7733893 | 0.01483171 |
| **#548** | RCN1 | 12.03451 | 0.7216795 | 0.01152449 |
| **#549** | CDC14A | 57.83547 | -0.5400336 | 0.03624726 |
| **#550** | GHITM | 110.6581 | 0.3134656 | 0.000534759 |
| **#552** | RNF13 | 61.40387 | 0.3880621 | 0.009357572 |
| **#553** | ECHS1 | 50.23571 | 0.3782532 | 0.002811325 |
| **#555** | FGFR1OP | 14.68074 | -0.73043 | 0.00032352 |

| | | | | |
|---|---|---|---|---|
| **#557** | ATP5F1B | 144.2301 | 0.3747397 | 0.02188681 |
| **#558** | IL15RA | 4.860121 | 1.240899 | 0.0114532 |
| **#559** | DERL1 | 42.75912 | 0.3937837 | 0.002494322 |
| **#560** | ERP44 | 61.36581 | 0.3491839 | 0.000803058 |
| **#561** | TRAF1 | 11.65373 | -0.9402093 | 0.002203366 |
| **#562** | SUPT4H1 | 70.49578 | 0.4113063 | 0.02106899 |
| **#563** | APOL1 | 14.42711 | 0.88042 | 0.04342443 |
| **#564** | WASHC2C | 33.26428 | 0.4771548 | 0.01519212 |
| **#565** | SLC5A3 | 26.81203 | -0.5109495 | 0.002036191 |
| **#567** | CHMP2B | 42.02477 | 0.4050993 | 0.005179428 |
| **#568** | SUB1 | 302.3573 | 0.2818526 | 0.001314474 |
| **#570** | RAB2A | 99.13928 | 0.3475594 | 0.008678367 |
| **#571** | CASP7 | 10.15581 | 0.8746354 | 0.02268816 |
| **#572** | POLE4 | 73.70036 | 0.3692676 | 0.01040561 |
| **#575** | CTSO | 16.22464 | 0.5681778 | 0.005547619 |
| **#576** | AKR1A1 | 52.68907 | 0.4778353 | 0.03375393 |
| **#577** | VPS29 | 81.93405 | 0.3221786 | 4.63495E-05 |
| **#578** | DENND5A | 24.96684 | 0.5284281 | 0.01693622 |
| **#579** | HELZ2 | 17.11314 | 0.7670255 | 0.03994739 |
| **#580** | STAT3 | 129.9576 | 0.38366 | 0.02535778 |
| **#582** | UNC93B1 | 52.46941 | 0.4769616 | 0.03433615 |
| **#583** | ARPC2 | 617.5552 | 0.4017614 | 0.04719762 |
| **#584** | TRIM52 | 37.76516 | -0.4800511 | 0.01058136 |
| **#585** | LDHA | 151.7048 | 0.3728353 | 0.02417459 |
| **#586** | ARL8B | 29.92178 | 0.4629107 | 0.009954823 |
| **#587** | SCAI | 17.75494 | -0.6363717 | 0.002018657 |
| **#588** | RP2 | 19.3277 | 0.5290592 | 0.007687159 |
| **#590** | TMEM167A | 61.58346 | 0.3471136 | 0.001780311 |
| **#591** | HCK | 71.21536 | 0.4896837 | 0.04895929 |
| **#594** | PDE4DIP | 20.71567 | -0.599741 | 0.004953126 |
| **#595** | AGAP2 | 13.96601 | -0.7377667 | 0.000809155 |
| **#596** | ATP6V1F | 123.7826 | 0.3939074 | 0.02888658 |

| | | | | |
|---|---|---|---|---|
| **#597** | SYK | 75.89594 | 0.3774081 | 0.01517835 |
| **#598** | SNAP23 | 63.62101 | 0.3682722 | 0.008329534 |
| **#599** | ZC3H8 | 34.1728 | -0.4597373 | 0.003653699 |
| **#600** | AP3S1 | 42.16325 | 0.4134141 | 0.009080226 |
| **#601** | RBP7 | 32.74254 | 0.5883833 | 0.04585956 |
| **#602** | FTX | 83.42009 | -0.4861916 | 0.04400875 |
| **#603** | NUP210 | 54.71121 | -0.4046694 | 0.007099618 |
| **#604** | S100PBP | 25.22752 | -0.5740719 | 0.01128977 |
| **#606** | ABCA5 | 15.12151 | -0.7432531 | 0.004779247 |
| **#607** | GLG1 | 113.7682 | -0.3173054 | 0.002079267 |
| **#608** | EHMT2 | 28.03921 | -0.4751683 | 0.000891047 |
| **#612** | TMEM71 | 34.06314 | -0.5068531 | 0.01439431 |
| **#613** | LRCH3 | 32.25328 | -0.4763915 | 0.006089436 |
| **#614** | TARSL2 | 34.68379 | -0.5598909 | 0.02499555 |
| **#615** | ACTR2 | 345.255 | 0.3332325 | 0.02272908 |
| **#617** | ILK | 38.82038 | 0.4673812 | 0.02380481 |
| **#618** | CA5B | 35.10257 | -0.4985977 | 0.01440485 |
| **#619** | BCL6 | 27.39996 | 0.524732 | 0.02474778 |
| **#620** | ATP6V1G1 | 261.1516 | 0.2844121 | 0.004048198 |
| **#621** | PML | 20.1388 | 0.6790726 | 0.04110939 |
| **#622** | SSBP2 | 36.60155 | -0.5732836 | 0.03061411 |
| **#623** | ACOT9 | 23.14544 | 0.5252607 | 0.01784481 |
| **#624** | RNF114 | 54.64224 | 0.3895538 | 0.0132472 |
| **#625** | ASGR2 | 11.87912 | 0.9500925 | 0.04913424 |
| **#626** | EIF6 | 59.57712 | 0.4370159 | 0.03144783 |
| **#628** | CBLB | 53.52154 | -0.4785927 | 0.02868377 |
| **#629** | SMAP2 | 189.1153 | 0.3817321 | 0.03617371 |
| **#630** | SEC61B | 193.3712 | 0.34477 | 0.02281161 |
| **#631** | CKAP4 | 25.2051 | 0.5064886 | 0.01778532 |
| **#632** | HRH2 | 28.3914 | 0.5007988 | 0.02191157 |
| **#634** | IRF3 | 82.16816 | -0.4277143 | 0.02953819 |
| **#635** | CIR1 | 89.64935 | 0.3160898 | 0.003055514 |

| | | | | |
|---|---|---|---|---|
| **#637** | PKM | 165.2425 | 0.3615005 | 0.02758456 |
| **#638** | PRCP | 38.6033 | 0.444969 | 0.01821364 |
| **#640** | FOXN2 | 34.7195 | 0.4530589 | 0.0167364 |
| **#641** | ENOSF1 | 18.28908 | -0.6443107 | 0.007317933 |
| **#642** | MTHFR | 26.88478 | 0.4995715 | 0.0199047 |
| **#643** | ATP6AP2 | 121.6391 | 0.3224048 | 0.009317938 |
| **#644** | TOB2 | 23.66232 | -0.6403041 | 0.02102893 |
| **#645** | KAT6B | 107.3516 | -0.4348636 | 0.0403063 |
| **#646** | TIMM8B | 55.87007 | 0.4061938 | 0.02096292 |
| **#648** | STAC3 | 9.800049 | 0.7509315 | 0.01241989 |
| **#649** | KIAA0930 | 36.17058 | 0.4654242 | 0.02276704 |
| **#650** | ABI2 | 24.08292 | -0.5134007 | 0.003465795 |
| **#651** | NEXN | 4.333201 | 1.675168 | 0.04625152 |
| **#652** | SNX6 | 115.5329 | 0.3171359 | 0.007669664 |
| **#653** | COQ10A | 14.2131 | -0.8566858 | 0.01311492 |
| **#654** | CENPT | 15.32253 | -0.6299716 | 2.47317E-05 |
| **#655** | ACPP | 9.946032 | 0.9342976 | 0.03695123 |
| **#656** | INTS6 | 68.19872 | -0.4078145 | 0.01930561 |
| **#657** | LTBR | 15.77355 | 0.6303325 | 0.02228879 |
| **#658** | CDC42 | 525.6729 | 0.2777566 | 0.007738208 |
| **#659** | ETV6 | 42.78966 | 0.4688245 | 0.03286289 |
| **#661** | NUMA1 | 67.49002 | -0.3881181 | 0.01319174 |
| **#663** | H2AFY | 162.2622 | 0.3661314 | 0.03112081 |
| **#664** | EPHB6 | 21.69828 | -0.5775297 | 0.008648558 |
| **#665** | BCKDK | 19.05384 | 0.6095825 | 0.02963061 |
| **#666** | KLHDC7B | 4.622393 | 1.066924 | 0.005221912 |
| **#667** | GRB2 | 189.0158 | 0.3566388 | 0.0301162 |
| **#668** | CBR1 | 22.9135 | 0.5621333 | 0.02895777 |
| **#669** | MKRN1 | 80.65173 | 0.3383812 | 0.00927301 |
| **#670** | HSPA6 | 8.140223 | 1.05428 | 0.03881863 |
| **#671** | PLIN3 | 32.22665 | 0.4904151 | 0.02749607 |
| **#672** | GAS5 | 393.2824 | -0.3273436 | 0.02513071 |

| | | | | |
|---|---|---|---|---|
| **#673** | SNHG8 | 241.2729 | -0.295701 | 0.008107162 |
| **#675** | WWP1 | 48.74584 | -0.4974776 | 0.03300303 |
| **#677** | PWWP3A | 24.18956 | -0.4904224 | 0.001736767 |
| **#679** | PRR13 | 183.3543 | 0.3099761 | 0.01302486 |
| **#681** | SYNRG | 91.8242 | -0.3982284 | 0.02795506 |
| **#684** | CD163 | 23.34452 | 0.4796223 | 0.01186392 |
| **#686** | PHF1 | 40.31558 | -0.4451162 | 0.01258968 |
| **#688** | RPL38 | 2,031 | -0.3411608 | 0.03975094 |
| **#690** | DSE | 23.43829 | 0.5932406 | 0.03973453 |
| **#691** | TRIM69 | 17.18274 | 0.7234473 | 0.04816984 |
| **#692** | TMEM116 | 11.10126 | -0.8140483 | 0.001619426 |
| **#693** | MFSD4B | 17.9945 | -0.5762433 | 0.002090974 |
| **#694** | TPM3 | 561.3033 | 0.3633138 | 0.04578454 |
| **#696** | SNF8 | 65.87484 | 0.3896154 | 0.02371276 |
| **#697** | EIF4A2 | 283.7182 | -0.29294 | 0.01054015 |
| **#698** | CMIP | 60.12447 | 0.4236609 | 0.03352151 |
| **#700** | KATNBL1 | 41.65441 | -0.4222463 | 0.008820427 |
| **#701** | CEP250 | 15.32839 | -0.6213165 | 0.001041506 |
| **#702** | NFAM1 | 22.6695 | 0.4885776 | 0.01384107 |
| **#703** | SYNGR2 | 69.41798 | 0.3605847 | 0.01537318 |
| **#704** | ZFAS1 | 462.6236 | -0.2450296 | 3.13593E-05 |
| **#705** | RRAGC | 24.78763 | 0.4161904 | 0.000749757 |
| **#706** | SLC25A42 | 18.11705 | -0.5784695 | 0.003220289 |
| **#707** | GRAMD1B | 9.412941 | 0.6716938 | 0.00476894 |
| **#708** | ARF1 | 175.9576 | 0.3331277 | 0.02368384 |
| **#709** | CEP126 | 12.5746 | -0.7907645 | 0.006094393 |
| **#710** | CCDC66 | 54.83261 | -0.4651672 | 0.03334007 |
| **#712** | PASK | 23.73032 | -0.6677311 | 0.03089022 |
| **#713** | EIF4E2 | 48.36849 | 0.3992381 | 0.01887202 |
| **#714** | SAP18 | 184.9894 | 0.2897771 | 0.008157751 |
| **#715** | PPP6R2 | 56.10546 | -0.4862843 | 0.04241619 |
| **#716** | MYDGF | 74.06264 | 0.3970221 | 0.03277353 |

| | | | | |
|---|---|---|---|---|
| #718 | DAZAP2 | 236.328 | 0.2923064 | 0.01189622 |
| #719 | TRAM1 | 116.098 | 0.2976528 | 0.005745551 |
| #722 | NAP1L4 | 108.9958 | -0.3224173 | 0.009592881 |
| #723 | NCF4 | 29.55474 | 0.5626929 | 0.04926099 |
| #724 | ARHGAP45 | 143.6839 | -0.3226316 | 0.01491091 |
| #726 | RTCB | 33.12567 | 0.3892972 | 0.004642477 |
| #728 | SERPINB9 | 47.04543 | 0.4695122 | 0.04542206 |
| #729 | RBM14 | 28.09937 | -0.4559265 | 0.003413043 |
| #733 | DRAM1 | 6.37593 | 0.9360582 | 0.01567822 |
| #734 | TMEM170B | 26.71686 | 0.5735451 | 0.04787184 |
| #735 | DTNBP1 | 31.15468 | 0.4241285 | 0.01241186 |
| #736 | ANXA4 | 33.61205 | 0.49716 | 0.03872081 |
| #737 | MINDY2 | 25.80092 | -0.4522853 | 0.000801115 |
| #738 | MBOAT7 | 32.42536 | 0.50947 | 0.04123035 |
| #739 | CD5 | 25.724 | -0.6321276 | 0.03422126 |
| #740 | SLC35B1 | 19.72803 | 0.4807924 | 0.00868898 |
| #741 | RALGAPA1 | 39.12724 | -0.5521858 | 0.04313012 |
| #742 | SHISAL2A | 18.69221 | -0.7519417 | 0.02905157 |
| #744 | SERF2 | 1,009 | 0.3150307 | 0.03366287 |
| #745 | SZRD1 | 51.844 | 0.3257919 | 0.000983999 |
| #746 | TECPR1 | 32.18036 | -0.4375693 | 0.00682368 |
| #747 | RSBN1L | 91.96856 | -0.3705428 | 0.02515465 |
| #748 | KCNQ1OT1 | 35.9641 | -0.5910972 | 0.04894219 |
| #749 | MRNIP | 20.14969 | -0.633161 | 0.020172 |
| #750 | UQCRFS1 | 96.52064 | 0.311281 | 0.01019235 |
| #751 | PLXND1 | 22.13502 | -0.5774708 | 0.01661189 |
| #752 | CD33 | 32.88514 | 0.4754115 | 0.03306035 |
| #753 | CABIN1 | 32.63283 | -0.4175157 | 0.003186577 |
| #754 | TTC39B | 23.71159 | -0.5056157 | 0.008156888 |
| #755 | RASGRP2 | 107.3413 | -0.292184 | 0.001866968 |
| #756 | ANKS3 | 10.6155 | -0.7808972 | 0.001111759 |
| #757 | IDH3A | 18.87635 | 0.4660987 | 0.004724637 |

| | | | | |
|---|---|---|---|---|
| **#760** | TXNDC17 | 43.11698 | 0.4025857 | 0.02014114 |
| **#761** | ZNF559 | 13.00424 | -0.6977624 | 0.004280574 |
| **#762** | ZNF44 | 28.3918 | -0.4737175 | 0.01013668 |
| **#769** | RBX1 | 115.3086 | 0.3445645 | 0.02761817 |
| **#770** | PCNX4 | 32.96836 | -0.4279435 | 0.00668644 |
| **#773** | SCARB2 | 38.86922 | 0.4722469 | 0.04274455 |
| **#774** | PPWD1 | 32.84269 | -0.5104732 | 0.02799595 |
| **#775** | GNB2 | 132.6172 | 0.3836118 | 0.04850458 |
| **#776** | MAGT1 | 32.60867 | 0.3838885 | 0.005295894 |
| **#777** | USP53 | 16.91223 | -0.7936377 | 0.03022123 |
| **#778** | RGS14 | 33.01642 | -0.529432 | 0.03322287 |
| **#780** | CYTH1 | 169.2744 | -0.325055 | 0.02311626 |
| **#781** | FOXP1 | 365.9371 | -0.3514056 | 0.0472494 |
| **#783** | LTBP4 | 14.48689 | -0.6942803 | 0.01044277 |
| **#784** | CYCS | 113.7637 | 0.3170562 | 0.01671556 |
| **#785** | FARP2 | 8.580268 | 0.8911064 | 0.0359081 |
| **#786** | COPE | 150.5056 | 0.350944 | 0.03633592 |
| **#787** | VAMP1 | 13.53779 | -0.6744824 | 0.005399939 |
| **#788** | MRPL44 | 26.54468 | 0.4115489 | 0.00627007 |
| **#789** | KDM1B | 27.33255 | 0.4944167 | 0.03321752 |
| **#791** | PGD | 49.54974 | 0.4488477 | 0.04684526 |
| **#794** | NSUN6 | 24.19363 | -0.5153945 | 0.01323931 |
| **#796** | MCTP1 | 21.29958 | 0.5969306 | 0.0482926 |
| **#799** | CAMK2G | 27.90252 | -0.4250255 | 0.001143219 |
| **#800** | SYNGAP1 | 11.68468 | -0.8230889 | 0.01105433 |
| **#801** | LAPTM4A | 79.66831 | 0.3501086 | 0.02411219 |
| **#804** | ZNF439 | 18.71576 | -0.6014919 | 0.01482361 |
| **#807** | ARHGAP15 | 89.80807 | -0.4021055 | 0.04267109 |
| **#809** | TMEM204 | 17.12446 | -0.9231409 | 0.04707472 |
| **#810** | FRMD3 | 2.127181 | 1.927622 | 0.02814579 |
| **#811** | CTC1 | 18.21675 | -0.7036229 | 0.02767473 |
| **#812** | PPIB | 247.488 | 0.3393435 | 0.04278643 |

| | | | | |
|---|---|---|---|---|
| **#813** | WDR37 | 23.22821 | -0.4597836 | 0.00163643 |
| **#814** | LPCAT4 | 11.46103 | -0.7376107 | 0.003564117 |
| **#815** | PSD4 | 34.43688 | -0.4649981 | 0.02110485 |
| **#816** | TNRC6A | 61.79264 | -0.3588265 | 0.01185998 |
| **#817** | CD96 | 64.56722 | -0.4481415 | 0.04761657 |
| **#818** | ARID1B | 195.6526 | -0.3089518 | 0.0215122 |
| **#819** | LAT2 | 51.01922 | 0.3778813 | 0.02216188 |
| **#821** | TMC6 | 95.40905 | -0.3577749 | 0.02681852 |
| **#822** | NFATC3 | 42.34432 | -0.404291 | 0.01380511 |
| **#824** | DNMT3A | 26.21037 | -0.4942496 | 0.01440603 |
| **#825** | KARS | 52.60993 | 0.4204188 | 0.04092583 |
| **#826** | BAK1 | 14.98674 | 0.5222925 | 0.009741089 |
| **#830** | ITPRIPL2 | 11.24804 | 0.7343292 | 0.03448856 |
| **#831** | RPL36A | 2,131 | -0.2437498 | 0.009968839 |
| **#833** | CYFIP2 | 96.59313 | -0.3224498 | 0.01420354 |
| **#834** | TMX1 | 40.81706 | 0.3391996 | 0.002255578 |
| **#835** | ZNF827 | 14.6093 | -0.6370462 | 0.007743702 |
| **#836** | SNRPB | 148.4543 | 0.3460685 | 0.03963864 |
| **#837** | GSTK1 | 195.7036 | 0.333035 | 0.03858775 |
| **#840** | PRKAR2B | 9.261571 | 0.7878193 | 0.03046308 |
| **#841** | GTF3C6 | 57.05632 | 0.3416712 | 0.01369748 |
| **#842** | UCHL3 | 20.37433 | 0.4144715 | 0.000211654 |
| **#843** | TMEM208 | 38.72643 | 0.3965934 | 0.0203878 |
| **#844** | PKD1 | 15.84984 | -0.6537189 | 0.01522874 |
| **#845** | NANS | 32.13741 | 0.4314712 | 0.02566339 |
| **#847** | ATP5F1C | 121.3709 | 0.2868458 | 0.01040853 |
| **#848** | SKI | 23.27864 | -0.4863135 | 0.008786548 |
| **#850** | HERC4 | 54.68758 | -0.4608935 | 0.04826985 |
| **#851** | KCTD7 | 12.81875 | -0.6399291 | 0.003212797 |
| **#852** | ANP32A | 132.3621 | 0.2550421 | 0.00101942 |
| **#853** | GDI2 | 168.3875 | 0.3308795 | 0.0360207 |
| **#854** | PCNX2 | 14.19885 | -0.7829146 | 0.02317514 |

| | | | | |
|---|---|---|---|---|
| **#856** | LRRK1 | 14.33309 | 0.6090679 | 0.03005026 |
| **#857** | FLYWCH1 | 9.906434 | -0.7978343 | 0.003323308 |
| **#860** | PRH1 | 26.7508 | -0.4365628 | 0.004828058 |
| **#861** | NDUFV2 | 83.48224 | 0.2816669 | 0.002234083 |
| **#862** | APBB1 | 13.84893 | -0.7749684 | 0.02106788 |
| **#864** | CNP | 16.17353 | 0.4695975 | 0.004095428 |
| **#867** | KMT2B | 34.33034 | -0.5300557 | 0.0443667 |
| **#868** | P4HTM | 15.21089 | -0.6464414 | 0.01362728 |
| **#869** | N4BP2L1 | 47.90343 | 0.3495135 | 0.01237062 |
| **#871** | ABCA2 | 14.26256 | -0.8306767 | 0.02927687 |
| **#872** | MFSD1 | 42.05422 | 0.3818888 | 0.01979528 |
| **#874** | PCED1A | 17.05837 | -0.5588904 | 0.007605826 |
| **#875** | GLB1 | 20.28281 | 0.4361654 | 0.006654699 |
| **#878** | RINL | 23.49307 | -0.5612792 | 0.02689418 |
| **#879** | FBRSL1 | 24.89085 | -0.4966641 | 0.01659509 |
| **#880** | RFX1 | 20.92495 | -0.5001397 | 0.008180683 |
| **#881** | SLC31A1 | 13.32318 | 0.5918874 | 0.02267446 |
| **#883** | USP20 | 11.47702 | -0.6769895 | 0.002646324 |
| **#886** | TRIM65 | 11.04323 | -0.676646 | 0.001151942 |
| **#888** | ZNF335 | 21.66114 | -0.4577029 | 0.002400878 |
| **#889** | IL11RA | 9.350103 | -0.8617333 | 0.006344707 |
| **#890** | SGSM3 | 35.11398 | -0.3864658 | 0.005370635 |
| **#891** | CARD8 | 80.63019 | -0.3452809 | 0.02267028 |
| **#892** | ARRDC2 | 41.31176 | -0.4295072 | 0.0264014 |
| **#894** | DNAJB11 | 37.40924 | 0.3574523 | 0.008674855 |
| **#898** | IMPAD1 | 21.71465 | 0.412235 | 0.004612846 |
| **#899** | DENND1A | 17.62075 | 0.5707987 | 0.03919166 |
| **#900** | SPICE1 | 10.86683 | -0.8621642 | 0.01648242 |
| **#901** | MPC1 | 49.22919 | 0.3557405 | 0.01755248 |
| **#902** | AP2A2 | 32.63091 | -0.3844177 | 0.002409149 |
| **#903** | ULK3 | 16.51351 | -0.5901571 | 0.01252997 |
| **#904** | PTPN6 | 130.6249 | 0.3056185 | 0.02392301 |

| | | | | |
|---|---|---|---|---|
| **#905** | ATXN7L1 | 20.09421 | -0.4784593 | 0.003468387 |
| **#906** | EMC7 | 44.85878 | 0.3948566 | 0.03049617 |
| **#907** | TRAPPC6B | 19.63708 | -0.5117095 | 0.008858944 |
| **#909** | ATP6V0E2 | 31.40361 | -0.491396 | 0.03189491 |
| **#910** | YWHAQ | 96.34886 | 0.3344257 | 0.03187276 |
| **#914** | PCSK7 | 160.689 | -0.3313192 | 0.03683995 |
| **#916** | ABCC5 | 11.06233 | -0.6591937 | 0.000853245 |
| **#918** | SNRNP70 | 105.8106 | -0.3206911 | 0.02237553 |
| **#919** | GCSAM | 10.03152 | -0.9359101 | 0.01700345 |
| **#920** | RABL2B | 15.98809 | -0.5104802 | 0.0006277 |
| **#924** | UPF2 | 90.68529 | -0.3110655 | 0.01423025 |
| **#925** | CYB561D1 | 11.47406 | -0.6612546 | 0.002808844 |
| **#926** | MCM7 | 20.12583 | -0.4732959 | 0.003414459 |
| **#929** | SFMBT1 | 14.12675 | -0.6391758 | 0.01287303 |
| **#931** | SLC25A45 | 40.41831 | -0.3990226 | 0.01752259 |
| **#932** | CFDP1 | 79.1363 | -0.2902592 | 0.0032153 |
| **#934** | CHSY1 | 11.79866 | 0.5985208 | 0.021032 |
| **#935** | HABP4 | 11.78607 | -0.8455986 | 0.02285905 |
| **#938** | SLC35E2A | 10.93102 | -0.7172625 | 0.006367394 |
| **#939** | ENSA | 107.761 | 0.2713135 | 0.008214043 |
| **#941** | CEP164 | 27.85821 | -0.4050803 | 0.003428398 |
| **#946** | ST3GAL5 | 38.77635 | 0.3936477 | 0.02823883 |
| **#947** | MICU1 | 34.47548 | 0.3548732 | 0.00834456 |
| **#951** | TJP2 | 7.662818 | 0.6496021 | 0.006258378 |
| **#952** | LIMK2 | 6.050541 | 0.9578991 | 0.03458625 |
| **#954** | WDR41 | 14.60203 | 0.468471 | 0.005014071 |
| **#956** | MFSD14A | 23.13295 | 0.3904297 | 0.00454687 |
| **#957** | ACAP3 | 16.19254 | -0.5081721 | 0.002995235 |
| **#958** | PAN3 | 53.22315 | -0.325647 | 0.006486122 |
| **#959** | HLA-A | 1,912 | 0.260744 | 0.02916339 |
| **#961** | ZNF721 | 48.23721 | -0.4313049 | 0.04268325 |
| **#962** | PSMB4 | 84.89999 | 0.2866803 | 0.01105136 |

| | | | | |
|---|---|---|---|---|
| **#967** | PPP1R3F | 7.982404 | -0.9631371 | 0.007838029 |
| **#968** | GNAQ | 52.07037 | 0.3271094 | 0.01404096 |
| **#969** | TNIK | 41.81052 | -0.4472614 | 0.04195291 |
| **#972** | CSAD | 24.81406 | -0.4275747 | 0.005997165 |
| **#973** | SH2B1 | 18.91487 | -0.4537641 | 0.000972768 |
| **#975** | APLP2 | 217.4719 | 0.2834914 | 0.02819846 |
| **#976** | UGCG | 37.07789 | 0.4236202 | 0.042409 |
| **#977** | SUPT20H | 42.03158 | -0.3621308 | 0.01099987 |
| **#978** | KIF22 | 28.52498 | -0.4774759 | 0.02844975 |
| **#981** | DEF6 | 55.17962 | -0.3982856 | 0.03756621 |
| **#982** | NAGA | 24.37961 | 0.4824538 | 0.04363437 |
| **#983** | RPL34 | 5,846 | -0.2551554 | 0.03027494 |
| **#986** | SURF6 | 23.9842 | -0.4614981 | 0.01352622 |
| **#987** | PTGES3 | 213.9067 | 0.2435287 | 0.009391772 |
| **#989** | PRMT2 | 187.5479 | -0.3292205 | 0.04890543 |
| **#992** | EVL | 302.0225 | -0.3021593 | 0.04291201 |
| **#993** | SNX17 | 71.35777 | 0.3320089 | 0.03002107 |
| **#996** | GABPB2 | 31.90422 | -0.4251036 | 0.01940686 |
| **#998** | SNRPG | 82.82313 | 0.3197775 | 0.02884848 |
| **#1000** | TOM1L2 | 12.81981 | -0.5660655 | 0.00186152 |
| **#1002** | DESI1 | 23.94518 | 0.4690899 | 0.03824098 |
| **#1003** | GON4L | 61.37859 | -0.3349898 | 0.01676804 |
| **#1005** | DANCR | 48.43843 | -0.3940735 | 0.03133076 |
| **#1006** | ENY2 | 110.4013 | 0.2616217 | 0.008904745 |
| **#1008** | ZC3H12D | 13.6828 | -0.6689398 | 0.02090159 |
| **#1009** | RAB1A | 43.16642 | 0.3307956 | 0.0115115 |
| **#1010** | MTA1 | 21.97143 | -0.6152933 | 0.04866977 |
| **#1011** | PATL1 | 22.54007 | 0.42431 | 0.01692757 |
| **#1012** | PHC1 | 14.1436 | -0.5318448 | 0.002489108 |
| **#1013** | TECR | 79.10684 | -0.3559221 | 0.03851001 |
| **#1014** | PTOV1 | 30.56242 | -0.3925441 | 0.008627524 |
| **#1015** | GTF2B | 32.70686 | 0.3940655 | 0.0268515 |

| | | | | |
|---|---|---|---|---|
| **#1016** | STAT5A | 27.12588 | 0.3624488 | 0.005222105 |
| **#1017** | THOC3 | 31.29495 | -0.4838229 | 0.04195521 |
| **#1019** | HEBP1 | 21.82749 | 0.426939 | 0.01675791 |
| **#1020** | SNTB1 | 11.71787 | 0.5422097 | 0.01434376 |
| **#1024** | AKIRIN2 | 59.74225 | 0.3589066 | 0.03970865 |
| **#1025** | EZH1 | 36.5758 | -0.4718249 | 0.0492591 |
| **#1026** | ATP5PB | 116.4973 | 0.2913855 | 0.02533955 |
| **#1027** | TPCN1 | 18.27006 | -0.4741514 | 0.005709705 |
| **#1028** | PPP4C | 73.39825 | 0.3354816 | 0.03602005 |
| **#1031** | RBM6 | 89.49235 | -0.2808672 | 0.008564395 |
| **#1032** | LCP2 | 92.76881 | 0.3099747 | 0.0306154 |
| **#1033** | ACSL6 | 10.46792 | -0.8665772 | 0.02500947 |
| **#1034** | SMIM27 | 41.83769 | -0.3936755 | 0.02609702 |
| **#1036** | AAMP | 33.25793 | 0.3850796 | 0.02520234 |
| **#1037** | MRPL14 | 45.21209 | 0.3232101 | 0.01201414 |
| **#1038** | CDK10 | 21.01942 | -0.4400637 | 0.004703888 |
| **#1039** | ZMYND11 | 30.58477 | -0.4447959 | 0.02698752 |
| **#1040** | ASPHD2 | 5.442996 | 0.8089395 | 0.01276624 |
| **#1041** | SORBS3 | 14.07228 | -0.5485297 | 0.005744489 |
| **#1043** | PPM1M | 30.68108 | -0.4213957 | 0.01907801 |
| **#1045** | NME1 | 25.11214 | 0.4025703 | 0.01769725 |
| **#1046** | CNPY4 | 10.17199 | -0.7907359 | 0.01621471 |
| **#1048** | TTC13 | 13.44947 | -0.5866891 | 0.009661531 |
| **#1049** | WDR6 | 25.94034 | -0.4182972 | 0.009377617 |
| **#1052** | LSM 10.00 | 55.79509 | 0.3490791 | 0.0340249 |
| **#1053** | PPARGC1B | 9.198837 | 0.6204579 | 0.01929781 |
| **#1054** | HNRNPH1 | 328.5916 | -0.285632 | 0.04005131 |
| **#1055** | YWHAE | 99.63072 | 0.3199315 | 0.04012986 |
| **#1056** | CCT5 | 78.87773 | 0.267794 | 0.006944565 |
| **#1057** | MRPL30 | 19.87528 | -0.44962 | 0.005290361 |
| **#1058** | BICDL1 | 10.4426 | -0.7987515 | 0.02024467 |
| **#1059** | FNIP2 | 19.15882 | 0.4458748 | 0.01894918 |

| | | | | |
|---|---|---|---|---|
| **#1060** | AKTIP | 17.38963 | -0.6447635 | 0.03979538 |
| **#1061** | METTL17 | 28.1528 | -0.4273442 | 0.01835465 |
| **#1064** | CARMIL2 | 9.984496 | -0.9049953 | 0.0273817 |
| **#1066** | CNBP | 242.9646 | 0.2191295 | 0.005665644 |
| **#1067** | CALCOCO1 | 35.78139 | -0.3690137 | 0.01159723 |
| **#1069** | AKAP17A | 64.96243 | -0.3576232 | 0.03527624 |
| **#1071** | ADARB1 | 15.5534 | -0.584712 | 0.02018581 |
| **#1073** | ZC3H6 | 29.41972 | -0.4908687 | 0.04497849 |
| **#1074** | FBXL20 | 33.80083 | -0.3948709 | 0.01756377 |
| **#1075** | KLHDC2 | 40.36194 | -0.4164085 | 0.03758593 |
| **#1076** | POLG2 | 15.14337 | -0.5568364 | 0.01391676 |
| **#1081** | STK38 | 84.37477 | -0.3564678 | 0.04911722 |
| **#1083** | HSP90AB1 | 472.373 | 0.2425384 | 0.0239236 |
| **#1084** | SDHB | 51.92785 | 0.2978939 | 0.008530503 |
| **#1085** | GRPEL1 | 32.00965 | 0.4017317 | 0.03529298 |
| **#1086** | AMMECR1 | 15.48332 | -0.4774154 | 0.001242523 |
| **#1087** | VPS8 | 23.87166 | -0.4259212 | 0.009730284 |
| **#1093** | TAZ | 35.85078 | -0.3399436 | 0.003398867 |
| **#1094** | MED17 | 46.76562 | -0.354894 | 0.02120593 |
| **#1095** | DPYSL2 | 61.05454 | 0.3545019 | 0.04585625 |
| **#1098** | TMEM164 | 8.978708 | 0.5295924 | 0.001915213 |
| **#1099** | BLOC1S2 | 54.74105 | 0.2686935 | 0.000577 |
| **#1100** | EEPD1 | 7.617324 | -0.7823764 | 0.00181597 |
| **#1102** | NAA40 | 11.46962 | -0.651644 | 0.01271868 |
| **#1103** | ZBTB4 | 23.79406 | -0.3889231 | 0.001435927 |
| **#1104** | SIL1 | 18.05955 | 0.4370418 | 0.01550408 |
| **#1105** | FES | 27.43348 | 0.3975802 | 0.02536224 |
| **#1106** | NFKBIE | 9.812414 | 0.664505 | 0.03736569 |
| **#1107** | SIN3B | 22.86539 | -0.4932919 | 0.02822814 |
| **#1108** | ORMDL1 | 120.4736 | -0.2347537 | 0.001776403 |
| **#1109** | TSC2 | 29.2353 | -0.4010942 | 0.01449736 |
| **#1110** | PCYOX1 | 15.03882 | -0.473904 | 0.000390438 |

| | | | | |
|---|---|---|---|---|
| **#1111** | ALKBH6 | 9.498872 | -1.006762 | 0.03521023 |
| **#1113** | GAMT | 13.59373 | -0.4997328 | 0.000276076 |
| **#1116** | ELMOD3 | 17.03661 | -0.4493287 | 0.001506708 |
| **#1117** | MAPRE1 | 55.23944 | 0.3300182 | 0.02846339 |
| **#1121** | NAPEPLD | 7.663359 | -0.8540957 | 0.007375533 |
| **#1124** | KIAA2026 | 69.21358 | -0.3603865 | 0.04626626 |
| **#1127** | USP28 | 30.86716 | -0.4433788 | 0.03495484 |
| **#1128** | BCR | 20.19293 | -0.456856 | 0.01228882 |
| **#1130** | VMA21 | 49.43177 | 0.2816787 | 0.002987803 |
| **#1131** | PRPS2 | 13.75609 | 0.532144 | 0.03142238 |
| **#1132** | ZNF397 | 18.82976 | -0.5820708 | 0.03921191 |
| **#1133** | PCMT1 | 56.84788 | 0.3330637 | 0.03330217 |
| **#1135** | NDUFS6 | 115.2748 | 0.2914355 | 0.03444067 |
| **#1136** | NIPSNAP3A | 17.27499 | 0.4583749 | 0.02255811 |
| **#1137** | DZIP3 | 15.6856 | -0.5518603 | 0.0192358 |
| **#1139** | TANK | 53.31583 | 0.3512568 | 0.04079628 |
| **#1140** | RNF44 | 42.82392 | -0.3102698 | 0.002850567 |
| **#1142** | BCLAF3 | 14.11407 | -0.5489452 | 0.01159843 |
| **#1147** | MUS81 | 16.99121 | -0.5070702 | 0.01545454 |
| **#1148** | AGO2 | 40.05779 | -0.3251815 | 0.005493156 |
| **#1150** | OGFR | 55.11611 | 0.3429989 | 0.03883205 |
| **#1152** | PLA2G12A | 16.60393 | -0.5196471 | 0.01704904 |
| **#1156** | SLC7A6 | 24.37077 | -0.4950984 | 0.03875808 |
| **#1157** | HNRNPR | 160.8814 | 0.2357209 | 0.01259825 |
| **#1158** | ZNF236 | 18.19589 | -0.4204766 | 0.000233093 |
| **#1159** | TMED10 | 97.81995 | 0.2814605 | 0.02584979 |
| **#1160** | BNIP3 | 23.53486 | -0.5204357 | 0.04522488 |
| **#1161** | DENND11 | 21.37629 | -0.5001607 | 0.0304447 |
| **#1162** | TAX1BP1 | 165.5309 | 0.2781216 | 0.03695599 |
| **#1163** | HCG18 | 37.04972 | -0.3992644 | 0.03296392 |
| **#1166** | FRA10AC1 | 30.6447 | -0.420931 | 0.02846748 |
| **#1167** | ZNF609 | 31.92428 | -0.4350089 | 0.03788973 |

| | | | | |
|---|---|---|---|---|
| **#1168** | SMARCD3 | 11.53451 | 0.5959516 | 0.0400807 |
| **#1169** | MAP9 | 11.33884 | -0.584246 | 0.005670826 |
| **#1170** | SELENOT | 69.39501 | 0.3294448 | 0.04380815 |
| **#1173** | CCNY | 57.79136 | 0.313735 | 0.0267485 |
| **#1174** | GATD1 | 13.39906 | -0.5063803 | 0.003114434 |
| **#1175** | SMURF2 | 23.8728 | -0.4230534 | 0.01430605 |
| **#1176** | P3H1 | 12.15932 | -0.5454392 | 0.004043877 |
| **#1177** | LRRC14 | 9.99534 | -0.7918156 | 0.02456896 |
| **#1178** | LIG1 | 15.32528 | -0.5374064 | 0.01760731 |
| **#1180** | TNNI2 | 8.699123 | 0.5886228 | 0.01626875 |
| **#1182** | DUSP23 | 29.37478 | 0.4201561 | 0.0482446 |
| **#1185** | SPG7 | 73.105 | -0.3244538 | 0.03416162 |
| **#1187** | NUDCD3 | 33.79792 | -0.3727087 | 0.01694042 |
| **#1188** | HIP1R | 15.97899 | -0.556556 | 0.02563592 |
| **#1189** | FASTKD1 | 9.479567 | -0.5970475 | 0.000528452 |
| **#1191** | FUNDC2 | 46.88655 | 0.2901927 | 0.007780999 |
| **#1192** | CCDC14 | 26.50818 | -0.4187419 | 0.02037959 |
| **#1195** | WDR59 | 16.56275 | -0.5011168 | 0.01645319 |
| **#1198** | MAPKAPK5 | 30.88266 | -0.3967602 | 0.02270976 |
| **#1199** | SMYD4 | 15.56717 | -0.5087073 | 0.01430189 |
| **#1200** | PGRMC1 | 18.7252 | 0.3642736 | 0.000291311 |
| **#1202** | SRSF11 | 197.8531 | -0.2686403 | 0.03586834 |
| **#1204** | TANC2 | 13.58341 | -0.5308389 | 0.01069008 |
| **#1205** | KIF3A | 13.81169 | -0.4764764 | 0.001230359 |
| **#1206** | DPH5 | 22.31361 | -0.4789456 | 0.03199394 |
| **#1207** | ZNF699 | 15.5213 | -0.4485439 | 0.00120464 |
| **#1208** | LCLAT1 | 10.69082 | -0.6665245 | 0.01637735 |
| **#1209** | CROCC | 8.56794 | -0.7013288 | 0.006888616 |
| **#1210** | RPS28 | 4,997 | -0.2408571 | 0.04346335 |
| **#1211** | PSME3 | 21.47495 | 0.3513962 | 0.002083787 |
| **#1214** | ZNF66 | 10.71266 | -0.9010093 | 0.04887524 |
| **#1215** | RGL2 | 20.00159 | -0.4028274 | 0.003221536 |

| | | | | |
|---|---|---|---|---|
| **#1221** | ZKSCAN1 | 51.284 | -0.3006961 | 0.01093275 |
| **#1222** | MRPL16 | 29.17517 | 0.3191263 | 0.004177424 |
| **#1224** | SRP72 | 90.27089 | 0.2848618 | 0.03223464 |
| **#1226** | ARHGEF9 | 21.35311 | -0.4627675 | 0.02522843 |
| **#1227** | PRDX6 | 104.1199 | 0.2238447 | 0.00376364 |
| **#1228** | SH2D3A | 12.21521 | -0.7095657 | 0.03700029 |
| **#1231** | TMEM245 | 28.9865 | -0.4474088 | 0.04295026 |
| **#1232** | TPP1 | 76.4903 | 0.2925564 | 0.03201215 |
| **#1233** | SGSM2 | 13.12235 | -0.5254708 | 0.009026922 |
| **#1234** | CBR4 | 19.00196 | -0.5049231 | 0.0307805 |
| **#1235** | APH1A | 59.5144 | 0.3236496 | 0.04149075 |
| **#1236** | ABT1 | 31.54994 | 0.3667803 | 0.03119999 |
| **#1238** | CUL9 | 11.34682 | -0.5772153 | 0.009258517 |
| **#1239** | TMED7 | 34.21352 | 0.3874877 | 0.04894836 |
| **#1242** | JOSD2 | 28.44105 | 0.3888099 | 0.03690643 |
| **#1244** | DENND4C | 28.37923 | -0.432832 | 0.03742114 |
| **#1245** | RAB13 | 5.292069 | 0.7636074 | 0.01881969 |
| **#1248** | DIDO1 | 48.22187 | -0.3327772 | 0.02594937 |
| **#1251** | RPL31 | 1,238 | -0.2402346 | 0.04372117 |
| **#1252** | DGKE | 23.50433 | -0.4777696 | 0.04089665 |
| **#1254** | HIBCH | 18.63505 | -0.4820432 | 0.02491274 |
| **#1255** | ANKRD26 | 21.31958 | -0.4956406 | 0.03969683 |
| **#1256** | ZNF692 | 8.491423 | -0.6494691 | 0.003973179 |
| **#1257** | ST3GAL1 | 51.09415 | -0.3120377 | 0.01919637 |
| **#1259** | AP1G2 | 27.54188 | -0.3994163 | 0.02252521 |
| **#1260** | RAP1A | 114.7714 | 0.2208017 | 0.006773786 |
| **#1261** | AGPAT2 | 24.20272 | 0.4018292 | 0.03428906 |
| **#1262** | PRR12 | 25.29053 | -0.3799305 | 0.01070029 |
| **#1263** | RSU1 | 36.47669 | 0.3517506 | 0.03503032 |
| **#1264** | CERT1 | 60.80443 | 0.3105136 | 0.03685812 |
| **#1265** | SETD5 | 48.68394 | -0.3245398 | 0.02356354 |
| **#1266** | MR1 | 22.02607 | 0.354921 | 0.007539038 |

| | | | | |
|---|---|---|---|---|
| **#1267** | CLIP1 | 73.79014 | 0.3060596 | 0.04388173 |
| **#1271** | LIAS | 17.09391 | -0.4054342 | 0.000241987 |
| **#1272** | DOCK9 | 13.58576 | -0.6034285 | 0.03277606 |
| **#1273** | HOOK2 | 13.17074 | -0.5588946 | 0.01932287 |
| **#1274** | UBN2 | 24.97482 | -0.4685616 | 0.04608444 |
| **#1276** | COX7A2 | 228.7525 | 0.2583677 | 0.04495754 |
| **#1277** | CNNM3 | 19.08106 | -0.3855049 | 0.000523905 |
| **#1279** | KLF12 | 39.41952 | -0.337515 | 0.02022433 |
| **#1280** | SIDT1 | 16.75088 | -0.5223286 | 0.03175791 |
| **#1281** | ITPR3 | 10.80427 | -0.7002294 | 0.03027274 |
| **#1282** | APOLD1 | 9.883575 | -0.5620519 | 0.003169653 |
| **#1283** | MAPK1IP1L | 56.89029 | 0.2503735 | 0.003480885 |
| **#1284** | ZNF431 | 46.09759 | -0.3605885 | 0.0429463 |
| **#1287** | MAST4 | 12.55806 | -0.6729463 | 0.04181242 |
| **#1289** | BECN1 | 44.27723 | 0.260938 | 0.001043694 |
| **#1292** | ZNF618 | 4.550465 | 0.678175 | 0.000865691 |
| **#1293** | RGP1 | 14.06247 | -0.4733093 | 0.006900678 |
| **#1296** | NFIX | 4.793503 | 0.8729397 | 0.03540186 |
| **#1297** | TMEM140 | 4.42987 | 0.9733261 | 0.04553145 |
| **#1299** | NSMF | 17.16538 | -0.546109 | 0.04243923 |
| **#1300** | ASAP2 | 2.12325 | 1.259188 | 0.01429864 |
| **#1303** | DNAJC24 | 16.77063 | -0.4553214 | 0.01423981 |
| **#1304** | ST6GALNA( | 17.32249 | -0.5468127 | 0.04397785 |
| **#1309** | ZDHHC3 | 26.9908 | 0.3283221 | 0.009711832 |
| **#1312** | PHF14 | 60.5968 | -0.3205387 | 0.0378991 |
| **#1313** | PRKCQ | 27.30104 | -0.4114401 | 0.03250862 |
| **#1315** | SELENOS | 66.24611 | 0.3071874 | 0.04613274 |
| **#1317** | PPP2R3B | 7.984748 | -0.6681961 | 0.005980738 |
| **#1318** | WSB2 | 14.52342 | 0.4118742 | 0.01073279 |
| **#1319** | GLMN | 14.65971 | -0.4526716 | 0.005469232 |
| **#1322** | CACTIN | 8.425524 | -0.5898761 | 0.00026728 |
| **#1323** | YWHAB | 428.9648 | 0.2304813 | 0.04066769 |

| | | | | |
|---|---|---|---|---|
| **#1324** | ZNF605 | 11.70767 | -0.619413 | 0.02724754 |
| **#1327** | ZNF557 | 12.53652 | -0.5047595 | 0.008677531 |
| **#1330** | RMND1 | 13.9345 | -0.4670271 | 0.006491514 |
| **#1332** | RALGDS | 28.36531 | -0.4275262 | 0.04546134 |
| **#1333** | INO80E | 28.05825 | -0.3398235 | 0.006511443 |
| **#1337** | XRCC6 | 131.2504 | 0.200328 | 0.003618172 |
| **#1338** | DDX5 | 1,236 | -0.1998728 | 0.02621953 |
| **#1339** | STAU1 | 47.96086 | 0.2584634 | 0.004305544 |
| **#1340** | TRAPPC2 | 32.67121 | -0.3574074 | 0.02256847 |
| **#1342** | RNF11 | 26.56772 | 0.3350586 | 0.01458215 |
| **#1344** | ZNF76 | 13.64971 | -0.5123154 | 0.01704151 |
| **#1345** | RPL22 | 2,174 | -0.2182316 | 0.04239176 |
| **#1346** | ZNF248 | 9.414115 | -0.7421146 | 0.02920778 |
| **#1348** | TUBGCP6 | 16.90056 | -0.4626983 | 0.01950298 |
| **#1351** | GCN1 | 15.30477 | -0.4208463 | 0.001891306 |
| **#1353** | FBXW4 | 24.06487 | -0.3667956 | 0.009607102 |
| **#1354** | ZNF783 | 9.68287 | -0.5723985 | 0.007440791 |
| **#1356** | PGAP1 | 4.311086 | 0.9693015 | 0.04826381 |
| **#1357** | INF2 | 13.08145 | -0.5141161 | 0.01576287 |
| **#1358** | PRKAA1 | 42.7912 | -0.2939753 | 0.008923958 |
| **#1361** | C1D | 48.1616 | 0.2741899 | 0.01349115 |
| **#1363** | KNOP1 | 25.84394 | -0.3763975 | 0.01802818 |
| **#1364** | ITFG2 | 32.03778 | -0.3346403 | 0.01322966 |
| **#1365** | TMED2 | 81.92273 | 0.2432637 | 0.01593075 |
| **#1366** | IKBKB | 38.98355 | -0.3209012 | 0.01726279 |
| **#1369** | DUSP10 | 7.323778 | 0.5282568 | 0.006042257 |
| **#1371** | JTB | 160.9971 | 0.2480419 | 0.04097871 |
| **#1373** | ISYNA1 | 8.251675 | -0.6307285 | 0.006853986 |
| **#1376** | RTF2 | 75.25716 | 0.2577631 | 0.02284786 |
| **#1377** | ADAT2 | 11.52696 | -0.5595541 | 0.01724144 |
| **#1378** | MZF1 | 15.23954 | -0.4515002 | 0.0111246 |
| **#1381** | NRAS | 22.77052 | 0.3920307 | 0.03671916 |

| | | | | |
|---|---|---|---|---|
| **#1382** | MECP2 | 85.18181 | -0.2778508 | 0.03326492 |
| **#1383** | RBM7 | 33.65113 | 0.3022062 | 0.01323181 |
| **#1384** | BBS2 | 12.49656 | -0.6202676 | 0.03927505 |
| **#1387** | GPR82 | 8.927619 | -0.6471849 | 0.01557556 |
| **#1388** | CFAP97 | 63.96121 | -0.3187628 | 0.04804831 |
| **#1389** | ZSCAN30 | 8.938756 | -0.7422229 | 0.02883055 |
| **#1391** | RPL23A | 3,319 | -0.186123 | 0.02508301 |
| **#1394** | SDHD | 51.86849 | 0.3176549 | 0.04853638 |
| **#1395** | BRF1 | 12.47741 | -0.5107457 | 0.01429141 |
| **#1396** | OAF | 10.12636 | 0.5597758 | 0.04343896 |
| **#1397** | ARHGEF1 | 157.773 | -0.2532967 | 0.04153534 |
| **#1398** | PINK1 | 14.65064 | -0.4995292 | 0.02324223 |
| **#1399** | RILP | 7.326263 | 0.5903308 | 0.0220589 |
| **#1400** | RAB14 | 83.93692 | 0.2093623 | 0.001319267 |
| **#1409** | URGCP | 9.464584 | -0.555377 | 0.005557153 |
| **#1411** | ZNF281 | 33.51589 | 0.3142807 | 0.02220956 |
| **#1412** | NME3 | 80.65593 | -0.2948461 | 0.04621503 |
| **#1413** | RPS15A | 4,946 | -0.2088609 | 0.04416525 |
| **#1414** | SRSF8 | 55.53295 | -0.3208893 | 0.04415287 |
| **#1416** | KRIT1 | 26.2107 | -0.3602441 | 0.0158738 |
| **#1417** | GNB1 | 118.7602 | 0.2107033 | 0.0117813 |
| **#1418** | ADAMTSL4 | 9.685462 | 0.5472297 | 0.03568732 |
| **#1419** | CCDC152 | 9.948818 | -0.5273528 | 0.004666508 |
| **#1420** | PRIM2 | 12.60137 | -0.4512668 | 0.002541822 |
| **#1421** | EIF5B | 134.6449 | -0.2142703 | 0.01350824 |
| **#1423** | GAB2 | 9.35996 | 0.5852211 | 0.04690681 |
| **#1424** | RPL5 | 2,710 | -0.2024296 | 0.03878757 |
| **#1425** | ZBTB44 | 63.30315 | -0.2770415 | 0.0229779 |
| **#1427** | ABRAXAS1 | 28.79117 | -0.3509567 | 0.01769906 |
| **#1428** | RPL35 | 2,030 | -0.1719558 | 0.01702438 |
| **#1429** | CCDC130 | 30.5514 | -0.3097797 | 0.004657938 |
| **#1430** | ATXN10 | 40.71096 | 0.2778544 | 0.01256515 |

| | | | | |
|---|---|---|---|---|
| **#1431** | COX14 | 87.59378 | 0.2757786 | 0.04763544 |
| **#1432** | RPL35A | 3,349 | -0.1962649 | 0.03529498 |
| **#1433** | DDX51 | 17.60421 | -0.4463188 | 0.02367135 |
| **#1434** | PSMB7 | 62.75765 | 0.2669383 | 0.02605066 |
| **#1435** | ZNF593 | 17.58792 | 0.4165207 | 0.03438032 |
| **#1436** | TMEM41B | 34.84834 | -0.3121551 | 0.01237298 |
| **#1437** | MINK1 | 19.55259 | -0.3810771 | 0.008151012 |
| **#1439** | NHP2 | 71.68587 | 0.2217024 | 0.00478393 |
| **#1441** | MAD2L1BP | 6.892707 | 0.5208437 | 0.004094567 |
| **#1442** | PMF1 | 39.87806 | 0.304837 | 0.02863002 |
| **#1443** | TSGA10 | 14.03416 | -0.559306 | 0.04082979 |
| **#1444** | CCDC57 | 43.1259 | -0.297128 | 0.01589161 |
| **#1446** | EID1 | 205.6208 | 0.2324218 | 0.04196392 |
| **#1448** | EML3 | 22.73671 | -0.3906331 | 0.02136923 |
| **#1449** | EVI5 | 22.74579 | 0.3825936 | 0.0371764 |
| **#1452** | STN 1.00 | 27.96981 | 0.3183952 | 0.01570625 |
| **#1454** | BDH1 | 11.10824 | -0.5956081 | 0.02776774 |
| **#1455** | MIIP | 23.87976 | 0.390457 | 0.04698441 |
| **#1456** | PRMT7 | 14.60188 | -0.4486678 | 0.01132025 |
| **#1460** | AGER | 9.146289 | -0.5820448 | 0.01033928 |
| **#1461** | TM7SF2 | 8.437767 | -0.549562 | 0.000586151 |
| **#1464** | PRPF38B | 165.5267 | -0.2015859 | 0.01308175 |
| **#1465** | HEMK1 | 14.4447 | -0.4710326 | 0.0177544 |
| **#1467** | PHB | 58.24891 | 0.2768391 | 0.03026286 |
| **#1468** | TMEM268 | 10.49037 | 0.4769958 | 0.02131792 |
| **#1470** | RXRB | 16.50532 | -0.4040205 | 0.007091766 |
| **#1471** | NUP93 | 16.49736 | 0.3439713 | 0.001679814 |
| **#1472** | WDR73 | 25.35611 | -0.4089746 | 0.04060167 |
| **#1476** | MRPL47 | 34.06688 | 0.3002484 | 0.01820961 |
| **#1478** | NISCH | 24.84096 | -0.332797 | 0.006168849 |
| **#1479** | SNHG10 | 11.35076 | -0.5288788 | 0.0166656 |
| **#1480** | PSMG2 | 54.99964 | 0.2415078 | 0.008249812 |

| | | | | |
|---|---|---|---|---|
| **#1481** | CASP6 | 12.39162 | -0.5370863 | 0.02570768 |
| **#1482** | COPZ1 | 51.07789 | 0.3077012 | 0.04916693 |
| **#1483** | SNX8 | 7.150637 | 0.5029158 | 0.004107188 |
| **#1486** | PFDN2 | 61.70443 | 0.2455987 | 0.01520933 |
| **#1487** | ZNF585A | 22.60945 | -0.3940098 | 0.02509171 |
| **#1489** | ESYT1 | 37.78949 | -0.3257327 | 0.02768119 |
| **#1490** | PSMA1 | 115.8641 | 0.1819194 | 0.000602428 |
| **#1495** | TMEM129 | 13.32508 | -0.5367181 | 0.03409676 |
| **#1496** | RBM8A | 126.7973 | 0.2442553 | 0.04362113 |
| **#1498** | TMEM181 | 25.34584 | -0.3672086 | 0.02266346 |
| **#1499** | HADHB | 53.06652 | 0.2406513 | 0.007601657 |
| **#1502** | PNMA1 | 9.627945 | -0.6892916 | 0.03770305 |
| **#1503** | OSER1-DT | 9.887304 | -0.6764117 | 0.03771669 |
| **#1504** | RIC8B | 7.807738 | -0.6009636 | 0.005783739 |
| **#1505** | KCTD13 | 11.08851 | -0.475716 | 0.005199584 |
| **#1506** | UBE2Z | 41.12948 | 0.3114285 | 0.04130842 |
| **#1507** | HCLS1 | 220.8556 | 0.2058587 | 0.02840053 |
| **#1509** | TRA2A | 125.881 | -0.24405 | 0.03841472 |
| **#1510** | KBTBD3 | 11.7049 | -0.4921111 | 0.01292269 |
| **#1511** | TOMM22 | 71.95855 | 0.262201 | 0.03591914 |
| **#1512** | ZNF33B | 19.4098 | -0.4654074 | 0.04816625 |
| **#1513** | NOP56 | 70.70626 | -0.2408231 | 0.0132559 |
| **#1514** | ZNF514 | 8.719561 | -0.5492929 | 0.00466329 |
| **#1516** | GDF11 | 9.813707 | -0.6211818 | 0.02830156 |
| **#1521** | TCHP | 27.15584 | -0.3769153 | 0.03485996 |
| **#1522** | PPOX | 8.709256 | -0.5373475 | 0.003387334 |
| **#1524** | COMMD9 | 42.72146 | 0.3039691 | 0.04031614 |
| **#1527** | CDC40 | 78.72114 | 0.2230295 | 0.01335564 |
| **#1528** | PELP1 | 18.06849 | -0.3888454 | 0.01127029 |
| **#1529** | MOSMO | 11.26128 | -0.6136733 | 0.04082077 |
| **#1530** | LYPLA1 | 66.08796 | 0.2325236 | 0.01326703 |
| **#1533** | TJAP1 | 9.986759 | -0.4907627 | 0.003531086 |

| | | | | |
|---|---|---|---|---|
| **#1534** | CDK11B | 39.70088 | -0.2891303 | 0.01388612 |
| **#1536** | AUH | 13.59537 | -0.5027079 | 0.02955472 |
| **#1539** | ATP7A | 13.55628 | -0.4306574 | 0.007563356 |
| **#1540** | ADPRHL2 | 10.69249 | 0.4056088 | 0.003336047 |
| **#1545** | ECHDC2 | 22.41248 | -0.4089034 | 0.03899004 |
| **#1547** | ZSWIM9 | 8.15049 | -0.5704074 | 0.006388106 |
| **#1548** | M6PR | 114.1756 | 0.2254171 | 0.03164572 |
| **#1549** | RNPC3 | 42.60768 | -0.3106905 | 0.03349856 |
| **#1550** | TBC1D17 | 16.077 | -0.4005607 | 0.009339886 |
| **#1551** | PYURF | 78.44462 | 0.2352098 | 0.023385 |
| **#1552** | ZNF302 | 26.11571 | -0.3279125 | 0.01143064 |
| **#1553** | GPS2 | 104.7477 | -0.2412733 | 0.03382347 |
| **#1557** | SPNS3 | 16.53776 | -0.4047591 | 0.01344958 |
| **#1559** | TSC1 | 20.54265 | -0.400859 | 0.02918555 |
| **#1560** | SEC31B | 10.35251 | -0.4617721 | 0.001609744 |
| **#1564** | SLC27A1 | 10.46239 | -0.5915066 | 0.03309625 |
| **#1566** | MIGA2 | 9.369316 | -0.4813717 | 0.000784598 |
| **#1568** | LZTR1 | 9.392176 | -0.4886166 | 0.002087944 |
| **#1571** | ETFA | 38.28583 | 0.2781447 | 0.01998904 |
| **#1572** | KIF21A | 12.31769 | -0.512014 | 0.02815388 |
| **#1573** | PCSK5 | 15.7616 | -0.4685998 | 0.03563537 |
| **#1575** | LARP1 | 56.54172 | 0.2741084 | 0.03956481 |
| **#1577** | CASP2 | 21.54867 | -0.3381225 | 0.006814496 |
| **#1579** | RBM5 | 89.48346 | -0.2570296 | 0.042001 |
| **#1580** | SMAD2 | 62.73761 | -0.2717712 | 0.03391135 |
| **#1581** | MFSD8 | 19.80848 | -0.3990035 | 0.02759616 |
| **#1582** | ABCB8 | 14.59398 | -0.3827891 | 0.001363337 |
| **#1583** | TMEM14A | 8.504392 | -0.620088 | 0.0217952 |
| **#1587** | ABHD13 | 21.5484 | 0.2961618 | 0.001981669 |
| **#1589** | IFFO1 | 13.5656 | -0.5173618 | 0.04079774 |
| **#1595** | KIZ | 20.83113 | -0.4273873 | 0.0464739 |
| **#1597** | SLC26A11 | 9.542836 | -0.5666271 | 0.02151944 |

| | | | | |
|---|---|---|---|---|
| **#1598** | SLF1 | 19.65624 | -0.4417073 | 0.04854681 |
| **#1602** | TRMT10B | 21.5555 | -0.3899467 | 0.03220039 |
| **#1604** | HAGHL | 10.32083 | -0.6121691 | 0.04106175 |
| **#1606** | HNRNPC | 279.1013 | 0.1935171 | 0.03312195 |
| **#1607** | ZNF224 | 27.58823 | -0.3307366 | 0.02016326 |
| **#1608** | UNKL | 18.29257 | -0.3978498 | 0.02232359 |
| **#1609** | MMS22L | 14.30728 | -0.4209122 | 0.01301594 |
| **#1610** | SERPINB8 | 14.08165 | 0.383811 | 0.01648057 |
| **#1616** | POLRMT | 11.98081 | -0.5290419 | 0.03453732 |
| **#1620** | KIFC2 | 7.481814 | -0.5379478 | 0.001405591 |
| **#1621** | CLK2 | 12.5835 | -0.4588931 | 0.01671285 |
| **#1623** | PAAF1 | 11.58174 | -0.484223 | 0.0193801 |
| **#1624** | LAX1 | 11.85673 | -0.5297307 | 0.03462731 |
| **#1627** | HGSNAT | 33.02944 | -0.3374281 | 0.03971012 |
| **#1629** | EEFSEC | 19.92197 | -0.331267 | 0.00286023 |
| **#1630** | WDR54 | 22.29078 | -0.4113467 | 0.04941229 |
| **#1634** | PFKFB2 | 4.511283 | 0.6093827 | 0.007727658 |
| **#1637** | CAPN10 | 14.77265 | -0.4503518 | 0.02817335 |
| **#1640** | SLC43A3 | 10.21853 | 0.4254537 | 0.01344734 |
| **#1641** | GTPBP3 | 10.79785 | -0.4687016 | 0.009878555 |
| **#1643** | NDUFAB1 | 73.97763 | 0.2530768 | 0.04396375 |
| **#1644** | METAP2 | 98.40518 | -0.189674 | 0.00417857 |
| **#1646** | IFT80 | 12.38417 | -0.4868443 | 0.02722709 |
| **#1647** | EYA3 | 12.25424 | -0.4056328 | 0.001899669 |
| **#1648** | CNOT6 | 21.46737 | -0.3959361 | 0.03868333 |
| **#1649** | NCAPD3 | 14.72871 | -0.4204713 | 0.01685682 |
| **#1651** | WDR27 | 8.047801 | -0.6232783 | 0.02244756 |
| **#1653** | FXR2 | 8.710556 | -0.5439521 | 0.01272111 |
| **#1655** | NFRKB | 16.2897 | -0.3821865 | 0.01059957 |
| **#1657** | FLVCR2 | 4.386565 | 0.7893075 | 0.04760241 |
| **#1658** | KRI1 | 28.09286 | -0.3628773 | 0.04452324 |
| **#1659** | PCBP1 | 263.6807 | 0.1805115 | 0.02765426 |

| | | | | |
|---|---|---|---|---|
| **#1665** | PPP6C | 30.8501 | 0.2531135 | 0.001714049 |
| **#1666** | SPIN2B | 9.545946 | -0.4649339 | 0.002571167 |
| **#1667** | ZNF568 | 9.279398 | -0.5349224 | 0.01650982 |
| **#1670** | MCEE | 11.18104 | -0.4393752 | 0.005690526 |
| **#1676** | APPL2 | 14.90637 | -0.4821994 | 0.04762307 |
| **#1677** | HTATIP2 | 31.95705 | 0.2949231 | 0.02945599 |
| **#1678** | CYP2R1 | 8.93758 | -0.4773106 | 0.002710268 |
| **#1686** | ADPRH | 4.016102 | 0.7381797 | 0.0271626 |
| **#1688** | UBE2E1 | 33.47623 | 0.2459901 | 0.002690126 |
| **#1689** | WASHC2A | 27.00437 | 0.331062 | 0.04430113 |
| **#1691** | UBE3C | 28.18801 | -0.3425518 | 0.03593631 |
| **#1692** | CLSTN1 | 38.42028 | -0.2998393 | 0.03229425 |
| **#1695** | PRNP | 47.93022 | 0.2533817 | 0.02580308 |
| **#1696** | STK4 | 303.727 | -0.1675959 | 0.02056517 |
| **#1697** | SPHK1 | 2.738616 | 0.7437222 | 0.002695213 |
| **#1698** | RPS6KA4 | 27.40772 | 0.3185087 | 0.03611095 |
| **#1701** | ZNF90 | 8.100832 | -0.6155545 | 0.02557928 |
| **#1702** | LEMD2 | 24.35926 | -0.3016386 | 0.005681112 |
| **#1705** | CLDND1 | 57.437 | -0.2245099 | 0.008496527 |
| **#1707** | COG8 | 9.468294 | -0.4489452 | 0.000995257 |
| **#1708** | FBXO25 | 26.55648 | -0.2806019 | 0.00148946 |
| **#1712** | RPRD1A | 27.70516 | -0.3266809 | 0.02682504 |
| **#1713** | TFG | 29.04924 | 0.2727232 | 0.01060796 |
| **#1714** | DDHD2 | 16.68794 | -0.3971311 | 0.0223705 |
| **#1715** | TCF3 | 24.69787 | -0.3519032 | 0.03210864 |
| **#1716** | DFFB | 8.709347 | -0.4882618 | 0.004940593 |
| **#1718** | PYGO2 | 10.41125 | -0.5001892 | 0.02159375 |
| **#1722** | LAMTOR5 | 88.93746 | 0.2387303 | 0.04963912 |
| **#1727** | IGF1R | 17.27888 | -0.3537326 | 0.007887043 |
| **#1728** | NAE1 | 41.11253 | -0.2632929 | 0.01542264 |
| **#1730** | MORF4L1 | 213.8223 | 0.1981432 | 0.04403389 |
| **#1731** | ATP6V1E2 | 10.17786 | -0.4355701 | 0.002539434 |

| | | | | |
|---|---|---|---|---|
| **#1733** | TRAPPC12 | 26.1732 | -0.3324239 | 0.02753112 |
| **#1735** | CEP68 | 18.93991 | -0.414995 | 0.0460294 |
| **#1738** | ZNF544 | 13.42032 | -0.4909486 | 0.04468651 |
| **#1740** | ACRBP | 4.987493 | 0.686577 | 0.04331218 |
| **#1742** | SIRT5 | 9.431736 | -0.4727067 | 0.007355509 |
| **#1743** | ARF4 | 41.52947 | 0.2491261 | 0.01735309 |
| **#1747** | MRFAP1 | 74.08763 | 0.2269306 | 0.03222855 |
| **#1748** | HIGD1A | 24.98245 | 0.3248924 | 0.03850481 |
| **#1749** | OARD1 | 43.86087 | -0.2619285 | 0.02030695 |
| **#1752** | CA2 | 4.093576 | 0.6511131 | 0.01589691 |
| **#1753** | ATP6V0A2 | 25.14298 | -0.3634712 | 0.04506052 |
| **#1755** | TMEM185A | 10.74109 | -0.50019 | 0.02774089 |
| **#1756** | CCT8 | 87.00633 | 0.1813195 | 0.006054263 |
| **#1759** | VAT1 | 7.650724 | -0.4993419 | 0.00237766 |
| **#1765** | IL21R | 9.579559 | -0.5464902 | 0.03039287 |
| **#1766** | PIGL | 11.69351 | -0.4829321 | 0.03090214 |
| **#1768** | TBK1 | 23.59086 | 0.321291 | 0.03339029 |
| **#1771** | PGAM5 | 13.48553 | -0.4803437 | 0.04478245 |
| **#1772** | UQCRB | 639.0335 | -0.1583577 | 0.03117071 |
| **#1773** | THRAP3 | 134.8211 | 0.1642877 | 0.008696222 |
| **#1776** | AGO3 | 51.8793 | -0.2555296 | 0.02862642 |
| **#1780** | KLHL36 | 33.09738 | -0.2956104 | 0.02570532 |
| **#1781** | ACTR1B | 27.32743 | -0.3364043 | 0.03796443 |
| **#1783** | ZNF333 | 12.83393 | -0.3724403 | 0.00118729 |
| **#1795** | DNAJB2 | 17.12765 | -0.3393068 | 0.006312293 |
| **#1796** | GALM | 16.07767 | -0.3698324 | 0.01433147 |
| **#1797** | ARFGAP3 | 50.27381 | 0.2638838 | 0.04856083 |
| **#1799** | MRPL21 | 45.36023 | 0.2172128 | 0.00592804 |
| **#1809** | ZNF529 | 11.995 | -0.4154275 | 0.01131477 |
| **#1811** | TARS2 | 10.59328 | -0.4194825 | 0.003941066 |
| **#1812** | ZNF567 | 13.34664 | -0.4518102 | 0.03457582 |
| **#1814** | TIMM44 | 19.72691 | -0.3137929 | 0.004673816 |

| | | | | |
|---|---|---|---|---|
| **#1818** | MOGS | 24.93328 | -0.2989035 | 0.0116566 |
| **#1823** | HK1 | 49.23927 | 0.2519758 | 0.03688719 |
| **#1827** | ZNF185 | 4.90443 | 0.5597407 | 0.01313847 |
| **#1833** | RABEPK | 9.609243 | -0.4400869 | 0.004818701 |
| **#1835** | E4F1 | 13.23818 | -0.3762209 | 0.005871242 |
| **#1840** | SH2D3C | 21.65623 | -0.3489957 | 0.03000145 |
| **#1841** | PGM2L1 | 13.52252 | -0.435683 | 0.0311542 |
| **#1843** | PECR | 7.801604 | -0.5089743 | 0.008925261 |
| **#1846** | BUD13 | 9.185028 | -0.4590226 | 0.007835829 |
| **#1848** | ANXA7 | 69.55041 | 0.1809959 | 0.003215012 |
| **#1849** | ZNF445 | 17.62093 | -0.3517784 | 0.01539761 |
| **#1854** | ING5 | 26.36499 | -0.3351543 | 0.04104825 |
| **#1860** | SLC35E3 | 25.26017 | -0.3231232 | 0.03024422 |
| **#1864** | IFNAR2 | 53.92985 | 0.209007 | 0.01145132 |
| **#1865** | UBE3D | 7.856929 | -0.5651573 | 0.02319953 |
| **#1866** | IRF2BP1 | 10.36421 | -0.5374155 | 0.04692035 |
| **#1867** | PARL | 37.91815 | 0.2413298 | 0.01521222 |
| **#1869** | ST3GAL2 | 20.34728 | -0.3185059 | 0.01111411 |
| **#1871** | ACP2 | 5.835148 | 0.4995812 | 0.01283574 |
| **#1872** | PIEZO1 | 36.05571 | -0.2757022 | 0.0245043 |
| **#1877** | DNAJC7 | 106.131 | 0.1876501 | 0.02395746 |
| **#1882** | TAMM41 | 11.24893 | -0.3949283 | 0.003722968 |
| **#1885** | TSG101 | 23.98524 | 0.2631618 | 0.005879274 |
| **#1888** | GALNT11 | 29.39218 | -0.2527591 | 0.001941061 |
| **#1893** | MOB4 | 32.30168 | 0.2811915 | 0.03684097 |
| **#1894** | BBS9 | 8.541195 | -0.5555766 | 0.03051726 |
| **#1902** | IL17RC | 2.942691 | 0.6048889 | 0.000166449 |
| **#1906** | REXO4 | 32.92955 | -0.3002266 | 0.03850083 |
| **#1907** | TMA16 | 26.09519 | -0.331956 | 0.0420324 |
| **#1909** | MLYCD | 11.4525 | -0.4205345 | 0.01525195 |
| **#1913** | SDR42E2 | 9.589847 | -0.5566751 | 0.04818127 |
| **#1918** | DPH3 | 20.15012 | 0.3072114 | 0.02362398 |

| | | | | |
|---|---|---|---|---|
| **#1928** | COQ7 | 27.10218 | 0.2693999 | 0.01865935 |
| **#1932** | RUFY2 | 22.36307 | -0.3388867 | 0.03537422 |
| **#1933** | THOC7 | 68.64618 | 0.1975389 | 0.01840668 |
| **#1934** | PI4KA | 32.52615 | -0.2491519 | 0.007326363 |
| **#1936** | LTO1 | 12.31898 | -0.3675386 | 0.003505416 |
| **#1939** | CYB561D2. | 8.26561 | -0.6031356 | 0.04665934 |
| **#1940** | TTI1 | 9.148297 | -0.4602866 | 0.013436 |
| **#1941** | FEZ2 | 26.30045 | 0.2781323 | 0.02366648 |
| **#1949** | DIS3L | 19.59593 | -0.3237793 | 0.01644241 |
| **#1950** | ACSL5 | 17.13698 | 0.3431657 | 0.03767555 |
| **#1951** | XPNPEP1 | 16.49042 | 0.3376717 | 0.02979297 |
| **#1953** | ILF2 | 56.54568 | 0.2146804 | 0.02460582 |
| **#1955** | CALM3 | 132.0657 | 0.1909088 | 0.04414431 |
| **#1958** | SUSD6 | 32.34014 | 0.2810162 | 0.04402238 |
| **#1960** | TTC31 | 16.89199 | -0.3038191 | 0.000680139 |
| **#1963** | AK9 | 13.06317 | -0.4215972 | 0.03284907 |
| **#1965** | OSGIN2 | 13.3635 | 0.3866462 | 0.04358084 |
| **#1967** | ATAD3B | 10.25869 | -0.4435131 | 0.01818419 |
| **#1968** | ELOC | 57.22715 | 0.215575 | 0.02734999 |
| **#1969** | HDAC5 | 23.20962 | -0.302303 | 0.0182841 |
| **#1973** | SEC24D | 14.077 | 0.3190293 | 0.008616963 |
| **#1976** | SCARF1 | 2.912475 | 0.7931249 | 0.03203655 |
| **#1977** | MMS19 | 11.74911 | -0.3592655 | 0.000701647 |
| **#1982** | TRMU | 16.06391 | -0.3298966 | 0.008492208 |
| **#1983** | CC2D1B | 11.41912 | -0.39146 | 0.008813136 |
| **#1984** | PPP1R37 | 7.733026 | -0.5201501 | 0.02033083 |
| **#1990** | TRIQK | 9.288948 | 0.4208794 | 0.02881141 |
| **#1994** | ACAD11 | 7.501441 | -0.6008881 | 0.03858547 |
| **#1996** | ZNF383 | 9.460467 | -0.4653494 | 0.02088292 |
| **#1997** | SMARCC2 | 68.69348 | -0.1994453 | 0.0167303 |
| **#2000** | GPATCH1 | 11.43381 | -0.4111062 | 0.01671835 |
| **#2002** | TMEM94 | 10.36597 | -0.3825751 | 0.001822145 |

| | | | | |
|---|---|---|---|---|
| **#2006** | ITGA5 | 18.77542 | -0.343275 | 0.02755296 |
| **#2009** | MAP7D3 | 20.45896 | -0.2941095 | 0.007215554 |
| **#2014** | CAPS | 9.968353 | -0.3972081 | 0.004247952 |
| **#2015** | INTS2 | 9.696743 | -0.4853879 | 0.03253574 |
| **#2018** | GNPAT | 29.75689 | -0.2539298 | 0.009276718 |
| **#2026** | PCTP | 5.03932 | 0.5261356 | 0.01948146 |
| **#2034** | SDHA | 53.90441 | 0.2040129 | 0.01815703 |
| **#2036** | TOP3B | 10.43443 | -0.3681192 | 0.000261916 |
| **#2037** | ZNF701 | 13.83273 | -0.3609591 | 0.01463657 |
| **#2038** | NBEAL1 | 27.2951 | -0.306375 | 0.04064184 |
| **#2040** | SCCPDH | 10.73716 | 0.3588911 | 0.01320973 |
| **#2041** | DDX50 | 45.90247 | -0.2493094 | 0.03725966 |
| **#2046** | DCTN6 | 19.88536 | 0.2677187 | 0.006044177 |
| **#2048** | TRIM26 | 21.7171 | 0.2862138 | 0.02382615 |
| **#2049** | FECH | 7.740299 | 0.4097782 | 0.01252882 |
| **#2051** | TYSND1 | 12.86913 | -0.421841 | 0.03964751 |
| **#2052** | MICB | 9.116942 | 0.4261067 | 0.03586207 |
| **#2055** | TOMM6 | 177.9861 | 0.1657241 | 0.03866947 |
| **#2056** | PMS2 | 22.04642 | -0.2997412 | 0.01842999 |
| **#2058** | DCPS | 19.69644 | 0.3127857 | 0.03828175 |
| **#2060** | KLHL22 | 6.708087 | -0.533562 | 0.01769118 |
| **#2064** | NDUFAF7 | 13.37854 | -0.3650584 | 0.0151993 |
| **#2066** | TNK2 | 28.71533 | -0.2982538 | 0.04073776 |
| **#2070** | RAD54L2 | 16.89642 | -0.3245727 | 0.01435757 |
| **#2071** | SND1 | 45.07916 | 0.214612 | 0.02014971 |
| **#2074** | IRF5 | 17.47362 | 0.3109908 | 0.02630241 |
| **#2075** | CRTC2 | 22.66287 | -0.3063562 | 0.02728752 |
| **#2092** | FOXN3 | 123.4352 | 0.1496488 | 0.01398036 |
| **#2098** | NDE1 | 38.49981 | -0.2477575 | 0.02767384 |
| **#2099** | TMEM265 | 7.131192 | -0.5096357 | 0.01990469 |
| **#2109** | SLC35A4 | 18.38573 | 0.3180027 | 0.04029905 |
| **#2111** | VAMP7 | 18.27279 | 0.2785247 | 0.01001319 |

| | | | | |
|---|---|---|---|---|
| **#2112** | USE1 | 33.4853 | -0.2464418 | 0.01788674 |
| **#2115** | TMEM170A | 36.85353 | 0.215468 | 0.01167177 |
| **#2116** | RBM41 | 17.76009 | -0.3492156 | 0.03551227 |
| **#2119** | SPTSSA | 21.81302 | 0.3010601 | 0.04386254 |
| **#2120** | CEP89 | 9.22097 | -0.387946 | 0.002881774 |
| **#2123** | FCHO1 | 16.20793 | -0.2993374 | 0.002937467 |
| **#2127** | CPNE2 | 9.728134 | -0.4596344 | 0.03265247 |
| **#2129** | RMC1 | 12.27346 | 0.3196136 | 0.008783204 |
| **#2134** | SMPD4 | 19.13882 | -0.318734 | 0.02459483 |
| **#2137** | SLU7 | 58.24902 | 0.1935981 | 0.02071558 |
| **#2139** | NUP188 | 13.54827 | -0.3603218 | 0.0187333 |
| **#2140** | APBA3 | 9.057404 | -0.4326683 | 0.01635014 |
| **#2142** | NUTF2 | 43.14469 | 0.2136259 | 0.02032843 |
| **#2145** | TMEM33 | 35.13283 | 0.2571039 | 0.04661358 |
| **#2148** | ZNF395 | 14.90172 | -0.3139251 | 0.005129355 |
| **#2156** | SLC38A6 | 9.81059 | -0.4270454 | 0.02183054 |
| **#2158** | TRNAU1AP | 26.62616 | -0.2828369 | 0.03018897 |
| **#2159** | OTUD5 | 40.25163 | -0.2452162 | 0.03325111 |
| **#2162** | COMMD10 | 22.38084 | 0.2461826 | 0.006049848 |
| **#2167** | AASDH | 15.33511 | -0.3638918 | 0.03436138 |
| **#2168** | ANKRD46 | 8.098147 | -0.4602357 | 0.01859662 |
| **#2169** | PTGS1 | 7.304146 | 0.4259363 | 0.0218236 |
| **#2173** | TIMM17A | 28.53309 | 0.2725168 | 0.04497356 |
| **#2175** | ANKMY1 | 7.121824 | -0.4317277 | 0.00347406 |
| **#2180** | CA11 | 9.975686 | -0.4449145 | 0.03197462 |
| **#2181** | DDX55 | 21.83645 | -0.2928253 | 0.02146155 |
| **#2182** | CRYZL1 | 24.29341 | -0.2648719 | 0.01213002 |
| **#2185** | ZNF286A | 9.772473 | -0.3774569 | 0.005279065 |
| **#2188** | COQ6 | 11.87633 | -0.339539 | 0.003429686 |
| **#2193** | CKS1B | 15.72283 | 0.3361331 | 0.04559569 |
| **#2194** | TIAL1 | 87.03219 | -0.1942765 | 0.04186783 |
| **#2196** | PSMC1 | 62.04279 | 0.2005302 | 0.03566481 |

| | | | | |
|---|---|---|---|---|
| **#2197** | CTRL | 4.116763 | 0.6053887 | 0.0392903 |
| **#2198** | TUBE1 | 9.097957 | -0.4640011 | 0.03275181 |
| **#2199** | PSMC2 | 25.84013 | 0.226826 | 0.004306472 |
| **#2200** | RANBP10 | 8.97026 | -0.3956673 | 0.006526104 |
| **#2206** | TAF1C | 9.542358 | -0.4478595 | 0.03118222 |
| **#2209** | ZNF324 | 9.201836 | -0.3963043 | 0.00894679 |
| **#2215** | SLC2A11 | 10.30729 | -0.4170815 | 0.02613904 |
| **#2219** | CRAMP1 | 9.634583 | -0.4128523 | 0.01901306 |
| **#2224** | NDUFB5 | 67.99613 | 0.1915095 | 0.03404896 |
| **#2226** | RPUSD4 | 14.10635 | -0.3118869 | 0.004101528 |
| **#2230** | ZNF337 | 8.588906 | -0.4581999 | 0.02728385 |
| **#2231** | STX12 | 34.14947 | 0.2437325 | 0.03727495 |
| **#2233** | ZNF738 | 10.74024 | -0.3699421 | 0.01124502 |
| **#2234** | RETREG3 | 33.21326 | -0.2472441 | 0.02599416 |
| **#2248** | BAHD1 | 6.47526 | -0.4307591 | 0.002513543 |
| **#2255** | NPRL3 | 19.98578 | -0.2774232 | 0.009999795 |
| **#2266** | HPRT1 | 26.3737 | 0.2262064 | 0.007871033 |
| **#2267** | NMNAT1 | 7.390881 | -0.4225857 | 0.006613619 |
| **#2268** | BCL2L2 | 4.815994 | 0.4441096 | 0.006480797 |
| **#2271** | SLC26A2 | 10.05155 | -0.4357397 | 0.03535717 |
| **#2272** | ELP1 | 10.50452 | -0.4436751 | 0.04478068 |
| **#2286** | CSE1L | 13.80024 | -0.3373468 | 0.01684046 |
| **#2289** | NCEH1 | 7.008233 | 0.4027366 | 0.0154421 |
| **#2292** | AMPD3 | 6.758054 | -0.5131894 | 0.02835195 |
| **#2294** | HAUS5 | 7.575224 | -0.4705074 | 0.02453678 |
| **#2298** | TICAM1 | 6.291295 | 0.4543514 | 0.02999653 |
| **#2302** | RFFL | 24.09591 | -0.2950827 | 0.04114554 |
| **#2303** | TTF1 | 32.76145 | -0.2194999 | 0.008652679 |
| **#2304** | RBMXL1 | 15.60111 | 0.3031302 | 0.02645357 |
| **#2309** | HSD17B7 | 11.09085 | -0.4071255 | 0.03501683 |
| **#2310** | ENGASE | 13.38049 | -0.392508 | 0.04992983 |
| **#2313** | COMMD1 | 30.4402 | 0.2351121 | 0.02630179 |

| | | | | |
|---|---|---|---|---|
| #2320 | DDX19A | 23.15466 | -0.2612564 | 0.01367381 |
| #2321 | LARP1B | 17.84867 | -0.2900869 | 0.01327153 |
| #2322 | DROSHA | 9.545655 | -0.4040594 | 0.02080782 |
| #2327 | TXNDC12 | 46.92526 | 0.2006958 | 0.02672403 |
| #2329 | VPS18 | 7.834945 | 0.4310922 | 0.0440035 |
| #2331 | FBXO7 | 65.43546 | 0.1914716 | 0.03949221 |
| #2336 | PARVG | 88.40123 | 0.1646843 | 0.02851174 |
| #2338 | ZNF621 | 13.46324 | -0.3251459 | 0.01198188 |
| #2351 | PLAGL1 | 11.8118 | 0.3217692 | 0.01799438 |
| #2358 | CAMKMT | 9.736317 | -0.4424104 | 0.04228625 |
| #2364 | TRMT6 | 20.39569 | 0.2387819 | 0.00524427 |
| #2375 | FUT4 | 6.037582 | 0.4457302 | 0.02846542 |
| #2380 | AMDHD2 | 12.17689 | -0.33606 | 0.01291103 |
| #2381 | TIGAR | 14.47562 | 0.3162953 | 0.03612709 |
| #2382 | LAMTOR3 | 20.6976 | 0.2656911 | 0.02801951 |
| #2385 | SNRNP48 | 21.44685 | -0.2786896 | 0.02408216 |
| #2388 | ZDHHC8 | 9.392004 | -0.4281113 | 0.03558637 |
| #2390 | TAOK2 | 10.69698 | -0.3851365 | 0.026903 |
| #2391 | ARL 3.00 | 14.57291 | -0.3291949 | 0.02502239 |
| #2395 | APOBEC3D | 11.96253 | -0.3904729 | 0.04254645 |
| #2402 | TAF6 | 16.47907 | -0.2872183 | 0.01095371 |
| #2409 | PIP4K2B | 15.09199 | -0.315124 | 0.02109405 |
| #2414 | COG1 | 14.77281 | -0.3029947 | 0.01234586 |
| #2417 | COG4 | 22.55022 | -0.2449303 | 0.00831306 |
| #2419 | NCK1-DT | 8.493702 | -0.4607853 | 0.04288371 |
| #2422 | ELP3 | 15.34198 | -0.3120228 | 0.02185138 |
| #2435 | EDEM2 | 19.62202 | 0.2504431 | 0.01483684 |
| #2442 | ZNF641 | 17.17557 | -0.2972399 | 0.0231234 |
| #2443 | NPLOC4 | 31.3926 | -0.2131982 | 0.009242027 |
| #2445 | SMU1 | 54.73812 | 0.1879767 | 0.03310992 |
| #2454 | MTFMT | 11.01419 | -0.349768 | 0.01635019 |
| #2455 | POLG | 30.11424 | -0.2591186 | 0.04733543 |

| | | | | |
|---|---|---|---|---|
| **#2456** | ST7L | 9.393839 | -0.4122063 | 0.03314192 |
| **#2464** | KLHL42 | 12.57944 | -0.3505062 | 0.02885839 |
| **#2466** | TUBGCP4 | 11.73583 | -0.3717942 | 0.03506258 |
| **#2467** | METAP1 | 18.40826 | -0.3019747 | 0.03518093 |
| **#2468** | ZNF346 | 10.97213 | -0.3986922 | 0.04367771 |
| **#2472** | MMD | 12.84117 | 0.2939537 | 0.01498605 |
| **#2475** | SLC35B3 | 9.061541 | -0.4174723 | 0.03370328 |
| **#2479** | PACS2 | 15.5661 | -0.2965365 | 0.01633223 |
| **#2481** | COQ8A | 16.33485 | -0.2997704 | 0.02296368 |
| **#2482** | TIMM21 | 7.935609 | -0.4587029 | 0.03988968 |
| **#2495** | RAB18 | 49.08548 | 0.1685532 | 0.0106047 |
| **#2499** | RBFA | 19.02044 | -0.2635883 | 0.01333705 |
| **#2500** | TAB1 | 8.584451 | -0.4220087 | 0.03257612 |
| **#2503** | NDST2 | 17.37489 | 0.287447 | 0.03926636 |
| **#2506** | ANAPC7 | 20.57771 | -0.2768089 | 0.0281737 |
| **#2510** | PIGQ | 7.20072 | -0.3849134 | 0.007317767 |
| **#2511** | CLYBL | 7.506916 | -0.4535005 | 0.03456906 |
| **#2513** | STRIP1 | 16.18349 | -0.283261 | 0.01312852 |
| **#2515** | PRELID3B | 31.16184 | 0.2103349 | 0.02132852 |
| **#2519** | BSDC1 | 38.71134 | -0.2084639 | 0.02403736 |
| **#2521** | ANKRD54 | 9.022264 | -0.3309113 | 0.001234652 |
| **#2524** | ZNF316 | 16.52857 | -0.3211755 | 0.04490488 |
| **#2529** | TIMM23 | 10.01027 | 0.2937956 | 0.002123075 |
| **#2535** | VILL | 8.199793 | -0.403688 | 0.02306093 |
| **#2544** | ADCK5 | 6.722964 | -0.4022216 | 0.01125624 |
| **#2547** | SMIM15 | 22.59785 | 0.2560771 | 0.04034395 |
| **#2557** | RENBP | 8.680298 | 0.3695839 | 0.03665304 |
| **#2558** | PPCS | 67.26222 | 0.1677397 | 0.0338729 |
| **#2565** | DXO | 9.62681 | -0.3867464 | 0.03166781 |
| **#2567** | RAI1 | 9.035531 | -0.3958617 | 0.02987573 |
| **#2569** | MUL1 | 7.690515 | 0.3871983 | 0.03642957 |
| **#2572** | PSMC6 | 66.07109 | -0.1870658 | 0.04849688 |

| | | | | |
|---|---|---|---|---|
| #2575 | RPAIN | 46.48075 | -0.1586063 | 0.000567519 |
| #2584 | EEF2K | 7.578705 | -0.4317949 | 0.03210464 |
| #2586 | AP1M1 | 41.75965 | -0.1927168 | 0.01932607 |
| #2587 | NUP85 | 10.90084 | -0.3054606 | 0.003322046 |
| #2591 | EXOC7 | 40.1084 | -0.187967 | 0.01374797 |
| #2592 | ZMIZ2 | 19.2574 | -0.2537017 | 0.01234064 |
| #2599 | MPHOSPH6 | 24.01893 | -0.2675604 | 0.04327042 |
| #2602 | ZNF821 | 10.72565 | -0.3371158 | 0.01646307 |
| #2605 | UBIAD1 | 14.27289 | -0.2948913 | 0.01488304 |
| #2609 | GBA2 | 12.78187 | -0.2911405 | 0.005848874 |
| #2612 | ABCD4 | 20.87569 | -0.2652079 | 0.02827234 |
| #2615 | IL18BP | 9.137481 | 0.3317667 | 0.01989434 |
| #2618 | SRD5A1 | 7.747314 | 0.3603915 | 0.02357147 |
| #2623 | ZNF426 | 9.653536 | -0.3364317 | 0.01057306 |
| #2628 | PPARA | 10.55382 | -0.3463908 | 0.0218825 |
| #2629 | TTPAL | 11.27046 | -0.3441794 | 0.02670477 |
| #2640 | UBE2K | 63.4093 | 0.1476787 | 0.01316686 |
| #2645 | PPME1 | 9.243668 | -0.3217721 | 0.003413169 |
| #2649 | ABRAXAS2 | 12.38458 | 0.2754911 | 0.009073255 |
| #2650 | RFC4 | 8.876039 | -0.3665613 | 0.01886143 |
| #2651 | ZNF480 | 26.34532 | -0.2435472 | 0.03271522 |
| #2653 | LONP1 | 15.88265 | -0.2669075 | 0.008897227 |
| #2655 | SIK2 | 8.279157 | -0.4135723 | 0.0361305 |
| #2656 | LETMD1 | 30.36536 | -0.2318963 | 0.03573319 |
| #2664 | UBE4B | 20.6691 | -0.261582 | 0.02731664 |
| #2665 | UVSSA | 13.67191 | -0.3409864 | 0.04834697 |
| #2670 | FLCN | 9.813796 | -0.3403438 | 0.01548656 |
| #2673 | MFSD5 | 8.622841 | 0.3727456 | 0.0466596 |
| #2676 | AP3M2 | 25.57586 | -0.2608979 | 0.04953597 |
| #2684 | MOCS2 | 20.51949 | -0.2745975 | 0.03872293 |
| #2690 | TNPO2 | 8.559569 | -0.3839815 | 0.02711896 |
| #2692 | ABAT | 9.924165 | -0.3426545 | 0.0184963 |

| | | | | |
|---|---|---|---|---|
| **#2696** | ARFGAP1 | 13.05624 | -0.2928842 | 0.01229133 |
| **#2697** | SPATA33 | 8.71249 | -0.4203917 | 0.04893923 |
| **#2701** | ARSB | 7.782306 | 0.381117 | 0.04322289 |
| **#2702** | COPS7B | 16.57756 | -0.2665523 | 0.014159 |
| **#2703** | TRIM39 | 8.237928 | -0.3534865 | 0.01167051 |
| **#2704** | ZNF789 | 9.774791 | -0.353367 | 0.02339149 |
| **#2705** | DGCR8 | 12.25812 | -0.3306484 | 0.03029767 |
| **#2707** | PIDD1 | 8.344147 | -0.3374944 | 0.006945207 |
| **#2710** | TBC1D19 | 9.105336 | -0.3580891 | 0.02033647 |
| **#2719** | OXCT1 | 19.19588 | -0.2589525 | 0.02262368 |
| **#2720** | MBD1 | 19.69867 | -0.2524107 | 0.0196088 |
| **#2724** | TEFM | 11.48338 | -0.3540158 | 0.04061724 |
| **#2726** | ZMAT2 | 72.19873 | 0.1468647 | 0.02466935 |
| **#2729** | SLC39A11 | 8.241805 | 0.3398439 | 0.02327984 |
| **#2731** | ABCC10 | 8.550362 | -0.3855004 | 0.03030201 |
| **#2732** | SNHG3 | 67.47291 | -0.1624581 | 0.03172933 |
| **#2740** | ELMOD2 | 13.57106 | -0.3049413 | 0.025757 |
| **#2741** | HCCS | 10.77065 | 0.2793626 | 0.007122412 |
| **#2746** | OSBPL1A | 6.515534 | 0.4009378 | 0.04134483 |
| **#2747** | NCSTN | 24.98569 | 0.2392124 | 0.04878599 |
| **#2748** | ATP5MC1 | 75.01086 | 0.1443637 | 0.02629388 |
| **#2750** | EFHC1 | 8.834173 | -0.3468671 | 0.01581082 |
| **#2753** | UBAP2 | 23.30452 | -0.2568153 | 0.04118021 |
| **#2754** | METTL3 | 14.02624 | -0.3072903 | 0.03138774 |
| **#2760** | SEC22C | 22.64655 | -0.2408587 | 0.02502267 |
| **#2767** | ACLY | 26.046 | 0.1820979 | 0.000818906 |
| **#2769** | BTBD2 | 17.17578 | -0.2629239 | 0.01952672 |
| **#2779** | TAF9 | 43.34784 | 0.154431 | 0.007004257 |
| **#2783** | RPL26L1 | 24.4345 | 0.2149315 | 0.02305459 |
| **#2784** | NSMCE2 | 19.62037 | -0.2449709 | 0.01844412 |
| **#2789** | NELFA | 16.94138 | -0.2539995 | 0.01305438 |
| **#2815** | PSMD1 | 38.20844 | 0.1655511 | 0.0107165 |

| | | | | |
|---|---|---|---|---|
| **#2818** | KPTN | 3.431795 | 0.5055552 | 0.04234595 |
| **#2830** | MBD3 | 21.1947 | -0.2642109 | 0.04837045 |
| **#2831** | SCAP | 14.91734 | -0.3052281 | 0.04518777 |
| **#2834** | ZFAND1 | 31.99609 | -0.2232742 | 0.04950417 |
| **#2855** | TBL1X | 36.14366 | -0.2042367 | 0.03995076 |
| **#2858** | PRIMPOL | 8.443616 | -0.3943586 | 0.04632327 |
| **#2866** | SLC25A12 | 10.42721 | -0.3104493 | 0.01645421 |
| **#2871** | MRTFA | 22.16697 | -0.2004632 | 0.001058563 |
| **#2872** | PHF21A | 40.34712 | -0.1837889 | 0.02841338 |
| **#2875** | LRRC23 | 9.478105 | -0.3478786 | 0.03076769 |
| **#2876** | MAIP1 | 8.101146 | 0.3374504 | 0.03179295 |
| **#2887** | SHC1 | 16.47617 | 0.260024 | 0.04036366 |
| **#2889** | USP19 | 11.34008 | -0.2834184 | 0.008392547 |
| **#2894** | CCDC28A | 23.75387 | -0.2114735 | 0.01425074 |
| **#2895** | SRPK1 | 28.49186 | -0.2230314 | 0.04078458 |
| **#2919** | MAP11 | 8.632483 | -0.3802115 | 0.04593795 |
| **#2930** | IMPDH2 | 40.74084 | -0.18003 | 0.02969845 |
| **#2943** | GPN2 | 11.37252 | -0.2960303 | 0.01932943 |
| **#2944** | CISD2 | 24.48937 | 0.224815 | 0.04900893 |
| **#2948** | WDTC1 | 21.52884 | -0.2176529 | 0.01534864 |
| **#2955** | TBP | 12.93774 | -0.2695501 | 0.01375939 |
| **#2956** | POLD1 | 6.564376 | -0.4033929 | 0.03796964 |
| **#2959** | PSMD11 | 39.14291 | 0.1598548 | 0.01555133 |
| **#2960** | BTF3L4 | 47.28717 | 0.1689513 | 0.0409481 |
| **#2970** | BFAR | 36.95199 | 0.1892741 | 0.04729574 |
| **#2973** | QRICH1 | 28.48283 | -0.2186596 | 0.04335261 |
| **#2977** | YIPF1 | 10.30517 | 0.2886134 | 0.02498456 |
| **#3002** | MROH1 | 13.9499 | -0.2865223 | 0.03513405 |
| **#3003** | VAPB | 17.30823 | -0.2705449 | 0.04673204 |
| **#3005** | CTNNBL1 | 48.61515 | 0.151373 | 0.0237001 |
| **#3009** | GET1 | 8.398527 | -0.3179956 | 0.01615061 |
| **#3012** | COG7 | 8.871181 | -0.2782432 | 0.001249846 |

| | | | | |
|---|---|---|---|---|
| **#3017** | CENPW | 3.887023 | 0.4458104 | 0.0435966 |
| **#3018** | DERA | 18.65445 | 0.2034235 | 0.007208686 |
| **#3025** | SCRIB | 9.006028 | -0.3323381 | 0.02961828 |
| **#3034** | TM9SF4 | 21.65237 | 0.1908969 | 0.008486061 |
| **#3040** | OGFOD2 | 6.755202 | -0.3319567 | 0.01174311 |
| **#3042** | CAD | 6.684687 | -0.3415932 | 0.01561682 |
| **#3052** | ZFYVE19 | 9.49929 | -0.3368795 | 0.03980408 |
| **#3061** | GLRX3 | 34.75817 | 0.1848339 | 0.04494331 |
| **#3066** | SMARCAL1 | 9.582504 | -0.3181319 | 0.02931918 |
| **#3073** | VPS13B-DT | 7.098399 | -0.3611527 | 0.03196433 |
| **#3099** | ACTR5 | 7.039322 | -0.3800658 | 0.04413388 |
| **#3102** | RDH11 | 32.06744 | -0.1977353 | 0.04220979 |
| **#3103** | ANKRD52 | 8.56829 | -0.3073182 | 0.01816881 |
| **#3109** | BICRA | 8.352854 | -0.3151475 | 0.0211107 |
| **#3118** | FBXW2 | 30.18339 | -0.170649 | 0.009360028 |
| **#3128** | NCAPH2 | 13.61667 | -0.2675757 | 0.02832214 |
| **#3135** | SASS6 | 8.957559 | -0.3225206 | 0.03269229 |
| **#3141** | EIF2B4 | 17.44998 | -0.2095105 | 0.00511927 |
| **#3143** | DHX30 | 33.30775 | -0.1921357 | 0.04224416 |
| **#3145** | ACAD10 | 8.624762 | -0.3483619 | 0.0471075 |
| **#3151** | MRPL13 | 23.18079 | 0.2113568 | 0.04471404 |
| **#3167** | MAP4K1 | 22.64877 | -0.1938518 | 0.0113555 |
| **#3176** | SLC25A51 | 6.751196 | -0.3501876 | 0.0291395 |
| **#3198** | RBBP5 | 15.36842 | -0.2331059 | 0.0148762 |
| **#3199** | PAXIP1 | 7.478086 | -0.3387545 | 0.03206139 |
| **#3200** | IMPACT | 14.91201 | 0.2341608 | 0.02690565 |
| **#3206** | MTHFSD | 6.779504 | -0.3462508 | 0.02947338 |
| **#3211** | SLC7A6OS | 27.63523 | -0.2011418 | 0.03929564 |
| **#3214** | CAMTA2 | 14.0232 | -0.2422781 | 0.01554043 |
| **#3223** | RTL8C | 13.01547 | 0.2655721 | 0.04859232 |
| **#3228** | PXMP4 | 7.51885 | -0.2722411 | 0.001019117 |
| **#3233** | TRIT1 | 8.418116 | -0.3217894 | 0.03375693 |

| | | | | |
|---|---|---|---|---|
| **#3235** | ZSWIM8 | 23.39959 | -0.1955235 | 0.01846802 |
| **#3267** | NRF1 | 11.93515 | -0.2668509 | 0.02557343 |
| **#3269** | MMADHC | 39.56942 | 0.1662373 | 0.04831642 |
| **#3271** | PHRF1 | 18.06915 | -0.2460321 | 0.04926465 |
| **#3272** | IMP4 | 29.0623 | 0.1874338 | 0.04660215 |
| **#3275** | KIAA0513 | 15.22154 | 0.2270806 | 0.02773873 |
| **#3280** | CCDC58 | 9.701399 | -0.3009299 | 0.03456514 |
| **#3282** | GPAT4 | 20.30502 | -0.2196478 | 0.0344619 |
| **#3286** | TBC1D20 | 23.85464 | -0.1910072 | 0.01914281 |
| **#3297** | BBS7 | 6.647818 | -0.3181962 | 0.01924345 |
| **#3299** | MRS2 | 15.24988 | -0.2466638 | 0.03325381 |
| **#3310** | ATL3 | 32.93987 | 0.1771457 | 0.04902387 |
| **#3315** | RRP1 | 11.66794 | -0.2771809 | 0.03456017 |
| **#3317** | EDC4 | 13.36815 | -0.2471888 | 0.0226573 |
| **#3321** | MYO19 | 13.57564 | -0.2733071 | 0.04829916 |
| **#3332** | DPM1 | 21.54409 | 0.1729466 | 0.006929711 |
| **#3348** | POLL | 12.44576 | -0.2682339 | 0.03693962 |
| **#3349** | ABCD1 | 8.631073 | 0.3003536 | 0.04873073 |
| **#3356** | DEPDC5 | 13.53492 | -0.2520938 | 0.03039065 |
| **#3379** | SLC33A1 | 19.0974 | -0.2298544 | 0.04757002 |
| **#3391** | PDCL | 16.46936 | 0.2168083 | 0.03287525 |
| **#3399** | FKRP | 6.152978 | -0.3549631 | 0.04207251 |
| **#3406** | AP4B1 | 7.279465 | -0.3034534 | 0.02349335 |
| **#3409** | ZNF24 | 58.83046 | 0.1330985 | 0.04565954 |
| **#3415** | SLC37A1 | 10.23606 | 0.2712609 | 0.04351328 |
| **#3422** | PCBD2 | 12.14667 | -0.2125677 | 0.001096374 |
| **#3453** | ZFYVE27 | 14.90753 | -0.2168118 | 0.01622986 |
| **#3491** | ALAD | 13.52765 | -0.2089633 | 0.007042441 |
| **#3497** | ZNF330 | 23.15334 | 0.1821757 | 0.03599924 |
| **#3505** | BTN2A1 | 23.42722 | -0.1975093 | 0.04340241 |
| **#3541** | PDRG1 | 13.33739 | 0.2280034 | 0.03824626 |
| **#3572** | CERS5 | 22.30797 | -0.1997942 | 0.0477285 |

| | | | | |
|---|---|---|---|---|
| **#3590** | TMTC4 | 7.206261 | -0.310045 | 0.04460755 |
| **#3614** | PRPF38A | 44.22572 | -0.1416476 | 0.04282082 |
| **#3648** | PKP4 | 5.53008 | -0.2715486 | 0.009804314 |
| **#3658** | XPO7 | 19.00085 | -0.1671296 | 0.004545069 |
| **#3662** | INAFM2 | 6.828233 | 0.283981 | 0.0422266 |
| **#3677** | TASP1 | 9.881993 | -0.2614315 | 0.04027327 |
| **#3689** | CANT1 | 11.17816 | 0.2221761 | 0.02809427 |
| **#3704** | UTP4 | 9.320289 | 0.2072335 | 0.005819518 |
| **#3743** | EME2 | 6.366983 | -0.2536126 | 0.01261695 |
| **#3754** | FAHD2A | 12.13829 | -0.2414125 | 0.04623928 |
| **#3769** | MRPS5 | 58.39856 | -0.1193831 | 0.04891215 |
| **#3772** | ATP13A2 | 14.0912 | -0.2208628 | 0.03994875 |
| **#3806** | XRCC1 | 14.49223 | -0.198911 | 0.02211916 |
| **#3816** | ENOPH1 | 18.74234 | 0.1788501 | 0.03493082 |
| **#3832** | POC1B | 15.18702 | -0.1932974 | 0.02202104 |
| **#3833** | SOCS7 | 6.939492 | -0.247292 | 0.01716535 |
| **#3849** | RBM4B | 7.154892 | -0.2493579 | 0.02126313 |
| **#3853** | TMEM199 | 13.97197 | 0.2063609 | 0.04361747 |
| **#3857** | TACO1 | 9.668796 | 0.235143 | 0.04313165 |
| **#3883** | TM2D2 | 9.558515 | 0.2277112 | 0.03825749 |
| **#3887** | WDR5 | 10.0217 | -0.2137054 | 0.01651405 |
| **#3897** | HSDL1 | 8.249523 | -0.2078003 | 0.005042334 |
| **#3923** | SNRNP35 | 22.83817 | 0.1651158 | 0.04640017 |
| **#3944** | PPP2R5D | 9.112748 | -0.2248466 | 0.02404329 |
| **#3979** | ZSCAN29 | 7.363394 | -0.2590622 | 0.0418554 |
| **#3981** | SLC38A9 | 9.297399 | -0.2430915 | 0.04563055 |
| **#4035** | HECTD3 | 7.649024 | -0.2033642 | 0.007799515 |
| **#4037** | SLC35B4 | 8.431907 | -0.2337245 | 0.03521228 |
| **#4072** | STRADA | 25.61698 | -0.1426711 | 0.0293987 |
| **#4117** | CCHCR1 | 6.348624 | -0.2572124 | 0.04587946 |
| **#4196** | C1RL | 11.67022 | -0.1940559 | 0.03630429 |
| **#4231** | MED24 | 13.29521 | -0.1859806 | 0.04198868 |

| | | | | |
|---|---|---|---|---|
| **#4243** | DHX33 | 9.611307 | -0.1831053 | 0.01737051 |
| **#4252** | SCLY | 6.972388 | -0.2296399 | 0.04211841 |
| **#4278** | AATF | 55.21052 | 0.0872772 | 0.03860913 |
| **#4291** | DHDDS | 12.81554 | -0.1672615 | 0.02126552 |
| **#4314** | MRPS10 | 32.11595 | 0.1143962 | 0.03580969 |
| **#4330** | AAR2 | 8.407262 | 0.204935 | 0.0457535 |
| **#4364** | VPS33A | 22.85636 | -0.1449555 | 0.04938394 |
| **#4386** | AAGAB | 15.29108 | 0.1535539 | 0.03559103 |
| **#4389** | MLLT1 | 15.3438 | -0.1604231 | 0.03649178 |
| **#4397** | BRCC3 | 21.79282 | -0.1440318 | 0.04701323 |
| **#4462** | MAN2B2 | 20.88994 | -0.1340434 | 0.03253858 |
| **#4475** | SS18L1 | 7.191391 | -0.2029839 | 0.03743825 |
| **#4498** | ESS2 | 8.557155 | -0.1927482 | 0.03977954 |
| **#4616** | MRPS27 | 16.36841 | -0.1336182 | 0.02537265 |
| **#4740** | LYRM2 | 42.58223 | 0.08312102 | 0.04299526 |
| **#4817** | MRPL45 | 16.49904 | -0.134382 | 0.0450107 |

Supplementary Table 5

| Position | Symbol | Avg Rank | Rank (p) | p M/F | Rank (Rel Chg) | Rel Chg M/F | Rank (Abs Chg) | Abs Chg M/F |
|---|---|---|---|---|---|---|---|---|
| #1 | TSC22D3 | 92.66667 | 74 | 0.64/0.0386 | 68 | −0.2138/0.6205 | 137 | −62.38442/158.9083 |
| #2 | BTG1 | 190.6667 | 391 | 0.291/0.772 | 70 | −0.7852/0.04096 | 112 | −373.0795/39.89105 |
| #3 | CEBPD | 200 | 383 | 0.972/0.0896 | 17 | −0.09634/1.041 | 201 | −2.981401/130.6674 |
| #4 | IGLC2 | 210 | 317 | 0.562/0.457 | 30 | −0.7242/0.2968 | 284 | −102.0969/102.7643 |
| #5 | DDIT4 | 219 | 412 | 0.976/0.102 | 47 | −0.05573/0.8736 | 199 | −2.023091/82.91532 |
| #6 | CXCR4 | 225.3333 | 196 | 0.79/0.102 | 235 | −0.1225/0.4356 | 246 | −20.68296/101.1385 |
| #7 | ZFP36L2 | 228.6667 | 529 | 0.122/0.984 | 56 | −0.8942/−0.01653 | 102 | −345.9343/2.073018 |
| #8 | CEBPB | 282 | 161 | 0.76/0.0917 | 94 | −0.004263/0.7533 | 592 | 28.46287/102.8803 |
| #9 | KCNE3 | 333 | 42 | 0.293/0.253 | 24 | 0.2825/−0.7866 | 934 | 4.615898/−21.26329 |
| #10 | IGHD | 375.3333 | 60 | 0.183/0.456 | 4 | 1.272/−0.5782 | 1063 | 98.81362/−24.89462 |
| #11 | PNRC1 | 394.6667 | 815 | 0.471/0.783 | 181 | −0.564/0.04132 | 189 | −105.985/29.1781 |
| #12 | ANXA1 | 421.3333 | 657 | 0.963/0.227 | 457 | −0.02263/0.4127 | 151 | −5.326782/115.5896 |
| #13 | STUB1 | 425 | 221 | 0.377/0.547 | 170 | −0.5288/0.09552 | 885 | −24.65415/5.628968 |
| #14 | YBX1 | 434 | 132 | 0.584/0.23 | 156 | −0.3833/0.273 | 1015 | −103.7944/147.0025 |
| #15 | KRT10 | 466.3333 | 472 | 0.533/0.579 | 92 | −0.7026/0.06155 | 836 | −33.948/5.169405 |
| #16 | ZFAND5 | 478 | 119 | 0.305/0.484 | 202 | −0.3969/0.187 | 1114 | −31.94935/24.18409 |
| #17 | IGKC | 486.3333 | 719 | 0.778/0.439 | 577 | −0.005124/0.3979 | 164 | 132.7663/345.1827 |
| #18 | TAF10 | 500 | 474 | 0.612/0.5 | 82 | −0.6054/0.1817 | 945 | −24.42961/15.60362 |
| #19 | ID2 | 511.6667 | 684 | 0.878/0.324 | 403 | −0.01129/0.4465 | 449 | −15.73668/110.7724 |
| #20 | MALT1 | 516 | 98 | 0.451/0.301 | 573 | −0.2865/0.118 | 878 | −8.833314/6.975856 |
| #21 | SRSF9 | 533.6667 | 235 | 0.769/0.174 | 157 | −0.4204/0.2331 | 1210 | −16.92879/24.17413 |
| #22 | IRS2 | 537.3333 | 92 | 0.679/0.0678 | 143 | −0.11/0.5574 | 1378 | −13.46179/25.57939 |
| #23 | CD22 | 543.6667 | 377 | 0.624/0.434 | 43 | 0.4788/−0.4748 | 1212 | 6.829494/−9.265638 |
| #24 | VIM | 562.3333 | 647 | 0.65/0.536 | 916 | −0.1106/0.2242 | 125 | −110.8776/238.4299 |
| #25 | ETS1 | 565.6667 | 498 | 0.547/0.577 | 644 | −0.2439/0.1426 | 556 | −52.47739/36.21376 |
| #26 | EFHD2 | 576 | 1191 | 0.987/0.378 | 267 | −0.05541/0.4813 | 271 | 2.537905/98.59493 |
| #27 | AREG | 587.3333 | 1164 | 0.809/0.167 | 11 | 0.4771/1.782 | 588 | −13.36104/71.02494 |
| #28 | TNFAIP3 | 595 | 401 | 0.974/0.0444 | 262 | 0.07237/0.6121 | 1123 | −1.213821/72.92234 |
| #29 | RIPOR2 | 618.3333 | 887 | 0.857/0.424 | 847 | 0.02793/−0.3167 | 122 | 10.32209/−93.29529 |
| #30 | MAF 1.00 | 623.6667 | 207 | 0.63/0.277 | 562 | −0.2478/0.1586 | 1103 | −9.773047/7.532456 |
| #31 | CLTB | 640.6667 | 210 | 0.629/0.281 | 105 | −0.451/0.2802 | 1608 | −11.48437/11.75842 |
| #32 | UTRN | 642.6667 | 385 | 0.81/0.252 | 1375 | 0.07044/−0.2063 | 169 | 6.009271/−37.64169 |
| #33 | GPR183 | 648.3333 | 187 | 0.762/0.123 | 65 | −0.1524/0.6844 | 1694 | −6.177054/74.65431 |
| #34 | TRADD | 655 | 805 | 0.392/0.859 | 74 | −0.8038/0.01594 | 1087 | −19.87919/0.5249725 |
| #35 | SH3BP5 | 672 | 200 | 0.486/0.409 | 675 | −0.2555/0.1249 | 1142 | −12.67259/7.244909 |
| #36 | EGR1 | 679.3333 | 408 | 0.845/0.231 | 9 | −0.5118/0.8541 | 1622 | 7.276196/18.24707 |
| #37 | PLCG2 | 689.6667 | 1735 | 0.688/0.171 | 225 | −0.255/−0.8199 | 110 | −14.44922/−150.2415 |
| #38 | HNRNPD | 705.6667 | 172 | 0.614/0.248 | 1198 | −0.216/0.07969 | 748 | −18.78298/12.48615 |
| #39 | SERP1 | 713.6667 | 164 | 0.704/0.153 | 1387 | −0.07016/0.2057 | 591 | −13.19945/44.34569 |
| #40 | KIRREL3 | 714.6667 | 102 | 0.185/0.575 | 1 | −3.216/0.1704 | 2042 | −14.72953/2.94772 |
| #41 | CTDNEP1 | 722 | 111 | 0.426/0.352 | 865 | −0.1098/0.2308 | 1191 | −6.619447/11.27828 |

| | | | | | | | | |
|---|---|---|---|---|---|---|---|---|
| #42 | JUNB | 732.6667 | 663 | 0.58/0.613 | 540 | −0.3093/0.1029 | 996 | −179.141/120.3342 |
| #43 | MT-ATP8 | 734 | 1255 | 0.732/0.649 | 864 | −0.1773/0.1634 | 84 | −6.709693/109.0652 |
| #44 | PTGER2 | 745.6667 | 337 | 0.888/0.143 | 362 | −0.03405/0.4431 | 1539 | −1.067248/8.434101 |
| #45 | RAB5IF | 755.3333 | 1372 | 0.633/0.0458 | 353 | 0.1895/0.6703 | 542 | 16.42812/49.34115 |
| #46 | CMTM3 | 761.6667 | 269 | 0.765/0.212 | 614 | −0.04291/0.3506 | 1403 | −2.359585/11.68916 |
| #47 | TRAPPC5 | 766.6667 | 435 | 0.933/0.155 | 768 | −0.001861/0.3581 | 1098 | 2.492976/42.9187 |
| #48 | OGFRL1 | 798 | 537 | 0.384/0.758 | 1363 | −0.1726/0.1061 | 495 | −9.689944/8.114907 |
| #49 | FOS | 798.6667 | 2027 | 0.556/0.0897 | 294 | 0.1216/0.6373 | 76 | 86.84876/715.2239 |
| #50 | C1QBP | 802.6667 | 63 | 0.519/0.125 | 41 | −0.7937/0.181 | 2305 | −27.01073/13.95423 |
| #51 | PTEN | 805 | 528 | 0.435/0.703 | 887 | −0.2903/0.04767 | 1001 | −14.73052/4.896909 |
| #52 | CHD9 | 815.3333 | 152 | 0.606/0.236 | 1428 | 0.1035/−0.1682 | 867 | 4.141067/−9.830424 |
| #53 | MKLN1 | 818.6667 | 94 | 0.732/0.016 | 1047 | 0.04885/−0.266 | 1316 | 1.459658/−9.960103 |
| #54 | BRI3 | 831.3333 | 1684 | 0.992/0.484 | 355 | −0.1655/0.314 | 456 | 0.9950533/53.50307 |
| #55 | ARF6 | 835 | 318 | 0.858/0.162 | 838 | −0.1162/0.2294 | 1350 | −7.334056/25.81052 |
| #56 | IGFBP7 | 838.3333 | 1031 | 0.861/0.462 | 52 | −0.283/0.6257 | 1433 | 7.818179/18.99259 |
| #57 | RTN4 | 844.6667 | 527 | 0.936/0.201 | 964 | −0.01194/0.3157 | 1044 | −1.772504/30.30898 |
| #58 | NFKBIA | 850 | 1632 | 0.714/0.181 | 610 | 0.1771/0.5713 | 309 | 29.02027/189.7781 |
| #59 | TM9SF3 | 855.3333 | 549 | 0.439/0.711 | 1114 | −0.2503/0.05444 | 904 | −10.4149/3.972786 |
| #60 | KLF9 | 856 | 39 | 0.513/0.0135 | 95 | −0.2176/0.5396 | 2435 | −9.815586/20.73176 |
| #61 | ASCL2 | 862.6667 | 116 | 0.77/0.0118 | 23 | −0.05773/1.028 | 2450 | 3.462601/13.35402 |
| #62 | ASCC3 | 863 | 574 | 0.0267/0.866 | 1186 | 0.3159/0.01857 | 830 | 7.240649/1.585222 |
| #63 | COBLL1 | 877.6667 | 1477 | 0.225/0.788 | 25 | 1.115/0.05766 | 1132 | 17.12544/4.042086 |
| #64 | ELF1 | 882.6667 | 773 | 0.815/0.0528 | 961 | 0.02208/0.3505 | 915 | 2.008766/45.07598 |
| #65 | TSR3 | 895.6667 | 700 | 0.365/0.843 | 133 | −0.6507/0.02728 | 1855 | −13.58081/1.099267 |
| #66 | PTGER4 | 900 | 667 | 0.836/0.359 | 1201 | −0.01857/0.277 | 833 | 3.067761/18.96639 |
| #67 | TMEM203 | 902 | 388 | 0.533/0.529 | 138 | −0.5451/0.128 | 2181 | −8.216382/2.444313 |
| #68 | ERN 1.00 | 906.3333 | 400 | 0.329/0.741 | 410 | −0.3942/0.0608 | 1910 | −10.08728/2.386453 |
| #69 | SAMD9 | 906.3333 | 1302 | 0.3/0.903 | 712 | 0.3865/0.01538 | 706 | 20.19205/3.815692 |
| #70 | MS4A1 | 911.3333 | 2189 | 0.961/0.518 | 271 | −0.2511/−0.7841 | 275 | −3.969349/−51.96244 |
| #71 | SAMSN1 | 912.3333 | 259 | 0.852/0.118 | 80 | −0.0136/0.7788 | 2399 | −2.258439/43.44919 |
| #72 | GSPT1 | 918 | 497 | 0.395/0.728 | 436 | −0.4151/0.02944 | 1822 | −12.32442/1.741724 |
| #73 | NENF | 923.3333 | 177 | 0.377/0.491 | 60 | −0.718/0.1447 | 2534 | −13.26323/4.423658 |
| #74 | CTNNB1 | 927.6667 | 611 | 0.63/0.542 | 1419 | −0.1199/0.1526 | 754 | −10.45175/18.96296 |
| #75 | EREG | 936.6667 | 284 | 0.876/0.115 | 136 | −0.04291/0.6319 | 2391 | −1.582068/21.08857 |
| #76 | PARP15 | 942.3333 | 1445 | 0.68/0.109 | 129 | −0.0828/−0.7682 | 1254 | −2.537288/−18.8743 |
| #77 | CDKN2D | 954.3333 | 1466 | 0.822/0.257 | 500 | 0.0009003/0.4235 | 898 | 13.56538/43.39746 |
| #78 | TMEM131L | 954.3333 | 512 | 0.268/0.864 | 1327 | 0.2408/−0.04087 | 1025 | 5.369002/−1.219296 |
| #79 | SIRT1 | 960.3333 | 592 | 0.142/0.976 | 536 | −0.4696/−0.05668 | 1754 | −4.842156/0.2602881 |
| #80 | KLHL6 | 960.3333 | 310 | 0.4/0.609 | 332 | 0.1937/−0.301 | 2240 | 1.89654/−3.768608 |
| #81 | BRWD3 | 964 | 80 | 0.468/0.231 | 1123 | 0.08505/−0.2186 | 1690 | 1.4069/−5.522763 |
| #82 | CCDC141 | 964.6667 | 888 | 0.887/0.168 | 290 | −0.2969/−0.8162 | 1717 | −0.5891305/−9.911013 |
| #83 | WDFY4 | 965 | 991 | 0.287/0.977 | 251 | 0.5644/0.01354 | 1654 | 7.671877/0.1359729 |
| #84 | TTC21B | 967.6667 | 712 | 0.83/0.0436 | 388 | −0.02013/−0.4888 | 1804 | −0.2665991/−8.18736 |
| #85 | CCDC149 | 976.6667 | 156 | 0.347/0.497 | 35 | 0.7602/−0.2426 | 2740 | 5.943563/−2.73839 |

| | | | | | | | | |
|---|---|---|---|---|---|---|---|---|
| **#86** | TMEM156 | 982.6667 | 852 | 0.773/0.0411 | 638 | −0.1213/−0.5089 | 1459 | −2.028226/−12.69378 |
| **#87** | SSU72 | 983 | 211 | 0.627/0.284 | 1925 | −0.1468/0.08211 | 814 | −10.0095/6.037681 |
| **#88** | IL2RB | 985.3333 | 1737 | 0.775/0.709 | 616 | 0.04614/−0.347 | 604 | 15.67676/−7.842736 |
| **#89** | ICE1 | 994.3333 | 1003 | 0.306/0.993 | 418 | −0.454/−0.002648 | 1563 | −8.293656/0.0418207 |
| **#90** | HLA-DRB5 | 997 | 1453 | 0.0339/0.602 | 851 | 0.4971/0.8405 | 688 | 139.5573/157.8253 |
| **#91** | EHBP1L1 | 1,009 | 1224 | 0.641/0.733 | 384 | −0.3628/0.1077 | 1421 | −10.48169/6.323071 |
| **#92** | NELL2 | 1,010 | 280 | 0.866/0.123 | 128 | 0.001124/−0.686 | 2622 | −1.380213/−12.72658 |
| **#93** | H2AFV | 1,012 | 626 | 0.459/0.721 | 1255 | −0.2738/0.01557 | 1156 | −20.77722/1.380398 |
| **#94** | CAST | 1,014 | 1540 | 0.975/0.476 | 1278 | −0.01564/0.2713 | 225 | 1.408705/68.55397 |
| **#95** | TMEM160 | 1,022 | 1468 | 0.524/0.911 | 63 | −0.8315/0.01295 | 1537 | −23.65856/1.453059 |
| **#96** | CD4 | 1,023 | 677 | 0.148/0.948 | 1071 | −0.3143/−0.003864 | 1323 | −15.94472/1.087953 |
| **#97** | MAP3K1 | 1,027 | 534 | 0.746/0.394 | 2147 | 0.05336/−0.1613 | 400 | 2.731445/−13.54576 |
| **#98** | CYC1 | 1,045 | 1012 | 0.876/0.192 | 646 | 0.01982/0.4062 | 1477 | 2.325074/16.42522 |
| **#99** | UBR1 | 1,049 | 629 | 0.873/0.0537 | 671 | −0.01665/−0.3978 | 1848 | −0.4338773/−8.242517 |
| **#100** | MT-ND4L | 1,050 | 1136 | 0.548/0.802 | 1973 | −0.1972/0.02855 | 42 | −34.54056/42.56464 |
| **#101** | MTRNR2L1 | 1,051 | 2077 | 0.585/0.127 | 1028 | −0.3224/−0.6413 | 49 | −104.6867/−378.4587 |
| **#102** | NR4A2 | 1,054 | 1598 | 0.876/0.338 | 700 | 0.02902/0.4041 | 866 | 2.802321/37.12582 |
| **#103** | AKR7A2 | 1,055 | 736 | 0.348/0.877 | 137 | −0.6459/0.02796 | 2292 | −12.21606/0.8262304 |
| **#104** | PPP1R10 | 1,055 | 1558 | 0.69/0.146 | 1037 | −0.0663/−0.383 | 571 | −2.219565/−20.8043 |
| **#105** | S100A10 | 1,065 | 1681 | 0.998/0.476 | 1361 | −0.0006185/0.2781 | 155 | 0.2674009/105.0944 |
| **#106** | HEBP2 | 1,068 | 59 | 0.399/0.238 | 100 | −0.587/0.1527 | 3045 | −27.29063/11.44636 |
| **#107** | PRDM1 | 1,083 | 1497 | 0.56/0.000883 | 67 | 0.2366/1.072 | 1686 | 5.037256/24.1978 |
| **#108** | IER5 | 1,095 | 1890 | 0.944/0.454 | 877 | −0.03259/−0.3716 | 518 | 2.0394/−24.40601 |
| **#109** | HNRNPAB | 1,096 | 1332 | 0.734/0.67 | 414 | −0.3702/0.08298 | 1544 | −8.66118/6.041795 |
| **#110** | RAD23B | 1,097 | 639 | 0.834/0.349 | 1367 | −0.09222/0.1857 | 1286 | −1.994724/10.33224 |
| **#111** | CD79B | 1,099 | 1041 | 0.861/0.186 | 941 | −0.1486/−0.4795 | 1315 | −7.870342/−33.35685 |
| **#112** | USP33 | 1,100 | 126 | 0.579/0.225 | 2244 | 0.05562/−0.1529 | 930 | 1.939624/−7.353325 |
| **#113** | ZFP91 | 1,103 | 697 | 0.984/0.221 | 926 | −0.008795/0.3247 | 1687 | −0.1594518/19.60532 |
| **#114** | NFE2L2 | 1,104 | 774 | 0.928/0.166 | 444 | 0.04535/0.4874 | 2096 | 1.018954/33.19442 |
| **#115** | MT-ATP6 | 1,105 | 1201 | 0.934/0.435 | 2106 | 0.02139/−0.1953 | 9 | 142.566/−1,198 |
| **#116** | BANK1 | 1,109 | 2511 | 0.711/0.887 | 475 | 0.1812/−0.2482 | 343 | 16.31269/11.02106 |
| **#117** | TNFSF12 | 1,111 | 863 | 0.43/0.843 | 709 | −0.304/0.06789 | 1762 | −7.593078/1.407563 |
| **#118** | WDR70 | 1,111 | 344 | 0.985/0.0228 | 1440 | −0.01759/−0.2884 | 1550 | −0.06898207/−8.48307 |
| **#119** | CXXC5 | 1,116 | 316 | 0.83/0.188 | 333 | −0.1754/0.3192 | 2701 | −4.867959/14.8983 |
| **#120** | LITAF | 1,129 | 1803 | 0.872/0.368 | 1260 | 0.01265/0.3011 | 326 | 8.992244/35.19245 |
| **#121** | CDCA7L | 1,131 | 727 | 0.902/0.318 | 390 | 0.03777/−0.4304 | 2277 | 0.95279/−6.29139 |
| **#122** | SRBD1 | 1,136 | 81 | 0.198/0.508 | 717 | 0.2087/−0.1614 | 2611 | 2.379535/−2.405651 |
| **#123** | METTL9 | 1,142 | 66 | 0.504/0.148 | 599 | −0.2078/0.1893 | 2761 | −12.22571/13.71205 |
| **#124** | SGK1 | 1,144 | 936 | 0.86/0.156 | 109 | 0.00855/0.7309 | 2389 | 1.034867/23.9565 |
| **#125** | POU2F2 | 1,146 | 1177 | 0.871/0.49 | 1636 | 0.03077/−0.2228 | 625 | 3.144893/−27.82817 |
| **#126** | ZC3H7A | 1,147 | 423 | 0.54/0.542 | 2036 | −0.1508/0.07045 | 983 | −3.804782/3.797584 |
| **#127** | BCL2 | 1,147 | 1996 | 0.374/0.845 | 625 | −0.4915/−0.1003 | 822 | −14.37329/−5.770694 |
| **#128** | CHST11 | 1,147 | 148 | 0.783/0.054 | 130 | −0.2351/0.4479 | 3165 | −2.896897/10.34837 |
| **#129** | TXNL4A | 1,157 | 332 | 0.931/0.0967 | 1275 | −0.04486/0.2422 | 1866 | −1.239121/12.94288 |

| | | | | | | | | |
|---|---|---|---|---|---|---|---|---|
| **#130** | ACAA1 | 1,157 | 415 | 0.874/0.205 | 565 | −0.05947/0.3465 | 2493 | −2.521236/15.95365 |
| **#131** | MCPH1 | 1,165 | 84 | 0.0191/0.698 | 1983 | 0.1795/−0.04537 | 1430 | 3.114358/−1.209586 |
| **#132** | GADD45GI | 1,171 | 231 | 0.872/0.0694 | 1838 | −0.03791/0.1988 | 1444 | −1.707998/10.56378 |
| **#133** | CHMP4B | 1,172 | 624 | 0.873/0.0519 | 912 | 0.01105/0.346 | 1982 | 1.319925/18.52 |
| **#134** | KDM5C | 1,175 | 69 | 0.442/0.213 | 1669 | 0.05594/−0.1952 | 1789 | 0.6924377/−4.05622 |
| **#135** | NEDD9 | 1,178 | 176 | 0.554/0.311 | 1676 | 0.1409/−0.1098 | 1683 | 3.226368/−3.136693 |
| **#136** | VSTM1 | 1,181 | 107 | 0.581/0.187 | 69 | 0.2949/−0.5357 | 3368 | 7.666033/−6.338959 |
| **#137** | HEXB | 1,183 | 25 | 0.145/0.314 | 607 | −0.1512/0.2433 | 2919 | −4.286486/5.870584 |
| **#138** | DDX52 | 1,189 | 43 | 0.524/0.0236 | 850 | 0.08209/−0.2616 | 2674 | 2.396384/−8.365586 |
| **#139** | CNTRL | 1,195 | 1676 | 0.817/0.292 | 1258 | −0.06014/−0.3488 | 652 | −2.071892/−20.8675 |
| **#140** | NIBAN3 | 1,198 | 701 | 0.79/0.419 | 97 | 0.3517/−0.4004 | 2798 | 3.397538/−6.51656 |
| **#141** | CCNI | 1,200 | 311 | 0.693/0.318 | 2978 | −0.0416/0.1299 | 312 | −13.741/38.28685 |
| **#142** | SNX9 | 1,205 | 127 | 0.429/0.377 | 375 | −0.2187/0.255 | 3115 | −4.114108/6.749356 |
| **#143** | KHDRBS1 | 1,215 | 299 | 0.66/0.342 | 834 | −0.2202/0.1266 | 2514 | −12.15392/13.08804 |
| **#144** | LGALS2 | 1,220 | 2373 | 0.783/0.364 | 755 | 0.09787/0.4611 | 532 | 11.10639/52.30522 |
| **#145** | TIMP1 | 1,220 | 2258 | 0.786/0.353 | 1070 | 0.1499/0.4607 | 334 | 17.48001/44.26648 |
| **#146** | NSF | 1,222 | 1269 | 0.965/0.351 | 1381 | −0.02602/−0.3024 | 1018 | −0.177219/−7.858689 |
| **#147** | GOLGA4 | 1,223 | 1988 | 0.688/0.216 | 1348 | −0.03271/−0.3129 | 333 | −2.663912/−33.94718 |
| **#148** | UTS2 | 1,224 | 1309 | 0.851/0.547 | 21 | 0.3843/−0.728 | 2342 | −1.427108/−0.7120987 |
| **#149** | TMEM165 | 1,225 | 195 | 0.546/0.346 | 649 | −0.2454/0.1404 | 2832 | −8.843128/6.820554 |
| **#150** | SET | 1,226 | 270 | 0.907/0.0721 | 2448 | −0.00129/0.1957 | 960 | 7.239555/57.35646 |
| **#151** | ANTKMT | 1,226 | 1277 | 0.568/0.821 | 46 | −0.9126/0.01751 | 2357 | −12.25216/0.5514738 |
| **#152** | FDX1 | 1,230 | 183 | 0.527/0.349 | 119 | −0.5239/0.1744 | 3389 | −8.688647/4.798374 |
| **#153** | LYST | 1,233 | 1187 | 0.563/0.8 | 2223 | 0.1251/−0.08437 | 291 | 11.32917/−8.502828 |
| **#154** | CLEC10A | 1,238 | 1169 | 0.87/0.229 | 176 | 0.2491/0.8637 | 2371 | −4.824289/29.45297 |
| **#155** | DBF4 | 1,239 | 465 | 0.549/0.561 | 1289 | 0.09369/−0.1917 | 1963 | 2.03639/−4.149277 |
| **#156** | ERP29 | 1,239 | 48 | 0.253/0.328 | 3089 | −0.0965/0.07046 | 582 | −18.39366/12.29742 |
| **#157** | R3HCC1L | 1,241 | 206 | 0.735/0.171 | 1015 | 0.05238/−0.2688 | 2502 | 0.7097079/−4.241647 |
| **#158** | RNASE2 | 1,241 | 753 | 0.0364/0.802 | 42 | 1.163/0.1936 | 2928 | 14.23126/3.072477 |
| **#159** | CTPS1 | 1,242 | 462 | 0.73/0.378 | 484 | 0.1533/−0.2745 | 2782 | 2.485887/−4.617252 |
| **#160** | TLE1 | 1,243 | 813 | 0.518/0.736 | 524 | −0.3785/0.03738 | 2394 | −9.302449/3.036352 |
| **#161** | TRIM44 | 1,243 | 492 | 0.343/0.777 | 2704 | 0.1259/−0.0582 | 535 | 7.607878/−4.243705 |
| **#162** | SAP30 | 1,253 | 149 | 0.41/0.428 | 33 | −0.5887/0.4166 | 3579 | −6.00552/8.326968 |
| **#163** | CST3 | 1,255 | 3086 | 0.48/0.14 | 316 | 0.3225/0.8279 | 363 | 191.2488/656.3507 |
| **#164** | NME4 | 1,258 | 1256 | 0.382/0.999 | 601 | −0.348/0.04861 | 1919 | −7.285394/−0.0046291 |
| **#165** | SETBP1 | 1,266 | 1087 | 0.745/0.593 | 492 | 0.2679/−0.1566 | 2220 | 1.260513/−2.898054 |
| **#166** | SFPQ | 1,267 | 600 | 0.725/0.444 | 2969 | −0.06641/0.1056 | 234 | −7.154654/20.44898 |
| **#167** | CMTM7 | 1,270 | 141 | 0.259/0.568 | 507 | −0.2599/0.1611 | 3162 | −7.204854/3.907818 |
| **#168** | YLPM1 | 1,280 | 1139 | 0.71/0.0615 | 1079 | −0.04292/−0.3526 | 1623 | −1.003722/−10.24366 |
| **#169** | LAGE3 | 1,281 | 180 | 0.502/0.369 | 621 | −0.2725/0.1192 | 3042 | −5.056332/2.334698 |
| **#170** | TP53BP2 | 1,284 | 749 | 0.75/0.481 | 994 | −0.1332/0.1903 | 2111 | −1.168884/4.79487 |
| **#171** | GABPB1-IT | 1,286 | 7 | 0.239/0.0137 | 174 | −0.4602/0.1572 | 3677 | −16.8432/7.287802 |
| **#172** | ABCF1 | 1,288 | 15 | 0.334/0.0642 | 1350 | 0.08893/−0.1912 | 2500 | 4.233266/−9.537012 |
| **#173** | GNAI2 | 1,290 | 2312 | 0.756/0.328 | 1064 | 0.07944/0.3908 | 494 | 19.71325/107.3833 |

| | | | | | | | | |
|---|---|---|---|---|---|---|---|---|
| **#174** | KLRG1 | 1,291 | 1849 | 0.548/0.0501 | 956 | −0.4453/−0.7743 | 1070 | −11.48997/−13.38697 |
| **#175** | LAMP1 | 1,292 | 426 | 0.808/0.276 | 1707 | −0.08832/0.1597 | 1745 | −4.025415/9.447756 |
| **#176** | YBX3 | 1,297 | 871 | 0.896/0.378 | 427 | −0.04003/0.407 | 2594 | −2.795244/31.17034 |
| **#177** | ARHGAP17 | 1,300 | 1125 | 0.381/0.967 | 1124 | 0.2849/−0.01861 | 1653 | 6.308044/0.2933791 |
| **#178** | TOB1 | 1,301 | 292 | 0.12/0.878 | 140 | −0.6525/0.0164 | 3472 | −32.52694/2.959069 |
| **#179** | ARL2BP | 1,302 | 490 | 0.388/0.731 | 1307 | −0.2142/0.06928 | 2110 | −5.972549/1.905881 |
| **#180** | FXYD5 | 1,302 | 460 | 0.0755/0.967 | 3266 | −0.1836/−0.02397 | 182 | −48.90202/3.168256 |
| **#181** | EIF3L | 1,309 | 410 | 0.451/0.626 | 2612 | −0.1043/0.08384 | 906 | −14.82888/12.03002 |
| **#182** | HIST1H2AC | 1,313 | 309 | 0.393/0.616 | 868 | 0.1168/−0.2236 | 2762 | 1.938849/−4.592509 |
| **#183** | ERVK13-1 | 1,315 | 601 | 0.902/0.0715 | 275 | −0.02405/−0.5535 | 3069 | −0.105631/−5.616977 |
| **#184** | ERVK3-1 | 1,315 | 214 | 0.806/0.107 | 1376 | 0.02625/−0.2504 | 2356 | 0.9030775/−6.905852 |
| **#185** | NLRP12 | 1,316 | 70 | 0.23/0.429 | 27 | 0.6527/−0.3963 | 3852 | 7.135831/−5.002041 |
| **#186** | TBC1D22A | 1,319 | 125 | 0.322/0.481 | 2197 | 0.1046/−0.1066 | 1637 | 2.536494/−3.289666 |
| **#187** | ITGB7 | 1,320 | 470 | 0.957/0.0685 | 2307 | 0.01721/0.2221 | 1185 | 0.3903012/7.086442 |
| **#188** | TMUB1 | 1,323 | 1101 | 0.405/0.938 | 107 | −0.723/0.002058 | 2763 | −9.667614/0.1999915 |
| **#189** | PTAFR | 1,324 | 192 | 0.133/0.756 | 83 | 0.6651/−0.1215 | 3698 | 15.47221/−2.716412 |
| **#190** | WASHC4 | 1,325 | 659 | 0.633/0.558 | 2360 | 0.1119/−0.08937 | 956 | 5.1284/−4.223231 |
| **#191** | BASP1 | 1,328 | 605 | 0.937/0.108 | 1491 | 0.02217/0.2883 | 1888 | 0.7831559/12.84792 |
| **#192** | PHB2 | 1,330 | 150 | 0.231/0.61 | 2902 | −0.08335/0.09213 | 938 | −6.730335/5.786634 |
| **#193** | TMEM248 | 1,332 | 163 | 0.644/0.211 | 1292 | −0.1864/0.09854 | 2541 | −4.753753/3.092028 |
| **#194** | SYNE2 | 1,333 | 3075 | 0.525/0.184 | 715 | −0.1339/−0.5046 | 211 | −21.53797/−87.05228 |
| **#195** | CCDC50 | 1,334 | 1217 | 0.651/0.0248 | 1917 | 0.3845/0.6144 | 869 | 7.638694/37.82234 |
| **#196** | KANSL2 | 1,334 | 1102 | 0.821/0.164 | 1158 | −0.09114/−0.392 | 1743 | −1.091471/−8.678858 |
| **#197** | PEX26 | 1,338 | 157 | 0.531/0.316 | 849 | 0.1027/−0.2417 | 3008 | 1.851773/−4.52737 |
| **#198** | IGLC3 | 1,339 | 1951 | 0.447/0.926 | 79 | −0.8352/−0.02044 | 1989 | −39.47104/2.842469 |
| **#199** | EIF3F | 1,341 | 158 | 0.253/0.594 | 3556 | −0.06352/0.08447 | 310 | −14.74266/17.80176 |
| **#200** | IL13RA1 | 1,343 | 1351 | 0.646/0.0545 | 55 | 0.1211/1.003 | 2623 | 3.66311/32.59613 |
