## Supplementary figures and images for "BNT162b2 mRNA vaccine-induced sex differences in the single-cell transcriptome of peripheral blood mononuclear cells in healthy adults"

### Supplemental Fig. S1

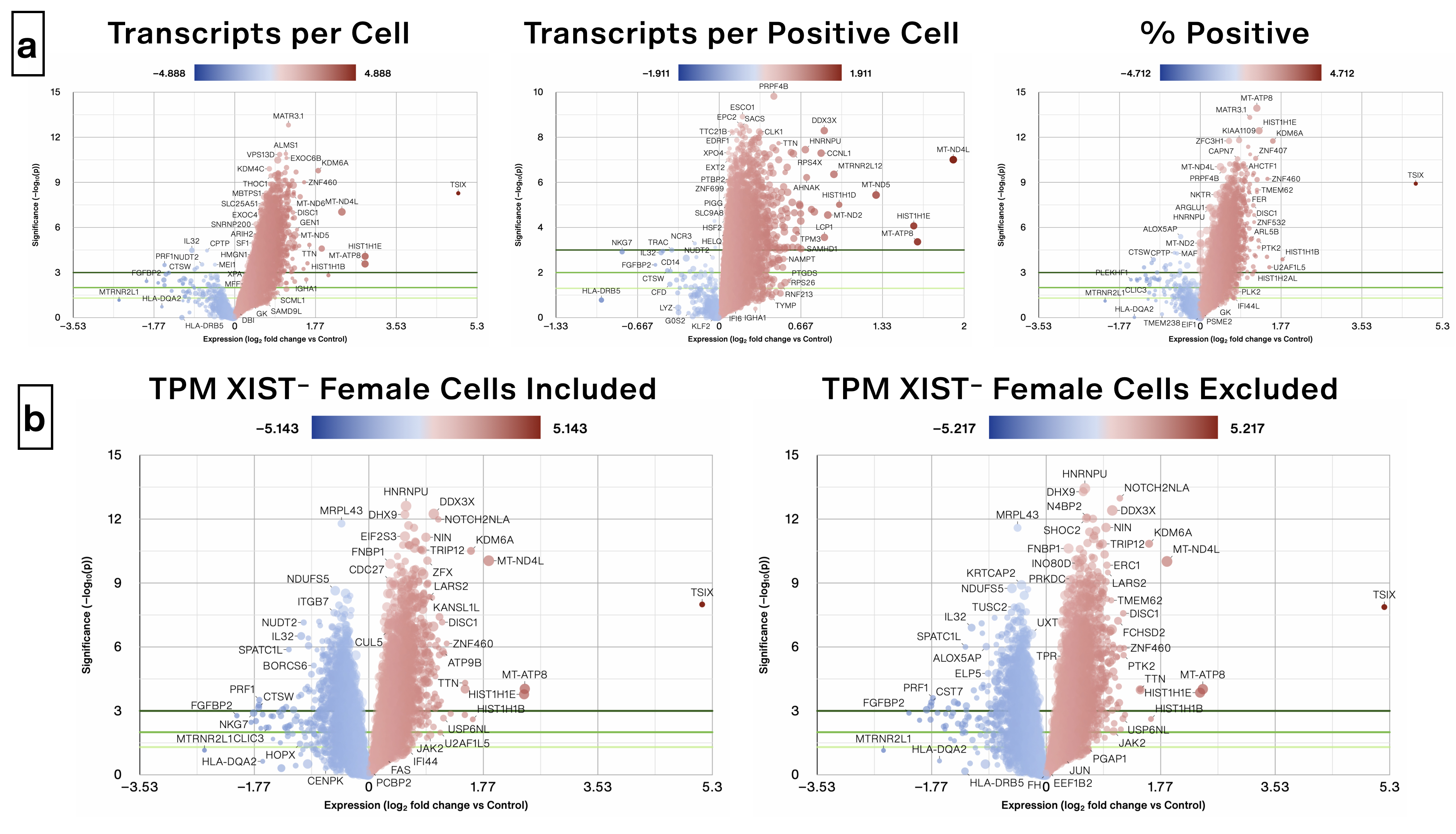

### Supplemental Fig. S2

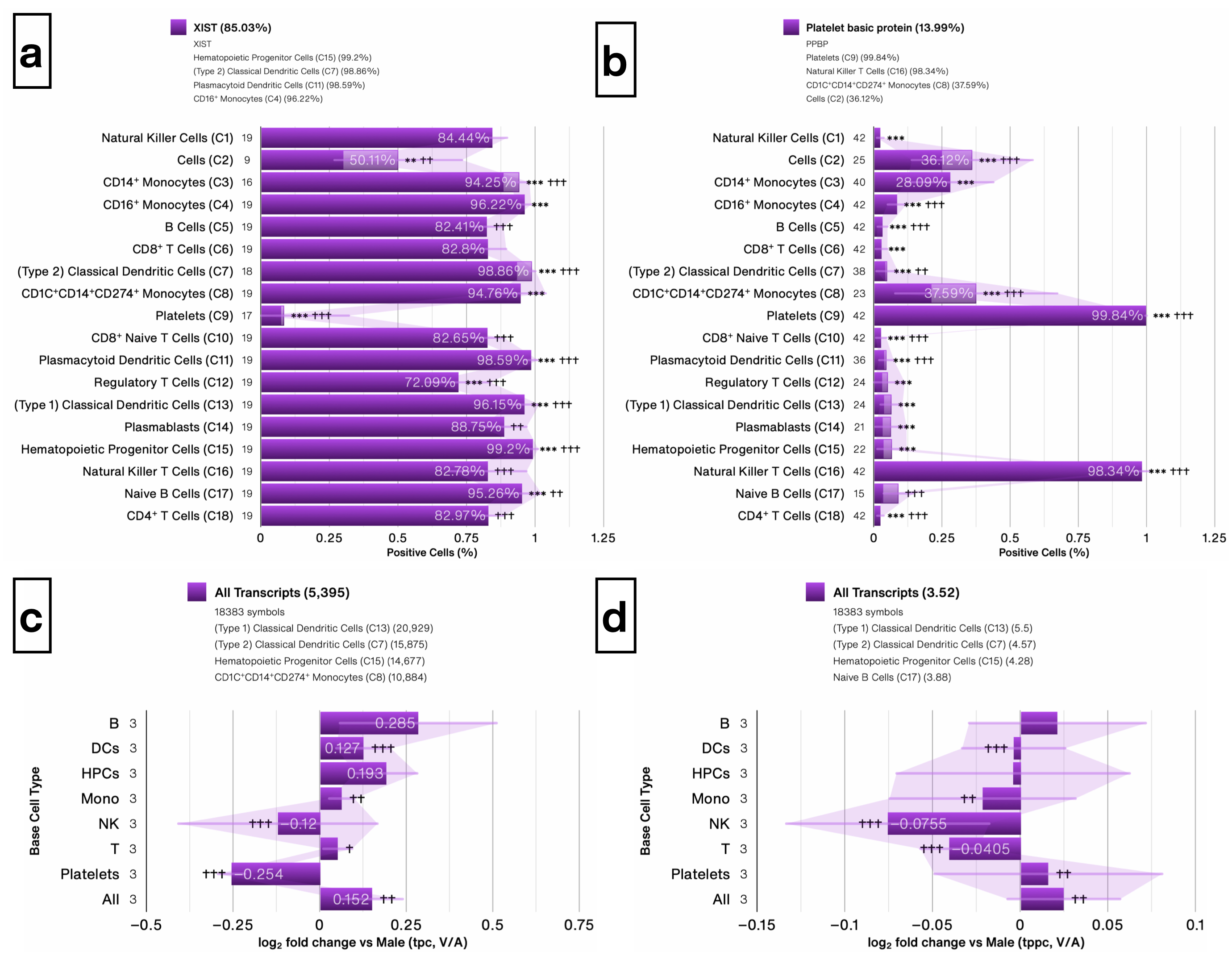

### Supplemental Fig. S3

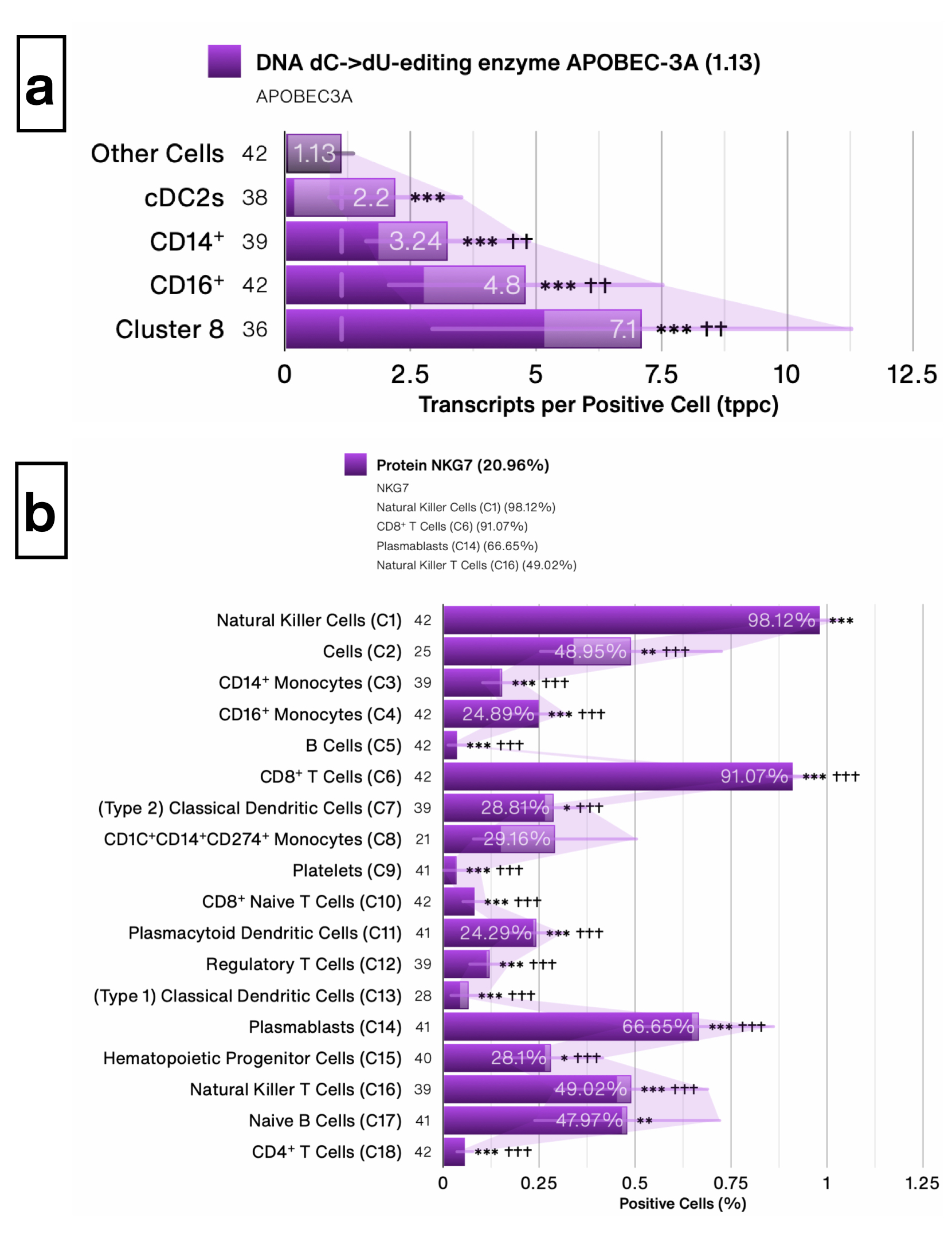

### Supplemental Fig. S4

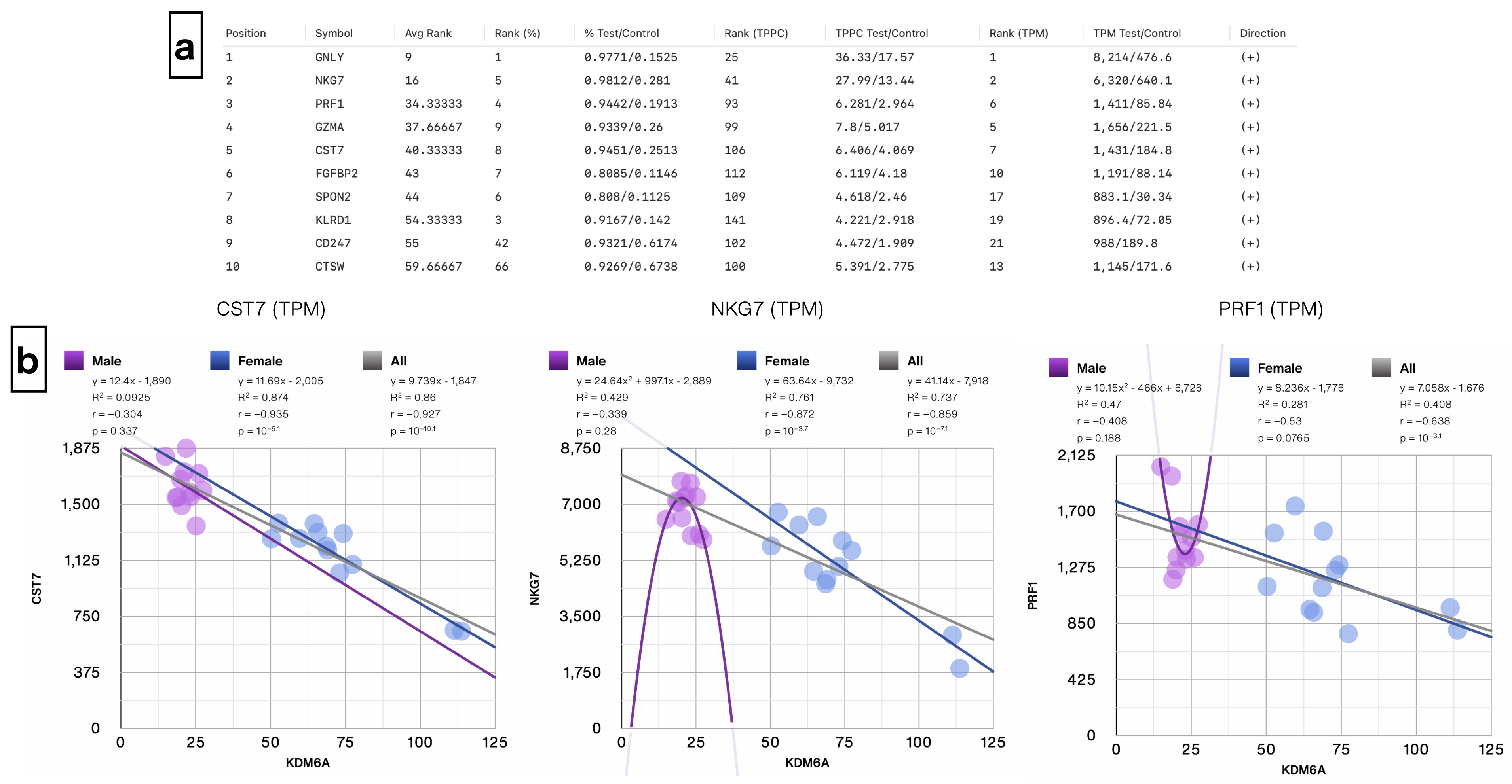
